# Using flux theory in dynamic omics data sets to identify differentially changing signals using DPoP

**DOI:** 10.1101/2024.07.29.605590

**Authors:** Harley Edwards, Joseph Zavorskas, Walker Huso, Alexander G. Doan, Caton Silbiger, Steven Harris, Ranjan Srivastava, Mark R. Marten

## Abstract

Derivative profiling (DP) is a novel approach to identify differential signals from dynamic omics data sets. This approach applies variable step-size differentiation to time dynamic omics data. This work assumes that there is a general omics derivative that is a useful and descriptive feature of dynamic omics experiments. We assert that this omics derivative, or omics flux, is a valuable descriptor that can be used instead of, or with, fold change calculations. The results of derivative profiling are compared to established methods such as Multivariate Adaptive Regression Splines (MARS), significance versus fold change analysis (Volcano), and an adjusted ratio over intensity (M/A) analysis to find that there is a statistically significant similarity between the results. This comparison is repeated for transcriptomic and phosphoproteomic expression profiles previously characterized in *Aspergillus nidulans*. This method has been packaged in an open-source, GUI-based MATLAB app, the Derivative Profiling omics Package (DPoP). Gene Ontology (GO) term enrichment has been included in the app so that a user can automatically/programmatically describe the over/under-represented GO terms in the derivative profiling results using domain specific knowledge found in their organism’s specific GO database file. The advantage of the DPoP analysis is that it is computationally inexpensive, it does not require fold change calculations, it describes both instantaneous as well as overall behavior, and it achieves statistical confidence with signal trajectories of a single bio-replicate over four or more points. While we apply this method to time dynamic transcriptomic and phosphoproteomic datasets, it is a numerically generalizable technique that can be applied to any organism and any field interested in time series data analysis. The app described in this work enables omics researchers with no computer science background to apply derivative profiling to their data sets, while also allowing multidisciplined users to build on the nascent idea of profiling derivatives in omics.

## Background

Data analysis is frequently the bottleneck in achieving biological interpretation of the results (*1, 2*). Thus, there is great interest in improved, generalizable, mathematical approaches to learn from large-scale omics data. Volcano plots(*3*) are the classic example of a numerically generalizable, omics-data-analysis technique. These plots of statistical significance versus fold change are ubiquitous in omics research, and can be found in genomic(*4*), transcriptomic(*5*), proteomic(*6*), phosphoproteomic (*7*), metabolomic(*8*), and lipidomic(*9*) studies. Their ubiquity is due to a numerical approach that is generalized to a format of data rather than a specific type of instrument or organism. Ratio versus intensity (M/A) plotting(*10*) is another type of generalizable statistical method used to identify differentially changing signals from a population of signals. It has been adjusted(*11, 12*) to include sample variance, creating an “adjusted p-value”(*13*) that results in a more robust false discovery rate. Multivariate adaptive regression splines (MARS) is a third method to identify differentially regulated signals from a population of trajectories(*14*).

These three methods each require fold change calculations which specifically have difficulty dealing with zero values. MARS considers the entire trajectory in its analysis, whereas Volcano and M/A plots look only at single endpoints. Hence, the latter two methods must be repeated for each time point of a signal trajectory. MARS is then the natural choice for dynamic time course data. Another difference is that MARS is applicable to omics trajectories of a single biological replicate, whereas Volcano and M/A plots require multiple bio-replicates to achieve statistical significance. All these methods introduce fold change bias, and none consider a derivative calculation.

The novel method proposed in this paper, which we will refer to as derivative profiling (DP), is numerically generalizable like these other methods. It can infer statistical significance from omics data using a single bio-replicate trajectory, similar to MARS analysis, but can also examine a particular end point, similar to Volcano plotting. DP determines statistical significance based on the z-score of the value of a normalized derivative in relation to a normal distribution of the population of normalized derivatives. DP is compatible with trajectories sampled using fixed or variable step sizes, however it cannot be used for single endpoint experiments since multiple points are needed to calculate a single derivative.

Derivatives in omics is not new, but have garnered attention under the theory and naming convention of RNA velocity, stated as “the time derivative of the gene expression state”(*15*). Despite over 1,500 citations of the original work (*15*), and success identifying useful features of biological systems, sources using “velocity” to describe behavior of biological macromolecules other than RNA are limited. A contemporary review of the state of RNA velocity research leaves out mention of any other omics entirely(*16*). There are few proteomic velocity papers(*17-19*), and to the best of the author’s knowledge, none regarding phosphoproteomic “velocity.” This is likely because important assumptions are made in regard to the underlying kinetics of transcript splicing. While these assumptions make RNA velocity a valuable descriptive model of transcriptomics, they make it difficult or impossible to apply this approach more broadly. These assumptions are the reason this method cannot be easily applied to other omics without rederivation of the underlying model. Phosphoproteomic kinetic interactions are less well characterized than transcript kinetics and do not undergo the same splicing phenomena. Published work has addressed that there are many assumptions limiting the original RNA velocity(*16*). We agree with the need for more generalized and simplified approaches and believe that DP represents a broader generalization of time derivatives in omics states such that it can be applied to any omics.

The theory behind DP is straightforward. All omics data yield peaks, which are transformed into counts and known to be proportional to an analyte’s relative abundance in the sample. This is stated mathematically below with Equation 1, where *N* stands for omics count per some unit area *A* of a sensor, *M* is the mass/mol abundance of some analyte over the sample unit area *A*′. We assume only that the sample area and sensor area are constant, and the omics sensor is operating in a linear dynamic range such that *P* is a unitless proportionality constant.

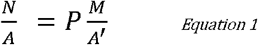

The classical definition of flux, Φ, is a change in mass per unit area per time (kg·m^-2^·s^-1^). By taking the time derivative of equation 1, equation 2 yields the dimensional units of flux.

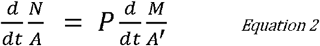

This can be restated as the derivative of the omics count is proportional to the flux of the analyte.

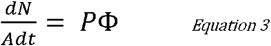

Deriving with respect to time again leads to an equation stating that the second derivative of the count signal is proportional to the change in flux.

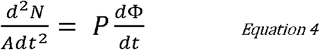

These mathematical generalizations of sensor/analyte behavior represent why this method is applicable to any omics, and why this work postulates that the derivative of an omics signal is appropriately called an omic flux. DP is founded on this omics flux theory that the derivative of an omics trajectory describes the flux of an analyte in a sample because the change in omics count is due to summative fluxes of the analyte for any number of reasons, reactive, diffusive, or other.

We propose a model for omics flux which only describes rates of total flux and does not assume to know anything about reaction, diffusion, or advection kinetics of the system. We use the definition of mass flux, where c is concentration (kg·m^-3^), and v is velocity (m·s^-1^).

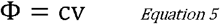

By multiplying both sides of the equation by a unit dimension of length, units of flux are conserved, and we simplify the units of concentration and velocity. The following equation results where *k*_*of*_ is the omics flux rate constant (s^-1^).

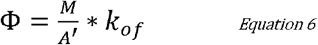

Equation 1 and 3 can be substituted into equation 6 to yield the final description of omics flux as described only by omics data, where *K*_*of*_ represents the omics flux rate constant incorporating the proportionality constant P.

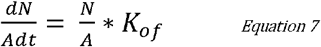

These combined ideas are what led to the theory that derivatives normalized by the average count signal yields rates, and these rates are useful features that describe omics signals without the need for fold changes. This omics flux rate constant is calculated for every signal in the population. Statistical significance is assigned based on a Z-score of the specific rate compared to the normal distribution of the population of rates.

The primary aim of this work is to advance all fields of omics by providing a new generalizable mathematical technique to discern differentially changing signals based on first principals of flux in engineering. This work distinguishes the generalizable idea of omics flux theory to be applicable in various types of omics data, and in multiple organisms. This paper will compare DP in its current state with the established methods used on previously published transcriptomic and phosphoproteomic datasets(*20-22*). We have packaged our DP method within a GUI-based MATLAB app, DPoP, or *Derivative Profiling omics Package*. This platform enables anyone with trajectories of dynamic omics data to easily perform this exact DP analysis without any coding experience. From within the app, one can run a GO term enrichment analysis(*23*) on their findings as long as that organism has a GO database file. The result is a field/organism specific description of the results. No MATLAB toolbox or expertise in installation is required since this app can be used with the free browser-based MATLAB Online terminal which automatically handles toolbox/version compatibility. Using this platform, DPoP enables researchers and technicians with no background in code or mathematics to perform this advanced analysis method and then do a bioinformatics screen on the results, all at the push of a button.

## Results

In practice, derivative profiling yields two features of interest: (i) the derivative estimation, a local rate estimation, and (ii) its slope, a regional rate estimation. The value of the derivative estimate describes the instantaneous behavior of the signal at each time point. The slope of the linear regression of the derivative is an approximation for the behavior of the derivative across the ENTIRE trajectory. These two features enable DP to offer a deeper level of understanding into cellular responses by investigating what is happening both at a point, and across the overall trajectory. Implementation will examine this behavior on single trajectories, followed by the results of the overall algorithm compared to other identification methods, and then the results of a GO term analysis of the output results of DPoP. After that, this work presents a comparison of DPoP results with published results from a phosphoproteomic experiment in S. cerevisiae and finally an overview of the GUI for DPoP. Exact descriptions leading to the variables used, such as 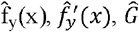, et cetera, can be found in the methods section.

### Behavior of DPoP on a Single Trajectory

An in-depth visualization of the behavior of the algorithm is detailed in Figures 1, 2, and 3. Figure 1 shows the same phosphoproteomic mass-spectrometry time course data in three different ways. Figure 1A shows the original phosphoproteomic time course data that has only been globally scaled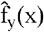. Figure 1B shows the data represented as relative fold change from the zero-time point. Figure 1C shows the inner workings/mathematics of DP, with mean-normalized differentiation data, 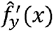, shown as open points and the linear regression approximation of the normalized estimation of the derivative, *Ĝ*, as a line.

**Figure 1.**
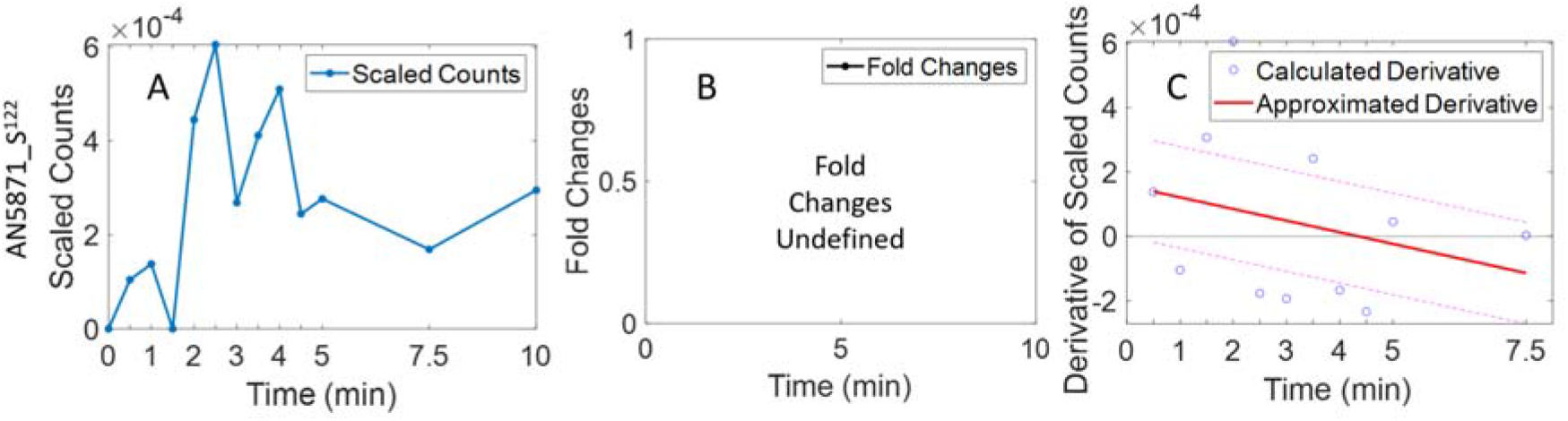
Visualization of the Data for AN5871_S^122^. A) Shows globally scaled Aspergillus nidulans phosphoproteomic mass spectrometry time course data over 10 minutes of micafungin exposure. B) Shows fold change calculations. C) Shows results of the differentiation calculation in blue as well as the linear regression approximation of that trend in solid red. Plus and minus 2 standard deviations is shown via the red dotted line

**Figure 2.**
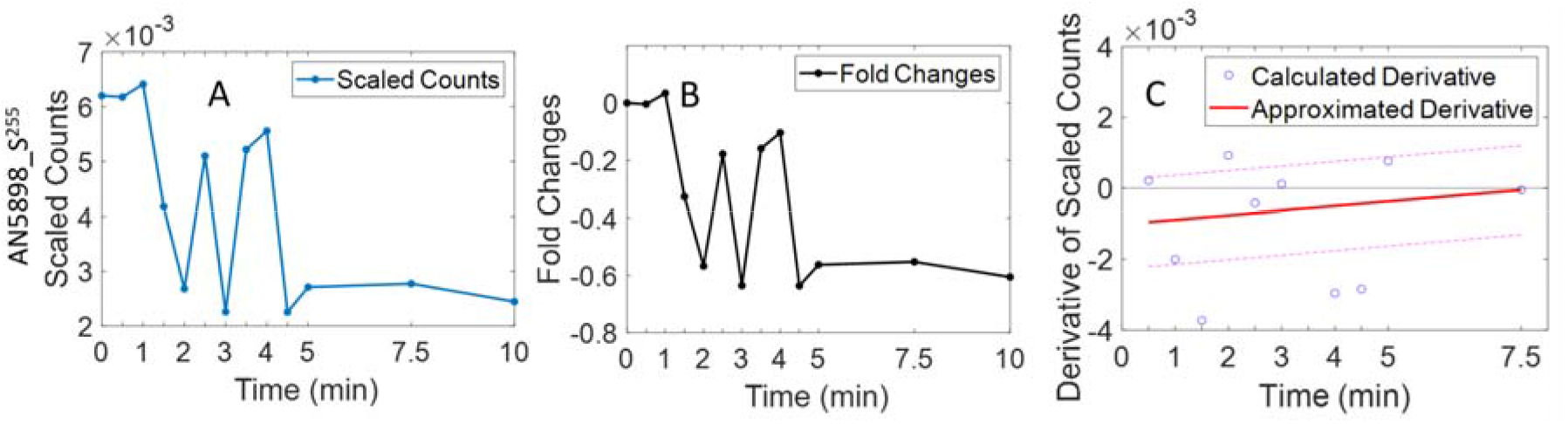
Visualization of the Data for AN5898_S^225^. A) Shows globally scaled Aspergillus nidulans phosphoproteomic mass spectrometry time course data over 10 minutes of micafungin exposure. B) Shows fold change calculations. C) Shows results of the differentiation calculation in blue as well as the linear regression approximation of that trend in red.

**Figure 3.**
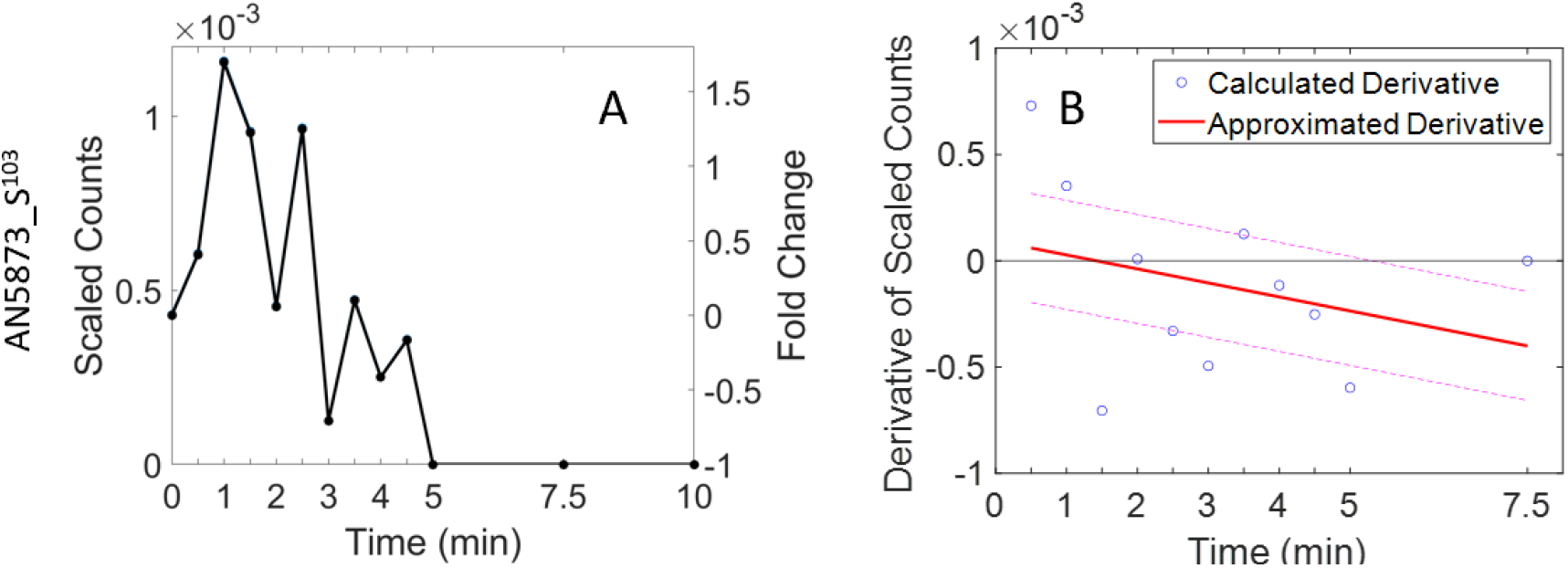
Visualizing the Data for AN5873_S^103^. A) Shows globally scaled Aspergillus nidulans phosphoproteomic mass spectrometry time course data and its fold change equivalent over 10 minutes of micafungin exposure. B) Shows results of the differentiation calculation in blue as well as the linear regession approximation of that trend in red.

Figure 1 demonstrates the difficulty in using only fold changes and demonstrates an ideal use case for DPoP where the experimentalist would benefit. For example, when an initial time point has a signal value of zero (e.g., Figure 1A), fold changes in 1B are undefined throughout the trajectory (i.e., it’s not possible to divide subsequent time points by the initial time point of zero). Thus, for fold changes, this leads to a bias toward on-to-off signals and against off-to-on signals. While traditional fold change calculations are unable to describe the behavior in Figure 1A, DP (Figure 1C) can describe the increase, the maximum, and the following decrease through derivative data that goes from positive to negative. While the maximum peaks at 2.5 minutes in the original data (Figure 1A), DP’s signal line (Figure 1C) crosses the x-axis near 4 minutes, closer to the second, smaller peak. Figure 1C also highlights how calculated derivatives can be quite scattered due to noise in the data, and how the linear regression smooths that behavior over. Figure 1C shows derivative data scattered above and below zero as the signal in 1A oscillates through its own trend. While the derivatives oscillate back and forth across zero, the regression approximation has a clear trend that starts positive and ends negative, congruent with the data in Figure 1A.

Figure 2 shows a different scaled phosphoproteomic mass-spectrometry time course from. Figure 2A shows data that has only been globally scaled. Figure 2B shows the same data represented as relative fold change from the zero-time point. Figure 2C shows the inner workings of DP, with mean-normalized differentiation data,, shown as open points and the linear regression approximation of the normalized estimation of the derivative,, as a line.

The first three time points in Figure 2A are flat-to-increasing and are followed by the significantly decreasing point at 1.5 min. Fold change behavior is mirrored identically, albeit with a different scale, in Figure 2B. Figure 2C shows the calculated and approximated derivative from the data shown in Figure 2A. We note that the differentiation algorithm uses a three-point sliding window, making a forward-looking property of the algorithm. As a result, the first calculated derivative includes data at (0, 0.5, 1min from 2A) and is slightly positive (0.5min in 2C). The second calculated derivative includes the fourth point (1.5 min in 2A) and accounts for the larger negative value at 1.0 min in 2C. The line in 2C is a regression of all the calculated derivative values and has the benefit that it immediately shows a negative value (i.e., at 0.5min) whereas the fold change plot (2B) does not show this until 1.5min. When we examine the last four points in Figure 2A, they are relatively flat compared to the rest of the trajectory. Fold change (2B) is range bound around –0.6 for the last four timepoints (i.e., 4.5, 5, 7.5,10mins). After 4.5 minutes, fold changes indicate the same level of expression. This is correct *in reference to the initial timepoint*, but it does not describe the current, instantaneous behavior within the trend. Without more calculations, fold changes make no distinction that 4.5 minutes is the end of the negative effect, and that after that it is at a new steady state. However, this behavior of negative-flux stopping and converging on a new steady-state is exactly what DP depicts. Looking at the approximated derivative of Figure 2C, one will observe a negative derivative at 4.5 minutes which increases toward zero at 7.5 minutes. We note that a zero derivative is the definition for non-differential in this method and the approximation line in 2C converged upon the correct answer, zero flux at a new steady state. This is more descriptive than fold changes (2B) which make no meaningful distinction between fold change of the data at points beyond 4.5 minutes.

A third and final trajectory is explained in Figure 3, which has a similar trajectory as Figure 2 but with a different outcome. In Figure 3A, data show an initial increase in the signal up to 1 min, followed by a noisy decrease through 5 min, and then a flat period. In Figure 3B, for times at and before 1min, the approximated derivative line has positive values. The slope of the regression line is negative and the derivative crosses into negatives. Together, this information implies the signal is increasing but increasing briefly to a maximum and then decreasing. The regression line in 3B then moves through zero at approximately 1.5min, indicating a maximum in the data, which occurred at 1 minute (Figure 3A). We note that fold changes in 3A do not indicate the transition to negative values until 3 minutes and at 2.5 minutes fold changes correctly identify a positive fold change at that point. While this is correct, based on the definition of fold changes (i.e., in reference to the initial time point) DP indicates this negative trend faster than fold changes in two ways. First, the derivative approximation of 3B is negative by 2 minutes, as it crossed the x-axis at 1.5 minutes, making anything after 1.5 minutes identifiable as being negatively affected. Second, the sign of the slope of the derivatives is immediately available as an observation about all time points. The negative slope of the derivative in 3B means increasing less, or decreasing more, both of which could be said about the data in Figure 3.

The approximation line in 3B is at a relative minimum at the end of the time course. We note that current methods rank this derivative estimation line, not the calculated derivatives. This means the end of the time course is when there is the highest chance this trajectory will be flagged as differential. However, it is unclear without the data at 5, 7.5 and 10 minutes that the trend is significantly down *overall*. While the end of 3A represented the points most negatively affected, the true derivative of the new “off” steady state *should be* zero. This is reflected in the calculated derivative at 7.5 min being zero but not the approximated derivative, which is what DPoP uses as its signal line. Since the signal line identifies differential signals, current methods would flag this phosphosite after it turned off. This result is useful, but not exactly phenomenologically correct, because it is not differential once it is off, it is differential when it is turning off. This divergence of the signal line from the true derivative calculation is a result of our underlying assumption of first order rates over the time span we are examining. Changing the regression order is not currently a feature of DPoP but changing the span of the overall regression is. By changing the span of the regression, the length of time can be adjusted to approximate over shorter intervals, leading to potentially more accurate local derivative estimations.

### DPoP Output Compared to Other Algorithms

While analyzing the datasets with DPoP, default settings were used, and the results from the analysis of the first two time points were retained. The first two time points were chosen because, as one can see from Figure 1, 2, and 3, the greatest rates of change occurred in the beginning of the experiments, and all trajectories approach a new steady state (flat trend) closer to the end. DPoP ranks instantaneous rates of change, whereas all other methods rank total change from the beginning (fold change). The resulting list of significantly differential analytes from DPoP is compared to the results from other established numerical techniques (Figure 4). Lists of differential signals as per the other analysis methods can be found in the supplementary data S2.1 and S2.2.

**Figure 4.**
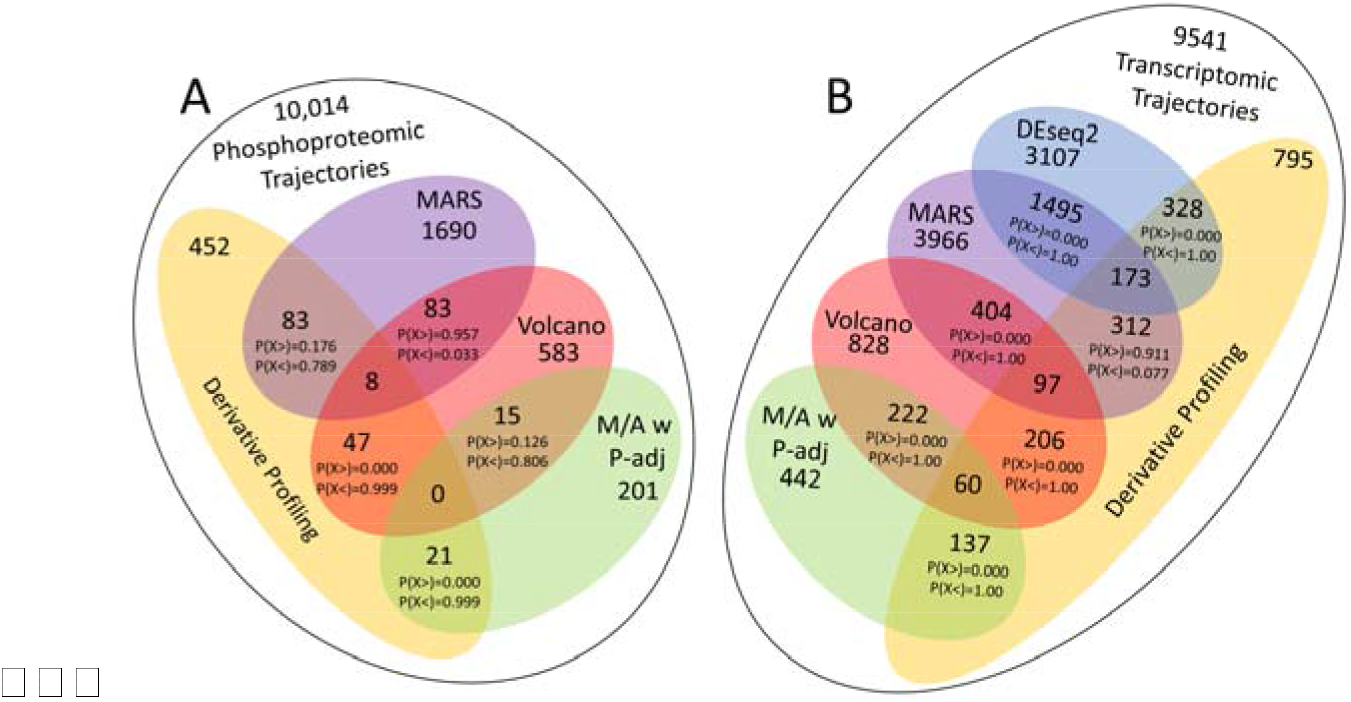
Comparison of Results from Various Methods used to Identify Differential Regulation/Abundance. Results from DP are compared to the results of a MARS analysis, significance vs fold change analysis (Volcano), ratio-over-intensity analysis with adjusted p-stat (M/A w P-adj), and DEseq2 for the transcript comparison only. Hypergeometric test was used to determine the significance of the overlap of names identified from two different methods applied to the same data. 4A) Shows comparison of various analyses performed on the phosphoproteomic mass spectrometry data. 4B) Shows comparison of various analysis performed on the transcriptomic RNA sequencing data.

Results in Figure 4 demonstrate that for both the phosphoproteomic dataset and the transcriptomic dataset from *A. nidulans*, DP has a statistically significant over representation of results when compared to volcano and M/A analysis, but not compared to MARS analysis. Take for example Figure 4A, at the intersection of DP and Volcano results. IFF the two sample sets were drawn at random from the same population, there would be a 99.9% chance that those two sets would have less than 47 in their intersection. This is a statistically significant overrepresentation of DP results within the volcano plot results. The same logic can be applied to any of the intersections with probabilities. We note that the behavior, (i.e., similar to volcano and M/A, dissimilar from MARS) holds similar for both the transcriptomic and phosphoproteomic data.

### Results of GO Analysis of DPoP Results

Typically, after any method has been used to identify significantly differential analytes, various approaches are used to discern biological trends relevant to the experiment. To simplify this step in DPoP, we have included integration of a GeneOntology Database file (.gaf) and GO term enrichment analysis. Thus, DPoP uses the enriched GO term analysis to programmatically describe the trends in a field-specific and human-interpretable manner. Any organisms .gaf file can be loaded into the app so that omics data in any organism can be analyzed and described in a way that is meaningful to the non-computational experimentalist. The GO term enrichment assigns statistical significance to GO terms that are over- or underrepresented in the differential signals as compared to the background of all GO terms in the database file. GO term enrichment analysis is performed on results from DPoP using the default settings and examining only one timepoint at a time. The resulting GO terms describe each individual timepoint and if the analysis is repeated over each timepoint, a description can be made of each point through the time course. This temporal GO term analysis was performed on phosphoproteomic and transcriptomic datasets, and a summary of the results are included in Figures 5 and 6, respectively.

**Figure 5.**
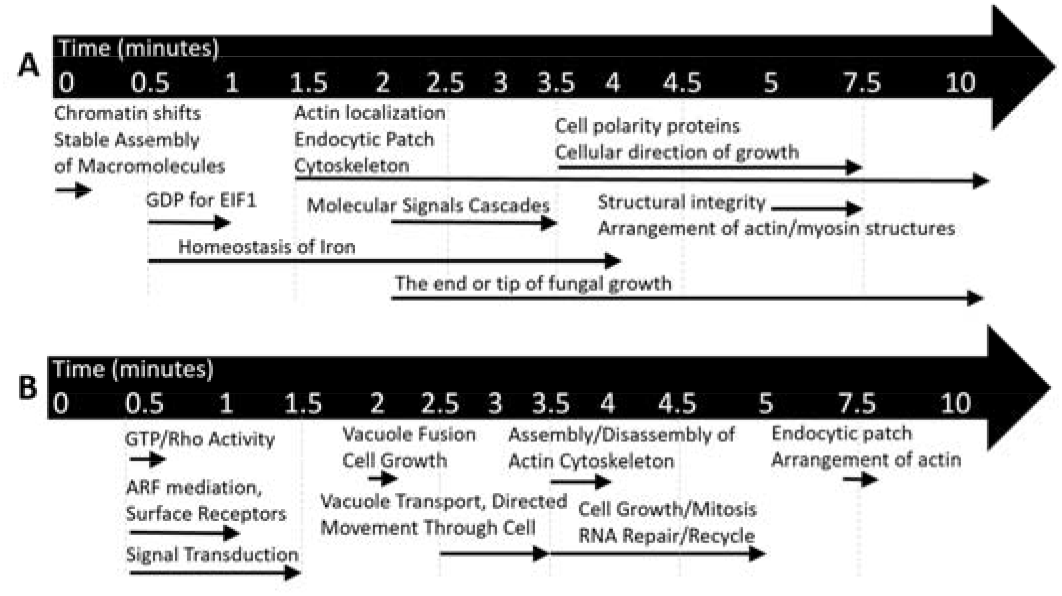
Temporal GO Term Enrichment Analysis on the Results of DP(5.A) and Volcano Analysis(5.B) Throughout the Phosphoproteomic Time Course. Descriptions in this figure were programmatically generated based on a GO term enrichment analysis of the results of DP(5.A) or the results of Volcano analysis(5.B), at each point in time. The base of any arrow represents when the particular GO term was first seen in the results. Likewise, the end of an arrow represents when this particular GO term was no longer seen in the results. These results use the 10 most significant GO term descriptors from the list of differential signals at each time point.

**Figure 6.**
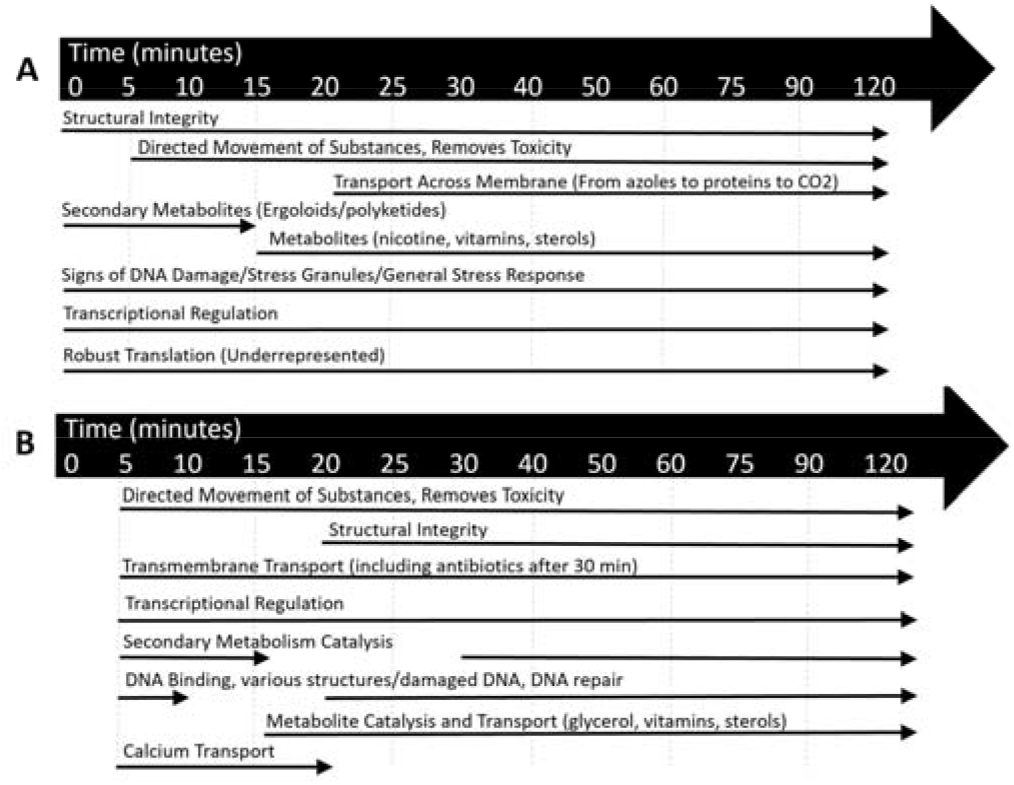
Temporal *GO Term Enrichment Analysis on the Results of DP(6.A) and DEseq2(6.B) Throughout the Transcriptomic Time Course*. Descriptions in this figure were programmatically generated based on a GO term enrichment analysis of the results of DP(6.A), or DEseq2(6.B), at each point in time. The base of any arrow represents when the GO term was first seen in the results. Likewise, the end of an arrow represents when this GO term was no longer seen in the results. These results were taken from the first 50 most significant GO term descriptors about the list of differential signals at that time point.

When developing a new method, it is important to compare results to known truths. With this in mind, our findings here are in close alignment with a previous publication describing this dataset(*20*). The GO term analysis of the *A. nidulans* phosphoproteomic dataset (Figure 5) yielded a resolved story of a cell responding to cell wall stress through signaling cascades in pathways involving structural integrity actin arrangement. It also identified the prevalence of iron homeostasis genes. There are even GO terms describing endocytosis and morphogenesis, thought to be the secondary effects of actin repurposing which are not as well characterized(*24-26*). The GO terms appear to be properly arranged temporally as well, moving from stable assembly of macromolecules in the beginning, to GDP activity, followed by a molecular signal cascade, and then by changes in structural integrity and arrangement of actin/myosin. DPoP, combined with GO term enrichment, was able to programmatically describe temporal slices of the phosphoproteomic mass spectrometry data with a striking congruence to current hypotheses regarding *A. nidulans* treated with micafungin. Comparing Figure 5A with figure 5B shows similar biological behaviors throughout the experiment. For example, both methods identified phosphosites involved with signal transduction. However, there are also differences. For example, GO analysis on the DPoP results identified iron, structural integrity and cell polarity terms, whereas GO analysis on the Volcano results yielded more phosphosites involved in endocytosis and vacuole mediation. Together, these temporal GO descriptions accurately describe what is expected in *A. nidulans* phosphoproteomic response to micafungin exposure (*20*). GO descriptions which produced these figures are available in the Supplementary data S3.1 and S3.2.

The transcriptomic dataset below (Figure 6) was less resolved via GO term enrichment analysis of the DP results through time. Many of the GO terms, once they appeared, would stay overrepresented throughout the trajectory. Structural integrity, transcriptional regulation and stress response were overrepresented in all timepoints. Secondary metabolites such as ergoloids and polyketides were overrepresented first, and primary metabolites like nicotinamide, vitamins, and sterols were overrepresented second. At the same time this is happening, transmembrane transporters for numerous compounds, from azoles to H^++^ ions, were expressed. This fits the theory that when *A. nidulans* encounters micafungin, it assumes it is under chemical attack from a foreign organism. By increasing abundance of complex secondary metabolites, A. *nidulans* is fighting back by producing known phytocides and bactericides. Transmembrane flux is also consistently overrepresented alluding to a cell trying to pump unwanted toxin outside of the cell. Comparing Figure 6A with 6B,shows similar results.. The GO analysis of the DEseq2 results identified genes involved with glycerol metabolism and calcium transport. While DPoP analysis did not identify glycerol or calcium descriptions, it did identify a wider range of secondary metabolites. Together, these temporal GO descriptions accurately describe what is expected in *A. nidulans* transcriptomic response to micafungin exposure (*20, 21*). GO descriptions which produced these figures are available in the Supplementary data S3.3 and S3.4.

### DPoP Results in S. cerevisiae

In another effort to compare the results of DPoP to known findings, the mass spectrometry dataset corresponding to the phosphoproteomic response to high osmolarity in *S. cerevisiae(22*) was analyzed with DPoP. This data set was included to demonstrate the flexibility of DPoP to other organisms and it is available in the default files of DPoP as an example problem. Of the 5,453 phosphosites identified in that work, the original authors identified 596 sites as differentially phosphorylated. Using the default settings, DPoP identifies 802 differential phosphosites. Between these two separate methods of identifying differential activity, there was an overlap of 116 phosphosites identified as differential by DPoP, as well as by the specific methods of the authors of that work. Using the hypergeometric probability distribution there is >99% confidence that there was a nonrandom overrepresentation of differential phosphosites identified by the original work that were also identified by DPoP. These phosphosites were analyzed via GO term enrichment and the resulting overrepresented GO descriptions included response to osmolarity shock, the subject of the paper/experiment, as well as numerous kinase cascades, ATP utilizing reactions and macromolecule transport/localization. These list of differential phosphosites in the *S. cerevisiae* dataset can be observed in the supplementary data S4 or replicated from included files.

### DPoP GUI

A significant result of this work is the app-based platform DPoP which can be used to apply this method or to build on it. DPoP was designed so that a scientist without a mathematic or computational background can perform this complex analysis on their own dynamic omics data with a point and click experience. This app is published under the MathWorks open-source licensing for anyone to use or build upon. The app screen in Figure 7 is what users see when analyzing their dataset using DPoP.

**Figure 7.**
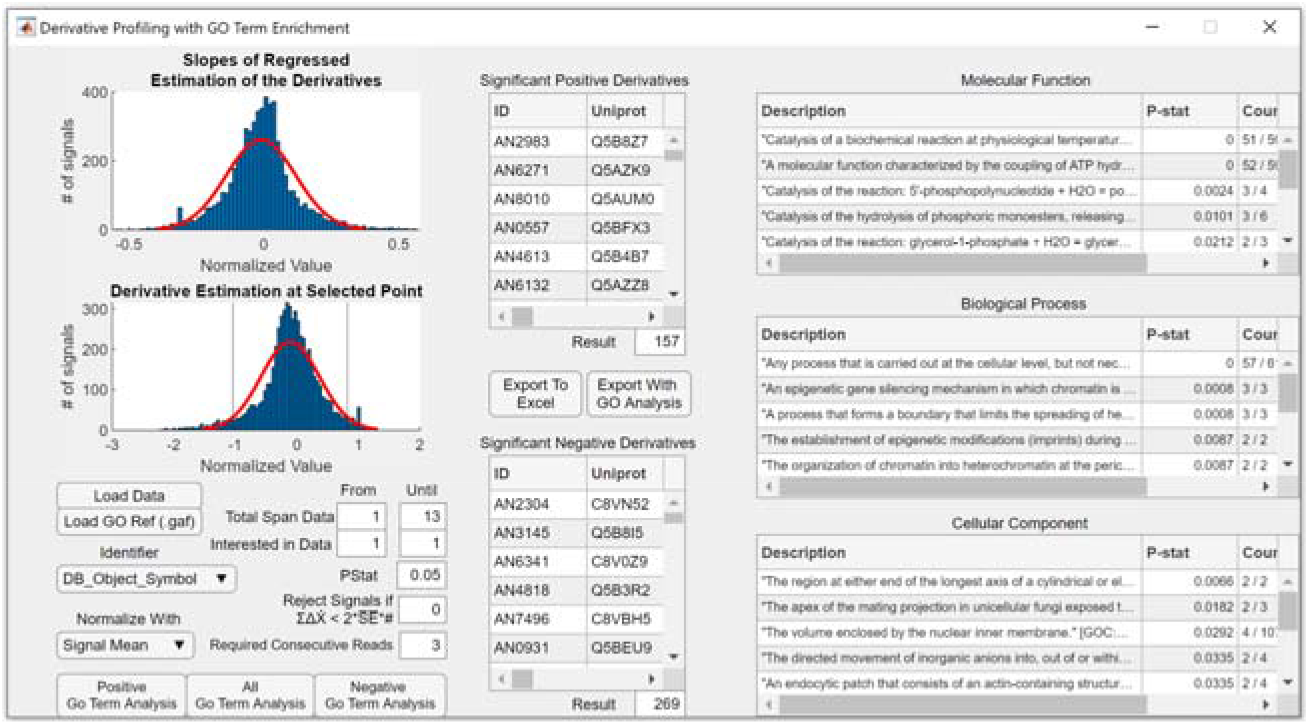
MATLAB App Based GUI DPoP: The distribution of derivatives at a point, as well as the distribution of slopes, is represented in the GUI on the top left. The inputs/user controls are at the bottom left. The list of signals with significantly positive and negative derivatives is in the column in the middle. The column section on the right displays the results of the GO term enrichment analysis applied to the list of significant signals, with respect to the population of signals. Controls are included in the GUI to select the statistical significance, the normalization factor, the GO database file, GO database field identifier, the first and final timepoint of interest in the regression, the first and final time point of interest within the regression, and the threshold for filtering based on noise or number of consecutive non-zero signals.

The histograms in the top left help visualize the population of signals behavior. The vertical grey bars in the derivative estimation histogram represent the cutoff values for significance and are adjustable with the Pstat field. One can observe the results appear in the middle column as they adjust settings. When preferred settings are found, and a .gaf file is loaded, GO term analysis buttons launch the GO term enrichment analysis on all, or only the positive, or negatively affected signals. This will populate the column to the right with GO term descriptions of the lists of differential signals found in the middle. These signals, their descriptions and information about where/how they are identified as differential, can be output to an excel file for further examination.

## Discussion

Essentially, DP is a methodology to be used with large lists, i.e., omics sized populations of dynamic signals, and make a smaller list of experimentally relevant omics signals. The researcher still must find meaning in the list or use the list to generate a hypothesis regarding what happened during the experiment. To this extent, the list of important genes, or phosphosites, is normally analyzed using a database like StringDB (*27*), or NetworKIN (*28*), respectively. We have included a bioinformatic description tool (GO term enrichment) in the app specifically so that a biologist doesn’t need to export lists and upload them to another database to get a human interpretable description of the results produced by DPoP. By doing so, DPoP can automatically describe the biology inherent to what was differential □ per the DP results.

A comparison between DP, and some other commonly used methods for omics data analysis is shown in Figure 4. This figure shows DP is a technique similar to volcano and M/A plotting, but quite different from MARS analysis. In both Figure 4A and 4B, there is a statistically significant overlap between DP, volcano, and M/A. Looking particularly at Figure 4B, at the overlap of DP, volcano, and M/A, DP identifies aspects of both other methods that singular methods do not identify by themselves. For instance, of the 447 genes identified via M/A plotting, 222 were also identified by Volcano plotting, leaving 225 genes as identified only by M/A plotting. DP identified 137 of those 225 M/A specific results, or over 50%. Similarly, of the 828 genes identified by Volcano plotting, 222 were also identified by M/A plotting, leaving 606 genes specific to Volcano plotting. Of those 606, 206, or ∼30%, are identified by DP. The same logic can be applied to the phosphoproteomic dataset of Figure 4A, albeit with statistics of 11% and 8%. This is still significant as the entire data sets are near 10,000 analytes each. Picking only 5% of the population, two separate times, but yielding 10-50% overlap, is highly convincing that the differential analyte selection process is similar, and the results are just as potentially meaningful as the other methods. This data suggests DP identifies differential signals as we currently understand them, but also identifies differential signals not included in the current methods. This method is similar to both, but also distinctly unique.

By using the value of the derivative, *as well as the trend of the derivative*, DP provides not just an alternative to fold change calculations, but an objectively more descriptive technique. Seen in Figures 1, 2 and 3, the derivative mathematically describes the data it came from. Fold changes do too, but neither should be taken as universally “correct.” Both are transformations of the same dataset. Fold changes are just a single feature of the system, as are derivatives. Fold changes reference initial points and they don’t project forward. Derivatives reference instantaneous points with directionality that can be projected to the next time point. By using the derivative estimate and also the slope of these estimates, researchers will be able to infer a deeper level of understanding than they could using fold changes alone. Now, instead of just “upregulated” or “downregulated”, a researcher, or program, can classify a line to say with empirical evidence “downregulated to a new steady state”, if the derivative starts negative and goes to zero, as in Figure 2C. The value of the derivative estimate gives the researcher a snapshot in time, (is increasing now), while the slope describes the whole trajectory, (but is increasing less over time). Mathematics around derivatives, like peaks, troughs, and zeros, add a further level of understanding by describing max or min rates of change and steady states, respectively. DP is also objectively less restrictive than any of the other methods used in this work due to the derivative calculations being able to handle zero readings in the first time point. All other methods compared in this paper required fold change calculations which are undefined if the first time point is zero. This is clear when examining Figure 1. Naturally, fold changes bias the researcher to see on-to-off and filters out off-to-on signals. DP does not have this bias. The only way to divide by zero if normalizing with the signal mean is if the entire signal trajectory was zero.

All omics signals from Figures 1, 2, and 3 had relatively flat trends at the end of the trajectory. These later timepoints would make the approximated derivative/signal line flatter. As a result, using fewer timepoints from the end of the trajectory would make the approximated derivative steeper. This is the first indication within the data that DP could perform more accurately on shorter time intervals. We propose this is because the derivatives are very likely not just simple linear trends as is the fundamental assumption of DP. However, over a sufficiently small window, nonlinear behavior can be accurately approximated with a linear trend, as is a fundamental assumption of calculus and modern algorithmic solvers. Extrapolating this property with DP means that only so much of the derivative can be accurately summarized with a linear trendline at one time. The lower resolution of the temporal GO description of the transcriptomic analysis (Figure 6), as compared to the phosphoproteomic temporal description (Figure 5), is likely the second indication in this work alluding to DP working best with smaller timesteps, or smaller total time spans. This is theoretically congruent for a method utilizing linear approximations of nonlinear phenomena.

Normalization by signal mean is the single most important factor in fitting the data to a normal distribution, as seen in Figure 9. Neither data set fits a normal distribution without preprocessing this way. Signal mean normalization is a crucial step in determining the statistical significance of the results, as the significance is based on the inverse cumulative probability of that distribution. Since only a normal distribution is applied, a fundamental assumption of this work is that a derivative rates of scaled and normalized omics dataset can be described by a normal distribution. This will not always be the case. DEseq2, the R package for differential omics analysis, has had wide success in examining RNAseq fold change calculations, not derivatives, under a negative binomial distribution instead(*29*). While the fit of the distribution is important, this paper does not test what the “best” probability distribution will be for all omics experiments analyzed via profiling the derivatives. That answer, or at least the functionality to test multiple probability distributions, represents a clear opportunity to explore in future work. Surely, the distribution of results will be highly dependent on the experiment, the normalization factor, the differentiation method, and the preprocessing steps. These numerous dependencies in the DPoP workflow are certainly non-trivial and worth future consideration, yet they do not fundamentally change the underlying process of profiling derivative rates as useful features of the -omics system.

**Figure 8.**
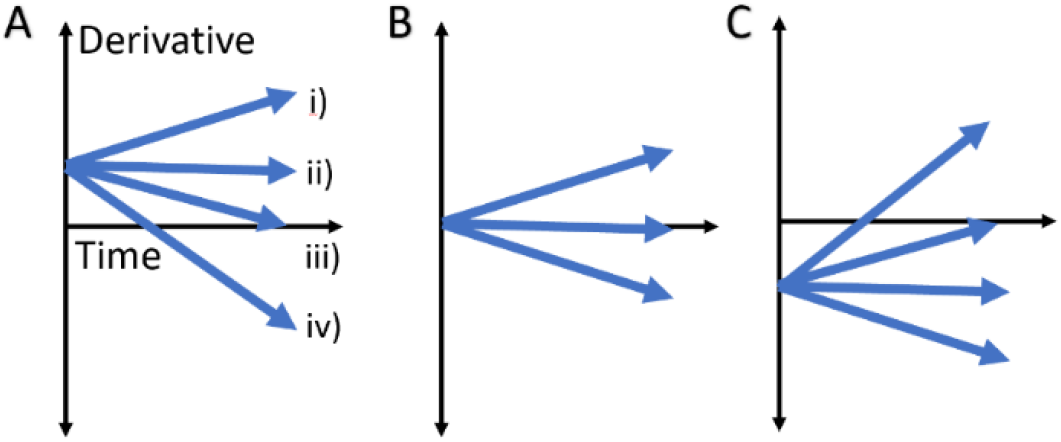
Range of All Real Number Approximations of Derivatives: All real number derivative profiles possible when the derivative trend is assumed to be a linear function. A) The derivative could start positive and (I) increase, (ii) remain constant, (iii) decrease to zero, or (iv) decrease through zero. The same trends would describe derivative regressions that start at approximately zero (B) or at a negative value (C).

**Figure 9.**
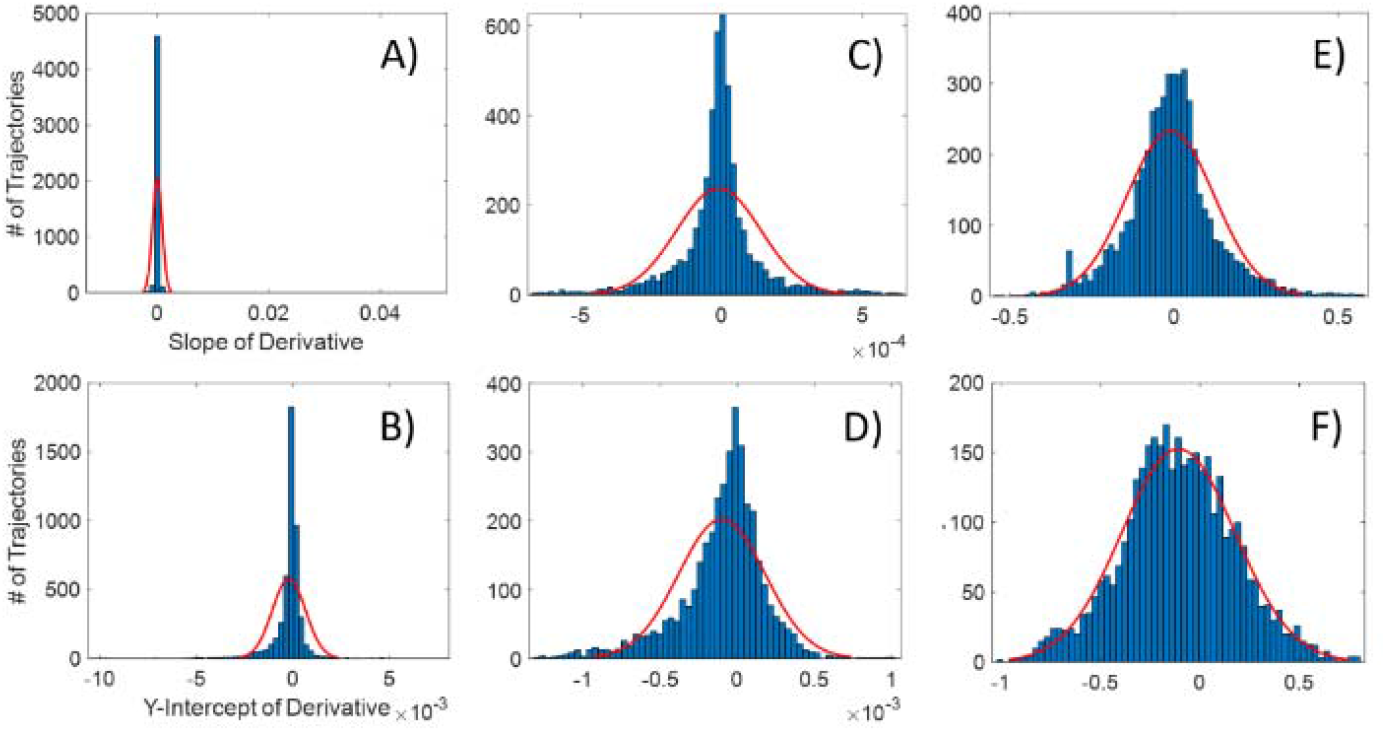
Distributions of Slopes and Intercepts of Linear Regression of Derivatives of Phosphoproteomic Mass Spectrometry Under Various Processing. The histograms represent the number of trajectories within a certain range of values for the derivative slopes (top) or derivatives (bottom). Red lines are normal distributions fit to the dataset. A, B) Show the distributions of slopes and Y-intercepts, respectively, of the linear regressions of derivative trajectories with no outlier discrimination nor normalization. C, D) Show the distributions of slopes and Y-intercepts, respectively, of the linear regressions of derivative trajectories with Grubbs test outlier discrimination and no normalization. E, F) Show the distributions of slopes and Y-intercepts, respectively, of the linear regressions of derivative trajectories with Grubs test and normalized with the non-derivative signal trajectory mean.

DP could be adjusted in several different ways and is without a doubt worthy of continued research. Different sized datasets should be tested to see if the method becomes increasingly useful over smaller and smaller incremental differences as is the case in examining datasets in this work. Variables in the method include the shape of the probability distribution, the normalizing factor, the regression method and order, the data exclusion criteria and derivation technique. These steps are represented by the arrows in Figure 10. Changing parameters within these steps, or even using a different but congruent algorithm in one of these steps, could optimize or change the technique in a variety of ways, but the overall framework of profiling derivatives as fit to a distribution is maintained. There is no requirement to use the time derivative. □ As DPoP is currently coded, it will differentiate with respect to whatever numbers are in the dynamic variable row of the data. While in this work the differential variable is always time, it is exciting to consider this method applied to data differential in any other variable, like concentration of a toxin. More work could be done applying if-then logic to sort the results into descriptive bins based on their derivative at different times. There is quite a lot of flexibility in the method, and much is still unknown regarding velocity/flux/derivatives in omics data.

**Figure 10.**
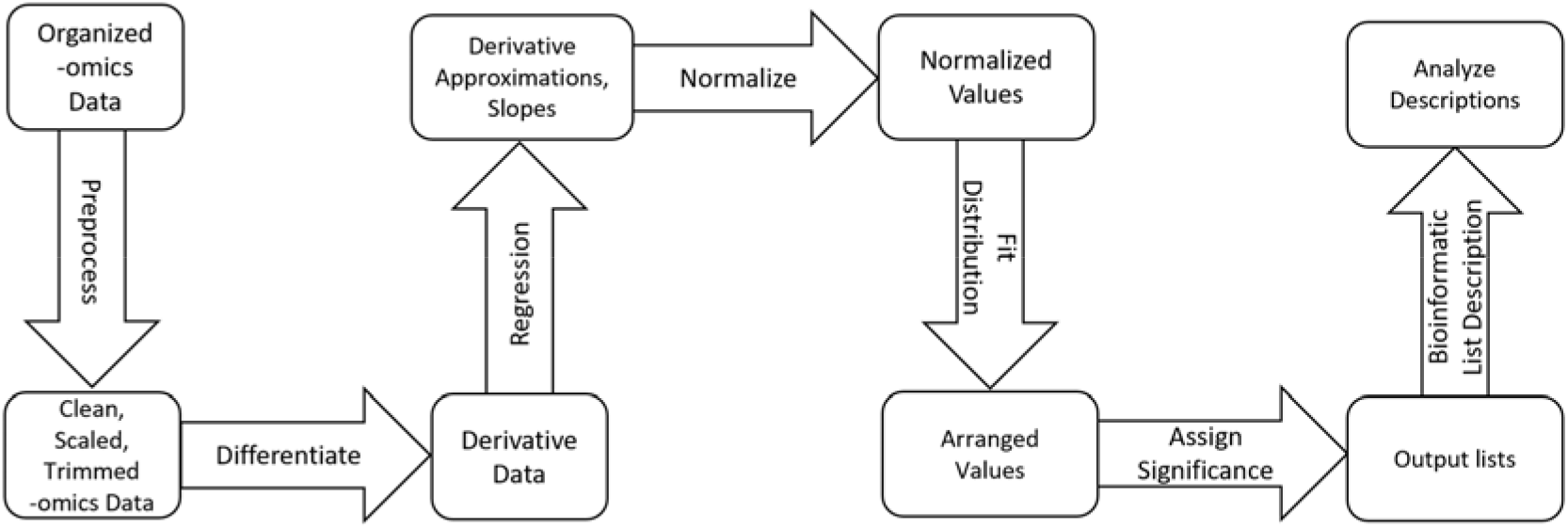
Flowchart of derivative profiling workflow. Data is input in the first block labeled “Organized -omics Data”. Any arrow is a tunable process operation of the overall workflow. Lists can be output and analyzed independently without bioinformatic analysis.

## Conclusions

The widespread applicability of this work provokes intriguing thoughts regarding the ubiquity of calculus and mathematics within biology. DP was partly inspired by advances in transcriptomic theory, particularly RNA velocity. Within classical chemical engineering, the idea that differences in concentrations govern particle behavior is the basic level of understanding reactive and diffusive flux. With this point of view, it is not surprising that transcripts have flux, rather it is encouraging to see fundamental assumptions of engineering continue to hold their merit in new and advanced applications. We encourage investigation into omics flux in a more general sense with this app, DPoP. We have applied this technique to whole cell phosphoproteomic mass spectrometry, and RNA sequencing data, and found that in both datasets DPoP can identify differential regulation significantly similar to what current, established methods are able to identify, seen in Figure 4. We also see that a GO term enrichment analysis of the DP results yields a temporally congruent description of what is known to be happening in the experiments from which the data was collected(*20-22*). This work expands and simplifies the idea of transcriptomic velocity and suggests there are flux values embedded in any dynamic -omics data set which can be used as descriptive features of the experiment without assuming any knowledge at all about reaction kinetics.

Just like the volcano plot analysis, this numerically generalizable method could be used ubiquitously to help numerous fields of researchers identify and describe signals of interest in dynamic omics experiments. To the best of the authors’ knowledge, this work is a novel application of the variable timestep LaGrange interpolation to omics data. This may also be the first consideration of phosphoproteomic “velocity,” or omics flux theory applied to phosphoproteomic mass spectrometry. To the best of the authors knowledge, DPoP is the first open-source tool that generalizes omics flux theory for application to any dynamic omics data set. We believe that the easy-to-use app implementation of these generalizable methods provides a beneficial contribution to all omic communities.

The case has been made that DP is less restrictive and more descriptive of omics trajectories than simple fold change calculations. This does not mean fold changes are meaningless. It has been said that “all models are wrong, yet some are useful” (*30*). Thus, it is more appropriate to identify which method is most useful given a specific application. For example, volcano plots work well for single endpoint experiments with multiple bio replicates, as do M/A plots. MARS analyzes a trajectory, not single endpoints. DP also requires trajectories, not single endpoints. While multiple bio-replicates should be used in practice, DP will work on a single bio-replicate of data but is restricted to experiments with at least four points in the trajectory. The experimentalist should decide whether to analyze bio-replicates individually or as one averaged data set. DP does not require fold change calculations, making it more widely applicable than those that do. It describes a snapshot along a trajectory, like volcano plots, but considers the whole trajectory in the final estimations of significance, like MARS. There is consistently a statistically significant overlap in the results of DP with the results of volcano and M/A analysis, but not with the results of MARS. DP identifies many results that would be identified by only volcano or M/A plots and identifies results that would not be identified by either. This work indicates DP does find differential regulation, but with a different bias than current methods. We believe this different view is a valuable addition to the toolkit of an omics researcher.

### Implementation

#### Data and Preprocessing

The omics data used in this work has been published previously (*20-22*). Such data includes phosphoproteomic(*20, 22*) and transcriptomic(*21*) time courses, taken during the exponential-growth phase of *A. nidulans* and *S. cerevisiae*, under cell-wall perturbation or high osmolarity conditions, respectively. Both phosphoproteomic and transcriptomic data from *A. nidulans* have 13 unevenly spaced timepoints and contain signals for 10013 phosphosites, and 9541 transcripts, respectively. The phosphoproteomic data from *S. cerevisiae* (*22*) has 13 evenly spaced time points describing 5453 phosphosites. DPoP is able to process bio-replicates when they are averaged at equal time points. For example, in the *A. nidulans*. phosphoproteomic dataset(*20*), all technical and biological replicates were averaged. In the *A. nidulans* transcriptomic set(*20, 21*), the average of all bio-replicates of the transcript-length normalized counts were used. In the *S. cerevisiae*. phosphoproteomic data set, the peptide intensity is used exactly as published in that literatures supplementary files(*22*). When biological replicates are carried out at different time points, DPoP must analyze the bio-replicates separately.

Data must be preprocessed for DPoP compatibility by removing strings or non-number values (i.e., Inf, NaN, or ERR). Count data should never be negative. If a specific signal trajectory has three or fewer signals (out of thirteen) it is removed. The 3 of 13 heuristic was specifically chosen for this work due to the length of the dataset and because the derivative calculations require three points. This step was programmed into DPoP so that researchers in the future may exclude non-consecutive data depending on the size of their own dataset.

Several computational techniques were used in this work. The results of Volcano(*3*), ratio-over-intensity (M/A) (*13*), Multivariate Adaptive Regressive Splines (MARS)(*14*) and GeneOntology (GO) term enrichment analysis(*23*) results are included for the *A. nidulans* transcriptomic and phosphoproteomic datasets in the <u>supplementary data S2.1 and S2.2</u>. The MARS analysis was completed previously with the open-source package AresLab(*14*). The code used to generate the Volcano and M/A plots was adapted from the MATLAB Bioinformatics Toolbox(*31*). DEseq2, a package in R, was used to analyze the transcriptomic set. All of the code necessary to run DPoP using MATLAB or MATLAB Online can be found, along with the omics data, in the <u>MathWorks File Exchange download</u>. A description of all files in that download can be found in the supplementary data S1. The application described here is packaged with these datasets (preformatted in .mat or .xlsx format) as well as the *A. nidulans* and *S. cerevisiae* GO annotations (.gaf file). All code was written using MATLAB (MathWorks; r2020a software license).

#### DPoP Workflow

Given a dynamic omics dataset with T timepoints, x, from t to T, and I number of trajectories, y, from i to I, DP analysis is carried out as follows. First, the data is globally scaled between zero and one, by dividing all data by the global maximum value in the dataset. We refer to any scaled trajectory data as fy(x), which is the scaled approximation of the original data fy(x). fy(x) is differentiated with respect to time using the modified LaGrange interpolation method((*32*)) to approximate fy ′ (x). fy′ (x) is a series of points that approximate the first order derivative of the signal trajectory, fy(x), at a given time point. This differentiation algorithm was chosen for its ability to calculate derivatives from data with unevenly spaced sub-intervals. This allows our approach to effectively deal with different time points or missing observations. To broadly characterize the overall behavior of the trajectory, 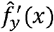 is fit with a straight line by linear regression to yield the function *g*_*y*_(*x*) where the slope of *g*_*y*_(*x*), represents a rough estimate for 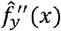. This process is repeated for all trajectories. Since the derivative estimation is calculated using three points, *f*_*y*_(*x*)is not calculated the first or last time point, *f*_*y*_(*t*) or *f*_*y*_(*T*). DPoP builds a matrix G, of size [I,T] out of approximated derivative values calculated from *g*_*y*_(*x*), for all y, extrapolating for x = t and x = T. Using the slope of *g*_*y*_(*x*) for all y, we build a column vector *G*′ [I,1] in size holding the 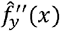 approximation for each trajectory, which was shown in equation 4 to be proportional to the change in flux for the analyte. The range of behavior of all possible real number derivatives is simplified and described in Figure 8.

To keep high intensity signals from being over-represented in the results, all estimates of the derivative, G, and their slopes, are normalized by the mean of the y^th^ trajectory from, yielding and, respectively. With this normalization, yields of the value K_of_ from equation 7 for all trajectories and at all timepoints. contains count size normalized approximations of. Normalization is crucial to avoid size bias, and it is this specific normalization factor which permits a zero initial value, unlike fold changes. Normalizing by intensity average has precedent in M/A plot analysis, which uses a fold-change-like “ratio” normalized by average signal intensity(*10, 13*).

Where I is the complete list of trajectories, we observe that for a specific timepoint (x=k), the rates contained in, for all valid i ∈ I, fall into a normal distribution after outlier removal via a Grubbs test. Similarly, so do the values in. Statistical significance is assigned by z-score to each trajectory. Whether or not a trajectory was an outlier to the normal distribution, as well as it’s statistical significance in the distribution are recorded in the output file from DPoP. A visualization of the described data manipulation is included as figure 9.

A post processing step was included to remove trajectories with a low signal to noise ratio from the derivative estimation. Trajectories are excluded when the total change in the derivative estimation is not greater than some multiple of the average standard error of the linear regression across the trajectory. The multiple of the standard error may vary in practice, so the option has been programmed as adjustable in DPoP.

The final analysis yields two results at any point. First, the normalized linear approximations of the derivative,, yields rates which describe the omic trajectories locally, at each point. Second, the normalized slope of that linear regression,, describes the change in flux over the entire course of the trajectory. This local and regional estimate offers two features to examine, as opposed to fold changes alone.

DP is highly customizable and computationally tractable. A flow chart describing the algorithm is shown in Figure 10 where each arrow is a customizable operation within the overall workflow. At this point in the description of the methods, DPoP is at the “output lists” step of the flow chart and DP is completed. GO term enrichment analysis(*23*) can then be performed on the resulting list, identifying over and underrepresented descriptors within the subset as compared to the genome wide background in the .gaf file. These GO terms are output so that DPoP can programmatically describe the experiment in human interpretable words using just the aforementioned “output list” from DP.

How the data is preprocessed, differentiated, or normalized can change. Negative binomial distributions have been used to model transcriptomic data(*29*), and there may be a better distribution for the derivatives of other experiments. Database tools other than GeneOntology, (e.g., NetworKIN(*28*), GSEA2(*33*)), could be used on the output lists. The mathematics for variable timestep differentiation used as a universal differentiation technique for omics data is, to the best of the authors knowledge, a novel application of that algorithm, but another would do. There is flexibility the workflow. The fundamental underlying assumption of DP is that ALL dynamic omics experiments will have an omics derivative or flux feature to observe. DPoP presents the method in an easy to use and flexible platform to test this assumption broadly.

## Declarations

### Ethics approval and consent to participate

Not applicable.

### Consent for publication

Not applicable.

### Availability of data and materials

Two database files, three data sets and four analysis techniques are used in this work.

The two database files are the GO database .gaf files for A. nidulans and S. cerevisiae made available here(<u>LINK</u>), and here(<u>LINK</u>), respectively.

The three data sets are characterized and available as follows. The transcriptomic response of A. nidulans(*20*) is made permanently available via the Gene Expression Omnibus (GSE136562). The phosphoproteomic response of A. nidulans(*20*) is made permanently available via the Proteomics IDEntifications (PRIDE) repository, PXD015038. The phosphoproteomic response of S. cerevisiae(*22*) is made permanently via that works supplementary files.

The four differential analysis techniques included in this work are Volcano plotting, M/A plotting, MARS analysis, and DPoP analysis. Only DPoP is novel in this work. Scripts for volcano analysis were taken from <u>Matlab documentation</u>(*3*) and the code are available upon request. Scripts for M/A analysis were taken from <u>Matlab documentation</u>(*10-13*) and are available upon request. Scripts for MARS analysis utilize the ARESlab(*14*) <u>open-source MATLAB package</u> and are available upon request. DPoP is compatible with Windows only, and available under BSD 3 open source licensing from the <u>MathWorks file exchange link</u> as a developable .mlapp file, or as a plug and play .mlappinstall file.

All data sets used in this work are formatted for the app and included in the <u>MathWorks file exchange download</u> which contains all necessary files to use our app and reproduce the differential analysis and GO term analysis of the data. Descriptions of all files found at that download can be found in supplementary data S1.

### Authors’ contributions

H.E. conceptualized and implemented the work, built the app DPoP, oversaw testing/debugging and was the primary author/writer of the manuscript. J.Z found/identified the LaGrange interpolation differentiation algorithm as well as proofread/edited the manuscript. A.D. wrote the code for the MARS analysis and proofread/edited the manuscript. W.H. aided in UX testing, helped design the output file format, and proofread/edited the manuscript. C.S. performed the temporal GO term enrichment analysis and proofread/edited the manuscript. S.H. and R.S. were a CO-PI’s on the grant/project who proofread/edited the manuscript. M.M. is the main PI of this work and supported with funding and oversight throughout while also proofreading/editing the manuscript.

## Acknowledgements

The authors would like to acknowledge NSF, CBEE at UMBC, CBE at UConn, PPEM at IAState, and MATLAB for their roles in this work.

