## Supplementary File 1 for "Using flux theory in dynamic omics data sets to identify differentially changing signals using DPoP"

Supplementary Data S3.1 – Temporal GO Description of DPoP analysis of Phosphoproteomic Data 91

Note: Use the table of contents to navigate this document easier.

### Downloading and Using DPoP

DPoP is available on MathWorks File Exchange as an open source-addition to MATLAB. The app is only functional on Windows PCs. The app can be installed and run locally on your PC with the propper toolboxes, or it can be installed on MATLABs cloud-based version, MATLAB Online which will assure toolbox compatibility for you. Below is a description of all data files present for download from that file.

#### S1- Supplementary Information About App Files

| File name | Description |
|---|---|
| Derivative_Profiling_App_wGO_V1_3.mlapp | Developable app file |
| DPoP.mlappinstall | MATLAB app installable |
| AppFormatData_original_WT1405_dyn_mica_phospho_AverageBRs.xlsx | Dynamic phosphoproteomic time course of *A. nidulans* under micafungin, example Excel file. |
| AppFormatData_original_WT1405_dyn_mica_phospho_AverageBRs.mat | Dynamic phosphoproteomic time course of *A. nidulans* exposed to micafungin, example MATLAB file |
| AppFormatData_original_WT1405_dyn_mica_Transcript_Average3BRs.xlsx | Dynamic transcriptomic time course of *A. nidulans* exposed to micafungin, example Excel file |
| Aspgd.gaf | GO database reference file for *A. nidulans* |
| AppFormatData_Kanshin2015_Phospho12and3P.xlsx | Dynamic phosphoproteomic time course of S. cerevisiae exposed to high-osmolarity conditions, example Excel file |
| Sgd.gaf | GO database reference for *S. cerevisiae* |
| README.txt | Technical instructions for operating the DPoP app |
| License.txt | Matlab Open-source licensing |

### Supplementary data

#### S2 – Lists of Results from Ven Diagram Comparison

##### Supplementary data S2.1- Phosphoproteomic Ven Diagram

| Method of Analysis → | DPoP 452 Total | Volcano 583 Total | M/A 202 Total | MARS 1690 Total |
|---|---|---|---|---|
| List of Significant Signals ↓ (Uniprot ID, P-site) | C8V891 13 | Q5BF26 1569 | Q5B3R2 485 | C8V1M3 15 |
| | Q5B1A4 407 | Q5BE73 14 | Q5BG90 183 | C8V2A5 288 |
| | Q5B8I5 478 | C8VIY2 77 | C8VN11 458 | C8VFE1 145 |
| | C8VA49 440 | C8VKD7 39 | C8V2U2 930 | Q5ARD4 1009 |
| | Q5AYS0 68 | C8VS54 634 | Q5B3F1 474 | Q5AXV5 1535 |
| | Q5B8Z7 680 | C8VB27 585 | Q5B712 510 | Q5AZA6 358 |
| | C8V2A5 966 | Q5BC36 382 | Q5AYL5 912 | Q5B010 1082 |
| | Q5ARI0 237 | Q5BEP8 186 | Q5B6C4 72 | Q5B568 238 |
| | Q5AYU8 6 | Q5B2S6 294 | Q5B1A3 714 | Q5BCM2 9 |
| | Q5B4N2 6 | Q5AW27 447 | Q5B5J3 716 | Q5BGL6 88 |
| | Q5B5B0 637 | Q5BB58 155 | C8V5G6 290 | Q5B6C4 72 |
| | Q5B9N1 718 | Q5BG69 346 | Q5B4Q3 59 | Q5B8S4 253 |
| | Q5B8N0 34 | Q5B8B7 107 | C8V1W0 43 | Q5B8Z7 687 |
| | Q5B3S5 351 | C8VTS7 467 | Q5BCF3 241 | C8V5G6 35 |
| | C8V858 304 | Q5B795 127 | Q5BBS5 166 | Q5B6U5 188 |
| | C8VA69 37 | Q5BD26 48 | C8VTF1 86 | Q5B8I5 507 |
| | Q5B5B0 628 | Q5BGL6 88 | Q5AWK9 413 | Q5BDW4 11 |
| | C8V794 106 | Q5B3B7 553 | Q5BBV0 42 | Q5AZQ2 1176 |
| | C8V5G6 825 | Q5BDK0 622 | C8V742 377 | Q5B0E9 631 |
| | Q5B4R9 231 | Q5BDK0 623 | Q5B5B0 640 | Q5BDP0 15 |
| | Q5AZL2 332 | Q5AUG3 738 | C8V4I7 152 | Q5AVG6 153 |
| | Q5B6C1 816 | Q5BDK0 619 | G5EB50 298 | Q5AYN2 312 |
| | Q5AVT6 40 | Q5B6P5 272 | Q5BCF3 236 | Q5AZQ2 1367 |
| | Q5B950 38 | Q5B4L2 704 | Q5B6U5 67 | Q5B0E9 533 |
| | C8VIG9 368 | Q5B7V5 156 | Q5B1T1 978 | Q5B1Y7 268 |
| | Q5B3I8 940 | Q5B3F1 177 | C8VFE5 39 | Q5B425 520 |
| | C8V9I8 159 | Q5B0E2 557 | Q5B161 99 | Q5BFV5 804 |
| | Q5B907 344 | Q5BG67 436 | C8VFE5 37 | Q5BG68 400 |
| | Q5AZJ6 654 | C8V1V6 92 | Q5BCN3 367 | C8V7N4 122 |
| | Q5B587 1274 | Q5AZL8 69 | Q5B3C8 863 | C8VTS7 80 |
| | Q5AXG7 637 | Q5B5J2 243 | Q5B5K9 5 | Q5B2V4 400 |
| | Q5B8S4 206 | Q5B194 160 | C8VR18 741 | Q5BE66 526 |
| | C8V9I8 156 | G5EB09 517 | C8V1P9 255 | Q5AYS0 62 |
| | Q5BBY2 145 | Q5BH96 83 | Q5AYS4 173 | Q5BB65 37 |
| | C8VK54 154 | Q5AVM1 16 | C8V1M3 23 | C8V7M9 21 |
| | Q5BAD8 403 | C8VMA9 797 | Q5B2V4 400 | Q5AYL5 1466 |
| | Q5ARL7 517 | Q5BFV7 573 | Q5B2N3 1047 | Q5B161 65 |
| | Q5BC55 138 | Q5BGS7 2 | Q5B2N3 1045 | Q5BB41 12 |
| | Q5B9K4 218 | Q5B7M3 347 | C8VTS7 468 | Q5BB49 499 |
| | Q5B434 412 | Q5ASC1 501 | C8V8Q8 95 | C8VFD0 151 |
| | Q5AU19 1634 | Q5AW57 11 | Q5BBV0 49 | C8VN52 82 |
| | Q5B6T4 34 | Q5AYP0 459 | Q5B436 312 | Q5AQI2 156 |
| | C8V766 31 | G5EB26 44 | C8V2U2 925 | Q5ASM0 275 |
| | Q5B7N3 398 | C8V1G2 44 | C8VJ53 455 | Q5AU54 54 |
| | C8V5G6 911 | Q5B434 194 | C8VJ53 453 | Q5AUF4 667 |
| | Q5BFQ3 83 | Q5AZW6 1033 | Q5B424 1764 | Q5B8I8 336 |
| | Q5AXK8 3 | Q5AS56 647 | Q5BDZ7 386 | Q5BCF3 52 |
| | Q5BH07 345 | Q5BAZ5 11 | Q5ARJ8 421 | Q5AZQ2 414 |
| | Q5B2S6 1292 | C8VBH5 517 | Q5ARJ3 451 | C8V2A5 966 |
| | Q5B712 638 | Q5B3E4 169 | C8V0C1 405 | C8V6A1 382 |
| | Q5B5B3 276 | Q5B152 214 | Q5AWR8 1457 | C8V7V5 365 |
| | Q5B1E9 545 | Q5B1U5 988 | C8VB12 318 | Q5AXG7 637 |

| | | | | |
|---|---|---|---|---|
| | Q5AXT4 560 | C8VRL9 111 | Q5BG69 344 | Q5B161 99 |
| | C8VAQ0 92 | Q5BF78 222 | Q5BF26 355 | Q5BBJ4 992 |
| | C8V9B5 76 | Q5BCK1 190 | Q5AZW5 301 | Q5BG55 653 |
| | Q5AYF4 233 | C8V1M3 3 | Q5AYL5 940 | Q5ASA5 697 |
| | C8V2Y7 537 | Q5BCF3 223 | Q5B8F6 38 | Q5AU61 541 |
| | C8VHB5 844 | C8VU21 214 | C8VMW9 615 | Q5AW75 1345 |
| | Q5B609 7 | C8VU21 210 | Q5BBJ8 139 | Q5B5B5 802 |
| | Q5AVJ9 255 | Q5AW90 107 | G5EB64 447 | Q5BB56 75 |
| | Q5BF63 15 | C8VJB5 595 | G5EB64 449 | G5EAY1 780 |
| | Q5B1Y4 207 | Q5AW39 214 | Q5BCQ6 58 | G5EB13 462 |
| | Q5B3F1 553 | Q5BFF5 372 | C8VBH5 520 | Q5AXQ8 298 |
| | Q5AUF4 511 | Q5B1I8 579 | C8VGM0 582 | Q5B195 155 |
| | C8V000 404 | Q5AWR8 832 | Q5B0E2 551 | Q5B4D3 124 |
| | Q5B9Y3 11 | Q5BBW4 2879 | C8VKT1 230 | Q5BCM2 8 |
| | Q5B0H4 86 | Q5BBJ4 909 | Q5B0E9 757 | Q5BF78 224 |
| | Q5ARD4 1049 | Q5B5V3 423 | Q5ASM0 278 | C8V3U9 222 |
| | Q5B9D0 351 | C8VFE1 330 | P17120 1006 | C8V7G9 224 |
| | Q5BH36 151 | Q5B568 347 | Q5BBZ5 117 | C8VUL0 87 |
| | Q5ARL7 563 | Q5AYC1 750 | Q5BB17 273 | Q5AZI9 288 |
| | Q5BB02 1207 | Q5B5B3 53 | Q5B8Z7 964 | Q5AZM3 1067 |
| | C8V858 302 | Q5B1D1 1051 | C8VUS6 2 | Q5B3E2 107 |
| | Q5B8G8 101 | C8VHJ1 737 | Q5B7M5 12 | Q5B4Q3 48 |
| | C8VGM0 578 | Q5BF78 209 | Q5B7M5 14 | Q5B6T4 44 |
| | C8V9K6 547 | C8VJ33 51 | Q5B1I8 579 | Q5B950 39 |
| | Q5AYF4 235 | C8VP26 526 | Q5BEG5 163 | C8V8P5 303 |
| | Q5BBY2 1295 | Q5BG91 506 | Q5ASM0 287 | C8VBH5 1039 |
| | Q5BEQ2 382 | C8VRW7 323 | Q5B0H6 53 | Q5AYN6 11 |
| | G5EAT8 216 | C8VCR2 304 | Q5BA83 18 | Q5B7N3 222 |
| | Q5B797 8 | Q5BEL2 754 | G5EAT8 216 | Q5B8J0 119 |
| | Q5B154 416 | Q5BC22 761 | Q5AXU1 22 | Q5B8J0 126 |
| | C8VKW2 600 | Q5BGR9 11 | Q5AXU1 24 | Q5BE05 264 |
| | Q5B7M3 347 | C8V9W5 149 | Q5B5B3 55 | Q5BE20 686 |
| | Q5BEF1 656 | C8V1P9 255 | Q5B6L5 563 | C8VUP2 151 |
| | C8VLU9 54 | Q5B0V1 657 | Q9P931 114 | C8V2Y7 525 |
| | O59922 406 | Q5B6L5 563 | Q5BB56 525 | C8VM09 193 |
| | C8VBH5 871 | Q5BG69 345 | Q5BC84 288 | Q5ARZ1 217 |
| | C8VFJ6 214 | Q5B6C0 83 | Q5B123 426 | Q5AZB2 6 |
| | Q5AXK8 201 | Q5AWR8 438 | C8VQV4 354 | Q5B4H2 167 |
| | C8V3U9 223 | C8VP27 452 | C8VUD8 52 | Q5BFB7 1112 |
| | Q5AUH4 1054 | C8VUP2 151 | Q5B4D3 115 | C8V4E0 348 |
| | Q5AQD2 137 | Q5B1F5 112 | Q5BBY2 1411 | C8VHJ2 136 |
| | C8VBI4 5 | Q5BBK4 686 | Q5AYC1 394 | C8VUL0 83 |
| | Q5AYY3 75 | Q5B3F1 293 | Q5B8M0 759 | Q5BBK4 801 |
| | Q5BBU6 22 | Q5BD34 420 | Q5AVD1 181 | Q5BHA7 455 |
| | Q5B4B7 700 | C8V097 361 | Q5BHA9 518 | Q5B0F7 450 |
| | Q5B8G1 164 | Q5BB58 162 | Q5B178 340 | C8V9G4 45 |
| | Q5BCH1 89 | C8VUP2 502 | C8V3Z7 84 | Q5AX03 132 |
| | Q5AWR8 966 | Q5BGE6 228 | Q5BGL3 846 | Q5AZQ2 534 |
| | Q5B549 302 | Q5B193 238 | C8VRX2 36 | Q5BDQ5 743 |
| | Q5B8R6 58 | G5EAU0 317 | Q5B7N3 267 | C8VAY4 143 |
| | Q5ATW1 372 | Q5BGR2 34 | Q5AZC6 1215 | C8V3U9 227 |
| | Q5AUF4 606 | Q5B3C8 685 | Q5B3V8 541 | C8VMA9 797 |
| | C8V371 285 | Q5AU61 541 | Q5B0Q8 63 | G5EB20 225 |
| | C8VRP6 418 | C8V2U2 494 | Q5B8X0 401 | P60204 82 |
| | Q5BFG5 660 | Q5B9C6 142 | Q5BGN7 40 | Q5ASC1 501 |
| | C8VUL9 97 | Q5BB13 580 | C8V179 54 | Q5BDL3 31 |
| | Q5ASA5 828 | Q5B5Z2 126 | Q5AVV8 1503 | Q5BEN0 11 |
| | Q5BG84 427 | C8VTT9 279 | C8VS08 1041 | Q5BHA9 319 |
| | P11837 600 | Q5BDX2 557 | Q5B3H8 268 | C8VTF1 80 |
| | Q5BGR2 762 | Q5AYV6 123 | Q5BBY2 1004 | G5EAY1 1436 |
| | C8V2A5 291 | C8VC15 23 | C8VB99 399 | Q5AUF4 1869 |
| | Q5BCM2 86 | Q5BFF5 319 | Q5BF26 356 | Q5AVQ9 458 |
| | C8VMG3 120 | C8V2U2 182 | Q5B6C0 83 | Q5B5F7 301 |
| | G5EAZ0 318 | C8VA69 48 | Q5AWR8 209 | Q5B7M3 347 |
| | Q5BCF7 374 | C8VA69 41 | Q5BBF2 748 | Q5BDC7 601 |
| | Q5BEV7 680 | C8V008 1624 | Q5AVN0 26 | Q5B2W5 138 |

| | | | | |
|---|---|---|---|---|
| | Q5B1E9 550<br>Q5B4W4 148<br>C8VA91 154<br>Q00757 6<br>C8VN24 360<br>Q5B549 301<br>Q5B434 174<br>Q5BGM8 283<br>Q5B9C3 473<br>C8VGM0 109<br>C8V0Z9 527<br>Q5BB18 314<br>C8VN24 364<br>Q5BH98 267<br>Q5AW75 36<br>C8V008 1659<br>Q5AU20 715<br>Q5AUM0 630<br>Q5B3R2 476<br>C8VDC9 831<br>Q5AXI3 165<br>Q5BEV8 255<br>Q5BCB0 148<br>C8VA69 35<br>Q5B8J0 128<br>Q5BE23 603<br>Q5B0E9 676<br>C8VUP2 238<br>Q5BCP0 113<br>Q00083 582<br>Q5BFM3 187<br>Q5BGL3 854<br>C8V9Z0 54<br>C8V097 715<br>Q5B9K7 94<br>Q5AS91 692<br>Q5B4N4 529<br>Q5ARW9 216<br>Q5AWR8 1043<br>C8VFE1 422<br>C8VHD0 141<br>Q5AS85 430<br>Q5AW75 35<br>Q5BEL2 533<br>Q5AS84 440<br>Q5BAD4 245<br>G5EB07 163<br>C8V5G6 39<br>Q5AXD2 46<br>C8VQM0 1079<br>Q5B4N4 161<br>Q5BDD7 501<br>C8VHB5 845<br>Q5B6C4 72<br>Q5BAY0 738<br>Q5AS72 54<br>C8VUP2 502<br>Q5AXU1 22<br>C8V007 136<br>C8VQL1 737<br>Q5AZI9 288<br>Q5AZW4 402<br>Q5BC57 214<br>Q5B4L1 541<br>Q5BEE1 336<br>G5EB41 14 | Q5AYN2 10<br>Q5AQG3 585<br>Q5BDX5 89<br>Q5B8H6 673<br>Q5B4W4 148<br>Q5AW26 844<br>C8VIC5 34<br>C8VIZ5 10<br>Q5BBY2 140<br>Q5BHA8 250<br>Q5B8V1 683<br>Q5AW72 220<br>Q5AS71 352<br>Q5BAA8 397<br>Q5AZK5 114<br>Q5BBK4 172<br>Q5ATW1 290<br>Q5B1A4 407<br>Q5B0Q8 16<br>C8V312 220<br>Q5B9C3 436<br>Q5AVV8 665<br>Q5AVN0 383<br>Q5AW75 49<br>Q5B8W5 53<br>Q5BE17 392<br>Q5BG67 486<br>Q5BF43 64<br>Q5B6C3 920<br>Q5B104 218<br>Q5B1P4 324<br>Q5B4Q8 274<br>Q5B0Y0 68<br>Q5BDC7 76<br>Q5ARH8 405<br>C8VNH4 247<br>Q5BH36 525<br>Q5BH36 527<br>C8VRW7 17<br>Q5ATX5 1298<br>Q5B093 217<br>Q5B3E4 168<br>Q5BFG5 511<br>Q5BAE0 2<br>C8V007 144<br>C8V007 148<br>Q5BFG5 513<br>C8VN24 294<br>C8V9T1 1016<br>Q5BC69 29<br>Q5B8A0 1765<br>Q5BEK5 397<br>Q5B5B5 791<br>C8VFJ6 233<br>C8VAQ7 383<br>C8VKT1 227<br>Q5BBW4 1205<br>C8V1S9 380<br>C8VIZ5 12<br>Q5B4G0 43<br>Q5AZN9 197<br>Q5B3Q0 331<br>C8VUS6 56<br>Q5AZU0 9<br>Q5B6S5 148<br>Q5BF78 192 | Q5ASA5 697<br>C8VQ27 2<br>Q5B7A2 62<br>Q5B7N3 412<br>C8VC15 25<br>C8VN52 80<br>Q5BDU2 184<br>Q5AS84 440<br>C8V9Z0 48<br>Q5BGP1 21<br>Q5AYN6 9<br>Q5B613 65<br>P56859 22<br>Q5BE72 1290<br>Q5BEE9 163<br>Q5BBK4 701<br>Q5AZH7 290<br>C8VRY6 120<br>Q5AZQ2 1367<br>Q5AYT7 213<br>C8V7F2 2<br>Q5B0H6 45<br>Q5B0H6 44<br>Q5BAE0 291<br>Q5AYL5 399<br>Q5B4W2 214<br>C8V7N4 120<br>Q5AYL5 402<br>Q5B5V7 7<br>Q5B9P6 302<br>Q5B0H6 46<br>Q5B1L2 6<br>Q5B2R8 309<br>C8VBH5 677<br>Q5B0E9 571<br>Q5BDB4 548<br>Q5BG69 346<br>C8VUL9 97<br>Q5AYS0 36<br>Q5B1A3 713<br>Q5B5J1 4<br>Q5ASM0 275<br>C8V0Z9 527<br>Q5BE74 829<br>Q5BEI5 48<br>Q5B104 83<br>C8VLW6 22<br>Q5AXT7 181<br>G5EAZ0 339<br>Q5BHB3 23<br>C8V1G2 543<br>Q5AS94 792<br>Q5AS94 780<br>C8V000 160<br>Q5AYL5 910<br>Q5BEN1 353<br>C8VGI5 1019<br>C8VR18 548<br>Q5B5K9 482<br>Q5AYX2 198<br>C8VRY1 378<br>Q5B3I8 645<br>Q5B411 469<br>C8VQ48 173<br>Q5B0E9 782<br>O93937 14 | Q5B608 621<br>C8V9A6 117<br>P0C5H8 213<br>Q5AW50 633<br>Q5BFF5 374<br>Q5AWR8 832<br>Q5AYF4 235<br>Q5B1A3 874<br>Q5BF99 141<br>Q5AYC8 8<br>Q5AYF4 23<br>Q5AZL8 90<br>Q5B2Q1 453<br>Q5BCF3 44<br>Q5BF43 502<br>Q5BGV2 46<br>C8V0C1 402<br>Q5AW50 631<br>Q5B1U5 1244<br>Q5BG55 788<br>Q5BHA9 512<br>Q5AX75 70<br>Q5AYS0 63<br>Q5B104 66<br>Q5B4N4 533<br>Q5B5Z6 146<br>C8VUL9 100<br>Q5AVF3 436<br>Q5AXS5 79<br>Q5B1L6 88<br>Q5B425 1004<br>Q5B566 594<br>Q5B692 50<br>C8VK54 286<br>Q5AUL7 602<br>Q5B063 13<br>Q5B1K2 676<br>Q5BBK9 584<br>Q5BDL3 25<br>Q5BDB4 501<br>C8VTS7 468<br>Q5AXH6 78<br>Q5AYF5 413<br>Q5B0H3 3<br>Q5B534 395<br>Q5AU20 325<br>Q5AVV8 649<br>Q5B441 280<br>Q5B5B0 1230<br>Q5BDC7 82<br>Q5BHA9 518<br>C8VA31 417<br>Q5B706 60<br>C8V9A6 99<br>C8VP26 299<br>Q5B224 62<br>Q5B224 1544<br>Q5B452 4<br>C8VMX1 51<br>C8VRX2 53<br>Q5AS56 647<br>Q5B2S6 234<br>Q5B534 19<br>Q5B7M3 316<br>Q5BDC7 632<br>Q5BEC4 262 |

| | | | | |
|---|---|---|---|---|
| | Q00083 1191 | Q5AVG3 279 | Q5BBK4 400 | C8V094 41 |
| | C8V2U2 925 | Q5B4Q3 25 | Q5BBK4 398 | P17429 642 |
| | Q5BF43 556 | Q5BEL2 751 | C8VBJ9 641 | Q5AVG6 144 |
| | C8VCL7 288 | Q5B377 10 | Q5B9L0 345 | Q5AXG1 124 |
| | C8VFD3 312 | Q5B377 11 | Q5BDY6 115 | Q5AYM6 290 |
| | Q5B5V3 460 | C8VHC5 204 | Q5B2S6 965 | Q5B4Q3 26 |
| | Q5BES9 1695 | Q5B3I8 940 | C8VA31 355 | Q5BAN3 611 |
| | C8V749 80 | C8VSE7 137 | Q5AVV4 688 | Q5BE66 112 |
| | P56859 15 | Q5ARD4 1049 | Q5B411 466 | G5EB06 392 |
| | Q5B425 588 | Q5BBY2 1310 | Q5B9K4 205 | Q5B2M5 592 |
| | C8VDC6 58 | Q5AYU2 80 | C8V2U2 1507 | Q5AZW5 298 |
| | Q5B215 196 | Q5BEI5 52 | C8V8Q8 96 | Q5B6C1 750 |
| | Q5BH98 265 | Q5B566 348 | C8VKT7 112 | Q5BF99 846 |
| | C8VAY4 261 | Q5B1L2 11 | C8VKT7 114 | Q5ATZ3 228 |
| | Q5BAL1 413 | C8VHD2 1135 | Q5BAJ7 90 | Q5B1K7 63 |
| | Q5B674 12 | Q5AVL0 247 | Q5BGL3 111 | Q5B5G9 999 |
| | C8VK49 1310 | C8VAQ0 92 | C8VA31 280 | Q5B8I8 334 |
| | Q5BGE6 304 | Q5B9P0 120 | | Q5BA33 66 |
| | Q5BF99 129 | Q5AWR8 1047 | | Q5BB06 395 |
| | Q5B1U5 1248 | Q5BBW4 309 | | Q5BBZ5 117 |
| | C8V4L4 930 | Q5B9P0 123 | | C8V2Y6 544 |
| | Q5BBD7 15 | Q5B8I5 510 | | C8VG47 117 |
| | Q5BFG5 657 | G5EAY1 1005 | | Q5AS85 5 |
| | Q5BF43 546 | Q5AW79 10 | | Q5AVW7 713 |
| | Q5AYC1 147 | Q5AZQ2 484 | | Q5B178 189 |
| | Q5BDD7 1287 | Q5AZC9 261 | | Q5B436 47 |
| | C8V3Q8 871 | Q5AWR8 436 | | Q5B4N2 58 |
| | Q5B1L2 6 | C8VFE1 334 | | Q5B9C3 520 |
| | Q5B606 551 | Q5BF26 403 | | Q5BCQ6 220 |
| | Q5B0B8 434 | Q5B0E9 344 | | Q5BDK5 359 |
| | Q5AXK0 259 | Q5BD26 49 | | Q5AUJ5 449 |
| | Q5AUM0 647 | Q5BB56 5 | | Q5B706 27 |
| | Q5AVQ0 109 | Q5BGE6 182 | | C8V494 12 |
| | Q5B010 200 | Q5BEZ7 171 | | P22022 430 |
| | C8V2A5 133 | Q5AZC6 1211 | | Q5B0P4 484 |
| | Q5BEV8 250 | Q5BE59 793 | | Q5B4I8 315 |
| | Q5BDQ5 743 | G5EB39 317 | | Q5AXH5 107 |
| | C8VKT6 373 | C8V3Z7 67 | | Q5B0E9 829 |
| | Q5BAD4 251 | Q5AVG3 229 | | Q5B3H7 11 |
| | Q5BBL0 235 | Q5B441 282 | | Q5B5J1 108 |
| | Q5B4W2 251 | Q5B2M5 73 | | Q5BCP6 101 |
| | Q5BF72 429 | Q5BBE5 313 | | Q5BGL3 701 |
| | Q5BHA9 362 | C8VCK1 302 | | Q5BGR2 34 |
| | C8VQ21 263 | C8VJ09 207 | | Q5BH53 286 |
| | Q5B0E9 344 | Q5B367 100 | | C8UZY7 13 |
| | C8VUG5 6 | Q5BEV8 251 | | C8V0Z7 231 |
| | Q5AYL5 823 | Q5BD94 266 | | C8VQJ9 905 |
| | Q5BBK4 701 | Q5B5R8 340 | | Q5B178 188 |
| | Q5BG69 344 | C8V007 819 | | Q5BDQ5 614 |
| | Q5B5R8 74 | P18696 16 | | Q5BFP3 58 |
| | Q5B0E2 1125 | C8UZY5 122 | | C8V008 1516 |
| | Q5BCF7 69 | C8V007 2798 | | C8V9W6 331 |
| | C8VG45 454 | C8V007 2794 | | Q5BBF0 749 |
| | Q5B0X8 165 | C8V996 8 | | Q5B224 1512 |
| | Q5B587 40 | Q5AVN0 23 | | Q5AT95 179 |
| | Q5B7A2 62 | Q5B674 16 | | Q5B9C6 293 |
| | Q5B1Y4 16 | Q5ATW1 449 | | C8V8H5 411 |
| | Q5B0E9 571 | G5EAY1 1423 | | P38093 418 |
| | Q5B608 619 | C8VSX4 166 | | Q5B3F1 848 |
| | C8VRY6 122 | Q5BGP1 23 | | Q5BH85 154 |
| | G5EAU0 315 | Q5BF43 170 | | Q5B4F9 348 |
| | Q5BCI4 28 | Q5B9N1 967 | | C8VNZ3 130 |
| | Q5AVN0 536 | C8VFJ6 235 | | Q5AS85 189 |
| | Q5B8L6 200 | C8V7V5 406 | | Q5AS91 413 |
| | C8VS08 4 | Q5B8W7 621 | | Q5B5Z2 395 |
| | Q5BGM8 287 | Q5BEM6 96 | | Q5BB02 1177 |

| | | | | |
|---|---|---|---|---|
| | G5EB90 52 | Q5AZU0 10 | | Q5B1T1 1626 |
| | Q5BDB5 50 | Q5BBY2 845 | | C8V0C1 461 |
| | Q5BDC7 140 | C8VRW7 93 | | C8VD17 5 |
| | Q5BC59 257 | C8VIQ3 438 | | Q5ASA7 377 |
| | C8V0J3 2 | Q5B5H4 451 | | Q5BE59 257 |
| | Q5AVN0 26 | C8VHD2 649 | | C8VAK9 51 |
| | Q5BBK4 397 | C8VHD2 652 | | Q5AYL5 441 |
| | Q5B1L6 88 | Q5B9M1 18 | | Q5B7N5 649 |
| | Q5BDG0 517 | Q5B705 172 | | Q5B9C6 298 |
| | Q5B7M3 345 | C8VB99 358 | | Q5BGV2 797 |
| | C8VQM0 591 | Q5BBK8 97 | | Q5BAE0 6 |
| | Q5B3H7 47 | Q5B5K9 500 | | Q9P4R5 467 |
| | Q5BBL2 62 | Q5BEP3 12 | | Q5B7V5 129 |
| | Q5BEU9 177 | Q5BEU9 179 | | C8VK54 266 |
| | Q5B4I8 84 | Q5ASM0 289 | | Q5B010 378 |
| | C8V0Z9 538 | Q5AS72 384 | | Q5B0B8 155 |
| | Q5AVK2 21 | Q5B511 359 | | Q5B1T1 978 |
| | Q5B434 964 | C8VP26 125 | | Q5BC57 214 |
| | Q5BF43 80 | Q5BDF8 163 | | Q5BCM2 110 |
| | Q5AU19 1623 | Q5BBH4 84 | | Q5BDK5 26 |
| | Q5ASM0 287 | Q5BDE3 5 | | Q5BDU2 49 |
| | C8VUB4 787 | G5EAY1 642 | | Q5BFF5 370 |
| | Q5BCG6 96 | G5EAY1 649 | | Q5AVD1 281 |
| | Q5BDK0 657 | G5EAY1 646 | | Q5B3I8 100 |
| | Q5BFX3 1038 | Q5B3X0 95 | | Q5BG68 337 |
| | C8VN52 80 | Q5B5J2 848 | | Q5BH85 98 |
| | C8VSE7 137 | Q5B3G8 303 | | C8V0Z9 485 |
| | Q5B1K7 153 | C8VUL9 345 | | C8V4L4 654 |
| | C8V116 61 | Q5AXG9 68 | | C8VQ45 563 |
| | C8VBH5 744 | Q5AXT8 510 | | Q5AUF4 511 |
| | Q5AYS4 173 | C8VJD9 274 | | Q5AWG3 343 |
| | Q5AVN0 306 | Q5AXV2 9 | | Q5B1Y2 331 |
| | C8V000 405 | Q5BB69 65 | | Q5B3S5 230 |
| | Q5BCP4 336 | Q5B6I6 788 | | Q5B5K9 6 |
| | Q5B161 61 | C8V9W5 154 | | C8V9T1 573 |
| | Q5B2S6 962 | Q5AVN0 644 | | Q5B018 191 |
| | Q5BFH3 452 | Q5AVN0 741 | | Q5BCP0 13 |
| | Q5AZL8 86 | C8VRM6 23 | | Q5BDZ7 572 |
| | Q5AXK8 197 | Q5B8J6 458 | | G5EB54 2 |
| | C8VH46 159 | C8V2A5 572 | | Q5B069 87 |
| | Q5AT83 24 | C8VJ88 314 | | Q5BBV0 452 |
| | Q5B7N3 401 | Q5AQL7 582 | | Q5BEU9 18 |
| | Q5B3F1 38 | C8VKH5 630 | | C8V9G4 121 |
| | Q5B7V5 142 | Q5AVC0 433 | | C8V9Z0 52 |
| | Q5BCP0 13 | Q5B7J9 6 | | C8VUP2 195 |
| | Q5AS94 512 | Q5B5L1 219 | | Q5B9L0 345 |
| | Q5B568 120 | Q5BEZ7 187 | | Q5AYL5 943 |
| | G5EB64 495 | Q5BFC7 373 | | Q5AZQ2 481 |
| | Q5AZZ8 219 | C8VGA0 635 | | Q5B3H8 303 |
| | C8VAQ7 383 | Q5B411 468 | | Q5BDA5 172 |
| | Q5B9A8 573 | Q5AZM3 936 | | G5EB77 15 |
| | Q5AUG3 368 | Q5B8G1 182 | | Q5AZZ8 219 |
| | Q5B6T4 38 | C8VU51 1069 | | C8VQT9 265 |
| | Q5B058 72 | Q5BDX6 221 | | Q5B7N3 398 |
| | C8VCW1 213 | C8V5G6 285 | | C8VHJ7 371 |
| | Q5AX75 193 | Q5AS85 436 | | Q5AU20 1193 |
| | C8VTV4 374 | C8V0A1 325 | | Q5BHB3 13 |
| | Q5BAD4 248 | C8VB25 31 | | Q5AZJ6 577 |
| | Q5B5B3 275 | Q5BF24 21 | | C8V094 331 |
| | C8VJ53 455 | Q5AYS0 33 | | C8V3J2 84 |
| | Q5B8R7 468 | Q5B797 480 | | Q5BCQ6 804 |
| | Q5BHA9 357 | Q5B4D3 134 | | C8VCW1 213 |
| | Q5B5F7 2183 | Q5B7V5 142 | | Q5AUJ9 648 |
| | Q5B4S4 498 | C8V1G2 169 | | Q5AWN0 441 |
| | Q5B4L2 574 | Q5B5B7 404 | | Q5B7V5 128 |
| | C8VJD2 761 | Q5BEZ2 180 | | Q5BBW4 2879 |

| | | | | |
|---|---|---|---|---|
| | C8V2A5 590<br>Q5BG84 429<br>Q5BAX8 70<br>C8VFE1 424<br>C8VJ72 136<br>Q5AU15 358<br>Q5BH85 98<br>G5EB07 160<br>Q5AVN0 544<br>C8VRX2 96<br>C8VHC5 1461<br>Q5BGL6 95<br>Q5BFG5 230<br>Q5B0H6 51<br>Q5AUH4 1050<br>Q5B2M5 73<br>C8V000 426<br>Q5B566 591<br>C8V2U2 557<br>Q5B6C3 124<br>Q5AWK4 2<br>Q5BBY2 642<br>Q5B5R8 127<br>C8VF70 16<br>Q5BFQ3 79<br>Q5B5F1 353<br>Q5B1Q9 822<br>Q5B6C3 8<br>C8VTF1 86<br>Q5B425 1021<br>Q5AU20 315<br>C8VLZ4 106<br>G5EB05 80<br>P0C153 8<br>Q5AXT8 510<br>Q5AXV5 715<br>Q5B7J9 102<br>C8V007 401<br>Q5B9C6 461<br>Q5BCN3 271<br>Q5B9L0 457<br>Q5B434 217<br>C8VK49 1290<br>Q5ASC1 508<br>Q5BCP0 9<br>C8VUU8 277<br>C8V7N4 120<br>Q5B5F1 351<br>C8VIC6 114<br>Q5AVV8 1931<br>C8V2U2 933<br>Q5BH83 908<br>Q5B8M8 126<br>Q5B916 183<br>C8VMQ0 440<br>Q5ARD4 1033<br>Q5BDK2 21<br>Q5BH07 349<br>Q5AYX3 754<br>Q5BBY2 661<br>Q5B6C3 7<br>Q5B4L8 151<br>Q5BCN3 288<br>C8V1M3 15<br>Q5BGL3 718<br>Q5BDW2 930 | Q5BC84 283<br>Q5B7M3 261<br>O59922 406<br>C8VA27 616<br>Q5B4N2 199<br>Q5AZQ2 292<br>P36011 423<br>Q5B7N3 452<br>Q5AVE9 1014<br>Q5AVE9 1017<br>Q5BG55 789<br>Q5AZQ2 381<br>C8V2U2 118<br>Q5BEJ2 72<br>Q5ASD3 25<br>C8V2Y7 532<br>Q5BG25 377<br>Q5B6S5 144<br>C8V002 234<br>C8V751 339<br>C8VT82 33<br>Q5AXX7 45<br>Q5B4N4 161<br>C8VRX2 35<br>Q5BDB5 47<br>Q5B0Q8 62<br>Q5AVZ2 789<br>Q5B950 38<br>Q5B3R6 392<br>Q5B0S3 397<br>Q5B1A4 420<br>Q5B1K7 259<br>C8VRW7 350<br>Q5ARJ3 75<br>Q5BEM6 92<br>Q5BEM6 90<br>Q5B0E9 730<br>Q5B9K7 94<br>Q5BGL3 713<br>C8V2U2 33<br>Q5B3R2 312<br>Q5B0V1 656<br>Q5BBK9 303<br>Q5BF43 556<br>Q5AZI5 313<br>Q5BAD4 795<br>Q5AVK2 512<br>Q5B6S0 243<br>Q5AVX6 10<br>C8VBH1 73<br>Q5AVY3 1482<br>C8VPX1 1095<br>Q5B8W5 18<br>C8V8H5 622<br>Q5B4K2 304<br>Q5B4P4 146<br>Q5BCN3 103<br>Q5B6T4 621<br>Q5AVX6 7<br>Q5BGE6 402<br>C8VIX9 3<br>Q5B2X2 240<br>Q5BB54 2045<br>Q5AVK2 419<br>Q5AYL5 404<br>Q5BEZ5 229 | | C8V8P5 295<br>C8VUS6 54<br>Q5AW90 107<br>Q5B5B0 1226<br>Q5B5Z6 18<br>Q5B8E7 127<br>Q5AVN0 753<br>Q5BEK5 357<br>Q5B436 271<br>C8V1W0 33<br>Q00757 6<br>Q5B7A1 388<br>C8VCK1 300<br>Q5AVN0 558<br>Q5BAA8 335<br>Q5BAE0 9<br>Q5BEQ3 115<br>Q5BFQ3 90<br>Q5AXT7 301<br>C8V139 384<br>C8V4D4 23<br>C8VBH5 517<br>C8VGH0 360<br>C8VUP2 584<br>Q5AZH7 1012<br>Q5B602 288<br>Q5BF43 173<br>C8V7V5 390<br>C8VLZ4 101<br>C8VUD8 52<br>Q5AVZ4 195<br>Q5AYQ5 234<br>Q5AYR3 388<br>Q5AZM3 1068<br>Q5B2V2 2<br>Q5B4N4 658<br>Q5B4Q3 52<br>Q5B362 327<br>C8V7N4 120<br>C8V9T1 719<br>Q5AVG3 234<br>Q5BDB4 380<br>Q5BDQ5 360<br>Q5BGB3 12<br>C8VR18 662<br>C8V007 821<br>C8VKH5 610<br>C8VRX2 101<br>Q5AYC1 390<br>Q5B178 194<br>Q5B9K9 14<br>Q5BBV0 225<br>Q5BBY2 427<br>Q5BDA5 369<br>C8V1P9 260<br>C8VIC5 387<br>C8VTZ0 183<br>Q5AS94 209<br>Q5B436 466<br>Q5BEN1 353<br>C8VTA5 113<br>Q5BBL0 390<br>C8V2R8 218<br>C8VB25 1089<br>C8VLZ0 265<br>Q5B7N3 267 |

| | | | | |
|---|---|---|---|---|
| | Q5BEN0 11<br>Q5AWH6 99<br>Q5B987 100<br>Q5BCM2 85<br>Q5AVV4 1173<br>C8V2A5 288<br>Q5B5R8 70<br>C8VS08 974<br>Q5AZF2 470<br>Q5AZQ2 528<br>Q5BBY2 664<br>Q5AYL5 1529<br>Q5AYS2 13<br>Q5B6C3 125<br>C8VKW2 450<br>Q5B2P4 195<br>Q5AW56 1015<br>P36011 423<br>Q5AUH4 1051<br>C8VJ53 453<br>Q5BBW4 1637<br>Q5AXF0 527<br>Q5B608 621<br>Q5B3H7 17<br>Q5B511 359<br>Q5BBM9 386<br>Q5B8V1 683<br>G5EB50 825<br>Q5AQM2 10<br>Q5B7N3 53<br>C8VT75 179<br>Q5AYC4 446<br>Q5AVV4 1172<br>C8V007 3030<br>Q5AQG3 585<br>C8VTZ5 8<br>Q5BGL3 701<br>Q5BAD4 254<br>C8VHD2 1135<br>C8V9T1 573<br>Q5AVW7 713<br>Q5B9C6 481<br>Q5B706 21<br>C8VCW1 225<br>C8V3I8 292<br>Q5AXU1 24<br>Q5BCB0 150<br>Q5AUG3 736<br>Q5AYN2 312<br>Q5BCP0 11<br>Q5B425 1024<br>Q5B0E9 629<br>Q5BBK0 151<br>Q5B0X8 167<br>C8V5G6 439<br>C8VDQ4 786<br>Q5BEP3 13<br>Q5BBY2 1310<br>Q00083 827<br>Q5AZK9 389<br>Q5B8H3 753<br>Q5ASC1 492<br>Q5B0E9 377<br>C8V9T1 568<br>C8VBH5 548<br>Q5B4Q3 173 | C8V2G4 126<br>Q5AS85 144<br>C8V3S3 54<br>C8V3S3 52<br>G5EB39 325<br>Q5BGL3 697<br>C8VJ78 78<br>Q5B4Q8 273<br>Q5B637 46<br>G5EB07 166<br>Q5B3F1 476<br>Q5BEL2 533<br>Q5BEL2 529<br>Q5B7Q7 137<br>Q5AXV5 715<br>Q5B7N3 220<br>Q5BG68 398<br>C8V2U2 557<br>Q5B2M6 251<br>Q5AVK0 190<br>Q5B8J3 261<br>Q5BB65 441<br>Q00202 516<br>Q5AUH4 971<br>Q5BDU2 187<br>Q5BDU2 75<br>C8VCK1 433<br>Q5BBY2 635<br>Q5AZL8 86<br>C8VKT1 226<br>G5EAZ0 330<br>Q5B9C3 470<br>Q5B7V5 128<br>Q5B8G1 185<br>Q5AZJ6 232<br>Q5BFG5 354<br>C8VP27 454<br>Q5BCF3 236<br>Q5BH68 1522<br>C8VBH5 520<br>Q5AZC6 1215<br>Q5BF43 1191<br>Q5AW92 1063<br>Q5AZE5 288<br>Q5AVG3 239<br>Q5BF43 67<br>Q5ARJ3 451<br>G5EB32 1206<br>Q5BH68 75<br>Q5BBH4 85<br>Q5AVV4 71<br>C8V7B0 10<br>Q5ARL7 559<br>Q5BB15 333<br>C8VS54 732<br>Q5ATW2 278<br>Q5B0Y5 757<br>C8V9B5 76<br>Q00647 1052<br>Q5BGP1 21<br>C8VG45 454<br>Q5AUF4 511<br>Q5BEM6 57<br>Q5B982 530<br>C8VF08 528<br>Q5AYQ5 279 | | Q5BAZ5 22<br>Q5BCI4 710<br>Q5BEV8 250<br>Q5BGS7 130<br>Q5BCF3 223<br>G5EAS6 856<br>Q5AS85 203<br>Q5B550 2<br>Q5B8R7 568<br>Q5BEB4 276<br>Q5BFV7 568<br>Q5B0E2 556<br>C8VCW1 2<br>C8VD16 108<br>Q5AVM1 16<br>Q5AVV4 1172<br>Q5B8I5 513<br>Q5BAG3 244<br>Q5BF43 62<br>Q5B9N1 719<br>Q5BH07 349<br>C8V7V5 280<br>Q5BEU9 15<br>C8VTS7 462<br>Q5ASK8 17<br>Q5B5B3 50<br>C8V097 708<br>G5EB62 15<br>Q5AVD1 18<br>Q5AZE9 700<br>Q5B5R8 524<br>Q5B950 40<br>Q5BAD8 144<br>Q5BCM2 112<br>Q5BDC6 49<br>Q5BBU6 3<br>C8VDC9 137<br>Q5B436 49<br>Q5B534 357<br>Q5BBM9 17<br>Q5BBW4 1203<br>Q5BH85 217<br>C8V4E0 50<br>C8V5G6 40<br>Q5BEZ2 306<br>Q5BH68 72<br>Q5AU65 267<br>C8V116 251<br>C8VCZ2 29<br>C8VHJ4 270<br>Q5AW72 56<br>Q5B4J3 555<br>C8V606 748<br>C8VFD0 147<br>Q5B0E9 377<br>Q5B9C3 510<br>Q5BAD4 254<br>C8V2Y7 526<br>C8VQJ9 904<br>Q5AUG9 313<br>Q5B0G4 403<br>Q5B122 384<br>Q5B5B0 640<br>Q5BB58 171<br>Q5AXH5 298<br>C8VHJ7 209 |

| | | | | |
|---|---|---|---|---|
| | Q5B2L4 584 | C8VTZ9 368 | | Q5B3B9 18 |
| | C8VHB5 848 | Q5BCN0 964 | | Q5B3B7 838 |
| | Q5BBK4 57 | C8V0Z9 486 | | Q5BAH3 440 |
| | Q5BFH3 526 | Q5AVV8 1503 | | Q5BGS7 215 |
| | | Q5B982 18 | | Q5BH46 532 |
| | | Q5B8Z7 862 | | C8VGA0 93 |
| | | Q5B5B0 640 | | C8VQM0 590 |
| | | C8V7N4 122 | | C8VRW7 341 |
| | | Q5BBY2 1402 | | C8VUL9 103 |
| | | Q5B7N3 454 | | Q5B3S5 234 |
| | | C8V2U2 488 | | Q5BE66 507 |
| | | Q5BBE5 330 | | Q5BH83 908 |
| | | C8VNZ3 2 | | Q5B1H7 160 |
| | | Q5AYC1 778 | | Q5B3I3 245 |
| | | C8V8P5 299 | | Q5B568 120 |
| | | Q5B3F1 899 | | Q5BCI0 399 |
| | | C8VTS7 462 | | Q5BDJ7 133 |
| | | C8V8Q8 45 | | C8V139 290 |
| | | G5EB07 169 | | C8V4L4 806 |
| | | Q5AV97 525 | | G5EB71 285 |
| | | Q5AS89 2 | | Q5BBD0 11 |
| | | C8VHC5 590 | | Q5BF26 1058 |
| | | Q5BBD7 1151 | | C8VTS7 78 |
| | | Q5BED1 371 | | Q5AWI8 629 |
| | | Q5B4N4 246 | | Q5BBY2 304 |
| | | Q5B5B5 784 | | C8VFD3 312 |
| | | C8VMX1 47 | | Q5B162 3 |
| | | C8V0V7 7 | | C8VQK3 5 |
| | | Q5BBJ8 399 | | C8VUA7 799 |
| | | Q5BDC7 600 | | Q5AYL5 823 |
| | | Q5AVY3 710 | | Q5AZC6 956 |
| | | C8VTS7 80 | | Q5B4W2 250 |
| | | Q5B5E4 3 | | Q5B7N3 409 |
| | | C8VPS9 11 | | Q5BF77 939 |
| | | Q5AVI4 574 | | Q5B0H6 44 |
| | | Q5B568 343 | | C8V742 377 |
| | | Q5B5K9 5 | | C8VF08 528 |
| | | Q5AUF4 508 | | G5EB66 16 |
| | | C8VHJ1 736 | | Q5AXW6 136 |
| | | Q5AZC9 505 | | Q5AYV1 19 |
| | | Q5B6P5 288 | | Q5BFV5 850 |
| | | Q5ASA5 85 | | Q5AW99 385 |
| | | Q5AZL8 93 | | Q9P4R5 575 |
| | | Q5AYC1 761 | | C8VHD2 235 |
| | | Q5BCR1 242 | | Q5B9A9 340 |
| | | Q5B5R8 409 | | Q5BDJ6 606 |
| | | Q5AVL9 141 | | Q5B8S4 259 |
| | | Q5ASA5 709 | | Q5AXF0 528 |
| | | Q5BEP4 152 | | Q5B9S9 1112 |
| | | C8VB27 568 | | Q5BB42 198 |
| | | Q5AUF4 543 | | Q5BCQ6 555 |
| | | C8V000 84 | | C8VJ53 453 |
| | | Q5B395 651 | | Q5B4S5 304 |
| | | Q5B716 163 | | Q5BE72 1213 |
| | | Q5BAJ7 98 | | C8V8D1 58 |
| | | Q5AWK5 26 | | C8V8L0 116 |
| | | C8V2G4 39 | | C8V9B5 44 |
| | | Q5BC69 11 | | Q5B2V3 35 |
| | | Q5B4W2 251 | | Q5B3J9 76 |
| | | C8VBH5 881 | | Q5BGR2 30 |
| | | Q5AT95 256 | | C8V4L4 930 |
| | | Q5B4F9 480 | | C8VHR2 353 |
| | | Q5B5K9 4 | | C8VJN4 140 |
| | | C8VBH5 677 | | Q5ASC1 532 |
| | | C8V1N7 320 | | Q5AVU3 271 |
| | | Q5AYN6 9 | | Q5AW04 33 |

| | | | | |
|---|---|---|---|---|
| | | Q5AYC1 777<br>Q5B2T9 362<br>Q5BEV8 259<br>C8V4D4 44<br>Q5B3I8 880<br>Q5AVV4 1173<br>C8VHJ1 732<br>Q5B6M4 4<br>Q5BBY2 709<br>C8V7Q0 127<br>Q5B323 487<br>Q5BF43 408<br>C8VQ48 272<br>Q5BGL3 701<br>Q5B3C8 697<br>Q5B1U5 914<br>Q5BAY0 736<br>Q5B5U1 272<br>Q5BFA9 57<br>C8VUP2 201<br>C8VA49 440<br>Q5B8X5 400<br>Q5AZF2 470<br>Q5B8N0 34<br>Q5BH68 72<br>Q5BH98 267<br>Q5AXH5 298<br>C8VEU7 516<br>Q5B434 174<br>Q5BAY0 738<br>Q5AYX3 754<br>C8V9Z0 54<br>C8VKH5 682<br>C8V8I3 201<br>C8V8I3 195<br>Q5B706 45<br>Q5BCM2 85<br>Q5BCM2 86<br>Q5BFG5 232<br>Q5B0Y5 770<br>Q5B6U5 345<br>C8VT05 103<br>Q5B6J6 39<br>Q5B4S4 517<br>Q5BFC7 60<br>Q5AZU3 416<br>Q5AYF4 235<br>Q00202 521<br>P38093 418<br>Q5B4K7 1768<br>Q5BCH1 89<br>Q5AUG3 53<br>Q5BDX5 97<br>Q5AYC1 149<br>Q5B154 415<br>C8VHJ2 470<br>C8V008 1516<br>Q5B1L6 88<br>Q5B5C5 453<br>Q5AZU0 11<br>Q5AXT7 234<br>Q5B0U1 68<br>C8VMQ0 439<br>Q5BCP6 82<br>Q5B6S0 245<br>Q5ARJ8 427 | | Q5B6U5 64<br>Q5BC59 261<br>Q5B706 268<br>C8VRY6 139<br>Q5B0V6 50<br>C8VFE5 39<br>C8VMF3 285<br>Q5ASM0 274<br>Q5AVJ9 261<br>Q5B0V3 221<br>Q5B7N3 278<br>Q5BAZ2 650<br>Q5BB49 183<br>Q5BDK2 52<br>Q5B6C4 1137<br>C8VHJ2 470<br>Q5AUF4 537<br>Q5AVG6 43<br>Q5B0Q8 63<br>Q5B323 43<br>Q5BBU0 441<br>Q5BCN0 956<br>C8V8P5 297<br>C8VGI4 193<br>C8VH61 25<br>C8VRY6 134<br>Q5B2R8 307<br>Q5B9A9 335<br>Q5BE74 827<br>Q5B1L6 29<br>Q5BBY2 1411<br>Q5BFF2 250<br>Q5B5B3 132<br>Q5BCG6 93<br>Q5BCR1 242<br>Q5BE59 625<br>Q5B1D1 1051<br>Q5B6S0 297<br>Q5BEW9 180<br>C8VFE1 475<br>Q5B8R7 513<br>Q5BBK4 460<br>Q5BC28 249<br>Q5AYR3 390<br>Q5B1K1 871<br>Q5B9C6 143<br>Q6SIF1 383<br>C8V3S7 230<br>Q5B0X8 165<br>Q5BB61 97<br>Q5BBL3 110<br>Q5B2S6 965<br>C8VHQ9 136<br>Q5B558 690<br>Q5B4N4 246<br>Q5B2S6 334<br>Q5BH27 22<br>Q5BH27 46<br>C8V097 706<br>G5EB06 523<br>P0C8Q0 262<br>Q5AWT0 930<br>Q5B550 693<br>Q5BAY0 739<br>Q5BE20 165<br>Q5B982 183 |

| | | | | |
|---|---|---|---|---|
| | | C8VQV4 341<br>Q5BGT8 633<br>Q5BDC6 165 | | Q5BBV0 222<br>Q5B6L5 25<br>Q5B9K4 225<br>Q5BAD4 794<br>C8VGM0 362<br>Q5AVZ2 1079<br>Q5B010 1289<br>Q5BGP1 795<br>Q5BEP3 10<br>Q5BEW1 232<br>Q5BFG5 660<br>Q96V52 4<br>Q5B6K9 288<br>C8V794 225<br>Q5AS84 432<br>Q5BG67 489<br>C8V2U2 572<br>C8V2U2 120<br>Q5BBL5 1121<br>C8V5C5 615<br>G5EAX9 29<br>Q5AUF4 470<br>Q5B0H6 51<br>Q5B9C6 28<br>C8VCK1 296<br>Q5AXV0 149<br>Q5B6C3 125<br>Q5BBE4 633<br>Q5BBY2 650<br>C8VRX2 47<br>Q5BEJ2 71<br>Q5BFU0 2<br>C8VHJ7 377<br>C8VNZ3 2<br>P56859 32<br>Q5BB49 299<br>Q5BBJ4 888<br>Q5BFU5 457<br>Q5BGX9 103<br>C8V1M3 23<br>C8V966 220<br>Q5AZQ2 1174<br>Q5B4B2 133<br>Q5BBL0 320<br>P17120 1006<br>Q5B154 401<br>Q5BEU9 177<br>C8VTT9 277<br>Q5AS84 390<br>Q5AVK0 92<br>Q5AYX2 198<br>Q5B152 855<br>Q5B6C4 1156<br>Q5BDX5 1075<br>Q5AZY7 822<br>Q5AUH4 604<br>Q5BEP1 350<br>Q5BHA7 254<br>Q5AW75 35<br>Q5BE66 422<br>Q5BGS7 97<br>C8V3J6 86<br>G5EAY1 1382<br>Q5AW90 9<br>Q5BCP6 82<br>C8V9W6 340 |

| | | | | |
|---|---|---|---|---|
| | | | | C8VBJ9 645<br>Q5B1E9 1065<br>Q5B8Z7 682<br>Q5BAG3 246<br>Q5B4N9 36<br>Q5BGT7 11<br>Q5AUJ2 144<br>Q5B8I5 398<br>Q5B9C3 349<br>Q5BEB4 154<br>C8V007 410<br>C8V9W6 217<br>Q5BGN7 26<br>Q5BGZ1 17<br>C8VG45 454<br>Q5AVY3 710<br>Q5B2S6 237<br>Q5BBF2 746<br>Q5BDL2 420<br>Q5B568 347<br>Q5B6A4 144<br>C8VQ21 302<br>Q5BB70 32<br>C8V2A5 138<br>C8VCW1 225<br>C8V5G6 850<br>P0C5H8 216<br>Q5BDG0 418<br>C8V9A6 96<br>C8VP27 307<br>C8VLY6 107<br>C8VTF1 82<br>Q5B8P8 671<br>Q5BE59 627<br>Q5BFI2 309<br>C8VAW6 11<br>Q5AXK8 4<br>Q5B0E9 632<br>Q5B1Q1 80<br>Q5B7M5 12<br>Q5BBY2 736<br>Q5BDU2 184<br>G5EB07 169<br>Q5BDB4 34<br>Q5BB13 584<br>Q5B4N4 526<br>Q5B6C3 170<br>Q00083 827<br>C8V141 27<br>Q5AVV8 977<br>Q5B7M3 263<br>C8VA31 321<br>C8VUP2 145<br>Q5B3I6 451<br>Q5B425 234<br>Q5BB65 538<br>Q5BEI5 74<br>C8VTT1 103<br>C8VH46 160<br>Q5AVJ9 305<br>Q5B2M6 247<br>Q5B5B0 628<br>Q5BBL0 608<br>C8V7M0 243<br>G5EAX9 27<br>Q5BF43 269 |

| | | | | |
|---|---|---|---|---|
| | | | | C8V7J7 18<br>Q5B0E9 782<br>Q5AXE9 164<br>Q5B9P6 302<br>Q5BGC1 345<br>Q5BGE6 294<br>Q5B8Z7 964<br>Q5AVJ9 156<br>Q5AWR8 1356<br>Q5BGE6 268<br>Q5BCI4 1080<br>C8VPS9 11<br>Q5BGZ7 271<br>Q5ARG4 334<br>Q5AZQ2 380<br>Q5BH36 431<br>Q5AWR8 815<br>Q5AXD2 16<br>Q5AYS9 1195<br>Q5B8V2 354<br>Q5BBK4 669<br>Q5B1L2 6<br>C8V2A5 627<br>C8VJB5 239<br>G5EAT2 2<br>Q5BDG0 622<br>C8VHB7 98<br>Q5AXV0 735<br>Q5B4R9 231<br>Q5BB64 320<br>C8V0V3 168<br>P30583 18<br>Q5BG67 492<br>Q5AWR8 775<br>Q5B1J1 32<br>Q5B8I5 294<br>Q5BGE6 537<br>Q5AW26 604<br>C8VUL9 385<br>Q5B164 605<br>C8VIG9 368<br>Q5B1Y4 16<br>Q5BDX0 412<br>C8V8B9 84<br>Q5AVD1 304<br>Q5B4N4 532<br>Q5BGE6 474<br>Q5B706 427<br>Q5B8Z7 971<br>C8VGM0 306<br>Q5AYL5 1469<br>C8VRX7 162<br>Q5AWN0 260<br>C8VB27 542<br>C8VQ48 170<br>C8VQ33 126<br>Q5AZJ6 239<br>Q5B194 160<br>Q5BDC6 57<br>Q5B4P4 150<br>Q5B7V5 461<br>Q5BE74 1313<br>Q5BEQ3 379<br>Q5BF43 80<br>Q5ASA7 216<br>Q5AUN6 16 |

| | | | | |
|---|---|---|---|---|
| | | | | Q5B323 433<br>Q5B6S0 316<br>Q5AX03 72<br>Q5B9K9 143<br>Q5BCU8 565<br>C8V1G2 44<br>Q5B6C3 124<br>Q5BAD8 140<br>C8V733 75<br>Q5AVG6 36<br>Q5AW56 971<br>Q5AZW5 564<br>Q5BBX0 212<br>Q5AS94 207<br>Q5B0H4 85<br>Q5B568 237<br>Q5BAH9 2<br>Q5BGL6 95<br>Q5AUG3 53<br>Q5AXD2 174<br>Q5B303 323<br>Q5AWN0 165<br>Q5BBQ9 1242<br>Q5BBV0 85<br>C8VCR2 341<br>Q5B1A3 853<br>Q5B5R8 337<br>Q5B9C6 461<br>Q5BF63 15<br>Q5B0E9 96<br>C8V7B0 58<br>C8V2U2 1509<br>Q5B8A0 1758<br>Q5B4N2 105<br>Q5BDC7 215<br>Q5BH68 59<br>Q5AWI8 519<br>Q5B8R7 468<br>Q5AWK5 29<br>C8VK94 56<br>Q5B8J0 128<br>Q5BA33 68<br>Q5BG69 346<br>Q5B3J9 75<br>C8VKH5 607<br>Q5AWK4 2<br>Q5B3P2 13<br>Q5B5R8 135<br>Q5B681 632<br>C8VGM0 313<br>Q5AZL8 59<br>Q5B3P2 12<br>Q5BDC6 74<br>Q5AW56 241<br>C8V1H8 187<br>Q5BFB4 560<br>C8VMW9 615<br>O93937 1908<br>C8V8H5 694<br>Q5B798 515<br>P80581 17<br>Q5AVG3 294<br>Q5AZF0 487<br>Q5B178 336<br>Q5AVZ2 252<br>Q5BBY2 1314 |

| | | | | |
|---|---|---|---|---|
| | | | | C8V4E0 349<br>C8VRX2 36<br>Q5AVL9 24<br>C8V9T1 983<br>Q5AW75 240<br>Q5B4M5 150<br>Q5BBE4 448<br>Q5BFU5 514<br>Q5AW24 1453<br>Q5BHA9 205<br>C8VB27 451<br>Q5B2V4 409<br>Q5B2N6 223<br>C8V008 1349<br>Q5AQ97 25<br>Q5AYN6 302<br>Q5AYJ8 6<br>C8V116 351<br>Q5B399 508<br>Q5B5Z3 3<br>Q5B4E9 179<br>Q5B6S9 289<br>C8VJ33 818<br>Q5AWN0 171<br>Q5B4C9 167<br>Q5BF43 593<br>C8VAW6 10<br>Q5B0E9 383<br>Q5B0J9 2<br>Q5B1A3 170<br>C8V116 388<br>P38091 9<br>Q5BGK8 26<br>C8V3R8 540<br>Q5B978 667<br>Q5BAX8 76<br>Q5BEP4 358<br>Q5AVH4 196<br>C8VS08 981<br>Q5BAA8 338<br>Q5BGR4 303<br>C8VAT2 19<br>Q5BFC7 468<br>Q5AVK0 94<br>Q5B436 320<br>Q5B4M5 2<br>Q5B8E5 443<br>Q5BFK2 74<br>G5EB39 319<br>Q5B5U1 537<br>Q5B7V5 460<br>C8VAK8 425<br>C8VUP2 148<br>Q5B1L6 84<br>Q5B425 244<br>Q5B5H4 117<br>Q5B0E9 575<br>Q5BBF0 55<br>Q5BC59 259<br>Q5BH53 290<br>C8V116 361<br>C8V4D4 45<br>G5EB50 314<br>Q5AUN6 22<br>Q5B8X5 85<br>C8VQT9 270 |

| | | | | |
|---|---|---|---|---|
| | | | | Q5AYD4 230<br>Q5B3F1 650<br>Q5BAY0 65<br>C8VC57 457<br>Q5B1A3 713<br>Q5B5B0 1039<br>Q5BBK9 304<br>Q5AZL2 332<br>C8VP27 317<br>Q5B1G3 39<br>Q5B706 290<br>Q5BEV6 408<br>C8VK94 1031<br>Q5B424 9<br>Q5BE59 628<br>C8V097 751<br>O43100 412<br>Q5AW49 528<br>Q5B6P5 275<br>Q5BDD7 444<br>Q5BE74 708<br>C8V9T2 783<br>Q5BBE4 292<br>Q5BDD7 449<br>Q5ATX5 659<br>Q5AZN7 210<br>Q5B0L9 494<br>G5EB00 561<br>Q5AUJ2 446<br>C8VN24 305<br>C8V010 567<br>C8V8H5 521<br>Q5B3I8 878<br>Q5BHB3 23<br>Q00757 64<br>Q5AS84 437<br>Q5AZL4 360<br>Q5B3A7 277<br>Q5B4F9 991<br>Q5B963 3<br>C8VHJ7 374<br>Q5AXG1 125<br>Q5B6Y4 209<br>Q5B615 12<br>C8V733 179<br>Q5AW92 1063<br>Q5AZQ2 482<br>Q5BGX6 481<br>Q5BC59 257<br>G5EAY1 1141<br>Q5ASU2 55<br>Q5BGR2 19<br>Q5B5B5 81<br>Q5B9N1 747<br>Q5BBZ5 114<br>C8VK94 202<br>Q5AS94 405<br>Q5AYD4 245<br>Q5B152 851<br>Q5B3E4 188<br>C8V1D8 1113<br>Q5B8Q8 371<br>C8V9E9 139<br>Q5AYL5 1528<br>Q5BBL2 682<br>Q5AW79 3 |

| | | | | |
|---|---|---|---|---|
| | | | | Q5B1X4 266<br>Q5BH93 210<br>C8VAK8 421<br>Q5AW49 100<br>Q5AT23 4<br>Q5B0H6 53<br>Q5B0X8 218<br>Q5B2S6 225<br>Q5BAY0 736<br>Q5BDU2 76<br>C8VUJ9 169<br>G5EAU0 337<br>Q5B7M3 306<br>Q5AWR8 1459<br>Q5AYD4 276<br>Q5BE20 169<br>Q5BEV6 414<br>Q5AZQ2 1091<br>Q5AWL9 38<br>Q5B2N6 226<br>Q5BD80 427<br>Q5BCC9 285<br>G5EB50 418<br>O42777 24<br>Q5AS94 780<br>Q5AVJ4 426<br>Q5AZM3 939<br>Q5B1A3 854<br>C8VK54 258<br>Q5BC17 588<br>Q5BD56 3<br>C8UZZ9 137<br>C8V3I8 292<br>C8VH46 159<br>Q5AQB5 99<br>Q5ASM0 328<br>Q5AVY3 708<br>Q5B8E7 125<br>C8V9T2 786<br>C8VHR4 296<br>C8VUU8 91<br>Q5B8E4 790<br>Q5BBH9 883<br>C8V4A6 135<br>C8V4D4 44<br>Q5AZM3 1093<br>Q5AV97 522<br>Q5AUG3 738<br>Q5B606 135<br>Q5AS96 183<br>C8V8H5 681<br>Q5B1A3 870<br>Q5ASC5 159<br>Q5AVY6 428<br>Q5AX03 137<br>C8VF13 184<br>Q5B436 77<br>Q5BFV7 248<br>C8V0I4 205<br>G5EAU0 589<br>Q5AT42 2<br>Q5B6U5 67<br>Q5B8X5 400<br>Q5BB65 41<br>Q5BBW4 3586<br>C8VFJ6 211 |

| | | | | |
|---|---|---|---|---|
| | | | | C8VG86 872<br>Q5AWK9 413<br>Q5BCI0 686<br>Q5BGT8 635<br>C8VF70 9<br>Q5ARL7 519<br>Q5B712 509<br>C8VCK1 430<br>C8VS08 2<br>Q5BC21 12<br>C8V4L4 401<br>Q5B4W2 212<br>Q5BE59 840<br>C8VQ27 2<br>Q5ASD3 9<br>Q5B1I5 363<br>O60039 891<br>Q5B3B7 793<br>Q5B8R7 562<br>Q5BDG0 523<br>Q5BER4 56<br>C8V116 295<br>C8VS54 732<br>Q5BEV6 411<br>Q5AUH4 973<br>C8VDQ4 876<br>Q5B2S6 966<br>Q5B5U8 416<br>Q5B7N3 454<br>Q5BAY0 738<br>Q5BB56 71<br>C8VRX2 48<br>Q5BF15 14<br>C8V9T1 1003<br>Q5B0Y5 62<br>Q5B8X5 405<br>Q5BFV7 569<br>Q5AVT0 230<br>C8VJB5 534<br>Q5BBJ7 353<br>Q5BF63 265<br>C8V3I8 80<br>Q5B0E9 733<br>Q5B9L0 349<br>Q5BH68 56<br>C8V116 62<br>Q5B5H0 1320<br>Q5B0E9 794<br>Q5B2P4 197<br>Q5B6L5 729<br>Q5BDK6 113<br>Q5BEE9 163<br>Q5BG90 462<br>Q5BDK0 622<br>C8VP26 526<br>Q5BBY2 630<br>C8V9K6 34<br>Q5AXK8 197<br>Q5B3F1 294<br>Q5B8J3 222<br>C8VUL9 391<br>Q5B1P4 326<br>Q5B3B7 787<br>Q5B9K4 205<br>Q5BBY2 705<br>C8V3I8 294 |

| | | | | |
|---|---|---|---|---|
| | | | | G5EB07 457<br>Q5AW75 32<br>Q5BF86 229<br>Q5BGE6 290<br>Q5B3Y5 26<br>Q5ASA5 699<br>Q5AS85 194<br>Q5B1A4 422<br>Q5B4D3 190<br>Q5B5Z2 126<br>Q5AYC1 750<br>Q5AZE9 690<br>Q5B550 6<br>Q5B706 470<br>Q5BB34 840<br>C8V7M9 16<br>C8V972 120<br>Q5B6U8 643<br>Q5B8J3 200<br>Q5BDZ7 383<br>Q5BHB6 249<br>Q5B7M3 107<br>C8VN44 850<br>C8VRL9 126<br>C8VU21 218<br>Q5ARH5 532<br>Q5BCS0 1481<br>Q5BDQ5 616<br>C8VHC5 591<br>C8VIQ4 131<br>Q5B5R8 260<br>Q5B5K8 239<br>C8V1P9 268<br>P0C8Q0 142<br>Q5ASA5 834<br>Q5B3T0 2<br>C8VF13 189<br>Q5BGM8 469<br>Q5BCQ9 314<br>Q5ARI5 188<br>Q5AVD1 284<br>C8V1W0 184<br>Q5AXF0 115<br>Q5B8G1 185<br>C8V9T1 1059<br>Q5AVD1 34<br>Q5AZU3 416<br>Q5B3J1 73<br>C8V179 123<br>Q5AS91 655<br>Q5B2T9 13<br>Q5BCF3 448<br>C8VGM0 608<br>Q5ASS0 197<br>Q5B0Y7 286<br>Q5B9K9 105<br>Q5BE20 105<br>Q5B0E2 559<br>Q5B0E9 757<br>C8V1M3 3<br>C8VB12 318<br>Q5AZF7 216<br>Q5B6P5 566<br>Q5B8K7 365<br>Q5BCL2 156<br>Q5BES9 276 |

| | | | | |
|---|---|---|---|---|
| | | | | C8V095 132<br>C8V9W6 221<br>Q5B1C8 1219<br>Q5AYL7 177<br>C8VUP2 588<br>Q5AYT7 213<br>Q5B1A3 701<br>Q5B8G1 167<br>C8V8I3 201<br>C8VLE8 278<br>Q5B0C3 258<br>Q5BBY2 709<br>Q5BCL2 158<br>Q5B4Q3 32<br>Q5BDL3 107<br>C8V9H0 255<br>Q5AS72 476<br>Q5ATL5 19<br>Q5AXX7 50<br>Q5B6P5 272<br>Q5B8X0 32<br>Q5BEK5 3<br>C8VCK1 333<br>Q5AUH4 868<br>Q5BB90 313<br>C8VBH4 166<br>Q00202 549<br>C8V9Z0 47<br>Q00647 1052<br>Q5BC28 447<br>Q5B5J1 3<br>Q5BDC7 165<br>C8V1T1 201<br>C8VH90 7<br>Q5BH96 82<br>Q5AVV2 23<br>Q5AUH4 1054<br>Q5B323 487<br>Q5B8P8 862<br>Q5BDD7 501<br>Q5BGR2 32<br>C8V0C1 467<br>C8VT03 194<br>Q5B392 345<br>Q5B8J0 131<br>C8VHJ2 483<br>C8VT75 174<br>Q5BBD7 20<br>Q5B2L4 586<br>C8V7F2 3<br>Q5AVT0 248<br>Q5B7N3 410<br>C8V8H5 670<br>C8VU21 145<br>C8VCK1 198<br>Q5AYC4 446<br>Q00202 525<br>C8V5G6 471<br>C8VTA0 45<br>Q5BD94 265<br>Q5BFG5 309<br>C8VGM0 109<br>C8VUP2 229<br>Q5BEN5 770<br>Q5AZN9 231<br>Q5B1A3 869 |

| | | | | |
|---|---|---|---|---|
| | | | | Q5B3I8 643<br>Q5B9M9 497<br>Q5BEY8 479<br>C8VAQ7 41<br>Q5B425 226<br>Q5B5E1 27<br>Q5BBY2 623<br>Q8TFN0 121<br>Q5B7M3 345<br>Q5BCH1 124<br>Q5BE66 334<br>Q5BBW4 320<br>Q5B7N2 559<br>Q5B6Y7 96<br>C8V5G6 825<br>C8VID5 31<br>Q5B4F9 178<br>Q5B4I4 442<br>Q5BGS7 132<br>Q5B3S8 145<br>C8VS08 970<br>Q5AZQ2 418<br>Q5BF43 131<br>Q5BE74 1170<br>C8VH53 22<br>P18819 2<br>Q5B122 70<br>Q5B224 1433<br>Q5B5E4 2<br>Q5B6C4 131<br>C8V9E9 144<br>Q5BCI4 714<br>C8V9V9 96<br>Q5AXU1 22<br>Q5B4S4 517<br>C8VLZ4 172<br>C8VM55 182<br>Q5AVG3 239<br>Q5BBK4 684<br>Q5BE74 1059<br>Q5BB58 170<br>C8V199 50<br>C8VFC8 922<br>Q5BBJ4 813<br>Q5BBY2 664<br>C8V9W6 223<br>Q5BGR2 18<br>C8VHJ1 751<br>C8VTU4 169<br>G5EAY1 784<br>C8V4P5 319<br>C8VHD2 237<br>Q5AVK0 624<br>Q5B5B3 41<br>Q5BE39 8<br>Q5BG55 794<br>C8VJB5 464<br>C8VTU6 249<br>O42631 203<br>Q5B9C6 33<br>Q5AYC1 760<br>Q5BD26 253<br>Q5BES9 279<br>Q5BC28 314<br>Q5BCF3 48<br>C8UZY7 260 |

| | | | | |
|---|---|---|---|---|
| | | | | Q5B2T9 356<br>Q5B9S9 1115<br>C8V9T2 775<br>C8VGM0 616<br>Q5BBZ5 112<br>Q5BGL3 684<br>Q5BDC7 153<br>C8V3Q8 30<br>Q5B136 93<br>Q5B6S0 315<br>Q5BH98 130<br>C8V1J5 15<br>Q5B1Y1 493<br>Q5B4Q3 115<br>Q5B8S4 157<br>Q5BHA9 912<br>Q5B0H6 45<br>Q5B3I9 272<br>Q5BF43 261<br>Q5B7N2 13<br>C8VBH3 470<br>Q5AVD1 70<br>Q5AVN0 752<br>Q5BAH3 442<br>Q5AVV8 1819<br>Q5BCC9 16<br>Q5AVV8 412<br>Q5AVW7 717<br>Q5BB33 528<br>C8VAK9 116<br>Q5B5B3 55<br>Q5B077 708<br>Q5BBE4 449<br>Q5BH53 285<br>C8VH90 8<br>Q5AVT0 265<br>Q5B6P5 288<br>Q5BDB4 8<br>C8VSS2 454<br>Q5B4J3 504<br>Q5B010 1297<br>Q5B1U5 984<br>Q5B4W2 254<br>Q5B7M5 552<br>Q5B3C8 697<br>Q00083 655<br>Q5B178 353<br>Q5BE74 1296<br>Q5AZC2 81<br>C8UZY7 128<br>C8V9W6 166<br>C8VU49 11<br>Q5ATX5 624<br>Q5B5B3 481<br>Q5BDK0 657<br>C8V0I4 190<br>Q5BE74 830<br>Q5BEC4 264<br>Q5BF78 190<br>Q5B5V7 7<br>Q5B425 248<br>C8VKT1 226<br>C8VUS6 2<br>P11837 510<br>Q5AWR8 1557<br>Q5BBK4 345 |

| | | | | |
|---|---|---|---|---|
| | | | | Q5BEV7 291<br>C8VNK3 644<br>Q5BDY6 128<br>Q5AUJ2 2<br>Q5AWN4 1478<br>Q5BE20 192<br>Q5BEB0 158<br>Q9P896 543<br>Q5B7V5 156<br>Q5AU19 1623<br>Q5BCR1 194<br>C8VIY6 250<br>C8VJ09 202<br>Q5AXD2 48<br>Q5AZC9 505<br>Q5B0E2 1125<br>Q5B3H7 9<br>Q5AYU2 1108<br>Q5B1K5 499<br>C8VAQ7 274<br>Q5ATX5 1123<br>Q5B178 340<br>Q5B3F1 899<br>Q5BBU6 204<br>Q5BBX0 218<br>C8VP26 91<br>Q5B3C8 694<br>C8VIZ6 48<br>P18819 397<br>Q5AZU0 11<br>Q5BHB1 334<br>Q5AUB3 313<br>Q5B9P0 105<br>C8VHB7 97<br>Q5ATZ6 219<br>Q5B5V7 2<br>Q5B7Q7 146<br>Q5BCQ6 208<br>Q5BB17 275<br>Q5BEN0 10<br>C8VMC8 360<br>Q5B1L0 505<br>Q5BBY2 685<br>Q5BFQ3 230<br>Q5B5U1 278<br>Q5AZB0 470<br>Q5BDY6 127<br>Q5BCI4 27<br>C8VGM0 583<br>C8V8H5 428<br>C8VJ33 302<br>C8VRL9 46<br>Q5B104 348<br>P48457 413<br>Q5BEJ2 1108<br>Q5BHB7 341<br>Q5AVN0 308<br>Q5B6L5 563<br>Q5B8I5 478<br>Q5BAY8 106<br>Q5ARK3 808<br>Q5B223 62<br>C8VHJ1 754<br>Q5BBL2 842<br>Q5B0E9 375<br>Q5B608 118 |

| | | | | |
|---|---|---|---|---|
| | | | | Q5B1Y7 266<br>Q5B6C4 69<br>C8V115 227<br>Q5B4F9 445<br>Q5B6T7 702<br>Q5BE74 1261<br>Q5BEK5 468<br>Q5BCQ6 318<br>Q5B125 255<br>Q5BDF8 161<br>Q5AVD1 181<br>Q5B4F9 352<br>Q5B9C6 44<br>Q5BB69 69<br>Q5BBT8 433<br>Q5ASI4 131<br>Q5AYD5 993<br>Q5BB13 613<br>Q5AW39 214<br>Q5B3C0 78<br>Q5B5I8 122<br>C8VCR2 304<br>Q5B6T7 705<br>C8V5J4 706<br>C8VB27 466<br>Q5B5Z2 373<br>Q5B4G9 220<br>Q5AZQ2 255<br>C8V1S9 385<br>Q5B5X9 222<br>Q5B9K7 600<br>Q5BAE0 15<br>C8VCK1 258<br>C8VRW7 300<br>Q5ASA5 50<br>C8VAW6 13<br>C8VTU7 728<br>Q5B3C8 654<br>C8V1J5 220<br>C8V3Z7 80<br>C8V733 198<br>C8VB27 445<br>P0C8Q0 139<br>C8VEV9 498<br>G5EB50 308<br>Q5AYN2 309<br>Q5BCK9 290<br>C8UZZ9 135<br>C8VCW1 11<br>Q5AWR8 897<br>C8VT05 103<br>O42777 750<br>Q5ARL7 46<br>Q5BDC7 102<br>Q5BF66 31<br>C8V179 93<br>Q5AX75 67<br>Q5BB49 257<br>Q5BHB6 266<br>C8V002 237<br>Q5B4B7 695<br>Q5B1E9 1066<br>Q5B4F0 52<br>Q5B613 65<br>C8V751 293<br>Q5AXT8 97 |

| | | | | |
|---|---|---|---|---|
| | | | | Q5BDJ1 268<br>Q5BFN6 5<br>Q5BGL6 344<br>Q5B1Y1 457<br>C8VKK1 187<br>Q5B1A3 169<br>Q5B178 342<br>C8VA01 1177<br>Q5BDY6 117<br>Q5BE20 162<br>C8V9H0 252<br>C8VIZ6 59<br>Q5AWL9 35<br>C8VF24 349<br>Q5AUF9 371<br>Q5B2V1 546<br>Q5BBJ4 808<br>Q5BDD7 453<br>Q5ARD4 1011<br>Q5B2Q3 49<br>Q5B8X5 202<br>Q5AXQ8 294<br>Q5AZC6 1313<br>Q5B2K1 303<br>Q5BF43 118<br>C8VCK1 250<br>Q5B1I5 23<br>Q5ASC1 495<br>Q5AWN3 573<br>Q5B1A3 876<br>Q5B549 302<br>Q5B5K9 33<br>Q5BE74 1033<br>C8V904 188<br>Q5AZW4 402<br>Q5AW39 2<br>Q5B3D5 110<br>Q5BBE5 219<br>Q5BCL2 142<br>Q5BFJ6 315<br>C8VK54 156<br>Q5BAD4 924<br>Q5AW99 375<br>Q5B3C0 49<br>Q5B534 420<br>Q5B9N6 3<br>C8VAY4 340<br>Q5BF63 17<br>C8VP27 362<br>Q5AXV5 793<br>Q5B1E0 325<br>Q5B602 1086<br>Q5B674 12<br>Q5BCP0 41<br>Q5AS83 19<br>Q5AVD1 280<br>Q5AVV8 1448<br>Q5B568 343<br>C8VIY5 97<br>Q5AYL5 817<br>Q5B0D4 28<br>Q5B4K7 1768<br>Q5B5B0 1233<br>C8VG45 325<br>C8VS08 1042<br>Q5AU15 362 |

| | | | | |
|---|---|---|---|---|
| | | | | Q5BE74 828<br>C8V2U2 875<br>Q5B706 266<br>C8VK50 25<br>C8V9G9 157<br>Q5B1Y4 563<br>C8VQL3 1231<br>C8VS08 1044<br>Q5BE62 125<br>Q5BGL3 1011<br>Q5B0U1 282<br>Q5B2L8 1184<br>Q5B943 90<br>Q5BFK2 63<br>Q5B7M3 266<br>Q9P896 534<br>C8V733 80<br>C8VMX1 47<br>Q5B0E2 549<br>Q5B982 756<br>Q5BDQ6 943<br>C8VBT0 191<br>C8VHC5 154<br>C8VA27 611<br>C8VQ45 561<br>C8VRP6 416<br>Q5AU15 358<br>Q5BFG5 339<br>Q5AVD1 185<br>Q5BE59 845<br>C8VA31 414<br>Q5ASD3 28<br>C8VA69 646<br>C8VFC8 672<br>G5EB26 44<br>G5EAZ5 56<br>Q5AYK0 92<br>Q5BBD7 1022<br>Q5AVV4 1173<br>Q5B1C8 658<br>C8V007 192<br>Q5BBV0 42<br>C8VIY2 76<br>Q5AX75 65<br>Q5B4B6 10<br>Q5B9A9 310<br>G5EB90 49<br>Q5B531 766<br>Q5BCP0 9<br>Q5B4G0 19<br>Q5AVN0 305<br>C8VHR4 293<br>Q5BCF3 236<br>Q6SIF1 508<br>G5EAZ0 339<br>Q5AVD1 313<br>Q5B323 466<br>Q5BBJ7 195<br>Q5BCF3 217<br>Q5BE72 1290<br>Q5BBD7 9<br>Q5B9C6 310<br>C8VSX4 145<br>G5EAT2 11<br>Q5AUI2 164<br>Q5B5R8 409 |

| | | | | |
|---|---|---|---|---|
| | | | | Q5BCS0 1229<br>G5EB05 302<br>Q5B104 72<br>Q5B392 289<br>Q5B010 1130<br>Q5B8R7 566<br>Q5B434 388<br>Q5BB58 173<br>Q5BE74 1300<br>Q5BEG9 4<br>C8VHD2 1138<br>Q5BBY2 633<br>C8V9H4 32<br>Q5B1D1 432<br>Q5B4Q3 54<br>Q5B154 413<br>C8VCK1 202<br>G5EB41 401<br>G5EB90 51<br>C8V997 239<br>C8VBJ9 641<br>Q5AW55 8<br>Q5BEC4 260<br>C8V4M8 16<br>Q5BE23 2<br>Q5BGE6 241<br>Q5AW90 7<br>C8VG90 562<br>P17120 1058<br>Q5BFV5 740<br>Q5AW75 1086<br>Q5B3F1 621<br>C8VEH4 282<br>Q5AW57 57<br>Q5AXR8 21<br>Q5BF26 151<br>C8VHC5 256<br>Q00202 511<br>Q5AVS9 329<br>Q5BGS7 105<br>C8V2X5 212<br>C8V9K6 547<br>C8VSQ5 109<br>Q5AW56 67<br>C8VRX2 15<br>Q5BBV0 245<br>Q5BFU7 60<br>Q5B8N7 87<br>C8V3I8 318<br>Q5BE74 829<br>Q5B3H7 42<br>C8VBH5 872<br>Q5BGB8 268<br>Q5BDE6 213 |

Supplementary Data S2.2- Transcriptomic Ven Diagram

| Method of Analysis → | DPoP 795 Total | Volcano 828 Total | M/A 442 Total | MARS 3966 Total | DEseq2 3107 |
|---|---|---|---|---|---|
| List of Significant Signals ↓ | AN0102 | AN3339 | AN3531 | AN0004 | AN0045 |
| | AN0489 | AN2469 | AN9375 | AN0005 | AN0314 |
| | AN0019 | AN1035 | AN1548 | AN0046 | AN0373 |
| | AN6414 | AN2116 | AN7826 | AN0047 | AN0471 |
| | AN5218 | AN8060 | AN2469 | AN0055 | AN0599 |
| | AN6958 | AN1548 | AN7702 | AN0078 | AN0600 |
| | AN7984 | AN4109 | AN1858 | AN0096 | AN0709 |
| | AN7319 | AN0216 | AN2698 | AN0104 | AN0742 |
| | AN5465 | AN1600 | AN5218 | AN0114 | AN0777 |
| | AN2712 | AN5841 | AN6963 | AN0128 | AN0859 |
| | AN3514 | AN2571 | AN8941 | AN0139 | AN0904 |
| | AN8411 | AN4106 | AN5558 | AN0147 | AN0930 |
| | AN3495 | AN8996 | AN1742 | AN0155 | AN1071 |
| | AN9299 | AN2562 | AN1312 | AN0157 | AN1250 |
| | AN5467 | AN2186 | AN9490 | AN0169 | AN1356 |
| | AN7412 | AN3571 | AN9299 | AN0178 | AN1549 |
| | AN9181 | AN2320 | AN2827 | AN0188 | AN1607 |
| | AN2352 | AN0523 | AN5467 | AN0213 | AN1620 |
| | AN8647 | AN4110 | AN0941 | AN0214 | AN1653 |
| | AN8052 | AN2960 | AN5977 | AN0229 | AN1871 |
| | AN2471 | AN2558 | AN4879 | AN0230 | AN1894 |
| | AN2806 | AN4111 | AN2118 | AN0233 | AN1935 |
| | AN0341 | AN2959 | AN5611 | AN0243 | AN1936 |
| | AN0100 | AN7859 | AN8158 | AN0260 | AN2011 |
| | AN7781 | AN8597 | AN0958 | AN0284 | AN2020 |
| | AN5509 | AN6873 | AN5652 | AN0296 | AN2421 |
| | AN9200 | AN7413 | AN5017 | AN0324 | AN2427 |
| | AN0723 | AN7598 | AN4109 | AN0331 | AN2477 |
| | AN5069 | AN3531 | AN5859 | AN0352 | AN2532 |
| | AN2605 | AN7053 | AN0553 | AN0357 | AN2581 |
| | AN8662 | AN0008 | AN1664 | AN0363 | AN2601 |
| | AN7232 | AN8132 | AN5424 | AN0365 | AN2685 |
| | AN7881 | AN7854 | AN0723 | AN0373 | AN2695 |
| | AN5823 | AN8551 | AN5069 | AN0377 | AN2776 |
| | AN6929 | AN9210 | AN3333 | AN0378 | AN2822 |
| | AN6234 | AN2028 | AN7232 | AN0392 | AN2911 |
| | AN1627 | AN4131 | AN7881 | AN0399 | AN2945 |
| | AN8582 | AN7066 | AN0020 | AN0418 | AN3085 |
| | AN8487 | AN8112 | AN0500 | AN0436 | AN3086 |
| | AN2031 | AN7531 | AN5171 | AN0455 | AN3101 |
| | AN4950 | AN0016 | AN2913 | AN0456 | AN3148 |
| | AN8371 | AN2706 | AN4142 | AN0466 | AN3287 |
| | AN0492 | AN1317 | AN4106 | AN0486 | AN3751 |
| | AN8728 | AN7237 | AN7683 | AN0487 | AN4088 |
| | AN6023 | AN3537 | AN6023 | AN0497 | AN4089 |
| | AN5289 | AN2840 | AN6847 | AN0499 | AN4104 |
| | AN3398 | AN6359 | AN2228 | AN0511 | AN4113 |
| | AN7378 | AN0552 | AN2656 | AN0520 | AN4266 |
| | AN3241 | AN0993 | AN6659 | AN0527 | AN4483 |
| | AN8609 | AN8157 | AN1677 | AN0529 | AN4566 |
| | AN3485 | AN7268 | AN3996 | AN0535 | AN4696 |
| | AN6947 | AN7333 | AN7866 | AN0552 | AN4712 |
| | AN2943 | AN4879 | AN9242 | AN0609 | AN4770 |
| | AN0993 | AN3305 | AN7167 | AN0638 | AN4824 |
| | AN1737 | AN8446 | AN7333 | AN0639 | AN4878 |
| | AN0972 | AN2393 | AN9001 | AN0674 | AN4879 |
| | AN2609 | AN4770 | AN2038 | AN0676 | AN4913 |
| | AN8611 | AN6936 | AN1574 | AN0692 | AN4920 |

| | | | | | |
|---|---|---|---|---|---|
| | AN7856 | AN6281 | AN2336 | AN0710 | AN4995 |
| | AN8442 | AN1664 | AN0993 | AN0722 | AN5017 |
| | AN9280 | AN3393 | AN7392 | AN0725 | AN5159 |
| | AN8927 | AN8349 | AN5421 | AN0749 | AN5213 |
| | AN0517 | AN3333 | AN9179 | AN0763 | AN5856 |
| | AN6281 | AN4257 | AN0488 | AN0767 | AN5858 |
| | AN9318 | AN1594 | AN2099 | AN0776 | AN6004 |
| | AN4049 | AN8594 | AN1616 | AN0777 | AN6071 |
| | AN8909 | AN0964 | AN0392 | AN0778 | AN6091 |
| | AN5444 | AN8968 | AN2197 | AN0787 | AN6114 |
| | AN3109 | AN5424 | AN0366 | AN0789 | AN6241 |
| | AN5845 | AN2719 | AN3888 | AN0832 | AN6277 |
| | AN2679 | AN7385 | AN0234 | AN0859 | AN6397 |
| | AN0609 | AN7668 | AN5053 | AN0868 | AN6404 |
| | AN6957 | AN1203 | AN6281 | AN0874 | AN6573 |
| | AN6273 | AN8949 | AN2557 | AN0875 | AN6634 |
| | AN2638 | AN2598 | AN5845 | AN0894 | AN6820 |
| | AN7753 | AN5546 | AN6833 | AN0911 | AN6835 |
| | AN4143 | AN3414 | AN0973 | AN0917 | AN6849 |
| | AN7322 | AN5449 | AN3433 | AN0922 | AN6871 |
| | AN8134 | AN3612 | AN7541 | AN0938 | AN6873 |
| | AN0220 | AN1677 | AN8446 | AN0939 | AN6982 |
| | AN1752 | AN0234 | AN0045 | AN0940 | AN7018 |
| | AN6089 | AN6963 | AN0694 | AN0949 | AN7170 |
| | AN3078 | AN5558 | AN7322 | AN0963 | AN7463 |
| | AN5029 | AN1042 | AN8134 | AN0972 | AN7530 |
| | AN3503 | AN9470 | AN3783 | AN0982 | AN7549 |
| | AN6377 | AN3314 | AN8095 | AN0991 | AN7604 |
| | AN3387 | AN1312 | AN7832 | AN0992 | AN7605 |
| | AN5435 | AN8341 | AN5015 | AN0998 | AN7670 |
| | AN3043 | AN8342 | AN1752 | AN1000 | AN7717 |
| | AN8554 | AN7083 | AN5338 | AN1014 | AN7992 |
| | AN8685 | AN9303 | AN9206 | AN1018 | AN8311 |
| | AN5937 | AN7683 | AN0215 | AN1027 | AN8534 |
| | AN3562 | AN7160 | AN7953 | AN1030 | AN8797 |
| | AN0336 | AN6018 | AN7870 | AN1040 | AN9168 |
| | AN7196 | AN0148 | AN3043 | AN1042 | AN0214 |
| | AN6691 | AN7170 | AN5271 | AN1091 | AN0644 |
| | AN8107 | AN0916 | AN5272 | AN1120 | AN0645 |
| | AN1623 | AN7825 | AN8587 | AN1138 | AN0651 |
| | AN2821 | AN3282 | AN8366 | AN1139 | AN0750 |
| | AN9183 | AN2704 | AN7557 | AN1143 | AN0789 |
| | AN4257 | AN6648 | AN0974 | AN1162 | AN0823 |
| | AN6959 | AN7225 | AN1893 | AN1181 | AN0833 |
| | AN7806 | AN2357 | AN2821 | AN1183 | AN0866 |
| | AN8139 | AN9263 | AN2392 | AN1188 | AN0983 |
| | AN3479 | AN6657 | AN4257 | AN1198 | AN1102 |
| | AN7095 | AN4888 | AN8082 | AN1209 | AN1251 |
| | AN9487 | AN2943 | AN7057 | AN1222 | AN1328 |
| | AN8338 | AN2488 | AN1203 | AN1228 | AN1535 |
| | AN0006 | AN3163 | AN5831 | AN1245 | AN1574 |
| | AN6021 | AN5130 | AN0467 | AN1259 | AN1580 |
| | AN3303 | AN7856 | AN9212 | AN1261 | AN1619 |
| | AN2402 | AN7867 | AN8639 | AN1290 | AN1622 |
| | AN4436 | AN8314 | AN7992 | AN1301 | AN1656 |
| | AN6668 | AN1378 | AN3303 | AN1302 | AN1676 |
| | AN2833 | AN6810 | AN8619 | AN1307 | AN1752 |
| | AN7065 | AN2286 | AN8781 | AN1311 | AN1797 |
| | AN2788 | AN5611 | AN9191 | AN1321 | AN1855 |
| | AN7867 | AN9031 | AN2404 | AN1327 | AN1870 |
| | AN9278 | AN7866 | AN1600 | AN1337 | AN1918 |
| | AN8365 | AN4119 | AN7600 | AN1377 | AN2010 |
| | AN4660 | AN7278 | AN2559 | AN1424 | AN2189 |
| | AN2787 | AN8554 | AN2833 | AN1432 | AN2286 |
| | AN3915 | AN9244 | AN9243 | AN1441 | AN2357 |
| | AN9223 | AN1034 | AN3570 | AN1445 | AN2391 |

| | | | | | |
|---|---|---|---|---|---|
| | AN9248 | AN9207 | AN2034 | AN1450 | AN2408 |
| | AN3243 | AN2821 | AN1898 | AN1471 | AN2488 |
| | AN3255 | AN2572 | AN3023 | AN1486 | AN2669 |
| | AN7179 | AN4376 | AN3271 | AN1490 | AN2682 |
| | AN1199 | AN9032 | AN8365 | AN1501 | AN2704 |
| | AN0342 | AN9042 | AN1582 | AN1505 | AN2749 |
| | AN8477 | AN2532 | AN3304 | AN1508 | AN2798 |
| | AN9182 | AN5370 | AN1631 | AN1509 | AN2826 |
| | AN5938 | AN7855 | AN3781 | AN1512 | AN2919 |
| | AN8953 | AN3698 | AN4111 | AN1540 | AN2943 |
| | AN6766 | AN8514 | AN2894 | AN1548 | AN3049 |
| | AN2196 | AN7916 | AN1199 | AN1555 | AN3163 |
| | AN8277 | AN8556 | AN5090 | AN1557 | AN3265 |
| | AN8140 | AN9269 | AN2655 | AN1562 | AN3415 |
| | AN0005 | AN6057 | AN9205 | AN1572 | AN3417 |
| | AN7519 | AN7865 | AN3571 | AN1598 | AN3420 |
| | AN2319 | AN8652 | AN8451 | AN1599 | AN3433 |
| | AN8404 | AN3276 | AN2588 | AN1603 | AN3526 |
| | AN8632 | AN7094 | AN2039 | AN1604 | AN3531 |
| | AN3472 | AN5659 | AN3776 | AN1605 | AN3552 |
| | AN4094 | AN7990 | AN6792 | AN1606 | AN3553 |
| | AN3891 | AN3285 | AN2941 | AN1607 | AN3665 |
| | AN7215 | AN3384 | AN5286 | AN1611 | AN3787 |
| | AN8241 | AN0393 | AN5270 | AN1612 | AN4051 |
| | AN7097 | AN5019 | AN2196 | AN1613 | AN4119 |
| | AN5280 | AN4642 | AN2198 | AN1615 | AN4120 |
| | AN7116 | AN2901 | AN8277 | AN1621 | AN4135 |
| | AN8220 | AN2611 | AN8333 | AN1629 | AN4170 |
| | AN9031 | AN5290 | AN9372 | AN1630 | AN4192 |
| | AN5385 | AN3195 | AN7519 | AN1637 | AN4252 |
| | AN7908 | AN9205 | AN2187 | AN1638 | AN4257 |
| | AN5651 | AN2798 | AN2321 | AN1645 | AN4365 |
| | AN3276 | AN2827 | AN5918 | AN1654 | AN4375 |
| | AN7518 | AN8008 | AN9176 | AN1655 | AN4609 |
| | AN1645 | AN3573 | AN8116 | AN1663 | AN4833 |
| | AN7323 | AN3496 | AN4173 | AN1669 | AN4837 |
| | AN4108 | AN1789 | AN3286 | AN1670 | AN4854 |
| | AN2033 | AN5465 | AN9031 | AN1672 | AN4888 |
| | AN2789 | AN3636 | AN9332 | AN1675 | AN4973 |
| | AN0971 | AN5359 | AN5651 | AN1676 | AN5130 |
| | AN8920 | AN4566 | AN7518 | AN1679 | AN5364 |
| | AN0612 | AN0031 | AN9219 | AN1685 | AN5385 |
| | AN3553 | AN5213 | AN4108 | AN1687 | AN5402 |
| | AN8452 | AN3206 | AN2387 | AN1691 | AN5465 |
| | AN5072 | AN6931 | AN0418 | AN1702 | AN5568 |
| | AN8627 | AN7654 | AN5269 | AN1709 | AN5611 |
| | AN1542 | AN7540 | AN0511 | AN1713 | AN5622 |
| | AN3581 | AN0983 | AN7058 | AN1722 | AN5667 |
| | AN3887 | AN5691 | AN8137 | AN1725 | AN5906 |
| | AN1314 | AN3513 | AN3361 | AN1741 | AN5929 |
| | AN8409 | AN8300 | AN7082 | AN1751 | AN5935 |
| | AN3858 | AN3310 | AN1895 | AN1756 | AN6012 |
| | AN2376 | AN1606 | AN1897 | AN1777 | AN6019 |
| | AN9284 | AN7549 | AN5074 | AN1780 | AN6132 |
| | AN8453 | AN5141 | AN3203 | AN1793 | AN6163 |
| | AN7843 | AN7130 | AN7406 | AN1794 | AN6249 |
| | AN2097 | AN9371 | AN2921 | AN1800 | AN6338 |
| | AN2817 | AN1884 | AN3414 | AN1811 | AN6405 |
| | AN7212 | AN0644 | AN3887 | AN1844 | AN6581 |
| | AN9178 | AN7732 | AN1941 | AN1883 | AN6625 |
| | AN6748 | AN4921 | AN2901 | AN1892 | AN6657 |
| | AN7488 | AN5468 | AN5885 | AN1894 | AN6701 |
| | AN7142 | AN1912 | AN6378 | AN1939 | AN6712 |
| | AN7809 | AN8062 | AN8445 | AN1940 | AN6739 |
| | AN7397 | AN3481 | AN1612 | AN1971 | AN6810 |
| | AN8459 | AN3218 | AN6024 | AN2009 | AN6869 |

| | | | | | |
|---|---|---|---|---|---|
| | AN0322 | AN8148 | AN8593 | AN2011 | AN6870 |
| | AN3574 | AN2582 | AN1605 | AN2013 | AN7053 |
| | AN3207 | AN1189 | AN8595 | AN2021 | AN7142 |
| | AN8526 | AN4920 | AN0230 | AN2022 | AN7166 |
| | AN5276 | AN3415 | AN8327 | AN2029 | AN7172 |
| | AN5487 | AN2408 | AN4590 | AN2030 | AN7173 |
| | AN8549 | AN2702 | AN1302 | AN2031 | AN7225 |
| | AN9343 | AN8994 | AN9178 | AN2034 | AN7337 |
| | AN1576 | AN1532 | AN6018 | AN2036 | AN7488 |
| | AN3248 | AN3998 | AN9175 | AN2037 | AN7510 |
| | AN5502 | AN2454 | AN7268 | AN2038 | AN7600 |
| | AN5564 | AN5402 | AN7142 | AN2042 | AN7759 |
| | AN7850 | AN7738 | AN7592 | AN2046 | AN7800 |
| | AN7398 | AN0171 | AN2959 | AN2063 | AN7816 |
| | AN5844 | AN0750 | AN7876 | AN2079 | AN7870 |
| | AN6671 | AN7191 | AN9453 | AN2091 | AN8062 |
| | AN7380 | AN0645 | AN7898 | AN2098 | AN8139 |
| | AN8471 | AN3279 | AN7859 | AN2109 | AN8140 |
| | AN2032 | AN3520 | AN8459 | AN2110 | AN8281 |
| | AN3984 | AN7941 | AN3239 | AN2111 | AN8314 |
| | AN8444 | AN7530 | AN0231 | AN2113 | AN8444 |
| | AN5259 | AN6801 | AN3194 | AN2117 | AN8524 |
| | AN5470 | AN8608 | AN9174 | AN2124 | AN8545 |
| | AN9364 | AN2826 | AN6398 | AN2153 | AN8558 |
| | AN5396 | AN1810 | AN2177 | AN2163 | AN8627 |
| | AN7233 | AN3566 | AN2823 | AN2167 | AN8641 |
| | AN6765 | AN7880 | AN5487 | AN2176 | AN8774 |
| | AN6936 | AN3777 | AN1614 | AN2178 | AN8907 |
| | AN1593 | AN2412 | AN9180 | AN2186 | AN8990 |
| | AN5092 | AN2682 | AN8008 | AN2193 | AN8994 |
| | AN0024 | AN7175 | AN1184 | AN2197 | AN8999 |
| | AN1307 | AN2022 | AN8547 | AN2199 | AN9105 |
| | AN4583 | AN4171 | AN7515 | AN2224 | AN9206 |
| | AN7066 | AN3497 | AN8996 | AN2227 | AN9263 |
| | AN8449 | AN8149 | AN5537 | AN2245 | AN9297 |
| | AN7075 | AN0364 | AN2778 | AN2301 | AN0215 |
| | AN3336 | AN4372 | AN7883 | AN2309 | AN0321 |
| | AN7409 | AN1586 | AN7021 | AN2320 | AN0330 |
| | AN9226 | AN6535 | AN3132 | AN2324 | AN0387 |
| | AN2030 | AN5944 | AN5664 | AN2328 | AN0409 |
| | AN4135 | AN7896 | AN3388 | AN2356 | AN0553 |
| | AN7812 | AN6772 | AN4769 | AN2357 | AN0604 |
| | AN4147 | AN8102 | AN3961 | AN2358 | AN0620 |
| | AN2357 | AN9262 | AN7170 | AN2359 | AN0732 |
| | AN8323 | AN9206 | AN8934 | AN2383 | AN0869 |
| | AN3163 | AN0539 | AN2186 | AN2392 | AN0918 |
| | AN2110 | AN4151 | AN2648 | AN2395 | AN0982 |
| | AN8550 | AN3347 | AN5935 | AN2406 | AN1037 |
| | AN5354 | AN8545 | AN9364 | AN2407 | AN1144 |
| | AN7865 | AN9271 | AN7233 | AN2411 | AN1487 |
| | AN0016 | AN3976 | AN7800 | AN2418 | AN1605 |
| | AN2603 | AN2683 | AN6401 | AN2431 | AN1864 |
| | AN8108 | AN7942 | AN0468 | AN2467 | AN1910 |
| | AN3504 | AN8395 | AN3234 | AN2468 | AN2030 |
| | AN4119 | AN2659 | AN1604 | AN2469 | AN2032 |
| | AN5542 | AN7178 | AN6936 | AN2483 | AN2034 |
| | AN7225 | AN3997 | AN2781 | AN2487 | AN2038 |
| | AN0516 | AN8316 | AN0010 | AN2493 | AN2197 |
| | AN8299 | AN7070 | AN1304 | AN2508 | AN2320 |
| | AN3347 | AN1575 | AN3315 | AN2515 | AN2343 |
| | AN0035 | AN1214 | AN0811 | AN2529 | AN2351 |
| | AN7827 | AN2714 | AN7882 | AN2548 | AN2655 |
| | AN3674 | AN1593 | AN5546 | AN2557 | AN2693 |
| | AN2729 | AN7414 | AN1613 | AN2561 | AN2760 |
| | AN8551 | AN2720 | AN2798 | AN2570 | AN3282 |
| | AN9276 | AN5896 | AN7409 | AN2573 | AN3512 |

| | | | | | |
|---|---|---|---|---|---|
| | AN6057 | AN1501 | AN1261 | AN2575 | AN3600 |
| | AN2606 | AN1307 | AN4121 | AN2581 | AN3957 |
| | AN6118 | AN2601 | AN9163 | AN2587 | AN3963 |
| | AN1747 | AN7886 | AN2608 | AN2608 | AN3971 |
| | AN0617 | AN8650 | AN5056 | AN2640 | AN3996 |
| | AN5944 | AN8665 | AN0610 | AN2644 | AN3998 |
| | AN2795 | AN9295 | AN4107 | AN2646 | AN4111 |
| | AN0223 | AN0546 | AN2110 | AN2648 | AN4127 |
| | AN3195 | AN9285 | AN7129 | AN2653 | AN4263 |
| | AN1030 | AN8754 | AN5354 | AN2655 | AN4287 |
| | AN7838 | AN6121 | AN8417 | AN2658 | AN4443 |
| | AN8354 | AN3255 | AN0016 | AN2667 | AN4749 |
| | AN8621 | AN2111 | AN3139 | AN2668 | AN4769 |
| | AN7517 | AN3343 | AN8157 | AN2697 | AN4844 |
| | AN7959 | AN3077 | AN0531 | AN2703 | AN4870 |
| | AN6942 | AN8455 | AN4119 | AN2726 | AN4950 |
| | AN3963 | AN7390 | AN3247 | AN2727 | AN5044 |
| | AN3551 | AN7137 | AN1813 | AN2773 | AN5051 |
| | AN2724 | AN4627 | AN5542 | AN2777 | AN5056 |
| | AN4127 | AN6389 | AN8400 | AN2778 | AN5198 |
| | AN6935 | AN6402 | AN5085 | AN2780 | AN5228 |
| | AN3260 | AN8451 | AN8109 | AN2825 | AN5354 |
| | AN3563 | AN8941 | AN0516 | AN2848 | AN5489 |
| | AN3615 | AN1582 | AN4770 | AN2850 | AN5694 |
| | AN2793 | AN5085 | AN5427 | AN2861 | AN5883 |
| | AN2542 | AN4125 | AN5083 | AN2866 | AN6057 |
| | AN7118 | AN6001 | AN1322 | AN2874 | AN6168 |
| | AN9267 | AN4136 | AN2640 | AN2890 | AN6377 |
| | AN9279 | AN9089 | AN8007 | AN2900 | AN6513 |
| | AN3536 | AN2710 | AN1607 | AN2918 | AN6534 |
| | AN2590 | AN3631 | AN2445 | AN2940 | AN6550 |
| | AN7264 | AN8364 | AN3238 | AN2942 | AN6753 |
| | AN9348 | AN9088 | AN6930 | AN2944 | AN6796 |
| | AN5843 | AN3316 | AN8657 | AN2945 | AN6801 |
| | AN2600 | AN8422 | AN9339 | AN2946 | AN7159 |
| | AN8408 | AN2534 | AN8548 | AN2950 | AN7295 |
| | AN7379 | AN8334 | AN3299 | AN2953 | AN7378 |
| | AN8910 | AN0572 | AN6118 | AN2954 | AN7392 |
| | AN0300 | AN1138 | AN1857 | AN2958 | AN7482 |
| | AN8612 | AN8299 | AN6941 | AN2980 | AN7493 |
| | AN0482 | AN3522 | AN1747 | AN2987 | AN7528 |
| | AN6411 | AN9172 | AN7668 | AN2988 | AN7557 |
| | AN8340 | AN0005 | AN8348 | AN2994 | AN7669 |
| | AN8324 | AN1631 | AN8341 | AN3016 | AN7743 |
| | AN5964 | AN3162 | AN8596 | AN3019 | AN7760 |
| | AN7868 | AN9033 | AN6869 | AN3020 | AN7774 |
| | AN8563 | AN4826 | AN1035 | AN3022 | AN7779 |
| | AN7418 | AN5360 | AN0473 | AN3026 | AN7922 |
| | AN7990 | AN9535 | AN8641 | AN3030 | AN7943 |
| | AN6292 | AN7136 | AN3511 | AN3042 | AN7954 |
| | AN2613 | AN6917 | AN3573 | AN3050 | AN8019 |
| | AN1928 | AN8901 | AN8354 | AN3078 | AN8067 |
| | AN9054 | AN7893 | AN2558 | AN3079 | AN8102 |
| | AN5846 | AN1925 | AN7989 | AN3099 | AN8337 |
| | AN9352 | AN5964 | AN7517 | AN3120 | AN8354 |
| | AN8422 | AN3511 | AN3963 | AN3129 | AN8362 |
| | AN8436 | AN3392 | AN9324 | AN3131 | AN8412 |
| | AN6525 | AN3475 | AN7053 | AN3135 | AN8421 |
| | AN8135 | AN7804 | AN5636 | AN3140 | AN8451 |
| | AN8508 | AN0214 | AN6935 | AN3142 | AN8530 |
| | AN2398 | AN6691 | AN7708 | AN3154 | AN8635 |
| | AN9350 | AN7827 | AN2199 | AN3161 | AN8651 |
| | AN1571 | AN0136 | AN3563 | AN3162 | AN8771 |
| | AN0950 | AN3223 | AN0009 | AN3174 | AN9031 |
| | AN3275 | AN3774 | AN0523 | AN3193 | AN9032 |
| | AN2379 | AN2612 | AN7278 | AN3221 | AN9039 |

| | | | | | |
|---|---|---|---|---|---|
| | AN6649 | AN7238 | AN7295 | AN3222 | AN9070 |
| | AN5046 | AN0200 | AN0606 | AN3225 | AN9205 |
| | AN7098 | AN6654 | AN5650 | AN3233 | AN9304 |
| | AN9502 | AN5608 | AN0300 | AN3238 | AN9470 |
| | AN2831 | AN0495 | AN5843 | AN3246 | AN0056 |
| | AN2841 | AN7215 | AN2351 | AN3264 | AN0210 |
| | AN7855 | AN9039 | AN0497 | AN3281 | AN0456 |
| | AN8607 | AN8999 | AN1581 | AN3284 | AN0482 |
| | AN0915 | AN7230 | AN2574 | AN3293 | AN0495 |
| | AN8433 | AN5044 | AN5555 | AN3297 | AN0617 |
| | AN8399 | AN4554 | AN9004 | AN3303 | AN0712 |
| | AN7805 | AN1623 | AN8592 | AN3308 | AN0723 |
| | AN1549 | AN0527 | AN6949 | AN3330 | AN1184 |
| | AN6236 | AN2697 | AN4051 | AN3347 | AN1199 |
| | AN3782 | AN8139 | AN8340 | AN3360 | AN1246 |
| | AN3200 | AN8052 | AN8659 | AN3384 | AN1278 |
| | AN4888 | AN4059 | AN8324 | AN3386 | AN1279 |
| | AN9247 | AN8398 | AN2393 | AN3414 | AN1378 |
| | AN3892 | AN0283 | AN6409 | AN3415 | AN1608 |
| | AN9032 | AN7928 | AN7391 | AN3451 | AN1941 |
| | AN0878 | AN2813 | AN0693 | AN3455 | AN2025 |
| | AN1029 | AN5383 | AN5422 | AN3496 | AN2029 |
| | AN9244 | AN8140 | AN9213 | AN3507 | AN2045 |
| | AN0860 | AN2841 | AN2116 | AN3508 | AN2277 |
| | AN6153 | AN2695 | AN5841 | AN3510 | AN2310 |
| | AN2677 | AN2025 | AN5976 | AN3512 | AN2312 |
| | AN3249 | AN6409 | AN7181 | AN3523 | AN2572 |
| | AN9295 | AN1581 | AN8135 | AN3525 | AN2636 |
| | AN5419 | AN6942 | AN2320 | AN3527 | AN2730 |
| | AN7237 | AN5408 | AN9190 | AN3529 | AN2813 |
| | AN9177 | AN4704 | AN3558 | AN3565 | AN2951 |
| | AN7393 | AN8697 | AN8968 | AN3570 | AN2979 |
| | AN2805 | AN7319 | AN8904 | AN3571 | AN3051 |
| | AN8465 | AN1644 | AN2360 | AN3580 | AN3072 |
| | AN7813 | AN5190 | AN1899 | AN3605 | AN3111 |
| | AN8302 | AN0513 | AN6871 | AN3607 | AN3162 |
| | AN3611 | AN5502 | AN7999 | AN3623 | AN3223 |
| | AN5130 | AN0760 | AN2719 | AN3643 | AN3290 |
| | AN7024 | AN9352 | AN7018 | AN3686 | AN3331 |
| | AN7962 | AN8477 | AN2560 | AN3695 | AN3344 |
| | AN3557 | AN7910 | AN9220 | AN3701 | AN3347 |
| | AN3344 | AN4507 | AN7110 | AN3713 | AN3356 |
| | AN1541 | AN0482 | AN7855 | AN3719 | AN3361 |
| | AN8994 | AN1279 | AN0528 | AN3747 | AN3392 |
| | AN5393 | AN5396 | AN3560 | AN3758 | AN3404 |
| | AN3272 | AN0006 | AN8594 | AN3763 | AN3491 |
| | AN6359 | AN0737 | AN2694 | AN3789 | AN3511 |
| | AN9304 | AN1580 | AN7385 | AN3804 | AN3585 |
| | AN2532 | AN5280 | AN0786 | AN3812 | AN3674 |
| | AN4669 | AN8611 | AN7955 | AN3852 | AN3738 |
| | AN8971 | AN9502 | AN7875 | AN3864 | AN3929 |
| | AN1786 | AN4621 | AN0148 | AN3885 | AN3975 |
| | AN7863 | AN2822 | AN3960 | AN3887 | AN3987 |
| | AN8113 | AN0694 | AN0964 | AN3888 | AN3988 |
| | AN7946 | AN9163 | AN0933 | AN3897 | AN4094 |
| | AN8342 | AN5071 | AN2576 | AN3931 | AN4136 |
| | AN6657 | AN4152 | AN9177 | AN3941 | AN4286 |
| | AN5556 | AN6397 | AN1803 | AN3942 | AN4583 |
| | AN1681 | AN5259 | AN9285 | AN3960 | AN4975 |
| | AN5449 | AN9364 | AN2561 | AN3995 | AN5129 |
| | AN5397 | AN2832 | AN5355 | AN3997 | AN5408 |
| | AN8892 | AN9174 | AN4642 | AN4017 | AN5480 |
| | AN8439 | AN9123 | AN7954 | AN4018 | AN5487 |
| | AN1034 | AN4806 | AN4110 | AN4023 | AN5509 |
| | AN9297 | AN5329 | AN3393 | AN4024 | AN5554 |
| | AN2488 | AN8934 | AN7880 | AN4040 | AN5564 |

| | | | | | |
|---|---|---|---|---|---|
| | AN9245 | AN4115 | AN3557 | AN4052 | AN5602 |
| | AN8962 | AN9196 | AN2783 | AN4054 | AN5845 |
| | AN8678 | AN0378 | AN0216 | AN4077 | AN5893 |
| | AN3008 | AN8082 | AN3872 | AN4080 | AN5975 |
| | AN8060 | AN7384 | AN2598 | AN4104 | AN6058 |
| | AN2568 | AN5301 | AN2891 | AN4105 | AN6089 |
| | AN8186 | AN8447 | AN8958 | AN4106 | AN6359 |
| | AN8353 | AN1969 | AN1594 | AN4108 | AN6409 |
| | AN2696 | AN8982 | AN6359 | AN4109 | AN6487 |
| | AN1901 | AN1815 | AN4806 | AN4110 | AN6541 |
| | AN4605 | AN9378 | AN9303 | AN4119 | AN6593 |
| | AN8730 | AN8630 | AN7020 | AN4125 | AN6636 |
| | AN2730 | AN0713 | AN7634 | AN4127 | AN6693 |
| | AN8907 | AN4423 | AN2571 | AN4134 | AN6718 |
| | AN5417 | AN5628 | AN8637 | AN4145 | AN6809 |
| | AN3285 | AN5030 | AN8597 | AN4150 | AN6973 |
| | AN6810 | AN7058 | AN6399 | AN4170 | AN7091 |
| | AN7384 | AN3960 | AN5273 | AN4194 | AN7226 |
| | AN3331 | AN8106 | AN3339 | AN4223 | AN7263 |
| | AN5093 | AN6384 | AN8463 | AN4229 | AN7266 |
| | AN7202 | AN6830 | AN4129 | AN4235 | AN7268 |
| | AN0535 | AN8438 | AN8342 | AN4248 | AN7278 |
| | AN8136 | AN2796 | AN1692 | AN4251 | AN7369 |
| | AN5261 | AN9351 | AN4105 | AN4256 | AN7577 |
| | AN2704 | AN2344 | AN5449 | AN4262 | AN7581 |
| | AN0330 | AN0477 | AN8370 | AN4272 | AN7683 |
| | AN9197 | AN8392 | AN8439 | AN4273 | AN7751 |
| | AN2649 | AN0588 | AN3321 | AN4294 | AN7796 |
| | AN3533 | AN9175 | AN9297 | AN4304 | AN7818 |
| | AN2811 | AN3769 | AN2562 | AN4327 | AN7866 |
| | AN5507 | AN4147 | AN6535 | AN4328 | AN8068 |
| | AN3531 | AN3627 | AN2200 | AN4344 | AN8269 |
| | AN8521 | AN6667 | AN7217 | AN4360 | AN8334 |
| | AN1548 | AN4480 | AN8084 | AN4362 | AN8376 |
| | AN5369 | AN0756 | AN7799 | AN4371 | AN8399 |
| | AN5304 | AN7912 | AN8060 | AN4383 | AN8540 |
| | AN1835 | AN9160 | AN9470 | AN4393 | AN8611 |
| | AN6098 | AN4481 | AN8349 | AN4407 | AN8689 |
| | AN2563 | AN8322 | AN3612 | AN4422 | AN8983 |
| | AN1807 | AN6642 | AN9123 | AN4449 | AN9124 |
| | AN0501 | AN1641 | AN8995 | AN4452 | AN9178 |
| | AN8011 | AN7069 | AN8392 | AN4469 | AN9243 |
| | AN5226 | AN5791 | AN5665 | AN4480 | AN9244 |
| | AN2564 | AN5864 | AN2730 | AN4481 | AN0365 |
| | AN7810 | AN6647 | AN8907 | AN4488 | AN0498 |
| | AN5301 | AN7022 | AN7266 | AN4513 | AN0561 |
| | AN4144 | AN4769 | AN3403 | AN4526 | AN0639 |
| | AN9170 | AN1316 | AN8136 | AN4534 | AN0654 |
| | AN9165 | AN5478 | AN7952 | AN4546 | AN0663 |
| | AN5611 | AN7669 | AN2704 | AN4549 | AN0756 |
| | AN8158 | AN5287 | AN8308 | AN4551 | AN0860 |
| | AN5652 | AN7406 | AN4829 | AN4560 | AN0993 |
| | AN0958 | AN3023 | AN2649 | AN4565 | AN1123 |
| | AN5466 | AN0732 | AN2960 | AN4577 | AN1198 |
| | AN5017 | AN2746 | | AN4607 | AN1412 |
| | AN8101 | AN8323 | | AN4608 | AN1451 |
| | AN8334 | AN1071 | | AN4610 | AN1557 |
| | AN5859 | AN0498 | | AN4641 | AN1572 |
| | AN9478 | AN7407 | | AN4654 | AN1581 |
| | AN2572 | AN4361 | | AN4701 | AN1600 |
| | AN0553 | AN2064 | | AN4732 | AN1644 |
| | AN8983 | AN1781 | | AN4744 | AN1901 |
| | AN4128 | AN1032 | | AN4768 | AN1923 |
| | AN0499 | AN8085 | | AN4769 | AN2024 |
| | AN5424 | AN7267 | | AN4773 | AN2118 |
| | AN1938 | AN4019 | | AN4794 | AN2230 |

| | | | | | |
|---|---|---|---|---|---|
| | AN7801 | AN8347 | | AN4799 | AN2326 |
| | AN3523 | AN1574 | | AN4800 | AN2336 |
| | AN3333 | AN6352 | | AN4805 | AN2481 |
| | AN1320 | AN8900 | | AN4820 | AN2571 |
| | AN1575 | AN7033 | | AN4847 | AN2649 |
| | AN7263 | AN8356 | | AN4855 | AN2801 |
| | AN5034 | AN4018 | | AN4861 | AN2805 |
| | AN1580 | AN6704 | | AN4866 | AN2821 |
| | AN6847 | AN6850 | | AN4870 | AN2837 |
| | AN1318 | AN6776 | | AN4887 | AN3043 |
| | AN7061 | AN6025 | | AN4900 | AN3100 |
| | AN6659 | AN8613 | | AN4904 | AN3129 |
| | AN7866 | AN7780 | | AN4908 | AN3255 |
| | AN9315 | AN6315 | | AN4909 | AN3312 |
| | AN6509 | AN6399 | | AN4924 | AN3329 |
| | AN9242 | AN6941 | | AN4929 | AN3336 |
| | AN1317 | AN1092 | | AN4950 | AN3387 |
| | AN7808 | AN5947 | | AN4978 | AN3393 |
| | AN6751 | AN6029 | | AN5017 | AN3396 |
| | AN2038 | AN8238 | | AN5018 | AN3401 |
| | AN8543 | AN8077 | | AN5020 | AN3870 |
| | AN1574 | AN6168 | | AN5027 | AN3888 |
| | AN9203 | AN0768 | | AN5049 | AN4025 |
| | AN8992 | AN1703 | | AN5050 | AN4076 |
| | AN2336 | AN9301 | | AN5055 | AN4117 |
| | AN1279 | AN3858 | | AN5068 | AN4128 |
| | AN8651 | AN1315 | | AN5076 | AN4363 |
| | AN5316 | AN0531 | | AN5078 | AN4405 |
| | AN7229 | AN0488 | | AN5084 | AN4615 |
| | AN2846 | AN0606 | | AN5087 | AN5034 |
| | AN4157 | AN3872 | | AN5109 | AN5037 |
| | AN1378 | AN0220 | | AN5129 | AN5093 |
| | AN6781 | AN1616 | | AN5130 | AN5141 |
| | AN2027 | AN7708 | | AN5142 | AN5155 |
| | AN8376 | AN3361 | | AN5143 | AN5181 |
| | AN8364 | AN8781 | | AN5155 | AN5215 |
| | AN9439 | AN6024 | | AN5167 | AN5218 |
| | AN3235 | AN3194 | | AN5168 | AN5226 |
| | AN7973 | AN7129 | | AN5176 | AN5256 |
| | AN7410 | AN3132 | | AN5182 | AN5270 |
| | AN3971 | AN3334 | | AN5192 | AN5271 |
| | AN3888 | AN9179 | | AN5214 | AN5273 |
| | AN2340 | AN1304 | | AN5217 | AN5276 |
| | AN2697 | AN1607 | | AN5219 | AN5311 |
| | AN3886 | AN5189 | | AN5227 | AN5378 |
| | AN6664 | AN8640 | | AN5232 | AN5383 |
| | AN9277 | AN0786 | | AN5257 | AN5424 |
| | AN8143 | AN7018 | | AN5268 | AN5435 |
| | AN6833 | AN3043 | | AN5269 | AN5449 |
| | AN9457 | AN8638 | | AN5270 | AN5542 |
| | AN2595 | AN1918 | | AN5271 | AN5775 |
| | AN7281 | AN2361 | | AN5272 | AN5843 |
| | AN8328 | AN1185 | | AN5273 | AN6018 |
| | AN3282 | AN0400 | | AN5285 | AN6025 |
| | AN7832 | AN3021 | | AN5310 | AN6412 |
| | AN1585 | AN8277 | | AN5328 | AN6506 |
| | AN2343 | AN3265 | | AN5334 | AN6850 |
| | AN0008 | AN7621 | | AN5337 | AN6918 |
| | AN7317 | AN2707 | | AN5338 | AN6921 |
| | AN8412 | AN8046 | | AN5371 | AN6936 |
| | AN8397 | AN8760 | | AN5376 | AN6954 |
| | AN3988 | AN8816 | | AN5378 | AN7041 |
| | AN5030 | AN4905 | | AN5395 | AN7042 |
| | AN7828 | AN6739 | | AN5400 | AN7051 |
| | AN3561 | AN0788 | | AN5408 | AN7074 |
| | AN7870 | AN0395 | | AN5415 | AN7136 |

| | | | | | |
|---|---|---|---|---|---|
| | AN7070 | AN6969 | | AN5423 | AN7183 |
| | AN8558 | AN1311 | | AN5434 | AN7232 |
| | AN0397 | AN0723 | | AN5436 | AN7261 |
| | AN6235 | AN8727 | | AN5441 | AN7385 |
| | AN9317 | AN8907 | | AN5443 | AN7437 |
| | AN2378 | AN5781 | | AN5447 | AN7543 |
| | AN5320 | AN8893 | | AN5448 | AN7580 |
| | AN5023 | AN7593 | | AN5456 | AN7593 |
| | AN2339 | AN8397 | | AN5466 | AN7786 |
| | AN6636 | AN7635 | | AN5489 | AN7805 |
| | AN5334 | AN1840 | | AN5508 | AN7825 |
| | AN8124 | AN0152 | | AN5523 | AN7856 |
| | AN9353 | AN9127 | | AN5545 | AN7865 |
| | AN3273 | AN5437 | | AN5546 | AN7867 |
| | AN9351 | AN9441 | | AN5548 | AN8046 |
| | AN2286 | AN2040 | | AN5555 | AN8060 |
| | AN4774 | AN7186 | | AN5557 | AN8132 |
| | AN3253 | AN9347 | | AN5603 | AN8150 |
| | AN8544 | AN7949 | | AN5607 | AN8242 |
| | AN1838 | AN6048 | | AN5610 | AN8308 |
| | AN9042 | AN2535 | | AN5612 | AN8323 |
| | AN4970 | AN6836 | | AN5613 | AN8338 |
| | AN5831 | AN8738 | | AN5632 | AN8402 |
| | AN8374 | AN3790 | | AN5635 | AN8479 |
| | AN8360 | AN3298 | | AN5644 | AN8544 |
| | AN9046 | AN5764 | | AN5648 | AN8561 |
| | AN7600 | AN6918 | | AN5663 | AN8597 |
| | AN8803 | AN8099 | | AN5674 | AN8663 |
| | AN9243 | AN5550 | | AN5685 | AN8697 |
| | AN3570 | AN5729 | | AN5725 | AN8754 |
| | AN2034 | AN1841 | | AN5763 | AN8789 |
| | AN7270 | AN9499 | | AN5780 | AN8800 |
| | AN1808 | AN3100 | | AN5793 | AN8892 |
| | AN7054 | AN4485 | | AN5813 | AN8909 |
| | AN2035 | AN1077 | | AN5814 | AN8944 |
| | AN7933 | AN8374 | | AN5822 | AN8982 |
| | AN1198 | AN5604 | | AN5840 | AN9139 |
| | AN5383 | AN5172 | | AN5842 | AN9165 |
| | AN7051 | AN4807 | | AN5884 | AN0005 |
| | AN0374 | AN1112 | | AN5894 | AN0006 |
| | AN2750 | AN2445 | | AN5896 | AN0079 |
| | AN4506 | AN2675 | | AN5944 | AN0097 |
| | AN8131 | AN7109 | | AN5971 | AN0269 |
| | AN3283 | AN9041 | | AN5976 | AN0318 |
| | AN0546 | AN7280 | | AN5980 | AN0329 |
| | AN5408 | AN8903 | | AN5987 | AN0332 |
| | AN8451 | AN6381 | | AN5988 | AN0393 |
| | AN2941 | AN5432 | | AN6013 | AN0418 |
| | AN5089 | AN8993 | | AN6045 | AN0572 |
| | AN5319 | AN1744 | | AN6048 | AN0575 |
| | AN8773 | AN6813 | | AN6051 | AN0595 |
| | AN2198 | AN8658 | | AN6057 | AN0627 |
| | AN6796 | AN0858 | | AN6131 | AN0694 |
| | AN3316 | AN2613 | | AN6155 | AN0726 |
| | AN2618 | AN1994 | | AN6167 | AN0737 |
| | AN8105 | AN6030 | | AN6168 | AN0771 |
| | AN6830 | AN2890 | | AN6183 | AN0773 |
| | AN5918 | AN7716 | | AN6184 | AN0791 |
| | AN3521 | AN8616 | | AN6195 | AN0822 |
| | AN7402 | AN7081 | | AN6203 | AN0825 |
| | AN7265 | AN2195 | | AN6232 | AN0856 |
| | AN8269 | AN2310 | | AN6245 | AN0941 |
| | AN7023 | AN7247 | | AN6276 | AN0964 |
| | AN5037 | AN4795 | | AN6277 | AN0979 |
| | AN0542 | AN4625 | | AN6280 | AN1025 |
| | AN1914 | AN5616 | | AN6306 | AN1089 |

| | | | | | |
|---|---|---|---|---|---|
| | AN6154 | AN0918 | | AN6313 | AN1092 |
| | AN8527 | AN6046 | | AN6314 | AN1274 |
| | AN8948 | AN5050 | | AN6323 | AN1373 |
| | AN5370 | AN1303 | | AN6336 | AN1401 |
| | AN9224 | AN7484 | | AN6343 | AN1508 |
| | AN7814 | AN4773 | | AN6345 | AN1510 |
| | AN0683 | AN4081 | | AN6350 | AN1541 |
| | AN8997 | AN0553 | | AN6354 | AN1551 |
| | AN7960 | AN2230 | | AN6360 | AN1556 |
| | AN7153 | AN1418 | | AN6365 | AN1573 |
| | AN0511 | AN7451 | | AN6376 | AN1681 |
| | AN7058 | AN4691 | | AN6377 | AN1754 |
| | AN7790 | AN1123 | | AN6386 | AN1783 |
| | AN8137 | AN5335 | | AN6397 | AN1898 |
| | AN2387 | AN2668 | | AN6398 | AN2003 |
| | AN1829 | AN4535 | | AN6400 | AN2036 |
| | AN0618 | AN6314 | | AN6413 | AN2042 |
| | AN8947 | AN6135 | | AN6505 | AN2051 |
| | AN3410 | AN8199 | | AN6524 | AN2111 |
| | AN5064 | AN0353 | | AN6600 | AN2169 |
| | AN2678 | AN1563 | | AN6617 | AN2241 |
| | AN7063 | AN0299 | | AN6620 | AN2243 |
| | AN8265 | AN8089 | | AN6632 | AN2314 |
| | AN2714 | AN3136 | | AN6637 | AN2385 |
| | AN1630 | AN0638 | | AN6639 | AN2404 |
| | AN7137 | AN2468 | | AN6641 | AN2407 |
| | AN1321 | AN3606 | | AN6644 | AN2424 |
| | AN6661 | AN7221 | | AN6652 | AN2469 |
| | AN0548 | AN1505 | | AN6654 | AN2587 |
| | AN3327 | AN2801 | | AN6656 | AN2604 |
| | AN8593 | AN3697 | | AN6700 | AN2648 |
| | AN5310 | AN5458 | | AN6703 | AN2664 |
| | AN0509 | AN2114 | | AN6720 | AN2684 |
| | AN3406 | AN4649 | | AN6732 | AN2727 |
| | AN8251 | AN6645 | | AN6775 | AN2751 |
| | AN3474 | AN8133 | | AN6788 | AN2855 |
| | AN1309 | AN4527 | | AN6791 | AN2898 |
| | AN3975 | AN5624 | | AN6808 | AN2955 |
| | AN9175 | AN5524 | | AN6809 | AN3080 |
| | AN7692 | AN2395 | | AN6814 | AN3140 |
| | AN2028 | AN0942 | | AN6846 | AN3191 |
| | AN3396 | AN1136 | | AN6849 | AN3200 |
| | AN8666 | AN7187 | | AN6857 | AN3206 |
| | AN3325 | AN0895 | | AN6860 | AN3249 |
| | AN9311 | AN8683 | | AN6865 | AN3285 |
| | AN7876 | AN3672 | | AN6880 | AN3390 |
| | AN5942 | AN7124 | | AN6890 | AN3406 |
| | AN5936 | AN0221 | | AN6898 | AN3425 |
| | AN3494 | AN4224 | | AN6899 | AN3496 |
| | AN6237 | AN0188 | | AN6917 | AN3497 |
| | AN2120 | AN0437 | | AN6920 | AN3518 |
| | AN2036 | AN7786 | | AN6923 | AN3628 |
| | AN2010 | AN6848 | | AN6935 | AN3643 |
| | AN9345 | AN6640 | | AN6940 | AN3667 |
| | AN3967 | AN1067 | | AN6943 | AN3758 |
| | AN0012 | AN8222 | | AN6950 | AN3815 |
| | AN8547 | AN7936 | | AN6961 | AN3848 |
| | AN7136 | AN4041 | | AN6968 | AN3905 |
| | AN7897 | AN4503 | | AN7019 | AN3961 |
| | AN8555 | AN7709 | | AN7020 | AN3983 |
| | AN6946 | AN1040 | | AN7032 | AN4019 |
| | AN0029 | AN5256 | | AN7034 | AN4187 |
| | AN0946 | AN5768 | | AN7050 | AN4299 |
| | AN4769 | AN9051 | | AN7052 | AN4328 |
| | AN0619 | AN0718 | | AN7057 | AN4345 |
| | AN0639 | AN6770 | | AN7073 | AN4349 |

| | | | | | |
|---|---|---|---|---|---|
| | AN2648 | AN1646 | | AN7080 | AN4361 |
| | AN7851 | AN3624 | | AN7111 | AN4395 |
| | AN8106 | AN0764 | | AN7113 | AN4463 |
| | AN7800 | AN8355 | | AN7133 | AN4515 |
| | AN8556 | AN7113 | | AN7137 | AN4527 |
| | AN7211 | AN0145 | | AN7148 | AN4547 |
| | AN5076 | AN2800 | | AN7189 | AN4592 |
| | AN0010 | AN5321 | | AN7190 | AN4621 |
| | AN0403 | AN7815 | | AN7193 | AN4808 |
| | AN7089 | AN3849 | | AN7197 | AN4812 |
| | AN7837 | AN2323 | | AN7232 | AN4862 |
| | AN2365 | AN8221 | | AN7248 | AN4901 |
| | AN3613 | AN8737 | | AN7262 | AN4909 |
| | AN7523 | AN8390 | | AN7265 | AN4921 |
| | AN4152 | AN2730 | | AN7266 | AN4940 |
| | AN9454 | AN3151 | | AN7267 | AN4988 |
| | AN2842 | AN7263 | | AN7277 | AN5069 |
| | AN4607 | AN8740 | | AN7294 | AN5088 |
| | AN3873 | AN9121 | | AN7301 | AN5092 |
| | AN8510 | AN4195 | | AN7321 | AN5205 |
| | AN9044 | AN7082 | | AN7332 | AN5207 |
| | AN7831 | AN2538 | | AN7384 | AN5225 |
| | AN8982 | AN0157 | | AN7385 | AN5259 |
| | AN2621 | AN5284 | | AN7406 | AN5335 |
| | AN2593 | AN8610 | | AN7427 | AN5338 |
| | AN0530 | AN0899 | | AN7483 | AN5370 |
| | AN4770 | AN5169 | | AN7501 | AN5396 |
| | AN5929 | AN4078 | | AN7510 | AN5423 |
| | AN9339 | AN7703 | | AN7511 | AN5467 |
| | AN8548 | AN2326 | | AN7514 | AN5478 |
| | AN2025 | AN9291 | | AN7529 | AN5493 |
| | AN1278 | AN7622 | | AN7567 | AN5636 |
| | AN9524 | AN1506 | | AN7576 | AN5690 |
| | AN8341 | AN5002 | | AN7590 | AN5768 |
| | AN8348 | AN6412 | | AN7591 | AN5776 |
| | AN3242 | AN2199 | | AN7592 | AN5823 |
| | AN8458 | AN8336 | | AN7618 | AN5832 |
| | AN3508 | AN2625 | | AN7620 | AN5940 |
| | AN6869 | AN1787 | | AN7621 | AN5944 |
| | AN2706 | AN5537 | | AN7624 | AN6005 |
| | AN6403 | AN4940 | | AN7656 | AN6017 |
| | AN8617 | AN3889 | | AN7657 | AN6052 |
| | AN5244 | AN2537 | | AN7660 | AN6118 |
| | AN5378 | AN8262 | | AN7667 | AN6156 |
| | AN8337 | AN6653 | | AN7668 | AN6208 |
| | AN7112 | AN8538 | | AN7682 | AN6234 |
| | AN4975 | AN1269 | | AN7701 | AN6281 |
| | AN0527 | AN8018 | | AN7702 | AN6286 |
| | AN7841 | AN5977 | | AN7707 | AN6361 |
| | AN7053 | AN1605 | | AN7708 | AN6525 |
| | AN6753 | AN1549 | | AN7710 | AN6527 |
| | AN7891 | AN5556 | | AN7713 | AN6621 |
| | AN0009 | AN3512 | | AN7720 | AN6680 |
| | AN7278 | AN7989 | | AN7759 | AN6691 |
| | AN9055 | AN4749 | | AN7775 | AN6704 |
| | AN1262 | AN7943 | | AN7778 | AN6713 |
| | AN5650 | AN2024 | | AN7779 | AN6734 |
| | AN2192 | AN1681 | | AN7784 | AN6761 |
| | AN0056 | AN8627 | | AN7792 | AN6847 |
| | AN8998 | AN2805 | | AN7793 | AN6875 |
| | AN2574 | AN8465 | | AN7795 | AN6930 |
| | AN8622 | AN7826 | | AN7796 | AN6941 |
| | AN5368 | AN9267 | | AN7798 | AN6942 |
| | AN4051 | AN3891 | | AN7799 | AN7211 |
| | AN1615 | AN3892 | | AN7804 | AN7246 |
| | AN6621 | AN8944 | | AN7831 | AN7251 |

| | | | | | |
|---|---|---|---|---|---|
| | AN0400 | AN6404 | | AN7864 | AN7296 |
| | AN6409 | AN3558 | | AN7876 | AN7335 |
| | AN2642 | AN8471 | | AN7877 | AN7390 |
| | AN3162 | AN5083 | | AN7879 | AN7459 |
| | AN0565 | AN6792 | | AN7882 | AN7480 |
| | AN8642 | AN5555 | | AN7924 | AN7514 |
| | AN7900 | AN7295 | | AN7935 | AN7558 |
| | AN2116 | AN7021 | | AN7941 | AN7590 |
| | AN3252 | AN2039 | | AN7951 | AN7594 |
| | AN2813 | AN9324 | | AN7952 | AN7708 |
| | AN3256 | AN7266 | | AN7953 | AN7709 |
| | AN3780 | AN9197 | | AN7954 | AN7724 |
| | AN2722 | AN3203 | | AN7988 | AN7838 |
| | AN3558 | AN0610 | | AN7992 | AN7999 |
| | AN7983 | AN7753 | | AN7994 | AN8156 |
| | AN8114 | AN3782 | | AN8004 | AN8262 |
| | AN9261 | AN5017 | | AN8006 | AN8277 |
| | AN2407 | AN5417 | | AN8007 | AN8278 |
| | AN0283 | AN1615 | | AN8042 | AN8342 |
| | AN4604 | AN1322 | | AN8045 | AN8439 |
| | AN2928 | AN2197 | | AN8084 | AN8482 |
| | AN0022 | AN2724 | | AN8093 | AN8495 |
| | AN2694 | AN7394 | | AN8115 | AN8547 |
| | AN5545 | AN5427 | | AN8145 | AN8548 |
| | AN8132 | AN3560 | | AN8149 | AN8666 |
| | AN9266 | AN8995 | | AN8154 | AN8714 |
| | AN7385 | AN0215 | | AN8165 | AN8739 |
| | AN3980 | AN2200 | | AN8221 | AN8798 |
| | AN7875 | AN4108 | | AN8227 | AN8825 |
| | AN0148 | AN4142 | | AN8242 | AN8929 |
| | AN8473 | AN2321 | | AN8245 | AN8943 |
| | AN7965 | AN4107 | | AN8261 | AN9003 |
| | AN2561 | AN4129 | | AN8264 | AN9042 |
| | AN8314 | AN4173 | | AN8276 | AN9088 |
| | AN7854 | AN6949 | | AN8307 | AN9160 |
| | AN8351 | AN9490 | | AN8313 | AN9163 |
| | AN9036 | AN3888 | | AN8326 | AN9172 |
| | AN2695 | AN6847 | | AN8333 | AN9192 |
| | AN8515 | AN5218 | | AN8339 | AN9262 |
| | AN7836 | AN7600 | | AN8340 | AN9285 |
| | AN6408 | AN1752 | | AN8341 | AN9296 |
| | AN6648 | AN3887 | | AN8347 | AN9348 |
| | AN8552 | AN7051 | | AN8348 | AN9364 |
| | AN8438 | AN1692 | | AN8352 | AN9412 |
| | AN7825 | AN2576 | | AN8355 | AN9510 |
| | AN9263 | AN8596 | | AN8365 | AN0009 |
| | AN8363 | AN9139 | | AN8366 | AN0011 |
| | AN7020 | AN2641 | | AN8367 | AN0200 |
| | AN8779 | AN1573 | | AN8369 | AN0232 |
| | AN8540 | AN7797 | | AN8373 | AN0268 |
| | AN4337 | AN0860 | | AN8391 | AN0300 |
| | AN8597 | AN9165 | | AN8398 | AN0382 |
| | AN0011 | AN3406 | | AN8444 | AN0403 |
| | AN5544 | AN3233 | | AN8445 | AN0523 |
| | AN8944 | AN0799 | | AN8454 | AN0562 |
| | AN3321 | AN5615 | | AN8458 | AN0609 |
| | AN6356 | AN5334 | | AN8461 | AN0660 |
| | AN2200 | AN8376 | | AN8462 | AN0693 |
| | AN0187 | AN3494 | | AN8477 | AN0815 |
| | AN9012 | AN3989 | | AN8478 | AN0830 |
| | AN7906 | AN7073 | | AN8529 | AN0879 |
| | AN6541 | AN0639 | | AN8537 | AN0887 |
| | AN3390 | AN1672 | | AN8546 | AN0950 |
| | AN7693 | AN2783 | | AN8547 | AN0973 |
| | AN3400 | AN3557 | | AN8567 | AN1061 |
| | AN9088 | AN7154 | | AN8590 | AN1182 |

| | | | | | |
|---|---|---|---|---|---|
| | AN7266 | AN6649 | | AN8593 | AN1195 |
| | AN4363 | AN7557 | | AN8595 | AN1220 |
| | AN2569 | AN2656 | | AN8596 | AN1262 |
| | AN9381 | AN8184 | | AN8607 | AN1322 |
| | AN8922 | AN9457 | | AN8609 | AN1326 |
| | AN3329 | AN6959 | | AN8612 | AN1338 |
| | AN0214 | AN6935 | | AN8615 | AN1385 |
| | AN2832 | AN3234 | | AN8618 | AN1427 |
| | AN7417 | AN2118 | | AN8625 | AN1528 |
| | AN0621 | AN7999 | | AN8629 | AN1542 |
| | AN8090 | AN8619 | | AN8633 | AN1561 |
| | | AN3247 | | AN8640 | AN1593 |
| | | AN9220 | | AN8650 | AN1631 |
| | | AN9219 | | AN8660 | AN1692 |
| | | AN8958 | | AN8678 | AN1694 |
| | | AN2655 | | AN8691 | AN1742 |
| | | AN9191 | | AN8692 | AN1781 |
| | | AN9190 | | AN8709 | AN1869 |
| | | AN2698 | | AN8725 | AN2015 |
| | | AN9372 | | AN8726 | AN2056 |
| | | AN9213 | | AN8739 | AN2058 |
| | | AN9375 | | AN8740 | AN2100 |
| | | AN8348 | | AN8763 | AN2116 |
| | | AN5976 | | AN8770 | AN2164 |
| | | AN8340 | | AN8780 | AN2228 |
| | | AN8595 | | AN8789 | AN2300 |
| | | AN2781 | | AN8796 | AN2390 |
| | | AN1199 | | AN8797 | AN2393 |
| | | AN2110 | | AN8830 | AN2398 |
| | | AN8904 | | AN8841 | AN2468 |
| | | AN4105 | | AN8872 | AN2471 |
| | | AN3239 | | AN8877 | AN2525 |
| | | AN8084 | | AN8892 | AN2534 |
| | | AN2560 | | AN8893 | AN2576 |
| | | AN2187 | | AN8903 | AN2596 |
| | | AN2559 | | AN8906 | AN2609 |
| | | AN9004 | | AN8909 | AN2641 |
| | | AN8593 | | AN8923 | AN2683 |
| | | AN7702 | | AN8933 | AN2746 |
| | | AN3996 | | AN8938 | AN2783 |
| | | AN8158 | | AN8940 | AN2867 |
| | | AN7962 | | AN8944 | AN2868 |
| | | AN3553 | | AN8945 | AN2924 |
| | | AN8220 | | AN8957 | AN2928 |
| | | | | AN8958 | AN2983 |
| | | | | AN8962 | AN3138 |
| | | | | AN8963 | AN3149 |
| | | | | AN8967 | AN3185 |
| | | | | AN8985 | AN3276 |
| | | | | AN8993 | AN3307 |
| | | | | AN8994 | AN3308 |
| | | | | AN9006 | AN3321 |
| | | | | AN9048 | AN3333 |
| | | | | AN9050 | AN3345 |
| | | | | AN9067 | AN3428 |
| | | | | AN9069 | AN3431 |
| | | | | AN9093 | AN3508 |
| | | | | AN9104 | AN3517 |
| | | | | AN9107 | AN3522 |
| | | | | AN9110 | AN3573 |
| | | | | AN9115 | AN3608 |
| | | | | AN9120 | AN3638 |
| | | | | AN9126 | AN3765 |
| | | | | AN9142 | AN3771 |
| | | | | AN9148 | AN3889 |
| | | | | AN9151 | AN3903 |

| | | | | | |
|---|---|---|---|---|---|
| | | | | AN9163<br>AN9173<br>AN9187<br>AN9190<br>AN9205<br>AN9219<br>AN9258<br>AN9284<br>AN9300<br>AN9301<br>AN9331<br>AN9339<br>AN9371<br>AN9372<br>AN9375<br>AN9378<br>AN9391<br>AN9392<br>AN9396<br>AN9465<br>AN9472<br>AN9480<br>AN9496<br>AN9498<br>AN9521<br>AN0008<br>AN0028<br>AN0030<br>AN0051<br>AN0056<br>AN0069<br>AN0083<br>AN0087<br>AN0144<br>AN0151<br>AN0167<br>AN0168<br>AN0174<br>AN0180<br>AN0184<br>AN0187<br>AN0220<br>AN0306<br>AN0329<br>AN0335<br>AN0391<br>AN0416<br>AN0437<br>AN0467<br>AN0479<br>AN0494<br>AN0502<br>AN0601<br>AN0653<br>AN0693<br>AN0708<br>AN0717<br>AN0724<br>AN0731<br>AN0769<br>AN0822<br>AN0857<br>AN0877<br>AN0883<br>AN0912<br>AN0921 | AN3979<br>AN4024<br>AN4049<br>AN4070<br>AN4159<br>AN4171<br>AN4270<br>AN4305<br>AN4376<br>AN4541<br>AN4622<br>AN4642<br>AN4671<br>AN4691<br>AN4733<br>AN4757<br>AN4774<br>AN4791<br>AN4842<br>AN4974<br>AN5019<br>AN5082<br>AN5123<br>AN5145<br>AN5168<br>AN5175<br>AN5185<br>AN5232<br>AN5272<br>AN5348<br>AN5444<br>AN5515<br>AN5537<br>AN5659<br>AN5665<br>AN5710<br>AN5885<br>AN5976<br>AN6001<br>AN6108<br>AN6115<br>AN6121<br>AN6128<br>AN6140<br>AN6235<br>AN6236<br>AN6237<br>AN6239<br>AN6315<br>AN6394<br>AN6399<br>AN6401<br>AN6530<br>AN6642<br>AN6653<br>AN6654<br>AN6791<br>AN6792<br>AN6816<br>AN6830<br>AN6857<br>AN6943<br>AN6949<br>AN7033<br>AN7128<br>AN7175 |

| | | | | | |
|---|---|---|---|---|---|
| | | | | AN0937<br>AN1034<br>AN1064<br>AN1065<br>AN1073<br>AN1114<br>AN1197<br>AN1268<br>AN1281<br>AN1341<br>AN1463<br>AN1539<br>AN1569<br>AN1571<br>AN1573<br>AN1582<br>AN1600<br>AN1646<br>AN1649<br>AN1656<br>AN1742<br>AN1817<br>AN1834<br>AN1840<br>AN1863<br>AN1881<br>AN1895<br>AN1908<br>AN1990<br>AN2000<br>AN2020<br>AN2032<br>AN2033<br>AN2071<br>AN2125<br>AN2188<br>AN2195<br>AN2229<br>AN2256<br>AN2285<br>AN2303<br>AN2316<br>AN2319<br>AN2342<br>AN2350<br>AN2384<br>AN2397<br>AN2403<br>AN2440<br>AN2479<br>AN2536<br>AN2555<br>AN2556<br>AN2558<br>AN2559<br>AN2560<br>AN2607<br>AN2674<br>AN2693<br>AN2700<br>AN2701<br>AN2745<br>AN2782<br>AN2797<br>AN2800<br>AN2826 | AN7215<br>AN7233<br>AN7267<br>AN7333<br>AN7407<br>AN7413<br>AN7537<br>AN7567<br>AN7570<br>AN7610<br>AN7634<br>AN7742<br>AN7797<br>AN7822<br>AN7832<br>AN7834<br>AN7881<br>AN7898<br>AN7936<br>AN7987<br>AN8085<br>AN8120<br>AN8123<br>AN8182<br>AN8200<br>AN8215<br>AN8222<br>AN8327<br>AN8341<br>AN8348<br>AN8365<br>AN8366<br>AN8390<br>AN8392<br>AN8438<br>AN8528<br>AN8536<br>AN8565<br>AN8608<br>AN8619<br>AN8650<br>AN8677<br>AN8779<br>AN8781<br>AN8854<br>AN8894<br>AN8930<br>AN9001<br>AN9089<br>AN9116<br>AN9177<br>AN9197<br>AN9375<br>AN9473<br>AN0112<br>AN0163<br>AN0229<br>AN0231<br>AN0234<br>AN0239<br>AN0261<br>AN0270<br>AN0285<br>AN0319<br>AN0404<br>AN0431 |

| | | | | | |
|---|---|---|---|---|---|
| | | | | AN2836 | AN0467 |
| | | | | AN2854 | AN0468 |
| | | | | AN2865 | AN0648 |
| | | | | AN2871 | AN0673 |
| | | | | AN2912 | AN0675 |
| | | | | AN2927 | AN0688 |
| | | | | AN2959 | AN0736 |
| | | | | AN2986 | AN0780 |
| | | | | AN3064 | AN0797 |
| | | | | AN3070 | AN0875 |
| | | | | AN3090 | AN0933 |
| | | | | AN3149 | AN0937 |
| | | | | AN3220 | AN0958 |
| | | | | AN3244 | AN0991 |
| | | | | AN3252 | AN1039 |
| | | | | AN3270 | AN1040 |
| | | | | AN3309 | AN1044 |
| | | | | AN3320 | AN1066 |
| | | | | AN3333 | AN1116 |
| | | | | AN3337 | AN1158 |
| | | | | AN3346 | AN1160 |
| | | | | AN3365 | AN1211 |
| | | | | AN3395 | AN1243 |
| | | | | AN3413 | AN1261 |
| | | | | AN3427 | AN1271 |
| | | | | AN3432 | AN1283 |
| | | | | AN3480 | AN1304 |
| | | | | AN3506 | AN1310 |
| | | | | AN3511 | AN1312 |
| | | | | AN3593 | AN1325 |
| | | | | AN3603 | AN1396 |
| | | | | AN3626 | AN1423 |
| | | | | AN3627 | AN1446 |
| | | | | AN3638 | AN1449 |
| | | | | AN3663 | AN1493 |
| | | | | AN3673 | AN1523 |
| | | | | AN3718 | AN1586 |
| | | | | AN3762 | AN1606 |
| | | | | AN3813 | AN1612 |
| | | | | AN3818 | AN1614 |
| | | | | AN3819 | AN1638 |
| | | | | AN3828 | AN1664 |
| | | | | AN3978 | AN1680 |
| | | | | AN4060 | AN1691 |
| | | | | AN4076 | AN1695 |
| | | | | AN4093 | AN1714 |
| | | | | AN4120 | AN1730 |
| | | | | AN4138 | AN1760 |
| | | | | AN4141 | AN1865 |
| | | | | AN4239 | AN1895 |
| | | | | AN4247 | AN1911 |
| | | | | AN4298 | AN1926 |
| | | | | AN4363 | AN1969 |
| | | | | AN4417 | AN1991 |
| | | | | AN4486 | AN2064 |
| | | | | AN4516 | AN2068 |
| | | | | AN4538 | AN2099 |
| | | | | AN4588 | AN2110 |
| | | | | AN4591 | AN2126 |
| | | | | AN4694 | AN2133 |
| | | | | AN4752 | AN2165 |
| | | | | AN4771 | AN2187 |
| | | | | AN4774 | AN2194 |
| | | | | AN4823 | AN2201 |
| | | | | AN4836 | AN2223 |
| | | | | AN4920 | AN2237 |

| | | | | | |
|---|---|---|---|---|---|
| | | | | AN4939 | AN2240 |
| | | | | AN4949 | AN2287 |
| | | | | AN4996 | AN2321 |
| | | | | AN5057 | AN2350 |
| | | | | AN5108 | AN2384 |
| | | | | AN5140 | AN2387 |
| | | | | AN5161 | AN2441 |
| | | | | AN5230 | AN2458 |
| | | | | AN5256 | AN2484 |
| | | | | AN5280 | AN2486 |
| | | | | AN5311 | AN2509 |
| | | | | AN5351 | AN2537 |
| | | | | AN5426 | AN2574 |
| | | | | AN5444 | AN2594 |
| | | | | AN5486 | AN2640 |
| | | | | AN5487 | AN2659 |
| | | | | AN5507 | AN2668 |
| | | | | AN5521 | AN2702 |
| | | | | AN5529 | AN2774 |
| | | | | AN5567 | AN2781 |
| | | | | AN5574 | AN2825 |
| | | | | AN5601 | AN2851 |
| | | | | AN5662 | AN2864 |
| | | | | AN5664 | AN2875 |
| | | | | AN5676 | AN2882 |
| | | | | AN5696 | AN2962 |
| | | | | AN5712 | AN2968 |
| | | | | AN5751 | AN2989 |
| | | | | AN5752 | AN2999 |
| | | | | AN5754 | AN3014 |
| | | | | AN5767 | AN3023 |
| | | | | AN5809 | AN3029 |
| | | | | AN5819 | AN3119 |
| | | | | AN5820 | AN3132 |
| | | | | AN5844 | AN3137 |
| | | | | AN5845 | AN3139 |
| | | | | AN5868 | AN3180 |
| | | | | AN5939 | AN3234 |
| | | | | AN5943 | AN3294 |
| | | | | AN5975 | AN3339 |
| | | | | AN5986 | AN3384 |
| | | | | AN6022 | AN3385 |
| | | | | AN6047 | AN3440 |
| | | | | AN6058 | AN3456 |
| | | | | AN6088 | AN3479 |
| | | | | AN6092 | AN3519 |
| | | | | AN6104 | AN3520 |
| | | | | AN6165 | AN3557 |
| | | | | AN6224 | AN3578 |
| | | | | AN6225 | AN3609 |
| | | | | AN6235 | AN3644 |
| | | | | AN6248 | AN3675 |
| | | | | AN6380 | AN3680 |
| | | | | AN6494 | AN3702 |
| | | | | AN6531 | AN3720 |
| | | | | AN6533 | AN3743 |
| | | | | AN6541 | AN3748 |
| | | | | AN6546 | AN3789 |
| | | | | AN6552 | AN3814 |
| | | | | AN6623 | AN3819 |
| | | | | AN6730 | AN3821 |
| | | | | AN6733 | AN3839 |
| | | | | AN6735 | AN3887 |
| | | | | AN6738 | AN3906 |
| | | | | AN6739 | AN3916 |
| | | | | AN6769 | AN3954 |

| | | | | | |
|---|---|---|---|---|---|
| | | | | AN6787 | AN4017 |
| | | | | AN6806 | AN4018 |
| | | | | AN6811 | AN4045 |
| | | | | AN6812 | AN4053 |
| | | | | AN6818 | AN4079 |
| | | | | AN6914 | AN4105 |
| | | | | AN6932 | AN4142 |
| | | | | AN6972 | AN4173 |
| | | | | AN6982 | AN4174 |
| | | | | AN6986 | AN4180 |
| | | | | AN7055 | AN4190 |
| | | | | AN7081 | AN4250 |
| | | | | AN7083 | AN4303 |
| | | | | AN7158 | AN4321 |
| | | | | AN7165 | AN4372 |
| | | | | AN7174 | AN4423 |
| | | | | AN7186 | AN4426 |
| | | | | AN7216 | AN4453 |
| | | | | AN7224 | AN4462 |
| | | | | AN7286 | AN4495 |
| | | | | AN7289 | AN4546 |
| | | | | AN7322 | AN4590 |
| | | | | AN7326 | AN4623 |
| | | | | AN7333 | AN4640 |
| | | | | AN7335 | AN4663 |
| | | | | AN7351 | AN4724 |
| | | | | AN7450 | AN4847 |
| | | | | AN7458 | AN4952 |
| | | | | AN7467 | AN5068 |
| | | | | AN7526 | AN5075 |
| | | | | AN7539 | AN5083 |
| | | | | AN7540 | AN5085 |
| | | | | AN7556 | AN5200 |
| | | | | AN7569 | AN5220 |
| | | | | AN7598 | AN5224 |
| | | | | AN7663 | AN5233 |
| | | | | AN7679 | AN5269 |
| | | | | AN7690 | AN5321 |
| | | | | AN7771 | AN5324 |
| | | | | AN7783 | AN5339 |
| | | | | AN7817 | AN5343 |
| | | | | AN7821 | AN5351 |
| | | | | AN7878 | AN5400 |
| | | | | AN7880 | AN5437 |
| | | | | AN7891 | AN5445 |
| | | | | AN7895 | AN5448 |
| | | | | AN7940 | AN5488 |
| | | | | AN7970 | AN5558 |
| | | | | AN7998 | AN5563 |
| | | | | AN8003 | AN5586 |
| | | | | AN8009 | AN5587 |
| | | | | AN8017 | AN5608 |
| | | | | AN8029 | AN5658 |
| | | | | AN8038 | AN5698 |
| | | | | AN8071 | AN5726 |
| | | | | AN8088 | AN5742 |
| | | | | AN8098 | AN5748 |
| | | | | AN8121 | AN5755 |
| | | | | AN8122 | AN5787 |
| | | | | AN8135 | AN5791 |
| | | | | AN8136 | AN5810 |
| | | | | AN8198 | AN5834 |
| | | | | AN8200 | AN5841 |
| | | | | AN8217 | AN5859 |
| | | | | AN8229 | AN5895 |
| | | | | AN8269 | AN5972 |

| | | | | | |
|---|---|---|---|---|---|
| | | | | AN8280 | AN5977 |
| | | | | AN8290 | AN6061 |
| | | | | AN8303 | AN6219 |
| | | | | AN8337 | AN6246 |
| | | | | AN8344 | AN6257 |
| | | | | AN8346 | AN6262 |
| | | | | AN8361 | AN6344 |
| | | | | AN8362 | AN6400 |
| | | | | AN8383 | AN6523 |
| | | | | AN8530 | AN6532 |
| | | | | AN8535 | AN6548 |
| | | | | AN8559 | AN6553 |
| | | | | AN8585 | AN6598 |
| | | | | AN8636 | AN6615 |
| | | | | AN8644 | AN6655 |
| | | | | AN8682 | AN6688 |
| | | | | AN8731 | AN6776 |
| | | | | AN8769 | AN6812 |
| | | | | AN8799 | AN6815 |
| | | | | AN8883 | AN6853 |
| | | | | AN8891 | AN6901 |
| | | | | AN8899 | AN6917 |
| | | | | AN9003 | AN6935 |
| | | | | AN9066 | AN6959 |
| | | | | AN9149 | AN7008 |
| | | | | AN9162 | AN7019 |
| | | | | AN9179 | AN7073 |
| | | | | AN9212 | AN7108 |
| | | | | AN9241 | AN7110 |
| | | | | AN9297 | AN7124 |
| | | | | AN9303 | AN7141 |
| | | | | AN9311 | AN7208 |
| | | | | AN9338 | AN7222 |
| | | | | AN9485 | AN7228 |
| | | | | AN0074 | AN7299 |
| | | | | AN0149 | AN7322 |
| | | | | AN0206 | AN7323 |
| | | | | AN0274 | AN7576 |
| | | | | AN0279 | AN7615 |
| | | | | AN0326 | AN7631 |
| | | | | AN0364 | AN7636 |
| | | | | AN0426 | AN7659 |
| | | | | AN0445 | AN7668 |
| | | | | AN0562 | AN7673 |
| | | | | AN0621 | AN7680 |
| | | | | AN0640 | AN7721 |
| | | | | AN0642 | AN7783 |
| | | | | AN0666 | AN7826 |
| | | | | AN0730 | AN7855 |
| | | | | AN0753 | AN7875 |
| | | | | AN0768 | AN7876 |
| | | | | AN0809 | AN7879 |
| | | | | AN1002 | AN7882 |
| | | | | AN1023 | AN7884 |
| | | | | AN1048 | AN7896 |
| | | | | AN1210 | AN7917 |
| | | | | AN1273 | AN7942 |
| | | | | AN1384 | AN7953 |
| | | | | AN1448 | AN7955 |
| | | | | AN1460 | AN8002 |
| | | | | AN1479 | AN8021 |
| | | | | AN1513 | AN8026 |
| | | | | AN1560 | AN8134 |
| | | | | AN1624 | AN8135 |
| | | | | AN1690 | AN8137 |
| | | | | AN1768 | AN8149 |

| | | | | | |
|---|---|---|---|---|---|
| | | | | AN1799<br>AN1804<br>AN1809<br>AN1812<br>AN1864<br>AN1872<br>AN1966<br>AN2041<br>AN2145<br>AN2150<br>AN2161<br>AN2164<br>AN2214<br>AN2325<br>AN2335<br>AN2386<br>AN2387<br>AN2389<br>AN2453<br>AN2519<br>AN2530<br>AN2537<br>AN2551<br>AN2597<br>AN2622<br>AN2761<br>AN2881<br>AN2891<br>AN2978<br>AN3033<br>AN3130<br>AN3187<br>AN3205<br>AN3263<br>AN3393<br>AN3481<br>AN3519<br>AN3611<br>AN3635<br>AN3649<br>AN3685<br>AN3724<br>AN3742<br>AN3831<br>AN3898<br>AN3905<br>AN3951<br>AN3962<br>AN3991<br>AN4006<br>AN4035<br>AN4048<br>AN4118<br>AN4126<br>AN4132<br>AN4179<br>AN4186<br>AN4192<br>AN4233<br>AN4257<br>AN4263<br>AN4316<br>AN4401<br>AN4414<br>AN4479<br>AN4518 | AN8158<br>AN8181<br>AN8203<br>AN8217<br>AN8230<br>AN8241<br>AN8340<br>AN8347<br>AN8370<br>AN8455<br>AN8538<br>AN8546<br>AN8596<br>AN8610<br>AN8613<br>AN8630<br>AN8653<br>AN8678<br>AN8709<br>AN8727<br>AN8790<br>AN8904<br>AN8941<br>AN8968<br>AN8995<br>AN9048<br>AN9079<br>AN9094<br>AN9121<br>AN9144<br>AN9161<br>AN9213<br>AN9219<br>AN9220<br>AN9242<br>AN9289<br>AN9369<br>AN9378<br>AN9438<br>AN9441<br>AN9490<br>AN9521<br>AN0075<br>AN0080<br>AN0084<br>AN0088<br>AN0115<br>AN0145<br>AN0156<br>AN0179<br>AN0184<br>AN0242<br>AN0275<br>AN0280<br>AN0286<br>AN0364<br>AN0366<br>AN0374<br>AN0378<br>AN0392<br>AN0402<br>AN0427<br>AN0488<br>AN0539<br>AN0546<br>AN0610 |

| | | | | | |
|---|---|---|---|---|---|
| | | | | AN4523<br>AN4690<br>AN4719<br>AN4778<br>AN4879<br>AN4919<br>AN4933<br>AN5032<br>AN5089<br>AN5131<br>AN5148<br>AN5170<br>AN5183<br>AN5293<br>AN5320<br>AN5324<br>AN5353<br>AN5402<br>AN5412<br>AN5549<br>AN5596<br>AN5615<br>AN5633<br>AN5666<br>AN5667<br>AN5694<br>AN5772<br>AN5885<br>AN5892<br>AN5989<br>AN5999<br>AN6001<br>AN6024<br>AN6074<br>AN6096<br>AN6120<br>AN6218<br>AN6243<br>AN6247<br>AN6270<br>AN6321<br>AN6385<br>AN6504<br>AN6506<br>AN6619<br>AN6718<br>AN6803<br>AN6847<br>AN7135<br>AN7138<br>AN7185<br>AN7242<br>AN7253<br>AN7368<br>AN7375<br>AN7446<br>AN7509<br>AN7568<br>AN7726<br>AN7850<br>AN7858<br>AN7903<br>AN7990<br>AN8022<br>AN8052<br>AN8134 | AN0622<br>AN0667<br>AN0725<br>AN0761<br>AN0770<br>AN0786<br>AN0819<br>AN0826<br>AN0840<br>AN0847<br>AN0912<br>AN0922<br>AN0926<br>AN0999<br>AN1024<br>AN1032<br>AN1115<br>AN1117<br>AN1120<br>AN1142<br>AN1152<br>AN1163<br>AN1167<br>AN1213<br>AN1269<br>AN1302<br>AN1314<br>AN1377<br>AN1380<br>AN1430<br>AN1461<br>AN1462<br>AN1475<br>AN1480<br>AN1491<br>AN1506<br>AN1540<br>AN1548<br>AN1554<br>AN1582<br>AN1587<br>AN1599<br>AN1604<br>AN1655<br>AN1657<br>AN1670<br>AN1673<br>AN1703<br>AN1723<br>AN1744<br>AN1857<br>AN1867<br>AN1896<br>AN2012<br>AN2014<br>AN2021<br>AN2022<br>AN2047<br>AN2055<br>AN2062<br>AN2112<br>AN2121<br>AN2159<br>AN2166<br>AN2167<br>AN2174 |

| | | | | | |
|---|---|---|---|---|---|
| | | | | AN8147<br>AN8174<br>AN8189<br>AN8273<br>AN8329<br>AN8354<br>AN8420<br>AN8526<br>AN8536<br>AN8592<br>AN8594<br>AN8610<br>AN8677<br>AN8713<br>AN8785<br>AN8807<br>AN8826<br>AN8935<br>AN8969<br>AN8995<br>AN8998<br>AN9064<br>AN9180<br>AN9218<br>AN9270<br>AN9374<br>AN9437<br>AN9514<br>AN0059<br>AN0084<br>AN0162<br>AN0245<br>AN0289<br>AN0376<br>AN0403<br>AN0557<br>AN0752<br>AN0755<br>AN0795<br>AN0801<br>AN0849<br>AN0887<br>AN0888<br>AN0925<br>AN0979<br>AN1173<br>AN1184<br>AN1260<br>AN1422<br>AN1492<br>AN1729<br>AN1746<br>AN1766<br>AN1854<br>AN1917<br>AN1944<br>AN2025<br>AN2028<br>AN2115<br>AN2120<br>AN2158<br>AN2179<br>AN2230<br>AN2239<br>AN2360<br>AN2424 | AN2186<br>AN2239<br>AN2392<br>AN2409<br>AN2412<br>AN2419<br>AN2425<br>AN2466<br>AN2513<br>AN2523<br>AN2536<br>AN2588<br>AN2608<br>AN2622<br>AN2623<br>AN2660<br>AN2719<br>AN2910<br>AN2921<br>AN2959<br>AN2960<br>AN2988<br>AN3026<br>AN3079<br>AN3089<br>AN3098<br>AN3126<br>AN3143<br>AN3175<br>AN3188<br>AN3194<br>AN3203<br>AN3244<br>AN3279<br>AN3286<br>AN3309<br>AN3403<br>AN3414<br>AN3452<br>AN3577<br>AN3592<br>AN3599<br>AN3636<br>AN3664<br>AN3691<br>AN3714<br>AN3764<br>AN3783<br>AN3849<br>AN3876<br>AN3917<br>AN4023<br>AN4039<br>AN4085<br>AN4106<br>AN4108<br>AN4115<br>AN4129<br>AN4146<br>AN4147<br>AN4183<br>AN4195<br>AN4215<br>AN4301<br>AN4418<br>AN4455 |

| | | | | | |
|---|---|---|---|---|---|
| | | | | AN2486<br>AN2512<br>AN2527<br>AN2574<br>AN2583<br>AN2584<br>AN2714<br>AN2887<br>AN2894<br>AN3088<br>AN3123<br>AN3125<br>AN3243<br>AN3275<br>AN3285<br>AN3327<br>AN3383<br>AN3392<br>AN3517<br>AN3559<br>AN3622<br>AN3642<br>AN3674<br>AN3702<br>AN3862<br>AN3902<br>AN3924<br>AN4038<br>AN4050<br>AN4084<br>AN4177<br>AN4322<br>AN4396<br>AN4456<br>AN4626<br>AN4633<br>AN4640<br>AN4741<br>AN4750<br>AN4888<br>AN5103<br>AN5145<br>AN5199<br>AN5235<br>AN5414<br>AN5479<br>AN5556<br>AN5629<br>AN5656<br>AN5709<br>AN5721<br>AN5744<br>AN5790<br>AN5923<br>AN6023<br>AN6032<br>AN6068<br>AN6080<br>AN6115<br>AN6329<br>AN6351<br>AN6658<br>AN6696<br>AN6933<br>AN7007<br>AN7041 | AN4473<br>AN4481<br>AN4514<br>AN4572<br>AN4579<br>AN4625<br>AN4627<br>AN4638<br>AN4742<br>AN4761<br>AN4806<br>AN4839<br>AN4867<br>AN4902<br>AN4919<br>AN4925<br>AN4958<br>AN4979<br>AN4987<br>AN4997<br>AN5053<br>AN5139<br>AN5162<br>AN5169<br>AN5221<br>AN5318<br>AN5329<br>AN5368<br>AN5427<br>AN5481<br>AN5534<br>AN5555<br>AN5604<br>AN5616<br>AN5644<br>AN5663<br>AN5677<br>AN5686<br>AN5695<br>AN5718<br>AN5722<br>AN5746<br>AN5764<br>AN5794<br>AN5807<br>AN5812<br>AN5814<br>AN5904<br>AN5973<br>AN5981<br>AN6000<br>AN6033<br>AN6046<br>AN6172<br>AN6179<br>AN6195<br>AN6210<br>AN6222<br>AN6278<br>AN6341<br>AN6352<br>AN6360<br>AN6366<br>AN6389<br>AN6495<br>AN6535 |

| | | | | | |
|---|---|---|---|---|---|
| | | | | AN7091<br>AN7159<br>AN7169<br>AN7176<br>AN7213<br>AN7227<br>AN7291<br>AN7298<br>AN7377<br>AN7474<br>AN7588<br>AN7653<br>AN7797<br>AN7837<br>AN7838<br>AN7869<br>AN7971<br>AN8076<br>AN8101<br>AN8102<br>AN8129<br>AN8201<br>AN8256<br>AN8338<br>AN8392<br>AN8399<br>AN8400<br>AN8411<br>AN8450<br>AN8638<br>AN8639<br>AN8706<br>AN8755<br>AN8864<br>AN8870<br>AN8902<br>AN8928<br>AN9122<br>AN9183<br>AN9204<br>AN9328<br>AN9469<br>AN0009<br>AN0131<br>AN0161<br>AN0225<br>AN0241<br>AN0298<br>AN0573<br>AN0599<br>AN0600<br>AN0615<br>AN0866<br>AN0934<br>AN1039<br>AN1119<br>AN1195<br>AN1310<br>AN1326<br>AN1359<br>AN1364<br>AN1367<br>AN1417<br>AN1476<br>AN1482<br>AN1586 | AN6588<br>AN6652<br>AN6833<br>AN6874<br>AN6888<br>AN6898<br>AN6900<br>AN6916<br>AN6934<br>AN6963<br>AN7020<br>AN7058<br>AN7082<br>AN7114<br>AN7137<br>AN7138<br>AN7139<br>AN7149<br>AN7186<br>AN7188<br>AN7229<br>AN7243<br>AN7303<br>AN7331<br>AN7341<br>AN7353<br>AN7367<br>AN7386<br>AN7434<br>AN7451<br>AN7466<br>AN7479<br>AN7500<br>AN7587<br>AN7591<br>AN7592<br>AN7635<br>AN7654<br>AN7664<br>AN7780<br>AN7791<br>AN7794<br>AN7859<br>AN7883<br>AN7945<br>AN7949<br>AN8001<br>AN8005<br>AN8007<br>AN8018<br>AN8041<br>AN8057<br>AN8072<br>AN8080<br>AN8084<br>AN8210<br>AN8246<br>AN8252<br>AN8285<br>AN8324<br>AN8356<br>AN8532<br>AN8556<br>AN8560<br>AN8566<br>AN8568 |

| | | | | | |
|---|---|---|---|---|---|
| | | | | AN1610<br>AN1782<br>AN1850<br>AN1910<br>AN1962<br>AN2043<br>AN2165<br>AN2196<br>AN2242<br>AN2266<br>AN2345<br>AN2478<br>AN2639<br>AN2691<br>AN2737<br>AN2755<br>AN2923<br>AN3420<br>AN3439<br>AN3465<br>AN3554<br>AN3569<br>AN3583<br>AN3595<br>AN3670<br>AN3684<br>AN3795<br>AN3871<br>AN3935<br>AN4098<br>AN4112<br>AN4135<br>AN4172<br>AN4173<br>AN4214<br>AN4279<br>AN4372<br>AN4394<br>AN4424<br>AN4473<br>AN4622<br>AN4687<br>AN4811<br>AN4838<br>AN4853<br>AN4918<br>AN5003<br>AN5082<br>AN5096<br>AN5124<br>AN5154<br>AN5166<br>AN5174<br>AN5275<br>AN5352<br>AN5459<br>AN5477<br>AN5511<br>AN5636<br>AN5795<br>AN5812<br>AN5925<br>AN6044<br>AN6408<br>AN6502<br>AN6692 | AN8587<br>AN8590<br>AN8595<br>AN8659<br>AN8718<br>AN8776<br>AN8786<br>AN8808<br>AN8819<br>AN8893<br>AN8945<br>AN8958<br>AN9106<br>AN9143<br>AN9157<br>AN9174<br>AN9175<br>AN9210<br>AN9271<br>AN9292<br>AN9298<br>AN9301<br>AN9314<br>AN9332<br>AN9371<br>AN9393<br>AN9434<br>AN9460<br>AN9497<br>AN0031<br>AN0048<br>AN0052<br>AN0071<br>AN0129<br>AN0140<br>AN0148<br>AN0157<br>AN0171<br>AN0189<br>AN0207<br>AN0226<br>AN0230<br>AN0279<br>AN0296<br>AN0313<br>AN0315<br>AN0353<br>AN0355<br>AN0417<br>AN0442<br>AN0464<br>AN0473<br>AN0487<br>AN0500<br>AN0516<br>AN0606<br>AN0626<br>AN0631<br>AN0711<br>AN0713<br>AN0722<br>AN0731<br>AN0740<br>AN0764<br>AN0768<br>AN0799 |

| | | | | | |
|---|---|---|---|---|---|
| | | | | AN6760<br>AN6889<br>AN6930<br>AN6977<br>AN7110<br>AN7182<br>AN7231<br>AN7263<br>AN7452<br>AN7532<br>AN7548<br>AN7731<br>AN7773<br>AN7897<br>AN8059<br>AN8133<br>AN8250<br>AN8268<br>AN8331<br>AN8421<br>AN8566<br>AN8589<br>AN8860<br>AN8904<br>AN8905<br>AN9178<br>AN9333<br>AN0044<br>AN0449<br>AN0550<br>AN0699<br>AN0818<br>AN0829<br>AN0863<br>AN0932<br>AN0999<br>AN1202<br>AN1223<br>AN1264<br>AN1534<br>AN1546<br>AN1550<br>AN1802<br>AN1922<br>AN2148<br>AN2198<br>AN2279<br>AN2294<br>AN2417<br>AN2521<br>AN2534<br>AN2546<br>AN2621<br>AN2681<br>AN2694<br>AN2845<br>AN2884<br>AN2917<br>AN2920<br>AN2967<br>AN3133<br>AN3492<br>AN3518<br>AN3530<br>AN3542<br>AN3592 | AN0846<br>AN0858<br>AN0867<br>AN0876<br>AN0938<br>AN0940<br>AN0942<br>AN0951<br>AN0974<br>AN0980<br>AN0989<br>AN0995<br>AN1002<br>AN1008<br>AN1015<br>AN1021<br>AN1034<br>AN1035<br>AN1055<br>AN1067<br>AN1107<br>AN1108<br>AN1145<br>AN1189<br>AN1296<br>AN1303<br>AN1311<br>AN1360<br>AN1454<br>AN1539<br>AN1560<br>AN1563<br>AN1594<br>AN1611<br>AN1613<br>AN1615<br>AN1625<br>AN1646<br>AN1688<br>AN1747<br>AN1803<br>AN1809<br>AN1813<br>AN1873<br>AN1884<br>AN1893<br>AN1931<br>AN1967<br>AN2031<br>AN2037<br>AN2039<br>AN2114<br>AN2142<br>AN2150<br>AN2162<br>AN2180<br>AN2196<br>AN2304<br>AN2319<br>AN2325<br>AN2328<br>AN2358<br>AN2360<br>AN2361<br>AN2388<br>AN2439 |

| | | | | | |
|---|---|---|---|---|---|
| | | | | AN3757<br>AN3838<br>AN3839<br>AN3848<br>AN3916<br>AN3919<br>AN3933<br>AN3974<br>AN4158<br>AN4178<br>AN4244<br>AN4404<br>AN4443<br>AN4550<br>AN4561<br>AN4600<br>AN4723<br>AN4788<br>AN4806<br>AN5011<br>AN5098<br>AN5117<br>AN5144<br>AN5197<br>AN5225<br>AN5238<br>AN5354<br>AN5430<br>AN5467<br>AN5689<br>AN5830<br>AN6004<br>AN6293<br>AN6326<br>AN6564<br>AN6618<br>AN6635<br>AN6649<br>AN6795<br>AN7036<br>AN7126<br>AN7184<br>AN7187<br>AN7328<br>AN7336<br>AN7388<br>AN7465<br>AN7506<br>AN7516<br>AN7593<br>AN7712<br>AN7758<br>AN7814<br>AN8066<br>AN8083<br>AN8168<br>AN8170<br>AN8543<br>AN8544<br>AN8552<br>AN8591<br>AN8680<br>AN8708<br>AN8818<br>AN8931<br>AN9000 | AN2445<br>AN2518<br>AN2519<br>AN2528<br>AN2538<br>AN2557<br>AN2562<br>AN2583<br>AN2585<br>AN2598<br>AN2656<br>AN2724<br>AN2733<br>AN2738<br>AN2778<br>AN2779<br>AN2856<br>AN2862<br>AN2894<br>AN2896<br>AN2901<br>AN2913<br>AN3020<br>AN3021<br>AN3083<br>AN3130<br>AN3136<br>AN3159<br>AN3178<br>AN3181<br>AN3207<br>AN3222<br>AN3239<br>AN3271<br>AN3298<br>AN3343<br>AN3357<br>AN3360<br>AN3378<br>AN3395<br>AN3416<br>AN3422<br>AN3464<br>AN3499<br>AN3509<br>AN3530<br>AN3558<br>AN3560<br>AN3598<br>AN3604<br>AN3606<br>AN3624<br>AN3626<br>AN3627<br>AN3676<br>AN3703<br>AN3725<br>AN3759<br>AN3773<br>AN3777<br>AN3829<br>AN3838<br>AN3851<br>AN3872<br>AN3882<br>AN3890 |

| | | | | | |
|---|---|---|---|---|---|
| | | | | AN9004<br>AN9127<br>AN9128<br>AN9147<br>AN9302<br>AN9400<br>AN9488<br>AN0010<br>AN0021<br>AN0052<br>AN0232<br>AN0325<br>AN0401<br>AN0452<br>AN0678<br>AN0788<br>AN0834<br>AN1108<br>AN1116<br>AN1415<br>AN1541<br>AN1857<br>AN2024<br>AN2035<br>AN2207<br>AN2373<br>AN2429<br>AN2457<br>AN2480<br>AN2547<br>AN2578<br>AN2593<br>AN2672<br>AN2696<br>AN2804<br>AN2982<br>AN2984<br>AN3025<br>AN3106<br>AN3458<br>AN3591<br>AN3708<br>AN3715<br>AN3729<br>AN3923<br>AN4010<br>AN4016<br>AN4085<br>AN4164<br>AN4261<br>AN4419<br>AN4433<br>AN4502<br>AN4807<br>AN4809<br>AN4846<br>AN4862<br>AN4940<br>AN5356<br>AN5367<br>AN5463<br>AN5599<br>AN5748<br>AN5792<br>AN5907<br>AN6034 | AN3914<br>AN3920<br>AN3943<br>AN3974<br>AN4007<br>AN4021<br>AN4031<br>AN4057<br>AN4078<br>AN4086<br>AN4107<br>AN4109<br>AN4110<br>AN4121<br>AN4130<br>AN4151<br>AN4161<br>AN4189<br>AN4193<br>AN4214<br>AN4234<br>AN4241<br>AN4258<br>AN4262<br>AN4312<br>AN4373<br>AN4470<br>AN4482<br>AN4530<br>AN4532<br>AN4539<br>AN4550<br>AN4561<br>AN4597<br>AN4653<br>AN4655<br>AN4720<br>AN4776<br>AN4801<br>AN4825<br>AN4863<br>AN4885<br>AN4905<br>AN4911<br>AN4914<br>AN4986<br>AN5004<br>AN5021<br>AN5028<br>AN5050<br>AN5057<br>AN5058<br>AN5077<br>AN5079<br>AN5090<br>AN5109<br>AN5125<br>AN5143<br>AN5171<br>AN5183<br>AN5184<br>AN5210<br>AN5286<br>AN5287<br>AN5301<br>AN5327 |

| | | | | | |
|---|---|---|---|---|---|
| | | | | AN6036<br>AN6158<br>AN6234<br>AN6835<br>AN7108<br>AN7109<br>AN7141<br>AN7250<br>AN7276<br>AN7315<br>AN7453<br>AN7634<br>AN7768<br>AN7825<br>AN7847<br>AN8060<br>AN8156<br>AN8219<br>AN8325<br>AN8350<br>AN8390<br>AN8597<br>AN8681<br>AN9288<br>AN0042<br>AN0079<br>AN0189<br>AN0266<br>AN0380<br>AN0672<br>AN0765<br>AN0835<br>AN0869<br>AN0990<br>AN1070<br>AN1330<br>AN1468<br>AN1547<br>AN1609<br>AN1915<br>AN1929<br>AN1992<br>AN2092<br>AN2535<br>AN2544<br>AN2634<br>AN2889<br>AN2956<br>AN3097<br>AN3288<br>AN3378<br>AN3394<br>AN3407<br>AN3425<br>AN3636<br>AN3671<br>AN3870<br>AN3872<br>AN3940<br>AN3970<br>AN4107<br>AN4146<br>AN4180<br>AN4182<br>AN4188<br>AN4264 | AN5457<br>AN5458<br>AN5468<br>AN5512<br>AN5528<br>AN5546<br>AN5575<br>AN5613<br>AN5624<br>AN5626<br>AN5634<br>AN5664<br>AN5744<br>AN5745<br>AN5752<br>AN5753<br>AN5757<br>AN5773<br>AN5781<br>AN5947<br>AN6007<br>AN6014<br>AN6048<br>AN6093<br>AN6141<br>AN6169<br>AN6196<br>AN6203<br>AN6204<br>AN6221<br>AN6230<br>AN6243<br>AN6307<br>AN6381<br>AN6385<br>AN6386<br>AN6398<br>AN6413<br>AN6503<br>AN6508<br>AN6536<br>AN6578<br>AN6589<br>AN6620<br>AN6624<br>AN6638<br>AN6645<br>AN6658<br>AN6661<br>AN6750<br>AN6770<br>AN6779<br>AN6807<br>AN6819<br>AN6831<br>AN6836<br>AN6915<br>AN6933<br>AN6952<br>AN6960<br>AN7057<br>AN7109<br>AN7135<br>AN7167<br>AN7177<br>AN7187 |

| | | | | | |
|---|---|---|---|---|---|
| | | | | AN4429<br>AN4712<br>AN4760<br>AN4830<br>AN4889<br>AN4913<br>AN5422<br>AN5503<br>AN5536<br>AN5587<br>AN5598<br>AN5623<br>AN5739<br>AN5775<br>AN5779<br>AN5823<br>AN6166<br>AN6209<br>AN6238<br>AN6497<br>AN6607<br>AN6640<br>AN6749<br>AN6845<br>AN6941<br>AN7168<br>AN7228<br>AN7279<br>AN7285<br>AN7390<br>AN7504<br>AN7513<br>AN7517<br>AN7522<br>AN7528<br>AN7874<br>AN7881<br>AN7916<br>AN7938<br>AN8143<br>AN8157<br>AN8253<br>AN8299<br>AN8343<br>AN8658<br>AN8668<br>AN8683<br>AN8688<br>AN8752<br>AN8937<br>AN9022<br>AN9031<br>AN9159<br>AN9171<br>AN9262<br>AN9295<br>AN0072<br>AN0077<br>AN0140<br>AN0224<br>AN0476<br>AN0544<br>AN0727<br>AN0739<br>AN0766<br>AN0784 | AN7207<br>AN7214<br>AN7217<br>AN7250<br>AN7292<br>AN7302<br>AN7326<br>AN7352<br>AN7406<br>AN7436<br>AN7462<br>AN7484<br>AN7502<br>AN7511<br>AN7538<br>AN7541<br>AN7575<br>AN7589<br>AN7595<br>AN7621<br>AN7622<br>AN7702<br>AN7703<br>AN7712<br>AN7784<br>AN7864<br>AN7939<br>AN7941<br>AN7976<br>AN7989<br>AN7990<br>AN8008<br>AN8023<br>AN8077<br>AN8082<br>AN8116<br>AN8122<br>AN8136<br>AN8146<br>AN8155<br>AN8166<br>AN8184<br>AN8208<br>AN8218<br>AN8223<br>AN8228<br>AN8268<br>AN8329<br>AN8330<br>AN8339<br>AN8349<br>AN8355<br>AN8386<br>AN8395<br>AN8440<br>AN8445<br>AN8463<br>AN8554<br>AN8591<br>AN8593<br>AN8604<br>AN8616<br>AN8634<br>AN8637<br>AN8638<br>AN8639 |

| | | | | | |
|---|---|---|---|---|---|
| | | | | AN1166<br>AN1358<br>AN1795<br>AN1821<br>AN1851<br>AN2044<br>AN2058<br>AN2139<br>AN2194<br>AN2211<br>AN2393<br>AN2751<br>AN2903<br>AN2976<br>AN3043<br>AN3111<br>AN3304<br>AN3349<br>AN3376<br>AN3387<br>AN3446<br>AN3584<br>AN4007<br>AN4167<br>AN4312<br>AN4558<br>AN4698<br>AN4789<br>AN4842<br>AN4960<br>AN5220<br>AN5457<br>AN5829<br>AN5941<br>AN6404<br>AN6707<br>AN6731<br>AN6752<br>AN7154<br>AN7404<br>AN7470<br>AN7575<br>AN7622<br>AN7936<br>AN8035<br>AN8056<br>AN8131<br>AN8499<br>AN8500<br>AN8792<br>AN8832<br>AN9181<br>AN9355<br>AN9383<br>AN0033<br>AN0108<br>AN0231<br>AN0332<br>AN0443<br>AN0719<br>AN0878<br>AN1193<br>AN1487<br>AN1504<br>AN1522<br>AN1583 | AN8683<br>AN8710<br>AN8726<br>AN8730<br>AN8740<br>AN8782<br>AN8787<br>AN8820<br>AN8859<br>AN8861<br>AN8888<br>AN8900<br>AN8903<br>AN8905<br>AN8915<br>AN8924<br>AN8934<br>AN8959<br>AN8996<br>AN8997<br>AN9002<br>AN9004<br>AN9076<br>AN9123<br>AN9156<br>AN9173<br>AN9180<br>AN9190<br>AN9191<br>AN9277<br>AN9303<br>AN9337<br>AN9376<br>AN9392<br>AN9499<br>AN0057<br>AN0077<br>AN0134<br>AN0152<br>AN0161<br>AN0168<br>AN0169<br>AN0186<br>AN0216<br>AN0289<br>AN0290<br>AN0324<br>AN0369<br>AN0437<br>AN0450<br>AN0472<br>AN0477<br>AN0493<br>AN0512<br>AN0563<br>AN0718<br>AN0788<br>AN0806<br>AN0883<br>AN0884<br>AN0889<br>AN0895<br>AN0901<br>AN0934<br>AN0939<br>AN0957 |

| | | | | | |
|---|---|---|---|---|---|
| | | | | AN1620<br>AN1743<br>AN1760<br>AN1816<br>AN1934<br>AN1968<br>AN1978<br>AN2166<br>AN2206<br>AN2331<br>AN2454<br>AN2893<br>AN3325<br>AN3355<br>AN3424<br>AN3430<br>AN3775<br>AN3880<br>AN4051<br>AN4133<br>AN4232<br>AN4240<br>AN4249<br>AN4268<br>AN4708<br>AN4770<br>AN4783<br>AN4868<br>AN4925<br>AN5067<br>AN5446<br>AN5791<br>AN5843<br>AN5870<br>AN5952<br>AN5972<br>AN6122<br>AN6221<br>AN6272<br>AN6339<br>AN6340<br>AN6535<br>AN6664<br>AN6734<br>AN6887<br>AN7357<br>AN7408<br>AN7442<br>AN7538<br>AN7574<br>AN7687<br>AN7696<br>AN7706<br>AN7780<br>AN7865<br>AN7937<br>AN7965<br>AN8096<br>AN8119<br>AN8428<br>AN8561<br>AN8738<br>AN8810<br>AN8851<br>AN8880<br>AN8987 | AN0992<br>AN1001<br>AN1003<br>AN1006<br>AN1023<br>AN1033<br>AN1050<br>AN1052<br>AN1070<br>AN1103<br>AN1174<br>AN1181<br>AN1201<br>AN1218<br>AN1282<br>AN1287<br>AN1289<br>AN1291<br>AN1333<br>AN1509<br>AN1570<br>AN1598<br>AN1610<br>AN1616<br>AN1686<br>AN1700<br>AN1702<br>AN1732<br>AN1800<br>AN1810<br>AN1839<br>AN1841<br>AN1843<br>AN1845<br>AN1848<br>AN1858<br>AN1897<br>AN1899<br>AN1912<br>AN1945<br>AN1995<br>AN2023<br>AN2035<br>AN2140<br>AN2151<br>AN2177<br>AN2179<br>AN2199<br>AN2200<br>AN2209<br>AN2222<br>AN2233<br>AN2323<br>AN2334<br>AN2359<br>AN2375<br>AN2395<br>AN2396<br>AN2454<br>AN2464<br>AN2530<br>AN2545<br>AN2582<br>AN2591<br>AN2597<br>AN2675 |

| | | | | | |
|---|---|---|---|---|---|
| | | | | AN9002<br>AN9307<br>AN9393<br>AN9449<br>AN9517<br>AN0247<br>AN0323<br>AN0354<br>AN0867<br>AN1113<br>AN1155<br>AN1266<br>AN1278<br>AN1288<br>AN1314<br>AN1402<br>AN1453<br>AN1589<br>AN1699<br>AN1718<br>AN2185<br>AN2208<br>AN2818<br>AN2855<br>AN3081<br>AN3082<br>AN3209<br>AN3298<br>AN3428<br>AN4269<br>AN4291<br>AN4300<br>AN4489<br>AN4573<br>AN4652<br>AN4667<br>AN5035<br>AN5114<br>AN5341<br>AN5372<br>AN5514<br>AN5708<br>AN5878<br>AN5917<br>AN6123<br>AN6174<br>AN6185<br>AN6399<br>AN6534<br>AN6736<br>AN6855<br>AN6893<br>AN6988<br>AN7142<br>AN7149<br>AN7311<br>AN7352<br>AN7700<br>AN7875<br>AN8018<br>AN8067<br>AN8181<br>AN8239<br>AN8342<br>AN8406<br>AN8460 | AN2731<br>AN2800<br>AN2823<br>AN2827<br>AN2853<br>AN2880<br>AN2970<br>AN2978<br>AN2987<br>AN3141<br>AN3152<br>AN3153<br>AN3155<br>AN3218<br>AN3221<br>AN3280<br>AN3293<br>AN3304<br>AN3320<br>AN3346<br>AN3348<br>AN3388<br>AN3429<br>AN3442<br>AN3524<br>AN3528<br>AN3541<br>AN3543<br>AN3566<br>AN3590<br>AN3612<br>AN3672<br>AN3686<br>AN3687<br>AN3709<br>AN3716<br>AN3769<br>AN3776<br>AN3877<br>AN3927<br>AN3976<br>AN3997<br>AN4011<br>AN4061<br>AN4069<br>AN4075<br>AN4081<br>AN4082<br>AN4160<br>AN4217<br>AN4236<br>AN4255<br>AN4280<br>AN4292<br>AN4310<br>AN4323<br>AN4402<br>AN4420<br>AN4480<br>AN4503<br>AN4521<br>AN4524<br>AN4535<br>AN4551<br>AN4589<br>AN4608 |

| | | | | | |
|---|---|---|---|---|---|
| | | | | AN8487<br>AN8554<br>AN8556<br>AN8617<br>AN8793<br>AN9068<br>AN9123<br>AN9130<br>AN9154<br>AN9161<br>AN9166<br>AN9324<br>AN9345<br>AN9376<br>AN9489<br>AN0065<br>AN0070<br>AN0281<br>AN0314<br>AN0442<br>AN0687<br>AN0873<br>AN1001<br>AN1078<br>AN1141<br>AN1151<br>AN1168<br>AN1590<br>AN1593<br>AN1798<br>AN2054<br>AN2323<br>AN2343<br>AN2675<br>AN2718<br>AN2957<br>AN2998<br>AN3151<br>AN3469<br>AN3604<br>AN4191<br>AN4205<br>AN4216<br>AN4225<br>AN4566<br>AN4864<br>AN5016<br>AN5163<br>AN5169<br>AN5403<br>AN5407<br>AN5554<br>AN5592<br>AN5679<br>AN5801<br>AN6077<br>AN6844<br>AN6912<br>AN7085<br>AN7566<br>AN7608<br>AN7698<br>AN7834<br>AN7872<br>AN7986<br>AN8061 | AN4626<br>AN4635<br>AN4641<br>AN4702<br>AN4758<br>AN4783<br>AN4795<br>AN4799<br>AN4800<br>AN4807<br>AN4810<br>AN4852<br>AN4869<br>AN4890<br>AN4918<br>AN4951<br>AN4961<br>AN4990<br>AN5033<br>AN5067<br>AN5099<br>AN5115<br>AN5116<br>AN5137<br>AN5167<br>AN5172<br>AN5239<br>AN5322<br>AN5346<br>AN5372<br>AN5379<br>AN5410<br>AN5416<br>AN5422<br>AN5447<br>AN5451<br>AN5494<br>AN5521<br>AN5524<br>AN5525<br>AN5550<br>AN5557<br>AN5600<br>AN5606<br>AN5645<br>AN5649<br>AN5675<br>AN5700<br>AN5705<br>AN5796<br>AN5848<br>AN5866<br>AN5886<br>AN5968<br>AN6031<br>AN6041<br>AN6075<br>AN6232<br>AN6314<br>AN6328<br>AN6351<br>AN6505<br>AN6518<br>AN6597<br>AN6608<br>AN6610 |

| | | | | | |
|---|---|---|---|---|---|
| | | | | AN8097<br>AN8191<br>AN8240<br>AN8388<br>AN8659<br>AN8711<br>AN8939<br>AN9074<br>AN9108<br>AN9206<br>AN9296<br>AN0066<br>AN0381<br>AN0495<br>AN0508<br>AN0605<br>AN0657<br>AN0825<br>AN0941<br>AN1049<br>AN1074<br>AN1092<br>AN1159<br>AN1182<br>AN1191<br>AN1585<br>AN1695<br>AN1909<br>AN1925<br>AN2055<br>AN2173<br>AN2215<br>AN2413<br>AN2638<br>AN2734<br>AN2879<br>AN3457<br>AN3743<br>AN3850<br>AN3853<br>AN3936<br>AN3982<br>AN4162<br>AN4314<br>AN4691<br>AN4740<br>AN4943<br>AN5652<br>AN5777<br>AN5824<br>AN5856<br>AN5859<br>AN5871<br>AN5909<br>AN6192<br>AN6236<br>AN6370<br>AN6496<br>AN7090<br>AN7283<br>AN7310<br>AN7461<br>AN7533<br>AN7666<br>AN7673<br>AN7782 | AN6628<br>AN6640<br>AN6646<br>AN6647<br>AN6667<br>AN6697<br>AN6721<br>AN6772<br>AN6813<br>AN6866<br>AN6899<br>AN6969<br>AN7010<br>AN7016<br>AN7021<br>AN7077<br>AN7081<br>AN7113<br>AN7129<br>AN7227<br>AN7311<br>AN7391<br>AN7443<br>AN7515<br>AN7540<br>AN7560<br>AN7617<br>AN7689<br>AN7704<br>AN7713<br>AN7729<br>AN7741<br>AN7793<br>AN7815<br>AN7833<br>AN7874<br>AN7877<br>AN7878<br>AN7880<br>AN7904<br>AN7908<br>AN7937<br>AN7952<br>AN7971<br>AN7972<br>AN7993<br>AN7995<br>AN8003<br>AN8013<br>AN8040<br>AN8045<br>AN8095<br>AN8129<br>AN8130<br>AN8157<br>AN8169<br>AN8220<br>AN8221<br>AN8243<br>AN8400<br>AN8416<br>AN8425<br>AN8428<br>AN8443<br>AN8446<br>AN8462 |

| | | | | | |
|---|---|---|---|---|---|
| | | | | AN7945 | AN8478 |
| | | | | AN8081 | AN8489 |
| | | | | AN8539 | AN8523 |
| | | | | AN8999 | AN8537 |
| | | | | AN9155 | AN8567 |
| | | | | AN9165 | AN8573 |
| | | | | AN9220 | AN8592 |
| | | | | AN9370 | AN8598 |
| | | | | AN9458 | AN8615 |
| | | | | AN0067 | AN8741 |
| | | | | AN0286 | AN8777 |
| | | | | AN0315 | AN8818 |
| | | | | AN0900 | AN8823 |
| | | | | AN0993 | AN8833 |
| | | | | AN1115 | AN8884 |
| | | | | AN1248 | AN8963 |
| | | | | AN1303 | AN8964 |
| | | | | AN1461 | AN8970 |
| | | | | AN1514 | AN9005 |
| | | | | AN1853 | AN9034 |
| | | | | AN1999 | AN9051 |
| | | | | AN2288 | AN9067 |
| | | | | AN2318 | AN9068 |
| | | | | AN2594 | AN9075 |
| | | | | AN2939 | AN9127 |
| | | | | AN3018 | AN9129 |
| | | | | AN3054 | AN9148 |
| | | | | AN3167 | AN9179 |
| | | | | AN3188 | AN9207 |
| | | | | AN3217 | AN9299 |
| | | | | AN3257 | AN9302 |
| | | | | AN3761 | AN9324 |
| | | | | AN4139 | AN9351 |
| | | | | AN4378 | AN9370 |
| | | | | AN4616 | AN9461 |
| | | | | AN4689 | AN9481 |
| | | | | AN4722 | AN0010 |
| | | | | AN4749 | AN0016 |
| | | | | AN4986 | AN0023 |
| | | | | AN5440 | AN0030 |
| | | | | AN5669 | AN0046 |
| | | | | AN5670 | AN0054 |
| | | | | AN5722 | AN0058 |
| | | | | AN5743 | AN0059 |
| | | | | AN5957 | AN0060 |
| | | | | AN6079 | AN0068 |
| | | | | AN6255 | AN0070 |
| | | | | AN6864 | AN0098 |
| | | | | AN6927 | AN0105 |
| | | | | AN7040 | AN0109 |
| | | | | AN7134 | AN0170 |
| | | | | AN7330 | AN0172 |
| | | | | AN7361 | AN0175 |
| | | | | AN7425 | AN0181 |
| | | | | AN7531 | AN0188 |
| | | | | AN7623 | AN0201 |
| | | | | AN7670 | AN0221 |
| | | | | AN7717 | AN0228 |
| | | | | AN7890 | AN0233 |
| | | | | AN7975 | AN0246 |
| | | | | AN8715 | AN0253 |
| | | | | AN8858 | AN0257 |
| | | | | AN8873 | AN0266 |
| | | | | AN9399 | AN0267 |
| | | | | AN0133 | AN0276 |
| | | | | AN0210 | AN0288 |

| | | | | | |
|---|---|---|---|---|---|
| | | | | AN0694<br>AN0978<br>AN1216<br>AN1636<br>AN1818<br>AN2590<br>AN3646<br>AN3797<br>AN4271<br>AN4491<br>AN4684<br>AN4772<br>AN4884<br>AN5171<br>AN5300<br>AN5413<br>AN5888<br>AN5914<br>AN5934<br>AN5953<br>AN6090<br>AN6539<br>AN6645<br>AN6675<br>AN6740<br>AN7015<br>AN7101<br>AN7128<br>AN7325<br>AN7563<br>AN7729<br>AN8981<br>AN9360<br>AN9363<br>AN9491<br>AN0212<br>AN0305<br>AN0608<br>AN0785<br>AN0956<br>AN0984<br>AN1058<br>AN1277<br>AN1431<br>AN1447<br>AN1553<br>AN1952<br>AN2002<br>AN2065<br>AN2221<br>AN2438<br>AN2586<br>AN2647<br>AN2752<br>AN2843<br>AN2846<br>AN3137<br>AN3441<br>AN3566<br>AN3582<br>AN3908<br>AN4195<br>AN4285<br>AN4386<br>AN4438<br>AN4563 | AN0299<br>AN0306<br>AN0327<br>AN0328<br>AN0333<br>AN0358<br>AN0361<br>AN0377<br>AN0379<br>AN0381<br>AN0400<br>AN0405<br>AN0407<br>AN0408<br>AN0413<br>AN0415<br>AN0419<br>AN0438<br>AN0466<br>AN0484<br>AN0491<br>AN0494<br>AN0496<br>AN0503<br>AN0509<br>AN0528<br>AN0531<br>AN0551<br>AN0558<br>AN0601<br>AN0607<br>AN0633<br>AN0638<br>AN0640<br>AN0641<br>AN0646<br>AN0649<br>AN0650<br>AN0677<br>AN0714<br>AN0720<br>AN0751<br>AN0753<br>AN0778<br>AN0779<br>AN0782<br>AN0783<br>AN0784<br>AN0785<br>AN0790<br>AN0798<br>AN0801<br>AN0802<br>AN0811<br>AN0814<br>AN0820<br>AN0835<br>AN0845<br>AN0849<br>AN0864<br>AN0878<br>AN0899<br>AN0935<br>AN0959<br>AN0967<br>AN0968 |

| | | | | | |
|---|---|---|---|---|---|
| | | | | AN4686<br>AN4745<br>AN5527<br>AN5562<br>AN5600<br>AN5621<br>AN5796<br>AN5847<br>AN6229<br>AN6929<br>AN6978<br>AN7136<br>AN7214<br>AN7518<br>AN7932<br>AN8039<br>AN8545<br>AN8555<br>AN8757<br>AN9099<br>AN9160<br>AN9298<br>AN9461<br>AN0057<br>AN0182<br>AN0254<br>AN0271<br>AN0665<br>AN0800<br>AN1052<br>AN1066<br>AN1190<br>AN1206<br>AN1242<br>AN2522<br>AN3388<br>AN3809<br>AN3876<br>AN3903<br>AN4041<br>AN4220<br>AN4466<br>AN4934<br>AN5317<br>AN5686<br>AN6066<br>AN6111<br>AN6390<br>AN6768<br>AN6873<br>AN7304<br>AN7434<br>AN7445<br>AN7606<br>AN7614<br>AN7749<br>AN7761<br>AN8020<br>AN8175<br>AN8459<br>AN8990<br>AN9377<br>AN0192<br>AN0760<br>AN0790<br>AN0895 | AN0985<br>AN1017<br>AN1028<br>AN1029<br>AN1041<br>AN1049<br>AN1053<br>AN1077<br>AN1079<br>AN1082<br>AN1091<br>AN1097<br>AN1112<br>AN1121<br>AN1136<br>AN1139<br>AN1166<br>AN1185<br>AN1191<br>AN1194<br>AN1196<br>AN1197<br>AN1203<br>AN1205<br>AN1207<br>AN1212<br>AN1232<br>AN1247<br>AN1249<br>AN1305<br>AN1315<br>AN1324<br>AN1331<br>AN1340<br>AN1341<br>AN1414<br>AN1418<br>AN1424<br>AN1426<br>AN1433<br>AN1464<br>AN1465<br>AN1468<br>AN1469<br>AN1486<br>AN1502<br>AN1505<br>AN1527<br>AN1546<br>AN1547<br>AN1569<br>AN1637<br>AN1639<br>AN1647<br>AN1671<br>AN1677<br>AN1710<br>AN1720<br>AN1725<br>AN1727<br>AN1736<br>AN1761<br>AN1767<br>AN1769<br>AN1773<br>AN1787 |

| | | | | | |
|---|---|---|---|---|---|
| | | | | AN1157<br>AN1403<br>AN2100<br>AN2858<br>AN2863<br>AN3010<br>AN3138<br>AN3235<br>AN3302<br>AN3335<br>AN3379<br>AN3557<br>AN3746<br>AN4073<br>AN4189<br>AN4289<br>AN4432<br>AN4462<br>AN4484<br>AN4747<br>AN4941<br>AN4954<br>AN4966<br>AN4979<br>AN5052<br>AN5349<br>AN5451<br>AN5493<br>AN5611<br>AN5618<br>AN6757<br>AN6874<br>AN6948<br>AN7440<br>AN7477<br>AN7494<br>AN7658<br>AN7832<br>AN8488<br>AN8611<br>AN8690<br>AN0054<br>AN0312<br>AN0337<br>AN0594<br>AN0865<br>AN1325<br>AN1480<br>AN1945<br>AN2886<br>AN3108<br>AN3110<br>AN3426<br>AN4388<br>AN4487<br>AN4501<br>AN4533<br>AN4639<br>AN4651<br>AN4655<br>AN4901<br>AN5139<br>AN5323<br>AN5852<br>AN5977<br>AN6233 | AN1801<br>AN1817<br>AN1818<br>AN1822<br>AN1834<br>AN1840<br>AN1851<br>AN1863<br>AN1882<br>AN1891<br>AN1909<br>AN1915<br>AN1942<br>AN1946<br>AN1953<br>AN1968<br>AN1972<br>AN1987<br>AN1997<br>AN2000<br>AN2001<br>AN2040<br>AN2043<br>AN2053<br>AN2086<br>AN2107<br>AN2119<br>AN2125<br>AN2130<br>AN2173<br>AN2188<br>AN2195<br>AN2203<br>AN2212<br>AN2238<br>AN2245<br>AN2246<br>AN2255<br>AN2257<br>AN2290<br>AN2295<br>AN2317<br>AN2337<br>AN2349<br>AN2382<br>AN2405<br>AN2414<br>AN2437<br>AN2470<br>AN2479<br>AN2485<br>AN2494<br>AN2516<br>AN2535<br>AN2546<br>AN2547<br>AN2548<br>AN2549<br>AN2558<br>AN2625<br>AN2646<br>AN2662<br>AN2663<br>AN2692<br>AN2698<br>AN2701 |

| | | | | | |
|---|---|---|---|---|---|
| | | | | AN6281<br>AN6324<br>AN6489<br>AN6515<br>AN6653<br>AN6797<br>AN6832<br>AN6838<br>AN6896<br>AN7678<br>AN8040<br>AN8199<br>AN8698<br>AN8854<br>AN8868<br>AN8932<br>AN9005<br>AN0136<br>AN0252<br>AN0779<br>AN1122<br>AN1153<br>AN1525<br>AN1635<br>AN1666<br>AN1976<br>AN2577<br>AN2902<br>AN3058<br>AN3199<br>AN3208<br>AN3416<br>AN4692<br>AN4840<br>AN4989<br>AN5106<br>AN5650<br>AN5651<br>AN5774<br>AN5983<br>AN6527<br>AN6592<br>AN6813<br>AN6875<br>AN6975<br>AN7260<br>AN7429<br>AN7466<br>AN7949<br>AN7955<br>AN8072<br>AN8334<br>AN8689<br>AN0453<br>AN0465<br>AN0743<br>AN1054<br>AN1322<br>AN1352<br>AN1930<br>AN2452<br>AN2731<br>AN2791<br>AN3013<br>AN3171<br>AN3278 | AN2737<br>AN2775<br>AN2777<br>AN2815<br>AN2824<br>AN2833<br>AN2835<br>AN2871<br>AN2876<br>AN2881<br>AN2888<br>AN2890<br>AN2892<br>AN2914<br>AN2917<br>AN2918<br>AN2935<br>AN2977<br>AN2980<br>AN2981<br>AN2985<br>AN2992<br>AN2996<br>AN3017<br>AN3019<br>AN3022<br>AN3024<br>AN3065<br>AN3066<br>AN3103<br>AN3104<br>AN3112<br>AN3113<br>AN3121<br>AN3124<br>AN3151<br>AN3154<br>AN3172<br>AN3179<br>AN3212<br>AN3220<br>AN3225<br>AN3226<br>AN3227<br>AN3247<br>AN3264<br>AN3278<br>AN3299<br>AN3303<br>AN3310<br>AN3314<br>AN3315<br>AN3351<br>AN3366<br>AN3380<br>AN3386<br>AN3398<br>AN3408<br>AN3413<br>AN3454<br>AN3480<br>AN3481<br>AN3482<br>AN3513<br>AN3527<br>AN3529 |

| | | | | | |
|---|---|---|---|---|---|
| | | | | AN3479<br>AN3598<br>AN3928<br>AN4275<br>AN4366<br>AN4571<br>AN4713<br>AN4869<br>AN5159<br>AN5267<br>AN5327<br>AN5502<br>AN5586<br>AN5701<br>AN6411<br>AN6518<br>AN6553<br>AN6646<br>AN6892<br>AN7171<br>AN7373<br>AN7478<br>AN7649<br>AN7705<br>AN7943<br>AN8140<br>AN8697<br>AN8702<br>AN8721<br>AN8768<br>AN8973<br>AN0199<br>AN0976<br>AN1196<br>AN1304<br>AN2143<br>AN3712<br>AN3956<br>AN4556<br>AN4589<br>AN4623<br>AN4881<br>AN5146<br>AN5150<br>AN5860<br>AN6083<br>AN6493<br>AN6547<br>AN6565<br>AN6587<br>AN6624<br>AN6647<br>AN7302<br>AN7358<br>AN7496<br>AN9016<br>AN0786<br>AN1622<br>AN1833<br>AN1856<br>AN2061<br>AN2080<br>AN2141<br>AN2181<br>AN2504<br>AN2545 | AN3593<br>AN3605<br>AN3610<br>AN3646<br>AN3661<br>AN3679<br>AN3685<br>AN3697<br>AN3706<br>AN3707<br>AN3713<br>AN3731<br>AN3737<br>AN3741<br>AN3747<br>AN3760<br>AN3781<br>AN3790<br>AN3796<br>AN3798<br>AN3803<br>AN3820<br>AN3823<br>AN3843<br>AN3847<br>AN3850<br>AN3853<br>AN3880<br>AN3881<br>AN3894<br>AN3899<br>AN3902<br>AN3912<br>AN3925<br>AN3928<br>AN3936<br>AN3941<br>AN3952<br>AN3960<br>AN4034<br>AN4035<br>AN4041<br>AN4047<br>AN4048<br>AN4055<br>AN4060<br>AN4073<br>AN4087<br>AN4099<br>AN4139<br>AN4140<br>AN4163<br>AN4168<br>AN4178<br>AN4182<br>AN4202<br>AN4216<br>AN4224<br>AN4248<br>AN4264<br>AN4265<br>AN4276<br>AN4285<br>AN4314<br>AN4329<br>AN4340 |

| | | | | | |
|---|---|---|---|---|---|
| | | | | AN2809 | AN4371 |
| | | | | AN2880 | AN4379 |
| | | | | AN3002 | AN4393 |
| | | | | AN3065 | AN4409 |
| | | | | AN4147 | AN4431 |
| | | | | AN4259 | AN4440 |
| | | | | AN4894 | AN4444 |
| | | | | AN4956 | AN4466 |
| | | | | AN5494 | AN4474 |
| | | | | AN5827 | AN4485 |
| | | | | AN5974 | AN4487 |
| | | | | AN6106 | AN4492 |
| | | | | AN6228 | AN4493 |
| | | | | AN6942 | AN4499 |
| | | | | AN7123 | AN4507 |
| | | | | AN7225 | AN4520 |
| | | | | AN7435 | AN4552 |
| | | | | AN7469 | AN4602 |
| | | | | AN7637 | AN4614 |
| | | | | AN7646 | AN4645 |
| | | | | AN7728 | AN4649 |
| | | | | AN7824 | AN4656 |
| | | | | AN8163 | AN4668 |
| | | | | AN8209 | AN4688 |
| | | | | AN8364 | AN4690 |
| | | | | AN0345 | AN4692 |
| | | | | AN0402 | AN4695 |
| | | | | AN0558 | AN4699 |
| | | | | AN0846 | AN4714 |
| | | | | AN0848 | AN4729 |
| | | | | AN1323 | AN4732 |
| | | | | AN1464 | AN4741 |
| | | | | AN2121 | AN4745 |
| | | | | AN2374 | AN4768 |
| | | | | AN2423 | AN4772 |
| | | | | AN3091 | AN4773 |
| | | | | AN3116 | AN4784 |
| | | | | AN3306 | AN4787 |
| | | | | AN3385 | AN4790 |
| | | | | AN3846 | AN4805 |
| | | | | AN3953 | AN4821 |
| | | | | AN4545 | AN4829 |
| | | | | AN4590 | AN4841 |
| | | | | AN6030 | AN4843 |
| | | | | AN6170 | AN4856 |
| | | | | AN6580 | AN4861 |
| | | | | AN6627 | AN4866 |
| | | | | AN6934 | AN4871 |
| | | | | AN7192 | AN4872 |
| | | | | AN8283 | AN4910 |
| | | | | AN8292 | AN4916 |
| | | | | AN8564 | AN4934 |
| | | | | AN8959 | AN4941 |
| | | | | AN9373 | AN4946 |
| | | | | AN0201 | AN4983 |
| | | | | AN0211 | AN4985 |
| | | | | AN0491 | AN5001 |
| | | | | AN0886 | AN5002 |
| | | | | AN0916 | AN5012 |
| | | | | AN1090 | AN5015 |
| | | | | AN1211 | AN5018 |
| | | | | AN1219 | AN5027 |
| | | | | AN1332 | AN5074 |
| | | | | AN1633 | AN5110 |
| | | | | AN1898 | AN5144 |
| | | | | AN2008 | AN5147 |

| | | | | | |
|---|---|---|---|---|---|
| | | | | AN2951<br>AN3380<br>AN3725<br>AN4974<br>AN5283<br>AN5876<br>AN6062<br>AN6128<br>AN6169<br>AN6201<br>AN6206<br>AN6660<br>AN6688<br>AN6790<br>AN6798<br>AN6897<br>AN8048<br>AN8065<br>AN8234<br>AN8368<br>AN8497<br>AN8498<br>AN8614<br>AN8637<br>AN8869<br>AN8894<br>AN9287<br>AN0019<br>AN0196<br>AN0480<br>AN1361<br>AN2348<br>AN2517<br>AN2554<br>AN2909<br>AN3973<br>AN4129<br>AN4280<br>AN4369<br>AN4381<br>AN4498<br>AN4828<br>AN6100<br>AN6217<br>AN6262<br>AN6993<br>AN7001<br>AN7314<br>AN7609<br>AN8138<br>AN8214<br>AN8442<br>AN8819<br>AN9322<br>AN0278<br>AN0434<br>AN0626<br>AN0770<br>AN1160<br>AN1407<br>AN1526<br>AN1708<br>AN1862<br>AN1891<br>AN1969<br>AN3071 | AN5157<br>AN5161<br>AN5170<br>AN5177<br>AN5187<br>AN5189<br>AN5201<br>AN5209<br>AN5231<br>AN5245<br>AN5258<br>AN5268<br>AN5284<br>AN5294<br>AN5312<br>AN5352<br>AN5353<br>AN5355<br>AN5357<br>AN5359<br>AN5399<br>AN5404<br>AN5415<br>AN5421<br>AN5426<br>AN5432<br>AN5436<br>AN5461<br>AN5483<br>AN5492<br>AN5495<br>AN5513<br>AN5526<br>AN5530<br>AN5548<br>AN5556<br>AN5566<br>AN5569<br>AN5621<br>AN5633<br>AN5660<br>AN5670<br>AN5691<br>AN5697<br>AN5713<br>AN5719<br>AN5737<br>AN5763<br>AN5766<br>AN5769<br>AN5778<br>AN5815<br>AN5821<br>AN5824<br>AN5830<br>AN5857<br>AN5860<br>AN5869<br>AN5901<br>AN5924<br>AN5945<br>AN5971<br>AN5989<br>AN5990<br>AN5996<br>AN6024 |

| | | | | | |
|---|---|---|---|---|---|
| | | | | AN3377 | AN6049 |
| | | | | AN3625 | AN6078 |
| | | | | AN3893 | AN6079 |
| | | | | AN3998 | AN6084 |
| | | | | AN4128 | AN6092 |
| | | | | AN4559 | AN6105 |
| | | | | AN4637 | AN6116 |
| | | | | AN5036 | AN6124 |
| | | | | AN5122 | AN6135 |
| | | | | AN5202 | AN6159 |
| | | | | AN5259 | AN6167 |
| | | | | AN5348 | AN6170 |
| | | | | AN5409 | AN6181 |
| | | | | AN5585 | AN6184 |
| | | | | AN6138 | AN6185 |
| | | | | AN6507 | AN6186 |
| | | | | AN7082 | AN6188 |
| | | | | AN7205 | AN6202 |
| | | | | AN7595 | AN6207 |
| | | | | AN7600 | AN6213 |
| | | | | AN7764 | AN6218 |
| | | | | AN7765 | AN6224 |
| | | | | AN7883 | AN6225 |
| | | | | AN8222 | AN6229 |
| | | | | AN8271 | AN6250 |
| | | | | AN9323 | AN6268 |
| | | | | AN1154 | AN6271 |
| | | | | AN2084 | AN6306 |
| | | | | AN2391 | AN6322 |
| | | | | AN3647 | AN6327 |
| | | | | AN3796 | AN6337 |
| | | | | AN3873 | AN6340 |
| | | | | AN3975 | AN6342 |
| | | | | AN4286 | AN6367 |
| | | | | AN4315 | AN6372 |
| | | | | AN4568 | AN6378 |
| | | | | AN5657 | AN6384 |
| | | | | AN5854 | AN6507 |
| | | | | AN6368 | AN6521 |
| | | | | AN6536 | AN6528 |
| | | | | AN6908 | AN6542 |
| | | | | AN7021 | AN6547 |
| | | | | AN7147 | AN6580 |
| | | | | AN7264 | AN6584 |
| | | | | AN7444 | AN6591 |
| | | | | AN8914 | AN6595 |
| | | | | AN9418 | AN6617 |
| | | | | AN9432 | AN6626 |
| | | | | AN0073 | AN6633 |
| | | | | AN0559 | AN6649 |
| | | | | AN0898 | AN6662 |
| | | | | AN1674 | AN6666 |
| | | | | AN2380 | AN6679 |
| | | | | AN2499 | AN6687 |
| | | | | AN2509 | AN6694 |
| | | | | AN2808 | AN6703 |
| | | | | AN3203 | AN6705 |
| | | | | AN3314 | AN6726 |
| | | | | AN3634 | AN6730 |
| | | | | AN3913 | AN6731 |
| | | | | AN4254 | AN6736 |
| | | | | AN4444 | AN6740 |
| | | | | AN4454 | AN6747 |
| | | | | AN5209 | AN6763 |
| | | | | AN5223 | AN6769 |
| | | | | AN5276 | AN6787 |

| | | | | | |
|---|---|---|---|---|---|
| | | | | AN5515 | AN6845 |
| | | | | AN5745 | AN6848 |
| | | | | AN6140 | AN6856 |
| | | | | AN6614 | AN6876 |
| | | | | AN6716 | AN6881 |
| | | | | AN6862 | AN6890 |
| | | | | AN7037 | AN6894 |
| | | | | AN7331 | AN6913 |
| | | | | AN7790 | AN6922 |
| | | | | AN0186 | AN6924 |
| | | | | AN0359 | AN6931 |
| | | | | AN0560 | AN6944 |
| | | | | AN0720 | AN6962 |
| | | | | AN0815 | AN6987 |
| | | | | AN0850 | AN6989 |
| | | | | AN1081 | AN6999 |
| | | | | AN1577 | AN7017 |
| | | | | AN1661 | AN7022 |
| | | | | AN1868 | AN7028 |
| | | | | AN1967 | AN7037 |
| | | | | AN2291 | AN7076 |
| | | | | AN3037 | AN7084 |
| | | | | AN3237 | AN7092 |
| | | | | AN3373 | AN7105 |
| | | | | AN3418 | AN7107 |
| | | | | AN3597 | AN7130 |
| | | | | AN3637 | AN7134 |
| | | | | AN3969 | AN7145 |
| | | | | AN4081 | AN7148 |
| | | | | AN4313 | AN7160 |
| | | | | AN4445 | AN7162 |
| | | | | AN4762 | AN7181 |
| | | | | AN4787 | AN7190 |
| | | | | AN5081 | AN7191 |
| | | | | AN6042 | AN7193 |
| | | | | AN6256 | AN7221 |
| | | | | AN7102 | AN7245 |
| | | | | AN7157 | AN7280 |
| | | | | AN7471 | AN7287 |
| | | | | AN7527 | AN7289 |
| | | | | AN7578 | AN7334 |
| | | | | AN7961 | AN7338 |
| | | | | AN9105 | AN7345 |
| | | | | AN9343 | AN7354 |
| | | | | AN0492 | AN7357 |
| | | | | AN1379 | AN7387 |
| | | | | AN1568 | AN7389 |
| | | | | AN1838 | AN7394 |
| | | | | AN2132 | AN7409 |
| | | | | AN2420 | AN7447 |
| | | | | AN2427 | AN7448 |
| | | | | AN2840 | AN7453 |
| | | | | AN3558 | AN7454 |
| | | | | AN3735 | AN7467 |
| | | | | AN3772 | AN7469 |
| | | | | AN3987 | AN7470 |
| | | | | AN4014 | AN7491 |
| | | | | AN4020 | AN7505 |
| | | | | AN4295 | AN7529 |
| | | | | AN5746 | AN7533 |
| | | | | AN5815 | AN7534 |
| | | | | AN5872 | AN7539 |
| | | | | AN6830 | AN7542 |
| | | | | AN6998 | AN7544 |
| | | | | AN7433 | AN7564 |
| | | | | AN7489 | AN7568 |

| | | | | | |
|---|---|---|---|---|---|
| | | | | AN7866<br>AN8001<br>AN8525<br>AN8676<br>AN9350<br>AN9439<br>AN0183<br>AN0721<br>AN0843<br>AN1083<br>AN1356<br>AN1416<br>AN1542<br>AN2129<br>AN2352<br>AN2582<br>AN2992<br>AN3004<br>AN4776<br>AN5342<br>AN5640<br>AN5732<br>AN5800<br>AN5926<br>AN6490<br>AN6523<br>AN7391<br>AN7423<br>AN7861<br>AN9299<br>AN9346<br>AN0150<br>AN0632<br>AN1200<br>AN1312<br>AN1587<br>AN1735<br>AN1982<br>AN2496<br>AN2883<br>AN2925<br>AN3842<br>AN4227<br>AN4529<br>AN4682<br>AN4882<br>AN5805<br>AN6014<br>AN6555<br>AN6710<br>AN7447<br>AN9070<br>AN9092<br>AN9138<br>AN0837<br>AN0973<br>AN1559<br>AN1772<br>AN1913<br>AN1916<br>AN2212<br>AN2339<br>AN2466<br>AN2702<br>AN2867<br>AN2934 | AN7579<br>AN7598<br>AN7619<br>AN7623<br>AN7624<br>AN7648<br>AN7666<br>AN7667<br>AN7691<br>AN7706<br>AN7711<br>AN7714<br>AN7715<br>AN7716<br>AN7718<br>AN7732<br>AN7738<br>AN7772<br>AN7776<br>AN7792<br>AN7799<br>AN7820<br>AN7847<br>AN7869<br>AN7873<br>AN7892<br>AN7913<br>AN7919<br>AN7925<br>AN7932<br>AN7944<br>AN7962<br>AN7985<br>AN8004<br>AN8009<br>AN8036<br>AN8043<br>AN8049<br>AN8050<br>AN8061<br>AN8069<br>AN8079<br>AN8089<br>AN8099<br>AN8103<br>AN8109<br>AN8110<br>AN8114<br>AN8148<br>AN8151<br>AN8167<br>AN8171<br>AN8176<br>AN8179<br>AN8180<br>AN8190<br>AN8191<br>AN8192<br>AN8199<br>AN8201<br>AN8235<br>AN8240<br>AN8248<br>AN8257<br>AN8258<br>AN8270 |

| | | | | | |
|---|---|---|---|---|---|
| | | | | AN3556<br>AN3609<br>AN3720<br>AN4096<br>AN4435<br>AN5464<br>AN5642<br>AN5659<br>AN6202<br>AN6315<br>AN6683<br>AN7392<br>AN7760<br>AN8166<br>AN8635<br>AN8778<br>AN0215<br>AN0228<br>AN0431<br>AN0565<br>AN0610<br>AN1194<br>AN1257<br>AN1446<br>AN1845<br>AN2932<br>AN3311<br>AN3463<br>AN3910<br>AN4062<br>AN4395<br>AN4867<br>AN5421<br>AN5717<br>AN5782<br>AN5834<br>AN5979<br>AN6017<br>AN6056<br>AN6894<br>AN7536<br>AN7537<br>AN7776<br>AN8210<br>AN8722<br>AN8822<br>AN9410<br>AN9447<br>AN9505<br>AN0141<br>AN0209<br>AN0307<br>AN0368<br>AN0523<br>AN1217<br>AN2523<br>AN2657<br>AN2905<br>AN2919<br>AN3224<br>AN3490<br>AN3581<br>AN3894<br>AN4114<br>AN4728<br>AN4735 | AN8300<br>AN8304<br>AN8322<br>AN8326<br>AN8333<br>AN8336<br>AN8344<br>AN8345<br>AN8374<br>AN8389<br>AN8403<br>AN8415<br>AN8430<br>AN8457<br>AN8461<br>AN8481<br>AN8493<br>AN8494<br>AN8496<br>AN8499<br>AN8500<br>AN8503<br>AN8525<br>AN8553<br>AN8557<br>AN8575<br>AN8594<br>AN8628<br>AN8636<br>AN8640<br>AN8645<br>AN8654<br>AN8657<br>AN8695<br>AN8722<br>AN8724<br>AN8738<br>AN8752<br>AN8753<br>AN8799<br>AN8801<br>AN8804<br>AN8810<br>AN8814<br>AN8816<br>AN8829<br>AN8831<br>AN8837<br>AN8838<br>AN8873<br>AN8906<br>AN8961<br>AN8986<br>AN8993<br>AN9000<br>AN9028<br>AN9049<br>AN9073<br>AN9108<br>AN9155<br>AN9159<br>AN9176<br>AN9212<br>AN9222<br>AN9286<br>AN9288 |

| | | | | | |
|---|---|---|---|---|---|
| | | | | AN4914<br>AN4993<br>AN5014<br>AN5432<br>AN6278<br>AN6657<br>AN6666<br>AN6742<br>AN6816<br>AN6959<br>AN7030<br>AN7244<br>AN7559<br>AN7912<br>AN8641<br>AN9291<br>AN0170<br>AN0173<br>AN0813<br>AN1150<br>AN1175<br>AN2596<br>AN2682<br>AN3103<br>AN3318<br>AN3329<br>AN3419<br>AN4944<br>AN5110<br>AN5390<br>AN6729<br>AN6837<br>AN7058<br>AN7104<br>AN7589<br>AN8492<br>AN8622<br>AN8813<br>AN8823<br>AN9512<br>AN0015<br>AN0062<br>AN0348<br>AN0554<br>AN0664<br>AN0819<br>AN1082<br>AN1147<br>AN1545<br>AN1608<br>AN1701<br>AN2588<br>AN2989<br>AN3495<br>AN3526<br>AN3659<br>AN4302<br>AN4851<br>AN8336<br>AN8751<br>AN8936<br>AN9448<br>AN9450<br>AN9467<br>AN0240<br>AN0374 | AN9316<br>AN9323<br>AN9342<br>AN9347<br>AN9373<br>AN9384<br>AN9386<br>AN9397<br>AN9449<br>AN9453<br>AN9466<br>AN9523<br>AN9531 |

| | | | | |
|---|---|---|---|---|
| | | | | AN0411<br>AN0797<br>AN0806<br>AN0824<br>AN0929<br>AN1243<br>AN1366<br>AN1692<br>AN1839<br>AN1847<br>AN3372<br>AN3879<br>AN3881<br>AN3986<br>AN5191<br>AN5491<br>AN6754<br>AN8943<br>AN0053<br>AN0592<br>AN0684<br>AN0796<br>AN1567<br>AN1932<br>AN2118<br>AN2292<br>AN2401<br>AN2598<br>AN3567<br>AN3820<br>AN4721<br>AN5309<br>AN5512<br>AN5760<br>AN6178<br>AN6182<br>AN6719<br>AN7499<br>AN7525<br>AN8405<br>AN8730<br>AN9209<br>AN0143<br>AN0320<br>AN0457<br>AN1053<br>AN1543<br>AN1761<br>AN2330<br>AN2774<br>AN3121<br>AN4078<br>AN4540<br>AN4922<br>AN5156<br>AN5903<br>AN6549<br>AN6717<br>AN7836<br>AN8215<br>AN8322<br>AN9102<br>AN9193<br>AN0311<br>AN1372<br>AN1619 |

| | | | | |
|---|---|---|---|---|
| | | | | AN1650<br>AN2168<br>AN2382<br>AN2735<br>AN3358<br>AN4159<br>AN4231<br>AN4448<br>AN4605<br>AN4937<br>AN5186<br>AN5229<br>AN6511<br>AN6662<br>AN6725<br>AN6999<br>AN7024<br>AN7703<br>AN8565<br>AN8603<br>AN0294<br>AN0571<br>AN1328<br>AN1592<br>AN1953<br>AN2334<br>AN4340<br>AN4437<br>AN4519<br>AN4531<br>AN4863<br>AN4953<br>AN5204<br>AN5208<br>AN5671<br>AN6151<br>AN6594<br>AN6888<br>AN7733<br>AN8110<br>AN8203<br>AN9327<br>AN0317<br>AN0712<br>AN0735<br>AN2272<br>AN2969<br>AN3143<br>AN3169<br>AN3328<br>AN3776<br>AN3875<br>AN3996<br>AN4463<br>AN5905<br>AN6286<br>AN6359<br>AN6378<br>AN6577<br>AN0304<br>AN1527<br>AN2963<br>AN3035<br>AN3585<br>AN3752<br>AN3878 |

| | | | | |
|---|---|---|---|---|
| | | | | AN3930<br>AN4030<br>AN4193<br>AN4329<br>AN4970<br>AN5321<br>AN5786<br>AN5921<br>AN5927<br>AN8836<br>AN9353<br>AN0154<br>AN0412<br>AN0481<br>AN1376<br>AN1491<br>AN1673<br>AN1896<br>AN2149<br>AN2160<br>AN2293<br>AN4624<br>AN5482<br>AN5590<br>AN6251<br>AN6364<br>AN6512<br>AN6938<br>AN7996<br>AN8148<br>AN9385<br>AN0406<br>AN1680<br>AN3152<br>AN3616<br>AN3821<br>AN3889<br>AN4936<br>AN4959<br>AN6103<br>AN7079<br>AN8014<br>AN8238<br>AN8976<br>AN1013<br>AN2191<br>AN2619<br>AN3164<br>AN3345<br>AN5525<br>AN6015<br>AN6681<br>AN7130<br>AN8241<br>AN8262<br>AN8285<br>AN8407<br>AN8714<br>AN9529<br>AN0283<br>AN0328<br>AN0451<br>AN0500<br>AN1602<br>AN1949<br>AN2057 |

| | | | | |
|---|---|---|---|---|
| | | | | AN3811<br>AN4408<br>AN4669<br>AN4850<br>AN5713<br>AN5877<br>AN6538<br>AN1135<br>AN2566<br>AN3179<br>AN5091<br>AN5092<br>AN5480<br>AN5541<br>AN5570<br>AN5811<br>AN5882<br>AN6304<br>AN6711<br>AN7254<br>AN7439<br>AN7791<br>AN8218<br>AN8260<br>AN0484<br>AN2039<br>AN2991<br>AN3201<br>AN4033<br>AN4796<br>AN5399<br>AN6936<br>AN7078<br>AN7223<br>AN8817<br>AN0441<br>AN0644<br>AN1329<br>AN1820<br>AN3343<br>AN4364<br>AN5883<br>AN6715<br>AN6778<br>AN9341<br>AN0604<br>AN1025<br>AN3160<br>AN3830<br>AN4375<br>AN4693<br>AN4865<br>AN5207<br>AN5237<br>AN5350<br>AN5438<br>AN6949<br>AN7023<br>AN7140<br>AN7920<br>AN8107<br>AN0138<br>AN0726<br>AN3074<br>AN3618<br>AN5768 |

| | | | | |
|---|---|---|---|---|
| | | | | AN6179<br>AN6915<br>AN7650<br>AN8058<br>AN8578<br>AN0413<br>AN0705<br>AN1960<br>AN2832<br>AN2926<br>AN3076<br>AN3782<br>AN4499<br>AN4731<br>AN4975<br>AN5073<br>AN5133<br>AN5151<br>AN5833<br>AN6804<br>AN6921<br>AN7722<br>AN7944<br>AN8259<br>AN8426<br>AN8439<br>AN8881<br>AN8895<br>AN9490<br>AN0509<br>AN2170<br>AN4785<br>AN5558<br>AN5821<br>AN6268<br>AN7027<br>AN7488<br>AN7512<br>AN8123<br>AN8837<br>AN9098<br>AN0343<br>AN0713<br>AN2455<br>AN2822<br>AN3681<br>AN6275<br>AN7601<br>AN7820<br>AN9426<br>AN9445<br>AN0804<br>AN0933<br>AN1126<br>AN1146<br>AN1421<br>AN1596<br>AN1893<br>AN2518<br>AN3028<br>AN3230<br>AN4082<br>AN5331<br>AN5831<br>AN5865<br>AN6121 |

| | | | | |
|---|---|---|---|---|
| | | | | AN6207<br>AN6363<br>AN6801<br>AN7210<br>AN7560<br>AN7714<br>AN8216<br>AN8587<br>AN0275<br>AN1032<br>AN1429<br>AN1694<br>AN1734<br>AN5305<br>AN5312<br>AN6529<br>AN6776<br>AN7457<br>AN0397<br>AN3573<br>AN3736<br>AN5565<br>AN1111<br>AN2460<br>AN5340<br>AN5365<br>AN7103<br>AN7436<br>AN9172<br>AN0267<br>AN2298<br>AN2461<br>AN4737<br>AN5906<br>AN6006<br>AN8128<br>AN8284<br>AN8805<br>AN1289<br>AN1958<br>AN2254<br>AN2310<br>AN3190<br>AN4230<br>AN5398<br>AN5804<br>AN6551<br>AN7240<br>AN7503<br>AN0950<br>AN2465<br>AN2781<br>AN2785<br>AN2922<br>AN6856<br>AN0339<br>AN0515<br>AN1521<br>AN1771<br>AN2236<br>AN2317<br>AN3344<br>AN4461<br>AN4528<br>AN4942<br>AN5429 |

| | | | | |
|---|---|---|---|---|
| | | | | AN5724<br>AN5997<br>AN6371<br>AN7587<br>AN7715<br>AN8775<br>AN9112<br>AN0116<br>AN0385<br>AN0561<br>AN1107<br>AN1397<br>AN1423<br>AN1753<br>AN1870<br>AN2003<br>AN2045<br>AN2688<br>AN3197<br>AN3528<br>AN4951<br>AN5001<br>AN6704<br>AN7756<br>AN8480<br>AN8560<br>AN8808<br>AN9321<br>AN9409<br>AN9479<br>AN0036<br>AN0596<br>AN2050<br>AN2750<br>AN4490<br>AN4584<br>AN4790<br>AN5762<br>AN6939<br>AN8435<br>AN1717<br>AN1866<br>AN2336<br>AN2337<br>AN2408<br>AN2686<br>AN4525<br>AN5625<br>AN6526<br>AN6638<br>AN7150<br>AN7424<br>AN8786<br>AN9305<br>AN1387<br>AN1689<br>AN2550<br>AN2738<br>AN2947<br>AN3368<br>AN3860<br>AN6223<br>AN7633<br>AN8402<br>AN8455<br>AN8942 |

| | | | | |
|---|---|---|---|---|
| | | | | AN9347<br>AN2463<br>AN3216<br>AN3849<br>AN4296<br>AN4798<br>AN5212<br>AN5916<br>AN6878<br>AN7934<br>AN8257<br>AN9242<br>AN0160<br>AN0295<br>AN1295<br>AN1724<br>AN2817<br>AN2977<br>AN3083<br>AN3577<br>AN4265<br>AN5172<br>AN5750<br>AN5858<br>AN6557<br>AN8161<br>AN8857<br>AN8867<br>AN8871<br>AN9134<br>AN2086<br>AN2264<br>AN2678<br>AN3067<br>AN4754<br>AN5893<br>AN6197<br>AN6743<br>AN6852<br>AN8178<br>AN8773<br>AN0097<br>AN0129<br>AN3156<br>AN4500<br>AN5396<br>AN5424<br>AN5704<br>AN5771<br>AN5797<br>AN7144<br>AN7162<br>AN8202<br>AN9515<br>AN0111<br>AN0931<br>AN0957<br>AN2144<br>AN2470<br>AN3339<br>AN3721<br>AN4241<br>AN4574<br>AN4663<br>AN6532<br>AN7129 |

| | | | | |
|---|---|---|---|---|
| | | | | AN7448<br>AN7454<br>AN1644<br>AN2014<br>AN2060<br>AN2262<br>AN2524<br>AN3823<br>AN5188<br>AN5386<br>AN5416<br>AN5572<br>AN5758<br>AN6007<br>AN7166<br>AN8457<br>AN8727<br>AN8941<br>AN2182<br>AN4803<br>AN5119<br>AN6761<br>AN7303<br>AN7743<br>AN9106<br>AN9434<br>AN0407<br>AN0619<br>AN2128<br>AN3722<br>AN4101<br>AN4283<br>AN5023<br>AN7619<br>AN8049<br>AN8840<br>AN1076<br>AN1556<br>AN1871<br>AN1994<br>AN2571<br>AN3104<br>AN3587<br>AN3633<br>AN4299<br>AN5051<br>AN5783<br>AN6556<br>AN8784<br>AN1158<br>AN1339<br>AN1744<br>AN2213<br>AN2659<br>AN3546<br>AN5099<br>AN5749<br>AN6287<br>AN6839<br>AN8753<br>AN1098<br>AN1176<br>AN2248<br>AN2760<br>AN2799<br>AN4092 |

| | | | | |
|---|---|---|---|---|
| | | | | AN4430<br>AN4644<br>AN6902<br>AN7177<br>AN7544<br>AN0903<br>AN1100<br>AN1517<br>AN1579<br>AN1697<br>AN2409<br>AN3359<br>AN4065<br>AN4841<br>AN7338<br>AN7498<br>AN8002<br>AN8311<br>AN9007<br>AN0327<br>AN0577<br>AN3640<br>AN4210<br>AN4825<br>AN7366<br>AN7635<br>AN0105<br>AN2138<br>AN3184<br>AN4338<br>AN5573<br>AN6753<br>AN6980<br>AN0649<br>AN3021<br>AN4034<br>AN4415<br>AN4614<br>AN5605<br>AN5649<br>AN5738<br>AN6129<br>AN6581<br>AN7547<br>AN7766<br>AN8528<br>AN9213<br>AN9429<br>AN1019<br>AN1059<br>AN1167<br>AN1712<br>AN1715<br>AN2828<br>AN3680<br>AN4027<br>AN4029<br>AN4471<br>AN4958<br>AN4990<br>AN5619<br>AN7584<br>AN7894<br>AN8908<br>AN9221<br>AN0454 |

| | | | | |
|---|---|---|---|---|
| | | | | AN1103<br>AN3159<br>AN4376<br>AN4662<br>AN5628<br>AN7484<br>AN7594<br>AN9075<br>AN9156<br>AN1101<br>AN2763<br>AN2857<br>AN3788<br>AN3840<br>AN4350<br>AN5687<br>AN5699<br>AN6568<br>AN8100<br>AN0265<br>AN1286<br>AN1727<br>AN2698<br>AN3279<br>AN3959<br>AN6789<br>AN8044<br>AN8345<br>AN2878<br>AN2990<br>AN3228<br>AN3945<br>AN6762<br>AN7239<br>AN8449<br>AN0075<br>AN0803<br>AN2289<br>AN2776<br>AN3783<br>AN4757<br>AN4832<br>AN5911<br>AN7721<br>AN9433<br>AN1703<br>AN1880<br>AN1897<br>AN2083<br>AN2898<br>AN6009<br>AN7500<br>AN7553<br>AN8627<br>AN8720<br>AN0414<br>AN0551<br>AN1178<br>AN1488<br>AN3276<br>AN3562<br>AN4373<br>AN5946<br>AN7246<br>AN7699<br>AN7855 |

| | | | | |
|---|---|---|---|---|
| | | | | AN8085<br>AN9198<br>AN0918<br>AN3600<br>AN6257<br>AN7340<br>AN7422<br>AN8900<br>AN9145<br>AN0383<br>AN3504<br>AN4539<br>AN5898<br>AN6842<br>AN7048<br>AN8265<br>AN9111<br>AN9119<br>AN0805<br>AN1419<br>AN1595<br>AN2390<br>AN4053<br>AN5240<br>AN6296<br>AN7709<br>AN8078<br>AN1986<br>AN3341<br>AN4818<br>AN5304<br>AN5984<br>AN7887<br>AN8737<br>AN0630<br>AN1350<br>AN2082<br>AN2481<br>AN3805<br>AN4116<br>AN4621<br>AN4910<br>AN6500<br>AN6799<br>AN8809<br>AN0782<br>AN0901<br>AN1340<br>AN2223<br>AN2231<br>AN2717<br>AN3331<br>AN3730<br>AN7732<br>AN7889<br>AN8063<br>AN0018<br>AN0477<br>AN1474<br>AN2177<br>AN2449<br>AN3150<br>AN4507<br>AN4792<br>AN4917<br>AN6267 |

| | | | | |
|---|---|---|---|---|
| | | | | AN6872<br>AN7180<br>AN8754<br>AN9146<br>AN0372<br>AN1241<br>AN2482<br>AN2997<br>AN3516<br>AN3679<br>AN5177<br>AN7022<br>AN8278<br>AN9072 |

#### S3- Temporal GO Description of Time Courses of *A. nidulans Exposed to Micafungin*

Following images were taken from the results of the output of DPoP after a GO term enrichment analysis of the results. Each image represents results of one timepoint, neglecting timepoint 13.

##### Supplementary Data S3.1 – Temporal GO Description of DPoP analysis of Phosphoproteomic Data

Each one of the following pictures is an aggregation of outputs of a GO term enrichment analysis on the results of DPoP at each specific point along the phosphoproteomic time course.

Supplementary Data S3.1.1

### Timepoint 1

Molecular Function

| | |
|---|---|
| 131 / 597 | "Catalysis of a biochemical reaction at physiological temperatures. In biologically catalyzed reactions, the reactants are known as substrates, and the catalysts are naturally occurring macromolecular substances kn |
| 12 / 114 | "Binding to an RNA molecule or a portion thereof." [GOC:jl, GOC:mah] |
| 9 / 73 | "Binding to a nucleic acid region that regulates a nucleic acid-based process. Such processes include transcription, DNA replication, and DNA repair." [GOC:txnOH] |
| 9 / 73 | "Binding to a methylated cytosine/guanine dinucleotide." [GOC:jl, PMID:11746232] |
| 9 / 73 | "Any selective and non-covalent interaction with a nucleic acid involved in the initiation, activation, perpetuation, repression or termination of polypeptide synthesis at the ribosome." [GOC:dph, GOC:tb, GOC:vw] |
| 9 / 72 | "Binding to a RNA/DNA hybrid." [GOC:ecd] |
| 8 / 66 | "Binding to a ribonucleotide, any compound consisting of a ribonucleoside that is esterified with (ortho)phosphate or an oligophosphate at any hydroxyl group on the ribose moiety." [GOC:mah] |
| 8 / 65 | "Binding to a nucleotide, any compound consisting of a nucleoside that is esterified with (ortho)phosphate or an oligophosphate at any hydroxyl group on the ribose or deoxyribose." [GOC:mah, ISBN:0198547684] |
| 20 / 60 | "Binding to damaged DNA." [GOC:jl] |
| 8 / 64 | "Binding to nicotinamide adenine dinucleotide, a coenzyme involved in many redox and biosynthetic reactions; binding may be to either the oxidized form, NAD+, or the reduced form, NADH." [GOC:ai] |

Biological Process

| | |
|---|---|
| 2 / 39 | "Any process that modulates the frequency, rate or extent of cellular DNA-templated transcription." [GOC:go_curators, GOC:txnOH] |
| 0 / 22 | "Any process involved in the conversion of a primary ribosomal RNA (rRNA) transcript into one or more mature rRNA molecules." [GOC:curators] |
| 0 / 21 | "Any process that results in the conversion of one or more primary non-coding RNA (ncRNA) transcripts into one or more mature ncRNA molecules." [GOC:mah] |
| 1 / 26 | "Any process involved in the maturation of a precursor Small SubUnit (SSU) ribosomal RNA (rRNA) molecule into a mature SSU-rRNA molecule." [GOC:curators] |
| 3 / 3 | "The conversion of a single-cell organism from one mating type to another by the precise replacement of a DNA sequence at the expressed mating type locus with a copy of a sequence from a donor locus." [PMID:99 |
| 3 / 3 | "A process that forms a boundary that limits the spreading of heterochromatin along a chromosome." [GOC:mah] |
| 0 / 17 | "Any process involved in the maturation of a precursor 5.8S ribosomal RNA (rRNA) molecule into a mature 5.8S rRNA molecule." [GOC:curators] |
| 0 / 17 | "Any process involved in forming the mature 5' end of an rRNA molecule." [GOC:krc] |
| 0 / 16 | "Any phosphodiester bond hydrolysis involved in the conversion of a primary ribosomal RNA (rRNA) transcript into a mature rRNA molecule." [GOC:curators] |
| 0 / 16 | "The chemical reactions and pathways involving rRNA, ribosomal RNA, a structural constituent of ribosomes." [ISBN:0198506732] |

Cellular Compartment

| | |
|---|---|
| 4 / 5 | "A region of a chromosome at which a DNA double-strand break has occurred. DNA damage signaling and repair proteins accumulate at the lesion to respond to the damage and repair the DNA to form a continuo |
| 4 / 5 | "A region of a chromosome at which DNA damage has occurred. DNA damage signaling and repair proteins accumulate at the lesion to respond to the damage and repair the DNA to form a continuous DNA helix." |
| 0 / 19 | "The chemical reactions and pathways involving an organic substance, any molecular entity containing carbon." [GOC:mah] |
| 16 / 107 | "The volume enclosed by the nuclear inner membrane." [GOC:mah, GOC:pz] |
| 0 / 15 | "The chemical reactions and pathways resulting in the formation of substances; typically the energy-requiring part of metabolism in which simpler substances are transformed into more complex ones." [GOC:curat |
| 0 / 14 | "The chemical reactions and pathways involving organic or inorganic compounds that contain nitrogen." [GOC:jl, ISBN:0198506732] |
| 0 / 14 | "The chemical reactions and pathways resulting in many of the chemical changes of compounds that are not necessarily required for growth and maintenance of cells, and are often unique to a taxon. In multicellul |
| 3 / 4 | "Any process that results in a change in state or activity of a cell (in terms of movement, secretion, enzyme production, gene expression, etc.) as a result of deprivation of iron ions." [GOC:mg] |
| 153 / 677 | "A stable assembly of two or more macromolecules, i.e. proteins, nucleic acids, carbohydrates or lipids, in which at least one component is a protein and the constituent parts function together." [GOC:dos, GOC:ma |
| 20 / 118 | "Any process that is carried out at the cellular level, but not necessarily restricted to a single cell. For example, cell communication occurs among more than one cell, but occurs at the cellular level." [GOC:go_curat |

Supplementary Data S3.1.2

### Timepoint 2

Molecular Function

| | |
|---|---|
| 133 / 597 | "Catalysis of a biochemical reaction at physiological temperatures. In biologically catalyzed reactions, the reactants are known as substrates, and the catalysts are naturally occurring macromolecular substances kn |
| 12 / 114 | "Binding to an RNA molecule or a portion thereof." [GOC:jl, GOC:mah] |
| 8 / 73 | "Binding to a methylated cytosine/guanine dinucleotide." [GOC:jl, PMID:11746232] |
| 7 / 66 | "Binding to a ribonucleotide, any compound consisting of a ribonucleoside that is esterified with (ortho)phosphate or an oligophosphate at any hydroxyl group on the ribose moiety." [GOC:mah] |
| 7 / 65 | "Binding to a nucleotide, any compound consisting of a nucleoside that is esterified with (ortho)phosphate or an oligophosphate at any hydroxyl group on the ribose or deoxyribose." [GOC:mah, ISBN:0198547684] |
| 7 / 64 | "Binding to nicotinamide adenine dinucleotide, a coenzyme involved in many redox and biosynthetic reactions; binding may be to either the oxidized form, NAD+, or the reduced form, NADH." [GOC:ai] |
| 7 / 63 | "Binding to a small molecule, any low molecular weight, monomeric, non-encoded molecule." [GOC:curators, GOC:pde, GOC:pm] |
| 7 / 63 | "Binding to FAD, flavin-adenine dinucleotide, the coenzyme or the prosthetic group of various flavoprotein oxidoreductase enzymes, in either the oxidized form, FAD, or the reduced form, FADH2." [GOC:ai, GOC:im |
| 7 / 63 | "Binding to nicotinamide-adenine dinucleotide phosphate, a coenzyme involved in many redox and biosynthetic reactions; binding may be to either the oxidized form, NADP+, or the reduced form, NADPH." [GOC:a |
| 9 / 73 | "Binding to a nucleic acid region that regulates a nucleic acid-based process. Such processes include transcription, DNA replication, and DNA repair." [GOC:txnOH] |

Biological Process

| | |
|---|---|
| 2 / 39 | "Any process that modulates the frequency, rate or extent of cellular DNA-templated transcription." [GOC:go_curators, GOC:txnOH] |
| 0 / 21 | "Any process that results in the conversion of one or more primary non-coding RNA (ncRNA) transcripts into one or more mature ncRNA molecules." [GOC:mah] |
| 3 / 3 | "The conversion of a single-cell organism from one mating type to another by the precise replacement of a DNA sequence at the expressed mating type locus with a copy of a sequence from a donor locus." [PMID:99 |
| 3 / 3 | "A process that forms a boundary that limits the spreading of heterochromatin along a chromosome." [GOC:mah] |
| 0 / 17 | "Any process involved in the maturation of a precursor 5.8S ribosomal RNA (rRNA) molecule into a mature 5.8S rRNA molecule." [GOC:curators] |
| 0 / 17 | "Any process involved in forming the mature 5' end of an rRNA molecule." [GOC:krc] |
| 0 / 16 | "Any phosphodiester bond hydrolysis involved in the conversion of a primary ribosomal RNA (rRNA) transcript into a mature rRNA molecule." [GOC:curators] |
| 0 / 16 | "The chemical reactions and pathways involving rRNA, ribosomal RNA, a structural constituent of ribosomes." [ISBN:0198506732] |
| 0 / 16 | "Any process involved in forming the mature 3' end of an rRNA molecule." [GOC:mah] |
| 2 / 30 | "Any process that modulates the frequency, rate or extent of transcription mediated by RNA polymerase I." [GOC:go_curators] |

Cellular Compartment

| | |
|---|---|
| 4 / 4 | "Any process that results in a change in state or activity of a cell (in terms of movement, secretion, enzyme production, gene expression, etc.) as a result of deprivation of iron ions." [GOC:mg] |
| 4 / 5 | "A region of a chromosome at which a DNA double-strand break has occurred. DNA damage signaling and repair proteins accumulate at the lesion to respond to the damage and repair the DNA to form a continuou |
| 4 / 5 | "A region of a chromosome at which DNA damage has occurred. DNA damage signaling and repair proteins accumulate at the lesion to respond to the damage and repair the DNA to form a continuous DNA helix." |
| 0 / 19 | "The chemical reactions and pathways involving an organic substance, any molecular entity containing carbon." [GOC:mah] |
| 3 / 3 | "Any process involved in the maintenance of an internal steady state of iron ions at the level of a cell." [GOC:ai, GOC:mah] |
| 16 / 107 | "The volume enclosed by the nuclear inner membrane." [GOC:mah, GOC:pz] |
| 10 / 25 | "A multisubunit guanine nucleotide exchange factor which catalyzes the exchange of GDP bound to initiation factor eIF2 for GTP, generating active eIF2-GTP. In humans, it is composed of five subunits, alpha, beta, |
| 0 / 15 | "The chemical reactions and pathways resulting in the formation of substances; typically the energy-requiring part of metabolism in which simpler substances are transformed into more complex ones." [GOC:curat |
| 9 / 23 | "A protein complex that possesses magnesium-dependent protein serine/threonine phosphatase (AMD phosphatase) activity, and consists of a catalytic subunit and one or more regulatory subunits that dictates th |
| 9 / 23 | "A complex consisting of methionyl- and glutamyl-tRNA synthetases. The tRNA synthetases present in the complex bind to their cognate tRNAs more efficiently than they do as monomers." [GOC:mcc, PMID:11069 |

Supplementary Data S3.1.3

### Timepoint 3

#### Molecular Function

| | |
|---|---|
| 133 / 597 | "Catalysis of a biochemical reaction at physiological temperatures. In biologically catalyzed reactions, the reactants are known as substrates, and the catalysts are naturally occurring macromolecular substances kn |
| 12 / 114 | "Binding to an RNA molecule or a portion thereof." [GOC:jl, GOC:mah] |
| 8 / 73 | "Binding to a methylated cytosine/guanine dinucleotide." [GOC:jl, PMID:11746232] |
| 7 / 65 | "Binding to a nucleotide, any compound consisting of a nucleoside that is esterified with (ortho)phosphate or an oligophosphate at any hydroxyl group on the ribose or deoxyribose." [GOC:mah, ISBN:0198547684] |
| 7 / 64 | "Binding to nicotinamide adenine dinucleotide, a coenzyme involved in many redox and biosynthetic reactions; binding may be to either the oxidized form, NAD+, or the reduced form, NADH." [GOC:ai] |
| 7 / 63 | "Binding to a small molecule, any low molecular weight, monomeric, non-encoded molecule." [GOC:curators, GOC:pde, GOC:pm] |
| 7 / 63 | "Binding to FAD, flavin-adenine dinucleotide, the coenzyme or the prosthetic group of various flavoprotein oxidoreductase enzymes, in either the oxidized form, FAD, or the reduced form, FADH2." [GOC:ai, GOC:im |
| 7 / 63 | "Binding to nicotinamide-adenine dinucleotide phosphate, a coenzyme involved in many redox and biosynthetic reactions; binding may be to either the oxidized form, NADP+, or the reduced form, NADPH." [GOC:a |
| 9 / 73 | "Binding to a nucleic acid region that regulates a nucleic acid-based process. Such processes include transcription, DNA replication, and DNA repair." [GOC:txnOH] |
| 9 / 73 | "Any selective and non-covalent interaction with a nucleic acid involved in the initiation, activation, perpetuation, repression or termination of polypeptide synthesis at the ribosome." [GOC:dph, GOC:tb, GOC:vw] |

#### Biological Process

| | |
|---|---|
| 2 / 39 | "Any process that modulates the frequency, rate or extent of cellular DNA-templated transcription." [GOC:go_curators, GOC:txnOH] |
| 0 / 21 | "Any process that results in the conversion of one or more primary non-coding RNA (ncRNA) transcripts into one or more mature ncRNA molecules." [GOC:mah] |
| 3 / 3 | "The conversion of a single-cell organism from one mating type to another by the precise replacement of a DNA sequence at the expressed mating type locus with a copy of a sequence from a donor locus." [PMID:99 |
| 3 / 3 | "A process that forms a boundary that limits the spreading of heterochromatin along a chromosome." [GOC:mah] |
| 0 / 17 | "Any process involved in the maturation of a precursor 5.8S ribosomal RNA (rRNA) molecule into a mature 5.8S rRNA molecule." [GOC:curators] |
| 0 / 17 | "Any process involved in forming the mature 5' end of an rRNA molecule." [GOC:krc] |
| 0 / 16 | "Any phosphodiester bond hydrolysis involved in the conversion of a primary ribosomal RNA (rRNA) transcript into a mature rRNA molecule." [GOC:curators] |
| 0 / 16 | "The chemical reactions and pathways involving rRNA, ribosomal RNA, a structural constituent of ribosomes." [ISBN:0198506732] |
| 0 / 16 | "Any process involved in forming the mature 3' end of an rRNA molecule." [GOC:mah] |
| 2 / 30 | "Any process that modulates the frequency, rate or extent of transcription mediated by RNA polymerase I." [GOC:go_curators] |

#### Cellular Compartment

| | |
|---|---|
| 4 / 4 | "Any process that results in a change in state or activity of a cell (in terms of movement, secretion, enzyme production, gene expression, etc.) as a result of deprivation of iron ions." [GOC:mg] |
| 0 / 19 | "The chemical reactions and pathways involving an organic substance, any molecular entity containing carbon." [GOC:mah] |
| 4 / 5 | "A region of a chromosome at which a DNA double-strand break has occurred. DNA damage signaling and repair proteins accumulate at the lesion to respond to the damage and repair the DNA to form a continuou |
| 4 / 5 | "A region of a chromosome at which DNA damage has occurred. DNA damage signaling and repair proteins accumulate at the lesion to respond to the damage and repair the DNA to form a continuous DNA helix." [ |
| 3 / 3 | "Any process involved in the maintenance of an internal steady state of iron ions at the level of a cell." [GOC:ai, GOC:mah] |
| 10 / 25 | "A multisubunit guanine nucleotide exchange factor which catalyzes the exchange of GDP bound to initiation factor eIF2 for GTP, generating active eIF2-GTP. In humans, it is composed of five subunits, alpha, beta, c |
| 0 / 15 | "The chemical reactions and pathways resulting in the formation of substances; typically the energy-requiring part of metabolism in which simpler substances are transformed into more complex ones." [GOC:curato |
| 19 / 118 | "Any process that is carried out at the cellular level, but not necessarily restricted to a single cell. For example, cell communication occurs among more than one cell, but occurs at the cellular level." [GOC:go_curato |
| 19 / 117 | "A biological process represents a specific objective that the organism is genetically programmed to achieve. Biological processes are often described by their outcome or ending state, e.g., the biological process of c |
| 19 / 117 | "The chemical reactions and pathways, including anabolism and catabolism, by which living organisms transform chemical substances. Metabolic processes typically transform small molecules, but also include macr |

Supplementary Data S3.1.4

### Timepoint 4

#### Molecular Function

| | |
|---|---|
| 14 / 114 | "Binding to an RNA molecule or a portion thereof." [GOC:jl, GOC:mah] |
| 8 / 73 | "Binding to a methylated cytosine/guanine dinucleotide." [GOC:jl, PMID:11746232] |
| 7 / 64 | "Binding to nicotinamide adenine dinucleotide, a coenzyme involved in many redox and biosynthetic reactions; binding may be to either the oxidized form, NAD+, or the reduced form, NADH." [GC |
| 7 / 63 | "Binding to a small molecule, any low molecular weight, monomeric, non-encoded molecule." [GOC:curators, GOC:pde, GOC:pm] |
| 7 / 63 | "Binding to FAD, flavin-adenine dinucleotide, the coenzyme or the prosthetic group of various flavoprotein oxidoreductase enzymes, in either the oxidized form, FAD, or the reduced form, FADH2.' |
| 7 / 62 | "Binding to a methylated cytosine/unspecified/guanine trinucleotide." [PMID:17239600] |
| 7 / 62 | "Binding to a methylated cytosine/unspecified/unspecified trinucleotide." [PMID:17239600] |
| 7 / 62 | "Binding to a purine nucleotide, a compound consisting of a purine nucleoside esterified with (ortho)phosphate." [GOC:ai] |
| 7 / 62 | "Binding to a pyrimidine nucleotide, a pyrimidine nucleoside esterified with (ortho)phosphate." [GOC:ai] |
| 7 / 62 | "Binding to a cyclic nucleotide, a nucleotide in which the phosphate group is in diester linkage to two positions on the sugar residue." [GOC:ai] |

#### Biological Process

| | |
|---|---|
| 6 / 9 | "The series of molecular signals generated as a consequence of the presence of unfolded proteins in the endoplasmic reticulum (ER) or other ER-related stress; results in changes in the regulation of transcri |
| 8 / 15 | "The cellular process in which a signal is conveyed to trigger a change in the activity or state of a cell. Signal transduction begins with reception of a signal (e.g. a ligand binding to a receptor or receptor activa |
| 3 / 39 | "Any process that modulates the frequency, rate or extent of cellular DNA-templated transcription." [GOC:go_curators, GOC:txnOH] |
| 3 / 3 | "The conversion of a single-cell organism from one mating type to another by the precise replacement of a DNA sequence at the expressed mating type locus with a copy of a sequence from a donor locus." [ |
| 3 / 3 | "A process that forms a boundary that limits the spreading of heterochromatin along a chromosome." [GOC:mah] |
| 3 / 3 | "Any process that modulates the frequency, rate or extent of filamentous growth of a population of unicellular organisms in response to biotic stimulus." [GOC:di, GOC:TermGenie] |
| 0 / 17 | "Any process involved in the maturation of a precursor 5.8S ribosomal RNA (rRNA) molecule into a mature 5.8S rRNA molecule." [GOC:curators] |
| 9 / 20 | "The process in which a signal is passed on to downstream components within the cell, which become activated themselves to further propagate the signal and finally trigger a change in the function or state |
| 5 / 8 | "The cascade of processes by which a signal interacts with a receptor, causing a change in the level or activity of a second messenger or other downstream target, and ultimately leading to the activation, per |
| 5 / 8 | "The series of molecular signals initiated by activation of a receptor on the surface of a cell. The pathway begins with binding of an extracellular ligand to a cell surface receptor, or for receptors that signal in |

#### Cellular Compartment

| | |
|---|---|
| 4 / 4 | "Any process that results in a change in state or activity of a cell (in terms of movement, secretion, enzyme production, gene expression, etc.) as a result of deprivation of iron ions." [GOC:mg] |
| 3 / 3 | "Any process involved in the maintenance of an internal steady state of iron ions at the level of a cell." [GOC:ai, GOC:mah] |
| 0 / 15 | "The chemical reactions and pathways resulting in the formation of substances; typically the energy-requiring part of metabolism in which simpler substances are transformed into more complex ones." [G |
| 19 / 118 | "Any process that is carried out at the cellular level, but not necessarily restricted to a single cell. For example, cell communication occurs among more than one cell, but occurs at the cellular level." [GOC: |
| 19 / 117 | "A biological process represents a specific objective that the organism is genetically programmed to achieve. Biological processes are often described by their outcome or ending state, e.g., the biological |
| 19 / 117 | "The chemical reactions and pathways, including anabolism and catabolism, by which living organisms transform chemical substances. Metabolic processes typically transform small molecules, but also inc |
| 3 / 4 | "An endocytic patch that consists of an actin-containing structure found at the plasma membrane in cells; formed of networks of branched actin filaments that lie just beneath the plasma membrane and a |
| 3 / 4 | "The portion of the actin cytoskeleton, comprising filamentous actin and associated proteins, that lies just beneath the plasma membrane." [GOC:mah] |
| 3 / 4 | "A complex that localizes to actin cortical patches at sites of endocytosis and negatively regulates barbed end F-actin assembly, resulting in the generation of free actin pools. The Aim21-Tda2 complex is n |
| 0 / 14 | "The chemical reactions and pathways involving organic or inorganic compounds that contain nitrogen." [GOC:jl, ISBN:0198506732] |

Supplementary Data S3.1.5

### Timepoint 5

#### Molecular Function

| | |
|---|---|
| 126 / 598 | "A molecular function characterized by the coupling of ATP hydrolysis to other steps of a reaction mechanism to make the reaction energetically favorable, for example to catalyze a reaction or drive transp |
| 13 / 114 | "Binding to an RNA molecule or a portion thereof." [GOC:jl, GOC:mah] |
| 10 / 20 | "Binds to and modulates the activity of a GTPase." [GOC:mah] |
| 5 / 7 | "Stimulates the exchange of GDP to GTP on a signaling GTPase, changing its conformation to its active form. Guanine nucleotide exchange factors (GEFs) act by stimulating the release of guanosine diphosp |
| 8 / 73 | "Binding to a methylated cytosine/guanine dinucleotide." [GOC:jl, PMID:11746232] |
| 7 / 64 | "Binding to nicotinamide adenine dinucleotide, a coenzyme involved in many redox and biosynthetic reactions; binding may be to either the oxidized form, NAD+, or the reduced form, NADH." [GOC:ai] |
| 7 / 63 | "Binding to a small molecule, any low molecular weight, monomeric, non-encoded molecule." [GOC:curators, GOC:pde, GOC:pm] |
| 7 / 63 | "Binding to FAD, flavin-adenine dinucleotide, the coenzyme or the prosthetic group of various flavoprotein oxidoreductase enzymes, in either the oxidized form, FAD, or the reduced form, FADH2." [GOC:a |
| 7 / 62 | "Binding to a methylated cytosine/unspecified/guanine trinucleotide." [PMID:17239600] |
| 7 / 62 | "Binding to a methylated cytosine/unspecified/unspecified trinucleotide." [PMID:17239600] |

#### Biological Process

| | |
|---|---|
| 11 / 15 | "The cellular process in which a signal is conveyed to trigger a change in the activity or state of a cell. Signal transduction begins with reception of a signal (e.g. a ligand binding to a receptor or receptor activa |
| 12 / 20 | "The process in which a signal is passed on to downstream components within the cell, which become activated themselves to further propagate the signal and finally trigger a change in the function or state |
| 7 / 9 | "The series of molecular signals generated as a consequence of the presence of unfolded proteins in the endoplasmic reticulum (ER) or other ER-related stress; results in changes in the regulation of transcri |
| 6 / 8 | "The cascade of processes by which a signal interacts with a receptor, causing a change in the level or activity of a second messenger or other downstream target, and ultimately leading to the activation, per |
| 6 / 8 | "The series of molecular signals initiated by activation of a receptor on the surface of a cell. The pathway begins with binding of an extracellular ligand to a cell surface receptor, or for receptors that signal in |
| 6 / 8 | "The series of molecular signals initiated by a ligand binding to its receptor, in which the activated receptor promotes the exchange of GDP for GTP on the alpha-subunit of an associated heterotrimeric G-pr |
| 6 / 8 | "The series of molecular signals generated as a consequence of a serotonin receptor binding to one of its physiological ligands." [GOC:mah] |
| 6 / 8 | "The series of molecular signals generated as a consequence of a dopamine receptor binding to one of its physiological ligands." [GOC:mah, PMID:21711983] |
| 6 / 8 | "The sequence of reactions within a cell required to convert absorbed photons into a molecular signal." [GOC:go_curators] |
| 6 / 8 | "The series of molecular signals mediated by the detection of a hormone." [GOC:sm] |

#### Cellular Compartment

| | |
|---|---|
| 4 / 4 | "Any process that results in a change in state or activity of a cell (in terms of movement, secretion, enzyme production, gene expression, etc.) as a result of deprivation of iron ions." [GOC:mg] |
| 4 / 5 | "The end, or tip, of a fungal hypha, where polarized growth occurs during hyphal elongation." [GOC:mcc] |
| 3 / 3 | "Any process involved in the maintenance of an internal steady state of iron ions at the level of a cell." [GOC:ai, GOC:mah] |
| 0 / 16 | "The chemical reactions and pathways by which individual cells transform chemical substances." [GOC:go_curators] |
| 18 / 118 | "Any process that is carried out at the cellular level, but not necessarily restricted to a single cell. For example, cell communication occurs among more than one cell, but occurs at the cellular level." [GOC:g |
| 18 / 118 | "The increase in size or mass of an entire organism, a part of an organism or a cell." [GOC:bf, GOC:ma] |
| 3 / 4 | "Any part of a cell where non-isotropic growth takes place." [GOC:mah] |
| 3 / 4 | "An endocytic patch that consists of an actin-containing structure found at the plasma membrane in cells; formed of networks of branched actin filaments that lie just beneath the plasma membrane and as |
| 3 / 4 | "The portion of the actin cytoskeleton, comprising filamentous actin and associated proteins, that lies just beneath the plasma membrane." [GOC:mah] |
| [illegible] | [illegible] |

Supplementary Data S3.1.6

### Timepoint 6

#### Molecular Function

| | |
|---|---|
| 132 / 598 | "A molecular function characterized by the coupling of ATP hydrolysis to other steps of a reaction mechanism to make the reaction energetically favorable, for example to catalyze a reaction or drive transp |
| 11 / 20 | "Binds to and modulates the activity of a GTPase." [GOC:mah] |
| 7 / 64 | "Binding to nicotinamide adenine dinucleotide, a coenzyme involved in many redox and biosynthetic reactions; binding may be to either the oxidized form, NAD+, or the reduced form, NADH." [GOC:ai] |
| 5 / 7 | "Stimulates the exchange of GDP to GTP on a signaling GTPase, changing its conformation to its active form. Guanine nucleotide exchange factors (GEFs) act by stimulating the release of guanosine diphosp |
| 7 / 63 | "Binding to a small molecule, any low molecular weight, monomeric, non-encoded molecule." [GOC:curators, GOC:pde, GOC:pm] |
| 7 / 63 | "Binding to FAD, flavin-adenine dinucleotide, the coenzyme or the prosthetic group of various flavoprotein oxidoreductase enzymes, in either the oxidized form, FAD, or the reduced form, FADH2." [GOC:a |
| 17 / 114 | "Binding to an RNA molecule or a portion thereof." [GOC:jl, GOC:mah] |
| 9 / 73 | "Binding to a methylated cytosine/guanine dinucleotide." [GOC:jl, PMID:11746232] |
| 7 / 62 | "Binding to a methylated cytosine/unspecified/guanine trinucleotide." [PMID:17239600] |
| 7 / 62 | "Binding to a methylated cytosine/unspecified/unspecified trinucleotide." [PMID:17239600] |

#### Biological Process

| | |
|---|---|
| 12 / 15 | "The cellular process in which a signal is conveyed to trigger a change in the activity or state of a cell. Signal transduction begins with reception of a signal (e.g. a ligand binding to a receptor or receptor activa |
| 13 / 20 | "The process in which a signal is passed on to downstream components within the cell, which become activated themselves to further propagate the signal and finally trigger a change in the function or state |
| 8 / 9 | "The series of molecular signals generated as a consequence of the presence of unfolded proteins in the endoplasmic reticulum (ER) or other ER-related stress; results in changes in the regulation of transcri |
| 7 / 8 | "The cascade of processes by which a signal interacts with a receptor, causing a change in the level or activity of a second messenger or other downstream target, and ultimately leading to the activation, per |
| 7 / 8 | "The series of molecular signals initiated by activation of a receptor on the surface of a cell. The pathway begins with binding of an extracellular ligand to a cell surface receptor, or for receptors that signal in |
| 7 / 8 | "The series of molecular signals initiated by a ligand binding to its receptor, in which the activated receptor promotes the exchange of GDP for GTP on the alpha-subunit of an associated heterotrimeric G-pr |
| 7 / 8 | "The series of molecular signals generated as a consequence of a serotonin receptor binding to one of its physiological ligands." [GOC:mah] |
| 7 / 8 | "The series of molecular signals generated as a consequence of a dopamine receptor binding to one of its physiological ligands." [GOC:mah, PMID:21711983] |
| 7 / 8 | "The sequence of reactions within a cell required to convert absorbed photons into a molecular signal." [GOC:go_curators] |
| 7 / 8 | "The series of molecular signals mediated by the detection of a hormone." [GOC:sm] |

#### Cellular Compartment

| | |
|---|---|
| 4 / 5 | "The end, or tip, of a fungal hypha, where polarized growth occurs during hyphal elongation." [GOC:mcc] |
| 0 / 16 | "The chemical reactions and pathways by which individual cells transform chemical substances." [GOC:go_curators] |
| 7 / 15 | "A lipid bilayer along with all the proteins and protein complexes embedded in it an attached to it." [GOC:dos, GOC:mah, ISBN:0815316194] |
| 0 / 15 | "The chemical reactions and pathways resulting in the formation of substances; typically the energy-requiring part of metabolism in which simpler substances are transformed into more complex ones." [GO |
| 0 / 15 | "The component of the organelle membrane consisting of the gene products having at least some part of their peptide sequence embedded in the hydrophobic region of the membrane." [GOC:dos, GOC:ma |
| 0 / 14 | "The chemical reactions and pathways involving organic or inorganic compounds that contain nitrogen." [GOC:jl, ISBN:0198506732] |
| 0 / 14 | "The chemical reactions and pathways resulting in many of the chemical changes of compounds that are not necessarily required for growth and maintenance of cells, and are often unique to a taxon. In mult |
| 3 / 4 | "The component of a plasma membrane consisting of gene products and protein complexes that are loosely bound to one of its surfaces, but not integrated into the hydrophobic region." [GOC:curators, GOC |
| 3 / 4 | "Any part of a cell where non-isotropic growth takes place." [GOC:mah] |
| [illegible] | [illegible] |

Supplementary Data S3.1.7

### Timepoint 7

#### Molecular Function

| | |
|---|---|
| 133 / 598 | "A molecular function characterized by the coupling of ATP hydrolysis to other steps of a reaction mechanism to make the reaction energetically favorable, for example to catalyze a reaction or drive transp |
| 12 / 20 | "Binds to and modulates the activity of a GTPase." [GOC:mah] |
| 7 / 64 | "Binding to nicotinamide adenine dinucleotide, a coenzyme involved in many redox and biosynthetic reactions; binding may be to either the oxidized form, NAD+, or the reduced form, NADH." [GOC:ai] |
| 5 / 7 | "Stimulates the exchange of GDP to GTP on a signaling GTPase, changing its conformation to its active form. Guanine nucleotide exchange factors (GEFs) act by stimulating the release of guanosine diphosp |
| 7 / 63 | "Binding to a small molecule, any low molecular weight, monomeric, non-encoded molecule." [GOC:curators, GOC:pde, GOC:pm] |
| 7 / 63 | "Binding to FAD, flavin-adenine dinucleotide, the coenzyme or the prosthetic group of various flavoprotein oxidoreductase enzymes, in either the oxidized form, FAD, or the reduced form, FADH2." [GOC:a |
| 9 / 73 | "Binding to a methylated cytosine/guanine dinucleotide." [GOC:jl, PMID:11746232] |
| 7 / 62 | "Binding to a methylated cytosine/unspecified/guanine trinucleotide." [PMID:17239600] |
| 7 / 62 | "Binding to a methylated cytosine/unspecified/unspecified trinucleotide." [PMID:17239600] |
| 7 / 62 | "Binding to a purine nucleotide, a compound consisting of a purine nucleoside esterified with (ortho)phosphate." [GOC:ai] |

#### Biological Process

| | |
|---|---|
| 12 / 15 | "The cellular process in which a signal is conveyed to trigger a change in the activity or state of a cell. Signal transduction begins with reception of a signal (e.g. a ligand binding to a receptor or receptor activa |
| 13 / 20 | "The process in which a signal is passed on to downstream components within the cell, which become activated themselves to further propagate the signal and finally trigger a change in the function or state |
| 8 / 9 | "The series of molecular signals generated as a consequence of the presence of unfolded proteins in the endoplasmic reticulum (ER) or other ER-related stress; results in changes in the regulation of transcri |
| 7 / 8 | "The cascade of processes by which a signal interacts with a receptor, causing a change in the level or activity of a second messenger or other downstream target, and ultimately leading to the activation, per |
| 7 / 8 | "The series of molecular signals initiated by activation of a receptor on the surface of a cell. The pathway begins with binding of an extracellular ligand to a cell surface receptor, or for receptors that signal in |
| 7 / 8 | "The series of molecular signals initiated by a ligand binding to its receptor, in which the activated receptor promotes the exchange of GDP for GTP on the alpha-subunit of an associated heterotrimeric G-pr |
| 7 / 8 | "The series of molecular signals generated as a consequence of a serotonin receptor binding to one of its physiological ligands." [GOC:mah] |
| 7 / 8 | "The series of molecular signals generated as a consequence of a dopamine receptor binding to one of its physiological ligands." [GOC:mah, PMID:21711983] |
| 7 / 8 | "The sequence of reactions within a cell required to convert absorbed photons into a molecular signal." [GOC:go_curators] |
| 7 / 8 | "The series of molecular signals mediated by the detection of a hormone." [GOC:sm] |

#### Cellular Compartment

| | |
|---|---|
| 4 / 4 | "An endocytic patch that consists of an actin-containing structure found at the plasma membrane in cells; formed of networks of branched actin filaments that lie just beneath the plasma membrane and ass |
| 4 / 4 | "The portion of the actin cytoskeleton, comprising filamentous actin and associated proteins, that lies just beneath the plasma membrane." [GOC:mah] |
| 4 / 4 | "A complex that localizes to actin cortical patches at sites of endocytosis and negatively regulates barbed end F-actin assembly, resulting in the generation of free actin pools. The Aim21-Tda2 complex is nec |
| 4 / 5 | "The end, or tip, of a fungal hypha, where polarized growth occurs during hyphal elongation." [GOC:mcc] |
| 0 / 16 | "The chemical reactions and pathways by which individual cells transform chemical substances." [GOC:go_curators] |
| 0 / 15 | "The chemical reactions and pathways resulting in the formation of substances; typically the energy-requiring part of metabolism in which simpler substances are transformed into more complex ones." [GO |
| 0 / 15 | "The component of the organelle membrane consisting of the gene products having at least some part of their peptide sequence embedded in the hydrophobic region of the membrane." [GOC:dos, GOC:ma |
| 7 / 15 | "A lipid bilayer along with all the proteins and protein complexes embedded in it an attached to it." [GOC:dos, GOC:mah, ISBN:0815316194] |
| 0 / 14 | "The chemical reactions and pathways involving organic or inorganic compounds that contain nitrogen." [GOC:jl, ISBN:0198506732] |
| 0 / 14 | "The chemical reactions and pathways resulting in many of the chemical changes of compounds that are not necessarily required for growth and maintenance of cells, and are often unique to a taxon. In mult |

Supplementary Data S3.1.8

### Timepoint 8

#### Molecular Function

| | |
|---|---|
| 121 / 597 | "Catalysis of a biochemical reaction at physiological temperatures. In biologically catalyzed reactions, the reactants are known as substrates, and the catalysts are naturally occurring macromolecular substa |
| 122 / 598 | "A molecular function characterized by the coupling of ATP hydrolysis to other steps of a reaction mechanism to make the reaction energetically favorable, for example to catalyze a reaction or drive transp |
| 10 / 20 | "Binds to and modulates the activity of a GTPase." [GOC:mah] |
| 116 / 586 | "A molecular process that can be carried out by the action of a single macromolecular machine, usually via direct physical interactions with other molecular entities. Function in this sense denotes an actior |
| 3 / 3 | "Binding to a histone H3 in which the lysine residue at position 27 has been modified by trimethylation." [GOC:dph, PMID:23948251] |
| 3 / 3 | "Binding to a histone H3 in which the lysine residue at position 9 has been modified by trimethylation." [PMID:30110338] |
| 3 / 3 | "Binding to a protein upon methylation of the target protein." [PMID:26060076] |
| 3 / 3 | "Binding to a histone H4 in which the lysine residue at position 20 has been modified by trimethylation." [PMID:22150589] |
| 24 / 79 | "Binding to a transition metal ions; a transition metal is an element whose atom has an incomplete d-subshell of extranuclear electrons, or which gives rise to a cation or cations with an incomplete d-subsl |
| 23 / 75 | "Binding to a zinc ion (Zn)." [GOC:ai] |

#### Biological Process

| | |
|---|---|
| 9 / 15 | "The cellular process in which a signal is conveyed to trigger a change in the activity or state of a cell. Signal transduction begins with reception of a signal (e.g. a ligand binding to a receptor or receptor activa |
| 10 / 20 | "The process in which a signal is passed on to downstream components within the cell, which become activated themselves to further propagate the signal and finally trigger a change in the function or state |
| 5 / 9 | "Any cellular process that results in the specification, formation or maintenance of polarized actin-based cytoskeletal structures." [GOC:mah] |
| 5 / 9 | "The series of molecular signals generated as a consequence of the presence of unfolded proteins in the endoplasmic reticulum (ER) or other ER-related stress; results in changes in the regulation of transcrip |
| 5 / 7 | "A process that is carried out at the cellular level which results in the assembly, arrangement of constituent parts, or disassembly of cytoskeletal structures." [GOC:dph, GOC:jl, GOC:mah] |
| 3 / 3 | "Any process that activates or increases the frequency, rate or extent of heterochromatin formation." [GOC:mah] |
| 3 / 3 | "Any process that modulates the direction of cell growth." [GOC:mah, GOC:vw] |
| 5 / 8 | "The cascade of processes by which a signal interacts with a receptor, causing a change in the level or activity of a second messenger or other downstream target, and ultimately leading to the activation, per |
| 5 / 8 | "The series of molecular signals initiated by activation of a receptor on the surface of a cell. The pathway begins with binding of an extracellular ligand to a cell surface receptor, or for receptors that signal in |
| 5 / 8 | "The series of molecular signals initiated by a ligand binding to its receptor, in which the activated receptor promotes the exchange of GDP for GTP on the alpha-subunit of an associated heterotrimeric G-prc |

#### Cellular Compartment

| | |
|---|---|
| 4 / 4 | "An endocytic patch that consists of an actin-containing structure found at the plasma membrane in cells; formed of networks of branched actin filaments that lie just beneath the plasma membrane and ass |
| 4 / 4 | "The portion of the actin cytoskeleton, comprising filamentous actin and associated proteins, that lies just beneath the plasma membrane." [GOC:mah] |
| 4 / 4 | "The region at the end of the longest axis of a cylindrical or elongated cell." [GOC:ai, GOC:mah] |
| 4 / 4 | "A complex that localizes to actin cortical patches at sites of endocytosis and negatively regulates barbed end F-actin assembly, resulting in the generation of free actin pools. The Aim21-Tda2 complex is nec |
| 4 / 5 | "The end, or tip, of a fungal hypha, where polarized growth occurs during hyphal elongation." [GOC:mcc] |
| 3 / 3 | "Structure within the hyphal tip of filamentous fungi that acts as an organizing center for hyphal tip growth; may function to supply vesicles to the elongating tip and/or to organize cytoskeletal microfilamen |
| 3 / 3 | "Protein complex that has a role in determining cell polarity, found at the tip of a growing fungal hypha." [PMID:15976451] |
| 0 / 16 | "The chemical reactions and pathways by which individual cells transform chemical substances." [GOC:go_curators] |
| 3 / 4 | "The component of a plasma membrane consisting of gene products and protein complexes that are loosely bound to one of its surfaces, but not integrated into the hydrophobic region." [GOC:curators, GOC |

Supplementary Data S3.1.9

### Timepoint 9

#### Molecular Function

| | |
|---|---|
| 109 / 598 | "A molecular function characterized by the coupling of ATP hydrolysis to other steps of a reaction mechanism to make the reaction energetically favorable, for example to catalyze a reaction or drive transp |
| 24 / 79 | "Binding to a transition metal ions; a transition metal is an element whose atom has an incomplete d-subshell of extranuclear electrons, or which gives rise to a cation or cations with an incomplete d-subsl |
| 23 / 75 | "Binding to a zinc ion (Zn)." [GOC:ai] |
| 23 / 75 | "Binding to and responding, e.g. by conformational change, to changes in the cellular level of zinc." [GOC:vw, PMID:31239353] |
| 3 / 3 | "Binding to a histone H3 in which the lysine residue at position 27 has been modified by trimethylation." [GOC:dph, PMID:23948251] |
| 3 / 3 | "Binding to a histone H3 in which the lysine residue at position 9 has been modified by trimethylation." [PMID:30110338] |
| 3 / 3 | "Binding to a protein upon methylation of the target protein." [PMID:26060076] |
| 3 / 3 | "Binding to a histone H4 in which the lysine residue at position 20 has been modified by trimethylation." [PMID:22150589] |
| 4 / 6 | "Catalysis of the formation of aminoacyl-tRNA from ATP, amino acid, and tRNA with the release of diphosphate and AMP." [ISBN:0198506732] |
| 8 / 20 | "Binds to and modulates the activity of a GTPase." [GOC:mah] |

#### Biological Process

| | |
|---|---|
| 6 / 9 | "Any cellular process that results in the specification, formation or maintenance of polarized actin-based cytoskeletal structures." [GOC:mah] |
| 0 / 24 | "The chemical reactions and pathways resulting in the breakdown of a protein or peptide by hydrolysis of its peptide bonds, initiated by the covalent attachment of ubiquitin, and mediated by the proteason |
| 5 / 7 | "A process that is carried out at the cellular level which results in the assembly, arrangement of constituent parts, or disassembly of cytoskeletal structures." [GOC:dph, GOC:jl, GOC:mah] |
| 4 / 5 | "The process of coupling alanine to alanyl-tRNA, catalyzed by alanyl-tRNA synthetase. The alanyl-tRNA synthetase is a class-II synthetases. The activated amino acid is transferred to the 3'-OH group of an ala |
| 4 / 5 | "The process of coupling arginine to arginyl-tRNA, catalyzed by arginyl-tRNA synthetase. The arginyl-tRNA synthetase is a class-I synthetase. The activated amino acid is transferred to the 2'-OH group of an a |
| 4 / 5 | "The process of coupling asparagine to asparaginyl-tRNA, catalyzed by asparaginyl-tRNA synthetase. The asparaginyl-tRNA synthetase is a class-II synthetase. The activated amino acid is transferred to the 3'- |
| 4 / 5 | "The process of coupling aspartate to aspartyl-tRNA, catalyzed by aspartyl-tRNA synthetase. The aspartyl-tRNA synthetase is a class-II synthetase. The activated amino acid is transferred to the 3'-OH group o |
| 4 / 5 | "The process of coupling cysteine to cysteinyl-tRNA, catalyzed by cysteinyl-tRNA synthetase. A cysteinyl-tRNA synthetase is a class-I synthetase. The activated amino acid is transferred to the 2'-OH group of |
| 4 / 5 | "The process of coupling glutamate to glutamyl-tRNA, catalyzed by glutamyl-tRNA synthetase. The glutamyl-tRNA synthetase is a class-I synthetase. The activated amino acid is transferred to the 2'-OH group |
| 4 / 5 | "The process of coupling glutamine to glutaminyl-tRNA, catalyzed by glutaminyl-tRNA synthetase. The glutaminyl-tRNA synthetase is a class-I synthetase. The activated amino acid is transferred to the 2'-OH |

#### Cellular Compartment

| | |
|---|---|
| 4 / 4 | "An endocytic patch that consists of an actin-containing structure found at the plasma membrane in cells; formed of networks of branched actin filaments that lie just beneath the plasma membrane and ass |
| 4 / 4 | "The portion of the actin cytoskeleton, comprising filamentous actin and associated proteins, that lies just beneath the plasma membrane." [GOC:mah] |
| 4 / 4 | "A complex that localizes to actin cortical patches at sites of endocytosis and negatively regulates barbed end F-actin assembly, resulting in the generation of free actin pools. The Aim21-Tda2 complex is nec |
| 16 / 54 | "A component of a cell contained within (but not including) the plasma membrane. In eukaryotes it includes the nucleus and cytoplasm." [ISBN:0198506732] |
| 3 / 4 | "The component of a plasma membrane consisting of gene products and protein complexes that are loosely bound to one of its surfaces, but not integrated into the hydrophobic region." [GOC:curators, GOC |
| 3 / 4 | "The region at the end of the longest axis of a cylindrical or elongated cell." [GOC:ai, GOC:mah] |
| 9 / 25 | "A complex of protein and RNA which facilitates translocation of proteins across membranes." [GOC:mlg] |
| 4 / 7 | "The part of the cell cortex consisting of an aggregation of proteins that will give rise to an endocytic vesicle." [GOC:dph, PMID:22949647] |
| 9 / 26 | "A ribonucleoprotein complex that contains the small ribosomal subunit, a translation initiation ternary complex (i.e. an initiator tRNA, GTP, and an IF2 or eIF2 complex), and an mRNA." [GOC:hjd, GOC:mah] |
| 0 / 16 | "The chemical reactions and pathways by which individual cells transform chemical substances." [GOC:go_curators] |

Supplementary Data S3.1.10

### Timepoint 10

#### Molecular Function

| | |
|---|---|
| 122 / 598 | "A molecular function characterized by the coupling of ATP hydrolysis to other steps of a reaction mechanism to make the reaction energetically favorable, for example to catalyze a reaction or drive transport agai |
| 26 / 79 | "Binding to a transition metal ions; a transition metal is an element whose atom has an incomplete d-subshell of extranuclear electrons, or which gives rise to a cation or cations with an incomplete d-subshell. Tra |
| 2 / 39 | "Catalysis of the reaction: a nucleoside triphosphate + H2O = nucleoside diphosphate + phosphate." [RHEA:23680] |
| 24 / 75 | "Binding to a zinc ion (Zn)." [GOC:ai] |
| 24 / 75 | "Binding to and responding, e.g. by conformational change, to changes in the cellular level of zinc." [GOC:vw, PMID:31239353] |
| 3 / 3 | "Binding to a histone H3 in which the lysine residue at position 27 has been modified by trimethylation." [GOC:dph, PMID:23948251] |
| 3 / 3 | "Binding to a histone H3 in which the lysine residue at position 9 has been modified by trimethylation." [PMID:30110338] |
| 3 / 3 | "Binding to a protein upon methylation of the target protein." [PMID:26060076] |
| 3 / 3 | "Binding to a histone H4 in which the lysine residue at position 20 has been modified by trimethylation." [PMID:22150589] |
| 120 / 587 | "The action of a molecule that contributes to the structural integrity of a complex or its assembly within or outside a cell." [GOC:mah, GOC:vw] |

#### Biological Process

| | |
|---|---|
| 6 / 9 | "Any cellular process that results in the specification, formation or maintenance of polarized actin-based cytoskeletal structures." [GOC:mah] |
| 5 / 7 | "A process that is carried out at the cellular level which results in the assembly, arrangement of constituent parts, or disassembly of cytoskeletal structures." [GOC:dph, GOC:jl, GOC:mah] |
| 3 / 3 | "Any process that modulates the direction of cell growth." [GOC:mah, GOC:vw] |
| 0 / 17 | "A cellular process that results in the biosynthesis of constituent macromolecules, assembly, and arrangement of constituent parts of a complex containing RNA and proteins. Includes the biosynthesis of the constitue |
| 1 / 24 | "The chemical reactions and pathways resulting in the breakdown of a protein or peptide by hydrolysis of its peptide bonds, initiated by the covalent attachment of ubiquitin, and mediated by the proteasome." [GOC |
| 4 / 6 | "The changes in the actin cytoskeleton that are necessary for the movement of cells along radial glial cells as a component of the process of cerebral cortex glial-mediated radial cell migration." [GO_REF:0000021, GC |
| 4 / 6 | "Any cellular process that depends upon or alters the actin cytoskeleton, that part of the cytoskeleton comprising actin filaments and their associated proteins." [GOC:mah] |
| 4 / 6 | "Covalent modification of an actin molecule." [GOC:mah] |
| 4 / 6 | "A process that is carried out at the cellular level which results in the assembly, arrangement of constituent parts, or disassembly of cytoskeletal structures containing both actin and myosin or paramyosin. The myosi |
| 4 / 6 | "A process that is carried out at the cellular level which results in the assembly, arrangement of constituent parts, or disassembly of a filament composed of myosin molecules." [GOC:mah] |

#### Cellular Compartment

| | |
|---|---|
| 4 / 4 | "An endocytic patch that consists of an actin-containing structure found at the plasma membrane in cells; formed of networks of branched actin filaments that lie just beneath the plasma membrane and assemble, |
| 4 / 4 | "The portion of the actin cytoskeleton, comprising filamentous actin and associated proteins, that lies just beneath the plasma membrane." [GOC:mah] |
| 4 / 4 | "A complex that localizes to actin cortical patches at sites of endocytosis and negatively regulates barbed end F-actin assembly, resulting in the generation of free actin pools. The Aim21-Tda2 complex is necessary fc |
| 18 / 54 | "A component of a cell contained within (but not including) the plasma membrane. In eukaryotes it includes the nucleus and cytoplasm." [ISBN:0198506732] |
| 11 / 32 | "Circular DNA structures that are not part of a chromosome." [GOC:ai] |
| 11 / 32 | "A membranous cellular structure that bears the photosynthetic pigments in plants, algae, and cyanobacteria. In cyanobacteria thylakoids are of various shapes and are attached to, or continuous with, the plasma m |
| 11 / 32 | "A discrete intracellular part formed of aggregated molecules such as proteins or other biopolymers." [GOC:mah, PMID:11121744] |
| 11 / 32 | "A multienzyme complex found in all multicellular eukaryotes composed of eight proteins with aminoacyl-tRNA synthetase activities (abbreviated as: ArgRS, AspRS, GluProRS, GlnRS, IleRS, LeuRS, LysRS, MetRS where |
| 11 / 32 | "A protein complex, that caps one or both ends of the proteasome core complex and regulates entry into, or exit from, the proteasome core complex." [GOC:mtg_sensu] |
| 11 / 32 | "A protein-pigment complex that may be closely or peripherally associated to photosynthetic reaction centers that participate in harvesting and transferring radiant energy to the reaction center." [GOC:lr] |

Supplementary Data S3.1.11

### Timepoint 11

#### Molecular Function

| | |
|---|---|
| 126 / 598 | "A molecular function characterized by the coupling of ATP hydrolysis to other steps of a reaction mechanism to make the reaction energetically favorable, for example to catalyze a reaction or drive transport agai |
| 2 / 39 | "Catalysis of the reaction: a nucleoside triphosphate + $H_2O$ = nucleoside diphosphate + phosphate." [RHEA:23680] |
| 25 / 79 | "Binding to a transition metal ions; a transition metal is an element whose atom has an incomplete d-subshell of extranuclear electrons, or which gives rise to a cation or cations with an incomplete d-subshell. Trai |
| 24 / 75 | "Binding to a zinc ion (Zn)." [GOC:ai] |
| 24 / 75 | "Binding to and responding, e.g. by conformational change, to changes in the cellular level of zinc." [GOC:vw, PMID:31239353] |
| 3 / 3 | "Binding to a histone H3 in which the lysine residue at position 27 has been modified by trimethylation." [GOC:dph, PMID:23948251] |
| 3 / 3 | "Binding to a histone H3 in which the lysine residue at position 9 has been modified by trimethylation." [PMID:30110338] |
| 3 / 3 | "Binding to a protein upon methylation of the target protein." [PMID:26060076] |
| 3 / 3 | "Binding to a histone H4 in which the lysine residue at position 20 has been modified by trimethylation." [PMID:22150589] |
| 2 / 31 | "Catalysis of the reaction: ATP + $H_2O$ = ADP + H+ Pi. ATP hydrolysis is used in some reactions as an energy source, for example to catalyze a reaction or drive transport against a concentration gradient." [RHEA:130 |

#### Biological Process

| | |
|---|---|
| 6 / 9 | "Any cellular process that results in the specification, formation or maintenance of polarized actin-based cytoskeletal structures." [GOC:mah] |
| 2 / 32 | "A cellular process that results in the biosynthesis of constituent macromolecules, assembly, and arrangement of constituent parts of ribosome subunits; includes transport to the sites of protein synthesis." [GOC:ma, |
| 5 / 7 | "A process that is carried out at the cellular level which results in the assembly, arrangement of constituent parts, or disassembly of cytoskeletal structures." [GOC:dph, GOC:jl, GOC:mah] |
| 0 / 17 | "A cellular process that results in the biosynthesis of constituent macromolecules, assembly, and arrangement of constituent parts of a complex containing RNA and proteins. Includes the biosynthesis of the constitue |
| 1 / 24 | "The chemical reactions and pathways resulting in the breakdown of a protein or peptide by hydrolysis of its peptide bonds, initiated by the covalent attachment of ubiquitin, and mediated by the proteasome." [GOC |
| 0 / 16 | "Endonucleolytic cleavage between the SSU-rRNA and the 5.8S rRNA of an rRNA molecule originally produced as a tricistronic rRNA transcript that contained the Small SubUnit (SSU) rRNA, the 5.8S rRNA, and the Lar |
| 1 / 23 | "Any process involved in the maturation of a precursor Large SubUnit (LSU) ribosomal RNA (rRNA) molecule into a mature LSU-rRNA molecule." [GOC:curators] |
| 2 / 28 | "Any cellular metabolic process involving deoxyribonucleic acid. This is one of the two main types of nucleic acid, consisting of a long, unbranched macromolecule formed from one, or more commonly, two, strands o |
| 1 / 21 | "The repair of double-strand breaks in DNA via homologous and nonhomologous mechanisms to reform a continuous DNA helix." [GOC:elh] |
| 1 / 21 | "The chemical reactions and pathways resulting in the breakdown of a protein or peptide by hydrolysis of its peptide bonds, initiated by the covalent attachment of a ubiquitin group, or multiple ubiquitin groups, to t |

#### Cellular Compartment

| | |
|---|---|
| 4 / 4 | "An endocytic patch that consists of an actin-containing structure found at the plasma membrane in cells; formed of networks of branched actin filaments that lie just beneath the plasma membrane and assemble, |
| 4 / 4 | "The portion of the actin cytoskeleton, comprising filamentous actin and associated proteins, that lies just beneath the plasma membrane." [GOC:mah] |
| 4 / 4 | "A complex that localizes to actin cortical patches at sites of endocytosis and negatively regulates barbed end F-actin assembly, resulting in the generation of free actin pools. The Aim21-Tda2 complex is necessary f |
| 15 / 107 | "The volume enclosed by the nuclear inner membrane." [GOC:mah, GOC:pz] |
| 14 / 97 | "Organized structure of distinctive morphology and function, bounded by a single or double lipid bilayer membrane and occurring within the cell. Includes the nucleus, mitochondria, plastids, vacuoles, and vesicles. I |
| 1 / 23 | "The component of a membrane consisting of the gene products having some covalently attached portion, for example part of a peptide sequence or some other covalently attached group such as a GPI anchor, whi |
| 0 / 15 | "The component of the organelle membrane consisting of the gene products having at least some part of their peptide sequence embedded in the hydrophobic region of the membrane." [GOC:dos, GOC:mah] |
| 3 / 4 | "The directed movement of inorganic anions into, out of or within a cell, or between cells, by means of some agent such as a transporter or pore. Inorganic anions are atoms or small molecules with a negative charg |
| 3 / 4 | "The component of a plasma membrane consisting of gene products and protein complexes that are loosely bound to one of its surfaces, but not integrated into the hydrophobic region." [GOC:curators, GOC:dos] |
| 4 / 7 | "The part of the cell cortex consisting of an aggregation of proteins that will give rise to an endocytic vesicle." [GOC:dph, PMID:22949647] |

Supplementary Data S3.1.12

### Timepoint 12

#### Molecular Function

| | |
|---|---|
| 129 / 597 | "Catalysis of a biochemical reaction at physiological temperatures. In biologically catalyzed reactions, the reactants are known as substrates, and the catalysts are naturally occurring macromolecular substances kno |
| 126 / 586 | "A molecular process that can be carried out by the action of a single macromolecular machine, usually via direct physical interactions with other molecular entities. Function in this sense denotes an action, or activ |
| 9 / 18 | "Binding to a lipid." [GOC:ai] |
| 9 / 18 | "Binding to a phosphatidylethanolamine, a class of glycerophospholipids in which a phosphatidyl group is esterified to the hydroxyl group of ethanolamine." [ISBN:0198506732] |
| 9 / 18 | "Binding to lysophosphatidic acid (LPA), a phospholipid derivative that acts as a potent mitogen due to its activation of high-affinity G protein-coupled receptors." [GOC:curators] |
| 9 / 18 | "Binding to phosphatidic acid, any of a class of glycerol phosphate in which both the remaining hydroxyl groups of the glycerol moiety are esterified with fatty acids." [CHEBI:16337, GOC:jp, ISBN:0198506732] |
| 9 / 18 | "Binding to phosphatidylglycerol." [GOC:kmv, GOC:TermGenie] |
| 9 / 18 | "Binding to ceramide 1-phosphate." [GOC:TermGenie, PMID:23863933] |
| 25 / 75 | "Binding to a zinc ion (Zn)." [GOC:ai] |
| 25 / 75 | "Binding to and responding, e.g. by conformational change, to changes in the cellular level of zinc." [GOC:vw, PMID:31239353] |

#### Biological Process

| | |
|---|---|
| 6 / 9 | "The series of molecular signals generated as a consequence of the presence of unfolded proteins in the endoplasmic reticulum (ER) or other ER-related stress; results in changes in the regulation of transcription and |
| 5 / 7 | "Any process that results in the specification, formation or maintenance of the physical structure of eukaryotic heterochromatin, a compact and highly condensed form of chromatin." [GOC:mah] |
| 0 / 16 | "Endonucleolytic cleavage between the SSU-rRNA and the 5.8S rRNA of an rRNA molecule originally produced as a tricistronic rRNA transcript that contained the Small SubUnit (SSU) rRNA, the 5.8S rRNA, and the Lar |
| 0 / 16 | "Any process involved in the maturation of a precursor Small SubUnit (SSU) ribosomal RNA (rRNA) molecule into a mature SSU-rRNA molecule from the pre-rRNA molecule originally produced as a tricistronic rRNA tr |
| 4 / 5 | "Any process that results in a change in state or activity of a cell or an organism (in terms of movement, secretion, enzyme production, gene expression, etc.) as a result of detection of, or exposure to, a hypotonic env |
| 3 / 3 | "Any process that modulates the frequency, rate or extent of any process in which a protein is transported to, or maintained in, a specific location." [GOC:dph, GOC:mah, GOC:tb] |
| 3 / 3 | "A process that forms a boundary that limits the spreading of heterochromatin along a chromosome." [GOC:mah] |
| 3 / 3 | "Any process that modulates the direction of cell growth." [GOC:mah, GOC:vw] |
| 3 / 3 | "Any process that modulates the frequency, rate or extent of protein localization to chromatin." [GO_REF:0000058, GOC:TermGenie, PMID:20889714] |
| 1 / 22 | "Any process involved in the conversion of a primary ribosomal RNA (rRNA) transcript into one or more mature rRNA molecules." [GOC:curators] |

#### Cellular Compartment

| | |
|---|---|
| 4 / 4 | "The directed movement of inorganic anions into, out of or within a cell, or between cells, by means of some agent such as a transporter or pore. Inorganic anions are atoms or small molecules with a negative charge |
| 4 / 4 | "An endocytic patch that consists of an actin-containing structure found at the plasma membrane in cells; formed of networks of branched actin filaments that lie just beneath the plasma membrane and assemble, n |
| 4 / 4 | "The portion of the actin cytoskeleton, comprising filamentous actin and associated proteins, that lies just beneath the plasma membrane." [GOC:mah] |
| 4 / 4 | "A complex that localizes to actin cortical patches at sites of endocytosis and negatively regulates barbed end F-actin assembly, resulting in the generation of free actin pools. The Aim21-Tda2 complex is necessary for |
| 4 / 5 | "The end, or tip, of a fungal hypha, where polarized growth occurs during hyphal elongation." [GOC:mcc] |
| 3 / 3 | "A compact and highly condensed form of chromatin that is refractory to transcription." [PMID:32017156] |
| 3 / 3 | "A membrane coat found on a coated vesicle." [GOC:mah] |
| 3 / 4 | "The component of a plasma membrane consisting of gene products and protein complexes that are loosely bound to one of its surfaces, but not integrated into the hydrophobic region." [GOC:curators, GOC:dos] |
| 3 / 4 | "Any part of a cell where non-isotropic growth takes place." [GOC:mah] |
| 3 / 4 | "The region at the end of the longest axis of a cylindrical or elongated cell." [GOC:ai, GOC:mah] |

#### Supplementary Data S3.2– Temporal GO Description of Volcano Analysis of Phosphoproteomic Data

Each one of the following pictures is an aggregation of outputs of a GO term enrichment analysis on the results of Volcano analysis at each specific point along the phosphoproteomic time course.

##### Supplementary Data S3.2.1

| GO_Term | P_val | Counts | MolFunc_Description |
|---|---|---|---|
| GO:0003824 | 0 | 38 / 825 | "Catalysis of a biochemical reaction at physiological temperatures. In biologically catalyzed reactions, the reactants are known as substrates, and the catalysts are naturally occurring macromolecular substance |
| GO:0004843 | 0.006745564 | 3 / 9 | "An thiol-dependent isopeptidase activity that cleaves ubiquitin from a target protein to which it is conjugated." [GOC:jh2, PMID:30783221] |
| GO:0061815 | 0.006745564 | 3 / 9 | "Catalysis of the hydrolysis of ubiquitin units from Met1-linked (or linear) polyubiquitin chains." [GOC:dph, PMID:26503766, PMID:27702987] |
| GO:0101005 | 0.006745564 | 3 / 9 | "An isopeptidase activity that cleaves ubiquitin from a target protein to which it is conjugated." [GOC:mec, PMID:30783221] |
| GO:0008234 | 0.00919957 | 3 / 10 | "Catalysis of the hydrolysis of peptide bonds in a polypeptide chain by a mechanism in which the sulfhydryl group of a cysteine residue at the active center acts as a nucleophile." [GOC:mah, https://www.ebi. |
| GO:0046914 | 0.009330992 | 1 / 124 | "Binding to a transition metal ions; a transition metal is an element whose atom has an incomplete d-subshell of extranuclear electrons, or which gives rise to a cation or cations with an incomplete d-subshell |
| GO:0008270 | 0.013112867 | 1 / 117 | "Binding to a zinc ion (Zn)." [GOC:ai] |
| GO:0003677 | 0.039001363 | 5 / 193 | "Any molecular function by which a gene product interacts selectively and non-covalently with DNA (deoxyribonucleic acid)." [GOC:dph, GOC:jl, GOC:tb, GOC:vw] |
| GO:0005515 | 0.047822674 | 4 / 32 | "Binding to a protein." [GOC:go_curators] |
| GO:0000403 | 0.048484848 | 1 / 1 | "Binding to a DNA segment shaped like a Y. This shape occurs when DNA contains a region of paired double-stranded DNA on one end and a region of unpaired DNA strands on the opposite end." [GOC:elh, PI |

| GO_Term | P_val | Counts | BioProc_Description |
|---|---|---|---|
| GO:0051056 | 0.00127099 | 4 / 11 | "Any process that modulates the frequency, rate or extent of small GTPase mediated signal transduction." [GOC:go_curators] |
| GO:0032012 | 0.003326913 | 3 / 7 | "Any process that modulates the frequency, rate or extent of ARF protein signal transduction." [GOC:mah] |
| GO:0048285 | 0.006915867 | 2 / 3 | "The creation of two or more organelles by division of one organelle." [GOC:jid] |
| GO:0051052 | 0.006915867 | 2 / 3 | "Any process that modulates the frequency, rate or extent of the chemical reactions and pathways involving DNA." [GOC:ai] |
| GO:0061389 | 0.006915867 | 2 / 3 | "Any process that modulates the direction of cell growth." [GOC:mah, GOC:vw] |
| GO:0035023 | 0.009883499 | 3 / 10 | "Any process that modulates the frequency, rate or extent of Rho protein signal transduction." [GOC:bf] |
| GO:0141124 | 0.012954317 | 3 / 11 | "An intracellular signaling module that is part of larger signaling pathways that can be initiated either intracellularly or by cell surface receptors. Intracellular signaling cassettes are discrete signaling units that |
| GO:0007006 | 0.013173864 | 2 / 4 | "A process that is carried out at the cellular level which results in the assembly, arrangement of constituent parts, or disassembly of a mitochondrial membrane, either of the lipid bilayer surrounding a mitoch |
| GO:0030382 | 0.013173864 | 2 / 4 | "A process that is carried out at the cellular level which results in the assembly, arrangement of constituent parts, or disassembly of sperm mitochondria; the process in which they take on their characteristic |
| GO:0033108 | 0.013173864 | 2 / 4 | "The aggregation, arrangement and bonding together of a set of components to form a mitochondrial respiratory chain complex." [GOC:mah] |

| GO_Term | P_val | Counts | CellComp_Description |
|---|---|---|---|
| GO:1901564 | 0.001587834 | 3 / 6 | "The chemical reactions and pathways involving organonitrogen compound." [GOC:pr, GOC:TermGenie] |
| GO:0006213 | 0.002059832 | 2 / 2 | "The chemical reactions and pathways involving any pyrimidine nucleoside, one of a family of organic molecules consisting of a pyrimidine base covalently bonded to ribose (a ribonucleoside) or deoxyribose ( |
| GO:0009116 | 0.002059832 | 2 / 2 | "The chemical reactions and pathways involving a nucleoside, a nucleobase linked to either beta-D-ribofuranose (a ribonucleoside) or 2-deoxy-beta-D-ribofuranose, (a deoxyribonucleoside), e.g. adenosine, gu |
| GO:0009119 | 0.002059832 | 2 / 2 | "The chemical reactions and pathways involving any ribonucleoside, a nucleoside in which purine or pyrimidine base is linked to a ribose (beta-D-ribofuranose) molecule." [GOC:jl] |
| GO:0009120 | 0.002059832 | 2 / 2 | "The chemical reactions and pathways involving any one of a family of organic molecules consisting of a purine or pyrimidine base covalently bonded to a sugar deoxyribose (a deoxyribonucleoside)." [GOC:jl, |
| GO:0009163 | 0.002059832 | 2 / 2 | "The chemical reactions and pathways resulting in the formation of any one of a family of organic molecules consisting of a purine or pyrimidine base covalently bonded to a sugar ribose (a ribonucleoside) or |
| GO:0009164 | 0.002059832 | 2 / 2 | "The chemical reactions and pathways resulting in the breakdown of any one of a family of organic molecules consisting of a purine or pyrimidine base covalently bonded to a sugar ribose (a ribonucleoside) o |
| GO:0042278 | 0.002059832 | 2 / 2 | "The chemical reactions and pathways involving one of a family of organic molecules consisting of a purine base covalently bonded to a sugar ribose (a ribonucleoside) or deoxyribose (a deoxyribonucleoside). |
| GO:0070637 | 0.002059832 | 2 / 2 | "The chemical reactions and pathways involving any pyridine nucleoside, a nucleoside in which a pyridine base covalently bonded to a sugar, usually ribose." [GOC:mah] |
| GO:1901072 | 0.002059832 | 2 / 2 | "The chemical reactions and pathways resulting in the breakdown of glucosamine-containing compounds (glucosamines)." [GOC:TermGenie] |

##### Supplementary Data S3.2.2

| GO_Term | P_val | Counts | MolFunc_Description |
|---|---|---|---|
| GO:0005096 | 7.04624E-05 | 5 / 15 | "Binds to and increases the activity of a GTPase, an enzyme that catalyzes the hydrolysis of GTP." [GOC:mah] |
| GO:0008047 | 0.000401464 | 5 / 21 | "Binds to and increases the activity of an enzyme." [GOC:dph, GOC:mah, GOC:tb] |
| GO:0008429 | 0.00264363 | 4 / 19 | "Binding to a phosphatidylethanolamine, a class of glycerophospholipids in which a phosphatidyl group is esterified to the hydroxyl group of ethanolamine." [ISBN:0198506732] |
| GO:0035727 | 0.00264363 | 4 / 19 | "Binding to lysophosphatidic acid (LPA), a phospholipid derivative that acts as a potent mitogen due to its activation of high-affinity G protein-coupled receptors." [GOC:curators] |
| GO:0070300 | 0.00264363 | 4 / 19 | "Binding to phosphatidic acid, any of a class of glycerol phosphate in which both the remaining hydroxyl groups of the glycerol moiety are esterified with fatty acids." [CHEBI:16337, GOC:jp, ISBN:0198506732] |
| GO:1901611 | 0.00264363 | 4 / 19 | "Binding to phosphatidylglycerol." [GOC:kmv, GOC:TermGenie] |
| GO:1902387 | 0.00264363 | 4 / 19 | "Binding to ceramide 1-phosphate." [GOC:TermGenie, PMID:23863933] |
| GO:0001786 | 0.00320614 | 4 / 20 | "Binding to phosphatidylserine, a class of glycophospholipids in which a phosphatidyl group is esterified to the hydroxyl group of L-serine." [ISBN:0198506732, PMID:12000961] |
| GO:0005544 | 0.00320614 | 4 / 20 | "Binding to a phospholipid, a class of lipids containing phosphoric acid as a mono- or diester, in the presence of calcium." [GOC:jl] |
| GO:0031210 | 0.00320614 | 4 / 20 | "Binding to a phosphatidylcholine, a glycophospholipid in which a phosphatidyl group is esterified to the hydroxyl group of choline." [GOC:mah, ISBN:0198506732] |

| GO_Term | P_val | Counts | BioProc_Description |
|---|---|---|---|
| GO:0032012 | 0.001510595 | 3 / 7 | "Any process that modulates the frequency, rate or extent of ARF protein signal transduction." [GOC:mah] |
| GO:0002764 | 0.00233321 | 3 / 8 | "The cascade of processes by which a signal interacts with a receptor, causing a change in the level or activity of a second messenger or other downstream target, and ultimately leading to the activation, perp |
| GO:0007166 | 0.00233321 | 3 / 8 | "The series of molecular signals initiated by activation of a receptor on the surface of a cell. The pathway begins with binding of an extracellular ligand to a cell surface receptor, or for receptors that signal in tl |
| GO:0007186 | 0.00233321 | 3 / 8 | "The series of molecular signals initiated by a ligand binding to its receptor, in which the activated receptor promotes the exchange of GDP for GTP on the alpha-subunit of an associated heterotrimeric G-prot |
| GO:0007210 | 0.00233321 | 3 / 8 | "The series of molecular signals generated as a consequence of a serotonin receptor binding to one of its physiological ligands." [GOC:mah] |
| GO:0007212 | 0.00233321 | 3 / 8 | "The series of molecular signals generated as a consequence of a dopamine receptor binding to one of its physiological ligands." [GOC:mah, PMID:21711983] |
| GO:0007602 | 0.00233321 | 3 / 8 | "The sequence of reactions within a cell required to convert absorbed photons into a molecular signal." [GOC:go_curators] |
| GO:0009755 | 0.00233321 | 3 / 8 | "The series of molecular signals mediated by the detection of a hormone." [GOC:sm] |
| GO:0009756 | 0.00233321 | 3 / 8 | "The series of molecular signals mediated by the detection of carbohydrate." [GOC:sm] |
| GO:0009863 | 0.00233321 | 3 / 8 | "The series of molecular signals mediated by salicylic acid." [GOC:jy] |

| GO_Term | P_val | Counts | CellComp_Description |
|---|---|---|---|
| GO:0031981 | 0.006964485 | 1 / 148 | "The volume enclosed by the nuclear inner membrane." [GOC:mah, GOC:pz] |
| GO:0140513 | 0.009829028 | 1 / 140 | "A stable assembly of two or more macromolecules, i.e. proteins, nucleic acids, carbohydrates or lipids, in which at least one component is a protein and the constituent parts function together in the nucleus |
| GO:0043231 | 0.012147807 | 1 / 135 | "Organized structure of distinctive morphology and function, bounded by a single or double lipid bilayer membrane and occurring within the cell. Includes the nucleus, mitochondria, plastids, vacuoles, and ve |
| GO:0000943 | 0.018398757 | 1 / 125 | "A complex of the retrotransposon RNA genome, reverse transcriptase, integrase, and associated molecules required for reproduction and integration of the retrotransposon into the host genome; the main s |
| GO:0005634 | 0.018398757 | 1 / 125 | "A membrane-bounded organelle of eukaryotic cells in which chromosomes are housed and replicated. In most cells, the nucleus contains all of the cell's chromosomes except the organellar chromosomes, ar |
| GO:0005880 | 0.018398757 | 1 / 125 | "Any microtubule in the nucleus of a cell." [GOC:mah] |
| GO:0031039 | 0.018398757 | 1 / 125 | "A membrane-bounded organelle of ciliated protozoan cells that contains polyploid copies of a portion of the cell's complete genome. Transcription of genes occurs in macronuclei. Some ciliate species may c |
| GO:0031040 | 0.018398757 | 1 / 125 | "A membrane-bounded organelle of ciliated protozoan cells that contains a diploid copy of the cell's complete genome. Sections of contiguous sequence in the macronucleus are often interrupted by internal |
| GO:0042405 | 0.018398757 | 1 / 125 | "An intranuclear focus at which aggregated proteins have been sequestered." [GOC:jl] |
| GO:0043073 | 0.018398757 | 1 / 125 | "The nucleus of a germ cell, a reproductive cell in multicellular organisms." [CL:0000586, GOC:go_curators] |

#### Supplementary Data S3.2.3

| GO_Term | P_val | Counts | MolFunc_Description |
|---|---|---|---|
| GO:0003824 | 0 | 33 / 825 | "Catalysis of a biochemical reaction at physiological temperatures. In biologically catalyzed reactions, the reactants are known as substrates, and the catalysts are naturally occurring macromolecular substance |
| GO:0005096 | 0.018350567 | 3 / 15 | "Binds to and increases the activity of a GTPase, an enzyme that catalyzes the hydrolysis of GTP." [GOC:mah] |
| GO:0000403 | 0.041212121 | 1 / 1 | "Binding to a DNA segment shaped like a Y. This shape occurs when DNA contains a region of paired double-stranded DNA on one end and a region of unpaired DNA strands on the opposite end." [GOC:elh, P |
| GO:0000405 | 0.041212121 | 1 / 1 | "Binding to DNA segment that contains a bubble. A bubble occurs when DNA contains a region of unpaired, single-stranded DNA flanked on both sides by regions of paired, double-stranded DNA." [GOC:elh, |
| GO:0000976 | 0.041212121 | 1 / 1 | "Binding to a specific sequence of DNA that is part of a regulatory region that controls transcription of that section of the DNA. The transcribed region might be described as a gene, cistron, or operon." [GOC: |
| GO:0001016 | 0.041212121 | 1 / 1 | "Binding to a DNA region that controls the transcription of a gene by RNA polymerase III. Binding may occur as a sequence specific interaction or as an interaction observed only once a factor has been recruit |
| GO:0001018 | 0.041212121 | 1 / 1 | "Binding to a DNA region that controls the transcription of the mitochondrial DNA." [GOC:txnOH, GOC:vw, PMID:20056105] |
| GO:0001019 | 0.041212121 | 1 / 1 | "Binding to a DNA region that controls transcription by a plastid RNA polymerase. Binding may occur as a sequence specific interaction or as an interaction observed only once a factor has been recruited to t |
| GO:0001046 | 0.041212121 | 1 / 1 | "Binding to a sequence of DNA that is part of a core promoter region. The core promoter is composed of the transcription start site and binding sites for the RNA polymerase and the basal transcription machi |
| GO:0001147 | 0.041212121 | 1 / 1 | "Binding to a sequence of DNA that promotes termination by RNA polymerase. The transcribed region might be described as a gene, cistron, or operon." [GOC:txnOH, PMID:18280161, PMID:18391175] |

| GO_Term | P_val | Counts | BioProc_Description |
|---|---|---|---|
| GO:0009987 | 0 | 37 / 844 | "Any process that is carried out at the cellular level, but not necessarily restricted to a single cell. For example, cell communication occurs among more than one cell, but occurs at the cellular level." [GOC:go_ |
| GO:0042539 | 0.002082947 | 2 / 2 | "Any process that results in a change in state or activity of a cell or an organism (in terms of movement, secretion, enzyme production, gene expression, etc.) as a result of detection of, or exposure to, a decre |
| GO:0046907 | 0.002588526 | 5 / 23 | "The directed movement of substances within a cell." [GOC:ai] |
| GO:1903468 | 0.005974248 | 2 / 3 | "Any process that activates or increases the frequency, rate or extent of DNA replication initiation involved in mitotic DNA replication." [GO_REF:0000058, GOC:mtg_cell_cycle, GOC:TermGenie, PMID:1234] |
| GO:0061951 | 0.011422818 | 2 / 4 | "The directed movement of a protein to a specific location in a plasma membrane." [GOC:dph, GOC:vw] |
| GO:0006888 | 0.013556648 | 3 / 12 | "The directed movement of substances from the endoplasmic reticulum (ER) to the Golgi, mediated by COP II vesicles. Small COP II coated vesicles form from the ER and then fuse directly with the cis-Golgi. L |
| GO:0000076 | 0.018199451 | 2 / 5 | "A signal transduction process that contributes to a DNA replication checkpoint, that prevents the initiation of nuclear division until DNA replication is complete, thereby ensuring that progeny inherit a full c |
| GO:0006890 | 0.018199451 | 2 / 5 | "The directed movement of substances from the Golgi back to the endoplasmic reticulum, mediated by vesicles bearing specific protein coats such as COPI or COG." [ISBN:0716731363, PMID:16510524] |
| GO:0006971 | 0.018199451 | 2 / 5 | "Any process that results in a change in state or activity of a cell or an organism (in terms of movement, secretion, enzyme production, gene expression, etc.) as a result of detection of, or exposure to, a hypo |
| GO:0033314 | 0.018199451 | 2 / 5 | "A signal transduction process that contributes to a mitotic DNA replication checkpoint." [GOC:mtg_cell_cycle] |

| GO_Term | P_val | Counts | CellComp_Description |
|---|---|---|---|
| GO:0009067 | 0.005662065 | 2 / 3 | "The chemical reactions and pathways resulting in the formation of amino acids of the aspartate family, comprising asparagine, aspartate, lysine, methionine and threonine." [GOC:ai] |
| GO:0008150 | 0.044691114 | 3 / 154 | "A biological process is the execution of a genetically-encoded biological module or program. It consists of all the steps required to achieve the specific biological objective of the module. A biological process |
| GO:0008152 | 0.044691114 | 3 / 154 | "The chemical reactions and pathways, including anabolism and catabolism, by which living organisms transform chemical substances. Metabolic processes typically transform small molecules, but also include |
| GO:0001100 | 0.044878049 | 1 / 1 | "Any process involved in the inhibition of progression from anaphase/telophase (high mitotic CDK activity) to G1 (low mitotic CDK activity)." [GOC:rn] |
| GO:0006044 | 0.044878049 | 1 / 1 | "The chemical reactions and pathways involving N-acetylglucosamine. The D isomer is a common structural unit of glycoproteins in plants, bacteria and animals; it is often the terminal sugar of an oligosacchar |
| GO:0006046 | 0.044878049 | 1 / 1 | "The chemical reactions and pathways resulting in the breakdown of N-acetylglucosamine. The D isomer is a common structural unit of glycoproteins in plants, bacteria and animals; it is often the terminal su |
| GO:0006528 | 0.044878049 | 1 / 1 | "The chemical reactions and pathways involving asparagine, 2-amino-3-carbamoylpropanoic acid." [GOC:go_curators] |
| GO:0006529 | 0.044878049 | 1 / 1 | "The chemical reactions and pathways resulting in the formation of asparagine, 2-amino-3-carbamoylpropanoic acid." [GOC:go_curators] |
| GO:0006553 | 0.044878049 | 1 / 1 | "The chemical reactions and pathways involving lysine, 2,6-diaminohexanoic acid." [GOC:go_curators] |
| GO:0009099 | 0.044878049 | 1 / 1 | "The chemical reactions and pathways resulting in the formation of valine, 2-amino-3-methylbutanoic acid." [GOC:ai] |

#### Supplementary Data S3.2.4

| GO_Term | P_val | Counts | MolFunc_Description |
|---|---|---|---|
| GO:0003824 | 0 | 36 / 825 | "Catalysis of a biochemical reaction at physiological temperatures. In biologically catalyzed reactions, the reactants are known as substrates, and the catalysts are naturally occurring macromolecular substance |
| GO:0016779 | 0.005933355 | 2 / 3 | "Catalysis of the transfer of a nucleotidyl group to a reactant." [ISBN:0198506732] |
| GO:0003674 | 0.015132917 | 34 / 802 | "A molecular process that can be carried out by the action of a single macromolecular machine, usually via direct physical interactions with other molecular entities. Function in this sense denotes an action, o |
| GO:0005096 | 0.024332542 | 3 / 15 | "Binds to and increases the activity of a GTPase, an enzyme that catalyzes the hydrolysis of GTP." [GOC:mah] |
| GO:0016301 | 0.025932821 | 2 / 6 | "Catalysis of the transfer of a phosphate group, usually from ATP, to a substrate molecule." [ISBN:0198506732] |
| GO:0005198 | 0.034543442 | 34 / 795 | "The action of a molecule that contributes to the structural integrity of a complex." [GOC:mah, GOC:vw] |
| GO:0016772 | 0.034710083 | 2 / 7 | "Catalysis of the transfer of a phosphorus-containing group from one compound (donor) to another (acceptor)." [GOC:jl, ISBN:0198506732] |
| GO:0016773 | 0.034710083 | 2 / 7 | "Catalysis of the transfer of a phosphorus-containing group from one compound (donor) to an alcohol group (acceptor)." [GOC:jl] |
| GO:0004673 | 0.046060606 | 1 / 1 | "Catalysis of the reaction: ATP + protein L-histidine = ADP + protein phospho-L-histidine." [EC:2.7.13.3, GOC:mah] |
| GO:0004712 | 0.046060606 | 1 / 1 | "Catalysis of the reactions: ATP + a protein serine = ADP + protein serine phosphate; ATP + a protein threonine = ADP + protein threonine phosphate; and ATP + a protein tyrosine = ADP + protein tyrosine pho |

| GO_Term | P_val | Counts | BioProc_Description |
|---|---|---|---|
| GO:0071396 | 0.005421648 | 3 / 8 | "Any process that results in a change in state or activity of a cell (in terms of movement, secretion, enzyme production, gene expression, etc.) as a result of a lipid stimulus." [GOC:mah] |
| GO:0001403 | 0.007244183 | 2 / 3 | "A growth pattern exhibited by budding haploid cells under certain growth conditions, in which cells retain the typical axial budding pattern of haploids, but become elongated and fail to separate after divisio |
| GO:0036267 | 0.007244183 | 2 / 3 | "The growth of colonies in filamentous chains of cells into a substrate." [GOC:di, PMID:22276126] |
| GO:0042144 | 0.007244183 | 2 / 3 | "The fusion of two vacuole membranes to form a single vacuole." [GOC:jl] |
| GO:0008104 | 0.007743358 | 3 / 9 | "Any process in which a protein is transported to, or maintained in, a specific location." [GOC:ai] |
| GO:0000282 | 0.013782036 | 2 / 4 | "The specification of the site where a daughter cell will form, in organisms that reproduce by budding. An example of this process is found in Saccharomyces cerevisiae." [GOC:mah] |
| GO:0000378 | 0.013782036 | 2 / 4 | "The RNA metabolic process that joins two exons, each of which has free ends that were generated by endonucleolytic cleavages, by a ligation reaction." [GOC:krc, ISBN:0879695897] |
| GO:0000959 | 0.013782036 | 2 / 4 | "The chemical reactions and pathways involving RNA transcribed from the mitochondrial genome and occurring in the mitochondrion." [GOC:krc, GOC:mah] |
| GO:0010501 | 0.013782036 | 2 / 4 | "The process in which a secondary structure of RNA are broken or 'melted'." [PMID:17169986] |
| GO:0034660 | 0.013782036 | 2 / 4 | "The chemical reactions and pathways involving non-coding RNA transcripts (ncRNAs)." [GOC:mah] |

| GO_Term | P_val | Counts | CellComp_Description |
|---|---|---|---|
| GO:0031981 | 0.001672108 | 1 / 148 | "The volume enclosed by the nuclear inner membrane." [GOC:mah, GOC:pz] |
| GO:0140513 | 0.00259759 | 1 / 140 | "A stable assembly of two or more macromolecules, i.e. proteins, nucleic acids, carbohydrates or lipids, in which at least one component is a protein and the constituent parts function together in the nucleus |
| GO:0043231 | 0.00340729 | 1 / 135 | "Organized structure of distinctive morphology and function, bounded by a single or double lipid bilayer membrane and occurring within the cell. Includes the nucleus, mitochondria, plastids, vacuoles, and ve |
| GO:0000943 | 0.00580691 | 1 / 125 | "A complex of the retrotransposon RNA genome, reverse transcriptase, integrase, and associated molecules required for reproduction and integration of the retrotransposon into the host genome; the main s |
| GO:0005634 | 0.00580691 | 1 / 125 | "A membrane-bounded organelle of eukaryotic cells in which chromosomes are housed and replicated. In most cells, the nucleus contains all of the cell's chromosomes except the organellar chromosomes, ar |
| GO:0005880 | 0.00580691 | 1 / 125 | "Any microtubule in the nucleus of a cell." [GOC:mah] |
| GO:0031039 | 0.00580691 | 1 / 125 | "A membrane-bounded organelle of ciliated protozoan cells that contains polyploid copies of a portion of the cell's complete genome. Transcription of genes occurs in macronuclei. Some ciliate species may c |
| GO:0031040 | 0.00580691 | 1 / 125 | "A membrane-bounded organelle of ciliated protozoan cells that contains a diploid copy of the cell's complete genome. Sections of contiguous sequence in the macronucleus are often interrupted by internal |
| GO:0042405 | 0.00580691 | 1 / 125 | "An intranuclear focus at which aggregated proteins have been sequestered." [GOC:jl] |
| GO:0043073 | 0.00580691 | 1 / 125 | "The nucleus of a germ cell, a reproductive cell in multicellular organisms." [CL:0000586, GOC:go_curators] |

#### Supplementary Data S3.2.5

| GO_Term | P_val | Counts | MolFunc_Description |
|---|---|---|---|
| GO:0003935 | 0.036363636 | 1 / 1 | "Catalysis of the reaction: GTP + 4 H2O = 2,5-diamino-6-hydroxy-4-(5-phosphoribosylamino)-pyrimidine + formate + 3 H+ + 2 phosphate." [EC:3.5.4.25, RHEA:23704] |
| GO:0004072 | 0.036363636 | 1 / 1 | "Catalysis of the reaction: L-aspartate + ATP = 4-phospho-L-aspartate + ADP + H+." [EC:2.7.2.4, RHEA:23776] |
| GO:0019202 | 0.036363636 | 1 / 1 | "Catalysis of the transfer of a phosphate group, usually from ATP, to an amino acid substrate." [GOC:jl] |
| GO:1990380 | 0.036363636 | 1 / 1 | "Hydrolysis of a ubiquitin unit from a ubiquitinated protein linked via the Lys48 residue of ubiquitin." [GOC:bf, GOC:PARL, PMID:22970133] |
| GO:0003912 | 0.070167696 | 1 / 2 | "Catalysis of the reaction: deoxynucleoside triphosphate + DNA(n) = diphosphate + DNA(n+1)." [EC:2.7.7.31] |
| GO:0003933 | 0.070167696 | 1 / 2 | "Catalysis of the hydrolysis of the imidazole ring of GTP, releasing formate. Two C-N bonds are hydrolyzed and the pentase unit is isomerized." [GOC:curators] |
| GO:0004103 | 0.070167696 | 1 / 2 | "Catalysis of the reaction: ATP + choline = ADP + choline phosphate + 2 H+." [EC:2.7.1.32, RHEA:12837] |
| GO:0004305 | 0.070167696 | 1 / 2 | "Catalysis of the reaction: ATP + ethanolamine = ADP + 2 H+ + phosphoethanolamine." [EC:2.7.1.82, RHEA:13069] |
| GO:0004654 | 0.070167696 | 1 / 2 | "Catalysis of the reaction: RNA(n+1) + phosphate = RNA(n) + a nucleoside diphosphate." [EC:2.7.7.8] |
| GO:0008195 | 0.070167696 | 1 / 2 | "Catalysis of the reaction: a 1,2-diacylglycerol 3-phosphate + H2O = a 1,2-diacyl-sn-glycerol + phosphate." [GOC:pr, PMID:25814022, RHEA:27429] |

| GO_Term | P_val | Counts | BioProc_Description |
|---|---|---|---|
| GO:0015031 | 0.001710648 | 4 / 14 | "The directed movement of proteins into, out of or within a cell, or between cells, by means of some agent such as a transporter or pore." [GOC:ai] |
| GO:0006890 | 0.014832795 | 2 / 5 | "The directed movement of substances from the Golgi back to the endoplasmic reticulum, mediated by vesicles bearing specific protein coats such as COPI or COG." [ISBN:0716731363, PMID:16510524] |
| GO:0048206 | 0.014832795 | 2 / 5 | "The process in which vesicles are directed to specific destination membranes during transport from the cis-Golgi to the rough ER." [GOC:jid, GOC:mah, ISBN:0716731363, PMID:10219233] |
| GO:1990669 | 0.014832795 | 2 / 5 | "The joining of the lipid bilayer membrane around an ERGIC-derived vesicle to the lipid bilayer membrane of the ER. Such vesicles include COPI-coated transport vesicles involved in retrograde transport." [GOC |
| GO:1990688 | 0.014832795 | 2 / 5 | "The joining of the lipid bilayer membrane around a Golgi vesicle to the lipid bilayer membrane of the ERGIC. Such vesicles include COPI-coated transport vesicles involved in retrograde transport." [GOC:bhm, P |
| GO:0070086 | 0.015531911 | 3 / 14 | "Endocytosis of a protein that requires the substrate to be modified by ubiquitination. Several plasma membrane proteins, including cell surface permeases and some receptors, are targeted for internalization |
| GO:0043328 | 0.022087172 | 3 / 16 | "The process of directing proteins towards the vacuole that contributes to protein catabolism via the multivesicular body (MVB) pathway." [GOC:jl, PMID:11511343] |
| GO:0043001 | 0.028745325 | 2 / 7 | "The directed movement of proteins from the Golgi to the plasma membrane in transport vesicles that move from the trans-Golgi network to the plasma membrane." [ISBN:0716731363] |
| GO:0046474 | 0.028745325 | 2 / 7 | "The chemical reactions and pathways resulting in the formation of glycerophospholipids, any derivative of glycerophosphate that contains at least one O-acyl, O-alkyl, or O-alkenyl group attached to the glycer |
| GO:0006611 | 0.036815996 | 2 / 8 | "The directed movement of a protein from the nucleus into the cytoplasm." [GOC:jl] |

| GO_Term | P_val | Counts | CellComp_Description |
|---|---|---|---|
| GO:0009987 | 0.011478007 | 1 / 153 | "Any process that is carried out at the cellular level, but not necessarily restricted to a single cell. For example, cell communication occurs among more than one cell, but occurs at the cellular level." [GOC:go_cu |
| GO:0040007 | 0.011921926 | 1 / 152 | "The increase in size or mass of an entire organism, a part of an organism or a cell." [GOC:bf, GOC:ma] |
| GO:0030120 | 0.012640969 | 2 / 5 | "A membrane coat found on a coated vesicle." [GOC:mah] |
| GO:0022414 | 0.014939954 | 1 / 146 | "A biological process that directly contributes to the process of producing new individuals by one or two organisms. The new individuals inherit some proportion of their genetic material from the parent or par |
| GO:0044419 | 0.015507258 | 1 / 145 | "Any process evolved to enable an interaction with an organism of a different species." [GOC:cc] |
| GO:0050896 | 0.015507258 | 1 / 145 | "Any process that results in a change in state or activity of a cell or an organism (in terms of movement, secretion, enzyme production, gene expression, etc.) as a result of a stimulus. The process begins with de |
| GO:0098754 | 0.015507258 | 1 / 145 | "Any process that reduces or removes the toxicity of a toxic substance. These may include transport of the toxic substance away from sensitive areas and to compartments or complexes whose purpose is sequ |
| GO:0000003 | 0.016094512 | 1 / 144 | "The production of new individuals that contain some portion of genetic material inherited from one or more parent organisms." [GOC:go_curators, GOC:isa_complete, GOC:jl, ISBN:0198506732] |
| GO:0002376 | 0.016094512 | 1 / 144 | "Any process involved in the development or functioning of the immune system, an organismal system for calibrated responses to potential internal or invasive threats." [GO_REF:0000022, GOC:add] |
| GO:0016032 | 0.016094512 | 1 / 144 | "A multi-organism process in which a virus is a participant. The other participant is the host. Includes infection of a host cell, replication of the viral genome, and assembly of progeny virus particles. In some cas |

#### Supplementary Data S3.2.7

| GO_Term | P_val | Counts | MolFunc_Description |
|---|---|---|---|
| GO:0003824 | 0 | 24 / 825 | "Catalysis of a biochemical reaction at physiological temperatures. In biologically catalyzed reactions, the reactants are known as substrates, and the catalysts are naturally occurring macromolecular substance |
| GO:0008429 | 0.002004887 | 4 / 19 | "Binding to a phosphatidylethanolamine, a class of glycerophospholipids in which a phosphatidyl group is esterified to the hydroxyl group of ethanolamine." [ISBN:0198506732] |
| GO:0035727 | 0.002004887 | 4 / 19 | "Binding to lysophosphatidic acid (LPA), a phospholipid derivative that acts as a potent mitogen due to its activation of high-affinity G protein-coupled receptors." [GOC:curators] |
| GO:0070300 | 0.002004887 | 4 / 19 | "Binding to phosphatidic acid, any of a class of glycerol phosphate in which both the remaining hydroxyl groups of the glycerol moiety are esterified with fatty acids." [CHEBI:16337, GOC:jp, ISBN:0198506732] |
| GO:1901611 | 0.002004887 | 4 / 19 | "Binding to phosphatidylglycerol." [GOC:kmv, GOC:TermGenie] |
| GO:1902387 | 0.002004887 | 4 / 19 | "Binding to ceramide 1-phosphate." [GOC:TermGenie, PMID:23863933] |
| GO:0001786 | 0.002437704 | 4 / 20 | "Binding to phosphatidylserine, a class of glycophospholipids in which a phosphatidyl group is esterified to the hydroxyl group of L-serine." [ISBN:0198506732, PMID:12000961] |
| GO:0005544 | 0.002437704 | 4 / 20 | "Binding to a phospholipid, a class of lipids containing phosphoric acid as a mono- or diester, in the presence of calcium." [GOC:jl] |
| GO:0031210 | 0.002437704 | 4 / 20 | "Binding to a phosphatidylcholine, a glycophospholipid in which a phosphatidyl group is esterified to the hydroxyl group of choline." [GOC:mah, ISBN:0198506732] |
| GO:0008289 | 0.002928985 | 4 / 21 | "Binding to a lipid." [GOC:ai] |

| GO_Term | P_val | Counts | BioProc_Description |
|---|---|---|---|
| GO:0009987 | 0 | 29 / 844 | "Any process that is carried out at the cellular level, but not necessarily restricted to a single cell. For example, cell communication occurs among more than one cell, but occurs at the cellular level." [GOC:go_ |
| GO:0061389 | 0.00354636 | 2 / 3 | "Any process that modulates the direction of cell growth." [GOC:mah, GOC:vw] |
| GO:0030950 | 0.003879839 | 3 / 10 | "Any cellular process that results in the specification, formation or maintenance of polarized actin-based cytoskeletal structures." [GOC:mah] |
| GO:0046907 | 0.006318908 | 4 / 23 | "The directed movement of substances within a cell." [GOC:ai] |
| GO:0030866 | 0.006659669 | 3 / 12 | "A process that is carried out at the cellular level which results in the assembly, arrangement of constituent parts, or disassembly of actin-based cytoskeletal structures in the cell cortex, i.e. just beneath the pl |
| GO:0098657 | 0.006659669 | 3 / 12 | "The directed movement of some substance from outside of a cell into a cell. This may occur via transport across the plasma membrane or via endocytosis." [GOC:dos] |
| GO:0000378 | 0.006856577 | 2 / 4 | "The RNA metabolic process that joins two exons, each of which has free ends that were generated by endonucleolytic cleavages, by a ligation reaction." [GOC:krc, ISBN:0879695897] |
| GO:0000959 | 0.006856577 | 2 / 4 | "The chemical reactions and pathways involving RNA transcribed from the mitochondrial genome and occurring in the mitochondrion." [GOC:krc, GOC:mah] |
| GO:0010501 | 0.006856577 | 2 / 4 | "The process in which a secondary structure of RNA are broken or 'melted'." [PMID:17169986] |
| GO:0030994 | 0.006856577 | 2 / 4 | "Dissolution of the primary septum during cell separation." [PMID:12665550] |

| GO_Term | P_val | Counts | CellComp_Description |
|---|---|---|---|
| GO:0000012 | 0.009512598 | 2 / 4 | "The repair of single strand breaks in DNA. Repair of such breaks is mediated by the same enzyme systems as are used in base excision repair." [PMID:18626472] |
| GO:0000725 | 0.009512598 | 2 / 4 | "A DNA repair process that involves the exchange, reciprocal or nonreciprocal, of genetic material between the broken DNA molecule and a homologous DNA region." [GOC:elh] |
| GO:0006284 | 0.009512598 | 2 / 4 | "In base excision repair, an altered base is removed by a DNA glycosylase enzyme, followed by excision of the resulting sugar phosphate. The small gap left in the DNA helix is filled in by the sequential action c |
| GO:0006289 | 0.009512598 | 2 / 4 | "A DNA repair process in which a small region of the strand surrounding the damage is removed from the DNA helix as an oligonucleotide. The small gap left in the DNA helix is filled in by the sequential action |
| GO:0006290 | 0.009512598 | 2 / 4 | "The repair of UV-induced T-T, C-T and C-C dimers." [ISBN:0815316194] |
| GO:0006298 | 0.009512598 | 2 / 4 | "A system for the correction of errors in which an incorrect base, which cannot form hydrogen bonds with the corresponding base in the parent strand, is incorporated into the daughter strand. The mismatch |
| GO:0006307 | 0.009512598 | 2 / 4 | "The repair of alkylation damage, e.g. the removal of the alkyl group at the O6-position of guanine by O6-alkylguanine-DNA alkyltransferase (AGT)." [PMID:10946226] |
| GO:0010213 | 0.009512598 | 2 / 4 | "A DNA repair process that is involved in repairing UV-induced DNA damage under non-photoreactivating conditions. The mechanism by which this repair process operates has not yet been completely elucid |
| GO:0036297 | 0.009512598 | 2 / 4 | "Removal of a DNA interstrand crosslink (a covalent attachment of DNA bases on opposite strands of the DNA) and restoration of the DNA. DNA interstrand crosslinks occur when both strands of duplex DNA a |
| GO:0043504 | 0.009512598 | 2 / 4 | "The process of restoring mitochondrial DNA after damage." [PMID:12565799, PMID:15189144, PMID:16050976] |

#### Supplementary Data S3.2.8

| GO_Term | P_val | Counts | MolFunc_Description |
|---|---|---|---|
| GO:0005096 | 0.013232923 | 3 / 15 | "Binds to and increases the activity of a GTPase, an enzyme that catalyzes the hydrolysis of GTP." [GOC:mah] |
| GO:0016301 | 0.016710394 | 2 / 6 | "Catalysis of the transfer of a phosphate group, usually from ATP, to a substrate molecule." [ISBN:0198506732] |
| GO:0016772 | 0.022594738 | 2 / 7 | "Catalysis of the transfer of a phosphorus-containing group from one compound (donor) to another (acceptor)." [GOC:jl, ISBN:0198506732] |
| GO:0016773 | 0.022594738 | 2 / 7 | "Catalysis of the transfer of a phosphorus-containing group from one compound (donor) to an alcohol group (acceptor)." [GOC:jl] |
| GO:0008047 | 0.031541176 | 3 / 21 | "Binds to and increases the activity of an enzyme." [GOC:dph, GOC:mah, GOC:tb] |
| GO:0004673 | 0.036363636 | 1 / 1 | "Catalysis of the reaction: ATP + protein L-histidine = ADP + protein phospho-L-histidine." [EC:2.7.13.3, GOC:mah] |
| GO:0004712 | 0.036363636 | 1 / 1 | "Catalysis of the reactions: ATP + a protein serine = ADP + protein serine phosphate; ATP + a protein threonine = ADP + protein threonine phosphate; and ATP + a protein tyrosine = ADP + protein tyrosine phosphate." [GC |
| GO:0004713 | 0.036363636 | 1 / 1 | "Catalysis of the reaction: ATP + a protein tyrosine = ADP + protein tyrosine phosphate." [RHEA:10596] |
| GO:0004740 | 0.036363636 | 1 / 1 | "Catalysis of the reaction: ATP + pyruvate dehydrogenase (acetyl-transferring) = ADP + pyruvate dehydrogenase (acetyl-transferring) phosphate." [EC:2.7.11.2] |
| GO:0008413 | 0.036363636 | 1 / 1 | "Catalysis of the reaction: 8-oxo-7,8-dihydroguanosine triphosphate (8-oxo-GTP) + H2O = 8-oxo-7,8-dihydroguanosine diphosphate (8-oxo-GDP) + phosphate. 8-oxo-7,8-dihydroguanosine triphosphate (8-oxo-GTP) is the |

| GO_Term | P_val | Counts | BioProc_Description |
|---|---|---|---|
| GO:0071396 | 0.00233321 | 3 / 8 | "Any process that results in a change in state or activity of a cell (in terms of movement, secretion, enzyme production, gene expression, etc.) as a result of a lipid stimulus." [GOC:mah] |
| GO:0006606 | 0.004658681 | 3 / 10 | "The directed movement of a protein from the cytoplasm to the nucleus." [GOC:jl] |
| GO:0000378 | 0.007779679 | 2 / 4 | "The RNA metabolic process that joins two exons, each of which has free ends that were generated by endonucleolytic cleavages, by a ligation reaction." [GOC:krc, ISBN:0879695897] |
| GO:0000959 | 0.007779679 | 2 / 4 | "The chemical reactions and pathways involving RNA transcribed from the mitochondrial genome and occurring in the mitochondrion." [GOC:krc, GOC:mah] |
| GO:0010501 | 0.007779679 | 2 / 4 | "The process in which a secondary structure of RNA are broken or 'melted'." [PMID:17169986] |
| GO:0034660 | 0.007779679 | 2 / 4 | "The chemical reactions and pathways involving non-coding RNA transcripts (ncRNAs)." [GOC:mah] |
| GO:0039703 | 0.007779679 | 2 / 4 | "The cellular metabolic process in which a cell duplicates one or more molecules of RNA." [GOC:bf, GOC:jl] |
| GO:0042245 | 0.007779679 | 2 / 4 | "Any process that results in the repair of damaged RNA." [PMID:11000254, PMID:11070075, UniProtKB-KW:KW-0692] |
| GO:0110154 | 0.007779679 | 2 / 4 | "Cleavage of the 5'-cap of an RNA." [GOC:sp, PMID:25533955, PMID:31101919] |
| GO:0006401 | 0.024411214 | 2 / 7 | "The chemical reactions and pathways resulting in the breakdown of RNA, ribonucleic acid, one of the two main type of nucleic acid, consisting of a long, unbranched macromolecule formed from ribonucleotides joined |

| GO_Term | P_val | Counts | CellComp_Description |
|---|---|---|---|
| GO:0008150 | 0.002549714 | 1 / 154 | "A biological process is the execution of a genetically-encoded biological module or program. It consists of all the steps required to achieve the specific biological objective of the module. A biological process is accompli |
| GO:0008152 | 0.002549714 | 1 / 154 | "The chemical reactions and pathways, including anabolism and catabolism, by which living organisms transform chemical substances. Metabolic processes typically transform small molecules, but also include macromc |
| GO:0009987 | 0.00268072 | 1 / 153 | "Any process that is carried out at the cellular level, but not necessarily restricted to a single cell. For example, cell communication occurs among more than one cell, but occurs at the cellular level." [GOC:go_curators, G |
| GO:0022414 | 0.003795161 | 1 / 146 | "A biological process that directly contributes to the process of producing new individuals by one or two organisms. The new individuals inherit some proportion of their genetic material from the parent or parents." [GC |
| GO:0044419 | 0.003986618 | 1 / 145 | "Any process evolved to enable an interaction with an organism of a different species." [GOC:cc] |
| GO:0050896 | 0.003986618 | 1 / 145 | "Any process that results in a change in state or activity of a cell or an organism (in terms of movement, secretion, enzyme production, gene expression, etc.) as a result of a stimulus. The process begins with detection o |
| GO:0098754 | 0.003986618 | 1 / 145 | "Any process that reduces or removes the toxicity of a toxic substance. These may include transport of the toxic substance away from sensitive areas and to compartments or complexes whose purpose is sequestration c |
| GO:0000003 | 0.004187261 | 1 / 144 | "The production of new individuals that contain some portion of genetic material inherited from one or more parent organisms." [GOC:go_curators, GOC:isa_complete, GOC:jl, ISBN:0198506732] |
| GO:0002376 | 0.004187261 | 1 / 144 | "Any process involved in the development or functioning of the immune system, an organismal system for calibrated responses to potential internal or invasive threats." [GO_REF:0000022, GOC:add] |
| GO:0016032 | 0.004187261 | 1 / 144 | "A multi-organism process in which a virus is a participant. The other participant is the host. Includes infection of a host cell, replication of the viral genome, and assembly of progeny virus particles. In some cases the vir |

#### Supplementary Data S3.2.9

| GO_Term | P_val | Counts | MolFunc_Description |
|---|---|---|---|
| GO:0008429 | 0.002309378 | 4 / 19 | "Binding to a phosphatidylethanolamine, a class of glycerophospholipids in which a phosphatidyl group is esterified to the hydroxyl group of ethanolamine." [ISBN:0198506732] |
| GO:0035727 | 0.002309378 | 4 / 19 | "Binding to lysophosphatidic acid (LPA), a phospholipid derivative that acts as a potent mitogen due to its activation of high-affinity G protein-coupled receptors." [GOC:curators] |
| GO:0070300 | 0.002309378 | 4 / 19 | "Binding to phosphatidic acid, any of a class of glycerol phosphate in which both the remaining hydroxyl groups of the glycerol moiety are esterified with fatty acids." [CHEBI:16337, GOC:jp, ISBN:0198506732] |
| GO:1901611 | 0.002309378 | 4 / 19 | "Binding to phosphatidylglycerol." [GOC:kmv, GOC:TermGenie] |
| GO:1902387 | 0.002309378 | 4 / 19 | "Binding to ceramide 1-phosphate." [GOC:TermGenie, PMID:23863933] |
| GO:0001786 | 0.002804347 | 4 / 20 | "Binding to phosphatidylserine, a class of glycophospholipids in which a phosphatidyl group is esterified to the hydroxyl group of L-serine." [ISBN:0198506732, PMID:12000961] |
| GO:0005544 | 0.002804347 | 4 / 20 | "Binding to a phospholipid, a class of lipids containing phosphoric acid as a mono- or diester, in the presence of calcium." [GOC:jl] |
| GO:0031210 | 0.002804347 | 4 / 20 | "Binding to a phosphatidylcholine, a glycophospholipid in which a phosphatidyl group is esterified to the hydroxyl group of choline." [GOC:mah, ISBN:0198506732] |
| GO:0008289 | 0.003365217 | 4 / 21 | "Binding to a lipid." [GOC:ai] |
| GO:0035091 | 0.003995381 | 4 / 22 | "Binding to an inositol-containing glycerophospholipid, i.e. phosphatidylinositol (PtdIns) and its phosphorylated derivatives." [GOC:bf, ISBN:0198506732, PMID:11395417] |

| GO_Term | P_val | Counts | BioProc_Description |
|---|---|---|---|
| GO:0030026 | 0.001141264 | 2 / 2 | "A homeostatic process involved in the maintenance of a steady state level of manganese ions within a cell." [GOC:mah] |
| GO:0055071 | 0.001141264 | 2 / 2 | "Any process involved in the maintenance of an internal steady state of manganese ions within an organism or cell." [GOC:jid, GOC:mah] |
| GO:1990659 | 0.001141264 | 2 / 2 | "The process of binding or confining manganese ions such that they are separated from other components of a biological system." [GOC:bhm, PMID:25417112] |
| GO:0030003 | 0.002544711 | 3 / 9 | "A homeostatic process involved in the maintenance of a steady state level of monoatomic cations within a cell. Monatomic cations (also called simple cations) are cations consisting of exactly one atom." [GOC |
| GO:0061389 | 0.003314002 | 2 / 3 | "Any process that modulates the direction of cell growth." [GOC:mah, GOC:vw] |
| GO:1903468 | 0.003314002 | 2 / 3 | "Any process that activates or increases the frequency, rate or extent of DNA replication initiation involved in mitotic DNA replication." [GO_REF:0000058, GOC:mtg_cell_cycle, GOC:TermGenie, PMID:1234] |
| GO:0001402 | 0.003498038 | 5 / 33 | "Relaying of environmental signals promoting filamentous growth." [GOC:mcc, PMID:9728395] |
| GO:0007015 | 0.006060629 | 3 / 12 | "A process that is carried out at the cellular level which results in the assembly, arrangement of constituent parts, or disassembly of cytoskeletal structures comprising actin filaments. Includes processes that cc |
| GO:0098662 | 0.006415215 | 2 / 4 | "A process in which an inorganic cation is transported from one side of a membrane to the other by means of some agent such as a transporter or pore." [GOC:mah] |
| GO:0030036 | 0.00941029 | 3 / 14 | "A process that is carried out at the cellular level which results in the assembly, arrangement of constituent parts, or disassembly of cytoskeletal structures comprising actin filaments and their associated protei |

| GO_Term | P_val | Counts | CellComp_Description |
|---|---|---|---|
| GO:0000041 | 0.03804878 | 1 / 1 | "The directed movement of transition metal ions into, out of or within a cell, or between cells, by means of some agent such as a transporter or pore. A transition metal is an element whose atom has an incom |
| GO:0001100 | 0.03804878 | 1 / 1 | "Any process involved in the inhibition of progression from anaphase/telophase (high mitotic CDK activity) to G1 (low mitotic CDK activity)." [GOC:rn] |
| GO:0005934 | 0.03804878 | 1 / 1 | "The end of a cellular bud distal to the site of attachment to the mother cell." [GOC:mah] |
| GO:0005935 | 0.03804878 | 1 / 1 | "The constriction between the mother cell and daughter cell (bud) in an organism that reproduces by budding." [GOC:mah] |
| GO:0006337 | 0.03804878 | 1 / 1 | "The controlled breakdown of nucleosomes, the beadlike structural units of eukaryotic chromatin composed of histones and DNA." [GOC:mah] |
| GO:0006825 | 0.03804878 | 1 / 1 | "The directed movement of copper (Cu) ions into, out of or within a cell, or between cells, by means of some agent such as a transporter or pore." [GOC:ai] |
| GO:0010876 | 0.03804878 | 1 / 1 | "Any process in which a lipid is transported to, or maintained in, a specific location." [GOC:BHF, GOC:dph, GOC:tb] |
| GO:0010877 | 0.03804878 | 1 / 1 | "The directed movement of lipids into cells that is part of their accumulation and maintenance." [GOC:BHF, GOC:dph, GOC:tb] |
| GO:0010878 | 0.03804878 | 1 / 1 | "The accumulation and maintenance in cells or tissues of cholesterol, cholest-5-en-3 beta-ol, the principal sterol of vertebrates and the precursor of many steroids, including bile acids and steroid hormones." [( |
| [illegible] | [illegible] | [illegible] | [illegible] |

#### Supplementary Data S3.2.10

| GO_Term | P_val | Counts | MolFunc_Description |
|---|---|---|---|
| GO:0003824 | 0 | 44 / 825 | "Catalysis of a biochemical reaction at physiological temperatures. In biologically catalyzed reactions, the reactants are known as substrates, and the catalysts are naturally occurring macromolecular substance |
| GO:0004815 | 0.025817397 | 2 / 5 | "Catalysis of the reaction: ATP + L-aspartate + tRNA(Asp) = AMP + diphosphate + L-aspartyl-tRNA(Asp)." [EC:6.1.1.12] |
| GO:0050560 | 0.025817397 | 2 / 5 | "Catalysis of the reaction: tRNA(Asx) + L-aspartate + ATP = aspartyl-tRNA(Asx) + diphosphate + AMP." [EC:6.1.1.23, MetaCyc:6.1.1.23-RXN] |
| GO:0003674 | 0.02748941 | 42 / 802 | "A molecular process that can be carried out by the action of a single macromolecular machine, usually via direct physical interactions with other molecular entities. Function in this sense denotes an action, o |
| GO:0005096 | 0.038698813 | 3 / 15 | "Binds to and increases the activity of a GTPase, an enzyme that catalyzes the hydrolysis of GTP." [GOC:mah] |
| GO:0004812 | 0.048550914 | 2 / 7 | "Catalysis of the formation of aminoacyl-tRNA from ATP, amino acid, and tRNA with the release of diphosphate and AMP." [ISBN:0198506732] |
| GO:0000976 | 0.055757576 | 1 / 1 | "Binding to a specific sequence of DNA that is part of a regulatory region that controls transcription of that section of the DNA. The transcribed region might be described as a gene, cistron, or operon." [GOC:t |
| GO:0001016 | 0.055757576 | 1 / 1 | "Binding to a DNA region that controls the transcription of a gene by RNA polymerase III. Binding may occur as a sequence specific interaction or as an interaction observed only once a factor has been recruite |
| GO:0001018 | 0.055757576 | 1 / 1 | "Binding to a DNA region that controls the transcription of the mitochondrial DNA." [GOC:txnOH, GOC:vw, PMID:20056105] |
| GO:0001019 | 0.055757576 | 1 / 1 | "Binding to a DNA region that controls transcription by a plastid RNA polymerase. Binding may occur as a sequence specific interaction or as an interaction observed only once a factor has been recruited to th |

| GO_Term | P_val | Counts | BioProc_Description |
|---|---|---|---|
| GO:0009987 | 0 | 43 / 844 | "Any process that is carried out at the cellular level, but not necessarily restricted to a single cell. For example, cell communication occurs among more than one cell, but occurs at the cellular level." [GOC:go_ |
| GO:0000281 | 0.002659201 | 2 / 2 | "A cell cycle process that results in the division of the cytoplasm of a cell after mitosis, resulting in the separation of the original cell into two daughter cells." [GOC:mtg_cell_cycle] |
| GO:0071396 | 0.005782034 | 3 / 8 | "Any process that results in a change in state or activity of a cell (in terms of movement, secretion, enzyme production, gene expression, etc.) as a result of a lipid stimulus." [GOC:mah] |
| GO:0000378 | 0.014402273 | 2 / 4 | "The RNA metabolic process that joins two exons, each of which has free ends that were generated by endonucleolytic cleavages, by a ligation reaction." [GOC:krc, ISBN:0879695897] |
| GO:0000425 | 0.014402273 | 2 / 4 | "The selective autophagy process in which a peroxisome is degraded by macroautophagy." [GOC:autophagy, PMID:12914914, PMID:16973210] |
| GO:0000959 | 0.014402273 | 2 / 4 | "The chemical reactions and pathways involving RNA transcribed from the mitochondrial genome and occurring in the mitochondrion." [GOC:krc, GOC:mah] |
| GO:0007006 | 0.014402273 | 2 / 4 | "A process that is carried out at the cellular level which results in the assembly, arrangement of constituent parts, or disassembly of a mitochondrial membrane, either of the lipid bilayer surrounding a mitoch |
| GO:0010501 | 0.014402273 | 2 / 4 | "The process in which a secondary structure of RNA are broken or 'melted'." [PMID:17169986] |
| GO:0016236 | 0.014402273 | 2 / 4 | "The autophagic process that proceeds via the formation of an autophagosome." [PMID:24366339] |
| GO:0030382 | 0.014402273 | 2 / 4 | "A process that is carried out at the cellular level which results in the assembly, arrangement of constituent parts, or disassembly of sperm mitochondria; the process in which they take on their characteristic |

| GO_Term | P_val | Counts | CellComp_Description |
|---|---|---|---|
| GO:0006554 | 0.003041159 | 2 / 2 | "The chemical reactions and pathways resulting in the breakdown of lysine, 2,6-diaminohexanoic acid." [GOC:go_curators] |
| GO:0046440 | 0.003041159 | 2 / 2 | "The chemical reactions and pathways involving L-lysine, the L-enantiomer of (S)-2,6-diaminohexanoic acid, i.e. (2S)-2,6-diaminohexanoic acid." [GOC:ai, GOC:jsg, GOC:mah] |
| GO:0046394 | 0.016336748 | 2 / 4 | "The chemical reactions and pathways resulting in the formation of carboxylic acids, any organic acid containing one or more carboxyl (-COOH) groups." [ISBN:0198506732] |
| GO:1901566 | 0.036558145 | 2 / 6 | "The chemical reactions and pathways resulting in the formation of organonitrogen compound." [GOC:pr, GOC:TermGenie] |
| GO:0140513 | 0.054712258 | 4 / 140 | "A stable assembly of two or more macromolecules, i.e. proteins, nucleic acids, carbohydrates or lipids, in which at least one component is a protein and the constituent parts function together in the nucleus |
| GO:0000745 | 0.055609756 | 1 / 1 | "The net movement of nuclei towards one another, leading to the bilateral transfer of genetic material in organisms undergoing conjugation without cellular fusion." [GOC:clt, GOC:mah] |
| GO:0006226 | 0.055609756 | 1 / 1 | "The chemical reactions and pathways resulting in the formation of dUMP, deoxyuridine monophosphate (2'-deoxyuridine 5'-phosphate)." [ISBN:0198506732] |
| GO:0006553 | 0.055609756 | 1 / 1 | "The chemical reactions and pathways involving lysine, 2,6-diaminohexanoic acid." [GOC:go_curators] |
| GO:0006558 | 0.055609756 | 1 / 1 | "The chemical reactions and pathways involving L-phenylalanine, the L-enantiomer of 2-amino-3-phenylpropanoic acid, i.e. (2S)-2-amino-3-phenylpropanoic acid." [GOC:jsg, GOC:mah] |
| GO:0006568 | 0.055609756 | 1 / 1 | "The chemical reactions and pathways involving tryptophan, the chiral amino acid 2-amino-3-(1H-indol-3-yl)propanoic acid." [ISBN:0198547684] |

#### Supplementary Data S3.2.11

| GO_Term | P_val | Counts | MolFunc_Description |
|---|---|---|---|
| GO:0003824 | 0 | 39 / 825 | "Catalysis of a biochemical reaction at physiological temperatures. In biologically catalyzed reactions, the reactants are known as substrates, and the catalysts are naturally occurring macromolecular substances |
| GO:0005096 | 0.003963102 | 4 / 15 | "Binds to and increases the activity of a GTPase, an enzyme that catalyzes the hydrolysis of GTP." [GOC:mah] |
| GO:0008047 | 0.013216808 | 4 / 21 | "Binds to and increases the activity of an enzyme." [GOC:dph, GOC:mah, GOC:tb] |
| GO:0030695 | 0.01784169 | 4 / 23 | "Binds to and modulates the activity of a GTPase." [GOC:mah] |
| GO:0004839 | 0.048484848 | 1 / 1 | "Catalysis of the reaction: E1 + ubiquitin + ATP--> E1-ubiquitin + AMP + PPi, where the E1-ubiquitin linkage is a thioester bond between the C-terminal glycine of Ub and a sulfhydryl side group of an E1 cysteine |
| GO:0005046 | 0.048484848 | 1 / 1 | "Binding to a KDEL sequence, the C terminus tetrapeptide sequence Lys-Asp-Glu-Leu found in proteins that are to be retained in the endoplasmic reticulum." [GOC:ai] |
| GO:0005338 | 0.048484848 | 1 / 1 | "Enables the transfer of a nucleotide-sugar from one side of a membrane to the other. A nucleotide-sugar is any nucleotide in which the distal phosphoric residue of a nucleoside 5'-diphosphate is in glycosidic |
| GO:0005456 | 0.048484848 | 1 / 1 | "Enables the transfer of a CMP-N-acetylneuraminate from one side of a membrane to the other." [GOC:ai, GOC:mtg_transport, ISBN:0815340729] |
| GO:0005460 | 0.048484848 | 1 / 1 | "Enables the transfer of a UDP-glucose from one side of a membrane to the other. UDP-glucose is a substance composed of glucose in glycosidic linkage with uridine diphosphate." [GOC:ai, GOC:mtg_transport |
| GO:0005461 | 0.048484848 | 1 / 1 | "Enables the transfer of a UDP-glucuronic acid from one side of a membrane to the other. UDP-glucuronic acid is a substance composed of glucuronic acid in glycosidic linkage with uridine diphosphate." [GOC: |

| GO_Term | P_val | Counts | BioProc_Description |
|---|---|---|---|
| GO:0007165 | 0.009547765 | 4 / 20 | "The cellular process in which a signal is conveyed to trigger a change in the activity or state of a cell. Signal transduction begins with reception of a signal (e.g. a ligand binding to a receptor or receptor activati |
| GO:0141124 | 0.010626745 | 3 / 11 | "An intracellular signaling module that is part of larger signaling pathways that can be initiated either intracellularly or by cell surface receptors. Intracellular signaling cassettes are discrete signaling units that a |
| GO:0000378 | 0.011422818 | 2 / 4 | "The RNA metabolic process that joins two exons, each of which has free ends that were generated by endonucleolytic cleavages, by a ligation reaction." [GOC:krc, ISBN:0879695897] |
| GO:0000959 | 0.011422818 | 2 / 4 | "The chemical reactions and pathways involving RNA transcribed from the mitochondrial genome and occurring in the mitochondrion." [GOC:krc, GOC:mah] |
| GO:0010501 | 0.011422818 | 2 / 4 | "The process in which a secondary structure of RNA are broken or 'melted'." [PMID:17169986] |
| GO:0034660 | 0.011422818 | 2 / 4 | "The chemical reactions and pathways involving non-coding RNA transcripts (ncRNAs)." [GOC:mah] |
| GO:0039703 | 0.011422818 | 2 / 4 | "The cellular metabolic process in which a cell duplicates one or more molecules of RNA." [GOC:bf, GOC:jl] |
| GO:0042245 | 0.011422818 | 2 / 4 | "Any process that results in the repair of damaged RNA." [PMID:11000254, PMID:11070075, UniProtKB-KW:KW-0692] |
| GO:0110154 | 0.011422818 | 2 / 4 | "Cleavage of the 5'-cap of an RNA." [GOC:sp, PMID:25533955, PMID:31101919] |
| GO:0006890 | 0.018199451 | 2 / 5 | "The directed movement of substances from the Golgi back to the endoplasmic reticulum, mediated by vesicles bearing specific protein coats such as COPI or COG." [ISBN:0716731363, PMID:16510524] |

| GO_Term | P_val | Counts | CellComp_Description |
|---|---|---|---|
| GO:0061645 | 0.001202535 | 4 / 11 | "The part of the cell cortex consisting of an aggregation of proteins that will give rise to an endocytic vesicle." [GOC:dph, PMID:22949647] |
| GO:0030479 | 0.004844612 | 3 / 8 | "An endocytic patch that consists of an actin-containing structure found at the plasma membrane in cells; formed of networks of branched actin filaments that lie just beneath the plasma membrane and assen |
| GO:0030864 | 0.004844612 | 3 / 8 | "The portion of the actin cytoskeleton, comprising filamentous actin and associated proteins, that lies just beneath the plasma membrane." [GOC:mah] |
| GO:0110131 | 0.004844612 | 3 / 8 | "A complex that localizes to actin cortical patches at sites of endocytosis and negatively regulates barbed end F-actin assembly, resulting in the generation of free actin pools. The Aim21-Tda2 complex is necess |
| GO:0000781 | 0.048780488 | 1 / 1 | "The end of a linear chromosome, required for the integrity and maintenance of the end. A chromosome telomere usually includes a region of telomerase-encoded repeats the length of which rarely exceeds 2 |
| GO:0001171 | 0.048780488 | 1 / 1 | "A DNA synthesis process that uses RNA as the initial template for synthesis of DNA, but which also includes an RNase activity to remove the RNA strand of an RNA-DNA heteroduplex produced by the RNA-dep |
| GO:0001666 | 0.048780488 | 1 / 1 | "Any process that results in a change in state or activity of a cell or an organism (in terms of movement, secretion, enzyme production, gene expression, etc.) as a result of a stimulus indicating lowered oxygen |
| GO:0002931 | 0.048780488 | 1 / 1 | "Any process that results in a change in state or activity of an organism (in terms of movement, secretion, enzyme production, gene expression, etc.) as a result of a inadequate blood supply." [GOC:hjd] |
| GO:0003299 | 0.048780488 | 1 / 1 | "The enlargement or overgrowth of all or part of a muscle organ or tissue due to an increase in the size of its muscle cells as a result of a disturbance in organismal or cellular homeostasis." [GOC:mtg_heart] |
| GO:0006044 | 0.048780488 | 1 / 1 | "The chemical reactions and pathways involving N-acetylglucosamine. The D isomer is a common structural unit of glycoproteins in plants, bacteria and animals; it is often the terminal sugar of an oligosacchari |
| GO:0006046 | 0.048780488 | 1 / 1 | "The chemical reactions and pathways resulting in the breakdown of N-acetylglucosamine. The D isomer is a common structural unit of glycoproteins in plants, bacteria and animals; it is often the terminal suga |

Supplementary Data S3.2.12

| GO_Term | P_val | Counts | MolFunc_Description |
|---|---|---|---|
| GO:0005096 | 0.007533272 | 4 / 15 | "Binds to and increases the activity of a GTPase, an enzyme that catalyzes the hydrolysis of GTP." [GOC:mah] |
| GO:0005545 | 0.017746809 | 2 / 4 | "Binding to a phosphatidylinositol, a glycophospholipid with its sn-glycerol 3-phosphate residue is esterified to the 1-hydroxyl group of 1D-myo-inositol." [ISBN:0198506732] |
| GO:0034235 | 0.017746809 | 2 / 4 | "Binding to a glycosylphosphatidylinositol anchor. GPI anchors serve to attach membrane proteins to the lipid bilayer of cell membranes." [GOC:vw] |
| GO:0008047 | 0.023600082 | 4 / 21 | "Binds to and increases the activity of an enzyme." [GOC:dph, GOC:mah, GOC:tb] |
| GO:0035091 | 0.027265987 | 4 / 22 | "Binding to an inositol-containing glycerophospholipid, i.e. phosphatidylinositol (PtdIns) and its phosphorylated derivatives." [GOC:bf, ISBN:0198506732, PMID:11395417] |
| GO:0030695 | 0.031197636 | 4 / 23 | "Binds to and modulates the activity of a GTPase." [GOC:mah] |
| GO:0001217 | 0.036204346 | 3 / 14 | "A DNA-binding transcription factor activity that represses or decreases the transcription of specific gene sets." [GOC:txnOH-2018] |
| GO:0016301 | 0.03953174 | 2 / 6 | "Catalysis of the transfer of a phosphate group, usually from ATP, to a substrate molecule." [ISBN:0198506732] |
| GO:0001216 | 0.042744342 | 3 / 15 | "A DNA-binding transcription factor activity that activates or increases transcription of specific gene sets." [GOC:txnOH-2018] |
| GO:0005543 | 0.044459475 | 4 / 26 | "Binding to a phospholipid, a class of lipids containing phosphoric acid as a mono- or diester." [ISBN:0198506732] |

| GO_Term | P_val | Counts | BioProc_Description |
|---|---|---|---|
| GO:0006984 | 0.001123911 | 4 / 9 | "The series of molecular signals that conveys information from the endoplasmic reticulum to the nucleus, usually resulting in a change in transcriptional regulation." [GOC:mah] |
| GO:0006996 | 0.001123911 | 4 / 9 | "A process that is carried out at the cellular level which results in the assembly, arrangement of constituent parts, or disassembly of an organelle within a cell. An organelle is an organized structure of distincti |
| GO:0097411 | 0.001767748 | 4 / 10 | "The series of molecular signals mediated by hypoxia-inducible factor (HIF1) in response to lowered oxygen levels (hypoxia). Under hypoxic conditions, the oxygen-sensitive alpha-subunit of hypoxia-inducible |
| GO:0006914 | 0.003496405 | 3 / 6 | "The cellular catabolic process in which cells digest cellular materials, such as organelles and other macromolecular constituents, or non-self materials such as intracellular pathogens. Autophagy serves to pro |
| GO:0097576 | 0.003496405 | 3 / 6 | "Merging of two or more vacuoles, or of vacuoles and vesicles within a cell to form a single larger vacuole." [GOC:pr, GOC:vw, Wikipedia:Vacuole] |
| GO:0072741 | 0.003584018 | 2 / 2 | "A cellular protein localization process in which a protein is transported to, or maintained at, the site of cell division." [GOC:mah, PMID:19756689] |
| GO:0030968 | 0.003710159 | 4 / 12 | "The series of molecular signals generated as a consequence of the presence of unfolded proteins in the endoplasmic reticulum (ER) or other ER-related stress; results in changes in the regulation of transcript |
| GO:0007165 | 0.004399866 | 5 / 20 | "The cellular process in which a signal is conveyed to trigger a change in the activity or state of a cell. Signal transduction begins with reception of a signal (e.g. a ligand binding to a receptor or receptor activat |
| GO:0007154 | 0.00505638 | 4 / 13 | "Any process that mediates interactions between a cell and its surroundings. Encompasses interactions such as signaling or attachment between one cell and another cell, between a cell and an extracellular r |
| GO:1905037 | 0.005768234 | 3 / 7 | "A process that is carried out at the cellular level which results in the assembly, arrangement of constituent parts, or disassembly of an autophagosome." [GOC:bf, GOC:PARL, GOC:TermGenie, PMID:22186024 |

| GO_Term | P_val | Counts | CellComp_Description |
|---|---|---|---|
| GO:0006554 | 0.004087271 | 2 / 2 | "The chemical reactions and pathways resulting in the breakdown of lysine, 2,6-diaminohexanoic acid." [GOC:go_curators] |
| GO:0046440 | 0.004087271 | 2 / 2 | "The chemical reactions and pathways involving L-lysine, the L-enantiomer of (S)-2,6-diaminohexanoic acid, i.e. (2S)-2,6-diaminohexanoic acid." [GOC:ai, GOC:jsg, GOC:mah] |
| GO:0046394 | 0.021549754 | 2 / 4 | "The chemical reactions and pathways resulting in the formation of carboxylic acids, any organic acid containing one or more carboxyl (-COOH) groups." [ISBN:0198506732] |
| GO:0005727 | 0.024387104 | 6 / 39 | "Circular DNA structures that are not part of a chromosome." [GOC:ai] |
| GO:0016234 | 0.024387104 | 6 / 39 | "A discrete intracellular part formed of aggregated molecules such as proteins or other biopolymers." [GOC:mah, PMID:11121744] |
| GO:0019774 | 0.024387104 | 6 / 39 | "The proteasome core subcomplex that constitutes the two inner rings of the proteasome core complex. An example of this component is found in Mus musculus." [GOC:jl, GOC:mtg_sensu, GOC:rb, PMID:10 |
| GO:0030076 | 0.024387104 | 6 / 39 | "A protein-pigment complex that may be closely or peripherally associated to photosynthetic reaction centers that participate in harvesting and transferring radiant energy to the reaction center." [GOC:lr] |
| GO:0032300 | 0.024387104 | 6 / 39 | "Any complex formed of proteins that act in mismatch repair." [GOC:mah] |
| GO:0034708 | 0.024387104 | 6 / 39 | "A protein complex that possesses methyltransferase activity." [GOC:mah] |
| GO:0043229 | 0.024387104 | 6 / 39 | "Organized structure of distinctive morphology and function, occurring within the cell. Includes the nucleus, mitochondria, plastids, vacuoles, vesicles, ribosomes and the cytoskeleton. Excludes the plasma m |

#### Supplementary Data S3.3– Temporal GO Description of DPoP analysis Transcriptomic Data

Each one of the following pictures is an aggregation of outputs of a GO term enrichment analysis on the results of DPoP at each specific point along the transcriptomic time course.

##### Supplementary Data S3.3.1

### Timepoint 1

###### Molecular Function

| | |
|---|---|
| 356 / 3514 | "The action of a molecule that contributes to the structural integrity of a complex or its assembly within or outside a cell." [GOC:mah, GOC:vw] |
| 18 / 510 | "Any molecular function by which a gene product interacts selectively and non-covalently with DNA (deoxyribonucleic acid)." [GOC:dph, GOC:jl, GOC:tb, GOC:vw] |
| 19 / 515 | "Binding to a nucleic acid." [GOC:jl] |
| 362 / 3521 | "A molecular function characterized by the coupling of ATP hydrolysis to other steps of a reaction mechanism to make the reaction energetically favorable, for example to catalyz |
| 121 / 632 | "Catalysis of an oxidation-reduction (redox) reaction, a reversible chemical reaction in which the oxidation state of an atom or atoms within a molecule is altered. One substrate a |
| 80 / 358 | "Catalysis of an oxidation-reduction (redox) reaction in which an aldehyde or ketone (oxo) group acts as a hydrogen or electron donor and reduces a hydrogen or electron accept |
| 78 / 349 | "Catalysis of an oxidation-reduction (redox) reaction in which a CH-NH group acts as a hydrogen or electron donor and reduces a hydrogen or electron acceptor." [GOC:ai] |
| 77 / 347 | "Catalysis of the reaction: 3,5,3'-L-triiodo-L-thyronine + iodide + acceptor + H+ = L-thyroxine + donor-H2." [EC:1.21.99.4] |
| 77 / 347 | "Catalysis of the hydroxylation of tyramine to form octopamine." [PMID:10745161] |
| 77 / 347 | "Catalysis of the reaction: 2 H2O = O2 + 4 H+ + 4 e-. The evolution of oxygen from oxidizing water is carried out by the oxygen evolving complex in photosystem II of plants. P680 |

###### Biological Process

| | |
|---|---|
| 397 / 3778 | "Any process that is carried out at the cellular level, but not necessarily restricted to a single cell. For example, cell communication occurs among more than one cell, but occurs at the cellular level." [GOC:go_cur |
| 1 / 150 | "The cellular metabolic process in which a protein is formed, using the sequence of a mature mRNA or circRNA molecule to specify the sequence of amino acids in a polypeptide chain. Translation is mediated by |
| 407 / 3664 | "The chemical reactions and pathways, including anabolism and catabolism, by which living organisms transform chemical substances. Metabolic processes typically transform small molecules, but also include m |
| 0 / 107 | "The chemical reactions and pathways resulting in the formation of a protein in the cytoplasm. This is a ribosome-mediated process in which the information in messenger RNA (mRNA) is used to specify the sequ |
| 0 / 106 | "The synthesis of aminoacyl tRNA by the formation of an ester bond between the 3'-hydroxyl group of the most 3' adenosine of the tRNA and the alpha carboxylic acid group of an amino acid, to be used in ribo |
| 73 / 431 | "OBSOLETE. A metabolic process that results in the removal or addition of one or more electrons to or from a substance, with or without the concomitant removal or addition of a proton or protons." [GOC:dhl, G |
| 17 / 52 | "The chemical reactions and pathways involving an ergot alkaloid, an indole alkaloid." [GOC:yaf] |
| 16 / 47 | "The chemical reactions and pathways involving melanins, pigments largely of animal origin. High molecular weight polymers of indole quinone, they are irregular polymeric structures and are divided into three |
| 16 / 47 | "The chemical reactions and pathways involving glycosinolates, substituted thioglycosides found in rapeseed products and related cruciferae." [GOC:mah, http://www.gardeneaters.net/family_characteristics.htn |
| 16 / 48 | "The chemical reactions and pathways resulting in the formation of verruculogen." [GOC:di, GOC:TermGenie, PMID:23649274] |

###### Cellular Compartment

| | |
|---|---|
| 13 / 494 | "A biological process represents a specific objective that the organism is genetically programmed to achieve. Biological processes are often described by their outcome or ending state, e.g., the biological process of c |
| 14 / 503 | "The chemical reactions and pathways, including anabolism and catabolism, by which living organisms transform chemical substances. Metabolic processes typically transform small molecules, but also include macr |
| 14 / 503 | "Any process that is carried out at the cellular level, but not necessarily restricted to a single cell. For example, cell communication occurs among more than one cell, but occurs at the cellular level." [GOC:go_curato |
| 13 / 476 | "The increase in size or mass of an entire organism, a part of an organism or a cell." [GOC:bf, GOC:ma] |
| 13 / 466 | "A biological process that directly contributes to the process of producing new individuals by one or two organisms. The new individuals inherit some proportion of their genetic material from the parent or parents." |
| 13 / 466 | "The entirety of a process in which information is transmitted within a biological system. This process begins with an active signal and ends when a cellular response has been triggered." [GOC:mtg_signal, GOC:mtg_ |
| 13 / 466 | "Any process that reduces or removes the toxicity of a toxic substance. These may include transport of the toxic substance away from sensitive areas and to compartments or complexes whose purpose is sequestrat |
| 13 / 464 | "Any process that results in a change in state or activity of a cell or an organism (in terms of movement, secretion, enzyme production, gene expression, etc.) as a result of a stimulus. The process begins with detecti |
| 13 / 463 | "The production of new individuals that contain some portion of genetic material inherited from one or more parent organisms." [GOC:go_curators, GOC:isa_complete, GOC:jl, ISBN:0198506732] |
| 13 / 463 | "Any process involved in the development or functioning of the immune system, an organismal system for calibrated responses to potential internal or invasive threats." [GO_REF:0000022, GOC:add] |

##### Supplementary Data S3.3.2

### Timepoint 2

###### Molecular Function

| | |
|---|---|
| 352 / 3514 | "The action of a molecule that contributes to the structural integrity of a complex or its assembly within or outside a cell." [GOC:mah, GOC:vw] |
| 18 / 510 | "Any molecular function by which a gene product interacts selectively and non-covalently with DNA (deoxyribonucleic acid)." [GOC:dph, GOC:jl, GOC:tb, GOC:vw] |
| 19 / 515 | "Binding to a nucleic acid." [GOC:jl] |
| 358 / 3521 | "A molecular function characterized by the coupling of ATP hydrolysis to other steps of a reaction mechanism to make the reaction energetically favorable, for example to catalyze a reaction or drive transport aga |
| 117 / 632 | "Catalysis of an oxidation-reduction (redox) reaction, a reversible chemical reaction in which the oxidation state of an atom or atoms within a molecule is altered. One substrate acts as a hydrogen or electron dor |
| 10 / 330 | "Binding to an RNA molecule or a portion thereof." [GOC:jl, GOC:mah] |
| 354 / 3464 | "The selective, non-covalent, often stoichiometric, interaction of a molecule with one or more specific sites on another molecule." [GOC:ceb, GOC:mah, ISBN:0198506732] |
| 80 / 386 | "Catalysis of an oxidation-reduction (redox) reaction in which a CH-OH group act as a hydrogen or electron donor and reduces a hydrogen or electron acceptor." [GOC:ai] |
| 74 / 349 | "Catalysis of an oxidation-reduction (redox) reaction in which a CH-NH group acts as a hydrogen or electron donor and reduces a hydrogen or electron acceptor." [GOC:ai] |
| 22 / 478 | "Binding to a zinc ion (Zn)." [GOC:ai] |

###### Biological Process

| | |
|---|---|
| 392 / 3778 | "Any process that is carried out at the cellular level, but not necessarily restricted to a single cell. For example, cell communication occurs among more than one cell, but occurs at the cellular level." [GOC:go_cu |
| 1 / 150 | "The cellular metabolic process in which a protein is formed, using the sequence of a mature mRNA or circRNA molecule to specify the sequence of amino acids in a polypeptide chain. Translation is mediated by |
| 403 / 3664 | "The chemical reactions and pathways, including anabolism and catabolism, by which living organisms transform chemical substances. Metabolic processes typically transform small molecules, but also include r |
| 0 / 111 | "The process preceding formation of the peptide bond between the first two amino acids of a protein. This includes the formation of a complex of the ribosome, mRNA or circRNA, and an initiation complex that |
| 62 / 342 | "The chemical reactions and pathways resulting in many of the chemical changes of compounds that are not necessarily required for growth and maintenance of cells, and are often unique to a taxon. In multice |
| 0 / 107 | "The chemical reactions and pathways resulting in the formation of a protein in the cytoplasm. This is a ribosome-mediated process in which the information in messenger RNA (mRNA) is used to specify the seq |
| 0 / 106 | "The synthesis of aminoacyl tRNA by the formation of an ester bond between the 3'-hydroxyl group of the most 3' adenosine of the tRNA and the alpha carboxylic acid group of an amino acid, to be used in ribo |
| 0 / 92 | "The successive addition of amino acid residues to a nascent polypeptide chain during protein biosynthesis." [GOC:ems] |
| 0 / 92 | "The chemical reactions and pathways resulting in the formation of a protein in a mitochondrion. This is a ribosome-mediated process in which the information in messenger RNA (mRNA) is used to specify the s |
| 16 / 336 | "Any process that modulates the frequency, rate or extent of transcription mediated by RNA polymerase II." [GOC:go_curators, GOC:txnOH] |

###### Cellular Compartment

| | |
|---|---|
| 13 / 494 | "A biological process represents a specific objective that the organism is genetically programmed to achieve. Biological processes are often described by their outcome or ending state, e.g., the biological process of c |
| 14 / 503 | "The chemical reactions and pathways, including anabolism and catabolism, by which living organisms transform chemical substances. Metabolic processes typically transform small molecules, but also include macr |
| 14 / 503 | "Any process that is carried out at the cellular level, but not necessarily restricted to a single cell. For example, cell communication occurs among more than one cell, but occurs at the cellular level." [GOC:go_curato |
| 13 / 476 | "The increase in size or mass of an entire organism, a part of an organism or a cell." [GOC:bf, GOC:ma] |
| 13 / 466 | "A biological process that directly contributes to the process of producing new individuals by one or two organisms. The new individuals inherit some proportion of their genetic material from the parent or parents." |
| 13 / 466 | "The entirety of a process in which information is transmitted within a biological system. This process begins with an active signal and ends when a cellular response has been triggered." [GOC:mtg_signal, GOC:mtg_ |
| 13 / 466 | "Any process that reduces or removes the toxicity of a toxic substance. These may include transport of the toxic substance away from sensitive areas and to compartments or complexes whose purpose is sequestrat |
| 13 / 464 | "Any process that results in a change in state or activity of a cell or an organism (in terms of movement, secretion, enzyme production, gene expression, etc.) as a result of a stimulus. The process begins with detecti |
| 13 / 463 | "The production of new individuals that contain some portion of genetic material inherited from one or more parent organisms." [GOC:go_curators, GOC:isa_complete, GOC:jl, ISBN:0198506732] |
| 13 / 463 | "Any process involved in the development or functioning of the immune system, an organismal system for calibrated responses to potential internal or invasive threats." [GO_REF:0000022, GOC:add] |

Supplementary Data S3.3.3

### Timepoint 3

#### Molecular Function

| | |
|---|---|
| 18 / 510 | "Any molecular function by which a gene product interacts selectively and non-covalently with DNA (deoxyribonucleic acid)." [GOC:dph, GOC:jl, GOC:tb, GOC:vw] |
| 19 / 515 | "Binding to a nucleic acid." [GOC:jl] |
| 359 / 3514 | "The action of a molecule that contributes to the structural integrity of a complex or its assembly within or outside a cell." [GOC:mah, GOC:vw] |
| 10 / 330 | "Binding to an RNA molecule or a portion thereof." [GOC:jl, GOC:mah] |
| 366 / 3521 | "A molecular function characterized by the coupling of ATP hydrolysis to other steps of a reaction mechanism to make the reaction energetically favorable, for example to catalyze a reaction or drive transport aga |
| 115 / 632 | "Catalysis of an oxidation-reduction (redox) reaction, a reversible chemical reaction in which the oxidation state of an atom or atoms within a molecule is altered. One substrate acts as a hydrogen or electron don |
| 23 / 478 | "Binding to a zinc ion (Zn)." [GOC:ai] |
| 23 / 477 | "Binding to and responding, e.g. by conformational change, to changes in the cellular level of zinc." [GOC:vw, PMID:31239353] |
| 11 / 315 | "Binding to DNA that is assembled into chromatin." [GOC:mah] |
| 0 / 123 | "The action of a molecule that contributes to the structural integrity of the ribosome." [GOC:mah] |

#### Biological Process

| | |
|---|---|
| 398 / 3778 | "Any process that is carried out at the cellular level, but not necessarily restricted to a single cell. For example, cell communication occurs among more than one cell, but occurs at the cellular level." [GOC:go_cura |
| 1 / 150 | "The cellular metabolic process in which a protein is formed, using the sequence of a mature mRNA or circRNA molecule to specify the sequence of amino acids in a polypeptide chain. Translation is mediated by tl |
| 409 / 3664 | "The chemical reactions and pathways, including anabolism and catabolism, by which living organisms transform chemical substances. Metabolic processes typically transform small molecules, but also include ma |
| 64 / 342 | "The chemical reactions and pathways resulting in many of the chemical changes of compounds that are not necessarily required for growth and maintenance of cells, and are often unique to a taxon. In multicellu |
| 0 / 111 | "The process preceding formation of the peptide bond between the first two amino acids of a protein. This includes the formation of a complex of the ribosome, mRNA or circRNA, and an initiation complex that c |
| 0 / 107 | "The chemical reactions and pathways resulting in the formation of a protein in the cytoplasm. This is a ribosome-mediated process in which the information in messenger RNA (mRNA) is used to specify the seque |
| 0 / 106 | "The synthesis of aminoacyl tRNA by the formation of an ester bond between the 3'-hydroxyl group of the most 3' adenosine of the tRNA and the alpha carboxylic acid group of an amino acid, to be used in ribosc |
| 17 / 52 | "The chemical reactions and pathways involving an ergot alkaloid, an indole alkaloid." [GOC:yaf] |
| 16 / 47 | "The chemical reactions and pathways involving melanins, pigments largely of animal origin. High molecular weight polymers of indole quinone, they are irregular polymeric structures and are divided into three g |
| 16 / 47 | "The chemical reactions and pathways involving glycosinolates, substituted thioglycosides found in rapeseed products and related cruciferae." [GOC:mah, http://www.gardeneaters.net/family_characteristics.html |

#### Cellular Compartment

| | |
|---|---|
| 14 / 503 | "The chemical reactions and pathways, including anabolism and catabolism, by which living organisms transform chemical substances. Metabolic processes typically transform small molecules, but also include macr |
| 14 / 494 | "A biological process represents a specific objective that the organism is genetically programmed to achieve. Biological processes are often described by their outcome or ending state, e.g., the biological process of |
| 15 / 503 | "Any process that is carried out at the cellular level, but not necessarily restricted to a single cell. For example, cell communication occurs among more than one cell, but occurs at the cellular level." [GOC:go_curato |
| 14 / 476 | "The increase in size or mass of an entire organism, a part of an organism or a cell." [GOC:bf, GOC:ma] |
| 21 / 575 | "Organized structure of distinctive morphology and function, bounded by a single or double lipid bilayer membrane and occurring within the cell. Includes the nucleus, mitochondria, plastids, vacuoles, and vesicles. |
| 14 / 466 | "A biological process that directly contributes to the process of producing new individuals by one or two organisms. The new individuals inherit some proportion of their genetic material from the parent or parents. |
| 14 / 466 | "The entirety of a process in which information is transmitted within a biological system. This process begins with an active signal and ends when a cellular response has been triggered." [GOC:mtg_signal, GOC:mtg_ |
| 14 / 466 | "Any process that reduces or removes the toxicity of a toxic substance. These may include transport of the toxic substance away from sensitive areas and to compartments or complexes whose purpose is sequestrat |
| 14 / 464 | "Any process that results in a change in state or activity of a cell or an organism (in terms of movement, secretion, enzyme production, gene expression, etc.) as a result of a stimulus. The process begins with detecti |
| 14 / 463 | "The production of new individuals that contain some portion of genetic material inherited from one or more parent organisms." [GOC:go_curators, GOC:isa_complete, GOC:jl, ISBN:0198506732] |

Supplementary Data S3.3.4

### Timepoint 4

#### Molecular Function

| | |
|---|---|
| 16 / 510 | "Any molecular function by which a gene product interacts selectively and non-covalently with DNA (deoxyribonucleic acid)." [GOC:dph, GOC:jl, GOC:tb, GOC:vw] |
| 18 / 515 | "Binding to a nucleic acid." [GOC:jl] |
| 353 / 3514 | "The action of a molecule that contributes to the structural integrity of a complex or its assembly within or outside a cell." [GOC:mah, GOC:vw] |
| 9 / 330 | "Binding to an RNA molecule or a portion thereof." [GOC:jl, GOC:mah] |
| 359 / 3521 | "A molecular function characterized by the coupling of ATP hydrolysis to other steps of a reaction mechanism to make the reaction energetically favorable, for example to catalyze a reaction or drive transport ag |
| 116 / 632 | "Catalysis of an oxidation-reduction (redox) reaction, a reversible chemical reaction in which the oxidation state of an atom or atoms within a molecule is altered. One substrate acts as a hydrogen or electron dor |
| 23 / 478 | "Binding to a zinc ion (Zn)." [GOC:ai] |
| 23 / 477 | "Binding to and responding, e.g. by conformational change, to changes in the cellular level of zinc." [GOC:vw, PMID:31239353] |
| 360 / 3487 | "Enables the directed movement of substances (such as macromolecules, small molecules, ions) into, out of or within a cell, or between cells." [GOC:ai, GOC:dgf] |
| 357 / 3464 | "The selective, non-covalent, often stoichiometric, interaction of a molecule with one or more specific sites on another molecule." [GOC:ceb, GOC:mah, ISBN:0198506732] |

#### Biological Process

| | |
|---|---|
| 392 / 3778 | "Any process that is carried out at the cellular level, but not necessarily restricted to a single cell. For example, cell communication occurs among more than one cell, but occurs at the cellular level." [GOC:go_cura |
| 0 / 150 | "The cellular metabolic process in which a protein is formed, using the sequence of a mature mRNA or circRNA molecule to specify the sequence of amino acids in a polypeptide chain. Translation is mediated by tl |
| 65 / 342 | "The chemical reactions and pathways resulting in many of the chemical changes of compounds that are not necessarily required for growth and maintenance of cells, and are often unique to a taxon. In multicellu |
| 405 / 3664 | "The chemical reactions and pathways, including anabolism and catabolism, by which living organisms transform chemical substances. Metabolic processes typically transform small molecules, but also include ma |
| 0 / 108 | "The chemical reactions and pathways involving a specific protein, rather than of proteins in general, occurring at the level of an individual cell. Includes cellular protein modification." [GOC:jl] |
| 0 / 107 | "The chemical reactions and pathways resulting in the formation of a protein in the cytoplasm. This is a ribosome-mediated process in which the information in messenger RNA (mRNA) is used to specify the seque |
| 0 / 106 | "The synthesis of aminoacyl tRNA by the formation of an ester bond between the 3'-hydroxyl group of the most 3' adenosine of the tRNA and the alpha carboxylic acid group of an amino acid, to be used in ribosc |
| 17 / 52 | "The chemical reactions and pathways involving an ergot alkaloid, an indole alkaloid." [GOC:yaf] |
| 16 / 47 | "The chemical reactions and pathways involving melanins, pigments largely of animal origin. High molecular weight polymers of indole quinone, they are irregular polymeric structures and are divided into three g |
| 16 / 47 | "The chemical reactions and pathways involving glycosinolates, substituted thioglycosides found in rapeseed products and related cruciferae." [GOC:mah, http://www.gardeneaters.net/family_characteristics.html |

#### Cellular Compartment

| | |
|---|---|
| 12 / 503 | "The chemical reactions and pathways, including anabolism and catabolism, by which living organisms transform chemical substances. Metabolic processes typically transform small molecules, but also include macr |
| 12 / 494 | "A biological process represents a specific objective that the organism is genetically programmed to achieve. Biological processes are often described by their outcome or ending state, e.g., the biological process of |
| 13 / 503 | "Any process that is carried out at the cellular level, but not necessarily restricted to a single cell. For example, cell communication occurs among more than one cell, but occurs at the cellular level." [GOC:go_curato |
| 12 / 476 | "The increase in size or mass of an entire organism, a part of an organism or a cell." [GOC:bf, GOC:ma] |
| 12 / 466 | "A biological process that directly contributes to the process of producing new individuals by one or two organisms. The new individuals inherit some proportion of their genetic material from the parent or parents.' |
| 12 / 466 | "The entirety of a process in which information is transmitted within a biological system. This process begins with an active signal and ends when a cellular response has been triggered." [GOC:mtg_signal, GOC:mtg_ |
| 12 / 466 | "Any process that reduces or removes the toxicity of a toxic substance. These may include transport of the toxic substance away from sensitive areas and to compartments or complexes whose purpose is sequestrat |
| 12 / 464 | "Any process that results in a change in state or activity of a cell or an organism (in terms of movement, secretion, enzyme production, gene expression, etc.) as a result of a stimulus. The process begins with detecti |
| 12 / 463 | "The production of new individuals that contain some portion of genetic material inherited from one or more parent organisms." [GOC:go_curators, GOC:isa_complete, GOC:jl, ISBN:0198506732] |
| 12 / 463 | "Any process involved in the development or functioning of the immune system, an organismal system for calibrated responses to potential internal or invasive threats." [GO_REF:0000022, GOC:add] |

Supplementary Data S3.3.5

### Timepoint 5

#### Molecular Function

| | |
|---|---|
| 15 / 510 | "Any molecular function by which a gene product interacts selectively and non-covalently with DNA (deoxyribonucleic acid)." [GOC:dph, GOC:jl, GOC:tb, GOC:vw] |
| 17 / 515 | "Binding to a nucleic acid." [GOC:jl] |
| 340 / 3514 | "The action of a molecule that contributes to the structural integrity of a complex or its assembly within or outside a cell." [GOC:mah, GOC:vw] |
| 8 / 330 | "Binding to an RNA molecule or a portion thereof." [GOC:jl, GOC:mah] |
| 344 / 3521 | "A molecular function characterized by the coupling of ATP hydrolysis to other steps of a reaction mechanism to make the reaction energetically favorable, for example to catalyze a reaction or drive transport aga |
| 20 / 478 | "Binding to a zinc ion (Zn)." [GOC:ai] |
| 20 / 477 | "Binding to and responding, e.g. by conformational change, to changes in the cellular level of zinc." [GOC:vw, PMID:31239353] |
| 109 / 632 | "Catalysis of an oxidation-reduction (redox) reaction, a reversible chemical reaction in which the oxidation state of an atom or atoms within a molecule is altered. One substrate acts as a hydrogen or electron don |
| 10 / 315 | "Binding to DNA that is assembled into chromatin." [GOC:mah] |
| 10 / 305 | "Binding to double-stranded DNA." [GOC:elh, GOC:vw] |

#### Biological Process

| | |
|---|---|
| 384 / 3778 | "Any process that is carried out at the cellular level, but not necessarily restricted to a single cell. For example, cell communication occurs among more than one cell, but occurs at the cellular level." [GOC:go_cura |
| 0 / 150 | "The cellular metabolic process in which a protein is formed, using the sequence of a mature mRNA or circRNA molecule to specify the sequence of amino acids in a polypeptide chain. Translation is mediated by tl |
| 62 / 342 | "The chemical reactions and pathways resulting in many of the chemical changes of compounds that are not necessarily required for growth and maintenance of cells, and are often unique to a taxon. In multicellu |
| 393 / 3664 | "The chemical reactions and pathways, including anabolism and catabolism, by which living organisms transform chemical substances. Metabolic processes typically transform small molecules, but also include ma |
| 0 / 108 | "The chemical reactions and pathways involving a specific protein, rather than of proteins in general, occurring at the level of an individual cell. Includes cellular protein modification." [GOC:jl] |
| 0 / 107 | "The chemical reactions and pathways resulting in the formation of a protein in the cytoplasm. This is a ribosome-mediated process in which the information in messenger RNA (mRNA) is used to specify the seque |
| 0 / 106 | "The synthesis of aminoacyl tRNA by the formation of an ester bond between the 3'-hydroxyl group of the most 3' adenosine of the tRNA and the alpha carboxylic acid group of an amino acid, to be used in ribosc |
| 14 / 336 | "Any process that modulates the frequency, rate or extent of transcription mediated by RNA polymerase II." [GOC:go_curators, GOC:txnOH] |
| 12 / 305 | "Any process that modulates the frequency, rate or extent of cellular DNA-templated transcription." [GOC:go_curators, GOC:txnOH] |
| 50 / 275 | "The chemical reactions and pathways involving pigment, any general or particular coloring matter in living organisms, e.g. melanin." [GOC:jl, ISBN:0198506732] |

#### Cellular Compartment

| | |
|---|---|
| 12 / 503 | "The chemical reactions and pathways, including anabolism and catabolism, by which living organisms transform chemical substances. Metabolic processes typically transform small molecules, but also include macro |
| 12 / 494 | "A biological process represents a specific objective that the organism is genetically programmed to achieve. Biological processes are often described by their outcome or ending state, e.g., the biological process of c |
| 13 / 503 | "Any process that is carried out at the cellular level, but not necessarily restricted to a single cell. For example, cell communication occurs among more than one cell, but occurs at the cellular level." [GOC:go_curator |
| 12 / 476 | "The increase in size or mass of an entire organism, a part of an organism or a cell." [GOC:bf, GOC:ma] |
| 12 / 466 | "A biological process that directly contributes to the process of producing new individuals by one or two organisms. The new individuals inherit some proportion of their genetic material from the parent or parents." |
| 12 / 466 | "The entirety of a process in which information is transmitted within a biological system. This process begins with an active signal and ends when a cellular response has been triggered." [GOC:mtg_signal, GOC:mtg_ |
| 12 / 466 | "Any process that reduces or removes the toxicity of a toxic substance. These may include transport of the toxic substance away from sensitive areas and to compartments or complexes whose purpose is sequestrati |
| 19 / 575 | "Organized structure of distinctive morphology and function, bounded by a single or double lipid bilayer membrane and occurring within the cell. Includes the nucleus, mitochondria, plastids, vacuoles, and vesicles. E |
| 12 / 464 | "Any process that results in a change in state or activity of a cell or an organism (in terms of movement, secretion, enzyme production, gene expression, etc.) as a result of a stimulus. The process begins with detectic |
| 12 / 463 | "The production of new individuals that contain some portion of genetic material inherited from one or more parent organisms." [GOC:go_curators, GOC:isa_complete, GOC:jl, ISBN:0198506732] |

Supplementary Data S3.3.6

### Timepoint 6

#### Molecular Function

| | |
|---|---|
| 16 / 510 | "Any molecular function by which a gene product interacts selectively and non-covalently with DNA (deoxyribonucleic acid)." [GOC:dph, GOC:jl, GOC:tb, GOC:vw] |
| 18 / 515 | "Binding to a nucleic acid." [GOC:jl] |
| 340 / 3514 | "The action of a molecule that contributes to the structural integrity of a complex or its assembly within or outside a cell." [GOC:mah, GOC:vw] |
| 8 / 330 | "Binding to an RNA molecule or a portion thereof." [GOC:jl, GOC:mah] |
| 344 / 3521 | "A molecular function characterized by the coupling of ATP hydrolysis to other steps of a reaction mechanism to make the reaction energetically favorable, for example to catalyze a reaction or drive transport aga |
| 19 / 478 | "Binding to a zinc ion (Zn)." [GOC:ai] |
| 19 / 477 | "Binding to and responding, e.g. by conformational change, to changes in the cellular level of zinc." [GOC:vw, PMID:31239353] |
| 108 / 632 | "Catalysis of an oxidation-reduction (redox) reaction, a reversible chemical reaction in which the oxidation state of an atom or atoms within a molecule is altered. One substrate acts as a hydrogen or electron don |
| 344 / 3464 | "The selective, non-covalent, often stoichiometric, interaction of a molecule with one or more specific sites on another molecule." [GOC:ceb, GOC:mah, ISBN:0198506732] |
| 11 / 315 | "Binding to DNA that is assembled into chromatin." [GOC:mah] |

#### Biological Process

| | |
|---|---|
| 0 / 150 | "The cellular metabolic process in which a protein is formed, using the sequence of a mature mRNA or circRNA molecule to specify the sequence of amino acids in a polypeptide chain. Translation is mediated by the |
| 13 / 336 | "Any process that modulates the frequency, rate or extent of transcription mediated by RNA polymerase II." [GOC:go_curators, GOC:txnOH] |
| 11 / 305 | "Any process that modulates the frequency, rate or extent of cellular DNA-templated transcription." [GOC:go_curators, GOC:txnOH] |
| 0 / 108 | "The chemical reactions and pathways involving a specific protein, rather than of proteins in general, occurring at the level of an individual cell. Includes cellular protein modification." [GOC:jl] |
| 0 / 107 | "The chemical reactions and pathways resulting in the formation of a protein in the cytoplasm. This is a ribosome-mediated process in which the information in messenger RNA (mRNA) is used to specify the sequenc |
| 0 / 106 | "The synthesis of aminoacyl tRNA by the formation of an ester bond between the 3'-hydroxyl group of the most 3' adenosine of the tRNA and the alpha carboxylic acid group of an amino acid, to be used in ribosom |
| 12 / 302 | "Any process that activates or increases the frequency, rate or extent of cellular DNA-templated transcription." [GOC:go_curators, GOC:txnOH] |
| 12 / 298 | "Any process that stops, prevents, or reduces the frequency, rate or extent of cellular DNA-templated transcription." [GOC:go_curators, GOC:txnOH] |
| 58 / 342 | "The chemical reactions and pathways resulting in many of the chemical changes of compounds that are not necessarily required for growth and maintenance of cells, and are often unique to a taxon. In multicellula |
| 11 / 278 | "Any process that regulates transcription such that the target genes are involved in the transition between G1 and S phase of the mitotic cell cycle." [GOC:mtg_cell_cycle] |

#### Cellular Compartment

| | |
|---|---|
| 6 / 503 | "The chemical reactions and pathways, including anabolism and catabolism, by which living organisms transform chemical substances. Metabolic processes typically transform small molecules, but also include macro |
| 6 / 494 | "A biological process represents a specific objective that the organism is genetically programmed to achieve. Biological processes are often described by their outcome or ending state, e.g., the biological process of ce |
| 7 / 503 | "Any process that is carried out at the cellular level, but not necessarily restricted to a single cell. For example, cell communication occurs among more than one cell, but occurs at the cellular level." [GOC:go_curators |
| 6 / 476 | "The increase in size or mass of an entire organism, a part of an organism or a cell." [GOC:bf, GOC:ma] |
| 6 / 466 | "A biological process that directly contributes to the process of producing new individuals by one or two organisms. The new individuals inherit some proportion of their genetic material from the parent or parents." |
| 6 / 466 | "The entirety of a process in which information is transmitted within a biological system. This process begins with an active signal and ends when a cellular response has been triggered." [GOC:mtg_signal, GOC:mtg_s |
| 6 / 466 | "Any process that reduces or removes the toxicity of a toxic substance. These may include transport of the toxic substance away from sensitive areas and to compartments or complexes whose purpose is sequestratic |
| 6 / 464 | "Any process that results in a change in state or activity of a cell or an organism (in terms of movement, secretion, enzyme production, gene expression, etc.) as a result of a stimulus. The process begins with detectio |
| 6 / 463 | "The production of new individuals that contain some portion of genetic material inherited from one or more parent organisms." [GOC:go_curators, GOC:isa_complete, GOC:jl, ISBN:0198506732] |
| 6 / 463 | "Any process involved in the development or functioning of the immune system, an organismal system for calibrated responses to potential internal or invasive threats." [GO_REF:0000022, GOC:add] |

Supplementary Data S3.3.7

### Timepoint 7

#### Molecular Function

| | |
|---|---|
| 16 / 510 | "Any molecular function by which a gene product interacts selectively and non-covalently with DNA (deoxyribonucleic acid)." [GOC:dph, GOC:jl, GOC:tb, GOC:vw] |
| 347 / 3514 | "The action of a molecule that contributes to the structural integrity of a complex or its assembly within or outside a cell." [GOC:mah, GOC:vw] |
| 18 / 515 | "Binding to a nucleic acid." [GOC:jl] |
| 351 / 3521 | "A molecular function characterized by the coupling of ATP hydrolysis to other steps of a reaction mechanism to make the reaction energetically favorable, for example to catalyze a reaction or drive transport aga |
| 7 / 330 | "Binding to an RNA molecule or a portion thereof." [GOC:jl, GOC:mah] |
| 121 / 632 | "Catalysis of an oxidation-reduction (redox) reaction, a reversible chemical reaction in which the oxidation state of an atom or atoms within a molecule is altered. One substrate acts as a hydrogen or electron don |
| 19 / 478 | "Binding to a zinc ion (Zn)." [GOC:ai] |
| 19 / 477 | "Binding to and responding, e.g. by conformational change, to changes in the cellular level of zinc." [GOC:vw, PMID:31239353] |
| 350 / 3464 | "The selective, non-covalent, often stoichiometric, interaction of a molecule with one or more specific sites on another molecule." [GOC:ceb, GOC:mah, ISBN:0198506732] |
| 348 / 3446 | "A molecular function that controls the rate, timing and/or magnitude of gene transcription. The function of transcriptional regulators is to modulate gene expression at the transcription step so that they are exp |

#### Biological Process

| | |
|---|---|
| 0 / 150 | "The cellular metabolic process in which a protein is formed, using the sequence of a mature mRNA or circRNA molecule to specify the sequence of amino acids in a polypeptide chain. Translation is mediated by the |
| 69 / 369 | "The process in which an organomercurial compound is transported across a membrane. Organomercurial substances are any organic compound containing a mercury atom." [GOC:ai, PMID:18793329] |
| 69 / 369 | "The directed movement of bicyclomycin across a lipid bilayer, from one side of a membrane to the other. Bicyclomycin (or bicozamycin) is an antibacterial drug often used as a livestock feed additive." [PMID:20067 |
| 69 / 369 | "The process in which carbon dioxide (CO2) is transported across a membrane." [GOC:yaf] |
| 69 / 369 | "The process in which urea, the water-soluble compound H2N-CO-NH2, is transported from one side of a membrane to the other by means of some agent such as a transporter or pore." [GOC:mah] |
| 69 / 369 | "The process in which a pyrimidine-containing compound is transported across a membrane. A pyrimidine-containing compound is any compound that contains pyrimidine or a formal derivative thereof." [GOC:mah] |
| 69 / 369 | "The directed movement of auxins from endoplasmic reticulum to cytosol." [PMID:19506555] |
| 69 / 369 | "The process in which hydrogen peroxide is transported across a membrane." [GOC:tb] |
| 69 / 369 | "The directed movement of a substance across the outer membrane in cells with two membranes." [PMID:15968039] |
| 69 / 369 | "The directed movement of benomyl across a membrane." [GOC:TermGenie] |

#### Cellular Compartment

| | |
|---|---|
| 7 / 503 | "The chemical reactions and pathways, including anabolism and catabolism, by which living organisms transform chemical substances. Metabolic processes typically transform small molecules, but also include macro |
| 7 / 494 | "A biological process represents a specific objective that the organism is genetically programmed to achieve. Biological processes are often described by their outcome or ending state, e.g., the biological process of ce |
| 8 / 503 | "Any process that is carried out at the cellular level, but not necessarily restricted to a single cell. For example, cell communication occurs among more than one cell, but occurs at the cellular level." [GOC:go_curators |
| 7 / 476 | "The increase in size or mass of an entire organism, a part of an organism or a cell." [GOC:bf, GOC:ma] |
| 7 / 466 | "A biological process that directly contributes to the process of producing new individuals by one or two organisms. The new individuals inherit some proportion of their genetic material from the parent or parents." [ |
| 7 / 466 | "The entirety of a process in which information is transmitted within a biological system. This process begins with an active signal and ends when a cellular response has been triggered." [GOC:mtg_signal, GOC:mtg_s |
| 7 / 466 | "Any process that reduces or removes the toxicity of a toxic substance. These may include transport of the toxic substance away from sensitive areas and to compartments or complexes whose purpose is sequestratio |
| 7 / 464 | "Any process that results in a change in state or activity of a cell or an organism (in terms of movement, secretion, enzyme production, gene expression, etc.) as a result of a stimulus. The process begins with detectio |
| 7 / 463 | "The production of new individuals that contain some portion of genetic material inherited from one or more parent organisms." [GOC:go_curators, GOC:isa_complete, GOC:jl, ISBN:0198506732] |
| 7 / 463 | "Any process involved in the development or functioning of the immune system, an organismal system for calibrated responses to potential internal or invasive threats." [GO_REF:0000022, GOC:add] |

Supplementary Data S3.3.8

### Timepoint 8

#### Molecular Function

| | |
|---|---|
| 346 / 3521 | "A molecular function characterized by the coupling of ATP hydrolysis to other steps of a reaction mechanism to make the reaction energetically favorable, for example to catalyze a reaction or drive transport aga |
| 345 / 3514 | "The action of a molecule that contributes to the structural integrity of a complex or its assembly within or outside a cell." [GOC:mah, GOC:vw] |
| 5 / 330 | "Binding to an RNA molecule or a portion thereof." [GOC:jl, GOC:mah] |
| 18 / 510 | "Any molecular function by which a gene product interacts selectively and non-covalently with DNA (deoxyribonucleic acid)." [GOC:dph, GOC:jl, GOC:tb, GOC:vw] |
| 19 / 515 | "Binding to a nucleic acid." [GOC:jl] |
| 5 / 294 | "Binding to a purine ribonucleoside triphosphate, a compound consisting of a purine base linked to a ribose sugar esterified with triphosphate on the sugar." [GOC:BHF, GOC:ebc, ISBN:0198506732] |
| 3 / 255 | "Binding to an adenyl ribonucleotide, any compound consisting of adenosine esterified with (ortho)phosphate or an oligophosphate at any hydroxyl group on the ribose moiety." [GOC:mah] |
| 120 / 632 | "Catalysis of an oxidation-reduction (redox) reaction, a reversible chemical reaction in which the oxidation state of an atom or atoms within a molecule is altered. One substrate acts as a hydrogen or electron don |
| 3 / 247 | "Binding to ATP, adenosine 5'-triphosphate, a universally important coenzyme and enzyme regulator." [ISBN:0198506732] |
| 348 / 3464 | "The selective, non-covalent, often stoichiometric, interaction of a molecule with one or more specific sites on another molecule." [GOC:ceb, GOC:mah, ISBN:0198506732] |

#### Biological Process

| | |
|---|---|
| 72 / 372 | "A process in which an ion is transported across a membrane." [GOC:mah] |
| 71 / 369 | "The process in which an organomercurial compound is transported across a membrane. Organomercurial substances are any organic compound containing a mercury atom." [GOC:ai, PMID:18793329] |
| 71 / 369 | "The directed movement of bicyclomycin across a lipid bilayer, from one side of a membrane to the other. Bicyclomycin (or bicozamycin) is an antibacterial drug often used as a livestock feed additive." [PMID:20067 |
| 71 / 369 | "The process in which carbon dioxide (CO2) is transported across a membrane." [GOC:yaf] |
| 71 / 369 | "The process in which urea, the water-soluble compound H2N-CO-NH2, is transported from one side of a membrane to the other by means of some agent such as a transporter or pore." [GOC:mah] |
| 71 / 369 | "The process in which a pyrimidine-containing compound is transported across a membrane. A pyrimidine-containing compound is any compound that contains pyrimidine or a formal derivative thereof." [GOC:mah] |
| 71 / 369 | "The directed movement of auxins from endoplasmic reticulum to cytosol." [PMID:19506555] |
| 71 / 369 | "The process in which hydrogen peroxide is transported across a membrane." [GOC:tb] |
| 71 / 369 | "The directed movement of a substance across the outer membrane in cells with two membranes." [PMID:15968039] |
| 71 / 369 | "The directed movement of benomyl across a membrane." [GOC:TermGenie] |

#### Cellular Compartment

| | |
|---|---|
| 9 / 503 | "The chemical reactions and pathways, including anabolism and catabolism, by which living organisms transform chemical substances. Metabolic processes typically transform small molecules, but also include macr |
| 9 / 494 | "A biological process represents a specific objective that the organism is genetically programmed to achieve. Biological processes are often described by their outcome or ending state, e.g., the biological process of c |
| 10 / 503 | "Any process that is carried out at the cellular level, but not necessarily restricted to a single cell. For example, cell communication occurs among more than one cell, but occurs at the cellular level." [GOC:go_curato |
| 9 / 476 | "The increase in size or mass of an entire organism, a part of an organism or a cell." [GOC:bf, GOC:ma] |
| 9 / 466 | "A biological process that directly contributes to the process of producing new individuals by one or two organisms. The new individuals inherit some proportion of their genetic material from the parent or parents." |
| 9 / 466 | "The entirety of a process in which information is transmitted within a biological system. This process begins with an active signal and ends when a cellular response has been triggered." [GOC:mtg_signal, GOC:mtg_ |
| 9 / 466 | "Any process that reduces or removes the toxicity of a toxic substance. These may include transport of the toxic substance away from sensitive areas and to compartments or complexes whose purpose is sequestrati |
| 9 / 464 | "Any process that results in a change in state or activity of a cell or an organism (in terms of movement, secretion, enzyme production, gene expression, etc.) as a result of a stimulus. The process begins with detecti |
| 9 / 463 | "The production of new individuals that contain some portion of genetic material inherited from one or more parent organisms." [GOC:go_curators, GOC:isa_complete, GOC:jl, ISBN:0198506732] |
| 9 / 463 | "Any process involved in the development or functioning of the immune system, an organismal system for calibrated responses to potential internal or invasive threats." [GO_REF:0000022, GOC:add] |

Supplementary Data S3.3.9

### Timepoint 9

#### Molecular Function

| | |
|---|---|
| 363 / 3521 | "A molecular function characterized by the coupling of ATP hydrolysis to other steps of a reaction mechanism to make the reaction energetically favorable, for example to catalyze a reaction or drive transport aga |
| 363 / 3514 | "The action of a molecule that contributes to the structural integrity of a complex or its assembly within or outside a cell." [GOC:mah, GOC:vw] |
| 17 / 515 | "Binding to a nucleic acid." [GOC:jl] |
| 17 / 510 | "Any molecular function by which a gene product interacts selectively and non-covalently with DNA (deoxyribonucleic acid)." [GOC:dph, GOC:jl, GOC:tb, GOC:vw] |
| 4 / 294 | "Binding to a purine ribonucleoside triphosphate, a compound consisting of a purine base linked to a ribose sugar esterified with triphosphate on the sugar." [GOC:BHF, GOC:ebc, ISBN:0198506732] |
| 2 / 255 | "Binding to an adenyl ribonucleotide, any compound consisting of adenosine esterified with (ortho)phosphate or an oligophosphate at any hydroxyl group on the ribose moiety." [GOC:mah] |
| 2 / 247 | "Binding to ATP, adenosine 5'-triphosphate, a universally important coenzyme and enzyme regulator." [ISBN:0198506732] |
| 130 / 632 | "Catalysis of an oxidation-reduction (redox) reaction, a reversible chemical reaction in which the oxidation state of an atom or atoms within a molecule is altered. One substrate acts as a hydrogen or electron don |
| 365 / 3464 | "The selective, non-covalent, often stoichiometric, interaction of a molecule with one or more specific sites on another molecule." [GOC:ceb, GOC:mah, ISBN:0198506732] |
| 8 / 330 | "Binding to an RNA molecule or a portion thereof." [GOC:jl, GOC:mah] |

#### Biological Process

| | |
|---|---|
| 0 / 150 | "The cellular metabolic process in which a protein is formed, using the sequence of a mature mRNA or circRNA molecule to specify the sequence of amino acids in a polypeptide chain. Translation is mediated by the |
| 74 / 372 | "A process in which an ion is transported across a membrane." [GOC:mah] |
| 73 / 369 | "The process in which an organomercurial compound is transported across a membrane. Organomercurial substances are any organic compound containing a mercury atom." [GOC:ai, PMID:18793329] |
| 73 / 369 | "The directed movement of bicyclomycin across a lipid bilayer, from one side of a membrane to the other. Bicyclomycin (or bicozamycin) is an antibacterial drug often used as a livestock feed additive." [PMID:20067 |
| 73 / 369 | "The process in which carbon dioxide (CO2) is transported across a membrane." [GOC:yaf] |
| 73 / 369 | "The process in which urea, the water-soluble compound H2N-CO-NH2, is transported from one side of a membrane to the other by means of some agent such as a transporter or pore." [GOC:mah] |
| 73 / 369 | "The process in which a pyrimidine-containing compound is transported across a membrane. A pyrimidine-containing compound is any compound that contains pyrimidine or a formal derivative thereof." [GOC:mah] |
| 73 / 369 | "The directed movement of auxins from endoplasmic reticulum to cytosol." [PMID:19506555] |
| 73 / 369 | "The process in which hydrogen peroxide is transported across a membrane." [GOC:tb] |
| 73 / 369 | "The directed movement of a substance across the outer membrane in cells with two membranes." [PMID:15968039] |

#### Cellular Compartment

| | |
|---|---|
| 11 / 503 | "The chemical reactions and pathways, including anabolism and catabolism, by which living organisms transform chemical substances. Metabolic processes typically transform small molecules, but also include macr |
| 11 / 494 | "A biological process represents a specific objective that the organism is genetically programmed to achieve. Biological processes are often described by their outcome or ending state, e.g., the biological process of c |
| 12 / 503 | "Any process that is carried out at the cellular level, but not necessarily restricted to a single cell. For example, cell communication occurs among more than one cell, but occurs at the cellular level." [GOC:go_curato |
| 11 / 476 | "The increase in size or mass of an entire organism, a part of an organism or a cell." [GOC:bf, GOC:ma] |
| 11 / 466 | "A biological process that directly contributes to the process of producing new individuals by one or two organisms. The new individuals inherit some proportion of their genetic material from the parent or parents." |
| 11 / 466 | "The entirety of a process in which information is transmitted within a biological system. This process begins with an active signal and ends when a cellular response has been triggered." [GOC:mtg_signal, GOC:mtg_ |
| 11 / 466 | "Any process that reduces or removes the toxicity of a toxic substance. These may include transport of the toxic substance away from sensitive areas and to compartments or complexes whose purpose is sequestrati |
| 11 / 464 | "Any process that results in a change in state or activity of a cell or an organism (in terms of movement, secretion, enzyme production, gene expression, etc.) as a result of a stimulus. The process begins with detecti |
| 11 / 463 | "The production of new individuals that contain some portion of genetic material inherited from one or more parent organisms." [GOC:go_curators, GOC:isa_complete, GOC:jl, ISBN:0198506732] |
| [illegible] | [illegible] |

Supplementary Data S3.3.10

### Timepoint 10

#### Molecular Function

| | |
|---|---|
| 12 / 510 | "Any molecular function by which a gene product interacts selectively and non-covalently with DNA (deoxyribonucleic acid)." [GOC:dph, GOC:jl, GOC:tb, GOC:vw] |
| 13 / 515 | "Binding to a nucleic acid." [GOC:jl] |
| 385 / 3521 | "A molecular function characterized by the coupling of ATP hydrolysis to other steps of a reaction mechanism to make the reaction energetically favorable, for example to catalyze a reaction or drive transport aga |
| 385 / 3514 | "The action of a molecule that contributes to the structural integrity of a complex or its assembly within or outside a cell." [GOC:mah, GOC:vw] |
| 100 / 386 | "Catalysis of an oxidation-reduction (redox) reaction in which a CH-OH group act as a hydrogen or electron donor and reduces a hydrogen or electron acceptor." [GOC:ai] |
| 139 / 632 | "Catalysis of an oxidation-reduction (redox) reaction, a reversible chemical reaction in which the oxidation state of an atom or atoms within a molecule is altered. One substrate acts as a hydrogen or electron don |
| 92 / 355 | "Catalysis of the incorporation of one atom from molecular oxygen into a compound and the reduction of the other atom of oxygen to water." [ISBN:0198506732] |
| 7 / 330 | "Binding to an RNA molecule or a portion thereof." [GOC:jl, GOC:mah] |
| 94 / 367 | "Catalysis of an oxidation-reduction (redox) reaction in which hydrogen or electrons are transferred from each of two donors, and molecular oxygen is reduced or incorporated into a donor." [GOC:mah] |
| 92 / 358 | "Catalysis of an oxidation-reduction (redox) reaction in which an aldehyde or ketone (oxo) group acts as a hydrogen or electron donor and reduces a hydrogen or electron acceptor." [GOC:ai] |

#### Biological Process

| | |
|---|---|
| 90 / 431 | "OBSOLETE. A metabolic process that results in the removal or addition of one or more electrons to or from a substance, with or without the concomitant removal or addition of a proton or protons." [GOC:dhl, GOC |
| 0 / 150 | "The cellular metabolic process in which a protein is formed, using the sequence of a mature mRNA or circRNA molecule to specify the sequence of amino acids in a polypeptide chain. Translation is mediated by the |
| 12 / 336 | "Any process that modulates the frequency, rate or extent of transcription mediated by RNA polymerase II." [GOC:go_curators, GOC:txnOH] |
| 70 / 342 | "The chemical reactions and pathways resulting in many of the chemical changes of compounds that are not necessarily required for growth and maintenance of cells, and are often unique to a taxon. In multicellula |
| 74 / 372 | "A process in which an ion is transported across a membrane." [GOC:mah] |
| 74 / 373 | "The process in which an oligopeptide is transported across a membrane. Oligopeptides are molecules that contain a small number (2 to 20) of amino-acid residues connected by peptide linkages." [GOC:vw, ISBN:01 |
| 73 / 369 | "The process in which an organomercurial compound is transported across a membrane. Organomercurial substances are any organic compound containing a mercury atom." [GOC:ai, PMID:18793329] |
| 73 / 369 | "The directed movement of bicyclomycin across a lipid bilayer, from one side of a membrane to the other. Bicyclomycin (or bicozamycin) is an antibacterial drug often used as a livestock feed additive." [PMID:20067 |
| 73 / 369 | "The process in which carbon dioxide (CO2) is transported across a membrane." [GOC:yaf] |
| 73 / 369 | "The process in which urea, the water-soluble compound H2N-CO-NH2, is transported from one side of a membrane to the other by means of some agent such as a transporter or pore." [GOC:mah] |

#### Cellular Compartment

| | |
|---|---|
| 18 / 575 | "Organized structure of distinctive morphology and function, bounded by a single or double lipid bilayer membrane and occurring within the cell. Includes the nucleus, mitochondria, plastids, vacuoles, and vesicles. E |
| 13 / 494 | "A biological process represents a specific objective that the organism is genetically programmed to achieve. Biological processes are often described by their outcome or ending state, e.g., the biological process of c |
| 14 / 503 | "The chemical reactions and pathways, including anabolism and catabolism, by which living organisms transform chemical substances. Metabolic processes typically transform small molecules, but also include macr |
| 15 / 507 | "A stable assembly of two or more macromolecules, i.e. proteins, nucleic acids, carbohydrates or lipids, in which at least one component is a protein and the constituent parts function together in the nucleus." [GOC |
| 13 / 476 | "The increase in size or mass of an entire organism, a part of an organism or a cell." [GOC:bf, GOC:ma] |
| 15 / 503 | "Any process that is carried out at the cellular level, but not necessarily restricted to a single cell. For example, cell communication occurs among more than one cell, but occurs at the cellular level." [GOC:go_curato |
| 15 / 497 | "The volume enclosed by the nuclear inner membrane." [GOC:mah, GOC:pz] |
| 13 / 466 | "A biological process that directly contributes to the process of producing new individuals by one or two organisms. The new individuals inherit some proportion of their genetic material from the parent or parents." |
| 13 / 466 | "The entirety of a process in which information is transmitted within a biological system. This process begins with an active signal and ends when a cellular response has been triggered." [GOC:mtg_signal, GOC:mtg_ |
| 13 / 466 | "Any process that reduces or removes the toxicity of a toxic substance. These may include transport of the toxic substance away from sensitive areas and to compartments or complexes whose purpose is sequestrati |

Supplementary Data S3.3.11

### Timepoint 11

#### Molecular Function

| | |
|---|---|
| 13 / 510 | "Any molecular function by which a gene product interacts selectively and non-covalently with DNA (deoxyribonucleic acid)." [GOC:dph, GOC:jl, GOC:tb, GOC:vw] |
| 14 / 515 | "Binding to a nucleic acid." [GOC:jl] |
| 385 / 3521 | "A molecular function characterized by the coupling of ATP hydrolysis to other steps of a reaction mechanism to make the reaction energetically favorable, for example to catalyze a reaction or drive transport aga |
| 384 / 3514 | "The action of a molecule that contributes to the structural integrity of a complex or its assembly within or outside a cell." [GOC:mah, GOC:vw] |
| 92 / 355 | "Catalysis of the incorporation of one atom from molecular oxygen into a compound and the reduction of the other atom of oxygen to water." [ISBN:0198506732] |
| 94 / 367 | "Catalysis of an oxidation-reduction (redox) reaction in which hydrogen or electrons are transferred from each of two donors, and molecular oxygen is reduced or incorporated into a donor." [GOC:mah] |
| 92 / 358 | "Catalysis of an oxidation-reduction (redox) reaction in which an aldehyde or ketone (oxo) group acts as a hydrogen or electron donor and reduces a hydrogen or electron acceptor." [GOC:ai] |
| 90 / 348 | "Catalysis of the hydroxylation of sphingolipid long chain bases." [PMID:9556590] |
| 90 / 348 | "Catalysis of the conversion of a fatty acid to an alpha-hydroxylated fatty acid. A hydroxyl group is added to the second carbon, counted from the carboxyl end, of a fatty acid chain." [PMID:19054355] |
| 91 / 354 | "Catalysis of an oxidation-reduction (redox) reaction in which NADH or NADPH acts as a hydrogen or electron donor and reduces a hydrogen or electron acceptor." [GOC:ai] |

#### Biological Process

| | |
|---|---|
| 0 / 150 | "The cellular metabolic process in which a protein is formed, using the sequence of a mature mRNA or circRNA molecule to specify the sequence of amino acids in a polypeptide chain. Translation is mediated by the |
| 12 / 336 | "Any process that modulates the frequency, rate or extent of transcription mediated by RNA polymerase II." [GOC:go_curators, GOC:txnOH] |
| 86 / 431 | "OBSOLETE. A metabolic process that results in the removal or addition of one or more electrons to or from a substance, with or without the concomitant removal or addition of a proton or protons." [GOC:dhl, GOC |
| 10 / 305 | "Any process that modulates the frequency, rate or extent of cellular DNA-templated transcription." [GOC:go_curators, GOC:txnOH] |
| 76 / 372 | "A process in which an ion is transported across a membrane." [GOC:mah] |
| 76 / 373 | "The process in which an oligopeptide is transported across a membrane. Oligopeptides are molecules that contain a small number (2 to 20) of amino-acid residues connected by peptide linkages." [GOC:vw, ISBN:01 |
| 75 / 369 | "The process in which an organomercurial compound is transported across a membrane. Organomercurial substances are any organic compound containing a mercury atom." [GOC:ai, PMID:18793329] |
| 75 / 369 | "The directed movement of bicyclomycin across a lipid bilayer, from one side of a membrane to the other. Bicyclomycin (or bicozamycin) is an antibacterial drug often used as a livestock feed additive." [PMID:20067 |
| 75 / 369 | "The process in which carbon dioxide (CO2) is transported across a membrane." [GOC:yaf] |
| 75 / 369 | "The process in which urea, the water-soluble compound H2N-CO-NH2, is transported from one side of a membrane to the other by means of some agent such as a transporter or pore." [GOC:mah] |

#### Cellular Compartment

| | |
|---|---|
| 12 / 494 | "A biological process represents a specific objective that the organism is genetically programmed to achieve. Biological processes are often described by their outcome or ending state, e.g., the biological process of c |
| 13 / 503 | "The chemical reactions and pathways, including anabolism and catabolism, by which living organisms transform chemical substances. Metabolic processes typically transform small molecules, but also include macr |
| 18 / 575 | "Organized structure of distinctive morphology and function, bounded by a single or double lipid bilayer membrane and occurring within the cell. Includes the nucleus, mitochondria, plastids, vacuoles, and vesicles. E |
| 12 / 476 | "The increase in size or mass of an entire organism, a part of an organism or a cell." [GOC:bf, GOC:ma] |
| 14 / 503 | "Any process that is carried out at the cellular level, but not necessarily restricted to a single cell. For example, cell communication occurs among more than one cell, but occurs at the cellular level." [GOC:go_curato |
| 12 / 466 | "A biological process that directly contributes to the process of producing new individuals by one or two organisms. The new individuals inherit some proportion of their genetic material from the parent or parents." |
| 12 / 466 | "The entirety of a process in which information is transmitted within a biological system. This process begins with an active signal and ends when a cellular response has been triggered." [GOC:mtg_signal, GOC:mtg_ |
| 12 / 466 | "Any process that reduces or removes the toxicity of a toxic substance. These may include transport of the toxic substance away from sensitive areas and to compartments or complexes whose purpose is sequestrati |
| 12 / 464 | "Any process that results in a change in state or activity of a cell or an organism (in terms of movement, secretion, enzyme production, gene expression, etc.) as a result of a stimulus. The process begins with detecti |
| 12 / 463 | "The production of new individuals that contain some portion of genetic material inherited from one or more parent organisms." [GOC:go_curators, GOC:isa_complete, GOC:jl, ISBN:0198506732] |

Supplementary Data S3.3.12

### Timepoint 12

#### Molecular Function

| | |
|---|---|
| 385 / 3521 | "A molecular function characterized by the coupling of ATP hydrolysis to other steps of a reaction mechanism to make the reaction energetically favorable, for example to catalyze a reaction or drive transport ag |
| 384 / 3514 | "The action of a molecule that contributes to the structural integrity of a complex or its assembly within or outside a cell." [GOC:mah, GOC:vw] |
| 16 / 510 | "Any molecular function by which a gene product interacts selectively and non-covalently with DNA (deoxyribonucleic acid)." [GOC:dph, GOC:jl, GOC:tb, GOC:vw] |
| 18 / 515 | "Binding to a nucleic acid." [GOC:jl] |
| 96 / 358 | "Catalysis of an oxidation-reduction (redox) reaction in which an aldehyde or ketone (oxo) group acts as a hydrogen or electron donor and reduces a hydrogen or electron acceptor." [GOC:ai] |
| 94 / 348 | "Catalysis of the hydroxylation of sphingolipid long chain bases." [PMID:9556590] |
| 94 / 348 | "Catalysis of the conversion of a fatty acid to an alpha-hydroxylated fatty acid. A hydroxyl group is added to the second carbon, counted from the carboxyl end, of a fatty acid chain." [PMID:19054355] |
| 95 / 354 | "Catalysis of an oxidation-reduction (redox) reaction in which NADH or NADPH acts as a hydrogen or electron donor and reduces a hydrogen or electron acceptor." [GOC:ai] |
| 94 / 349 | "Catalysis of an oxidation-reduction (redox) reaction in which a CH-NH group acts as a hydrogen or electron donor and reduces a hydrogen or electron acceptor." [GOC:ai] |
| 95 / 355 | "Catalysis of the incorporation of one atom from molecular oxygen into a compound and the reduction of the other atom of oxygen to water." [ISBN:0198506732] |

#### Biological Process

| | |
|---|---|
| 89 / 431 | "OBSOLETE. A metabolic process that results in the removal or addition of one or more electrons to or from a substance, with or without the concomitant removal or addition of a proton or protons." [GOC:dhl, GOC |
| 0 / 150 | "The cellular metabolic process in which a protein is formed, using the sequence of a mature mRNA or circRNA molecule to specify the sequence of amino acids in a polypeptide chain. Translation is mediated by the |
| 74 / 372 | "A process in which an ion is transported across a membrane." [GOC:mah] |
| 74 / 373 | "The process in which an oligopeptide is transported across a membrane. Oligopeptides are molecules that contain a small number (2 to 20) of amino-acid residues connected by peptide linkages." [GOC:vw, ISBN:01 |
| 14 / 336 | "Any process that modulates the frequency, rate or extent of transcription mediated by RNA polymerase II." [GOC:go_curators, GOC:txnOH] |
| 73 / 369 | "The process in which an organomercurial compound is transported across a membrane. Organomercurial substances are any organic compound containing a mercury atom." [GOC:ai, PMID:18793329] |
| 73 / 369 | "The directed movement of bicyclomycin across a lipid bilayer, from one side of a membrane to the other. Bicyclomycin (or bicozamycin) is an antibacterial drug often used as a livestock feed additive." [PMID:20067 |
| 73 / 369 | "The process in which carbon dioxide (CO2) is transported across a membrane." [GOC:yaf] |
| 73 / 369 | "The process in which urea, the water-soluble compound H2N-CO-NH2, is transported from one side of a membrane to the other by means of some agent such as a transporter or pore." [GOC:mah] |
| 73 / 369 | "The process in which a pyrimidine-containing compound is transported across a membrane. A pyrimidine-containing compound is any compound that contains pyrimidine or a formal derivative thereof." [GOC:mah] |

#### Cellular Compartment

| | |
|---|---|
| 12 / 494 | "A biological process represents a specific objective that the organism is genetically programmed to achieve. Biological processes are often described by their outcome or ending state, e.g., the biological process of c |
| 13 / 503 | "The chemical reactions and pathways, including anabolism and catabolism, by which living organisms transform chemical substances. Metabolic processes typically transform small molecules, but also include macr |
| 13 / 503 | "Any process that is carried out at the cellular level, but not necessarily restricted to a single cell. For example, cell communication occurs among more than one cell, but occurs at the cellular level." [GOC:go_curato |
| 12 / 476 | "The increase in size or mass of an entire organism, a part of an organism or a cell." [GOC:bf, GOC:ma] |
| 12 / 466 | "A biological process that directly contributes to the process of producing new individuals by one or two organisms. The new individuals inherit some proportion of their genetic material from the parent or parents." |
| 12 / 466 | "The entirety of a process in which information is transmitted within a biological system. This process begins with an active signal and ends when a cellular response has been triggered." [GOC:mtg_signal, GOC:mtg_ |
| 12 / 466 | "Any process that reduces or removes the toxicity of a toxic substance. These may include transport of the toxic substance away from sensitive areas and to compartments or complexes whose purpose is sequestrati |
| 12 / 464 | "Any process that results in a change in state or activity of a cell or an organism (in terms of movement, secretion, enzyme production, gene expression, etc.) as a result of a stimulus. The process begins with detecti |
| 12 / 463 | "The production of new individuals that contain some portion of genetic material inherited from one or more parent organisms." [GOC:go_curators, GOC:isa_complete, GOC:jl, ISBN:0198506732] |
| 12 / 463 | "Any process involved in the development or functioning of the immune system, an organismal system for calibrated responses to potential internal or invasive threats." [GO_REF:0000022, GOC:add] |

#### Supplementary Data S3.4– Temporal GO Description of Transcriptomics

**Each one of the following pictures is an aggregation of outputs of a GO term enrichment analysis on the results of DEseq2 at each specific point along the transcriptomic time course.**

##### Supplementary Data S3.4.1

| GO_Term | P_val | Counts | MolFunc_Description |
|---|---|---|---|
| GO:0008327 | 0.003734252 | 7 / 211 | "Binding to a methylated cytosine/guanine dinucleotide." [GOC:jl, PMID:11746232] |
| GO:0045322 | 0.003992516 | 3 / 33 | "Binding to uan nmethylated CpG motif. Unmethylated CpG dinucleotides are often associated with gene promoters." [GOC:ai, PMID:10688657] |
| GO:0141094 | 0.003992516 | 3 / 33 | "Binding to a DNA loop anchor, a region of chromosome that defines the ends of a DNA loop where two parts of same DNA strand are held in close physical proximity. For example, in animals, the anchors of DNA loops are convergently oriented CTCF binding sites during interph |
| GO:0042162 | 0.005076913 | 3 / 36 | "Binding to a telomere, a specific structure at the end of a linear chromosome required for the integrity and maintenance of the end." [GOC:jl, SO:0000624] |
| GO:0008086 | 0.010142923 | 1 / 1 | "Enables the transmembrane transfer of a calcium ion by a voltage-gated channel that is activated in response to light. A voltage-gated channel is a channel whose open state is dependent on the voltage across the membrane in which it is embedded." [GOC:mtg_transport] |
| GO:0008331 | 0.010142923 | 1 / 1 | "Enables the transmembrane transfer of a calcium ion by a high voltage-gated channel. A high voltage-gated channel is a channel whose open state is dependent on high voltage across the membrane in which it is embedded." [GOC:mtg_transport, ISBN:0815340729, PMID:16382 |
| GO:0008332 | 0.010142923 | 1 / 1 | "Enables the transmembrane transfer of a calcium ion by a low voltage-gated channel. A low voltage-gated channel is a channel whose open state is dependent on low voltage across the membrane in which it is embedded." [GOC:mtg_transport, ISBN:0815340729, PMID:1638209 |
| GO:0022843 | 0.010142923 | 1 / 1 | "Enables the transmembrane transfer of a cation by a voltage-gated channel. A cation is a positively charged ion. A voltage-gated channel is a channel whose open state is dependent on the voltage across the membrane in which it is embedded." [GOC:mtg_transport, ISBN:0815 |
| GO:0047498 | 0.010142923 | 1 / 1 | "Catalysis of the reaction: phosphatidylcholine + H2O = 1-acylglycerophosphocholine + a carboxylate. This reaction requires Ca2+." [EC:3.1.1.4] |
| GO:0051575 | 0.010142923 | 1 / 1 | "Catalysis of the beta-elimination of the 5' deoxyribose-5-phosphate at an abasic site in DNA where a DNA-(apurinic or apyrimidinic site) lyase has already cleaved the C-O-P bond 3' to the apurinic or apyrimidinic site." [PMID:11251121, PMID:16120966] |
| GO:0086007 | 0.010142923 | 1 / 1 | "Enables the transmembrane transfer of a calcium ion by a voltage-gated channel across the plasma membrane of a cardiac muscle cell that contributes to the depolarization phase of an action potential. A voltage-gated channel is a channel whose open state is dependent on th |
| GO:0099511 | 0.010142923 | 1 / 1 | "Regulation of cytosolic calcium ion concentrations via the directed movement of calcium ions across the plasma-membrane into the cytosol via the action of a voltage-gated calcium ion channel." [GOC:dos] |
| GO:0099635 | 0.010142923 | 1 / 1 | "Positive regulation of presynaptic cytosolic calcium ion concentrations via the directed movement of calcium ions across the plasma-membrane into the cytosol via the action of voltage-gated calcium ion channels. This is the first step in synaptic transmission." [GOC:dos] |
| GO:1990028 | 0.010142923 | 1 / 1 | "Enables the transmembrane transfer of a calcium ion by an intermediate voltage-gated channel. An intermediate voltage-gated channel is a channel whose open state is dependent on intermediate voltage across the membrane in which it is embedded." [GOC:BHF, GOC:rl, PMID |
| GO:0098847 | 0.019324111 | 3 / 60 | "Binding to single-stranded DNA of a specific nucleotide composition." [PMID:9531483] |
| GO:0017128 | 0.020084718 | 1 / 2 | "Catalysis of the movement of phospholipids from one membrane bilayer leaflet to the other, by an ATP-independent mechanism." [GOC:cjm, PMID:20043909, PMID:20302864] |
| GO:0047499 | 0.020084718 | 1 / 2 | "Catalysis of the reaction: phosphatidylcholine + H2O = 1-acylglycerophosphocholine + a carboxylate. This reaction does not require Ca2+." [EC:3.1.1.4] |
| GO:0140110 | 0.020130555 | 42 / 367 | "A molecular function that controls the rate, timing and/or magnitude of gene transcription. The function of transcriptional regulators is to modulate gene expression at the transcription step so that they are expressed in the right cell at the right time and in the right amount th |
| GO:0016758 | 0.020729288 | 2 / 23 | "Catalysis of the transfer of a hexosyl group from one compound (donor) to another (acceptor)." [GOC:jl] |
| GO:1990837 | 0.02528736 | 3 / 67 | "Binding to double-stranded DNA of a specific nucleotide composition, e.g. GC-rich DNA binding, or with a specific sequence motif or type of DNA, e.g. promotor binding or rDNA binding." [GOC:dos, GOC:sl] |
| GO:0005544 | 0.029492137 | 2 / 28 | "Binding to a phospholipid, a class of lipids containing phosphoric acid as a mono- or diester, in the presence of calcium." [GOC:jl] |
| GO:0003847 | 0.029828308 | 1 / 3 | "Catalysis of the reaction: 2-acetyl-1-alkyl-sn-glycero-3-phosphocholine + H2O = 1-alkyl-sn-glycero-3-phosphocholine + acetate." [EC:3.1.1.47] |
| GO:0005245 | 0.029828308 | 1 / 3 | "Enables the transmembrane transfer of a calcium ion by a voltage-gated channel. A voltage-gated channel is a channel whose open state is dependent on the voltage across the membrane in which it is embedded." [GOC:mtg_transport, GOC:tb, ISBN:0815340729] |
| GO:0005274 | 0.029828308 | 1 / 3 | "Enables the transfer of a solute or solutes from one side of a membrane to the other according to the reaction: allantoin(out) + H+(out) = allantoin(in) + H+(in) by secondary active transport." [GOC:mtg_transport, ISBN:0815340729, TC:2.A.39.3.1] |
| GO:0005280 | 0.029828308 | 1 / 3 | "Enables the transfer of a solute or solutes from one side of a membrane to the other according to the reaction: amino acid(out) + H+(out) = amino acid(in) + H+(in)." [GOC:ai] |
| GO:0005366 | 0.029828308 | 1 / 3 | "Enables the transfer of a solute or solutes from one side of a membrane to the other according to the reaction: myo-inositol(out) + H+(out) = myo-inositol(in) + H+(in)." [TC:2.A.1.1.8] |
| GO:0008512 | 0.029828308 | 1 / 3 | "Enables the transfer of a solute or solutes from one side of a membrane to the other according to the reaction: sulfate(out) + H+(out) = sulfate(in) + H+(in)." [TC:2.A.53.-.-] |
| GO:0009671 | 0.029828308 | 1 / 3 | "Enables the transfer of a solute or solutes from one side of a membrane to the other according to the reaction: nitrate(out) + H+(out) = nitrate(in) + H+(in)." [GOC:mah, PMID:10066586, PMID:1990981] |
| GO:0009672 | 0.029828308 | 1 / 3 | "Enables the transfer of a solute or solutes from one side of a membrane to the other according to the reaction: auxin(out) + H+(out) = auxin(in) + H+(in)." [PMID:8688077] |
| GO:0015345 | 0.029828308 | 1 / 3 | "Enables the transfer of a solute or solutes from one side of a membrane to the other according to the reaction: ferric enterobactin(out) + H+(out) = ferric enterobactin(in) + H+(in)." [TC:2.A.1.16.2] |
| GO:0015346 | 0.029828308 | 1 / 3 | "Enables the transfer of a solute or solutes from one side of a membrane to the other according to the reaction: ferric triacetylfusarinine C(out) + H+(out) = ferric triacetylfusarinine C(in) + H+(in)." [TC:2.A.1.16.3] |
| GO:0015360 | 0.029828308 | 1 / 3 | "Enables the transfer of a solute or solutes from one side of a membrane to the other according to the reaction: acetate(out) + H+(out) = acetate(in) + H+(in)." [TC:2.A.44.4.1] |
| GO:0015387 | 0.029828308 | 1 / 3 | "Enables the transfer of a solute or solutes from one side of a membrane to the other according to the reaction: K+(out) + H+(out) = K+(in) + H+(in)." [TC:2.A.38.-.-] |
| GO:0015504 | 0.029828308 | 1 / 3 | "Enables the transfer of a solute or solutes from one side of a membrane to the other according to the reaction: cytosine(out) + H+(out) = cytosine(in) + H+(in)." [TC:2.A.39.1.1] |
| GO:0015506 | 0.029828308 | 1 / 3 | "Enables the transfer of a solute or solutes from one side of a membrane to the other according to the reaction: nucleoside(out) + H+(out) = nucleoside(in) + H+(in)." [TC:2.A.1.10.1, TC:2.A.41.1.1] |
| GO:0015531 | 0.029828308 | 1 / 3 | "Enables the transfer of a solute or solutes from one side of a membrane to the other according to the reaction: citrate(out) + H+(out) = citrate(in) + H+(in)." [TC:2.A.1.6.1] |
| GO:0015532 | 0.029828308 | 1 / 3 | "Enables the transfer of a solute or solutes from one side of a membrane to the other according to the reaction: alpha-ketoglutarate(out) + H+(out) = alpha-ketoglutarate(in) + H+(in)." [TC:2.A.1.6.2] |
| GO:0015533 | 0.029828308 | 1 / 3 | "Enables the transfer of a solute or solutes from one side of a membrane to the other according to the reaction: shikimate(out) + H+(out) = shikimate(in) + H+(in)." [TC:2.A.1.6.6] |
| GO:0015538 | 0.029828308 | 1 / 3 | "Enables the transfer of a solute or solutes from one side of a membrane to the other according to the reaction: sialate(out) + H+(out) = sialate(in) + H+(in)." [TC:2.A.1.12.1] |
| GO:0015540 | 0.029828308 | 1 / 3 | "Enables the transfer of a solute or solutes from one side of a membrane to the other according to the reaction: 3-hydroxyphenyl propionate(out) + H+(out) = 3-hydroxyphenyl propionate(in) + H+(in)." [TC:2.A.1.15.2] |
| GO:0015650 | 0.029828308 | 1 / 3 | "Enables the transfer of a solute or solutes from one side of a membrane to the other according to the reaction: lactate (out) + H+ (out) = lactate (in) + H+ (in)." [TC:2.A.14.1.1] |
| GO:0015652 | 0.029828308 | 1 / 3 | "Enables the transfer of a solute or solutes from one side of a membrane to the other according to the reaction: quaternary ammonium group(out) + H+(out) = quaternary ammonium group(in) + H+(in)." [GOC:ai] |
| GO:0016151 | 0.029828308 | 1 / 3 | "Binding to a nickel (Ni) cation." [GOC:ai] |
| GO:0034215 | 0.029828308 | 1 / 3 | "Enables the transfer of a solute or solutes from one side of a membrane to the other according to the reaction: thiamine(out) + H+(out) = thiamine(in) + H+(in)." [GOC:mah] |
| GO:0140211 | 0.029828308 | 1 / 3 | "Enables the transfer of a solute or solutes from one side of a membrane to the other according to the reaction: folic acid(out) + H+(out) = folic acid(in) + H+(in). The main folic acid symporter is the Proton-Coupled Folate Transporter (PCFT/SLC46A1), which has similar affinity for |
| GO:0005215 | 0.030473027 | 42 / 372 | "Enables the directed movement of substances (such as macromolecules, small molecules, ions) into, out of or within a cell, or between cells." [GOC:ai, GOC:dgf] |
| GO:0003774 | 0.036352776 | 41 / 362 | "Generation of force resulting in movement, for example along a microfilament or microtubule, or in torque resulting in membrane scission or rotation of a flagellum. The energy required is obtained either from the hydrolysis of a nucleoside triphosphate or by an electrochemic |
| GO:0008970 | 0.039376577 | 1 / 4 | "Catalysis of the reaction: phosphatidylcholine + H2O = 2-acylglycerophosphocholine + a carboxylate." [EC:3.1.1.32] |
| GO:0015250 | 0.039376577 | 1 / 4 | "Transport systems of this type enable facilitated diffusion of water (by an energy-independent process) by passage through a transmembrane aqueous pore or channel without evidence for a carrier-mediated mechanism." [GOC:mtg_transport, ISBN:0815340729] |
| GO:0015254 | 0.039376577 | 1 / 4 | "Enables the facilitated diffusion of glycerol (by an energy-independent process) involving passage through a transmembrane aqueous pore or channel without evidence for a carrier-mediated mechanism." [GOC:mtg_transport, ISBN:0815340729] |

| GO_Term | P_val | Counts | BioProc_Description |
|---|---|---|---|
| GO:0017062 | 0.001272452 | 2 / 5 | "The aggregation, arrangement and bonding together of a set of components to form the cytochrome bc(1) complex, a transmembrane lipoprotein complex that it catalyzes the reduction of cytochrome c by accepting reducing equivalents from Coenzyme Q, by the aggregation, a |
| GO:0034551 | 0.001272452 | 2 / 5 | "The aggregation, arrangement and bonding together of a set of components to form the cytochrome bc(1) complex (also known as ubiquinol-cytochrome c reductase), in the mitochondrial inner membrane." [GOC:dgf, GOC:mcc] |
| GO:0010960 | 0.011593488 | 1 / 1 | "Any process involved in the maintenance of an internal steady state of magnesium ions within an organism or cell." [GOC:dph, GOC:tb] |
| GO:0010961 | 0.011593488 | 1 / 1 | "A homeostatic process involved in the maintenance of a steady state level of magnesium ions within a cell." [GOC:dph, GOC:tb] |
| GO:1902657 | 0.011593488 | 1 / 1 | "A process in which a protein is transported to, or maintained in, a location within a prospore membrane." [GO_REF:0000087, GOC:dph, GOC:TermGenie, PMID:24036347] |
| GO:0008152 | 0.021380691 | 42 / 390 | "The chemical reactions and pathways, including anabolism and catabolism, by which living organisms transform chemical substances. Metabolic processes typically transform small molecules, but also include macromolecular processes such as DNA repair and replication, and pr |
| GO:0000050 | 0.022923813 | 1 / 2 | "The sequence of reactions by which arginine is synthesized from ornithine, then cleaved to yield urea and regenerate ornithine. The overall reaction equation is NH3 + CO2 + aspartate + 3 ATP + 2 H2O = urea + fumarate + 2 ADP + 2 phosphate + AMP + diphosphate." [GOC:pde, G |
| GO:0003052 | 0.022923813 | 1 / 2 | "Any process in which an organism modulates its blood pressure at different values with a regularity of approximately 24 hours." [GOC:mtg_cardio, GOC:rl] |
| GO:0003053 | 0.022923813 | 1 / 2 | "Any process in which an organism modulates its heart rate at different values with a regularity of approximately 24 hours." [GOC:mtg_cardio, GOC:rl] |
| GO:0006685 | 0.022923813 | 1 / 2 | "The chemical reactions and pathways resulting in the breakdown of sphingomyelin, N-acyl-4-sphingenyl-1-O-phosphorylcholine." [ISBN:0198506732] |
| GO:0010617 | 0.022923813 | 1 / 2 | "Any process that modulates the concentration of cytosolic free calcium ion [Ca2+]cyt with a regularity of approximately 24 hours." [PMID:17982000] |
| GO:0015793 | 0.022923813 | 1 / 2 | "The directed movement of glycerol across a membrane. Glycerol is 1,2,3-propanetriol, a sweet, hygroscopic, viscous liquid, widely distributed in nature as a constituent of many lipids." [GOC:ai] |
| GO:0045016 | 0.022923813 | 1 / 2 | "The process in which a magnesium ion (Mg2+) is transported across a mitochondrial membrane, into or out of the mitochondrion." [GOC:ai, PMID:11254124] |
| GO:0045339 | 0.022923813 | 1 / 2 | "The chemical reactions and pathways resulting in the breakdown of farnesyl diphosphate." [GOC:go_curators] |
| GO:0046434 | 0.022923813 | 1 / 2 | "The chemical reactions and pathways resulting in the breakdown of organophosphates, any phosphate-containing organic compound." [GOC:ai] |
| GO:0048512 | 0.022923813 | 1 / 2 | "The specific behavior of an organism that recurs with a regularity of approximately 24 hours." [GOC:bf, GOC:go_curators, GOC:pr] |
| GO:0051874 | 0.022923813 | 1 / 2 | "The chemical reactions and pathways resulting in the breakdown of sphinganine-1-phosphate, the phosphorylated derivative of D-erythro-2-amino-1,3-octadecanediol." [GOC:ai] |
| GO:0060086 | 0.022923813 | 1 / 2 | "Any homeostatic process in which an organism modulates its internal body temperature at different values with a regularity of approximately 24 hours." [GOC:dph, GOC:tb] |
| GO:0070146 | 0.022923813 | 1 / 2 | "The process of coupling aspartate to aspartyl-tRNA in a mitochondrion, catalyzed by aspartyl-tRNA synthetase. In tRNA aminoacylation, the amino acid is first activated by linkage to AMP and then transferred to either the 2'- or the 3'-hydroxyl group of the 3'-adenosine residue |
| GO:1901948 | 0.022923813 | 1 / 2 | "The chemical reactions and pathways resulting in the breakdown of 5alpha,9alpha,10beta-labda-8(20),13-dien-15-yl diphosphate." [GOC:TermGenie, PMID:22027823] |
| GO:1902242 | 0.022923813 | 1 / 2 | "The chemical reactions and pathways resulting in the breakdown of copal-8-ol diphosphate(3-)." [GOC:TermGenie, PMID:22672125] |
| GO:1902247 | 0.022923813 | 1 / 2 | "The chemical reactions and pathways resulting in the breakdown of geranylgeranyl diphosphate." [GOC:TermGenie, PMID:22672125] |
| GO:1905372 | 0.022923813 | 1 / 2 | "The chemical reactions and pathways resulting in the breakdown of ceramide phosphoethanolamine." [GO_REF:0000068, GOC:hjd, GOC:TermGenie, PMID:25667419] |
| GO:0034220 | 0.023796375 | 9 / 403 | "A process in which a monoatomic ion is transported across a membrane. Monatomic ions (also called simple ions) are ions consisting of exactly one atom." [GOC:mah] |
| GO:0055085 | 0.029962123 | 9 / 423 | "The process in which a solute is transported across a lipid bilayer, from one side of a membrane to the other." [GOC:dph, GOC:jid] |
| GO:0007623 | 0.033995358 | 1 / 3 | "Any biological process in an organism that recurs with a regularity of approximately 24 hours." [GOC:bf, GOC:go_curators] |
| GO:0010248 | 0.033995358 | 1 / 3 | "The directed movement of ions to establish or maintain an electrochemical gradient across a membrane by means of some agent such as a transporter or pore." [GOC:mah, GOC:sm] |
| GO:0019623 | 0.033995358 | 1 / 3 | "The chemical reactions and pathways resulting in the breakdown of atrazine, a triazine ring-containing herbicide, into urea." [GOC:jl] |
| GO:0019627 | 0.033995358 | 1 / 3 | "The chemical reactions and pathways involving urea, the water soluble compound O=C-(NH2)2." [ISBN:0198506732] |
| GO:0030259 | 0.033995358 | 1 / 3 | "Covalent attachment of a glycosyl residue to a lipid molecule." [GOC:mah] |
| GO:0043419 | 0.033995358 | 1 / 3 | "The chemical reactions and pathways resulting in the breakdown of urea, the water soluble compound O=C-(NH2)2." [GOC:jl] |
| GO:0009395 | 0.034904991 | 2 / 27 | "The chemical reactions and pathways resulting in the breakdown of phospholipids, any lipid containing phosphoric acid as a mono- or diester." [ISBN:0198506732] |
| GO:0006811 | 0.041823935 | 2 / 30 | "The directed movement of a monoatomic ion into, out of or within a cell, or between cells, by means of some agent such as a transporter or pore. Monatomic ions (also called simple ions) are ions consisting of exactly one atom." [GOC:ai] |
| GO:0036294 | 0.044812444 | 1 / 4 | "Any process that results in a change in state or activity of a cell (in terms of movement, secretion, enzyme production, gene expression, etc.) as a result of a stimulus reflecting a decline in the level of oxygen." [GOC:al] |
| GO:0042542 | 0.044812444 | 1 / 4 | "Any process that results in a change in state or activity of a cell or an organism (in terms of movement, secretion, enzyme production, gene expression, etc.) as a result of a hydrogen peroxide (H2O2) stimulus." [GOC:jl] |
| GO:0061692 | 0.044812444 | 1 / 4 | "Any process that reduces or removes the toxicity of hydrogen peroxide in a cell. These include transport of hydrogen peroxide away from sensitive areas and to compartments or complexes whose purpose is sequestration." [GOC:dph, GOC:vw] |
| GO:0071456 | 0.044812444 | 1 / 4 | "Any process that results in a change in state or activity of a cell (in terms of movement, secretion, enzyme production, gene expression, etc.) as a result of a stimulus indicating lowered oxygen tension. Hypoxia, defined as a decline in O2 levels below normoxic levels of 20.8 - 20. |
| GO:0097167 | 0.044812444 | 1 / 4 | "Any process that modulates the frequency, rate or extent of mRNA translation with a regularity of approximately 24 hours." [GOC:ans, PMID:17264215] |
| GO:0140329 | 0.044812444 | 1 / 4 | "The movement of a lysophospholipid molecule from one leaflet of a membrane bilayer to the opposite leaflet." [PMID:15661733] |
| GO:1990144 | 0.044812444 | 1 / 4 | "The series of molecular signals in which an intracellular signal is conveyed to trigger the apoptotic death of a cell. The pathway is induced in response to hypoxia (lowered oxygen tension). Hypoxia, defined as a decline in O2 levels below normoxic levels of 20.8 - 20.95%, results |
| GO:1901564 | 0.045954775 | 3 / 76 | "The chemical reactions and pathways involving organonitrogen compound." [GOC:pr, GOC:TermGenie] |
| GO:0001407 | 0.048837625 | 8 / 400 | "The process in which a glycerophosphodiester is transported across a membrane. Glycerophosphodiesters are small molecules composed of glycerol-3-phosphate and an alcohol, for example, glycerophosphoinositol." [GOC:mcc, PMID:12912892] |
| GO:0015794 | 0.048837625 | 8 / 400 | "The process in which glycerol-3-phosphate is transported across a membrane. Glycerol-3-phosphate is a phosphoric monoester of glycerol." [GOC:ai] |
| GO:0015897 | 0.048837625 | 8 / 400 | "The process in which an organomercurial compound is transported across a membrane. Organomercurial substances are any organic compound containing a mercury atom." [GOC:ai, PMID:18793329] |
| GO:0015904 | 0.048837625 | 8 / 400 | "The directed movement of tetracycline from one side of a membrane to the other. Tetracycline is a broad spectrum antibiotic that blocks binding of aminoacyl tRNA to the ribosomes of both Gram-positive and Gram-negative organisms (and those of organelles)." [GOC:curators |
| GO:0015905 | 0.048837625 | 8 / 400 | "The directed movement of bicyclomycin across a lipid bilayer, from one side of a membrane to the other. Bicyclomycin (or bicozamycin) is an antibacterial drug often used as a livestock feed additive." [PMID:20067529] |
| GO:0035378 | 0.048837625 | 8 / 400 | "The process in which carbon dioxide (CO2) is transported across a membrane." [GOC:yaf] |
| GO:0035436 | 0.048837625 | 8 / 400 | "The process in which triose phosphate (glyceraldehyde 3-phosphate) is transported across a membrane. Glyceraldehyde 3-phosphate is any organic three carbon compound phosphate ester." [GOC:bf, ISBN:0198506732] |
| GO:0070811 | 0.048837625 | 8 / 400 | "The process in which glycerol-2-phosphate is transported across a membrane. Glycerol-2-phosphate is a phosphoric monoester of glycerol." [GOC:mah] |
| GO:0071918 | 0.048837625 | 8 / 400 | "The process in which urea, the water-soluble compound H2N-CO-NH2, is transported from one side of a membrane to the other by means of some agent such as a transporter or pore." [GOC:mah] |

| GO_Term | P_val | Counts | CellComp_Description |
|---|---|---|---|
| GO:0060194 | 0.010481366 | 1 / 1 | "Any process that modulates the frequency, rate or extent of the synthesis of antisense RNA, an RNA molecule complementary in sequence to another RNA or DNA molecule, which, by binding the latter, acts to inhibit its function and/or completion of synthesis, on a template |
| GO:0140472 | 0.010481366 | 1 / 1 | "The region directly beneath the plasma membrane at the cell tip at which no growth takes place." [PMID:17895368] |
| GO:1902716 | 0.010481366 | 1 / 1 | "Any cell cortex that is part of a growing cell tip." [GO_REF:0000064, GOC:TermGenie, PMID:24146635] |
| GO:0005886 | 0.010864618 | 6 / 192 | "The membrane surrounding a cell that separates the cell from its external environment. It consists of a phospholipid bilayer and associated proteins." [ISBN:0716731363] |
| GO:0098590 | 0.011104701 | 6 / 193 | "A membrane that is a (regional) part of the plasma membrane." [GOC:dos] |
| GO:0016020 | 0.013972861 | 6 / 204 | "A lipid bilayer along with all the proteins and protein complexes embedded in it and attached to it." [GOC:dos, GOC:mah, ISBN:0815316194] |
| GO:0060195 | 0.020747042 | 1 / 2 | "Any process that decreases the frequency, rate or extent of the synthesis of antisense RNA, an RNA molecule complementary in sequence to another RNA or DNA molecule, which, by binding the latter, acts to inhibit its function and/or completion of synthesis, on a template c |
| GO:0065004 | 0.020747042 | 1 / 2 | "The aggregation, arrangement and bonding together of proteins and DNA molecules to form a protein-DNA complex." [GOC:jl] |
| GO:1903145 | 0.020747042 | 1 / 2 | "Any actin filament that is part of a cell cortex of cell tip." [GO_REF:0000064, GOC:TermGenie, PMID:20807799, PMID:24954052] |
| GO:0005905 | 0.041458763 | 4 / 138 | "A part of the endomembrane system in the form of an invagination of a membrane upon which a clathrin coat forms, and that can be converted by vesicle budding into a clathrin-coated vesicle. Coated pits form on the plasma membrane, where they are involved in receptor-m |
| GO:0034357 | 0.041458763 | 4 / 138 | "A membrane enriched in complexes formed of reaction centers, accessory pigments and electron carriers, in which photosynthetic reactions take place." [GOC:ds, GOC:mah] |
| GO:0036362 | 0.041458763 | 4 / 138 | "A double layer of lipid molecules that surrounds an ascus, a capsule containing the sexual spores in some fungi." [GOC:mcc, GOC:vw, PMID:21900489] |
| GO:0070469 | 0.041458763 | 4 / 138 | "The protein complexes that form the electron transport system (the respiratory chain), associated with a cell membrane, usually the plasma membrane (in prokaryotes) or the inner mitochondrial membrane (on eukaryotes). The respiratory chain complexes transfer electrons f |
| GO:0097478 | 0.041458763 | 4 / 138 | "Any of the two layers of lipid molecules that constitute a membrane." [GOC:cjm] |
| GO:0098552 | 0.041458763 | 4 / 138 | "A cellular component consisting of one leaflet of a membrane bilayer and any proteins embedded or anchored in it or attached to its surface." [GOC:dos] |
| GO:0140221 | 0.041458763 | 4 / 138 | "Host-derived membrane of a pathogen-containing vacuole." [PMID:10560000, PMID:26842840] |
| GO:0140549 | 0.041458763 | 4 / 138 | "The membrane surrounding the spore core (endospore core) that separates it from its external environment." [PMID:23202530] |
| GO:0005642 | 0.042261491 | 4 / 139 | "Stacks of endoplasmic reticulum (ER) membranes containing a high density of nuclear pores, thought to form from excess nuclear membrane components, that have been described in a number of different cells. Annulate lamellar membranes are continuous with and embedde |
| GO:0019867 | 0.042261491 | 4 / 139 | "The external membrane of Gram-negative bacteria or certain organelles such as mitochondria and chloroplasts; freely permeable to most ions and metabolites." [GOC:go_curators] |
| GO:0034045 | 0.042261491 | 4 / 139 | "A cellular membrane associated with the phagophore assembly site." [GOC:mah, GOC:rph, PMID:16874040, PMID:17382324] |
| GO:0098857 | 0.042261491 | 4 / 139 | "A membrane region with a lipid composition that is distinct from that of the membrane regions that surround it." [PMID:20044567, PMID:26253820] |
| GO:0005628 | 0.043885875 | 4 / 141 | "The prospore membrane is a double-membraned structure that extends from the cytoplasmic face of the spindle pole bodies to encompass the spindle pole bodies and the four nuclear lobes that are formed during meiosis. It helps isolate the meiotic nuclei from the cytoplasr |
| GO:0048475 | 0.043885875 | 4 / 141 | "A single or double lipid bilayer with any of several different proteinaceous coats that can associate with membranes. Membrane coats include those formed by clathrin plus an adaptor complex, the COPI and COPII complexes." [GOC:jid] |
| GO:0070056 | 0.043885875 | 4 / 141 | "The region of the prospore membrane that extends to surround the spore nucleus; coated with specific proteins that are thought to play a role in prospore membrane organization." [GOC:mah, PMID:14702385] |
| GO:0019898 | 0.048052777 | 4 / 146 | "The component of a membrane consisting of gene products and protein complexes that are loosely bound to one of its surfaces, but not integrated into the hydrophobic region." [GOC:dos, GOC:jl, GOC:mah] |
| GO:0042175 | 0.056797948 | 4 / 156 | "The continuous network of membranes encompassing the nuclear outer membrane and the endoplasmic reticulum membrane." [GOC:bf, GOC:jl, GOC:mah, GOC:mcc, GOC:pr, GOC:vw] |
| GO:0098796 | 0.059516183 | 4 / 159 | "Any protein complex that is part of a membrane." [GOC:dos] |
| GO:0005872 | 0.059717524 | 1 / 6 | "Any complex that includes a dimer of molecules from the kinesin superfamily and any associated proteins, and moves towards the minus end of a microtubule." [GOC:mah] |
| GO:0005873 | 0.059717524 | 1 / 6 | "Any complex that includes a dimer of molecules from the kinesin superfamily and any associated proteins, and moves towards the plus end of a microtubule." [GOC:mah] |
| GO:0016938 | 0.059717524 | 1 / 6 | "A complex of two kinesin heavy chains and two kinesin light chains." [http://www.csfic.mi.cnr.it/centro/lines/8/ric.html] |
| GO:0016939 | 0.059717524 | 1 / 6 | "A complex consisting of two distinct motor subunits that form a heterodimer complexed with a third non-motor accessory subunit, the kinesin associated protein or KAP; the KIF3 heterodimer interacts via its C-terminal portion with KAP, which is thought to regulate the bindin |
| GO:0051285 | 0.059717524 | 1 / 6 | "The region directly beneath the plasma membrane at the cell tip. The cell tip is the region at either end of the longest axis of a cylindrical or elongated cell." [GOC:ai] |
| GO:0009987 | 0.062343753 | 2 / 508 | "Any process that is carried out at the cellular level, but not necessarily restricted to a single cell. For example, cell communication occurs among more than one cell, but occurs at the cellular level." [GOC:go_curators, GOC:isa_complete] |
| GO:0008152 | 0.063724704 | 2 / 505 | "The chemical reactions and pathways, including anabolism and catabolism, by which living organisms transform chemical substances. Metabolic processes typically transform small molecules, but also include macromolecular processes such as DNA repair and replication, and p |
| GO:0031090 | 0.06694477 | 4 / 167 | "A membrane that is one of the two lipid bilayers of an organelle envelope or the outermost membrane of single membrane bound organelle." [GOC:dos, GOC:mah] |
| GO:0008150 | 0.068015778 | 2 / 496 | "A biological process is the execution of a genetically-encoded biological module or program. It consists of all the steps required to achieve the specific biological objective of the module. A biological process is accomplished by a particular set of molecular functions carried out l |
| GO:0006334 | 0.06895289 | 1 / 7 | "The aggregation, arrangement and bonding together of a nucleosome, the beadlike structural units of eukaryotic chromatin composed of histones and DNA." [GOC:mah] |
| GO:0040007 | 0.077283578 | 2 / 478 | "The increase in size or mass of an entire organism, a part of an organism or a cell." [GOC:bf, GOC:ma] |
| GO:0006907 | 0.07799153 | 1 / 8 | "An endocytosis process that results in the uptake of liquid material by cells from their external environment; literally 'cell drinking'. Liquid is enclosed in vesicles, called pinosomes, formed by invagination of the plasma membrane." [ISBN:0198506732] |
| GO:0006909 | 0.07799153 | 1 / 8 | "A vesicle-mediated transport process that results in the engulfment of external particulate material by phagocytes and their delivery to the lysosome. The particles are initially contained within phagocytic vacuoles (phagosomes), which then fuse with primary lysosomes to effe |
| GO:0035037 | 0.07799153 | 1 / 8 | "An endocytosis process that results in penetration of the egg shell through the micropyle (a specialized anterior opening in the vitelline envelope) and entry of the entire sperm, including the surrounding plasma membrane and the sperm tail, into the egg cytoplasm. This step |
| GO:0072584 | 0.07799153 | 1 / 8 | "An endocytosis process that begins when material is taken up into plasma membrane caveolae, which then pinch off to form endocytic caveolar carriers." [GOC:BHF, GOC:mah, PMID:17318224, PMID:18498251, PMID:8970738, PMID:9234965] |
| GO:0075509 | 0.07799153 | 1 / 8 | "Any endocytosis that is involved in the uptake of a virus into a host cell." [GOC:bf, GOC:jl, VZ:977] |
| GO:0099051 | 0.07799153 | 1 / 8 | "The membrane scission process that is the final step in the formation of an endocytic vesicle: separation from the plasma membrane." [PMID:21779028] |
| GO:0120207 | 0.07799153 | 1 / 8 | "The process of selecting and or marking the position where endocytosis will occur." [GOC:vw, PMID:30044717] |
| GO:0140238 | 0.07799153 | 1 / 8 | "A vesicle-mediated transport process in which the presynapse take up external materials or membrane constituents by the invagination of a small region of the plasma membrane to form a new membrane-bounded vesicle." [PMID:24719103] |
| GO:0140239 | 0.07799153 | 1 / 8 | "A vesicle-mediated transport process in which the postsynapse take up external materials or membrane constituents by the invagination of a small region of the plasma membrane to form a new membrane-bounded vesicle." [PMID:12839988] |
| GO:0043231 | 0.080053951 | 3 / 579 | "Organized structure of distinctive morphology and function, bounded by a single or double lipid bilayer membrane and occurring within the cell. Includes the nucleus, mitochondria, plastids, vacuoles, and vesicles. Excludes the plasma membrane." [GOC:go_curators] |
| GO:0022414 | 0.082838827 | 2 / 468 | "A biological process that directly contributes to the process of producing new individuals by one or two organisms. The new individuals inherit some proportion of their genetic material from the parent or parents." [GOC:dph, GOC:isa_complete] |
| GO:0098754 | 0.082838827 | 2 / 468 | "Any process that reduces or removes the toxicity of a toxic substance. These may include transport of the toxic substance away from sensitive areas and to compartments or complexes whose purpose is sequestration of the toxic substance." [GOC:dos] |

#### Supplementary Data S3.4.2

| GO_Term | P_val | Counts | MolFunc_Description |
|---|---|---|---|
| GO:0004053 | 0.003027345 | 2 / 5 | "Catalysis of the reaction: L-arginine + H2O = L-ornithine + urea." [EC:3.5.3.1] |
| GO:0004668 | 0.003027345 | 2 / 5 | "Catalysis of the reaction: H2O + L-arginyl-[protein] = L-citrullyl-[protein] + NH4+, resulting in citrullination of the target protein. This reaction is calcium-dependent." [PMID:27393304] |
| GO:0008783 | 0.003027345 | 2 / 5 | "Catalysis of the reaction: agmatine + H2O = putrescine + urea." [EC:3.5.3.11, RHEA:13929] |
| GO:0009015 | 0.003027345 | 2 / 5 | "Catalysis of the reaction: N(2)-succinyl-L-arginine + 2 H2O + 2 H+ = N(2)-succinyl-L-ornithine + CO2 + 2 NH4." [EC:3.5.3.23, RHEA:19533] |
| GO:0016403 | 0.003027345 | 2 / 5 | "Catalysis of the reaction: N(G),N(G)-dimethyl-L-arginine + H2O = dimethylamine + L-citrulline." [EC:3.5.3.18] |
| GO:0016980 | 0.003027345 | 2 / 5 | "Catalysis of the reaction: creatine + H2O = sarcosine + urea." [EC:3.5.3.3, RHEA:22456] |
| GO:0016990 | 0.003027345 | 2 / 5 | "Catalysis of the reaction: L-arginine + H2O = L-citrulline + NH3." [EC:3.5.3.6] |
| GO:0018754 | 0.003027345 | 2 / 5 | "Catalysis of the reaction: ammelide + H2O = cyanuric acid + NH3." [PMID:1991731] |
| GO:0018755 | 0.003027345 | 2 / 5 | "Catalysis of the reaction: 2-chloro-4-hydroxy-6-amino-1,3,5-triazine + OH- = 2,4-dihydroxy-6-amino-1,3,5-triazine + Cl-." [UM-BBD_reactionID:r1414] |
| GO:0018756 | 0.003027345 | 2 / 5 | "Catalysis of the reaction: ammeline + H2O = ammelide + NH3." [PMID:1991731] |
| GO:0018757 | 0.003027345 | 2 / 5 | "Catalysis of the reaction: deisopropylhydroxyatrazine + H2O = NH3 + 2,4-dihydroxy-6-(N'-ethyl)amino-1,3,5-triazine." [UM-BBD_reactionID:r0121] |
| GO:0018758 | 0.003027345 | 2 / 5 | "Catalysis of the reaction: 2,4-dihydroxy-6-(N'-ethyl)amino-1,3,5-triazine + H2O = CH3CH2NH2 + cyanuric acid." [UM-BBD_reactionID:r0122] |
| GO:0033972 | 0.003027345 | 2 / 5 | "Catalysis of the reaction: amidinoproclavaminate + H2O = proclavaminate + urea." [EC:3.5.3.22, RHEA:17001] |
| GO:0043730 | 0.003027345 | 2 / 5 | "Catalysis of the reaction: 5-ureido-4-imidazole carboxylate + H2O = 5-amino-4-imidazole carboxylate + NH3 + CO2." [GOC:jl] |
| GO:0043920 | 0.003027345 | 2 / 5 | "Catalysis of the reaction: N1-aminopropylagmatine + H2O = spermidine + urea." [GOC:jl, PMID:15983049, RHEA:35827] |
| GO:0047632 | 0.003027345 | 2 / 5 | "Catalysis of the reaction: agmatine + H2O = N-carbamoylputrescine + NH3." [EC:3.5.3.12, MetaCyc:AGMATINE-DEIMINASE-RXN] |
| GO:0047652 | 0.003027345 | 2 / 5 | "Catalysis of the reaction: allantoate + H2O + H+ = CO2 + NH3 + ureidoglycine." [EC:3.5.3.9, MetaCyc:ALLANTOATE-DEIMINASE-RXN] |
| GO:0047660 | 0.003027345 | 2 / 5 | "Catalysis of the reaction: N-amidino-L-aspartate + H2O = L-aspartate + urea." [EC:3.5.3.14, RHEA:14849] |
| GO:0047817 | 0.003027345 | 2 / 5 | "Catalysis of the reaction: D-arginine + H2O = D-ornithine + urea." [EC:3.5.3.10, RHEA:12901] |
| GO:0047854 | 0.003027345 | 2 / 5 | "Catalysis of the reaction: 1,4-diguanidinobutane + H2O = agmatine + urea." [EC:3.5.3.20, RHEA:13597] |
| GO:0047970 | 0.003027345 | 2 / 5 | "Catalysis of the reaction: guanidinoacetate + H2O = glycine + urea." [EC:3.5.3.2, RHEA:23268] |
| GO:0047971 | 0.003027345 | 2 / 5 | "Catalysis of the reaction: 4-guanidinobutanoate + H2O = 4-aminobutanoate + urea." [EC:3.5.3.7, RHEA:19501] |
| GO:0047972 | 0.003027345 | 2 / 5 | "Catalysis of the reaction: 3-guanidinopropanoate + H2O = beta-alanine + urea." [EC:3.5.3.17, RHEA:16029] |
| GO:0050098 | 0.003027345 | 2 / 5 | "Catalysis of the reaction: H2O + methylguanidine = methylammonium + urea." [EC:3.5.3.16, RHEA:11764] |
| GO:0050414 | 0.003027345 | 2 / 5 | "Catalysis of the reaction: N-formimidoyl-L-aspartate + H2O = N-formyl-L-aspartate + NH3." [EC:3.5.3.5, MetaCyc:FORMIMINOASPARTATE-DEIMINASE-RXN] |
| GO:0050415 | 0.003027345 | 2 / 5 | "Catalysis of the reaction: N-formimidoyl-L-glutamate + H2O = L-glutamate + formamide." [EC:3.5.3.8, RHEA:22492] |
| GO:0050416 | 0.003027345 | 2 / 5 | "Catalysis of the reaction: N-formimidoyl-L-glutamate + H2O = N-formyl-L-glutamate + NH3." [EC:3.5.3.13, MetaCyc:FORMIMINOGLUTAMATE-DEIMINASE-RXN] |
| GO:0071522 | 0.003027345 | 2 / 5 | "Catalysis of the reaction: ureidoglycine + H2O = S-ureidoglycolate + NH3." [MetaCyc:URUR-RXN, PMID:19935661, PMID:20038185] |
| GO:0004037 | 0.004461369 | 2 / 6 | "Catalysis of the reaction: allantoate + H2O = (S)-ureidoglycolate + urea." [EC:3.5.3.4, RHEA:11016] |
| GO:0008373 | 0.004461369 | 2 / 6 | "Catalysis of the transfer of sialic acid to an acceptor molecule, typically the terminal portions of the sialylated glycolipids (gangliosides) or to the N- or O-linked sugar chains of glycoproteins." [GOC:cjm, PMID:26192491] |
| GO:0008920 | 0.004461369 | 2 / 6 | "Catalysis of the reaction: a lipopolysaccharide + ADP-L-glycero-beta-D-manno-heptose = a heptosylated lipopolysaccharide + ADP + H+." [MetaCyc:RXN0-5061, MetaCyc:RXN0-5122, MetaCyc:RXN0-5127] |
| GO:0008961 | 0.004461369 | 2 / 6 | "Catalysis of the transfer of the diacylglyceryl group from phosphatidylglycerol to the sulfhydryl group of the prospective N-terminal cysteine residue in an unmodified prolipoprotein." [PMID:8051048, RHEA:56712] |
| GO:0043842 | 0.004461369 | 2 / 6 | "Catalysis of the reactions: (KDO)-lipid IVA + CMP-3-deoxy-D-manno-octulosonate = KDO2-lipid IVA + CMP, and lipid IVA + CMP-3-deoxy-D-manno-octulosonate = (KDO)-lipid IVA + CMP." [PMID:1577828, PMID:2033061, PMID:9195966] |
| GO:0051075 | 0.00613634 | 2 / 7 | "Catalysis of the reaction: S-adenosylmethionine + 7-(aminomethyl)-7-deazaguanosine-tRNA = adenine + methionine + epoxyqueuosine-tRNA. 7-(aminomethyl)-7-deazaguanosine-tRNA is also known as preQ1-tRNA, and epoxyqueuosine-tRNA as oQ-tRNA." [PMID:12731872] |
| GO:0008199 | 0.007637368 | 4 / 46 | "Binding to a ferric iron ion, Fe(III)." [GOC:ai] |
| GO:0008194 | 0.008038213 | 2 / 8 | "Catalysis of the transfer of a glycosyl group from a UDP-sugar to a small hydrophobic molecule." [InterPro:IPR004224, PMID:11846783] |
| GO:0008198 | 0.009411969 | 4 / 49 | "Binding to a ferrous iron ion, Fe(II)." [GOC:ai] |
| GO:0005506 | 0.010053892 | 4 / 50 | "Binding to an iron (Fe) ion." [GOC:ai] |
| GO:0047424 | 0.010153449 | 2 / 9 | "Catalysis of the reaction: 2 H2O + methylenediurea = CO2 + 2 NH3 + N-hydroxymethylurea." [EC:3.5.3.21, MetaCyc:3.5.3.21-RXN, RHEA:15929] |
| GO:0036094 | 0.012468993 | 2 / 10 | "Binding to a small molecule, any low molecular weight, monomeric, non-encoded molecule." [GOC:curators, GOC:pde, GOC:pm] |
| GO:0043273 | 0.013167563 | 5 / 84 | "Catalysis of the reaction: CTP + H2O = CDP + H+ + phosphate. May or may not be coupled to another reaction." [RHEA:29387] |
| GO:0050339 | 0.013167563 | 5 / 84 | "Catalysis of the reaction: dTTP + H2O = dTDP + H+ + phosphate." [RHEA:19013] |
| GO:0103023 | 0.013167563 | 5 / 84 | "Catalysis of the reaction: ITP + H2O = IDP + H+ + phosphate." [GOC:pz, PMID:16216582, RHEA:28330] |
| GO:0004109 | 0.017980636 | 1 / 1 | "Catalysis of the reaction: coproporphyrinogen III + 2 H+ + O2 = 2 CO2 + 2 H2O + protoporphyrinogen IX." [EC:1.3.3.3, RHEA:18257] |
| GO:0004738 | 0.017980636 | 1 / 1 | "Catalysis of the oxidative decarboxylation of pyruvate." [ISBN:0716720094] |
| GO:0004758 | 0.017980636 | 1 / 1 | "Catalysis of the reaction: L-serine + H+ + palmitoyl-CoA = 3-dehydrosphinganine + CO2 + CoA." [EC:2.3.1.50, RHEA:14761] |
| GO:0004783 | 0.017980636 | 1 / 1 | "Catalysis of the reaction: hydrogen sulfide + 3 NADP+ + 3 H2O = sulfite + 3 NADPH + 3 H+." [RHEA:13801] |
| GO:0005353 | 0.017980636 | 1 / 1 | "Enables the transfer of fructose from one side of a membrane to the other. Fructose exists in a open chain form or as a ring compound. D-fructose is the sweetest of the sugars and is found free in a large number of fruits and honey." [GOC:ai, GOC:mtg_transport, ISBN:08153407 |
| GO:0015284 | 0.017980636 | 1 / 1 | "Enables the transfer of a solute or solutes from one side of a membrane to the other according to the reaction: fructose(out) = fructose(in)." [TC:2.A.1.1.13] |
| GO:0015578 | 0.017980636 | 1 / 1 | "Enables the transfer of mannose from one side of a membrane to the other. Mannose is the aldohexose manno-hexose, the C-2 epimer of glucose. The D-(+)-form is widely distributed in mannans and hemicelluloses and is of major importance in the core oligosaccharide of N-li |

| GO_Term | P_val | Counts | BioProc_Description |
|---|---|---|---|
| GO:0070813 | 0.000365532 | 2 / 2 | "The chemical reactions and pathways involving hydrogen sulfide, H2S." [GOC:mah] |
| GO:0000103 | 0.00211168 | 2 / 4 | "The pathways by which inorganic sulfate is processed and incorporated into sulfated compounds." [GOC:jl] |
| GO:0010134 | 0.00211168 | 2 / 4 | "The pathway by which inorganic sulfate is activated, reduced and incorporated into sulfated compounds, where the activated sulfate, adenylyl-sulfate, is reduced to sulfite by the activity of adenylyl-sulfate reductase." [EC:1.8.99.2] |
| GO:0019379 | 0.00211168 | 2 / 4 | "The pathway by which inorganic sulfate is processed and incorporated into sulfated compounds, where the phosphoadenylyl sulfate reduction step is catalyzed by the enzyme phosphoadenylyl-sulfate reductase (thioredoxin) (EC:1.8.4.8)." [GOC:jl] |
| GO:0017062 | 0.003453422 | 2 / 5 | "The aggregation, arrangement and bonding together of a set of components to form the cytochrome bc(1) complex, a transmembrane lipoprotein complex that it catalyzes the reduction of cytochrome c by accepting reducing equivalents from Coenzyme Q, by the aggregation, ar |
| GO:0019420 | 0.003453422 | 2 / 5 | "The reduction of sulfate to hydrogen sulfide, which acts as a terminal electron acceptor. Sulfate is activated to adenosine-phosphosulfate (APS) which is then reduced to sulfite, which is in turn reduced to hydrogen sulfide." [PMID:25400653, PMID:26680199] |
| GO:0034551 | 0.003453422 | 2 / 5 | "The aggregation, arrangement and bonding together of a set of components to form the cytochrome bc(1) complex (also known as ubiquinol-cytochrome c reductase), in the mitochondrial inner membrane." [GOC:dgf, GOC:mcc] |
| GO:0018868 | 0.005082902 | 2 / 6 | "The chemical reactions and pathways involving 2-aminobenzenesulfonate, aniline-o-sulfonic acid, an aromatic sulfonate used in organic synthesis and in the manufacture of various dyes and medicines." [UM-BBD_pathwayID:abs] |
| GO:0001666 | 0.006509159 | 3 / 21 | "Any process that results in a change in state or activity of a cell or an organism (in terms of movement, secretion, enzyme production, gene expression, etc.) as a result of a stimulus indicating lowered oxygen tension. Hypoxia, defined as a decline in O2 levels below normoxic lev |
| GO:1901564 | 0.01164057 | 5 / 76 | "The chemical reactions and pathways involving organonitrogen compound." [GOC:pr, GOC:TermGenie] |
| GO:0006213 | 0.014057471 | 3 / 28 | "The chemical reactions and pathways involving any pyrimidine nucleoside, one of a family of organic molecules consisting of a pyrimidine base covalently bonded to ribose (a ribonucleoside) or deoxyribose (a deoxyribonucleoside)." [GOC:jl, ISBN:0140512713] |
| GO:0009119 | 0.014057471 | 3 / 28 | "The chemical reactions and pathways involving any ribonucleoside, a nucleoside in which purine or pyrimidine base is linked to a ribose (beta-D-ribofuranose) molecule." [GOC:jl] |
| GO:0009120 | 0.014057471 | 3 / 28 | "The chemical reactions and pathways involving any one of a family of organic molecules consisting of a purine or pyrimidine base covalently bonded to a sugar deoxyribose (a deoxyribonucleoside)." [GOC:jl, ISBN:0140512713] |
| GO:0009163 | 0.014057471 | 3 / 28 | "The chemical reactions and pathways resulting in the formation of any one of a family of organic molecules consisting of a purine or pyrimidine base covalently bonded to a sugar ribose (a ribonucleoside) or deoxyribose (a deoxyribonucleoside)." [GOC:jl, ISBN:0140512713] |
| GO:0042278 | 0.014057471 | 3 / 28 | "The chemical reactions and pathways involving one of a family of organic molecules consisting of a purine base covalently bonded to a sugar ribose (a ribonucleoside) or deoxyribose (a deoxyribonucleoside)." [GOC:jl, ISBN:0140512713] |
| GO:0070637 | 0.014057471 | 3 / 28 | "The chemical reactions and pathways involving any pyridine nucleoside, a nucleoside in which a pyridine base covalently bonded to a sugar, usually ribose." [GOC:mah] |
| GO:1901657 | 0.015387395 | 3 / 29 | "The chemical reactions and pathways involving glycosyl compound." [GOC:pr, GOC:TermGenie] |
| GO:0009116 | 0.016778526 | 3 / 30 | "The chemical reactions and pathways involving a nucleoside, a nucleobase linked to either beta-D-ribofuranose (a ribonucleoside) or 2-deoxy-beta-D-ribofuranose, (a deoxyribonucleoside), e.g. adenosine, guanosine, inosine, cytidine, uridine and deoxyadenosine, deoxyguanosi |
| GO:0009164 | 0.016778526 | 3 / 30 | "The chemical reactions and pathways resulting in the breakdown of any one of a family of organic molecules consisting of a purine or pyrimidine base covalently bonded to a sugar ribose (a ribonucleoside) or deoxyribose (a deoxyribonucleoside)." [GOC:jl] |
| GO:0000096 | 0.016951427 | 2 / 11 | "The chemical reactions and pathways involving amino acids containing sulfur, comprising cysteine, homocysteine, methionine and selenocysteine." [GOC:ai] |
| GO:0006061 | 0.019240257 | 1 / 1 | "The chemical reactions and pathways resulting in the formation of sorbitol (D-glucitol), one of the ten stereoisomeric hexitols. It can be derived from glucose by reduction of the aldehyde group." [ISBN:0198506732] |
| GO:0009094 | 0.019240257 | 1 / 1 | "The chemical reactions and pathways resulting in the formation of L-phenylalanine, the L-enantiomer of 2-amino-3-phenylpropanoic acid, i.e. (2S)-2-amino-3-phenylpropanoic acid." [GOC:jsg, GOC:mah] |
| GO:0009732 | 0.019240257 | 1 / 1 | "The series of events in which a stimulus from a hexose is received and converted into a molecular signal." [GOC:sm] |
| GO:0009757 | 0.019240257 | 1 / 1 | "The series of molecular signals mediated by the detection of hexose." [GOC:sm] |
| GO:0010126 | 0.019240257 | 1 / 1 | "The chemical reactions and pathways involving mycothiol, which consists of N-acetyl-L-cysteine linked to a pseudodisaccharide, D-glucosamine and myo-inositol. Mycothiol is produced in actinomycetes like mycobacteria and serves similar functions to glutathione." [GOC:pz] |
| GO:0010619 | 0.019240257 | 1 / 1 | "An adenylate cyclase-activating G protein-coupled receptor signaling pathway initiated by glucose binding to its receptor on the surface of the target cell, and ending with the regulation of a downstream cellular process." [GOC:dph, GOC:signaling, GOC:tb] |
| GO:0015671 | 0.019240257 | 1 / 1 | "The directed movement of oxygen (O2) into, out of or within a cell, or between cells, by means of some agent such as a transporter or pore." [GOC:ai] |
| GO:0016143 | 0.019240257 | 1 / 1 | "The chemical reactions and pathways involving S-glycosides, any compound in which a glycosyl group has been substituted into a thiol group." [ISBN:0198506732] |
| GO:0018892 | 0.019240257 | 1 / 1 | "The chemical reactions and pathways involving cyclohexylsulfamate, also known as cyclamic acid. Sodium cyclohexylsulfamate (CHS-Na) was a widely used sweetening agent but was banned because of the suspicion of carcinogenicity and metabolic conversion to cyclohexylamine |
| GO:0018895 | 0.019240257 | 1 / 1 | "The chemical reactions and pathways involving dibenzothiophene, a substance composed of two benzene rings linked by one sulfide bond and one carbon-carbon bond. Dibenzothiophene derivatives can be detected in diesel oil following hydrodesulfurization treatment to rem |
| GO:0018907 | 0.019240257 | 1 / 1 | "The chemical reactions and pathways involving dimethyl sulfoxide, DMSO (C2H6OS), an alkyl sulfoxide that is practically odorless in its purified form. As a highly polar organic liquid, it is a powerful solvent. Its biological activities include the ability to penetrate plant and animal |
| GO:0018909 | 0.019240257 | 1 / 1 | "The chemical reactions and pathways involving dodecyl sulfate, commonly found as sodium dodecyl sulfate (SDS), a component of a variety of synthetic surfactants." [UM-BBD_pathwayID:dds] |
| GO:0018969 | 0.019240257 | 1 / 1 | "The chemical reactions and pathways involving thiocyanate, the anion of thiocyanic acid, a toxic cyanide derivative commonly formed as a by-product in the production of gas for fuel, coke, and substances for chemical industries." [GOC:jl] |
| GO:0018972 | 0.019240257 | 1 / 1 | "The chemical reactions and pathways involving toluene-4-sulfonate, the anion of 4-toluene sulfonic acid, a white crystalline solid which is highly hygroscopic and soluble in water." [GOC:ai] |
| GO:0018984 | 0.019240257 | 1 / 1 | "The chemical reactions and pathways involving naphthalenesulfonate, sulfonated derivatives of naphthalene." [GOC:ai] |
| GO:0019296 | 0.019240257 | 1 / 1 | "The chemical reactions and pathways involving coenzyme M (2-thioethansulfonate), a coenzyme involved in the utilization of methane by methanogenic prokaryotes." [GOC:go_curators] |
| GO:0019416 | 0.019240257 | 1 / 1 | "The chemical reactions and pathways resulting in the conversion of thiosulfate to tetrathionate, using cytochrome c as an electron acceptor." [MetaCyc:THIOSULFOX-PWY] |
| GO:0019417 | 0.019240257 | 1 / 1 | "The chemical reactions and pathways resulting the addition of oxygen to elemental sulfur." [GOC:jl, MetaCyc:FESULFOX-PWY, MetaCyc:SULFUROX-PWY] |
| GO:0019418 | 0.019240257 | 1 / 1 | "The chemical reactions and pathways resulting in the conversion of sulfide to elemental sulfur in a higher oxidation state, or to sulfite or sulfate." [MetaCyc:P222-PWY, MetaCyc:PWY-5274, MetaCyc:PWY-5285] |
| GO:0019422 | 0.019240257 | 1 / 1 | "The process in which sulfur compounds with an intermediate oxidation state serve as both electron donors and electron acceptors in an energy-generating redox process. The reaction takes place anaerobically, in light and in the absence of CO2." [MetaCyc:P203-PWY] |
| GO:0019473 | 0.019240257 | 1 / 1 | "The chemical reactions and pathways resulting in the breakdown of L-lysine into other compounds, including glutarate, by acetylation." [GOC:go_curators] |
| GO:0019477 | 0.019240257 | 1 / 1 | "The chemical reactions and pathways resulting in the breakdown of L-lysine, the L-enantiomer of (S)-2,6-diaminohexanoic acid." [GOC:go_curators, GOC:jsg, GOC:mah] |
| GO:0019694 | 0.019240257 | 1 / 1 | "The chemical reactions and pathways involving alkanesulfonates, the anion of alkanesulfonic acids, sulfonic acid derivatives containing an aliphatic hydrocarbon group." [GOC:ai] |
| GO:0030202 | 0.019240257 | 1 / 1 | "The chemical reactions and pathways involving heparin, any member of a group of glycosaminoglycans found mainly as an intracellular component of mast cells. They are similar to heparan sulfates but are of somewhat higher average Mr (6000-20000) and contain fewer N-acety |
| GO:0030204 | 0.019240257 | 1 / 1 | "The chemical reactions and pathways involving chondroitin sulfate, any member of a group of 10-60 kDa glycosaminoglycans, widely distributed in cartilage and other mammalian connective tissues, the repeat units of which consist of beta-(1,4)-linked D-glucuronyl beta-(1,3)-N- |
| GO:0030205 | 0.019240257 | 1 / 1 | "The chemical reactions and pathways involving dermatan sulfate, any of a group of glycosaminoglycans with repeats consisting of beta-(1,4)-linked L-iduronyl-beta-(1,3)-N-acetyl-D-galactosamine 4-sulfate units. They are important components of ground substance or intercellula |
| GO:0031162 | 0.019240257 | 1 / 1 | "The incorporation of exogenous sulfur into a metallo-sulfur cluster." [GOC:mah] |
| GO:0033477 | 0.019240257 | 1 / 1 | "The chemical reactions and pathways involving S-methyl-methionine (SMM). SMM can be converted to methionine by donating a methyl group to homocysteine, and concurrent operation of this reaction and that mediated by MMT sets up the SMM cycle." [GOC:mah, PMID:126 |
| GO:0033585 | 0.019240257 | 1 / 1 | "The chemical reactions and pathways resulting in the formation of L-phenylalanine from other compounds, including chorismate, via the intermediate phenylpyruvate." [GOC:go_curators] |
| GO:0033586 | 0.019240257 | 1 / 1 | "The chemical reactions and pathways resulting in the formation of L-phenylalanine from other compounds, including chorismate, via the intermediate L-arogenate." [GOC:go_curators] |

| GO_Term | P_val | Counts | CellComp_Description |
|---|---|---|---|
| GO:0008152 | 0.010162804 | 3 / 505 | "The chemical reactions and pathways, including anabolism and catabolism, by which living organisms transform chemical substances. Metabolic processes typically transform small molecules, but also include macromolecular processes such as DNA repair and replication, and pr |
| GO:0008150 | 0.011501493 | 3 / 496 | "A biological process is the execution of a genetically-encoded biological module or program. It consists of all the steps required to achieve the specific biological objective of the module. A biological process is accomplished by a particular set of molecular functions carried out b |
| GO:0005737 | 0.013348258 | 2 / 408 | "The contents of a cell excluding the plasma membrane and nucleus, but including other subcellular structures." [ISBN:0198547684] |
| GO:0040007 | 0.014671547 | 3 / 478 | "The increase in size or mass of an entire organism, a part of an organism or a cell." [GOC:bf, GOC:ma] |
| GO:0022414 | 0.016755544 | 3 / 468 | "A biological process that directly contributes to the process of producing new individuals by one or two organisms. The new individuals inherit some proportion of their genetic material from the parent or parents." [GOC:dph, GOC:isa_complete] |
| GO:0098754 | 0.016755544 | 3 / 468 | "Any process that reduces or removes the toxicity of a toxic substance. These may include transport of the toxic substance away from sensitive areas and to compartments or complexes whose purpose is sequestration of the toxic substance." [GOC:dos] |
| GO:0050896 | 0.017202917 | 3 / 466 | "Any process that results in a change in state or activity of a cell or an organism (in terms of movement, secretion, enzyme production, gene expression, etc.) as a result of a stimulus. The process begins with detection of the stimulus and ends with a change in state or activity or |
| GO:0000003 | 0.017430591 | 3 / 465 | "The production of new individuals that contain some portion of genetic material inherited from one or more parent organisms." [GOC:go_curators, GOC:isa_complete, GOC:jl, ISBN:0198506732] |
| GO:0002376 | 0.017430591 | 3 / 465 | "Any process involved in the development or functioning of the immune system, an organismal system for calibrated responses to potential internal or invasive threats." [GO_REF:0000022, GOC:add] |
| GO:0016032 | 0.017430591 | 3 / 465 | "A multi-organism process in which a virus is a participant. The other participant is the host. Includes infection of a host cell, replication of the viral genome, and assembly of progeny virus particles. In some cases the viral genetic material may integrate into the host genome and |
| GO:0032501 | 0.017430591 | 3 / 465 | "Any biological process, occurring at the level of a multicellular organism, pertinent to its function." [GOC:curators, GOC:dph, GOC:isa_complete, GOC:tb] |
| GO:0032502 | 0.017430591 | 3 / 465 | "A biological process whose specific outcome is the progression of an integrated living unit: an anatomical structure (which may be a subcellular structure, cell, tissue, or organ), or organism over time from an initial condition to a later condition." [GOC:isa_complete] |
| GO:0040011 | 0.017430591 | 3 / 465 | "Self-propelled movement of a cell or organism from one location to another." [GOC:dgh] |
| GO:0042592 | 0.017430591 | 3 / 465 | "Any biological process involved in the maintenance of an internal steady state." [GOC:jl, ISBN:0395825172] |
| GO:0043473 | 0.017430591 | 3 / 465 | "The accumulation of pigment in an organism, tissue or cell, either by increased deposition or by increased number of cells." [GOC:jl] |
| GO:0044419 | 0.017430591 | 3 / 465 | "Any process evolved to enable an interaction with an organism of a different species." [GOC:cc] |
| GO:0044848 | 0.017430591 | 3 / 465 | "A distinct period or stage in a biological process or cycle." [GOC:jl] |
| GO:0048511 | 0.017430591 | 3 / 465 | "Any process pertinent to the generation and maintenance of rhythms in the physiology of an organism." [GOC:jid] |
| GO:0051179 | 0.017430591 | 3 / 465 | "Any process in which a cell, a substance, or a cellular entity, such as a protein complex or organelle, is transported, tethered to or otherwise maintained in a specific location. In the case of substances, localization may also be achieved via selective degradation." [GOC:ai, GOC:d |
| GO:0051703 | 0.017430591 | 3 / 465 | "Any process in which an organism has an effect on an organism of the same species." [GOC:ai] |
| GO:0065007 | 0.017430591 | 3 / 465 | "Any process that modulates a measurable attribute of any biological process, quality or function." [GOC:dph, GOC:isa_complete, GOC:mah, GOC:pr, GOC:vw] |
| GO:0000338 | 0.018245342 | 1 / 1 | "The removal of a ubiquitin-like protein of the NEDD8 type from a protein." [GOC:krc] |
| GO:0009279 | 0.018245342 | 1 / 1 | "A lipid bilayer that forms the outermost membrane of the cell envelope; enriched in polysaccharide and protein; the outer leaflet of the membrane contains specific lipopolysaccharide structures." [GOC:md, GOC:mtg_sensu, ISBN:0135712254] |
| GO:0009337 | 0.018245342 | 1 / 1 | "A multisubunit iron flavoprotein, which in yeast is composed of 2 alpha and 2 beta subunits. Catalyzes the reduction of sulfite to sulfide." [BRENDA:1.8.1.2, GOC:jl] |
| GO:0043594 | 0.018245342 | 1 / 1 | "The outer membrane around a bacterial endospore, located between the endospore cortex and endospore coat." [GOC:mlg] |
| GO:0051666 | 0.018245342 | 1 / 1 | "Any process in which actin cortical patches are transported to, or maintained in, a specific location. An actin cortical patch is a discrete actin-containing structure found just beneath the plasma membrane in fungal cells." [GOC:mah] |
| GO:0140472 | 0.018245342 | 1 / 1 | "The region directly beneath the plasma membrane at the cell tip at which no growth takes place." [PMID:17895368] |
| GO:1902716 | 0.018245342 | 1 / 1 | "Any cell cortex that is part of a growing cell tip." [GO_REF:0000064, GOC:TermGenie, PMID:24146635] |
| GO:0005739 | 0.022922335 | 0 / 201 | "A semiautonomous, self replicating organelle that occurs in varying numbers, shapes, and sizes in the cytoplasm of virtually all eukaryotic cells. It is notably the site of tissue respiration." [GOC:giardia, ISBN:0198506732] |
| GO:0009987 | 0.024595356 | 4 / 508 | "Any process that is carried out at the cellular level, but not necessarily restricted to a single cell. For example, cell communication occurs among more than one cell, but occurs at the cellular level." [GOC:go_curators, GOC:isa_complete] |
| GO:0007120 | 0.035831853 | 1 / 2 | "The process of defining the next site of bud emergence adjacent to the last site of bud emergence on a budding cell." [GOC:clt] |
| GO:0008302 | 0.035831853 | 1 / 2 | "Recruitment and organization of actin filaments in female germline ring canals." [ISBN:0879694238] |
| GO:0030240 | 0.035831853 | 1 / 2 | "The aggregation, arrangement and bonding together of proteins to form the actin-based thin filaments of myofibrils in skeletal muscle." [GOC:ef, GOC:mah, GOC:mtg_muscle] |
| GO:0051639 | 0.035831853 | 1 / 2 | "The assembly of a network of actin filaments; actin filaments on different axes and with differing orientations are crosslinked together to form a mesh of filaments." [GOC:ai] |
| GO:0051764 | 0.035831853 | 1 / 2 | "The process in which two or more actin filaments are connected together by proteins that act as crosslinks between the filaments. The crosslinked filaments may be on the same or differing axes." [GOC:ai] |
| GO:0071689 | 0.035831853 | 1 / 2 | "The aggregation, arrangement and bonding together of proteins to form the actin-based thin filaments of myofibrils in striated muscle." [GOC:mah] |
| GO:0071691 | 0.035831853 | 1 / 2 | "The aggregation, arrangement and bonding together of proteins to form the actin-based thin filaments of myofibrils in cardiac muscle." [GOC:mah] |
| GO:0090135 | 0.035831853 | 1 / 2 | "The formation of daughter actin filament branches at an angle on the sides of preexisting mother filaments." [GOC:ascb_2009, GOC:dph, GOC:tb] |
| GO:0097435 | 0.035831853 | 1 / 2 | "A process that is carried out at the cellular level which results in the assembly, arrangement of constituent parts, or disassembly of a supramolecular fiber, a polymer consisting of an indefinite number of protein or protein complex subunits that have polymerised to form a fibe |
| GO:1903145 | 0.035831853 | 1 / 2 | "Any actin filament that is part of a cell cortex of cell tip." [GO_REF:0000064, GOC:TermGenie, PMID:20807799, PMID:24954052] |
| GO:2000689 | 0.035831853 | 1 / 2 | "An actin filament organization process that contributes to actomyosin contractile ring assembly during cytokinesis." [GOC:mah] |
| GO:0034657 | 0.052777189 | 1 / 3 | "A protein complex with ubiquitin ligase activity that, in Saccharomyces cerevisiae, is involved in proteasomal degradation of fructose-1,6-bisphosphatase (FBPase) and phosphoenolpyruvate carboxykinase during the transition from gluconeogenic to glycolytic growth conditions. |
| GO:0044695 | 0.052777189 | 1 / 3 | "An E3 ubiquitin ligase complex localized to the ER and Golgi membrane. In fission yeast comprises Dsc1, 2, 3 and 4. Involved in the processes of fission yeast sre1 (human SREBP) transcriptional activator proteolytic cleavage, the multivesicular body (MVB) pathway, and a post-er |
| GO:0061572 | 0.052777189 | 1 / 3 | "A process that results in the assembly, arrangement of constituent parts, or disassembly of an actin filament bundle." [GOC:dph] |
| GO:0071797 | 0.052777189 | 1 / 3 | "A ubiquitin ligase complex that catalyzes linear head-to-tail polyubiquitin conjugation on its targets. In human the complex consists of RBCK1, RNF31 and SHARPIN, and has an MW of approximately 600 kDa, suggesting a heteromultimeric assembly of its subunits. LUBAC stands |
| GO:0097505 | 0.052777189 | 1 / 3 | "A ubiquitin ligase complex found to be involved in post-replicative bypass of UV-damaged DNA and UV mutagenesis. In S. cerevisiae, the complex contains the ubiquitin conjugating enzyme Rad6 and Rad18, a protein containing a RING finger motif and a nucleotide binding mot |
| GO:1990352 | 0.052777189 | 1 / 3 | "A homodimeric protein complex composed of the E3 ubiquitin-protein ligase BRE1. Plays a role in regulating association of RNA polymerase II with active genes." [GOC:bhm, PMID:19531475] |
| GO:1990353 | 0.052777189 | 1 / 3 | "A ubiquitin ligase complex. In D. melanogaster, it regulates ubiquitination and proteolysis of the BMP receptor Thickveins in cystoblasts, potentially by controlling Tkv ubiquitination and degradation." [GOC:bhm, PMID:21145463] |
| GO:1990452 | 0.052777189 | 1 / 3 | "A ubiquitin ligase complex containing Parkin (PARK2), the F-box protein FBXW7 (also called SEL-10) and a cullin from the Cul1 subfamily; substrate specificity is conferred by the F-box protein." [GOC:bf, GOC:PARL, PMID:12628165] |
| GO:0005759 | 0.058359214 | 0 / 152 | "The gel-like material, with considerable fine structure, that lies in the matrix space, or lumen, of a mitochondrion. It contains the enzymes of the tricarboxylic acid cycle and, in some organisms, the enzymes concerned with fatty acid oxidation." [GOC:as, ISBN:0198506732] |

#### Supplementary Data S3.4.3

| GO_Term | P_val | Counts | MolFunc_Description |
|---|---|---|---|
| GO:0016716 | 2.05993E-05 | 5 / 15 | "Catalysis of an oxidation-reduction (redox) reaction in which hydrogen or electrons are transferred from each of two donors, and one atom of oxygen is incorporated into one donor." [GOC:mah] |
| GO:0016709 | 4.04482E-05 | 5 / 17 | "Catalysis of an oxidation-reduction (redox) reaction in which hydrogen or electrons are transferred from NADH or NADPH and one other donor, and one atom of oxygen is incorporated into one donor." [GOC:mah] |
| GO:0001567 | 0.000101244 | 4 / 11 | "Catalysis of the reaction: AH(2) + cholesterol + O2 = 25-hydroxycholesterol + A + H2O." [EC:1.14.99.38, RHEA:21104] |
| GO:0004501 | 0.000101244 | 4 / 11 | "Catalysis of the reaction: AH(2) + Ecdysone + O2 = 20-hydroxyecdysone + A + H2O." [EC:1.14.99.22, RHEA:14021] |
| GO:0004508 | 0.000101244 | 4 / 11 | "Catalysis of the reaction: a steroid + AH2 + O2 = a 17a-hydroxysteroid + A + H2O." [EC:1.14.14.19] |
| GO:0004509 | 0.000101244 | 4 / 11 | "Catalysis of the reaction: A C(21) steroid + [reduced NADPH--hemoprotein reductase] + O2 = a 21-hydroxy-C(21)-steroid + [oxidized NADPH--hemoprotein reductase] + H2O." [RHEA:65612] |
| GO:0004666 | 0.000101244 | 4 / 11 | "Catalysis of the reaction: arachidonate + donor-H2 + 2 O2 = prostaglandin H2 + acceptor + H2O." [EC:1.14.99.1] |
| GO:0008387 | 0.000101244 | 4 / 11 | "Catalysis of the reaction: a steroid + donor-H2 + O2 = 7-alpha-hydroxysteroid + H2O." [GOC:mah] |
| GO:0008388 | 0.000101244 | 4 / 11 | "Catalysis of the reaction: testosterone + donor-H2 + O2 = 15-alpha-hydroxytestosterone + H2O." [GOC:ai] |
| GO:0008389 | 0.000101244 | 4 / 11 | "Catalysis of the reaction: coumarin + O2 + NADPH + H+ = hydroxycoumarin + H2O + NADP+." [Reactome:163103] |
| GO:0008390 | 0.000101244 | 4 / 11 | "Catalysis of the reaction: testosterone + donor-H2 + O2 = 16-alpha-hydroxytestosterone + H2O." [GOC:ai] |
| GO:0008391 | 0.000101244 | 4 / 11 | "Catalysis of the incorporation of one atom from molecular oxygen into arachidonic acid and the reduction of the other atom of oxygen to water." [GOC:mah] |
| GO:0008396 | 0.000101244 | 4 / 11 | "Catalysis of the reaction: an oxysterol + NADPH + O2 = 7-alpha-hydroxylated oxysterol + NADP+ + H2O." [PMID:10882791] |
| GO:0008399 | 0.000101244 | 4 / 11 | "Catalysis of the conversion of naphthalene to naphthalene 1,2-oxide." [PMID:1742282, PMID:1981702] |
| GO:0008401 | 0.000101244 | 4 / 11 | "Catalysis of the conversion of retinoic acid to 4-hydroxy-retinoic acid." [PMID:19519282, PMID:9250660] |
| GO:0008403 | 0.000101244 | 4 / 11 | "Catalysis of the hydroxylation of C-24 of 25-hydroxycholecalciferol (25-hydroxyvitamin D3) to form 24(R),25-dihydroxycholecalciferol." [ISBN:0471331309] |
| GO:0008681 | 0.000101244 | 4 / 11 | "Catalysis of the reaction: 2-octaprenyl-6-methoxyphenol + O2 + 4 H+ = 2-octaprenyl-6-methoxy-1,4-benzoquinol + H2O." [PMID:27822549, PMID:4572721, RHEA:29407] |
| GO:0016712 | 0.000101244 | 4 / 11 | "Catalysis of an oxidation-reduction (redox) reaction in which hydrogen or electrons are transferred from reduced flavin or flavoprotein and one other donor, and one atom of oxygen is incorporated into one donor." [GOC:mah] |
| GO:0016713 | 0.000101244 | 4 / 11 | "Catalysis of an oxidation-reduction (redox) reaction in which hydrogen or electrons are transferred from reduced iron-sulfur protein and one other donor, and one atom of oxygen is incorporated into one donor." [GOC:mah] |
| GO:0016714 | 0.000101244 | 4 / 11 | "Catalysis of an oxidation-reduction (redox) reaction in which hydrogen or electrons are transferred from reduced pteridine and one other donor, and one atom of oxygen is incorporated into one donor." [GOC:mah] |
| GO:0016715 | 0.000101244 | 4 / 11 | "Catalysis of an oxidation-reduction (redox) reaction in which hydrogen or electrons are transferred from reduced ascorbate and one other donor, and one atom of oxygen is incorporated into one donor." [GOC:mah] |
| GO:0016720 | 0.000101244 | 4 / 11 | "Catalysis of the reaction: AH(2) + linoleate + O2 = A + crepenynate + 2 H2O." [EC:1.14.99.33, RHEA:23456] |
| GO:0018585 | 0.000101244 | 4 / 11 | "Catalysis of the reaction: fluorene + 2 H+ + 2 e- + O2 = H2O + 9-fluorenol." [UM-BBD_reactionID:r0407] |
| GO:0018586 | 0.000101244 | 4 / 11 | "Catalysis of the reaction: butyltin + O2 + 2 H+ + 2 e- = H2O + beta-hydroxybutyltin." [UM-BBD_reactionID:r0647] |
| GO:0018588 | 0.000101244 | 4 / 11 | "Catalysis of the reaction: tri-n-butyltin + O2 + 2 H+ + 2 e- = H2O + beta-hydroxybutyldibutyltin." [UM-BBD_reactionID:r0643] |
| GO:0018589 | 0.000101244 | 4 / 11 | "Catalysis of the reaction: dibutyltin + O2 + 2 H+ + 2 e- = H2O + beta-hydroxybutylbutyltin." [UM-BBD_reactionID:r0645] |
| GO:0018590 | 0.000101244 | 4 / 11 | "Catalysis of the reaction: methylsilanetriol + O2 + 2 H+ + 2 e- = H2O + hydroxymethylsilanetriol." [UM-BBD_reactionID:r0640] |
| GO:0018591 | 0.000101244 | 4 / 11 | "Catalysis of the reaction: methyl tert-butyl ether + 1/2 O2 = tert-butyl alcohol + formaldehyde." [UM-BBD_reactionID:r1023] |
| GO:0018592 | 0.000101244 | 4 / 11 | "Catalysis of the reaction: 4-nitrocatechol + O2 + 4 e- + 3 H+ = H2O + nitrite + 1,2,4-benzenetriol." [UM-BBD_reactionID:r0231] |
| GO:0018593 | 0.000101244 | 4 / 11 | "Catalysis of the reaction: 4-chlorophenoxyacetate + 1/2 O2 = glyoxylate + 4-chlorophenol." [UM-BBD_reactionID:r0281] |
| GO:0018594 | 0.000101244 | 4 / 11 | "Catalysis of the reaction: tert-butanol + 1/2 O2 = 2-methyl-2-hydroxy-1-propanol." [UM-BBD_reactionID:r0615] |
| GO:0018596 | 0.000101244 | 4 / 11 | "Catalysis of the reaction: dimethylsilanediol + O2 + 2 H+ + 2 e- = H2O + hydroxymethylmethylsilanediol." [UM-BBD_reactionID:r0637] |
| GO:0018597 | 0.000101244 | 4 / 11 | "Catalysis of the oxidation of alkanes (up to C8) to alcohols and alkenes (up to C5) to epoxides and alcohols in the presence of ammonium ions." [PMID:16347810] |
| GO:0018598 | 0.000101244 | 4 / 11 | "Catalysis of the reaction: hydroxymethylsilanetriol + O2 + 2 H+ + 2 e- = 2 H2O + formylsilanetriol." [UM-BBD_reactionID:r0641] |
| GO:0018599 | 0.000101244 | 4 / 11 | "Catalysis of the reaction: 2-hydroxyisobutyrate + 1/2 O2 = 2,3-dihydroxy-2-methyl propionate." [UM-BBD_reactionID:r0619] |
| GO:0018679 | 0.000101244 | 4 / 11 | "Catalysis of the reaction: dibenzothiophene-5,5-dioxide + O2 + 2 NADH + H+ = 2 NAD+ + H2O + 2'-hydroxybiphenyl-2-sulfinate." [UM-BBD_reactionID:r0235] |
| GO:0018689 | 0.000101244 | 4 / 11 | "Catalysis of the reaction: (1/2)-unsubstituted naphthalenoid-(2/1)-sulfonate + 2 H+ + 2 e- + O2 = 1,2-dihydroxynaphthalene derivative + HSO3(-). Substrates include naphthalene-1,6-disulfonate (forms 1,2-dihydroxynaphthalene-6-sulfonate), naphthalene-1-sulfonate (forms 1,2- |
| GO:0018690 | 0.000101244 | 4 / 11 | "Catalysis of the reaction: 4-methoxybenzoate + AH(2) + O2 = 4-hydroxybenzoate + A + formaldehyde + H2O." [EC:1.14.99.15, RHEA:18613] |
| GO:0034559 | 0.000101244 | 4 / 11 | "Catalysis of the reaction: bisphenol A + NADH + H+ + O2 = 1,2-bis(4-hydroxyphenyl)-2-propanol + NAD+ + H2O." [UM-BBD_reactionID:r0860] |
| GO:0034562 | 0.000101244 | 4 / 11 | "Catalysis of the reaction: 2,2-bis(4-hydroxyphenyl)-1-propanol + NADH + H+ + O2 = 2,3-bis(4-hydroxyphenyl)-1,2-propanediol + NAD+ + H2O." [UM-BBD_reactionID:r0864] |
| GO:0034786 | 0.000101244 | 4 / 11 | "Catalysis of the reaction: 9-fluorenone + O2 + 2 H+ + 2 e- = 1-hydro-1,1a-dihydroxy-9-fluorenone." [UM-BBD_reactionID:r1039] |
| GO:0034804 | 0.000101244 | 4 / 11 | "Catalysis of the reaction: benzo(a)pyrene + O2 = benzo(a)pyrene-11,12-epoxide." [UM-BBD_reactionID:r1119] |
| GO:0034816 | 0.000101244 | 4 / 11 | "Catalysis of the reaction: anthracene + 2 H2O = cis-9,10-dihydroanthracene-9,10-diol." [UM-BBD_reactionID:r1141] |
| GO:0034857 | 0.000101244 | 4 / 11 | "Catalysis of the reaction: 2-(methylthio)benzothiazole + 1/2 O2 = 2-(methylsulfinyl)benzothiazole." [UM-BBD_reactionID:r1287] |
| GO:0034858 | 0.000101244 | 4 / 11 | "Catalysis of the reaction: 2-hydroxybenzothiazole + 1/2 O2 = 2,6-dihydroxybenzothiazole." [UM-BBD_reactionID:r1291] |
| GO:0034859 | 0.000101244 | 4 / 11 | "Catalysis of the reaction: benzothiazole + 1/2 O2 = 2-hydroxybenzothiazole." [UM-BBD_reactionID:r1292] |
| GO:0034862 | 0.000101244 | 4 / 11 | "Catalysis of the reaction: 2,6-dihydroxybenzothiazole + 1/2 O2 = 2,6,7-trihydroxybenzothiazole." [UM-BBD_reactionID:r1294] |
| GO:0034870 | 0.000101244 | 4 / 11 | "Catalysis of the reaction: 2,2-dimethyl-3-pentanone + 1/2 O2 = 1-hydroxy-4,4-dimethylpentan-3-one." [UM-BBD_reactionID:r12979] |
| GO:0034873 | 0.000101244 | 4 / 11 | "Catalysis of the reaction: thioacetamide + O2 + 2 H+ + 2 e- = thioacetamide S-oxide + H2O." [UM-BBD_reactionID:r1312] |
| GO:0034874 | 0.000101244 | 4 / 11 | "Catalysis of the reaction: thioacetamide S-oxide + O2 + 2 H+ + 2 e- = thioacetamide S,S-dioxide + H2O." [UM-BBD_reactionID:r1313] |

| GO_Term | P_val | Counts | BioProc_Description |
|---|---|---|---|
| GO:0055114 | 8.45649E-05 | 24 / 453 | "OBSOLETE. A metabolic process that results in the removal or addition of one or more electrons to or from a substance, with or without the concomitant removal or addition of a proton or protons." [GOC:dhl, GOC:ecd, GOC:jh2, GOC:jid, GOC:mlg, GOC:rph] |
| GO:0070813 | 0.000578545 | 2 / 2 | "The chemical reactions and pathways involving hydrogen sulfide, H2S." [GOC:mah] |
| GO:2001308 | 0.001694516 | 2 / 3 | "The chemical reactions and pathways involving the epipolythiodioxopiperazine gliotoxin, a poisonous substance produced by some species of fungi." [PMID:16333108, PMID:17574915, PMID:18272357] |
| GO:0000103 | 0.003308718 | 2 / 4 | "The pathways by which inorganic sulfate is processed and incorporated into sulfated compounds." [GOC:jl] |
| GO:0008206 | 0.003308718 | 2 / 4 | "The chemical reactions and pathways involving bile acids, a group of steroid carboxylic acids occurring in bile, where they are present as the sodium salts of their amides with glycine or taurine." [GOC:go_curators] |
| GO:0008209 | 0.003308718 | 2 / 4 | "The chemical reactions and pathways involving androgens, C19 steroid hormones that can stimulate the development of male sexual characteristics." [ISBN:0198506732] |
| GO:0008210 | 0.003308718 | 2 / 4 | "The chemical reactions and pathways involving estrogens, C18 steroid hormones that can stimulate the development of female sexual characteristics. Also found in plants." [ISBN:0198506732] |
| GO:0008211 | 0.003308718 | 2 / 4 | "The chemical reactions and pathways involving glucocorticoids, hormonal C21 corticosteroids synthesized from cholesterol. Glucocorticoids act primarily on carbohydrate and protein metabolism, and have anti-inflammatory effects." [ISBN:0198506732] |
| GO:0010134 | 0.003308718 | 2 / 4 | "The pathway by which inorganic sulfate is activated, reduced and incorporated into sulfated compounds, where the activated sulfate, adenylyl-sulfate, is reduced to sulfite by the activity of adenylyl-sulfate reductase." [EC:1.8.99.2] |
| GO:0019379 | 0.003308718 | 2 / 4 | "The pathway by which inorganic sulfate is processed and incorporated into sulfated compounds, where the phosphoadenylyl sulfate reduction step is catalyzed by the enzyme phosphoadenylyl-sulfate reductase (thioredoxin) (EC:1.8.4.8)." [GOC:jl] |
| GO:0042359 | 0.003308718 | 2 / 4 | "The chemical reactions and pathways involving vitamin D, any of a group of related, fat-soluble compounds that are derived from delta-5,7 steroids and play a central role in calcium metabolism. Specific forms of vitamin D include calciferol (ergocalciferol; vitamin D2) and chol |
| GO:0044272 | 0.004689309 | 3 / 15 | "The chemical reactions and pathways resulting in the formation of compounds that contain sulfur, such as the amino acids methionine and cysteine or the tripeptide glutathione." [GOC:jl] |
| GO:0017062 | 0.005383816 | 2 / 5 | "The aggregation, arrangement and bonding together of a set of components to form the cytochrome bc(1) complex, a transmembrane lipoprotein complex that it catalyzes the reduction of cytochrome c by accepting reducing equivalents from Coenzyme Q, by the aggregation, |
| GO:0019420 | 0.005383816 | 2 / 5 | "The reduction of sulfate to hydrogen sulfide, which acts as a terminal electron acceptor. Sulfate is activated to adenosine-phosphosulfate (APS) which is then reduced to sulfite, which is in turn reduced to hydrogen sulfide." [PMID:25400653, PMID:26680199] |
| GO:0034551 | 0.005383816 | 2 / 5 | "The aggregation, arrangement and bonding together of a set of components to form the cytochrome bc(1) complex (also known as ubiquinol-cytochrome c reductase), in the mitochondrial inner membrane." [GOC:dgf, GOC:mcc] |
| GO:0045455 | 0.005383816 | 2 / 5 | "The chemical reactions and pathways involving ecdysteroids, a group of polyhydroxylated ketosteroids ubiquitous in insects and other arthropods, in which they initiate post-embryonic development, including the metamorphosis of immature forms and the development of th |
| GO:0062173 | 0.005383816 | 2 / 5 | "The chemical reactions and pathways by which living organisms transform brexanolone." [PMID:24390875] |
| GO:1904461 | 0.005383816 | 2 / 5 | "The chemical reactions and pathways involving ergosteryl 3-beta-D-glucoside." [GO_REF:0000068, GOC:TermGenie, PMID:26116408] |
| GO:1900579 | 0.005981762 | 5 / 51 | "The chemical reactions and pathways involving (17Z)-protosta-17(20),24-dien-3beta-ol." [GOC:di, GOC:TermGenie] |
| GO:1900810 | 0.007504688 | 5 / 54 | "The chemical reactions and pathways involving helvolic acid." [GOC:di, GOC:TermGenie] |
| GO:0016128 | 0.007884252 | 2 / 6 | "The chemical reactions and pathways involving phytosteroids, steroids of higher plants that differ from animal steroids in having substitutions at C24 and/or a double bond at C22. Phytosteroids are so named because they occur in higher plants; some, notably ergosterol, are |
| GO:0018868 | 0.007884252 | 2 / 6 | "The chemical reactions and pathways involving 2-aminobenzenesulfonate, aniline-o-sulfonic acid, an aromatic sulfonate used in organic synthesis and in the manufacture of various dyes and medicines." [UM-BBD_pathwayID:abs] |
| GO:0001575 | 0.010776183 | 2 / 7 | "The chemical reactions and pathways involving globosides, globotetraosylceramides, ceramides containing a core structure of GalNAc-beta-(1->3)-Gal-alpha-(1->4)-Glc(I). Globosides are the major neutral glycosphingolipid in normal kidneys and erythrocytes." [ISBN:019850673 |
| GO:0042081 | 0.010776183 | 2 / 7 | "The chemical reactions and pathways involving glycosylsphingolipidinositol (GSI) anchors, which attach membrane proteins to the lipid bilayer of the cell membrane." [GOC:go_curators] |
| GO:0046479 | 0.010776183 | 2 / 7 | "The chemical reactions and pathways resulting in the breakdown of glycosphingolipid, a compound with residues of sphingoid and at least one monosaccharide." [ISBN:0198506732] |
| GO:1902056 | 0.010776183 | 2 / 7 | "The chemical reactions and pathways involving (25S)-Delta(7)-dafachronate." [GOC:TermGenie, PMID:22505847] |
| GO:1902057 | 0.010776183 | 2 / 7 | "The chemical reactions and pathways involving (25S)-Delta(4)-dafachronate." [GOC:TermGenie, PMID:20178781] |
| GO:0001666 | 0.011881973 | 3 / 21 | "Any process that results in a change in state or activity of a cell or an organism (in terms of movement, secretion, enzyme production, gene expression, etc.) as a result of a stimulus indicating lowered oxygen tension. Hypoxia, defined as a decline in O2 levels below normoxic l |
| GO:0001573 | 0.014027411 | 2 / 8 | "The chemical reactions and pathways involving ceramide oligosaccharides carrying in addition to other sugar residues, one or more sialic acid residues." [ISBN:0198506732] |
| GO:0006675 | 0.014027411 | 2 / 8 | "The chemical reactions and pathways involving mannosyl-inositol phosphorylceramide, any lipid with a phosphodiester bridge between an inositol residue and the ceramide group which contains a phosphoryl (-P(O)=) groups and a mannose derivative." [GOC:ai, MetaCyc:MIP( |
| GO:0006677 | 0.014027411 | 2 / 8 | "The chemical reactions and pathways involving glycosylceramides, any compound formed by the replacement of the glycosidic hydroxyl group of a cyclic form of a monosaccharide (or derivative) by a ceramide group." [GOC:ai, ISBN:0198506732] |
| GO:0046478 | 0.014027411 | 2 / 8 | "The chemical reactions and pathways involving lactosylceramides, Gal-beta-(1->4)-Glc-beta-(1->1') ceramides, any compound formed by the replacement of the glycosidic C1 hydroxyl group of lactose by a ceramide group. They are the precursors of both gangliosides and globo: |
| GO:0051999 | 0.017607318 | 2 / 9 | "The chemical reactions and pathways resulting in the formation of mannosyl-inositol phosphorylceramide, any lipid with a phosphodiester bridge between an inositol residue and the ceramide group which contains a phosphoryl (-P(O)=) groups and a mannose derivative." [G( |
| GO:0019757 | 0.022132283 | 4 / 48 | "The chemical reactions and pathways involving glycosinolates, substituted thioglycosides found in rapeseed products and related cruciferae." [GOC:mah, http://www.gardeneaters.net/family_characteristics.html] |
| GO:0001576 | 0.024173656 | 1 / 1 | "The chemical reactions and pathways resulting in the formation of a ceramide with a core structure of GalNAc-beta-(1->3)-Gal-alpha-(1->4)-Glc(I)." [ISBN:0198506732] |
| GO:0009732 | 0.024173656 | 1 / 1 | "The series of events in which a stimulus from a hexose is received and converted into a molecular signal." [GOC:sm] |
| GO:0009757 | 0.024173656 | 1 / 1 | "The series of molecular signals mediated by the detection of hexose." [GOC:sm] |
| GO:0010126 | 0.024173656 | 1 / 1 | "The chemical reactions and pathways involving mycothiol, which consists of N-acetyl-L-cysteine linked to a pseudodisaccharide, D-glucosamine and myo-inositol. Mycothiol is produced in actinomycetes like mycobacteria and serves similar functions to glutathione." [GOC:pz] |
| GO:0010619 | 0.024173656 | 1 / 1 | "An adenylate cyclase-activating G protein-coupled receptor signaling pathway initiated by glucose binding to its receptor on the surface of the target cell, and ending with the regulation of a downstream cellular process." [GOC:dph, GOC:signaling, GOC:tb] |
| GO:0015671 | 0.024173656 | 1 / 1 | "The directed movement of oxygen (O2) into, out of or within a cell, or between cells, by means of some agent such as a transporter or pore." [GOC:ai] |
| GO:0016143 | 0.024173656 | 1 / 1 | "The chemical reactions and pathways involving S-glycosides, any compound in which a glycosyl group has been substituted into a thiol group." [ISBN:0198506732] |
| GO:0018892 | 0.024173656 | 1 / 1 | "The chemical reactions and pathways involving cyclohexylsulfamate, also known as cyclamic acid. Sodium cyclohexylsulfamate (CHS-Na) was a widely used sweetening agent but was banned because of the suspicion of carcinogenicity and metabolic conversion to cyclohexylami |
| GO:0018895 | 0.024173656 | 1 / 1 | "The chemical reactions and pathways involving dibenzothiophene, a substance composed of two benzene rings linked by one sulfide bond and one carbon-carbon bond. Dibenzothiophene derivatives can be detected in diesel oil following hydrodesulfurization treatment to re |
| GO:0018907 | 0.024173656 | 1 / 1 | "The chemical reactions and pathways involving dimethyl sulfoxide, DMSO (C2H6OS), an alkyl sulfoxide that is practically odorless in its purified form. As a highly polar organic liquid, it is a powerful solvent. Its biological activities include the ability to penetrate plant and anima |
| GO:0018909 | 0.024173656 | 1 / 1 | "The chemical reactions and pathways involving dodecyl sulfate, commonly found as sodium dodecyl sulfate (SDS), a component of a variety of synthetic surfactants." [UM-BBD_pathwayID:dds] |
| GO:0018969 | 0.024173656 | 1 / 1 | "The chemical reactions and pathways involving thiocyanate, the anion of thiocyanic acid, a toxic cyanide derivative commonly formed as a by-product in the production of gas for fuel, coke, and substances for chemical industries." [GOC:jl] |
| GO:0018972 | 0.024173656 | 1 / 1 | "The chemical reactions and pathways involving toluene-4-sulfonate, the anion of 4-toluene sulfonic acid, a white crystalline solid which is highly hygroscopic and soluble in water." [GOC:ai] |
| GO:0018984 | 0.024173656 | 1 / 1 | "The chemical reactions and pathways involving naphthalenesulfonate, sulfonated derivatives of naphthalene." [GOC:ai] |
| GO:0019296 | 0.024173656 | 1 / 1 | "The chemical reactions and pathways involving coenzyme M (2-thioethansulfonate), a coenzyme involved in the utilization of methane by methanogenic prokaryotes." [GOC:go_curators] |
| GO:0019416 | 0.024173656 | 1 / 1 | "The chemical reactions and pathways resulting in the conversion of thiosulfate to tetrathionate, using cytochrome c as an electron acceptor." [MetaCyc:THIOSULFOX-PWY] |

| GO_Term | P_val | Counts | CellComp_Description |
|---|---|---|---|
| GO:0005727 | 0.000946137 | 10 / 127 | "Circular DNA structures that are not part of a chromosome." [GOC:ai] |
| GO:0016234 | 0.000946137 | 10 / 127 | "A discrete intracellular part formed of aggregated molecules such as proteins or other biopolymers." [GOC:mah, PMID:11121744] |
| GO:0030076 | 0.000946137 | 10 / 127 | "A protein-pigment complex that may be closely or peripherally associated to photosynthetic reaction centers that participate in harvesting and transferring radiant energy to the reaction center." [GOC:lr] |
| GO:0044100 | 0.000946137 | 10 / 127 | "The complex infective apparatus corresponding to the central mass of cytoplasm within a spore that is injected into a host cell by various parasitic microorganisms." [GOC:mf, PMID:12076771, PMID:16004371, PMID:9723921] |
| GO:0045261 | 0.000946137 | 10 / 127 | "The sector of a hydrogen-transporting ATP synthase complex in which the catalytic activity resides; it comprises the catalytic core and central stalk, and is peripherally associated with a membrane, such as the plasma membrane or the mitochondrial inner membrane, when th |
| GO:0045281 | 0.000946137 | 10 / 127 | "A multimeric complex which consists of flavoprotein (subunit A ; InterPro:IPR003952), iron-sulfur protein (subunit B) and membrane-bound cytochrome b560 (subunit C; InterPro:IPR000701). In some Archaea, the membrane-bound subunits (C or C and D) do not necessarily co |
| GO:0046821 | 0.000946137 | 10 / 127 | "DNA structures that are not part of a chromosome." [GOC:ai] |
| GO:0060171 | 0.000946137 | 10 / 127 | "The portion of the plasma membrane surrounding a stereocilium." [GOC:dph, GOC:rph] |
| GO:0070090 | 0.000946137 | 10 / 127 | "The intracellular plane, located halfway between the poles of the spindle, where chromosomes align during metaphase of mitotic or meiotic nuclear division." [GOC:mah] |
| GO:0070477 | 0.000946137 | 10 / 127 | "An intracellular part that represents the innermost portion of an endospore; the endospore core is dehydrated, enriched in dipicolinic acid and divalent cations, and metabolically inactive." [GOC:mah, PMID:15035041, PMID:18035610] |
| GO:0070725 | 0.000946137 | 10 / 127 | "A cytoplasmic part that appears as an electron-dense sphere of around 1.5 micron diameter containing Yb protein found in somatic cells of ovary and testis. There are one to two Yb bodies per cell." [GOC:sart, PMID:19433453] |
| GO:0071152 | 0.000946137 | 10 / 127 | "A protein complex formed by the association of the guanine nucleotide exchange factor synembrin with the alpha(q) subunit of a heterotrimeric G protein." [GOC:mah, PMID:12509430] |
| GO:0071153 | 0.000946137 | 10 / 127 | "A protein complex formed by the association of the guanine nucleotide exchange factor synembrin with the alpha(o) subunit of a heterotrimeric G protein." [GOC:mah, PMID:12509430] |
| GO:1990302 | 0.000946137 | 10 / 127 | "A ubiquitin ligase complex consisting of Bre1 and Rad6 that mediates monoubiquitination of histone H2B to form H2BK123ub1. H2BK123ub1 gives a specific tag for epigenetic transcriptional activation, elongation by RNA polymerase II, telomeric silencing, and is also a prerequ |
| GO:0022624 | 0.001001809 | 10 / 128 | "A protein complex, that caps one or both ends of the proteasome core complex and regulates entry into, or exit from, the proteasome core complex." [GOC:mtg_sensu] |
| GO:0032300 | 0.001001809 | 10 / 128 | "Any complex formed of proteins that act in mismatch repair." [GOC:mah] |
| GO:0034708 | 0.001001809 | 10 / 128 | "A protein complex that possesses methyltransferase activity." [GOC:mah] |
| GO:0043229 | 0.001001809 | 10 / 128 | "Organized structure of distinctive morphology and function, occurring within the cell. Includes the nucleus, mitochondria, plastids, vacuoles, vesicles, ribosomes and the cytoskeleton. Excludes the plasma membrane." [GOC:go_curators] |
| GO:0045263 | 0.001001809 | 10 / 128 | "All non-F1 subunits of a hydrogen-transporting ATP synthase, including integral and peripheral membrane proteins." [PMID:10838056] |
| GO:0000178 | 0.001251067 | 10 / 132 | "A ribonuclease complex that has 3-prime to 5-prime exoribonuclease activity and possibly endoribonuclease activity, producing 5-prime-phosphomonoesters. Participates in a multitude of cellular RNA processing and degradation events preventing nuclear export and/or trans |
| GO:0019774 | 0.001251067 | 10 / 132 | "The proteasome core subcomplex that constitutes the two inner rings of the proteasome core complex. An example of this component is found in Mus musculus." [GOC:jl, GOC:mtg_sensu, GOC:rb, PMID:10854779] |
| GO:0019773 | 0.001320442 | 10 / 133 | "The proteasome core subcomplex that constitutes the two outer rings of the proteasome core complex. An example of this component is found in Mus musculus." [GOC:jl, GOC:mtg_sensu, GOC:rb, PMID:10854779] |
| GO:0008152 | 0.006113146 | 5 / 505 | "The chemical reactions and pathways, including anabolism and catabolism, by which living organisms transform chemical substances. Metabolic processes typically transform small molecules, but also include macromolecular processes such as DNA repair and replication, and p |
| GO:0005622 | 0.006661891 | 12 / 225 | "A component of a cell contained within (but not including) the plasma membrane. In eukaryotes it includes the nucleus and cytoplasm." [ISBN:0198506732] |
| GO:0008150 | 0.007137107 | 5 / 496 | "A biological process is the execution of a genetically-encoded biological module or program. It consists of all the steps required to achieve the specific biological objective of the module. A biological process is accomplished by a particular set of molecular functions carried out |
| GO:0022414 | 0.011394755 | 5 / 468 | "A biological process that directly contributes to the process of producing new individuals by one or two organisms. The new individuals inherit some proportion of their genetic material from the parent or parents." [GOC:dph, GOC:isa_complete] |
| GO:0098754 | 0.011394755 | 5 / 468 | "Any process that reduces or removes the toxicity of a toxic substance. These may include transport of the toxic substance away from sensitive areas and to compartments or complexes whose purpose is sequestration of the toxic substance." [GOC:dos] |
| GO:0050896 | 0.011772052 | 5 / 466 | "Any process that results in a change in state or activity of a cell or an organism (in terms of movement, secretion, enzyme production, gene expression, etc.) as a result of a stimulus. The process begins with detection of the stimulus and ends with a change in state or activity c |
| GO:0000003 | 0.011964839 | 5 / 465 | "The production of new individuals that contain some portion of genetic material inherited from one or more parent organisms." [GOC:go_curators, GOC:isa_complete, GOC:jl, ISBN:0198506732] |
| GO:0002376 | 0.011964839 | 5 / 465 | "Any process involved in the development or functioning of the immune system, an organismal system for calibrated responses to potential internal or invasive threats." [GO_REF:0000022, GOC:add] |
| GO:0016032 | 0.011964839 | 5 / 465 | "A multi-organism process in which a virus is a participant. The other participant is the host. Includes infection of a host cell, replication of the viral genome, and assembly of progeny virus particles. In some cases the viral genetic material may integrate into the host genome ar |
| GO:0032501 | 0.011964839 | 5 / 465 | "Any biological process, occurring at the level of a multicellular organism, pertinent to its function." [GOC:curators, GOC:dph, GOC:isa_complete, GOC:tb] |
| GO:0032502 | 0.011964839 | 5 / 465 | "A biological process whose specific outcome is the progression of an integrated living unit: an anatomical structure (which may be a subcellular structure, cell, tissue, or organ), or organism over time from an initial condition to a later condition." [GOC:isa_complete] |
| GO:0040011 | 0.011964839 | 5 / 465 | "Self-propelled movement of a cell or organism from one location to another." [GOC:dgh] |
| GO:0042592 | 0.011964839 | 5 / 465 | "Any biological process involved in the maintenance of an internal steady state." [GOC:jl, ISBN:0395825172] |
| GO:0043473 | 0.011964839 | 5 / 465 | "The accumulation of pigment in an organism, tissue or cell, either by increased deposition or by increased number of cells." [GOC:jl] |
| GO:0044419 | 0.011964839 | 5 / 465 | "Any process evolved to enable an interaction with an organism of a different species." [GOC:cc] |
| GO:0044848 | 0.011964839 | 5 / 465 | "A distinct period or stage in a biological process or cycle." [GOC:jl] |
| GO:0048511 | 0.011964839 | 5 / 465 | "Any process pertinent to the generation and maintenance of rhythms in the physiology of an organism." [GOC:jid] |
| GO:0051179 | 0.011964839 | 5 / 465 | "Any process in which a cell, a substance, or a cellular entity, such as a protein complex or organelle, is transported, tethered to or otherwise maintained in a specific location. In the case of substances, localization may also be achieved via selective degradation." [GOC:ai, GOC: |
| GO:0051703 | 0.011964839 | 5 / 465 | "Any process in which an organism has an effect on an organism of the same species." [GOC:ai] |
| GO:0065007 | 0.011964839 | 5 / 465 | "Any process that modulates a measurable attribute of any biological process, quality or function." [GOC:dph, GOC:isa_complete, GOC:mah, GOC:pr, GOC:vw] |
| GO:0009987 | 0.013453391 | 6 / 508 | "Any process that is carried out at the cellular level, but not necessarily restricted to a single cell. For example, cell communication occurs among more than one cell, but occurs at the cellular level." [GOC:go_curators, GOC:isa_complete] |
| GO:0005759 | 0.019381141 | 0 / 152 | "The gel-like material, with considerable fine structure, that lies in the matrix space, or lumen, of a mitochondrion. It contains the enzymes of the tricarboxylic acid cycle and, in some organisms, the enzymes concerned with fatty acid oxidation." [GOC:as, ISBN:0198506732] |
| GO:0040007 | 0.020933006 | 6 / 478 | "The increase in size or mass of an entire organism, a part of an organism or a cell." [GOC:bf, GOC:ma] |
| GO:0006107 | 0.025232919 | 1 / 1 | "The chemical reactions and pathways involving oxaloacetate, the anion of oxobutanedioic acid, an important intermediate in metabolism, especially as a component of the TCA cycle." [ISBN:0198506732] |
| GO:0009279 | 0.025232919 | 1 / 1 | "A lipid bilayer that forms the outermost membrane of the cell envelope; enriched in polysaccharide and protein; the outer leaflet of the membrane contains specific lipopolysaccharide structures." [GOC:md, GOC:mtg_sensu, ISBN:0135712254] |
| GO:0009337 | 0.025232919 | 1 / 1 | "A multisubunit iron flavoprotein, which in yeast is composed of 2 alpha and 2 beta subunits. Catalyzes the reduction of sulfite to sulfide." [BRENDA:1.8.1.2, GOC:jl] |
| GO:0019554 | 0.025232919 | 1 / 1 | "The chemical reactions and pathways resulting in the breakdown of glutamate into other compounds, including oxaloacetate." [GOC:go_curators] |
| GO:0019617 | 0.025232919 | 1 / 1 | "The chemical reactions and pathways resulting in the breakdown of protocatechuate, the anion of 3,4-dihydroxybenzoic acid, to yield oxaloacetate and pyruvate." [MetaCyc:P184-PWY] |

#### Supplementary Data S3.4.4

| GO_Term | P_val | Counts | MolFunc_Description |
|---|---|---|---|
| GO:0031420 | 6.96702E-05 | 7 / 48 | "Binding to an alkali metal ion; alkali metals are those elements in group Ia of the periodic table, with the exception of hydrogen." [GOC:mah] |
| GO:0032791 | 6.96702E-05 | 7 / 48 | "Binding to lead (Pb) ions." [GOC:mah] |
| GO:0046911 | 6.96702E-05 | 7 / 48 | "The formation of bonds from two or more atoms within the same ligand to a metal atom in complexes in which the metal is part of a ring." [ISBN:0198506732, ISBN:0716731363] |
| GO:0043169 | 0.000147554 | 7 / 54 | "Binding to a cation, a charged atom or group of atoms with a net positive charge." [GOC:jl] |
| GO:0005509 | 0.000193822 | 8 / 75 | "Binding to a calcium ion (Ca2+)." [GOC:ai] |
| GO:0016716 | 0.000252549 | 4 / 15 | "Catalysis of an oxidation-reduction (redox) reaction in which hydrogen or electrons are transferred from each of two donors, and one atom of oxygen is incorporated into one donor." [GOC:mah] |
| GO:0000287 | 0.000380732 | 7 / 63 | "Binding to a magnesium (Mg) ion." [GOC:ai] |
| GO:0008199 | 0.000427084 | 6 / 46 | "Binding to a ferric iron ion, Fe(III)." [GOC:ai] |
| GO:0008198 | 0.000597896 | 6 / 49 | "Binding to a ferrous iron ion, Fe(II)." [GOC:ai] |
| GO:0005506 | 0.000665015 | 6 / 50 | "Binding to an iron (Fe) ion." [GOC:ai] |
| GO:0046872 | 0.000799761 | 8 / 93 | "Binding to a metal ion." [GOC:ai] |
| GO:0016740 | 0.001091657 | 15 / 289 | "Catalysis of the transfer of a group, e.g. a methyl group, glycosyl group, acyl group, phosphorus-containing, or other groups, from one compound (generally regarded as the donor) to another compound (generally regarded as the acceptor). Transferase is the systematic name |
| GO:0051087 | 0.001115327 | 3 / 10 | "Binding to a chaperone protein, a class of proteins that bind to nascent or unfolded polypeptides and ensure correct folding or transport." [PMID:10585443] |
| GO:0005198 | 0.001356247 | 73 / 3742 | "The action of a molecule that contributes to the structural integrity of a complex." [GOC:mah, GOC:vw] |
| GO:0000254 | 0.001451665 | 2 / 3 | "Catalysis of the reaction: 4,4-dimethyl-5alpha-cholest-7-en-3beta-ol + 6 Fe(II)-[cytochrome b5] + 5 H(+) + 3 O2 = 4alpha-carboxy-4beta-methyl-5alpha-cholest-7-ene-3beta-ol + 6 Fe(III)-[cytochrome b5] + 4 H2O." [PMID:9811880, RHEA:55220] |
| GO:0005215 | 0.001966197 | 73 / 3724 | "Enables the directed movement of substances (such as macromolecules, small molecules, ions) into, out of or within a cell, or between cells." [GOC:ai, GOC:dgf] |
| GO:0016491 | 0.002235432 | 25 / 658 | "Catalysis of an oxidation-reduction (redox) reaction, a reversible chemical reaction in which the oxidation state of an atom or atoms within a molecule is altered. One substrate acts as a hydrogen or electron donor and becomes oxidized, while the other acts as hydrogen or el |
| GO:0030544 | 0.003098649 | 3 / 14 | "Binding to a Hsp70 protein, heat shock proteins around 70kDa in size." [ISBN:0198506732] |
| GO:0004133 | 0.00355709 | 13 / 265 | "Catalysis of the cleavage of branch points in branched glycogen polymers." [ISBN:0198506732] |
| GO:0016218 | 0.00355709 | 13 / 265 | "Catalysis of a multistep reaction that produce polyketides through decarboxylative condensation of carboxylic acids. The key chain-building reaction, a C-N bond-forming reaction, involves the generation of the characteristic peptide bond by nucleophilic attack of the amino g |
| GO:0032451 | 0.00355709 | 13 / 265 | "Catalysis of the removal of a methyl group from a substrate." [GOC:mah] |
| GO:1904091 | 0.00355709 | 13 / 265 | "Catalysis of a multistep reaction that produce non-ribosomal peptides. The key chain-building reaction, a C-N bond-forming reaction, involves the generation of the characteristic peptide bond by nucleophilic attack of the amino group of an amino-acyl donor unit covalently b |
| GO:0009975 | 0.003662755 | 13 / 266 | "Catalysis of a ring closure reaction." [ISBN:0198547684] |
| GO:0061783 | 0.003770813 | 13 / 267 | "A catalytic activity that contributes to the degradation of peptidoglycan." [GOC:dph, GOC:jh, PMID:22748813] |
| GO:0016853 | 0.003881298 | 13 / 268 | "Catalysis of the geometric or structural changes within one molecule. Isomerase is the systematic name for any enzyme of EC class 5." [ISBN:0198506732] |
| GO:0016874 | 0.004471273 | 13 / 273 | "Catalysis of the joining of two molecules, or two groups within a single molecule, using the energy from the hydrolysis of ATP, a similar triphosphate, or a pH gradient." [GOC:mah] |
| GO:0004053 | 0.004629145 | 2 / 5 | "Catalysis of the reaction: L-arginine + H2O = L-ornithine + urea." [EC:3.5.3.1] |
| GO:0004668 | 0.004629145 | 2 / 5 | "Catalysis of the reaction: H2O + L-arginyl-[protein] = L-citrullyl-[protein] + NH4+, resulting in citrullination of the target protein. This reaction is calcium-dependent." [PMID:27393304] |
| GO:0008783 | 0.004629145 | 2 / 5 | "Catalysis of the reaction: agmatine + H2O = putrescine + urea." [EC:3.5.3.11, RHEA:13929] |
| GO:0009015 | 0.004629145 | 2 / 5 | "Catalysis of the reaction: N(2)-succinyl-L-arginine + 2 H2O + 2 H+ = N(2)-succinyl-L-ornithine + CO2 + 2 NH4." [EC:3.5.3.23, RHEA:19533] |
| GO:0016403 | 0.004629145 | 2 / 5 | "Catalysis of the reaction: N(G),N(G)-dimethyl-L-arginine + H2O = dimethylamine + L-citrulline." [EC:3.5.3.18] |
| GO:0016980 | 0.004629145 | 2 / 5 | "Catalysis of the reaction: creatine + H2O = sarcosine + urea." [EC:3.5.3.3, RHEA:22456] |
| GO:0016990 | 0.004629145 | 2 / 5 | "Catalysis of the reaction: L-arginine + H2O = L-citrulline + NH3." [EC:3.5.3.6] |
| GO:0018754 | 0.004629145 | 2 / 5 | "Catalysis of the reaction: ammelide + H2O = cyanuric acid + NH3." [PMID:1991731] |
| GO:0018755 | 0.004629145 | 2 / 5 | "Catalysis of the reaction: 2-chloro-4-hydroxy-6-amino-1,3,5-triazine + OH- = 2,4-dihydroxy-6-amino-1,3,5-triazine + Cl-." [UM-BBD_reactionID:r1414] |
| GO:0018756 | 0.004629145 | 2 / 5 | "Catalysis of the reaction: ammeline + H2O = ammelide + NH3." [PMID:1991731] |
| GO:0018757 | 0.004629145 | 2 / 5 | "Catalysis of the reaction: deisopropylhydroxyatrazine + H2O = NH3 + 2,4-dihydroxy-6-(N'-ethyl)amino-1,3,5-triazine." [UM-BBD_reactionID:r0121] |
| GO:0018758 | 0.004629145 | 2 / 5 | "Catalysis of the reaction: 2,4-dihydroxy-6-(N'-ethyl)amino-1,3,5-triazine + H2O = CH3CH2NH2 + cyanuric acid." [UM-BBD_reactionID:r0122] |
| GO:0033972 | 0.004629145 | 2 / 5 | "Catalysis of the reaction: amidinoproclavaminate + H2O = proclavaminate + urea." [EC:3.5.3.22, RHEA:17001] |
| GO:0043730 | 0.004629145 | 2 / 5 | "Catalysis of the reaction: 5-ureido-4-imidazole carboxylate + H2O = 5-amino-4-imidazole carboxylate + NH3 + CO2." [GOC:jl] |
| GO:0043920 | 0.004629145 | 2 / 5 | "Catalysis of the reaction: N1-aminopropylagmatine + H2O = spermidine + urea." [GOC:jl, PMID:15983049, RHEA:35827] |
| GO:0047632 | 0.004629145 | 2 / 5 | "Catalysis of the reaction: agmatine + H2O = N-carbamoylputrescine + NH3." [EC:3.5.3.12, MetaCyc:AGMATINE-DEIMINASE-RXN] |
| GO:0047652 | 0.004629145 | 2 / 5 | "Catalysis of the reaction: allantoate + H2O + H+ = CO2 + NH3 + ureidoglycine." [EC:3.5.3.9, MetaCyc:ALLANTOATE-DEIMINASE-RXN] |
| GO:0047660 | 0.004629145 | 2 / 5 | "Catalysis of the reaction: N-amidino-L-aspartate + H2O = L-aspartate + urea." [EC:3.5.3.14, RHEA:14849] |
| GO:0047817 | 0.004629145 | 2 / 5 | "Catalysis of the reaction: D-arginine + H2O = D-ornithine + urea." [EC:3.5.3.10, RHEA:12901] |
| GO:0047854 | 0.004629145 | 2 / 5 | "Catalysis of the reaction: 1,4-diguanidinobutane + H2O = agmatine + urea." [EC:3.5.3.20, RHEA:13597] |
| GO:0047970 | 0.004629145 | 2 / 5 | "Catalysis of the reaction: guanidinoacetate + H2O = glycine + urea." [EC:3.5.3.2, RHEA:23268] |
| GO:0047971 | 0.004629145 | 2 / 5 | "Catalysis of the reaction: 4-guanidinobutanoate + H2O = 4-aminobutanoate + urea." [EC:3.5.3.7, RHEA:19501] |
| GO:0047972 | 0.004629145 | 2 / 5 | "Catalysis of the reaction: 3-guanidinopropanoate + H2O = beta-alanine + urea." [EC:3.5.3.17, RHEA:16029] |
| GO:0050098 | 0.004629145 | 2 / 5 | "Catalysis of the reaction: H2O + methylguanidine = methylammonium + urea." [EC:3.5.3.16, RHEA:11764] |

| GO_Term | P_val | Counts | BioProc_Description |
|---|---|---|---|
| GO:0009987 | 0 | 93 / 4055 | "Any process that is carried out at the cellular level, but not necessarily restricted to a single cell. For example, cell communication occurs among more than one cell, but occurs at the cellular level." [GOC:go_curators, GOC:isa_complete] |
| GO:0070813 | 0.000543221 | 2 / 2 | "The chemical reactions and pathways involving hydrogen sulfide, H2S." [GOC:mah] |
| GO:0035080 | 0.00171121 | 3 / 11 | "The decondensing (loosening) and swelling of the chromosomal sites of heat shock genes on polytene chromosomes in response to a heat shock stimulus." [GOC:bf, PMID:12543962] |
| GO:0070414 | 0.00171121 | 3 / 11 | "The chemical reactions and pathways involving trehalose that occur as a result of a heat stimulus, a temperature stimulus above the optimal temperature for that organism." [GOC:jp, GOC:mah, PMID:9797333] |
| GO:0008202 | 0.002112866 | 6 / 60 | "The chemical reactions and pathways involving steroids, compounds with a 1,2,cyclopentanoperhydrophenanthrene nucleus." [ISBN:0198547684] |
| GO:0070370 | 0.00283266 | 3 / 13 | "Any process that increases heat tolerance of a cell in response to high temperatures." [GOC:jp] |
| GO:0000103 | 0.003111449 | 2 / 4 | "The pathways by which inorganic sulfate is processed and incorporated into sulfated compounds." [GOC:jl] |
| GO:0010134 | 0.003111449 | 2 / 4 | "The pathway by which inorganic sulfate is activated, reduced and incorporated into sulfated compounds, where the activated sulfate, adenylyl-sulfate, is reduced to sulfite by the activity of adenylyl-sulfate reductase." [EC:1.8.99.2] |
| GO:0019379 | 0.003111449 | 2 / 4 | "The pathway by which inorganic sulfate is processed and incorporated into sulfated compounds, where the phosphoadenylyl sulfate reduction step is catalyzed by the enzyme phosphoadenylyl-sulfate reductase (thioredoxin) (EC:1.8.4.8)." [GOC:jl] |
| GO:0006213 | 0.003353772 | 4 / 28 | "The chemical reactions and pathways involving any pyrimidine nucleoside, one of a family of organic molecules consisting of a pyrimidine base covalently bonded to ribose (a ribonucleoside) or deoxyribose (a deoxyribonucleoside)." [GOC:jl, ISBN:0140512713] |
| GO:0009119 | 0.003353772 | 4 / 28 | "The chemical reactions and pathways involving any ribonucleoside, a nucleoside in which purine or pyrimidine base is linked to a ribose (beta-D-ribofuranose) molecule." [GOC:jl] |
| GO:0009120 | 0.003353772 | 4 / 28 | "The chemical reactions and pathways involving any one of a family of organic molecules consisting of a purine or pyrimidine base covalently bonded to a sugar deoxyribose (a deoxyribonucleoside)." [GOC:jl, ISBN:0140512713] |
| GO:0009163 | 0.003353772 | 4 / 28 | "The chemical reactions and pathways resulting in the formation of any one of a family of organic molecules consisting of a purine or pyrimidine base covalently bonded to a sugar ribose (a ribonucleoside) or deoxyribose (a deoxyribonucleoside)." [GOC:jl, ISBN:0140512713] |
| GO:0042278 | 0.003353772 | 4 / 28 | "The chemical reactions and pathways involving one of a family of organic molecules consisting of a purine base covalently bonded to a sugar ribose (a ribonucleoside) or deoxyribose (a deoxyribonucleoside)." [GOC:jl, ISBN:0140512713] |
| GO:0070637 | 0.003353772 | 4 / 28 | "The chemical reactions and pathways involving any pyridine nucleoside, a nucleoside in which a pyridine base covalently bonded to a sugar, usually ribose." [GOC:mah] |
| GO:0031990 | 0.003523146 | 3 / 14 | "The directed movement of mRNA from the nucleus to the cytoplasm during a heat stimulus, a temperature stimulus above the optimal temperature for the organism; in particular, a process that enables an organism withstand exposure to temperatures that would otherwise |
| GO:1901657 | 0.003802463 | 4 / 29 | "The chemical reactions and pathways involving glycosyl compound." [GOC:pr, GOC:TermGenie] |
| GO:0009116 | 0.004288287 | 4 / 30 | "The chemical reactions and pathways involving a nucleoside, a nucleobase linked to either beta-D-ribofuranose (a ribonucleoside) or 2-deoxy-beta-D-ribofuranose, (a deoxyribonucleoside), e.g. adenosine, guanosine, inosine, cytidine, uridine and deoxyadenosine, deoxyguano |
| GO:0009164 | 0.004288287 | 4 / 30 | "The chemical reactions and pathways resulting in the breakdown of any one of a family of organic molecules consisting of a purine or pyrimidine base covalently bonded to a sugar ribose (a ribonucleoside) or deoxyribose (a deoxyribonucleoside)." [GOC:jl] |
| GO:0006189 | 0.005066698 | 2 / 5 | "The chemical reactions and pathways resulting in the formation of IMP, inosine monophosphate, by the stepwise assembly of a purine ring on ribose 5-phosphate." [GOC:mah, ISBN:0716720094] |
| GO:0019420 | 0.005066698 | 2 / 5 | "The reduction of sulfate to hydrogen sulfide, which acts as a terminal electron acceptor. Sulfate is activated to adenosine-phosphosulfate (APS) which is then reduced to sulfite, which is in turn reduced to hydrogen sulfide." [PMID:25400653, PMID:26680199] |
| GO:0018868 | 0.007425528 | 2 / 6 | "The chemical reactions and pathways involving 2-aminobenzenesulfonate, aniline-o-sulfonic acid, an aromatic sulfonate used in organic synthesis and in the manufacture of various dyes and medicines." [UM-BBD_pathwayID:abs] |
| GO:0055086 | 0.00821152 | 5 / 57 | "The cellular chemical reactions and pathways involving a nucleobase-containing small molecule: a nucleobase, a nucleoside, or a nucleotide." [GOC:vw] |
| GO:0034605 | 0.009609177 | 3 / 20 | "Any process that results in a change in state or activity of a cell (in terms of movement, secretion, enzyme production, gene expression, etc.) as a result of a heat stimulus, a temperature stimulus above the optimal temperature for that organism." [GOC:mah] |
| GO:0006059 | 0.010156963 | 2 / 7 | "The chemical reactions and pathways involving hexitols, any alditol with a chain of six carbon atoms in the molecule." [ISBN:0198506732] |
| GO:0016125 | 0.010474215 | 4 / 39 | "The chemical reactions and pathways involving sterols, steroids with one or more hydroxyl groups and a hydrocarbon side-chain in the molecule." [ISBN:0198547684] |
| GO:0006790 | 0.015556423 | 3 / 24 | "The chemical reactions and pathways involving the nonmetallic element sulfur or compounds that contain sulfur, such as the amino acids methionine and cysteine or the tripeptide glutathione." [GOC:ai] |
| GO:0006188 | 0.016620976 | 2 / 9 | "The chemical reactions and pathways resulting in the formation of IMP, inosine monophosphate." [ISBN:0198506732] |
| GO:0007323 | 0.016620976 | 2 / 9 | "The generation of a mature, active peptide pheromone via processes unique to its processing and modification. An example of this process is found in Saccharomyces cerevisiae." [GOC:elh] |
| GO:0006816 | 0.017274292 | 3 / 25 | "The directed movement of calcium (Ca) ions into, out of or within a cell, or between cells, by means of some agent such as a transporter or pore." [GOC:ai] |
| GO:1900555 | 0.018554707 | 5 / 71 | "The chemical reactions and pathways involving emericellamide." [GOC:di, GOC:TermGenie] |
| GO:0019757 | 0.020155877 | 4 / 48 | "The chemical reactions and pathways involving glycosinolates, substituted thioglycosides found in rapeseed products and related cruciferae." [GOC:mah, http://www.gardeneaters.net/family_characteristics.html] |
| GO:0051604 | 0.020155877 | 4 / 48 | "Any process leading to the attainment of the full functional capacity of a protein." [GOC:ai] |
| GO:0006458 | 0.020976697 | 3 / 27 | "The process of assisting in the folding of a nascent peptide chain into its correct tertiary structure." [GOC:mb] |
| GO:0022417 | 0.022957297 | 3 / 28 | "The process of assisting in the covalent and noncovalent assembly of single chain polypeptides or multisubunit complexes into the correct tertiary structure that results in the attainment of the full functional capacity of a protein." [GOC:isa_complete] |
| GO:0006061 | 0.023427867 | 1 / 1 | "The chemical reactions and pathways resulting in the formation of sorbitol (D-glucitol), one of the ten stereoisomeric hexitols. It can be derived from glucose by reduction of the aldehyde group." [ISBN:0198506732] |
| GO:0010126 | 0.023427867 | 1 / 1 | "The chemical reactions and pathways involving mycothiol, which consists of N-acetyl-L-cysteine linked to a pseudodisaccharide, D-glucosamine and myo-inositol. Mycothiol is produced in actinomycetes like mycobacteria and serves similar functions to glutathione." [GOC:pz] |
| GO:0015671 | 0.023427867 | 1 / 1 | "The directed movement of oxygen (O2) into, out of or within a cell, or between cells, by means of some agent such as a transporter or pore." [GOC:ai] |
| GO:0016143 | 0.023427867 | 1 / 1 | "The chemical reactions and pathways involving S-glycosides, any compound in which a glycosyl group has been substituted into a thiol group." [ISBN:0198506732] |
| GO:0018892 | 0.023427867 | 1 / 1 | "The chemical reactions and pathways involving cyclohexylsulfamate, also known as cyclamic acid. Sodium cyclohexylsulfamate (CHS-Na) was a widely used sweetening agent but was banned because of the suspicion of carcinogenicity and metabolic conversion to cyclohexylam |
| GO:0018895 | 0.023427867 | 1 / 1 | "The chemical reactions and pathways involving dibenzothiophene, a substance composed of two benzene rings linked by one sulfide bond and one carbon-carbon bond. Dibenzothiophene derivatives can be detected in diesel oil following hydrodesulfurization treatment to r |
| GO:0018907 | 0.023427867 | 1 / 1 | "The chemical reactions and pathways involving dimethyl sulfoxide, DMSO (C2H6OS), an alkyl sulfoxide that is practically odorless in its purified form. As a highly polar organic liquid, it is a powerful solvent. Its biological activities include the ability to penetrate plant and anim |
| GO:0018909 | 0.023427867 | 1 / 1 | "The chemical reactions and pathways involving dodecyl sulfate, commonly found as sodium dodecyl sulfate (SDS), a component of a variety of synthetic surfactants." [UM-BBD_pathwayID:dds] |
| GO:0018969 | 0.023427867 | 1 / 1 | "The chemical reactions and pathways involving thiocyanate, the anion of thiocyanic acid, a toxic cyanide derivative commonly formed as a by-product in the production of gas for fuel, coke, and substances for chemical industries." [GOC:jl] |
| GO:0018972 | 0.023427867 | 1 / 1 | "The chemical reactions and pathways involving toluene-4-sulfonate, the anion of 4-toluene sulfonic acid, a white crystalline solid which is highly hygroscopic and soluble in water." [GOC:ai] |
| GO:0018984 | 0.023427867 | 1 / 1 | "The chemical reactions and pathways involving naphthalenesulfonate, sulfonated derivatives of naphthalene." [GOC:ai] |
| GO:0019296 | 0.023427867 | 1 / 1 | "The chemical reactions and pathways involving coenzyme M (2-thioethansulfonate), a coenzyme involved in the utilization of methane by methanogenic prokaryotes." [GOC:go_curators] |
| GO:0019416 | 0.023427867 | 1 / 1 | "The chemical reactions and pathways resulting in the conversion of thiosulfate to tetrathionate, using cytochrome c as an electron acceptor." [MetaCyc:THIOSULFOX-PWY] |
| GO:0019417 | 0.023427867 | 1 / 1 | "The chemical reactions and pathways resulting the addition of oxygen to elemental sulfur." [GOC:jl, MetaCyc:FESULFOX-PWY, MetaCyc:SULFUROX-PWY] |
| GO:0019418 | 0.023427867 | 1 / 1 | "The chemical reactions and pathways resulting in the conversion of sulfide to elemental sulfur in a higher oxidation state, or to sulfite or sulfate." [MetaCyc:P222-PWY, MetaCyc:PWY-5274, MetaCyc:PWY-5285] |

| GO_Term | P_val | Counts | CellComp_Description |
|---|---|---|---|
| GO:0008152 | 0.002077811 | 4 / 504 | "The chemical reactions and pathways, including anabolism and catabolism, by which living organisms transform chemical substances. Metabolic processes typically transform small molecules, but also include macromolecular processes such as DNA repair and replication, and |
| GO:0040007 | 0.003444682 | 4 / 478 | "The increase in size or mass of an entire organism, a part of an organism or a cell." [GOC:bf, GOC:ma] |
| GO:0051641 | 0.003658458 | 2 / 4 | "A cellular localization process whereby a substance or cellular entity, such as a protein complex or organelle, is transported to, and/or maintained in, a specific location within a cell including the localization of substances or cellular entities to the cell membrane." [GOC:tb, GC |
| GO:0042175 | 0.004120898 | 10 / 156 | "The continuous network of membranes encompassing the nuclear outer membrane and the endoplasmic reticulum membrane." [GOC:bf, GOC:jl, GOC:mah, GOC:mcc, GOC:pr, GOC:vw] |
| GO:0022414 | 0.004166516 | 4 / 468 | "A biological process that directly contributes to the process of producing new individuals by one or two organisms. The new individuals inherit some proportion of their genetic material from the parent or parents." [GOC:dph, GOC:isa_complete] |
| GO:0098754 | 0.004166516 | 4 / 468 | "Any process that reduces or removes the toxicity of a toxic substance. These may include transport of the toxic substance away from sensitive areas and to compartments or complexes whose purpose is sequestration of the toxic substance." [GOC:dos] |
| GO:0050896 | 0.004326853 | 4 / 466 | "Any process that results in a change in state or activity of a cell or an organism (in terms of movement, secretion, enzyme production, gene expression, etc.) as a result of a stimulus. The process begins with detection of the stimulus and ends with a change in state or activity o |
| GO:0000003 | 0.004409159 | 4 / 465 | "The production of new individuals that contain some portion of genetic material inherited from one or more parent organisms." [GOC:go_curators, GOC:isa_complete, GOC:jl, ISBN:0198506732] |
| GO:0002376 | 0.004409159 | 4 / 465 | "Any process involved in the development or functioning of the immune system, an organismal system for calibrated responses to potential internal or invasive threats." [GO_REF:0000022, GOC:add] |
| GO:0016032 | 0.004409159 | 4 / 465 | "A multi-organism process in which a virus is a participant. The other participant is the host. Includes infection of a host cell, replication of the viral genome, and assembly of progeny virus particles. In some cases the viral genetic material may integrate into the host genome ar |
| GO:0032501 | 0.004409159 | 4 / 465 | "Any biological process, occurring at the level of a multicellular organism, pertinent to its function." [GOC:curators, GOC:dph, GOC:isa_complete, GOC:tb] |
| GO:0032502 | 0.004409159 | 4 / 465 | "A biological process whose specific outcome is the progression of an integrated living unit: an anatomical structure (which may be a subcellular structure, cell, tissue, or organ), or organism over time from an initial condition to a later condition." [GOC:isa_complete] |
| GO:0040011 | 0.004409159 | 4 / 465 | "Self-propelled movement of a cell or organism from one location to another." [GOC:dgh] |
| GO:0042592 | 0.004409159 | 4 / 465 | "Any biological process involved in the maintenance of an internal steady state." [GOC:jl, ISBN:0395825172] |
| GO:0043473 | 0.004409159 | 4 / 465 | "The accumulation of pigment in an organism, tissue or cell, either by increased deposition or by increased number of cells." [GOC:jl] |
| GO:0044419 | 0.004409159 | 4 / 465 | "Any process evolved to enable an interaction with an organism of a different species." [GOC:cc] |
| GO:0044848 | 0.004409159 | 4 / 465 | "A distinct period or stage in a biological process or cycle." [GOC:jl] |
| GO:0048511 | 0.004409159 | 4 / 465 | "Any process pertinent to the generation and maintenance of rhythms in the physiology of an organism." [GOC:jid] |
| GO:0051179 | 0.004409159 | 4 / 465 | "Any process in which a cell, a substance, or a cellular entity, such as a protein complex or organelle, is transported, tethered to or otherwise maintained in a specific location. In the case of substances, localization may also be achieved via selective degradation." [GOC:ai, GOC: |
| GO:0051703 | 0.004409159 | 4 / 465 | "Any process in which an organism has an effect on an organism of the same species." [GOC:ai] |
| GO:0065007 | 0.004409159 | 4 / 465 | "Any process that modulates a measurable attribute of any biological process, quality or function." [GOC:dph, GOC:isa_complete, GOC:mah, GOC:pr, GOC:vw] |
| GO:0005905 | 0.00561593 | 9 / 138 | "A part of the endomembrane system in the form of an invagination of a membrane upon which a clathrin coat forms, and that can be converted by vesicle budding into a clathrin-coated vesicle. Coated pits form on the plasma membrane, where they are involved in receptor- |
| GO:0034357 | 0.00561593 | 9 / 138 | "A membrane enriched in complexes formed of reaction centers, accessory pigments and electron carriers, in which photosynthetic reactions take place." [GOC:ds, GOC:mah] |
| GO:0036362 | 0.00561593 | 9 / 138 | "A double layer of lipid molecules that surrounds an ascus, a capsule containing the sexual spores in some fungi." [GOC:mcc, GOC:vw, PMID:21900489] |
| GO:0070469 | 0.00561593 | 9 / 138 | "The protein complexes that form the electron transport system (the respiratory chain), associated with a cell membrane, usually the plasma membrane (in prokaryotes) or the inner mitochondrial membrane (on eukaryotes). The respiratory chain complexes transfer electrons |
| GO:0097478 | 0.00561593 | 9 / 138 | "Any of the two layers of lipid molecules that constitute a membrane." [GOC:cjm] |
| GO:0098552 | 0.00561593 | 9 / 138 | "A cellular component consisting of one leaflet of a membrane bilayer and any proteins embedded or anchored in it or attached to its surface." [GOC:dos] |
| GO:0140221 | 0.00561593 | 9 / 138 | "Host-derived membrane of a pathogen-containing vacuole." [PMID:10560000, PMID:26842840] |
| GO:0140549 | 0.00561593 | 9 / 138 | "The membrane surrounding the spore core (endospore core) that separates it from its external environment." [PMID:23202530] |
| GO:0005642 | 0.005858648 | 9 / 139 | "Stacks of endoplasmic reticulum (ER) membranes containing a high density of nuclear pores, thought to form from excess nuclear membrane components, that have been described in a number of different cells. Annulate lamellar membranes are continuous with and embedd |
| GO:0019867 | 0.005858648 | 9 / 139 | "The external membrane of Gram-negative bacteria or certain organelles such as mitochondria and chloroplasts; freely permeable to most ions and metabolites." [GOC:go_curators] |
| GO:0034045 | 0.005858648 | 9 / 139 | "A cellular membrane associated with the phagophore assembly site." [GOC:mah, GOC:rph, PMID:16874040, PMID:17382324] |
| GO:0098857 | 0.005858648 | 9 / 139 | "A membrane region with a lipid composition that is distinct from that of the membrane regions that surround it." [PMID:20044567, PMID:26253820] |
| GO:0031090 | 0.006342791 | 10 / 167 | "A membrane that is one of the two lipid bilayers of an organelle envelope or the outermost membrane of single membrane bound organelle." [GOC:dos, GOC:mah] |
| GO:0005628 | 0.00636651 | 9 / 141 | "The prospore membrane is a double-membraned structure that extends from the cytoplasmic face of the spindle pole bodies to encompass the spindle pole bodies and the four nuclear lobes that are formed during meiosis. It helps isolate the meiotic nuclei from the cytoplas |
| GO:0048475 | 0.00636651 | 9 / 141 | "A single or double lipid bilayer with any of several different proteinaceous coats that can associate with membranes. Membrane coats include those formed by clathrin plus an adaptor complex, the COPI and COPII complexes." [GOC:jid] |
| GO:0070056 | 0.00636651 | 9 / 141 | "The region of the prospore membrane that extends to surround the spore nucleus; coated with specific proteins that are thought to play a role in prospore membrane organization." [GOC:mah, PMID:14702385] |
| GO:0008150 | 0.006706085 | 5 / 496 | "A biological process is the execution of a genetically-encoded biological module or program. It consists of all the steps required to achieve the specific biological objective of the module. A biological process is accomplished by a particular set of molecular functions carried out |
| GO:0019898 | 0.007771893 | 9 / 146 | "The component of a membrane consisting of gene products and protein complexes that are loosely bound to one of its surfaces, but not integrated into the hydrophobic region." [GOC:dos, GOC:jl, GOC:mah] |
| GO:0005872 | 0.008693558 | 2 / 6 | "Any complex that includes a dimer of molecules from the kinesin superfamily and any associated proteins, and moves towards the minus end of a microtubule." [GOC:mah] |
| GO:0005873 | 0.008693558 | 2 / 6 | "Any complex that includes a dimer of molecules from the kinesin superfamily and any associated proteins, and moves towards the plus end of a microtubule." [GOC:mah] |
| GO:0007266 | 0.008693558 | 2 / 6 | "An intracellular signaling cassette in which a small monomeric GTPase of the Rho subfamily relays a signal." [GOC:bf] |
| GO:0016601 | 0.008693558 | 2 / 6 | "An intracellular signaling cassette in which a small monomeric GTPase of the Rac subfamily relays a signal." [GOC:bf] |
| GO:0016938 | 0.008693558 | 2 / 6 | "A complex of two kinesin heavy chains and two kinesin light chains." [http://www.csfic.mi.cnr.it/centro/lines/8/ric.html] |
| GO:0016939 | 0.008693558 | 2 / 6 | "A complex consisting of two distinct motor subunits that form a heterodimer complexed with a third non-motor accessory subunit, the kinesin associated protein or KAP; the KIF3 heterodimer interacts via its C-terminal portion with KAP, which is thought to regulate the bindi |
| GO:0031028 | 0.008693558 | 2 / 6 | "The series of molecular signals, mediated by the small GTPase Ras, that results in the initiation of contraction of the contractile ring, at the beginning of cytokinesis and cell division by septum formation. The pathway coordinates chromosome segregation with mitotic exit and |
| GO:0031291 | 0.008693558 | 2 / 6 | "An intracellular signaling cassette in which a small monomeric GTPase of the Ran subfamily relays a signal." [GOC:mah] |
| GO:0032011 | 0.008693558 | 2 / 6 | "An intracellular signaling cassette in which a small monomeric GTPase of the ARF subfamily relays a signal." [GOC:mah] |
| GO:0032482 | 0.008693558 | 2 / 6 | "An intracellular signaling cassette in which a small monomeric GTPase of the Rab subfamily relays a signal." [GOC:mah] |
| GO:0032484 | 0.008693558 | 2 / 6 | "An intracellular signaling cassette in which a small monomeric GTPase of the Ral subfamily relays a signal." [GOC:mah] |

#### Supplementary Data S3.4.5

| GO_Term | P_val | Counts | MolFunc_Description |
|---|---|---|---|
| GO:0005198 | 3.19632E-06 | 108 / 374 | "The action of a molecule that contributes to the structural integrity of a complex." [GOC:mah, GOC:vw] |
| GO:0005215 | 1.57083E-05 | 109 / 372 | "Enables the directed movement of substances (such as macromolecules, small molecules, ions) into, out of or within a cell, or between cells." [GOC:ai, GOC:dgf] |
| GO:0016491 | 7.07305E-05 | 40 / 658 | "Catalysis of an oxidation-reduction (redox) reaction, a reversible chemical reaction in which the oxidation state of an atom or atoms within a molecule is altered. One substrate acts as a hydrogen or electron donor and becomes oxidized, while the other acts as hydrogen or e |
| GO:0140110 | 8.86339E-05 | 109 / 367 | "A molecular function that controls the rate, timing and/or magnitude of gene transcription. The function of transcriptional regulators is to modulate gene expression at the transcription step so that they are expressed in the right cell at the right time and in the right amount |
| GO:0005488 | 0.000138235 | 108 / 363 | "The selective, non-covalent, often stoichiometric, interaction of a molecule with one or more specific sites on another molecule." [GOC:ceb, GOC:mah, ISBN:0198506732] |
| GO:0060090 | 0.000160326 | 108 / 363 | "The binding activity of a molecule that brings together two or more molecules through a selective, non-covalent, often stoichiometric interaction, permitting those molecules to function in a coordinated way." [GOC:mtg_MIT_16mar07, GOC:vw] |
| GO:0009055 | 0.000169207 | 109 / 365 | "A molecular function representing the directed movement of electrons from one molecular entity to another, typically mediated by electron carriers or acceptors, resulting in the transfer of energy and/or the reduction-oxidation (redox) transformation of chemical species. Th |
| GO:0140223 | 0.000202244 | 108 / 362 | "A molecular function required for core promoter activity that mediates the assembly of the RNA polymerase holoenzyme at promoter DNA to form the pre-initiation complex (PIC). General transcription factors (GTFs) bind to and open promoter DNA, initiate RNA synthesis ar |
| GO:0016209 | 0.000208114 | 108 / 362 | "Inhibition of the reactions brought about by dioxygen (O2) or peroxides. Usually the antioxidant is effective because it can itself be more easily oxidized than the substance protected. The term is often applied to components that can trap free radicals, thereby breaking the c |
| GO:0098772 | 0.000220309 | 108 / 361 | "A molecular function regulator regulates the activity of its target via non-covalent binding that does not result in covalent modification to the target. Examples of molecular function regulators include regulatory subunits of multimeric enzymes and channels. Mechanisms of r |
| GO:0045735 | 0.00022664 | 108 / 361 | "Functions in the storage of nutritious substrates." [GOC:ai] |
| GO:0038024 | 0.000239788 | 108 / 361 | "Binding specifically to a substance (cargo) to deliver it to a transport vesicle. Cargo receptors span a membrane (either the plasma membrane or a vesicle membrane), binding simultaneously to cargo molecules and coat adaptors, to efficiently recruit soluble proteins to nasce |
| GO:0044183 | 0.000246611 | 108 / 361 | "Binding to a protein or a protein-containing complex to assist the protein folding process." [GOC:mtg_cambridge_2009] |
| GO:0045182 | 0.000246611 | 108 / 361 | "Any molecular function involved in the initiation, activation, perpetuation, repression or termination of polypeptide synthesis at the ribosome." [GOC:ai] |
| GO:0140104 | 0.000253605 | 108 / 361 | "Directly binding to a specific ion or molecule and delivering it either to an acceptor molecule or to a specific location." [GOC:molecular_function_refactoring, GOC:pdt] |
| GO:0060089 | 0.000260773 | 108 / 361 | "A compound molecular function in which an effector function is controlled by one or more regulatory components." [GOC:dos, GOC:pdt] |
| GO:0090729 | 0.000260773 | 108 / 361 | "Interacting selectively with one or more biological molecules in another (target) organism, initiating pathogenesis (leading to an abnormal, generally detrimental state) in the target organism. The activity should refer to an evolved function of the active gene product, i.e. one |
| GO:0140299 | 0.000260773 | 108 / 361 | "Binding to a small molecule and eliciting a change in the protein's activity in response to the intracellular level of that small molecule." [PMID:26328879] |
| GO:0140313 | 0.000260773 | 108 / 361 | "Binding to a specific molecule to prevent it from interacting with other partners or to inhibit its localization to the area of the cell or complex where it is active." [PMID:13130076] |
| GO:0140489 | 0.000260773 | 108 / 361 | "The action of a molecule that provides a shape or a sequence mimicking or complementary to the final product, providing template for copying the original molecule's shape or sequence." [GOC:pg] |
| GO:0140522 | 0.000260773 | 108 / 361 | "The activity of joining two lipid bilayers to form a single membrane." [PMID:10332732, PMID:11493675, PMID:12600315, PMID:32100701, PMID:32641474] |
| GO:0140691 | 0.000260773 | 108 / 361 | "Binding to an RNA or an RNA-containing complex to assist the folding process." [PMID:31165735] |
| GO:0140776 | 0.000260773 | 108 / 361 | "A molecular function that involves direct binding to one of the subunits of a protein-containing complex and promoting the dissociation of one or many subunits. This often happens by changing the conformation of the protein being bound, which decreases its affinity for th |
| GO:0140777 | 0.000260773 | 108 / 361 | "A molecular function that involves direct binding to one of the subunits of a protein-containing complex, thus preventing an interaction with a factor that would promote dissociation of the complex." [PMID:34040253] |
| GO:0140911 | 0.000260773 | 108 / 361 | "An activity in which a protein is inserted into the membrane of another cell where it forms transmembrane pores. Pores disrupts the integrity of the cell membrane, resulting in deregulated ion homeostasis, cellular dysfunction, and can result in cell death." [PMID:1406491, P |
| GO:0140912 | 0.000260773 | 108 / 361 | "Binding to a membrane and increasing its permeability. This may lead to cell membrane lysis and cell content release." [PMID:34496967] |
| GO:0141047 | 0.000260773 | 108 / 361 | "A molecular function exhibited by a protein that is covalently attached (AKA tagged or conjugated) to another molecule (for example a protein or a lipid) where it acts as a marker, recognized by the cellular apparatus to target the tagged protein for some cellular process sucl |
| GO:0180020 | 0.000260773 | 108 / 361 | "The activity of bending or deforming a membrane. This activity can occur by multiple mechanisms including the insertion amphipathic domains into one or both leaflets." [PMID:19780639] |
| GO:0180024 | 0.000260773 | 108 / 361 | "An activity in which a protein, or group of proteins, acts to restrict the size of a fenestration in a membrane or membrane system (i.e. restrict the diameter of a hole in a membrane). An example of this is the molecular function is performed by ESCRT-III proteins at the nuclea |
| GO:0140657 | 0.000280301 | 114 / 375 | "A molecular function characterized by the coupling of ATP hydrolysis to other steps of a reaction mechanism to make the reaction energetically favorable, for example to catalyze a reaction or drive transport against a concentration gradient." [PMID:24878343, PMID:25750732 |
| GO:0031490 | 0.000593117 | 2 / 324 | "Binding to DNA that is assembled into chromatin." [GOC:mah] |
| GO:0003774 | 0.00072763 | 110 / 362 | "Generation of force resulting in movement, for example along a microfilament or microtubule, or in torque resulting in membrane scission or rotation of a flagellum. The energy required is obtained either from the hydrolysis of a nucleoside triphosphate or by an electrochen |
| GO:0003690 | 0.000808487 | 2 / 314 | "Binding to double-stranded DNA." [GOC:elh, GOC:vw] |
| GO:0003697 | 0.000808487 | 2 / 314 | "Binding to single-stranded DNA." [GOC:elh, GOC:vw, PMID:22976174] |
| GO:0003676 | 0.000897463 | 7 / 531 | "Binding to a nucleic acid." [GOC:jl] |
| GO:0003677 | 0.001066305 | 7 / 524 | "Any molecular function by which a gene product interacts selectively and non-covalently with DNA (deoxyribonucleic acid)." [GOC:dph, GOC:jl, GOC:tb, GOC:vw] |
| GO:0003684 | 0.001098784 | 2 / 304 | "Binding to damaged DNA." [GOC:jl] |
| GO:0050660 | 0.001290835 | 21 / 315 | "Binding to FAD, flavin-adenine dinucleotide, the coenzyme or the prosthetic group of various flavoprotein oxidoreductase enzymes, in either the oxidized form, FAD, or the reduced form, FADH2." [GOC:ai, GOC:imk, ISBN:0198506732] |
| GO:0000217 | 0.001534311 | 2 / 293 | "Binding to a DNA secondary structure element such as a four-way junction, a bubble, a loop, Y-form DNA, or a double-strand/single-strand junction." [GOC:krc] |
| GO:0043273 | 0.001605077 | 9 / 84 | "Catalysis of the reaction: CTP + H2O = CDP + H+ + phosphate. May or may not be coupled to another reaction." [RHEA:29387] |
| GO:0050339 | 0.001605077 | 9 / 84 | "Catalysis of the reaction: dTTP + H2O = dTDP + H+ + phosphate." [RHEA:19013] |
| GO:0103023 | 0.001605077 | 9 / 84 | "Catalysis of the reaction: ITP + H2O = IDP + H+ + phosphate." [GOC:pz, PMID:16216582, RHEA:28330] |
| GO:0008301 | 0.001629653 | 2 / 291 | "The activity of binding selectively and non-covalently to and distorting the original structure of DNA, typically a straight helix, into a bend, or increasing the bend if the original structure was intrinsically bent due to its sequence." [GOC:krc, GOC:vw, PMID:10710711, PMID:1903 |
| GO:0045027 | 0.001629653 | 2 / 291 | "Binding to DNA ends exposed by the creation of double-strand breaks (DSBs)." [GOC:jl] |
| GO:0070336 | 0.001629653 | 2 / 291 | "Binding to a flap structure in DNA. A DNA flap structure is one in which a single-stranded length of DNA or RNA protrudes from a double-stranded DNA molecule." [GOC:mah, PMID:15189154] |
| GO:0000497 | 0.00167944 | 2 / 290 | "Binding to nucleic acid via hydrogen bonds between the bases of a gene product molecule and the bases of a target DNA molecule." [GOC:krc] |
| GO:0003681 | 0.00167944 | 2 / 290 | "Binding to DNA in a bent conformation." [GOC:jl, PMID:12627977] |
| GO:0010844 | 0.00167944 | 2 / 290 | "Binding to a genomic region which promotes recombination." [GOC:dph, GOC:tb] |
| GO:0045142 | 0.00167944 | 2 / 290 | "Binding to a DNA triple helix. The formation of triple helical DNA has been evoked in several cellular processes including transcription, replication, and recombination." [PMID:10681538] |
| GO:0051880 | 0.00167944 | 2 / 290 | "Binding to G-quadruplex DNA structures, in which groups of four guanines adopt a flat, cyclic Hoogsteen hydrogen-bonding arrangement known as a guanine tetrad. The stacking of guanine tetrads results in G-quadruplex DNA structures. G-quadruplex DNA can form under pl |

| GO_Term | P_val | Counts | BioProc_Description |
|---|---|---|---|
| GO:0009987 | 0 | 133 / 40 | "Any process that is carried out at the cellular level, but not necessarily restricted to a single cell. For example, cell communication occurs among more than one cell, but occurs at the cellular level." [GOC:go_curators, GOC:isa_complete] |
| GO:0019748 | 0.000897342 | 23 / 358 | "The chemical reactions and pathways resulting in many of the chemical changes of compounds that are not necessarily required for growth and maintenance of cells, and are often unique to a taxon. In multicellular organisms secondary metabolism is generally carried out in |
| GO:0055114 | 0.000985451 | 27 / 453 | "OBSOLETE. A metabolic process that results in the removal or addition of one or more electrons to or from a substance, with or without the concomitant removal or addition of a proton or protons." [GOC:dhl, GOC:ecd, GOC:jh2, GOC:jid, GOC:mlg, GOC:rph] |
| GO:0070813 | 0.001117409 | 2 / 2 | "The chemical reactions and pathways involving hydrogen sulfide, H2S." [GOC:mah] |
| GO:1903785 | 0.001117409 | 2 / 2 | "The directed movement of L-valine across a membrane." [GO_REF:0000069, GOC:TermGenie, PMID:20944394] |
| GO:1903826 | 0.001117409 | 2 / 2 | "The directed movement of L-arginine across a membrane." [GO_REF:0000069, GOC:TermGenie, PMID:18357653, PMID:22822152, PMID:8195186] |
| GO:0044281 | 0.001485884 | 20 / 305 | "The chemical reactions and pathways involving small molecules, any low molecular weight, monomeric, non-encoded molecule." [GOC:curators, GOC:pde, GOC:vw] |
| GO:0016999 | 0.001929961 | 19 / 290 | "The chemical reactions and pathways involving an antibiotic, a substance produced by or derived from certain fungi, bacteria, and other organisms, that can destroy or inhibit the growth of other microorganisms." [GOC:cab2] |
| GO:0018933 | 0.001929961 | 19 / 290 | "The chemical reactions and pathways involving nicotine, (S)(-)-3-(1-methyl-2-pyrrolidinyl)pyridine." [GOC:sm, ISBN:0198547684] |
| GO:0032963 | 0.001929961 | 19 / 290 | "The chemical reactions and pathways involving collagen, any of a group of fibrous proteins of very high tensile strength that form the main component of connective tissue in animals. Collagen is highly enriched in glycine (some regions are 33% glycine) and proline, occurring |
| GO:0042414 | 0.001929961 | 19 / 290 | "The chemical reactions and pathways involving epinephrine, a hormone produced by the medulla of the adrenal glands that increases heart activity, improves the power and prolongs the action of muscles, and increases the rate and depth of breathing. It is synthesized by th |
| GO:0042445 | 0.001929961 | 19 / 290 | "The chemical reactions and pathways involving any hormone, naturally occurring substances secreted by specialized cells that affects the metabolism or behavior of other cells possessing functional receptors for the hormone." [GOC:jl] |
| GO:0045730 | 0.001929961 | 19 / 290 | "A phase of elevated metabolic activity, during which oxygen consumption increases; this leads to the production, by an NADH dependent system, of hydrogen peroxide (H2O2), superoxide anions and hydroxyl radicals." [ISBN:0198506732] |
| GO:0140651 | 0.001929961 | 19 / 290 | "The phosphorylation and dephosphorylation of creatine in a futile cycle, which dissipates the high energy charge of phosphocreatine as heat without performing any mechanical or chemical work. The futile creatine cycle takes place in thermogenic fat cells and is part of adap |
| GO:1901275 | 0.001929961 | 19 / 290 | "The chemical reactions and pathways involving tartrate." [GOC:TermGenie, GOC:yaf, UniPathway:UPA00839] |
| GO:1901286 | 0.001929961 | 19 / 290 | "The chemical reactions and pathways involving iron-sulfur-molybdenum cofactor." [GOC:TermGenie, GOC:yaf, UniPathway:UPA00782] |
| GO:1901764 | 0.001929961 | 19 / 290 | "The chemical reactions and pathways involving phosphinothricin." [GOC:TermGenie, GOC:yaf] |
| GO:1901902 | 0.001929961 | 19 / 290 | "The chemical reactions and pathways involving tyrocidine." [GOC:TermGenie, GOC:yaf, PMID:9352938] |
| GO:1902421 | 0.001929961 | 19 / 290 | "The chemical reactions and pathways involving H2 (dihydrogen)." [GOC:mengo_curators, GOC:TermGenie, PMID:20395274, PMID:20692761] |
| GO:1990845 | 0.001929961 | 19 / 290 | "The regulated production of heat in response to short term environmental changes, such as stress, diet or reduced temperature." [PMID:17260010, PMID:20363363] |
| GO:0009056 | 0.002000593 | 19 / 291 | "The chemical reactions and pathways resulting in the breakdown of substances, including the breakdown of carbon compounds with the liberation of energy for use by the cell or organism." [ISBN:0198547684] |
| GO:0051189 | 0.002000593 | 19 / 291 | "The chemical reactions and pathways involving a prosthetic group, the non-amino acid portion of certain protein molecules. Prosthetic groups may be inorganic or organic and are usually required for the biological activity of the protein." [GOC:ai] |
| GO:0042440 | 0.002073296 | 19 / 292 | "The chemical reactions and pathways involving pigment, any general or particular coloring matter in living organisms, e.g. melanin." [GOC:jl, ISBN:0198506732] |
| GO:0070085 | 0.002073296 | 19 / 292 | "The covalent attachment and further modification of carbohydrate residues to a substrate molecule." [GOC:hjd, GOC:mah] |
| GO:0070988 | 0.002073296 | 19 / 292 | "The process of removing one or more methyl groups from a molecule." [GOC:BHF, GOC:rl] |
| GO:0032259 | 0.002148113 | 19 / 293 | "The process in which a methyl group is covalently attached to a molecule." [GOC:mah] |
| GO:0006807 | 0.002734612 | 19 / 300 | "The chemical reactions and pathways involving organic or inorganic compounds that contain nitrogen." [GOC:jl, ISBN:0198506732] |
| GO:0000954 | 0.003455785 | 4 / 20 | "The chemical reactions and pathways involving the catabolism of amino acids to produce carboxylic acids with one carbon less than the starting amino acid. In S. cerevisiae, this is known to occur for leucine, isoleucine, valine, methionine, phenylalanine, tyrosine, or tryptoph |
| GO:0006636 | 0.003455785 | 4 / 20 | "The chemical reactions and pathways resulting in the formation of an unsaturated fatty acid, any fatty acid containing one or more double bonds between carbon atoms." [GOC:mah, MetaCyc:PWY-762, MetaCyc:PWY-782] |
| GO:0042759 | 0.003455785 | 4 / 20 | "The chemical reactions and pathways resulting in the formation of a long-chain fatty acid. A long-chain fatty acid has an aliphatic tail containing 13 to 22 carbons." [PMID:18390550] |
| GO:0051790 | 0.003455785 | 4 / 20 | "The chemical reactions and pathways resulting in the formation of a short-chain fatty acid. A short-chain fatty acid has an aliphatic tail containing fewer than 6 carbons." [Wikipedia:Fatty_acid_metabolism] |
| GO:0071768 | 0.003455785 | 4 / 20 | "The chemical reactions and pathways resulting in the formation of mycolic acids, beta-hydroxy fatty acids with a long alpha-alkyl side chain." [GOC:mah, MetaCyc:PWYG-321, PMID:15653820] |
| GO:1902321 | 0.003455785 | 4 / 20 | "The chemical reactions and pathways resulting in the formation of methyl-branched fatty acid." [GOC:kmv, GOC:TermGenie, PMID:15340492] |
| GO:1903966 | 0.003455785 | 4 / 20 | "The chemical reactions and pathways resulting in the formation of monounsaturated fatty acid." [GO_REF:0000068, GOC:hjd, GOC:TermGenie, PMID:16443825] |
| GO:0044578 | 0.004129224 | 4 / 21 | "The chemical reactions and pathway resulting in the formation of butyryl-CoA." [GOC:jl] |
| GO:0051792 | 0.004129224 | 4 / 21 | "The chemical reactions and pathways resulting in the formation of a medium-chain fatty acid. A medium-chain fatty acid has an aliphatic tail containing 6 to 12 carbons." [Wikipedia:Fatty_acid_metabolism] |
| GO:0009107 | 0.004881647 | 4 / 22 | "The chemical reactions and pathways resulting in the formation of lipoate, 1,2-dithiolane-3-pentanoate, the anion derived from lipoic acid." [GOC:ai, ISBN:0198506732] |
| GO:0008152 | 0.005425986 | 136 / 39 | "The chemical reactions and pathways, including anabolism and catabolism, by which living organisms transform chemical substances. Metabolic processes typically transform small molecules, but also include macromolecular processes such as DNA repair and replication, and |
| GO:0030497 | 0.005715901 | 4 / 23 | "The elongation of a fatty acid chain by the sequential addition of two-carbon units." [ISBN:0716720094] |
| GO:0042761 | 0.005715901 | 4 / 23 | "The chemical reactions and pathways resulting in the formation of a very long-chain fatty acid. A very long-chain fatty acid has an aliphatic tail containing more than 22 carbons." [PMID:7744868] |
| GO:0071704 | 0.006084307 | 23 / 423 | "The chemical reactions and pathways involving an organic substance, any molecular entity containing carbon." [GOC:mah] |
| GO:0000103 | 0.006268302 | 2 / 4 | "The pathways by which inorganic sulfate is processed and incorporated into sulfated compounds." [GOC:jl] |
| GO:0010134 | 0.006268302 | 2 / 4 | "The pathway by which inorganic sulfate is activated, reduced and incorporated into sulfated compounds, where the activated sulfate, adenylyl-sulfate, is reduced to sulfite by the activity of adenylyl-sulfate reductase." [EC:1.8.99.2] |
| GO:0019379 | 0.006268302 | 2 / 4 | "The pathway by which inorganic sulfate is processed and incorporated into sulfated compounds, where the phosphoadenylyl sulfate reduction step is catalyzed by the enzyme phosphoadenylyl-sulfate reductase (thioredoxin) (EC:1.8.4.8)." [GOC:jl] |
| GO:0089709 | 0.006268302 | 2 / 4 | "The directed movement of L-histidine across a membrane." [PMID:21307582] |
| GO:0006412 | 0.006477584 | 0 / 145 | "The cellular metabolic process in which a protein is formed, using the sequence of a mature mRNA or circRNA molecule to specify the sequence of amino acids in a polypeptide chain. Translation is mediated by the ribosome, and begins with the formation of a ternary comple |
| GO:0044237 | 0.006489318 | 22 / 401 | "The chemical reactions and pathways by which individual cells transform chemical substances." [GOC:go_curators] |
| GO:0006357 | 0.007165077 | 4 / 332 | "Any process that modulates the frequency, rate or extent of transcription mediated by RNA polymerase II." [GOC:go_curators, GOC:txnOH] |
| GO:0006631 | 0.008188868 | 5 / 40 | "The chemical reactions and pathways involving fatty acids, aliphatic monocarboxylic acids liberated from naturally occurring fats and oils by hydrolysis." [ISBN:0198547684] |
| GO:0009058 | 0.008195255 | 19 / 337 | "The chemical reactions and pathways resulting in the formation of substances; typically the energy-requiring part of metabolism in which simpler substances are transformed into more complex ones." [GOC:curators, ISBN:0198547684] |

| GO_Term | P_val | Counts | CellComp_Description |
|---|---|---|---|
| GO:0008152 | 0.000428929 | 6 / 505 | "The chemical reactions and pathways, including anabolism and catabolism, by which living organisms transform chemical substances. Metabolic processes typically transform small molecules, but also include macromolecular processes such as DNA repair and replication, and |
| GO:0008150 | 0.000545646 | 6 / 496 | "A biological process is the execution of a genetically-encoded biological module or program. It consists of all the steps required to achieve the specific biological objective of the module. A biological process is accomplished by a particular set of molecular functions carried ou |
| GO:0040007 | 0.000875863 | 6 / 478 | "The increase in size or mass of an entire organism, a part of an organism or a cell." [GOC:bf, GOC:ma] |
| GO:0140513 | 0.001096454 | 7 / 509 | "A stable assembly of two or more macromolecules, i.e. proteins, nucleic acids, carbohydrates or lipids, in which at least one component is a protein and the constituent parts function together in the nucleus." [GOC:pg] |
| GO:0009987 | 0.001123971 | 7 / 508 | "Any process that is carried out at the cellular level, but not necessarily restricted to a single cell. For example, cell communication occurs among more than one cell, but occurs at the cellular level." [GOC:go_curators, GOC:isa_complete] |
| GO:0022414 | 0.001133725 | 6 / 468 | "A biological process that directly contributes to the process of producing new individuals by one or two organisms. The new individuals inherit some proportion of their genetic material from the parent or parents." [GOC:dph, GOC:isa_complete] |
| GO:0098754 | 0.001133725 | 6 / 468 | "Any process that reduces or removes the toxicity of a toxic substance. These may include transport of the toxic substance away from sensitive areas and to compartments or complexes whose purpose is sequestration of the toxic substance." [GOC:dos] |
| GO:0050896 | 0.001193261 | 6 / 466 | "Any process that results in a change in state or activity of a cell or an organism (in terms of movement, secretion, enzyme production, gene expression, etc.) as a result of a stimulus. The process begins with detection of the stimulus and ends with a change in state or activity |
| GO:0000003 | 0.001224125 | 6 / 465 | "The production of new individuals that contain some portion of genetic material inherited from one or more parent organisms." [GOC:go_curators, GOC:isa_complete, GOC:jl, ISBN:0198506732] |
| GO:0002376 | 0.001224125 | 6 / 465 | "Any process involved in the development or functioning of the immune system, an organismal system for calibrated responses to potential internal or invasive threats." [GO_REF:0000022, GOC:add] |
| GO:0016032 | 0.001224125 | 6 / 465 | "A multi-organism process in which a virus is a participant. The other participant is the host. Includes infection of a host cell, replication of the viral genome, and assembly of progeny virus particles. In some cases the viral genetic material may integrate into the host genome a |
| GO:0032501 | 0.001224125 | 6 / 465 | "Any biological process, occurring at the level of a multicellular organism, pertinent to its function." [GOC:curators, GOC:dph, GOC:isa_complete, GOC:tb] |
| GO:0032502 | 0.001224125 | 6 / 465 | "A biological process whose specific outcome is the progression of an integrated living unit: an anatomical structure (which may be a subcellular structure, cell, tissue, or organ), or organism over time from an initial condition to a later condition." [GOC:isa_complete] |
| GO:0040011 | 0.001224125 | 6 / 465 | "Self-propelled movement of a cell or organism from one location to another." [GOC:dgh] |
| GO:0042592 | 0.001224125 | 6 / 465 | "Any biological process involved in the maintenance of an internal steady state." [GOC:jl, ISBN:0395825172] |
| GO:0043473 | 0.001224125 | 6 / 465 | "The accumulation of pigment in an organism, tissue or cell, either by increased deposition or by increased number of cells." [GOC:jl] |
| GO:0044419 | 0.001224125 | 6 / 465 | "Any process evolved to enable an interaction with an organism of a different species." [GOC:cc] |
| GO:0044848 | 0.001224125 | 6 / 465 | "A distinct period or stage in a biological process or cycle." [GOC:jl] |
| GO:0048511 | 0.001224125 | 6 / 465 | "Any process pertinent to the generation and maintenance of rhythms in the physiology of an organism." [GOC:jid] |
| GO:0051179 | 0.001224125 | 6 / 465 | "Any process in which a cell, a substance, or a cellular entity, such as a protein complex or organelle, is transported, tethered to or otherwise maintained in a specific location. In the case of substances, localization may also be achieved via selective degradation." [GOC:ai, GOC |
| GO:0051703 | 0.001224125 | 6 / 465 | "Any process in which an organism has an effect on an organism of the same species." [GOC:ai] |
| GO:0065007 | 0.001224125 | 6 / 465 | "Any process that modulates a measurable attribute of any biological process, quality or function." [GOC:dph, GOC:isa_complete, GOC:mah, GOC:pr, GOC:vw] |
| GO:0031981 | 0.001335582 | 7 / 501 | "The volume enclosed by the nuclear inner membrane." [GOC:mah, GOC:pz] |
| GO:0043231 | 0.002895536 | 10 / 579 | "Organized structure of distinctive morphology and function, bounded by a single or double lipid bilayer membrane and occurring within the cell. Includes the nucleus, mitochondria, plastids, vacuoles, and vesicles. Excludes the plasma membrane." [GOC:go_curators] |
| GO:0005635 | 0.004434012 | 7 / 450 | "The double lipid bilayer enclosing the nucleus and separating its contents from the rest of the cytoplasm; includes the intermembrane space, a gap of width 20-40 nm (also called the perinuclear space)." [ISBN:0198547684] |
| GO:0005634 | 0.004534742 | 7 / 449 | "A membrane-bounded organelle of eukaryotic cells in which chromosomes are housed and replicated. In most cells, the nucleus contains all of the cell's chromosomes except the organellar chromosomes, and is the site of RNA synthesis and processing. In some species, or in |
| GO:0005880 | 0.004849603 | 7 / 446 | "Any microtubule in the nucleus of a cell." [GOC:mah] |
| GO:0000943 | 0.004958907 | 7 / 445 | "A complex of the retrotransposon RNA genome, reverse transcriptase, integrase, and associated molecules required for reproduction and integration of the retrotransposon into the host genome; the main structural molecule of the nucleocapsid is often a gag protein homol |
| GO:0031039 | 0.004958907 | 7 / 445 | "A membrane-bounded organelle of ciliated protozoan cells that contains polyploid copies of a portion of the cell's complete genome. Transcription of genes occurs in macronuclei. Some ciliate species may contain multiple macronuclei per cell." [GOC:ns] |
| GO:0031040 | 0.004958907 | 7 / 445 | "A membrane-bounded organelle of ciliated protozoan cells that contains a diploid copy of the cell's complete genome. Sections of contiguous sequence in the macronucleus are often interrupted by internal eliminated sequences (IES), and may be permuted, in micronuclei. G |
| GO:0042405 | 0.004958907 | 7 / 445 | "An intranuclear focus at which aggregated proteins have been sequestered." [GOC:jl] |
| GO:0043073 | 0.004958907 | 7 / 445 | "The nucleus of a germ cell, a reproductive cell in multicellular organisms." [CL:0000586, GOC:go_curators] |
| GO:0043076 | 0.004958907 | 7 / 445 | "The nucleus of a megasporocyte, a diploid cell that undergoes meiosis to produce four megaspores, and its descendents." [GOC:jl, ISBN:0618254153] |
| GO:0045120 | 0.004958907 | 7 / 445 | "The nucleus of either the ovum or the spermatozoon following fertilization. Thus, in the fertilized ovum, there are two pronuclei, one originating from the ovum, the other from the spermatozoon that brought about fertilization; they approach each other, but do not fuse un |
| GO:0046818 | 0.004958907 | 7 / 445 | "A location in the host cell nucleus where viral proteins colocalize late in infection prior to the onset of viral DNA synthesis. More than one site can be present simultaneously." [PMID:10233976] |
| GO:0048353 | 0.004958907 | 7 / 445 | "Nucleus resulting from the fusion of the male gamete and two polar nuclei in the central cell of the embryo sac." [ISBN:0471245208] |
| GO:0048555 | 0.004958907 | 7 / 445 | "The nucleus of the generative cell, a cell contained within the pollen grain that will divide to produce two haploid sperm cells." [GOC:tair_curators] |
| GO:0048556 | 0.004958907 | 7 / 445 | "The nucleus of the microsporocyte. The microsporocyte is a diploid cell in which meiosis will occur, resulting in four microspores. A microspore is a spore that, in vascular plants, gives rise to a male gametophyte." [GOC:tair_curators, ISBN:047186840X] |
| GO:0097165 | 0.004958907 | 7 / 445 | "A dense aggregation in the nucleus composed of proteins and RNAs that appear when the cell is under stress." [GOC:ans, PMID:10359787, PMID:12865437] |
| GO:0097571 | 0.004958907 | 7 / 445 | "One of the two nuclei found in Giardia species (trophozoite stage). It is located on the left side of the cell when viewed from the dorsal side." [GOC:giardia, ISBN:0-444-81258-X] |
| GO:0097572 | 0.004958907 | 7 / 445 | "One of the two nuclei found in Giardia species (trophozoite stage). It is located on the right side of the cell when viewed from the dorsal side." [GOC:giardia, ISBN:0-444-81258-X] |
| GO:0110092 | 0.004958907 | 7 / 445 | "The area of a motile nucleus closest to the direction of movement." [GOC:al, GOC:mah, GOC:vw, PMID:15030757, PMID:24335254] |
| GO:0110093 | 0.004958907 | 7 / 445 | "The area of a motile nucleus furthest from the direction of movement." [GOC:kmv, PMID:24335254] |
| GO:0140510 | 0.004958907 | 7 / 445 | "A narrow constricted region of the nucleus that forms around the anaphase spindle during closed mitosis, and connects the main portions of the newly forming daughter nuclei." [PMID:32848252] |
| GO:1905754 | 0.004958907 | 7 / 445 | "Any nucleus that is part of a ascospore-type prospore." [GO_REF:0000064, GOC:TermGenie, PMID:26942678] |
| GO:0005872 | 0.016314545 | 2 / 6 | "Any complex that includes a dimer of molecules from the kinesin superfamily and any associated proteins, and moves towards the minus end of a microtubule." [GOC:mah] |
| GO:0005873 | 0.016314545 | 2 / 6 | "Any complex that includes a dimer of molecules from the kinesin superfamily and any associated proteins, and moves towards the plus end of a microtubule." [GOC:mah] |
| GO:0007266 | 0.016314545 | 2 / 6 | "An intracellular signaling cassette in which a small monomeric GTPase of the Rho subfamily relays a signal." [GOC:bf] |
| GO:0016601 | 0.016314545 | 2 / 6 | "An intracellular signaling cassette in which a small monomeric GTPase of the Rac subfamily relays a signal." [GOC:bf] |
| GO:0016938 | 0.016314545 | 2 / 6 | "A complex of two kinesin heavy chains and two kinesin light chains." [http://www.csfic.mi.cnr.it/centro/lines/8/ric.html] |

#### Supplementary Data S3.4.6

| GO_Term | P_val | Counts | MolFunc_Description |
|---|---|---|---|
| GO:0016491 | 4.41867E-05 | 57 / 658 | "Catalysis of an oxidation-reduction (redox) reaction, a reversible chemical reaction in which the oxidation state of an atom or atoms within a molecule is altered. One substrate acts as a hydrogen or electron donor and becomes oxidized, while the other acts as hydrogen or |
| GO:0005198 | 0.000159136 | 182 / 374 | "The action of a molecule that contributes to the structural integrity of a complex." [GOC:mah, GOC:vw] |
| GO:0043169 | 0.000382856 | 10 / 54 | "Binding to a cation, a charged atom or group of atoms with a net positive charge." [GOC:jl] |
| GO:0008199 | 0.000490818 | 9 / 46 | "Binding to a ferric iron ion, Fe(III)." [GOC:ai] |
| GO:0016717 | 0.000527906 | 5 / 14 | "Catalysis of an oxidation-reduction (redox) reaction in which hydrogen or electrons are transferred from each of two donors, and molecular oxygen is reduced to two molecules of water." [GOC:mah] |
| GO:0005215 | 0.000549383 | 183 / 372 | "Enables the directed movement of substances (such as macromolecules, small molecules, ions) into, out of or within a cell, or between cells." [GOC:ai, GOC:dgf] |
| GO:0008198 | 0.000779281 | 9 / 49 | "Binding to a ferrous iron ion, Fe(II)." [GOC:ai] |
| GO:0005506 | 0.00090071 | 9 / 50 | "Binding to an iron (Fe) ion." [GOC:ai] |
| GO:0043168 | 0.001201468 | 20 / 186 | "Binding to an anion, a charged atom or group of atoms with a net negative charge." [GOC:jl] |
| GO:0050660 | 0.001294359 | 29 / 315 | "Binding to FAD, flavin-adenine dinucleotide, the coenzyme or the prosthetic group of various flavoprotein oxidoreductase enzymes, in either the oxidized form, FAD, or the reduced form, FADH2." [GOC:ai, GOC:imk, ISBN:0198506732] |
| GO:0140110 | 0.00160204 | 182 / 367 | "A molecular function that controls the rate, timing and/or magnitude of gene transcription. The function of transcriptional regulators is to modulate gene expression at the transcription step so that they are expressed in the right cell at the right time and in the right amour |
| GO:0005488 | 0.001854863 | 180 / 363 | "The selective, non-covalent, often stoichiometric, interaction of a molecule with one or more specific sites on another molecule." [GOC:ceb, GOC:mah, ISBN:0198506732] |
| GO:0000166 | 0.002093188 | 25 / 267 | "Binding to a nucleotide, any compound consisting of a nucleoside that is esterified with (ortho)phosphate or an oligophosphate at any hydroxyl group on the ribose or deoxyribose." [GOC:mah, ISBN:0198547684] |
| GO:0060090 | 0.002125328 | 180 / 363 | "The binding activity of a molecule that brings together two or more molecules through a selective, non-covalent, often stoichiometric interaction, permitting those molecules to function in a coordinated way." [GOC:mtg_MIT_16mar07, GOC:vw] |
| GO:0140223 | 0.002624909 | 180 / 362 | "A molecular function required for core promoter activity that mediates the assembly of the RNA polymerase holoenzyme at promoter DNA to form the pre-initiation complex (PIC). General transcription factors (GTFs) bind to and open promoter DNA, initiate RNA synthesis a |
| GO:0016209 | 0.002693567 | 180 / 362 | "Inhibition of the reactions brought about by dioxygen (O2) or peroxides. Usually the antioxidant is effective because it can itself be more easily oxidized than the substance protected. The term is often applied to components that can trap free radicals, thereby breaking the |
| GO:0031420 | 0.002729075 | 8 / 48 | "Binding to an alkali metal ion; alkali metals are those elements in group Ia of the periodic table, with the exception of hydrogen." [GOC:mah] |
| GO:0032791 | 0.002729075 | 8 / 48 | "Binding to lead (Pb) ions." [GOC:mah] |
| GO:0046911 | 0.002729075 | 8 / 48 | "The formation of bonds from two or more atoms within the same ligand to a metal atom in complexes in which the metal is part of a ring." [ISBN:0198506732, ISBN:0716731363] |
| GO:0016614 | 0.002818525 | 34 / 411 | "Catalysis of an oxidation-reduction (redox) reaction in which a CH-OH group act as a hydrogen or electron donor and reduces a hydrogen or electron acceptor." [GOC:ai] |
| GO:0098772 | 0.002835249 | 180 / 361 | "A molecular function regulator regulates the activity of its target via non-covalent binding that does not result in covalent modification to the target. Examples of molecular function regulators include regulatory subunits of multimeric enzymes and channels. Mechanisms of |
| GO:0009055 | 0.002853275 | 182 / 365 | "A molecular function representing the directed movement of electrons from one molecular entity to another, typically mediated by electron carriers or acceptors, resulting in the transfer of energy and/or the reduction-oxidation (redox) transformation of chemical species. T |
| GO:0000248 | 0.002873194 | 2 / 2 | "Catalysis of the reaction: 5,7,24(28)-ergostatrienol + O2 + NADPH = 5,7,22,24(28)-ergostatetraenol + 2 H2O + NADP+." [MetaCyc:RXN3O-227] |
| GO:0042284 | 0.002873194 | 2 / 2 | "Catalysis of the reaction: a dihydroceramide + 2 ferrocytochrome b5 + O2 + 2 H+ -> a sphingosine ceramide (aka (4E)-sphing-4-enine ceramide) + 2 ferricytochrome b5 + 2 H2O." [PMID:12417141, RHEA:46544] |
| GO:0050613 | 0.002873194 | 2 / 2 | "Catalysis of the reaction: NADP+ + 4,4-dimethyl-5-alpha-cholesta-8,24-dien-3-beta-ol = NADPH + H+ + 4,4-dimethyl-5-alpha-cholesta-8,14,24-trien-3-beta-ol." [EC:1.3.1.70] |
| GO:0045735 | 0.002908315 | 180 / 361 | "Functions in the storage of nutritious substrates." [GOC:ai] |
| GO:0038024 | 0.003059002 | 180 / 361 | "Binding specifically to a substance (cargo) to deliver it to a transport vesicle. Cargo receptors span a membrane (either the plasma membrane or a vesicle membrane), binding simultaneously to cargo molecules and coat adaptors, to efficiently recruit soluble proteins to nasc |
| GO:0044183 | 0.003136664 | 180 / 361 | "Binding to a protein or a protein-containing complex to assist the protein folding process." [GOC:mtg_cambridge_2009] |
| GO:0045182 | 0.003136664 | 180 / 361 | "Any molecular function involved in the initiation, activation, perpetuation, repression or termination of polypeptide synthesis at the ribosome." [GOC:ai] |
| GO:0140104 | 0.003215899 | 180 / 361 | "Directly binding to a specific ion or molecule and delivering it either to an acceptor molecule or to a specific location." [GOC:molecular_function_refactoring, GOC:pdt] |
| GO:0060089 | 0.00329673 | 180 / 361 | "A compound molecular function in which an effector function is controlled by one or more regulatory components." [GOC:dos, GOC:pdt] |
| GO:0090729 | 0.00329673 | 180 / 361 | "Interacting selectively with one or more biological molecules in another (target) organism, initiating pathogenesis (leading to an abnormal, generally detrimental state) in the target organism. The activity should refer to an evolved function of the active gene product, i.e. one |
| GO:0140299 | 0.00329673 | 180 / 361 | "Binding to a small molecule and eliciting a change in the protein's activity in response to the intracellular level of that small molecule." [PMID:26328879] |
| GO:0140313 | 0.00329673 | 180 / 361 | "Binding to a specific molecule to prevent it from interacting with other partners or to inhibit its localization to the area of the cell or complex where it is active." [PMID:13130076] |
| GO:0140489 | 0.00329673 | 180 / 361 | "The action of a molecule that provides a shape or a sequence mimicking or complementary to the final product, providing template for copying the original molecule's shape or sequence." [GOC:pg] |
| GO:0140522 | 0.00329673 | 180 / 361 | "The activity of joining two lipid bilayers to form a single membrane." [PMID:10332732, PMID:11493675, PMID:12600315, PMID:32100701, PMID:32641474] |
| GO:0140691 | 0.00329673 | 180 / 361 | "Binding to an RNA or an RNA-containing complex to assist the folding process." [PMID:31165735] |
| GO:0140776 | 0.00329673 | 180 / 361 | "A molecular function that involves direct binding to one of the subunits of a protein-containing complex and promoting the dissociation of one or many subunits. This often happens by changing the conformation of the protein being bound, which decreases its affinity for t |
| GO:0140777 | 0.00329673 | 180 / 361 | "A molecular function that involves direct binding to one of the subunits of a protein-containing complex, thus preventing an interaction with a factor that would promote dissociation of the complex." [PMID:34040253] |
| GO:0140911 | 0.00329673 | 180 / 361 | "An activity in which a protein is inserted into the membrane of another cell where it forms transmembrane pores. Pores disrupts the integrity of the cell membrane, resulting in deregulated ion homeostasis, cellular dysfunction, and can result in cell death." [PMID:1406491, |
| GO:0140912 | 0.00329673 | 180 / 361 | "Binding to a membrane and increasing its permeability. This may lead to cell membrane lysis and cell content release." [PMID:34496967] |
| GO:0141047 | 0.00329673 | 180 / 361 | "A molecular function exhibited by a protein that is covalently attached (AKA tagged or conjugated) to another molecule (for example a protein or a lipid) where it acts as a marker, recognized by the cellular apparatus to target the tagged protein for some cellular process su |
| GO:0180020 | 0.00329673 | 180 / 361 | "The activity of bending or deforming a membrane. This activity can occur by multiple mechanisms including the insertion amphipathic domains into one or both leaflets." [PMID:19780639] |
| GO:0180024 | 0.00329673 | 180 / 361 | "An activity in which a protein, or group of proteins, acts to restrict the size of a fenestration in a membrane or membrane system (i.e. restrict the diameter of a hole in a membrane). An example of this is the molecular function is performed by ESCRT-III proteins at the nucle |
| GO:0070704 | 0.003560813 | 4 / 13 | "Catalysis of the introduction of a double bond into a sterol molecule." [GOC:mah, GOC:vw] |
| GO:0016705 | 0.00413068 | 32 / 391 | "Catalysis of an oxidation-reduction (redox) reaction in which hydrogen or electrons are transferred from each of two donors, and molecular oxygen is reduced or incorporated into a donor." [GOC:mah] |
| GO:0003676 | 0.004179767 | 17 / 531 | "Binding to a nucleic acid." [GOC:jl] |
| GO:0140657 | 0.004192244 | 189 / 375 | "A molecular function characterized by the coupling of ATP hydrolysis to other steps of a reaction mechanism to make the reaction energetically favorable, for example to catalyze a reaction or drive transport against a concentration gradient." [PMID:24878343, PMID:2575073 |
| GO:0000287 | 0.004231629 | 9 / 63 | "Binding to a magnesium (Mg) ion." [GOC:ai] |
| GO:0008556 | 0.00430743 | 3 / 7 | "Enables the transfer of a solute or solutes from one side of a membrane to the other according to the reaction: ATP + H2O + K+(out) = ADP + phosphate + K+(in)." [RHEA:16777] |

| GO_Term | P_val | Counts | BioProc_Description |
|---|---|---|---|
| GO:0009987 | 0 | 212 / 405 | "Any process that is carried out at the cellular level, but not necessarily restricted to a single cell. For example, cell communication occurs among more than one cell, but occurs at the cellular level." [GOC:go_curators, GOC:isa_complete] |
| GO:0008152 | 0.000231236 | 216 / 390 | "The chemical reactions and pathways, including anabolism and catabolism, by which living organisms transform chemical substances. Metabolic processes typically transform small molecules, but also include macromolecular processes such as DNA repair and replication, and |
| GO:0055114 | 0.002039141 | 37 / 453 | "OBSOLETE. A metabolic process that results in the removal or addition of one or more electrons to or from a substance, with or without the concomitant removal or addition of a proton or protons." [GOC:dhl, GOC:ecd, GOC:jh2, GOC:jid, GOC:mlg, GOC:rph] |
| GO:0017121 | 0.00253873 | 3 / 6 | "The movement of a population of phospholipid molecules from one leaflet of the plasma membrane bilayer to the opposite leaflet, resulting in loss of lipid asymmetry and surface exposure of phosphatidylserine (PS) and phosphatidylethanolamine (PE)." [GOC:cjm, PMID:200 |
| GO:0018868 | 0.00253873 | 3 / 6 | "The chemical reactions and pathways involving 2-aminobenzenesulfonate, aniline-o-sulfonic acid, an aromatic sulfonate used in organic synthesis and in the manufacture of various dyes and medicines." [UM-BBD_pathwayID:abs] |
| GO:0006412 | 0.002605421 | 1 / 145 | "The cellular metabolic process in which a protein is formed, using the sequence of a mature mRNA or circRNA molecule to specify the sequence of amino acids in a polypeptide chain. Translation is mediated by the ribosome, and begins with the formation of a ternary comple |
| GO:0002181 | 0.002635433 | 0 / 107 | "The chemical reactions and pathways resulting in the formation of a protein in the cytoplasm. This is a ribosome-mediated process in which the information in messenger RNA (mRNA) is used to specify the sequence of amino acids in the protein." [GOC:hjd] |
| GO:0006418 | 0.002787966 | 0 / 106 | "The synthesis of aminoacyl tRNA by the formation of an ester bond between the 3'-hydroxyl group of the most 3' adenosine of the tRNA and the alpha carboxylic acid group of an amino acid, to be used in ribosome-mediated polypeptide synthesis." [GOC:ma] |
| GO:0019572 | 0.002826389 | 2 / 2 | "The chemical reactions and pathways resulting in the breakdown of L-arabinose, the L-enantiomer of arabino-pentose." [GOC:jsg, GOC:mah] |
| GO:0042843 | 0.002826389 | 2 / 2 | "The chemical reactions and pathways resulting in the breakdown of D-xylose, a naturally occurring plant polysaccharide." [ISBN:0198506732] |
| GO:0044577 | 0.002826389 | 2 / 2 | "The anaerobic chemical reactions and pathways resulting in the breakdown of xylose, an aldopentose, where one of the resulting products is ethanol." [GOC:mengo_curators, GOC:tt] |
| GO:0070813 | 0.002826389 | 2 / 2 | "The chemical reactions and pathways involving hydrogen sulfide, H2S." [GOC:mah] |
| GO:0019748 | 0.003450676 | 30 / 358 | "The chemical reactions and pathways resulting in many of the chemical changes of compounds that are not necessarily required for growth and maintenance of cells, and are often unique to a taxon. In multicellular organisms secondary metabolism is generally carried out in |
| GO:0051189 | 0.005085319 | 25 / 291 | "The chemical reactions and pathways involving a prosthetic group, the non-amino acid portion of certain protein molecules. Prosthetic groups may be inorganic or organic and are usually required for the biological activity of the protein." [GOC:ai] |
| GO:0006414 | 0.005785902 | 0 / 93 | "The successive addition of amino acid residues to a nascent polypeptide chain during protein biosynthesis." [GOC:ems] |
| GO:0032543 | 0.006119526 | 0 / 92 | "The chemical reactions and pathways resulting in the formation of a protein in a mitochondrion. This is a ribosome-mediated process in which the information in messenger RNA (mRNA) is used to specify the sequence of amino acids in the protein; the mitochondrion has its |
| GO:0043043 | 0.007656697 | 0 / 88 | "The chemical reactions and pathways resulting in the formation of peptides, compounds of 2 or more (but usually less than 100) amino acids where the alpha carboxyl group of one is bound to the alpha amino group of another. This may include the translation of a precurso |
| GO:0019323 | 0.008031352 | 2 / 3 | "The chemical reactions and pathways resulting in the breakdown of a pentose, any monosaccharide with a chain of five carbon atoms in the molecule." [ISBN:0198506732] |
| GO:0019571 | 0.008031352 | 2 / 3 | "The chemical reactions and pathways resulting in the breakdown of D-arabinose, the D-enantiomer of arabino-pentose." [GOC:jsg, GOC:mah] |
| GO:0042732 | 0.008031352 | 2 / 3 | "The chemical reactions and pathways involving D-xylose, a naturally occurring plant polysaccharide." [ISBN:0198506732] |
| GO:0044237 | 0.008184031 | 31 / 401 | "The chemical reactions and pathways by which individual cells transform chemical substances." [GOC:go_curators] |
| GO:0044281 | 0.008336055 | 25 / 305 | "The chemical reactions and pathways involving small molecules, any low molecular weight, monomeric, non-encoded molecule." [GOC:curators, GOC:pde, GOC:vw] |
| GO:0006415 | 0.008563778 | 0 / 86 | "The process resulting in the release of a polypeptide chain from the ribosome, usually in response to a termination codon (UAA, UAG, or UGA in the universal genetic code)." [GOC:hjd, ISBN:019879276X] |
| GO:0016999 | 0.008564729 | 24 / 290 | "The chemical reactions and pathways involving an antibiotic, a substance produced by or derived from certain fungi, bacteria, and other organisms, that can destroy or inhibit the growth of other microorganisms." [GOC:cab2] |
| GO:0018933 | 0.008564729 | 24 / 290 | "The chemical reactions and pathways involving nicotine, (S)(-)-3-(1-methyl-2-pyrrolidinyl)pyridine." [GOC:sm, ISBN:0198547684] |
| GO:0032963 | 0.008564729 | 24 / 290 | "The chemical reactions and pathways involving collagen, any of a group of fibrous proteins of very high tensile strength that form the main component of connective tissue in animals. Collagen is highly enriched in glycine (some regions are 33% glycine) and proline, occurring |
| GO:0042414 | 0.008564729 | 24 / 290 | "The chemical reactions and pathways involving epinephrine, a hormone produced by the medulla of the adrenal glands that increases heart activity, improves the power and prolongs the action of muscles, and increases the rate and depth of breathing. It is synthesized by th |
| GO:0042445 | 0.008564729 | 24 / 290 | "The chemical reactions and pathways involving any hormone, naturally occurring substances secreted by specialized cells that affects the metabolism or behavior of other cells possessing functional receptors for the hormone." [GOC:jl] |
| GO:0045730 | 0.008564729 | 24 / 290 | "A phase of elevated metabolic activity, during which oxygen consumption increases; this leads to the production, by an NADH dependent system, of hydrogen peroxide (H2O2), superoxide anions and hydroxyl radicals." [ISBN:0198506732] |
| GO:0140651 | 0.008564729 | 24 / 290 | "The phosphorylation and dephosphorylation of creatine in a futile cycle, which dissipates the high energy charge of phosphocreatine as heat without performing any mechanical or chemical work. The futile creatine cycle takes place in thermogenic fat cells and is part of adap |
| GO:1901275 | 0.008564729 | 24 / 290 | "The chemical reactions and pathways involving tartrate." [GOC:TermGenie, GOC:yaf, UniPathway:UPA00839] |
| GO:1901286 | 0.008564729 | 24 / 290 | "The chemical reactions and pathways involving iron-sulfur-molybdenum cofactor." [GOC:TermGenie, GOC:yaf, UniPathway:UPA00782] |
| GO:1901764 | 0.008564729 | 24 / 290 | "The chemical reactions and pathways involving phosphinothricin." [GOC:TermGenie, GOC:yaf] |
| GO:1901902 | 0.008564729 | 24 / 290 | "The chemical reactions and pathways involving tyrocidine." [GOC:TermGenie, GOC:yaf, PMID:9352938] |
| GO:1902421 | 0.008564729 | 24 / 290 | "The chemical reactions and pathways involving H2 (dihydrogen)." [GOC:mengo_curators, GOC:TermGenie, PMID:20395274, PMID:20692761] |
| GO:1990845 | 0.008564729 | 24 / 290 | "The regulated production of heat in response to short term environmental changes, such as stress, diet or reduced temperature." [PMID:17260010, PMID:20363363] |
| GO:0009056 | 0.008858438 | 24 / 291 | "The chemical reactions and pathways resulting in the breakdown of substances, including the breakdown of carbon compounds with the liberation of energy for use by the cell or organism." [ISBN:0198547684] |
| GO:1990145 | 0.009056658 | 0 / 85 | "Suppression of the occurrence of translational errors, such as codon-anticodon mis-paring, during the process of translation of a protein using an mRNA template." [GOC:hjd, ISBN:9781936113460, PMID:21841312] |
| GO:0042440 | 0.00915927 | 24 / 292 | "The chemical reactions and pathways involving pigment, any general or particular coloring matter in living organisms, e.g. melanin." [GOC:jl, ISBN:0198506732] |
| GO:0070085 | 0.00915927 | 24 / 292 | "The covalent attachment and further modification of carbohydrate residues to a substrate molecule." [GOC:hjd, GOC:mah] |
| GO:0070988 | 0.00915927 | 24 / 292 | "The process of removing one or more methyl groups from a molecule." [GOC:BHF, GOC:rl] |
| GO:0032259 | 0.009467289 | 24 / 293 | "The process in which a methyl group is covalently attached to a molecule." [GOC:mah] |
| GO:0099638 | 0.009485196 | 6 / 37 | "The directed movement of proteins from the endosome to the plasma membrane in transport vesicles." [GOC:dos] |
| GO:0008104 | 0.01067483 | 6 / 38 | "Any process in which a protein is transported to, or maintained in, a specific location." [GOC:ai] |
| GO:0032544 | 0.011327007 | 0 / 81 | "The chemical reactions and pathways resulting in the formation of a protein in a plastid. This is a ribosome-mediated process in which the information in messenger RNA (mRNA) is used to specify the sequence of amino acids in the protein; the plastid has its own ribosomes |
| GO:0071891 | 0.011327007 | 0 / 81 | "An N-terminal peptidyl-proline dimethylation process that contributes to translation." [GOC:mah] |
| GO:0140241 | 0.011327007 | 0 / 81 | "Translation that occurs at the synapse." [PMID:23083742] |
| GO:0006213 | 0.011750529 | 5 / 28 | "The chemical reactions and pathways involving any pyrimidine nucleoside, one of a family of organic molecules consisting of a pyrimidine base covalently bonded to ribose (a ribonucleoside) or deoxyribose (a deoxyribonucleoside)." [GOC:jl, ISBN:0140512713] |
| GO:0009119 | 0.011750529 | 5 / 28 | "The chemical reactions and pathways involving any ribonucleoside, a nucleoside in which purine or pyrimidine base is linked to a ribose (beta-D-ribofuranose) molecule." [GOC:jl] |
| GO:0009120 | 0.011750529 | 5 / 28 | "The chemical reactions and pathways involving any one of a family of organic molecules consisting of a purine or pyrimidine base covalently bonded to a sugar deoxyribose (a deoxyribonucleoside)." [GOC:jl, ISBN:0140512713] |

| GO_Term | P_val | Counts | CellComp_Description |
|---|---|---|---|
| GO:0008152 | 0.000683625 | 14 / 505 | "The chemical reactions and pathways, including anabolism and catabolism, by which living organisms transform chemical substances. Metabolic processes typically transform small molecules, but also include macromolecular processes such as DNA repair and replication, and |
| GO:0008150 | 0.000916651 | 14 / 496 | "A biological process is the execution of a genetically-encoded biological module or program. It consists of all the steps required to achieve the specific biological objective of the module. A biological process is accomplished by a particular set of molecular functions carried ou |
| GO:0022414 | 0.001080282 | 13 / 468 | "A biological process that directly contributes to the process of producing new individuals by one or two organisms. The new individuals inherit some proportion of their genetic material from the parent or parents." [GOC:dph, GOC:isa_complete] |
| GO:0098754 | 0.001080282 | 13 / 468 | "Any process that reduces or removes the toxicity of a toxic substance. These may include transport of the toxic substance away from sensitive areas and to compartments or complexes whose purpose is sequestration of the toxic substance." [GOC:dos] |
| GO:0050896 | 0.001152855 | 13 / 466 | "Any process that results in a change in state or activity of a cell or an organism (in terms of movement, secretion, enzyme production, gene expression, etc.) as a result of a stimulus. The process begins with detection of the stimulus and ends with a change in state or activity |
| GO:0000003 | 0.001190817 | 13 / 465 | "The production of new individuals that contain some portion of genetic material inherited from one or more parent organisms." [GOC:go_curators, GOC:isa_complete, GOC:jl, ISBN:0198506732] |
| GO:0002376 | 0.001190817 | 13 / 465 | "Any process involved in the development or functioning of the immune system, an organismal system for calibrated responses to potential internal or invasive threats." [GO_REF:0000022, GOC:add] |
| GO:0016032 | 0.001190817 | 13 / 465 | "A multi-organism process in which a virus is a participant. The other participant is the host. Includes infection of a host cell, replication of the viral genome, and assembly of progeny virus particles. In some cases the viral genetic material may integrate into the host genome a |
| GO:0032501 | 0.001190817 | 13 / 465 | "Any biological process, occurring at the level of a multicellular organism, pertinent to its function." [GOC:curators, GOC:dph, GOC:isa_complete, GOC:tb] |
| GO:0032502 | 0.001190817 | 13 / 465 | "A biological process whose specific outcome is the progression of an integrated living unit: an anatomical structure (which may be a subcellular structure, cell, tissue, or organ), or organism over time from an initial condition to a later condition." [GOC:isa_complete] |
| GO:0040011 | 0.001190817 | 13 / 465 | "Self-propelled movement of a cell or organism from one location to another." [GOC:dgh] |
| GO:0042592 | 0.001190817 | 13 / 465 | "Any biological process involved in the maintenance of an internal steady state." [GOC:jl, ISBN:0395825172] |
| GO:0043473 | 0.001190817 | 13 / 465 | "The accumulation of pigment in an organism, tissue or cell, either by increased deposition or by increased number of cells." [GOC:jl] |
| GO:0044419 | 0.001190817 | 13 / 465 | "Any process evolved to enable an interaction with an organism of a different species." [GOC:cc] |
| GO:0044848 | 0.001190817 | 13 / 465 | "A distinct period or stage in a biological process or cycle." [GOC:jl] |
| GO:0048511 | 0.001190817 | 13 / 465 | "Any process pertinent to the generation and maintenance of rhythms in the physiology of an organism." [GOC:jid] |
| GO:0051179 | 0.001190817 | 13 / 465 | "Any process in which a cell, a substance, or a cellular entity, such as a protein complex or organelle, is transported, tethered to or otherwise maintained in a specific location. In the case of substances, localization may also be achieved via selective degradation." [GOC:ai, GOC |
| GO:0051703 | 0.001190817 | 13 / 465 | "Any process in which an organism has an effect on an organism of the same species." [GOC:ai] |
| GO:0065007 | 0.001190817 | 13 / 465 | "Any process that modulates a measurable attribute of any biological process, quality or function." [GOC:dph, GOC:isa_complete, GOC:mah, GOC:pr, GOC:vw] |
| GO:0140513 | 0.00124415 | 15 / 509 | "A stable assembly of two or more macromolecules, i.e. proteins, nucleic acids, carbohydrates or lipids, in which at least one component is a protein and the constituent parts function together in the nucleus." [GOC:pg] |
| GO:0009987 | 0.001283195 | 15 / 508 | "Any process that is carried out at the cellular level, but not necessarily restricted to a single cell. For example, cell communication occurs among more than one cell, but occurs at the cellular level." [GOC:go_curators, GOC:isa_complete] |
| GO:0031981 | 0.001589774 | 15 / 501 | "The volume enclosed by the nuclear inner membrane." [GOC:mah, GOC:pz] |
| GO:0040007 | 0.003130527 | 15 / 478 | "The increase in size or mass of an entire organism, a part of an organism or a cell." [GOC:bf, GOC:ma] |
| GO:0015804 | 0.003136338 | 2 / 2 | "The directed movement of neutral amino acids, amino acids with no net charge, into, out of or within a cell, or between cells, by means of some agent such as a transporter or pore." [GOC:ai] |
| GO:0015807 | 0.003136338 | 2 / 2 | "The directed movement of L-enantiomer amino acids into, out of or within a cell, or between cells, by means of some agent such as a transporter or pore." [GOC:ai, GOC:jsg, GOC:mah] |
| GO:0043231 | 0.003675756 | 20 / 579 | "Organized structure of distinctive morphology and function, bounded by a single or double lipid bilayer membrane and occurring within the cell. Includes the nucleus, mitochondria, plastids, vacuoles, and vesicles. Excludes the plasma membrane." [GOC:go_curators] |
| GO:0005880 | 0.007477262 | 15 / 446 | "Any microtubule in the nucleus of a cell." [GOC:mah] |
| GO:0000943 | 0.007672404 | 15 / 445 | "A complex of the retrotransposon RNA genome, reverse transcriptase, integrase, and associated molecules required for reproduction and integration of the retrotransposon into the host genome; the main structural molecule of the nucleocapsid is often a gag protein homol |
| GO:0031039 | 0.007672404 | 15 / 445 | "A membrane-bounded organelle of ciliated protozoan cells that contains polyploid copies of a portion of the cell's complete genome. Transcription of genes occurs in macronuclei. Some ciliate species may contain multiple macronuclei per cell." [GOC:ns] |
| GO:0031040 | 0.007672404 | 15 / 445 | "A membrane-bounded organelle of ciliated protozoan cells that contains a diploid copy of the cell's complete genome. Sections of contiguous sequence in the macronucleus are often interrupted by internal eliminated sequences (IES), and may be permuted, in micronuclei. G |
| GO:0042405 | 0.007672404 | 15 / 445 | "An intranuclear focus at which aggregated proteins have been sequestered." [GOC:jl] |
| GO:0043073 | 0.007672404 | 15 / 445 | "The nucleus of a germ cell, a reproductive cell in multicellular organisms." [CL:0000586, GOC:go_curators] |
| GO:0043076 | 0.007672404 | 15 / 445 | "The nucleus of a megasporocyte, a diploid cell that undergoes meiosis to produce four megaspores, and its descendents." [GOC:jl, ISBN:0618254153] |
| GO:0045120 | 0.007672404 | 15 / 445 | "The nucleus of either the ovum or the spermatozoon following fertilization. Thus, in the fertilized ovum, there are two pronuclei, one originating from the ovum, the other from the spermatozoon that brought about fertilization; they approach each other, but do not fuse un |
| GO:0046818 | 0.007672404 | 15 / 445 | "A location in the host cell nucleus where viral proteins colocalize late in infection prior to the onset of viral DNA synthesis. More than one site can be present simultaneously." [PMID:10233976] |
| GO:0048353 | 0.007672404 | 15 / 445 | "Nucleus resulting from the fusion of the male gamete and two polar nuclei in the central cell of the embryo sac." [ISBN:0471245208] |
| GO:0048555 | 0.007672404 | 15 / 445 | "The nucleus of the generative cell, a cell contained within the pollen grain that will divide to produce two haploid sperm cells." [GOC:tair_curators] |
| GO:0048556 | 0.007672404 | 15 / 445 | "The nucleus of the microsporocyte. The microsporocyte is a diploid cell in which meiosis will occur, resulting in four microspores. A microspore is a spore that, in vascular plants, gives rise to a male gametophyte." [GOC:tair_curators, ISBN:047186840X] |
| GO:0097165 | 0.007672404 | 15 / 445 | "A dense aggregation in the nucleus composed of proteins and RNAs that appear when the cell is under stress." [GOC:ans, PMID:10359787, PMID:12865437] |
| GO:0097571 | 0.007672404 | 15 / 445 | "One of the two nuclei found in Giardia species (trophozoite stage). It is located on the left side of the cell when viewed from the dorsal side." [GOC:giardia, ISBN:0-444-81258-X] |
| GO:0097572 | 0.007672404 | 15 / 445 | "One of the two nuclei found in Giardia species (trophozoite stage). It is located on the right side of the cell when viewed from the dorsal side." [GOC:giardia, ISBN:0-444-81258-X] |
| GO:0110092 | 0.007672404 | 15 / 445 | "The area of a motile nucleus closest to the direction of movement." [GOC:al, GOC:mah, GOC:vw, PMID:15030757, PMID:24335254] |
| GO:0110093 | 0.007672404 | 15 / 445 | "The area of a motile nucleus furthest from the direction of movement." [GOC:kmv, PMID:24335254] |
| GO:0140510 | 0.007672404 | 15 / 445 | "A narrow constricted region of the nucleus that forms around the anaphase spindle during closed mitosis, and connects the main portions of the newly forming daughter nuclei." [PMID:32848252] |
| GO:1905754 | 0.007672404 | 15 / 445 | "Any nucleus that is part of a ascospore-type prospore." [GO_REF:0000064, GOC:TermGenie, PMID:26942678] |
| GO:0000101 | 0.008884668 | 2 / 3 | "The directed movement of amino acids containing sulfur (cystine, methionine and their derivatives) into, out of or within a cell, or between cells, by means of some agent such as a transporter or pore." [GOC:ai] |
| GO:0006869 | 0.008884668 | 2 / 3 | "The directed movement of lipids into, out of or within a cell, or between cells, by means of some agent such as a transporter or pore. Lipids are compounds soluble in an organic solvent but not, or sparingly, in an aqueous solvent." [ISBN:0198506732] |
| GO:0005871 | 0.010418152 | 3 / 9 | "Any complex that includes a dimer of molecules from the kinesin superfamily, a group of related proteins that contain an extended region of predicted alpha-helical coiled coil in the main chain that likely produces dimerization. The native complexes of several kinesin family |
| GO:0005635 | 0.011334636 | 16 / 450 | "The double lipid bilayer enclosing the nucleus and separating its contents from the rest of the cytoplasm; includes the intermembrane space, a gap of width 20-40 nm (also called the perinuclear space)." [ISBN:0198547684] |
| GO:0005634 | 0.011605291 | 16 / 449 | "A membrane-bounded organelle of eukaryotic cells in which chromosomes are housed and replicated. In most cells, the nucleus contains all of the cell's chromosomes except the organellar chromosomes, and is the site of RNA synthesis and processing. In some species, or in |

#### Supplementary Data S3.4.7

| GO_Term | P_val | Counts | MolFunc_Description |
|---|---|---|---|
| GO:0016491 | 5.7711E-10 | 72 / 658 | "Catalysis of an oxidation-reduction (redox) reaction, a reversible chemical reaction in which the oxidation state of an atom or atoms within a molecule is altered. One substrate acts as a hydrogen or electron donor and becomes oxidized, while the other acts as hydrogen or electron acceptor and becomes |
| GO:0005215 | 8.56075E-09 | 172 / 372 | "Enables the directed movement of substances (such as macromolecules, small molecules, ions) into, out of or within a cell, or between cells." [GOC:ai, GOC:dgf] |
| GO:0005198 | 9.38338E-08 | 176 / 374 | "The action of a molecule that contributes to the structural integrity of a complex." [GOC:mah, GOC:vw] |
| GO:0140110 | 1.82475E-07 | 172 / 367 | "A molecular function that controls the rate, timing and/or magnitude of gene transcription. The function of transcriptional regulators is to modulate gene expression at the transcription step so that they are expressed in the right cell at the right time and in the right amount throughout the life of the cell ar |
| GO:0060090 | 8.41781E-07 | 171 / 363 | "The binding activity of a molecule that brings together two or more molecules through a selective, non-covalent, often stoichiometric interaction, permitting those molecules to function in a coordinated way." [GOC:mtg_MIT_16mar07, GOC:vw] |
| GO:0140657 | 1.25092E-06 | 180 / 375 | "A molecular function characterized by the coupling of ATP hydrolysis to other steps of a reaction mechanism to make the reaction energetically favorable, for example to catalyze a reaction or drive transport against a concentration gradient." [PMID:24878343, PMID:25750732, PMID:32933017, PMID: |
| GO:0140223 | 1.26037E-06 | 171 / 362 | "A molecular function required for core promoter activity that mediates the assembly of the RNA polymerase holoenzyme at promoter DNA to form the pre-initiation complex (PIC). General transcription factors (GTFs) bind to and open promoter DNA, initiate RNA synthesis and stimulate the escape of the |
| GO:0016209 | 1.32465E-06 | 171 / 362 | "Inhibition of the reactions brought about by dioxygen (O2) or peroxides. Usually the antioxidant is effective because it can itself be more easily oxidized than the substance protected. The term is often applied to components that can trap free radicals, thereby breaking the chain reaction that normally lea |
| GO:0005488 | 1.37834E-06 | 172 / 363 | "The selective, non-covalent, often stoichiometric, interaction of a molecule with one or more specific sites on another molecule." [GOC:ceb, GOC:mah, ISBN:0198506732] |
| GO:0098772 | 1.46251E-06 | 171 / 361 | "A molecular function regulator regulates the activity of its target via non-covalent binding that does not result in covalent modification to the target. Examples of molecular function regulators include regulatory subunits of multimeric enzymes and channels. Mechanisms of regulation include allosteric cha |
| GO:0045735 | 1.53638E-06 | 171 / 361 | "Functions in the storage of nutritious substrates." [GOC:ai] |
| GO:0038024 | 1.69469E-06 | 171 / 361 | "Binding specifically to a substance (cargo) to deliver it to a transport vesicle. Cargo receptors span a membrane (either the plasma membrane or a vesicle membrane), binding simultaneously to cargo molecules and coat adaptors, to efficiently recruit soluble proteins to nascent vesicles." [PMID:1523995 |
| GO:0044183 | 1.77945E-06 | 171 / 361 | "Binding to a protein or a protein-containing complex to assist the protein folding process." [GOC:mtg_cambridge_2009] |
| GO:0045182 | 1.77945E-06 | 171 / 361 | "Any molecular function involved in the initiation, activation, perpetuation, repression or termination of polypeptide synthesis at the ribosome." [GOC:ai] |
| GO:0140104 | 1.86816E-06 | 171 / 361 | "Directly binding to a specific ion or molecule and delivering it either to an acceptor molecule or to a specific location." [GOC:molecular_function_refactoring, GOC:pdt] |
| GO:0060089 | 1.96099E-06 | 171 / 361 | "A compound molecular function in which an effector function is controlled by one or more regulatory components." [GOC:dos, GOC:pdt] |
| GO:0090729 | 1.96099E-06 | 171 / 361 | "Interacting selectively with one or more biological molecules in another (target) organism, initiating pathogenesis (leading to an abnormal, generally detrimental state) in the target organism. The activity should refer to an evolved function of the active gene product, i.e. one that was selected for. Example |
| GO:0140299 | 1.96099E-06 | 171 / 361 | "Binding to a small molecule and eliciting a change in the protein's activity in response to the intracellular level of that small molecule." [PMID:26328879] |
| GO:0140313 | 1.96099E-06 | 171 / 361 | "Binding to a specific molecule to prevent it from interacting with other partners or to inhibit its localization to the area of the cell or complex where it is active." [PMID:13130076] |
| GO:0140489 | 1.96099E-06 | 171 / 361 | "The action of a molecule that provides a shape or a sequence mimicking or complementary to the final product, providing template for copying the original molecule's shape or sequence." [GOC:pg] |
| GO:0140522 | 1.96099E-06 | 171 / 361 | "The activity of joining two lipid bilayers to form a single membrane." [PMID:10332732, PMID:11493675, PMID:12600315, PMID:32100701, PMID:32641474] |
| GO:0140691 | 1.96099E-06 | 171 / 361 | "Binding to an RNA or an RNA-containing complex to assist the folding process." [PMID:31165735] |
| GO:0140776 | 1.96099E-06 | 171 / 361 | "A molecular function that involves direct binding to one of the subunits of a protein-containing complex and promoting the dissociation of one or many subunits. This often happens by changing the conformation of the protein being bound, which decreases its affinity for the rest of the complex." [PMID: |
| GO:0140777 | 1.96099E-06 | 171 / 361 | "A molecular function that involves direct binding to one of the subunits of a protein-containing complex, thus preventing an interaction with a factor that would promote dissociation of the complex." [PMID:34040253] |
| GO:0140911 | 1.96099E-06 | 171 / 361 | "An activity in which a protein is inserted into the membrane of another cell where it forms transmembrane pores. Pores disrupts the integrity of the cell membrane, resulting in deregulated ion homeostasis, cellular dysfunction, and can result in cell death." [PMID:1406491, PMID:25157079, PMID:3438 |
| GO:0140912 | 1.96099E-06 | 171 / 361 | "Binding to a membrane and increasing its permeability. This may lead to cell membrane lysis and cell content release." [PMID:34496967] |
| GO:0141047 | 1.96099E-06 | 171 / 361 | "A molecular function exhibited by a protein that is covalently attached (AKA tagged or conjugated) to another molecule (for example a protein or a lipid) where it acts as a marker, recognized by the cellular apparatus to target the tagged protein for some cellular process such as modification, sequestratio |
| GO:0180020 | 1.96099E-06 | 171 / 361 | "The activity of bending or deforming a membrane. This activity can occur by multiple mechanisms including the insertion amphipathic domains into one or both leaflets." [PMID:19780639] |
| GO:0180024 | 1.96099E-06 | 171 / 361 | "An activity in which a protein, or group of proteins, acts to restrict the size of a fenestration in a membrane or membrane system (i.e. restrict the diameter of a hole in a membrane). An example of this is the molecular function is performed by ESCRT-III proteins at the nuclear envelope of fission yeast dur |
| GO:0140096 | 2.36406E-06 | 37 / 311 | "Catalytic activity that acts to modify a protein." [GOC:molecular_function_refactoring, GOC:pdt] |
| GO:0009055 | 2.56818E-06 | 174 / 365 | "A molecular function representing the directed movement of electrons from one molecular entity to another, typically mediated by electron carriers or acceptors, resulting in the transfer of energy and/or the reduction-oxidation (redox) transformation of chemical species. This activity is fundamental to v |
| GO:0003774 | 5.03186E-06 | 173 / 362 | "Generation of force resulting in movement, for example along a microfilament or microtubule, or in torque resulting in membrane scission or rotation of a flagellum. The energy required is obtained either from the hydrolysis of a nucleoside triphosphate or by an electrochemical proton gradient (proton-r |
| GO:0016740 | 7.80473E-06 | 34 / 289 | "Catalysis of the transfer of a group, e.g. a methyl group, glycosyl group, acyl group, phosphorus-containing, or other groups, from one compound (generally regarded as the donor) to another compound (generally regarded as the acceptor). Transferase is the systematic name for any enzyme of EC class |
| GO:0016829 | 8.83888E-06 | 33 / 278 | "Catalysis of the cleavage of C-C, C-O, C-N and other bonds by other means than by hydrolysis or oxidation, or conversely adding a group to a double bond. They differ from other enzymes in that two substrates are involved in one reaction direction, but only one in the other direction. When acting on the |
| GO:0016614 | 9.51726E-06 | 43 / 411 | "Catalysis of an oxidation-reduction (redox) reaction in which a CH-OH group act as a hydrogen or electron donor and reduces a hydrogen or electron acceptor." [GOC:ai] |
| GO:0004133 | 2.18845E-05 | 31 / 265 | "Catalysis of the cleavage of branch points in branched glycogen polymers." [ISBN:0198506732] |
| GO:0016218 | 2.18845E-05 | 31 / 265 | "Catalysis of a multistep reaction that produce polyketides through decarboxylative condensation of carboxylic acids. The key chain-building reaction, a C-N bond-forming reaction, involves the generation of the characteristic peptide bond by nucleophilic attack of the amino group of an amino-acyl donor |
| GO:0032451 | 2.18845E-05 | 31 / 265 | "Catalysis of the removal of a methyl group from a substrate." [GOC:mah] |
| GO:1904091 | 2.18845E-05 | 31 / 265 | "Catalysis of a multistep reaction that produce non-ribosomal peptides. The key chain-building reaction, a C-N bond-forming reaction, involves the generation of the characteristic peptide bond by nucleophilic attack of the amino group of an amino-acyl donor unit covalently bound to a downstream peptid |
| GO:0009975 | 2.35124E-05 | 31 / 266 | "Catalysis of a ring closure reaction." [ISBN:0198547684] |
| GO:0061783 | 2.52487E-05 | 31 / 267 | "A catalytic activity that contributes to the degradation of peptidoglycan." [GOC:dph, GOC:jh, PMID:22748813] |
| GO:0016853 | 2.70995E-05 | 31 / 268 | "Catalysis of the geometric or structural changes within one molecule. Isomerase is the systematic name for any enzyme of EC class 5." [ISBN:0198506732] |
| GO:0016874 | 3.83137E-05 | 31 / 273 | "Catalysis of the joining of two molecules, or two groups within a single molecule, using the energy from the hydrolysis of ATP, a similar triphosphate, or a pH gradient." [GOC:mah] |
| GO:0003943 | 4.1842E-05 | 4 / 5 | "Catalysis of the hydrolysis of the 4-sulfate groups of the N-acetyl-D-galactosamine 4-sulfate units of chondroitin sulfate and dermatan sulfate." [EC:3.1.6.12] |
| GO:0004065 | 4.1842E-05 | 4 / 5 | "Catalysis of the reaction: a phenol sulfate + H2O = a phenol + sulfate." [EC:3.1.6.1] |
| GO:0004098 | 4.1842E-05 | 4 / 5 | "Catalysis of the reaction: a cerebroside 3-sulfate + H2O = a cerebroside + sulfate." [EC:3.1.6.8] |
| GO:0004423 | 4.1842E-05 | 4 / 5 | "Catalysis of the hydrolysis of the 2-sulfate groups of the L-iduronate 2-sulfate units of dermatan sulfate, heparan sulfate and heparin." [EC:3.1.6.13] |
| GO:0004773 | 4.1842E-05 | 4 / 5 | "Catalysis of the reaction: 3-beta-hydroxyandrost-5-en-17-one 3-sulfate + H2O = 3-beta-hydroxyandrost-5-en-17-one + sulfate." [EC:3.1.6.2] |
| GO:0008449 | 4.1842E-05 | 4 / 5 | "Catalysis of the hydrolysis of the 6-sulfate group of the N-acetyl-D-glucosamine 6-sulfate units of heparan sulfate and keratan sulfate." [EC:3.1.6.14] |
| GO:0015024 | 4.1842E-05 | 4 / 5 | "Catalysis of the hydrolysis of the 2-sulfate groups of the 2-O-sulfo-D-glucuronate residues of chondroitin sulfate, heparin and heparitin sulfate." [EC:3.1.6.18] |

| GO_Term | P_val | Counts | BioProc_Description |
|---|---|---|---|
| GO:0055114 | 6.38959E-07 | 48 / 453 | "OBSOLETE. A metabolic process that results in the removal or addition of one or more electrons to or from a substance, with or without the concomitant removal or addition of a proton or protons." [GOC:dhl, GOC:ecd, GOC:jh2, GOC:jid, GOC:mlg, GOC:rph] |
| GO:0006865 | 6.46199E-05 | 6 / 15 | "The directed movement of amino acids, organic acids containing one or more amino substituents, into, out of or within a cell, or between cells, by means of some agent such as a transporter or pore." [GOC:ai] |
| GO:0019571 | 0.000147198 | 3 / 3 | "The chemical reactions and pathways resulting in the breakdown of D-arabinose, the D-enantiomer of arabino-pentose." [GOC:jsg, GOC:mah] |
| GO:0089718 | 0.000497266 | 5 / 14 | "The directed movement of an amino acid from outside of a cell, across the plasma membrane and into the cytosol." [GOC:krc, PMID:8195186] |
| GO:1990822 | 0.000497266 | 5 / 14 | "The directed movement of basic amino acids from one side of a membrane to the other." [GOC:dph, GOC:vw] |
| GO:0019568 | 0.000557979 | 3 / 4 | "The chemical reactions and pathways resulting in the breakdown of arabinose, arabino-pentose." [GOC:ai] |
| GO:1903713 | 0.000707127 | 5 / 15 | "The directed movement of asparagine across a membrane." [GO_REF:0000069, GOC:TermGenie, PMID:18503766] |
| GO:0032975 | 0.000975071 | 5 / 16 | "The directed movement of amino acids into the vacuole across the vacuolar membrane." [GOC:mah] |
| GO:0042316 | 0.001239961 | 9 / 53 | "The chemical reactions and pathways involving any antibiotic that contains the condensed beta-lactamthiazolidine ring system. Penicillins are produced naturally during the growth of various microfungi of the genera Penicillium and Aspergillus." [GOC:jl, ISBN:0198506732] |
| GO:0140115 | 0.001914686 | 34 / 406 | "The directed movement of some substance from inside of a cell, across the plasma membrane and into the extracellular region." [GOC:pg] |
| GO:0040007 | 0.00217134 | 178 / 359 | "The increase in size or mass of an entire organism, a part of an organism or a cell." [GOC:bf, GOC:ma] |
| GO:0099638 | 0.002259082 | 7 / 37 | "The directed movement of proteins from the endosome to the plasma membrane in transport vesicles." [GOC:dos] |
| GO:0032973 | 0.002484393 | 4 / 12 | "The directed movement of amino acids from inside of a cell, across the plasma membrane and into the extracellular region." [GOC:jl] |
| GO:0034487 | 0.002484393 | 4 / 12 | "The process in which an amino acid is transported from one side of the vacuolar membrane to the other." [GOC:mah] |
| GO:0035524 | 0.002484393 | 4 / 12 | "The directed movement of proline, pyrrolidine-2-carboxylic acid, across a membrane by means of some agent such as a transporter or pore." [GOC:vw] |
| GO:0140300 | 0.002484393 | 4 / 12 | "The process in which serine is transported from the cytosol into the mitochondrial matrix." [PMID:30442778] |
| GO:1903712 | 0.002484393 | 4 / 12 | "The directed movement of cysteine across a membrane." [GO_REF:0000069, GOC:TermGenie, PMID:17435223] |
| GO:0015802 | 0.002505428 | 3 / 6 | "The directed movement of basic amino acids, amino acids with a pH above 7, into, out of or within a cell, or between cells, by means of some agent such as a transporter or pore." [GOC:ai] |
| GO:0018868 | 0.002505428 | 3 / 6 | "The chemical reactions and pathways involving 2-aminobenzenesulfonate, aniline-o-sulfonic acid, an aromatic sulfonate used in organic synthesis and in the manufacture of various dyes and medicines." [UM-BBD_pathwayID:abs] |
| GO:0019757 | 0.002524052 | 8 / 48 | "The chemical reactions and pathways involving glycosinolates, substituted thioglycosides found in rapeseed products and related cruciferae." [GOC:mah, http://www.gardeneaters.net/family_characteristics.html] |
| GO:0006412 | 0.002694426 | 1 / 145 | "The cellular metabolic process in which a protein is formed, using the sequence of a mature mRNA or circRNA molecule to specify the sequence of amino acids in a polypeptide chain. Translation is mediated by the ribosome, and begins with the formation of a ternary complex between aminoacylated initi |
| GO:0002181 | 0.002710994 | 0 / 107 | "The chemical reactions and pathways resulting in the formation of a protein in the cytoplasm. This is a ribosome-mediated process in which the information in messenger RNA (mRNA) is used to specify the sequence of amino acids in the protein." [GOC:hjd] |
| GO:0019572 | 0.002800218 | 2 / 2 | "The chemical reactions and pathways resulting in the breakdown of L-arabinose, the L-enantiomer of arabino-pentose." [GOC:jsg, GOC:mah] |
| GO:0042843 | 0.002800218 | 2 / 2 | "The chemical reactions and pathways resulting in the breakdown of D-xylose, a naturally occurring plant polysaccharide." [ISBN:0198506732] |
| GO:0044577 | 0.002800218 | 2 / 2 | "The anaerobic chemical reactions and pathways resulting in the breakdown of xylose, an aldopentose, where one of the resulting products is ethanol." [GOC:mengo_curators, GOC:tt] |
| GO:0070813 | 0.002800218 | 2 / 2 | "The chemical reactions and pathways involving hydrogen sulfide, H2S." [GOC:mah] |
| GO:0045117 | 0.002803807 | 33 / 401 | "The directed movement of azoles, heterocyclic compounds found in many biologically important substances, across a lipid bilayer, across a membrane." [GOC:go_curators, ISBN:3527307206, Wikipedia:Azole] |
| GO:0006418 | 0.002867132 | 0 / 106 | "The synthesis of aminoacyl tRNA by the formation of an ester bond between the 3'-hydroxyl group of the most 3' adenosine of the tRNA and the alpha carboxylic acid group of an amino acid, to be used in ribosome-mediated polypeptide synthesis." [GOC:ma] |
| GO:0006855 | 0.002902369 | 33 / 402 | "The process in which a xenobiotic, a compound foreign to the organim exposed to it, is transported across a membrane. It may be synthesized by another organism (like ampicilin) or it can be a synthetic chemical." [GOC:ai, GOC:bf, GOC:krc] |
| GO:0046618 | 0.003003687 | 33 / 403 | "The directed movement of a xenobiotic from a cell, into the extracellular region. A xenobiotic is a compound foreign to the organim exposed to it. It may be synthesized by another organism (like ampicilin) or it can be a synthetic chemical." [GOC:go_curators, GOC:krc] |
| GO:1902047 | 0.003003687 | 33 / 403 | "The process in which a polyamine macromolecule is transported across a membrane." [GOC:TermGenie, GOC:vw] |
| GO:0090487 | 0.003229549 | 8 / 50 | "The chemical reactions and pathways resulting in the breakdown of secondary metabolites, the compounds that are not necessarily required for growth and maintenance of cells, and are often unique to a taxon." [GOC:tb] |
| GO:0019748 | 0.003249007 | 30 / 358 | "The chemical reactions and pathways resulting in many of the chemical changes of compounds that are not necessarily required for growth and maintenance of cells, and are often unique to a taxon. In multicellular organisms secondary metabolism is generally carried out in specific cell types, and may be |
| GO:0015810 | 0.003401238 | 4 / 13 | "The process in which aspartate is transported across a lipid bilayer, from one side of a membrane to the other." [GOC:go_curators, ISBN:0198506732] |
| GO:0090455 | 0.003401238 | 4 / 13 | "The directed movement of ornithine into the vacuole across the vacuolar membrane." [GOC:tb] |
| GO:0055085 | 0.003441295 | 34 / 423 | "The process in which a solute is transported across a lipid bilayer, from one side of a membrane to the other." [GOC:dph, GOC:jid] |
| GO:0006357 | 0.003619514 | 8 / 332 | "Any process that modulates the frequency, rate or extent of transcription mediated by RNA polymerase II." [GOC:go_curators, GOC:txnOH] |
| GO:0006355 | 0.003627306 | 7 / 307 | "Any process that modulates the frequency, rate or extent of cellular DNA-templated transcription." [GOC:go_curators, GOC:txnOH] |
| GO:0010467 | 0.003687755 | 5 / 255 | "The process in which a gene's sequence is converted into a mature gene product (protein or RNA). This includes the production of an RNA transcript and its processing, as well as translation and maturation for protein-coding genes." [GOC:txnOH-2018, PMID:25934543, PMID:31580950] |
| GO:0009237 | 0.004070816 | 8 / 52 | "The chemical reactions and pathways involving siderophores, low molecular weight Fe(III)-chelating substances made by aerobic or facultatively anaerobic bacteria, especially when growing under iron deficient conditions. The complexes of Fe(3+)-siderophores have very high stability constants and are |
| GO:0034982 | 0.004154876 | 3 / 7 | "The peptide cleavage of mitochondrial proteins, including cleavage contributing to their import." [GOC:curators] |
| GO:0098659 | 0.004154876 | 3 / 7 | "The directed movement of inorganic cations from outside of a cell, across the plasma membrane and into the cytosol." [GOC:dos] |
| GO:0045893 | 0.004296143 | 7 / 302 | "Any process that activates or increases the frequency, rate or extent of cellular DNA-templated transcription." [GOC:go_curators, GOC:txnOH] |
| GO:1903714 | 0.0045131 | 4 / 14 | "The directed movement of isoleucine across a membrane." [GO_REF:0000069, GOC:TermGenie, PMID:18503766] |
| GO:0006810 | 0.004572692 | 34 / 432 | "The directed movement of substances (such as macromolecules, small molecules, ions) or cellular components (such as complexes and organelles) into, out of or within a cell, or between cells, or within a multicellular organism by means of some agent such as a transporter or a transporter complex, a po |
| GO:0001407 | 0.004607179 | 32 / 400 | "The process in which a glycerophosphodiester is transported across a membrane. Glycerophosphodiesters are small molecules composed of glycerol-3-phosphate and an alcohol, for example, glycerophosphoinositol." [GOC:mcc, PMID:12912892] |
| GO:0015794 | 0.004607179 | 32 / 400 | "The process in which glycerol-3-phosphate is transported across a membrane. Glycerol-3-phosphate is a phosphoric monoester of glycerol." [GOC:ai] |
| GO:0015897 | 0.004607179 | 32 / 400 | "The process in which an organomercurial compound is transported across a membrane. Organomercurial substances are any organic compound containing a mercury atom." [GOC:ai, PMID:18793329] |
| GO:0015904 | 0.004607179 | 32 / 400 | "The directed movement of tetracycline from one side of a membrane to the other. Tetracycline is a broad spectrum antibiotic that blocks binding of aminoacyl tRNA to the ribosomes of both Gram-positive and Gram-negative organisms (and those of organelles)." [GOC:curators] |
| GO:0015905 | 0.004607179 | 32 / 400 | "The directed movement of bicyclomycin across a lipid bilayer, from one side of a membrane to the other. Bicyclomycin (or bicozamycin) is an antibacterial drug often used as a livestock feed additive." [PMID:20067529] |

| GO_Term | P_val | Counts | CellComp_Description |
|---|---|---|---|
| GO:0008150 | 0.001159003 | 15 / 496 | "A biological process is the execution of a genetically-encoded biological module or program. It consists of all the steps required to achieve the specific biological objective of the module. A biological process is accomplished by a particular set of molecular functions carried out by specific gene products (or |
| GO:0022414 | 0.001410262 | 14 / 468 | "A biological process that directly contributes to the process of producing new individuals by one or two organisms. The new individuals inherit some proportion of their genetic material from the parent or parents." [GOC:dph, GOC:isa_complete] |
| GO:0098754 | 0.001410262 | 14 / 468 | "Any process that reduces or removes the toxicity of a toxic substance. These may include transport of the toxic substance away from sensitive areas and to compartments or complexes whose purpose is sequestration of the toxic substance." [GOC:dos] |
| GO:0050896 | 0.001503839 | 14 / 466 | "Any process that results in a change in state or activity of a cell or an organism (in terms of movement, secretion, enzyme production, gene expression, etc.) as a result of a stimulus. The process begins with detection of the stimulus and ends with a change in state or activity or the cell or organism." [GOC |
| GO:0140513 | 0.001521671 | 16 / 509 | "A stable assembly of two or more macromolecules, i.e. proteins, nucleic acids, carbohydrates or lipids, in which at least one component is a protein and the constituent parts function together in the nucleus." [GOC:pg] |
| GO:0000003 | 0.001552745 | 14 / 465 | "The production of new individuals that contain some portion of genetic material inherited from one or more parent organisms." [GOC:go_curators, GOC:isa_complete, GOC:jl, ISBN:0198506732] |
| GO:0002376 | 0.001552745 | 14 / 465 | "Any process involved in the development or functioning of the immune system, an organismal system for calibrated responses to potential internal or invasive threats." [GO_REF:0000022, GOC:add] |
| GO:0016032 | 0.001552745 | 14 / 465 | "A multi-organism process in which a virus is a participant. The other participant is the host. Includes infection of a host cell, replication of the viral genome, and assembly of progeny virus particles. In some cases the viral genetic material may integrate into the host genome and only subsequently, under p |
| GO:0032501 | 0.001552745 | 14 / 465 | "Any biological process, occurring at the level of a multicellular organism, pertinent to its function." [GOC:curators, GOC:dph, GOC:isa_complete, GOC:tb] |
| GO:0032502 | 0.001552745 | 14 / 465 | "A biological process whose specific outcome is the progression of an integrated living unit: an anatomical structure (which may be a subcellular structure, cell, tissue, or organ), or organism over time from an initial condition to a later condition." [GOC:isa_complete] |
| GO:0040011 | 0.001552745 | 14 / 465 | "Self-propelled movement of a cell or organism from one location to another." [GOC:dgh] |
| GO:0042592 | 0.001552745 | 14 / 465 | "Any biological process involved in the maintenance of an internal steady state." [GOC:jl, ISBN:0395825172] |
| GO:0043473 | 0.001552745 | 14 / 465 | "The accumulation of pigment in an organism, tissue or cell, either by increased deposition or by increased number of cells." [GOC:jl] |
| GO:0044419 | 0.001552745 | 14 / 465 | "Any process evolved to enable an interaction with an organism of a different species." [GOC:cc] |
| GO:0044848 | 0.001552745 | 14 / 465 | "A distinct period or stage in a biological process or cycle." [GOC:jl] |
| GO:0048511 | 0.001552745 | 14 / 465 | "Any process pertinent to the generation and maintenance of rhythms in the physiology of an organism." [GOC:jid] |
| GO:0051179 | 0.001552745 | 14 / 465 | "Any process in which a cell, a substance, or a cellular entity, such as a protein complex or organelle, is transported, tethered to or otherwise maintained in a specific location. In the case of substances, localization may also be achieved via selective degradation." [GOC:ai, GOC:dos] |
| GO:0051703 | 0.001552745 | 14 / 465 | "Any process in which an organism has an effect on an organism of the same species." [GOC:ai] |
| GO:0065007 | 0.001552745 | 14 / 465 | "Any process that modulates a measurable attribute of any biological process, quality or function." [GOC:dph, GOC:isa_complete, GOC:mah, GOC:pr, GOC:vw] |
| GO:0009987 | 0.001569121 | 16 / 508 | "Any process that is carried out at the cellular level, but not necessarily restricted to a single cell. For example, cell communication occurs among more than one cell, but occurs at the cellular level." [GOC:go_curators, GOC:isa_complete] |
| GO:0008152 | 0.00171973 | 16 / 505 | "The chemical reactions and pathways, including anabolism and catabolism, by which living organisms transform chemical substances. Metabolic processes typically transform small molecules, but also include macromolecular processes such as DNA repair and replication, and protein synthesis and degra |
| GO:0031981 | 0.001941127 | 16 / 501 | "The volume enclosed by the nuclear inner membrane." [GOC:mah, GOC:pz] |
| GO:0040007 | 0.002036663 | 15 / 478 | "The increase in size or mass of an entire organism, a part of an organism or a cell." [GOC:bf, GOC:ma] |
| GO:0015804 | 0.003335073 | 2 / 2 | "The directed movement of neutral amino acids, amino acids with no net charge, into, out of or within a cell, or between cells, by means of some agent such as a transporter or pore." [GOC:ai] |
| GO:0015807 | 0.003335073 | 2 / 2 | "The directed movement of L-enantiomer amino acids into, out of or within a cell, or between cells, by means of some agent such as a transporter or pore." [GOC:ai, GOC:jsg, GOC:mah] |
| GO:0005886 | 0.003796638 | 20 / 192 | "The membrane surrounding a cell that separates the cell from its external environment. It consists of a phospholipid bilayer and associated proteins." [ISBN:0716731363] |
| GO:0098590 | 0.00399816 | 20 / 193 | "A membrane that is a (regional) part of the plasma membrane." [GOC:dos] |
| GO:0043231 | 0.006497305 | 22 / 579 | "Organized structure of distinctive morphology and function, bounded by a single or double lipid bilayer membrane and occurring within the cell. Includes the nucleus, mitochondria, plastids, vacuoles, and vesicles. Excludes the plasma membrane." [GOC:go_curators] |
| GO:0016020 | 0.006793701 | 20 / 204 | "A lipid bilayer along with all the proteins and protein complexes embedded in it and attached to it." [GOC:dos, GOC:mah, ISBN:0815316194] |
| GO:0000943 | 0.009176706 | 16 / 445 | "A complex of the retrotransposon RNA genome, reverse transcriptase, integrase, and associated molecules required for reproduction and integration of the retrotransposon into the host genome; the main structural molecule of the nucleocapsid is often a gag protein homolog." [GOC:clt, PMID:1086190 |
| GO:0031039 | 0.009176706 | 16 / 445 | "A membrane-bounded organelle of ciliated protozoan cells that contains polyploid copies of a portion of the cell's complete genome. Transcription of genes occurs in macronuclei. Some ciliate species may contain multiple macronuclei per cell." [GOC:ns] |
| GO:0031040 | 0.009176706 | 16 / 445 | "A membrane-bounded organelle of ciliated protozoan cells that contains a diploid copy of the cell's complete genome. Sections of contiguous sequence in the macronucleus are often interrupted by internal eliminated sequences (IES), and may be permuted, in micronuclei. Genic transcription is not found |
| GO:0042405 | 0.009176706 | 16 / 445 | "An intranuclear focus at which aggregated proteins have been sequestered." [GOC:jl] |
| GO:0043073 | 0.009176706 | 16 / 445 | "The nucleus of a germ cell, a reproductive cell in multicellular organisms." [CL:0000586, GOC:go_curators] |
| GO:0043076 | 0.009176706 | 16 / 445 | "The nucleus of a megasporocyte, a diploid cell that undergoes meiosis to produce four megaspores, and its descendents." [GOC:jl, ISBN:0618254153] |
| GO:0045120 | 0.009176706 | 16 / 445 | "The nucleus of either the ovum or the spermatozoon following fertilization. Thus, in the fertilized ovum, there are two pronuclei, one originating from the ovum, the other from the spermatozoon that brought about fertilization; they approach each other, but do not fuse until just before the first cleavage |
| GO:0046818 | 0.009176706 | 16 / 445 | "A location in the host cell nucleus where viral proteins colocalize late in infection prior to the onset of viral DNA synthesis. More than one site can be present simultaneously." [PMID:10233976] |
| GO:0048353 | 0.009176706 | 16 / 445 | "Nucleus resulting from the fusion of the male gamete and two polar nuclei in the central cell of the embryo sac." [ISBN:0471245208] |
| GO:0048555 | 0.009176706 | 16 / 445 | "The nucleus of the generative cell, a cell contained within the pollen grain that will divide to produce two haploid sperm cells." [GOC:tair_curators] |
| GO:0048556 | 0.009176706 | 16 / 445 | "The nucleus of the microsporocyte. The microsporocyte is a diploid cell in which meiosis will occur, resulting in four microspores. A microspore is a spore that, in vascular plants, gives rise to a male gametophyte." [GOC:tair_curators, ISBN:047186840X] |
| GO:0097165 | 0.009176706 | 16 / 445 | "A dense aggregation in the nucleus composed of proteins and RNAs that appear when the cell is under stress." [GOC:ans, PMID:10359787, PMID:12865437] |
| GO:0097571 | 0.009176706 | 16 / 445 | "One of the two nuclei found in Giardia species (trophozoite stage). It is located on the left side of the cell when viewed from the dorsal side." [GOC:giardia, ISBN:0-444-81258-X] |
| GO:0097572 | 0.009176706 | 16 / 445 | "One of the two nuclei found in Giardia species (trophozoite stage). It is located on the right side of the cell when viewed from the dorsal side." [GOC:giardia, ISBN:0-444-81258-X] |
| GO:0110092 | 0.009176706 | 16 / 445 | "The area of a motile nucleus closest to the direction of movement." [GOC:al, GOC:mah, GOC:vw, PMID:15030757, PMID:24335254] |
| GO:0110093 | 0.009176706 | 16 / 445 | "The area of a motile nucleus furthest from the direction of movement." [GOC:kmv, PMID:24335254] |
| GO:0140510 | 0.009176706 | 16 / 445 | "A narrow constricted region of the nucleus that forms around the anaphase spindle during closed mitosis, and connects the main portions of the newly forming daughter nuclei." [PMID:32848252] |
| GO:1905754 | 0.009176706 | 16 / 445 | "Any nucleus that is part of a ascospore-type prospore." [GO_REF:0000064, GOC:TermGenie, PMID:26942678] |
| GO:0000101 | 0.009430161 | 2 / 3 | "The directed movement of amino acids containing sulfur (cystine, methionine and their derivatives) into, out of or within a cell, or between cells, by means of some agent such as a transporter or pore." [GOC:ai] |
| GO:0006869 | 0.009430161 | 2 / 3 | "The directed movement of lipids into, out of or within a cell, or between cells, by means of some agent such as a transporter or pore. Lipids are compounds soluble in an organic solvent but not, or sparingly, in an aqueous solvent." [ISBN:0198506732] |
| GO:0005761 | 0.011086345 | 0 / 75 | "A ribosome found in the mitochondrion of a eukaryotic cell; contains a characteristic set of proteins distinct from those of cytosolic ribosomes." [GOC:mah, ISBN:0198506732] |

#### Supplementary Data S3.4.8

| GO_Term | P_val | Counts | MolFunc_Description |
|---|---|---|---|
| GO:0005198 | 1.31514E-09 | 272 / 3741 | "The action of a molecule that contributes to the structural integrity of a complex." [GOC:mah, GOC:vw] |
| GO:0005215 | 2.79228E-08 | 274 / 3723 | "Enables the directed movement of substances (such as macromolecules, small molecules, ions) into, out of or within a cell, or between cells." [GOC:ai, GOC:dgf] |
| GO:0140110 | 3.18626E-08 | 269 / 3673 | "A molecular function that controls the rate, timing and/or magnitude of gene transcription. The function of transcriptional regulators is to modulate gene expression at the transcription step so that they are expressed in the right cell at the right time and in the right amou |
| GO:0016491 | 7.96505E-08 | 91 / 658 | "Catalysis of an oxidation-reduction (redox) reaction, a reversible chemical reaction in which the oxidation state of an atom or atoms within a molecule is altered. One substrate acts as a hydrogen or electron donor and becomes oxidized, while the other acts as hydrogen o |
| GO:0009055 | 1.43754E-07 | 269 / 3652 | "A molecular function representing the directed movement of electrons from one molecular entity to another, typically mediated by electron carriers or acceptors, resulting in the transfer of energy and/or the reduction-oxidation (redox) transformation of chemical species. |
| GO:0060090 | 1.66929E-07 | 267 / 3630 | "The binding activity of a molecule that brings together two or more molecules through a selective, non-covalent, often stoichiometric interaction, permitting those molecules to function in a coordinated way." [GOC:mtg_MIT_16mar07, GOC:vw] |
| GO:0005488 | 2.32709E-07 | 268 / 3635 | "The selective, non-covalent, often stoichiometric, interaction of a molecule with one or more specific sites on another molecule." [GOC:ceb, GOC:mah, ISBN:0198506732] |
| GO:0140223 | 2.84457E-07 | 267 / 3622 | "A molecular function required for core promoter activity that mediates the assembly of the RNA polymerase holoenzyme at promoter DNA to form the pre-initiation complex (PIC). General transcription factors (GTFs) bind to and open promoter DNA, initiate RNA synthesis |
| GO:0016209 | 3.03731E-07 | 267 / 3621 | "Inhibition of the reactions brought about by dioxygen (O2) or peroxides. Usually the antioxidant is effective because it can itself be more easily oxidized than the substance protected. The term is often applied to components that can trap free radicals, thereby breaking th |
| GO:0098772 | 3.46043E-07 | 267 / 3619 | "A molecular function regulator regulates the activity of its target via non-covalent binding that does not result in covalent modification to the target. Examples of molecular function regulators include regulatory subunits of multimeric enzymes and channels. Mechanisms c |
| GO:0045735 | 3.69231E-07 | 267 / 3618 | "Functions in the storage of nutritious substrates." [GOC:ai] |
| GO:0038024 | 4.20079E-07 | 267 / 3616 | "Binding specifically to a substance (cargo) to deliver it to a transport vesicle. Cargo receptors span a membrane (either the plasma membrane or a vesicle membrane), binding simultaneously to cargo molecules and coat adaptors, to efficiently recruit soluble proteins to na |
| GO:0044183 | 4.47915E-07 | 267 / 3615 | "Binding to a protein or a protein-containing complex to assist the protein folding process." [GOC:mtg_cambridge_2009] |
| GO:0045182 | 4.47915E-07 | 267 / 3615 | "Any molecular function involved in the initiation, activation, perpetuation, repression or termination of polypeptide synthesis at the ribosome." [GOC:ai] |
| GO:0140657 | 4.55159E-07 | 281 / 3753 | "A molecular function characterized by the coupling of ATP hydrolysis to other steps of a reaction mechanism to make the reaction energetically favorable, for example to catalyze a reaction or drive transport against a concentration gradient." [PMID:24878343, PMID:257507 |
| GO:0140104 | 4.77486E-07 | 267 / 3614 | "Directly binding to a specific ion or molecule and delivering it either to an acceptor molecule or to a specific location." [GOC:molecular_function_refactoring, GOC:pdt] |
| GO:0060089 | 5.08893E-07 | 267 / 3613 | "A compound molecular function in which an effector function is controlled by one or more regulatory components." [GOC:dos, GOC:pdt] |
| GO:0090729 | 5.08893E-07 | 267 / 3613 | "Interacting selectively with one or more biological molecules in another (target) organism, initiating pathogenesis (leading to an abnormal, generally detrimental state) in the target organism. The activity should refer to an evolved function of the active gene product, i.e. o |
| GO:0140299 | 5.08893E-07 | 267 / 3613 | "Binding to a small molecule and eliciting a change in the protein's activity in response to the intracellular level of that small molecule." [PMID:26328879] |
| GO:0140313 | 5.08893E-07 | 267 / 3613 | "Binding to a specific molecule to prevent it from interacting with other partners or to inhibit its localization to the area of the cell or complex where it is active." [PMID:13130076] |
| GO:0140489 | 5.08893E-07 | 267 / 3613 | "The action of a molecule that provides a shape or a sequence mimicking or complementary to the final product, providing template for copying the original molecule's shape or sequence." [GOC:pg] |
| GO:0140522 | 5.08893E-07 | 267 / 3613 | "The activity of joining two lipid bilayers to form a single membrane." [PMID:10332732, PMID:11493675, PMID:12600315, PMID:32100701, PMID:32641474] |
| GO:0140691 | 5.08893E-07 | 267 / 3613 | "Binding to an RNA or an RNA-containing complex to assist the folding process." [PMID:31165735] |
| GO:0140776 | 5.08893E-07 | 267 / 3613 | "A molecular function that involves direct binding to one of the subunits of a protein-containing complex and promoting the dissociation of one or many subunits. This often happens by changing the conformation of the protein being bound, which decreases its affinity for |
| GO:0140777 | 5.08893E-07 | 267 / 3613 | "A molecular function that involves direct binding to one of the subunits of a protein-containing complex, thus preventing an interaction with a factor that would promote dissociation of the complex." [PMID:34040253] |
| GO:0140911 | 5.08893E-07 | 267 / 3613 | "An activity in which a protein is inserted into the membrane of another cell where it forms transmembrane pores. Pores disrupts the integrity of the cell membrane, resulting in deregulated ion homeostasis, cellular dysfunction, and can result in cell death." [PMID:1406491 |
| GO:0140912 | 5.08893E-07 | 267 / 3613 | "Binding to a membrane and increasing its permeability. This may lead to cell membrane lysis and cell content release." [PMID:34496967] |
| GO:0141047 | 5.08893E-07 | 267 / 3613 | "A molecular function exhibited by a protein that is covalently attached (AKA tagged or conjugated) to another molecule (for example a protein or a lipid) where it acts as a marker, recognized by the cellular apparatus to target the tagged protein for some cellular process s |
| GO:0180020 | 5.08893E-07 | 267 / 3613 | "The activity of bending or deforming a membrane. This activity can occur by multiple mechanisms including the insertion amphipathic domains into one or both leaflets." [PMID:19780639] |
| GO:0180024 | 5.08893E-07 | 267 / 3613 | "An activity in which a protein, or group of proteins, acts to restrict the size of a fenestration in a membrane or membrane system (i.e. restrict the diameter of a hole in a membrane). An example of this is the molecular function is performed by ESCRT-III proteins at the nucl |
| GO:0003774 | 9.61698E-07 | 269 / 3623 | "Generation of force resulting in movement, for example along a microfilament or microtubule, or in torque resulting in membrane scission or rotation of a flagellum. The energy required is obtained either from the hydrolysis of a nucleoside triphosphate or by an electroch |
| GO:0016787 | 4.38213E-05 | 56 / 410 | "Catalysis of the hydrolysis of various bonds, e.g. C-O, C-N, C-C, phosphoric anhydride bonds, etc." [ISBN:0198506732] |
| GO:0047536 | 4.7759E-05 | 4 / 4 | "Catalysis of the reaction: 2-oxoglutarate + L-2-aminoadipate = 2-oxoadipate + L-glutamate." [EC:2.6.1.39, RHEA:12601] |
| GO:0043168 | 6.92067E-05 | 31 / 186 | "Binding to an anion, a charged atom or group of atoms with a net negative charge." [GOC:jl] |
| GO:0016705 | 8.38337E-05 | 53 / 391 | "Catalysis of an oxidation-reduction (redox) reaction in which hydrogen or electrons are transferred from each of two donors, and molecular oxygen is reduced or incorporated into a donor." [GOC:mah] |
| GO:0016614 | 8.67968E-05 | 55 / 411 | "Catalysis of an oxidation-reduction (redox) reaction in which a CH-OH group act as a hydrogen or electron donor and reduces a hydrogen or electron acceptor." [GOC:ai] |
| GO:0016740 | 9.98795E-05 | 42 / 289 | "Catalysis of the transfer of a group, e.g. a methyl group, glycosyl group, acyl group, phosphorus-containing, or other groups, from one compound (generally regarded as the donor) to another compound (generally regarded as the acceptor). Transferase is the systematic nar |
| GO:0016722 | 0.000168872 | 50 / 373 | "Catalysis of an oxidation-reduction in which the oxidation state of metal ion is altered." [GOC:mah] |
| GO:0016903 | 0.000171395 | 51 / 383 | "Catalysis of an oxidation-reduction (redox) reaction in which an aldehyde or ketone (oxo) group acts as a hydrogen or electron donor and reduces a hydrogen or electron acceptor." [GOC:ai] |
| GO:0016667 | 0.000202859 | 50 / 376 | "Catalysis of an oxidation-reduction (redox) reaction in which a sulfur-containing group acts as a hydrogen or electron donor and reduces a hydrogen or electron acceptor." [GOC:ai] |
| GO:0003676 | 0.000207931 | 25 / 531 | "Binding to a nucleic acid." [GOC:jl] |
| GO:0003943 | 0.00021907 | 4 / 5 | "Catalysis of the hydrolysis of the 4-sulfate groups of the N-acetyl-D-galactosamine 4-sulfate units of chondroitin sulfate and dermatan sulfate." [EC:3.1.6.12] |
| GO:0004065 | 0.00021907 | 4 / 5 | "Catalysis of the reaction: a phenol sulfate + H2O = a phenol + sulfate." [EC:3.1.6.1] |
| GO:0004098 | 0.00021907 | 4 / 5 | "Catalysis of the reaction: a cerebroside 3-sulfate + H2O = a cerebroside + sulfate." [EC:3.1.6.8] |
| GO:0004423 | 0.00021907 | 4 / 5 | "Catalysis of the hydrolysis of the 2-sulfate groups of the L-iduronate 2-sulfate units of dermatan sulfate, heparan sulfate and heparin." [EC:3.1.6.13] |
| GO:0004773 | 0.00021907 | 4 / 5 | "Catalysis of the reaction: 3-beta-hydroxyandrost-5-en-17-one 3-sulfate + H2O = 3-beta-hydroxyandrost-5-en-17-one + sulfate." [EC:3.1.6.2] |
| GO:0008449 | 0.00021907 | 4 / 5 | "Catalysis of the hydrolysis of the 6-sulfate group of the N-acetyl-D-glucosamine 6-sulfate units of heparan sulfate and keratan sulfate." [EC:3.1.6.14] |
| GO:0015024 | 0.00021907 | 4 / 5 | "Catalysis of the hydrolysis of the 2-sulfate groups of the 2-O-sulfo-D-glucuronate residues of chondroitin sulfate, heparin and heparitin sulfate." [EC:3.1.6.18] |
| GO:0018741 | 0.00021907 | 4 / 5 | "Catalysis of the reaction: a primary linear alkyl sulfate ester + H2O =>a primary alcohol + H(+) + sulfate." [RHEA:67908, UM-BBD_reactionID:r0602] |
| GO:0033886 | 0.00021907 | 4 / 5 | "Catalysis of the hydrolysis of the 2- and 3-sulfate groups of the polysulfates of cellulose and charonin." [EC:3.1.6.7] |

| GO_Term | P_val | Counts | BioProc_Description |
|---|---|---|---|
| GO:0040007 | 2.77477E-06 | 262 / 3597 | "The increase in size or mass of an entire organism, a part of an organism or a cell." [GOC:bf, GOC:ma] |
| GO:0006810 | 7.82761E-06 | 59 / 432 | "The directed movement of substances (such as macromolecules, small molecules, ions) or cellular components (such as complexes and organelles) into, out of or within a cell, or between cells, or within a multicellular organism by means of some agent such as a transporte |
| GO:0055085 | 8.32899E-06 | 58 / 423 | "The process in which a solute is transported across a lipid bilayer, from one side of a membrane to the other." [GOC:dph, GOC:jid] |
| GO:0022414 | 1.44678E-05 | 258 / 3532 | "A biological process that directly contributes to the process of producing new individuals by one or two organisms. The new individuals inherit some proportion of their genetic material from the parent or parents." [GOC:dph, GOC:isa_complete] |
| GO:0055114 | 1.65491E-05 | 60 / 453 | "OBSOLETE. A metabolic process that results in the removal or addition of one or more electrons to or from a substance, with or without the concomitant removal or addition of a proton or protons." [GOC:dhl, GOC:ecd, GOC:jh2, GOC:jid, GOC:mlg, GOC:rph] |
| GO:0034627 | 1.67015E-05 | 6 / 9 | "The chemical reactions and pathways resulting in the formation of nicotinamide adenine dinucleotide (NAD), beginning with the synthesis of tryptophan or aspartate from simpler precursors; biosynthesis may be of either the oxidized form, NAD, or the reduced form, NAD |
| GO:0140115 | 2.01461E-05 | 55 / 406 | "The directed movement of some substance from inside of a cell, across the plasma membrane and into the extracellular region." [GOC:pg] |
| GO:0044419 | 2.79124E-05 | 258 / 3521 | "Any process evolved to enable an interaction with an organism of a different species." [GOC:cc] |
| GO:0045117 | 2.81582E-05 | 54 / 401 | "The directed movement of azoles, heterocyclic compounds found in many biologically important substances, across a lipid bilayer, across a membrane." [GOC:go_curators, ISBN:3527307206, Wikipedia:Azole] |
| GO:0097080 | 2.81582E-05 | 54 / 401 | "The directed movement of inorganic selenite (HSeO3-1 at physiological pH) across a plasma membrane." [GOC:mcc, PMID:20861301] |
| GO:0000003 | 2.95783E-05 | 258 / 3520 | "The production of new individuals that contain some portion of genetic material inherited from one or more parent organisms." [GOC:go_curators, GOC:isa_complete, GOC:jl, ISBN:0198506732] |
| GO:0032502 | 2.95783E-05 | 258 / 3520 | "A biological process whose specific outcome is the progression of an integrated living unit: an anatomical structure (which may be a subcellular structure, cell, tissue, or organ), or organism over time from an initial condition to a later condition." [GOC:isa_complete] |
| GO:0051703 | 2.95783E-05 | 258 / 3520 | "Any process in which an organism has an effect on an organism of the same species." [GOC:ai] |
| GO:0006855 | 3.01145E-05 | 54 / 402 | "The process in which a xenobiotic, a compound foreign to the organim exposed to it, is transported across a membrane. It may be synthesized by another organism (like ampicilin) or it can be a synthetic chemical." [GOC:ai, GOC:bf, GOC:krc] |
| GO:0007015 | 3.0451E-05 | 8 / 18 | "A process that is carried out at the cellular level which results in the assembly, arrangement of constituent parts, or disassembly of cytoskeletal structures comprising actin filaments. Includes processes that control the spatial distribution of actin filaments, such as organizi |
| GO:0002376 | 3.13344E-05 | 258 / 3519 | "Any process involved in the development or functioning of the immune system, an organismal system for calibrated responses to potential internal or invasive threats." [GO_REF:0000022, GOC:add] |
| GO:0016032 | 3.13344E-05 | 258 / 3519 | "A multi-organism process in which a virus is a participant. The other participant is the host. Includes infection of a host cell, replication of the viral genome, and assembly of progeny virus particles. In some cases the viral genetic material may integrate into the host genom |
| GO:0032501 | 3.13344E-05 | 258 / 3519 | "Any biological process, occurring at the level of a multicellular organism, pertinent to its function." [GOC:curators, GOC:dph, GOC:isa_complete, GOC:tb] |
| GO:0040011 | 3.13344E-05 | 258 / 3519 | "Self-propelled movement of a cell or organism from one location to another." [GOC:dgh] |
| GO:0042592 | 3.13344E-05 | 258 / 3519 | "Any biological process involved in the maintenance of an internal steady state." [GOC:jl, ISBN:0395825172] |
| GO:0043473 | 3.13344E-05 | 258 / 3519 | "The accumulation of pigment in an organism, tissue or cell, either by increased deposition or by increased number of cells." [GOC:jl] |
| GO:0044848 | 3.13344E-05 | 258 / 3519 | "A distinct period or stage in a biological process or cycle." [GOC:jl] |
| GO:0051179 | 3.13344E-05 | 258 / 3519 | "Any process in which a cell, a substance, or a cellular entity, such as a protein complex or organelle, is transported, tethered to or otherwise maintained in a specific location. In the case of substances, localization may also be achieved via selective degradation." [GOC:ai, G |
| GO:0065007 | 3.13344E-05 | 258 / 3519 | "Any process that modulates a measurable attribute of any biological process, quality or function." [GOC:dph, GOC:isa_complete, GOC:mah, GOC:pr, GOC:vw] |
| GO:0034220 | 3.21936E-05 | 54 / 403 | "A process in which a monoatomic ion is transported across a membrane. Monatomic ions (also called simple ions) are ions consisting of exactly one atom." [GOC:mah] |
| GO:0046618 | 3.21936E-05 | 54 / 403 | "The directed movement of a xenobiotic from a cell, into the extracellular region. A xenobiotic is a compound foreign to the organim exposed to it. It may be synthesized by another organism (like ampicilin) or it can be a synthetic chemical." [GOC:go_curators, GOC:krc] |
| GO:0046967 | 3.21936E-05 | 54 / 403 | "The directed movement of substances from the cytosol to the endoplasmic reticulum of a cell." [GOC:ai] |
| GO:1902047 | 3.21936E-05 | 54 / 403 | "The process in which a polyamine macromolecule is transported across a membrane." [GOC:TermGenie, GOC:vw] |
| GO:0098754 | 3.59383E-05 | 259 / 3527 | "Any process that reduces or removes the toxicity of a toxic substance. These may include transport of the toxic substance away from sensitive areas and to compartments or complexes whose purpose is sequestration of the toxic substance." [GOC:dos] |
| GO:0030047 | 3.84608E-05 | 6 / 10 | "Covalent modification of an actin molecule." [GOC:mah] |
| GO:0031032 | 3.84608E-05 | 6 / 10 | "A process that is carried out at the cellular level which results in the assembly, arrangement of constituent parts, or disassembly of cytoskeletal structures containing both actin and myosin or paramyosin. The myosin may be organized into filaments." [GOC:dph, GOC:jl, GC |
| GO:0031033 | 3.84608E-05 | 6 / 10 | "A process that is carried out at the cellular level which results in the assembly, arrangement of constituent parts, or disassembly of a filament composed of myosin molecules." [GOC:mah] |
| GO:0061204 | 3.84608E-05 | 6 / 10 | "The formation or disassembly of a filament composed of paramyosin molecules." [GOC:dph, GOC:kmv] |
| GO:0098974 | 3.84608E-05 | 6 / 10 | "A process that is carried out at the cellular level which results in the assembly, arrangement of constituent parts, or disassembly of cytoskeletal structures comprising actin filaments and their associated proteins in the postsynaptic actin cytoskeleton." [GOC:dos] |
| GO:0099140 | 3.84608E-05 | 6 / 10 | "A process that is carried out at the cellular level which results in the assembly, arrangement of constituent parts, or disassembly of cytoskeletal structures comprising actin filaments and their associated proteins in the presynaptic actin cytoskeleton." [GOC:dos] |
| GO:1904600 | 3.84608E-05 | 6 / 10 | "The aggregation, arrangement and bonding together of a set of components to form an actin fusion focus." [GO_REF:0000079, GOC:TermGenie, PMID:25825517] |
| GO:0050896 | 3.8895E-05 | 260 / 3536 | "Any process that results in a change in state or activity of a cell or an organism (in terms of movement, secretion, enzyme production, gene expression, etc.) as a result of a stimulus. The process begins with detection of the stimulus and ends with a change in state or activ |
| GO:0048511 | 5.04513E-05 | 259 / 3521 | "Any process pertinent to the generation and maintenance of rhythms in the physiology of an organism." [GOC:jid] |
| GO:1990542 | 5.07988E-05 | 54 / 410 | "The process in which a solute is transported from one side of a membrane to the other into, out of or within a mitochondrion." [PMID:20533899] |
| GO:0001407 | 5.09394E-05 | 53 / 400 | "The process in which a glycerophosphodiester is transported across a membrane. Glycerophosphodiesters are small molecules composed of glycerol-3-phosphate and an alcohol, for example, glycerophosphoinositol." [GOC:mcc, PMID:12912892] |
| GO:0015794 | 5.09394E-05 | 53 / 400 | "The process in which glycerol-3-phosphate is transported across a membrane. Glycerol-3-phosphate is a phosphoric monoester of glycerol." [GOC:ai] |
| GO:0015897 | 5.09394E-05 | 53 / 400 | "The process in which an organomercurial compound is transported across a membrane. Organomercurial substances are any organic compound containing a mercury atom." [GOC:ai, PMID:18793329] |
| GO:0015904 | 5.09394E-05 | 53 / 400 | "The directed movement of tetracycline from one side of a membrane to the other. Tetracycline is a broad spectrum antibiotic that blocks binding of aminoacyl tRNA to the ribosomes of both Gram-positive and Gram-negative organisms (and those of organelles)." [GOC:cur |
| GO:0015905 | 5.09394E-05 | 53 / 400 | "The directed movement of bicyclomycin across a lipid bilayer, from one side of a membrane to the other. Bicyclomycin (or bicozamycin) is an antibacterial drug often used as a livestock feed additive." [PMID:20067529] |
| GO:0035378 | 5.09394E-05 | 53 / 400 | "The process in which carbon dioxide (CO2) is transported across a membrane." [GOC:yaf] |
| GO:0035436 | 5.09394E-05 | 53 / 400 | "The process in which triose phosphate (glyceraldehyde 3-phosphate) is transported across a membrane. Glyceraldehyde 3-phosphate is any organic three carbon compound phosphate ester." [GOC:bf, ISBN:0198506732] |
| GO:0070811 | 5.09394E-05 | 53 / 400 | "The process in which glycerol-2-phosphate is transported across a membrane. Glycerol-2-phosphate is a phosphoric monoester of glycerol." [GOC:mah] |
| GO:0071918 | 5.09394E-05 | 53 / 400 | "The process in which urea, the water-soluble compound H2N-CO-NH2, is transported from one side of a membrane to the other by means of some agent such as a transporter or pore." [GOC:mah] |
| GO:0072489 | 5.09394E-05 | 53 / 400 | "The process in which methylammonium is transported across a membrane." [GOC:mah] |
| GO:0072531 | 5.09394E-05 | 53 / 400 | "The process in which a pyrimidine-containing compound is transported across a membrane. A pyrimidine-containing compound is any compound that contains pyrimidine or a formal derivative thereof." [GOC:mah] |

| GO_Term | P_val | Counts | CellComp_Description |
|---|---|---|---|
| GO:0042176 | 0.000657896 | 3 / 3 | "Any process that modulates the frequency, rate or extent of the chemical reactions and pathways resulting in the breakdown of a protein by the destruction of the native, active configuration, with or without the hydrolysis of peptide bonds." [GOC:go_curators, GOC:jl] |
| GO:0005761 | 0.001017123 | 0 / 75 | "A ribosome found in the mitochondrion of a eukaryotic cell; contains a characteristic set of proteins distinct from those of cytosolic ribosomes." [GOC:mah, ISBN:0198506732] |
| GO:0009987 | 0.002230876 | 29 / 508 | "Any process that is carried out at the cellular level, but not necessarily restricted to a single cell. For example, cell communication occurs among more than one cell, but occurs at the cellular level." [GOC:go_curators, GOC:isa_complete] |
| GO:0030120 | 0.00277821 | 4 / 8 | "A membrane coat found on a coated vesicle." [GOC:mah] |
| GO:0031981 | 0.002861538 | 29 / 501 | "The volume enclosed by the nuclear inner membrane." [GOC:mah, GOC:pz] |
| GO:0140513 | 0.003423174 | 30 / 509 | "A stable assembly of two or more macromolecules, i.e. proteins, nucleic acids, carbohydrates or lipids, in which at least one component is a protein and the constituent parts function together in the nucleus." [GOC:pg] |
| GO:0005886 | 0.003860362 | 27 / 192 | "The membrane surrounding a cell that separates the cell from its external environment. It consists of a phospholipid bilayer and associated proteins." [ISBN:0716731363] |
| GO:0098590 | 0.004106389 | 27 / 193 | "A membrane that is a (regional) part of the plasma membrane." [GOC:dos] |
| GO:0016020 | 0.004520903 | 28 / 204 | "A lipid bilayer along with all the proteins and protein complexes embedded in it and attached to it." [GOC:dos, GOC:mah, ISBN:0815316194] |
| GO:0008150 | 0.005252294 | 30 / 496 | "A biological process is the execution of a genetically-encoded biological module or program. It consists of all the steps required to achieve the specific biological objective of the module. A biological process is accomplished by a particular set of molecular functions carried c |
| GO:0022414 | 0.005601637 | 28 / 468 | "A biological process that directly contributes to the process of producing new individuals by one or two organisms. The new individuals inherit some proportion of their genetic material from the parent or parents." [GOC:dph, GOC:isa_complete] |
| GO:0098754 | 0.005601637 | 28 / 468 | "Any process that reduces or removes the toxicity of a toxic substance. These may include transport of the toxic substance away from sensitive areas and to compartments or complexes whose purpose is sequestration of the toxic substance." [GOC:dos] |
| GO:0050896 | 0.005976474 | 28 / 466 | "Any process that results in a change in state or activity of a cell or an organism (in terms of movement, secretion, enzyme production, gene expression, etc.) as a result of a stimulus. The process begins with detection of the stimulus and ends with a change in state or activi |
| GO:0040007 | 0.0061648 | 29 / 478 | "The increase in size or mass of an entire organism, a part of an organism or a cell." [GOC:bf, GOC:ma] |
| GO:0000003 | 0.006171736 | 28 / 465 | "The production of new individuals that contain some portion of genetic material inherited from one or more parent organisms." [GOC:go_curators, GOC:isa_complete, GOC:jl, ISBN:0198506732] |
| GO:0002376 | 0.006171736 | 28 / 465 | "Any process involved in the development or functioning of the immune system, an organismal system for calibrated responses to potential internal or invasive threats." [GO_REF:0000022, GOC:add] |
| GO:0016032 | 0.006171736 | 28 / 465 | "A multi-organism process in which a virus is a participant. The other participant is the host. Includes infection of a host cell, replication of the viral genome, and assembly of progeny virus particles. In some cases the viral genetic material may integrate into the host genome |
| GO:0032501 | 0.006171736 | 28 / 465 | "Any biological process, occurring at the level of a multicellular organism, pertinent to its function." [GOC:curators, GOC:dph, GOC:isa_complete, GOC:tb] |
| GO:0032502 | 0.006171736 | 28 / 465 | "A biological process whose specific outcome is the progression of an integrated living unit: an anatomical structure (which may be a subcellular structure, cell, tissue, or organ), or organism over time from an initial condition to a later condition." [GOC:isa_complete] |
| GO:0040011 | 0.006171736 | 28 / 465 | "Self-propelled movement of a cell or organism from one location to another." [GOC:dgh] |
| GO:0042592 | 0.006171736 | 28 / 465 | "Any biological process involved in the maintenance of an internal steady state." [GOC:jl, ISBN:0395825172] |
| GO:0043473 | 0.006171736 | 28 / 465 | "The accumulation of pigment in an organism, tissue or cell, either by increased deposition or by increased number of cells." [GOC:jl] |
| GO:0044419 | 0.006171736 | 28 / 465 | "Any process evolved to enable an interaction with an organism of a different species." [GOC:cc] |
| GO:0044848 | 0.006171736 | 28 / 465 | "A distinct period or stage in a biological process or cycle." [GOC:jl] |
| GO:0048511 | 0.006171736 | 28 / 465 | "Any process pertinent to the generation and maintenance of rhythms in the physiology of an organism." [GOC:jid] |
| GO:0051179 | 0.006171736 | 28 / 465 | "Any process in which a cell, a substance, or a cellular entity, such as a protein complex or organelle, is transported, tethered to or otherwise maintained in a specific location. In the case of substances, localization may also be achieved via selective degradation." [GOC:ai, G |
| GO:0051703 | 0.006171736 | 28 / 465 | "Any process in which an organism has an effect on an organism of the same species." [GOC:ai] |
| GO:0065007 | 0.006171736 | 28 / 465 | "Any process that modulates a measurable attribute of any biological process, quality or function." [GOC:dph, GOC:isa_complete, GOC:mah, GOC:pr, GOC:vw] |
| GO:0032991 | 0.006758506 | 391 / 4637 | "A stable assembly of two or more macromolecules, i.e. proteins, nucleic acids, carbohydrates or lipids, in which at least one component is a protein and the constituent parts function together." [GOC:dos, GOC:mah] |
| GO:0000018 | 0.007579786 | 2 / 2 | "Any process that modulates the frequency, rate or extent of DNA recombination, a DNA metabolic process in which a new genotype is formed by reassortment of genes resulting in gene combinations different from those that were present in the parents." [GOC:go_curator |
| GO:0006012 | 0.007579786 | 2 / 2 | "The chemical reactions and pathways involving galactose, the aldohexose galacto-hexose. D-galactose is widely distributed in combined form in plants, animals and microorganisms as a constituent of oligo- and polysaccharides; it also occurs in galactolipids and as its gluco |
| GO:0006820 | 0.007579786 | 2 / 2 | "The directed movement of a monoatomic anion, into, out of or within a cell, or between cells, by means of some agent such as a transporter or pore. Monatomic anions (also called simple anions) are negatively charged ions consisting of exactly one atom." [GOC:ai] |
| GO:0006821 | 0.007579786 | 2 / 2 | "The directed movement of chloride into, out of or within a cell, or between cells, by means of some agent such as a transporter or pore." [GOC:krc] |
| GO:0007537 | 0.007579786 | 2 / 2 | "The inactivation of recombination at sequences around a mating type donor locus, with the consequence that the other donor is the only one available for mating type switching; exemplified by the HML locus and surrounding sequences on Chromosome III in Saccharomyc |
| GO:0015804 | 0.007579786 | 2 / 2 | "The directed movement of neutral amino acids, amino acids with no net charge, into, out of or within a cell, or between cells, by means of some agent such as a transporter or pore." [GOC:ai] |
| GO:0015807 | 0.007579786 | 2 / 2 | "The directed movement of L-enantiomer amino acids into, out of or within a cell, or between cells, by means of some agent such as a transporter or pore." [GOC:ai, GOC:jsg, GOC:mah] |
| GO:0019388 | 0.007579786 | 2 / 2 | "The chemical reactions and pathways resulting in the breakdown of galactose, the aldohexose galacto-hexose." [ISBN:0198506732] |
| GO:0045732 | 0.007579786 | 2 / 2 | "Any process that activates or increases the frequency, rate or extent of the chemical reactions and pathways resulting in the breakdown of a protein by the destruction of the native, active configuration, with or without the hydrolysis of peptide bonds." [GOC:go_curators] |
| GO:0045829 | 0.007579786 | 2 / 2 | "Any process that stops, prevents, or reduces the frequency, rate or extent of isotype switching." [GOC:go_curators] |
| GO:0045910 | 0.007579786 | 2 / 2 | "Any process that stops, prevents, or reduces the frequency, rate or extent of DNA recombination." [GOC:go_curators] |
| GO:0045950 | 0.007579786 | 2 / 2 | "Any process that inhibits or decreases the rate of DNA recombination during mitosis." [GOC:go_curators, GOC:hjd] |
| GO:0048239 | 0.007579786 | 2 / 2 | "Any process that stops, prevents, or reduces the frequency, rate or extent of genetic recombination within the telomere." [GOC:jid, PMID:9635193] |
| GO:0051053 | 0.007579786 | 2 / 2 | "Any process that stops, prevents, or reduces the frequency, rate or extent of the chemical reactions and pathways involving DNA." [GOC:ai] |
| GO:0060543 | 0.007579786 | 2 / 2 | "Any process that decreases the rate, frequency or extent of strand invasion. Strand invasion is the process in which the nucleoprotein complex (composed of the broken single-strand DNA and the recombinase) searches and identifies a region of homology in intact duplex I |
| GO:0061136 | 0.007579786 | 2 / 2 | "Any process that modulates the rate, frequency, or extent of the chemical reactions and pathways resulting in the breakdown of a protein or peptide by hydrolysis of its peptide bonds that is mediated by the proteasome." [GOC:dph, GOC:tb] |
| GO:0061808 | 0.007579786 | 2 / 2 | "Any process that stops, prevents, or reduces the frequency, rate or extent of genetic recombination at the centromere." [GOC:dph, GOC:mah] |
| GO:0120105 | 0.007579786 | 2 / 2 | "The region of the mitotic actomyosin ring in between the proximal layer and the actin filament layer. This region contains the accessory protein network, some actin filaments and connections between the proximal layer and the actin filament layer." [GOC:krc, GOC:vw, PM |
| GO:1901800 | 0.007579786 | 2 / 2 | "Any process that activates or increases the frequency, rate or extent of proteasomal protein catabolic process." [GOC:BHF, GOC:rl, GOC:TermGenie, PMID:21669198] |
| GO:1903111 | 0.007579786 | 2 / 2 | "Any process that stops, prevents or reduces the frequency, rate or extent of single-strand break repair via homologous recombination." [GO_REF:0000058, GOC:bhm, GOC:TermGenie, PMID:24339919] |
| GO:2000042 | 0.007579786 | 2 / 2 | "Any process that stops, prevents, or reduces the frequency, rate or extent of double-strand break repair via homologous recombination." [GOC:vw] |

#### Supplementary Data S3.4.9

| GO_Term | P_val | Counts | MolFunc_Description |
|---|---|---|---|
| GO:0005198 | 9.81447E-07 | 369 / 3741 | "The action of a molecule that contributes to the structural integrity of a complex." [GOC:mah, GOC:vw] |
| GO:0003676 | 1.01353E-06 | 28 / 531 | "Binding to a nucleic acid." [GOC:jl] |
| GO:0140657 | 1.24062E-05 | 375 / 3753 | "A molecular function characterized by the coupling of ATP hydrolysis to other steps of a reaction mechanism to make the reaction energetically favorable, for example to catalyze a reaction or drive transport against a concentration gradient." [PMID:24878343, PMID:25750732, PMID:32933017, PMI |
| GO:0016491 | 1.3493E-05 | 102 / 658 | "Catalysis of an oxidation-reduction (redox) reaction, a reversible chemical reaction in which the oxidation state of an atom or atoms within a molecule is altered. One substrate acts as a hydrogen or electron donor and becomes oxidized, while the other acts as hydrogen or electron acceptor and becon |
| GO:0003943 | 1.43375E-05 | 5 / 5 | "Catalysis of the hydrolysis of the 4-sulfate groups of the N-acetyl-D-galactosamine 4-sulfate units of chondroitin sulfate and dermatan sulfate." [EC:3.1.6.12] |
| GO:0004065 | 1.43375E-05 | 5 / 5 | "Catalysis of the reaction: a phenol sulfate + H2O = a phenol + sulfate." [EC:3.1.6.1] |
| GO:0004098 | 1.43375E-05 | 5 / 5 | "Catalysis of the reaction: a cerebroside 3-sulfate + H2O = a cerebroside + sulfate." [EC:3.1.6.8] |
| GO:0004423 | 1.43375E-05 | 5 / 5 | "Catalysis of the hydrolysis of the 2-sulfate groups of the L-iduronate 2-sulfate units of dermatan sulfate, heparan sulfate and heparin." [EC:3.1.6.13] |
| GO:0004773 | 1.43375E-05 | 5 / 5 | "Catalysis of the reaction: 3-beta-hydroxyandrost-5-en-17-one 3-sulfate + H2O = 3-beta-hydroxyandrost-5-en-17-one + sulfate." [EC:3.1.6.2] |
| GO:0008449 | 1.43375E-05 | 5 / 5 | "Catalysis of the hydrolysis of the 6-sulfate group of the N-acetyl-D-glucosamine 6-sulfate units of heparan sulfate and keratan sulfate." [EC:3.1.6.14] |
| GO:0015024 | 1.43375E-05 | 5 / 5 | "Catalysis of the hydrolysis of the 2-sulfate groups of the 2-O-sulfo-D-glucuronate residues of chondroitin sulfate, heparin and heparitin sulfate." [EC:3.1.6.18] |
| GO:0018741 | 1.43375E-05 | 5 / 5 | "Catalysis of the reaction: a primary linear alkyl sulfate ester + H2O =>a primary alcohol + H(+) + sulfate." [RHEA:67908, UM-BBD_reactionID:r0602] |
| GO:0033886 | 1.43375E-05 | 5 / 5 | "Catalysis of the hydrolysis of the 2- and 3-sulfate groups of the polysulfates of cellulose and charonin." [EC:3.1.6.7] |
| GO:0033887 | 1.43375E-05 | 5 / 5 | "Catalysis of the reaction: 4-deoxy-beta-D-gluc-4-enuronosyl-(1->3)-N-acetyl-D-galactosamine 4-sulfate + H2O = 4-deoxy-beta-D-gluc-4-enuronosyl-(1->3)-N-acetyl-D-galactosamine + H+ + sulfate." [EC:3.1.6.9, RHEA:11444] |
| GO:0033888 | 1.43375E-05 | 5 / 5 | "Catalysis of the reaction: 4-deoxy-beta-D-gluc-4-enuronosyl-(1->3)-N-acetyl-D-galactosamine 6-sulfate + H2O = 4-deoxy-beta-D-gluc-4-enuronosyl-(1->3)-N-acetyl-D-galactosamine + H+ + sulfate." [EC:3.1.6.10, RHEA:10536] |
| GO:0033889 | 1.43375E-05 | 5 / 5 | "Catalysis of the hydrolysis of the 3-sulfate groups of the N-sulfo-D-glucosamine 3-O-sulfate units of heparin." [EC:3.1.6.15] |
| GO:0034889 | 1.43375E-05 | 5 / 5 | "Catalysis of the reaction: endosulfan hemisulfate + H2O = endosulfan monoalcohol + 2 H+ + sulfate." [UM-BBD_reactionID:r1384] |
| GO:0034902 | 1.43375E-05 | 5 / 5 | "Catalysis of the reaction: endosulfan sulfate + H2O = endosulfan diol + sulfite." [UM-BBD_reactionID:r1387] |
| GO:0043890 | 1.43375E-05 | 5 / 5 | "Catalysis of the hydrolysis of the 6-sulfate groups of the N-acetyl-D-galactosamine 6-sulfate units of chondroitin sulfate and of the D-galactose 6-sulfate units of keratan sulfate." [EC:3.1.6.4] |
| GO:0047753 | 1.43375E-05 | 5 / 5 | "Catalysis of the reaction: choline sulfate + H2O = choline + H+ + sulfate." [EC:3.1.6.6, RHEA:20820] |
| GO:0047825 | 1.43375E-05 | 5 / 5 | "Catalysis of the reaction: (R)-2-O-sulfolactate + H2O = (R)-lactate + H+ + sulfate." [EC:3.1.6.17, RHEA:20337] |
| GO:0047871 | 1.43375E-05 | 5 / 5 | "Catalysis of the reaction: N(2),6-disulfo-D-glucosamine + H2O = N-sulfo-D-glucosamine + H+ + sulfate." [EC:3.1.6.11, RHEA:15517] |
| GO:0047966 | 1.43375E-05 | 5 / 5 | "Catalysis of the reaction: D-glucose 6-sulfate + H2O = D-glucose + H+ + sulfate." [EC:3.1.6.3, RHEA:19145] |
| GO:0050106 | 1.43375E-05 | 5 / 5 | "Catalysis of the reaction: H2O + monomethyl sulfate = H+ + methanol + sulfate." [EC:3.1.6.16, RHEA:14221] |
| GO:0005215 | 3.08961E-05 | 373 / 3723 | "Enables the directed movement of substances (such as macromolecules, small molecules, ions) into, out of or within a cell, or between cells." [GOC:ai, GOC:dgf] |
| GO:0003677 | 3.21947E-05 | 32 / 524 | "Any molecular function by which a gene product interacts selectively and non-covalently with DNA (deoxyribonucleic acid)." [GOC:dph, GOC:jl, GOC:tb, GOC:vw] |
| GO:0140110 | 3.40528E-05 | 367 / 3673 | "A molecular function that controls the rate, timing and/or magnitude of gene transcription. The function of transcriptional regulators is to modulate gene expression at the transcription step so that they are expressed in the right cell at the right time and in the right amount throughout the life of the ce |
| GO:0009055 | 0.000110994 | 367 / 3652 | "A molecular function representing the directed movement of electrons from one molecular entity to another, typically mediated by electron carriers or acceptors, resulting in the transfer of energy and/or the reduction-oxidation (redox) transformation of chemical species. This activity is fundamental t |
| GO:0031490 | 0.000129552 | 17 / 324 | "Binding to DNA that is assembled into chromatin." [GOC:mah] |
| GO:0000099 | 0.000142657 | 7 / 13 | "Enables the transfer of sulfur amino acids from one side of a membrane to the other. Sulphur amino acids contain sulfur in the form of cystine, methionine or their derivatives." [GOC:ai, GOC:mtg_transport, ISBN:0815340729] |
| GO:0140223 | 0.000145696 | 364 / 3622 | "A molecular function required for core promoter activity that mediates the assembly of the RNA polymerase holoenzyme at promoter DNA to form the pre-initiation complex (PIC). General transcription factors (GTFs) bind to and open promoter DNA, initiate RNA synthesis and stimulate the escape of |
| GO:0016209 | 0.000153289 | 364 / 3621 | "Inhibition of the reactions brought about by dioxygen (O2) or peroxides. Usually the antioxidant is effective because it can itself be more easily oxidized than the substance protected. The term is often applied to components that can trap free radicals, thereby breaking the chain reaction that normally |
| GO:0015179 | 0.000153707 | 8 / 17 | "Enables the transfer of an L-amino acid from one side of a membrane to the other. L-amino acids are the L-enantiomers of amino acids." [GOC:ai, GOC:jsg, GOC:mah, GOC:mtg_transport, ISBN:0815340729] |
| GO:0098772 | 0.000169545 | 364 / 3619 | "A molecular function regulator regulates the activity of its target via non-covalent binding that does not result in covalent modification to the target. Examples of molecular function regulators include regulatory subunits of multimeric enzymes and channels. Mechanisms of regulation include allosteric |
| GO:0045735 | 0.000178237 | 364 / 3618 | "Functions in the storage of nutritious substrates." [GOC:ai] |
| GO:0044183 | 0.000206753 | 364 / 3615 | "Binding to a protein or a protein-containing complex to assist the protein folding process." [GOC:mtg_cambridge_2009] |
| GO:0045182 | 0.000206753 | 364 / 3615 | "Any molecular function involved in the initiation, activation, perpetuation, repression or termination of polypeptide synthesis at the ribosome." [GOC:ai] |
| GO:0003774 | 0.00021006 | 365 / 3623 | "Generation of force resulting in movement, for example along a microfilament or microtubule, or in torque resulting in membrane scission or rotation of a flagellum. The energy required is obtained either from the hydrolysis of a nucleoside triphosphate or by an electrochemical proton gradient (protc |
| GO:0140104 | 0.000217124 | 364 / 3614 | "Directly binding to a specific ion or molecule and delivering it either to an acceptor molecule or to a specific location." [GOC:molecular_function_refactoring, GOC:pdt] |
| GO:0019843 | 0.000225611 | 2 / 112 | "Binding to a ribosomal RNA." [GOC:jl] |
| GO:0060089 | 0.000227955 | 364 / 3613 | "A compound molecular function in which an effector function is controlled by one or more regulatory components." [GOC:dos, GOC:pdt] |
| GO:0090729 | 0.000227955 | 364 / 3613 | "Interacting selectively with one or more biological molecules in another (target) organism, initiating pathogenesis (leading to an abnormal, generally detrimental state) in the target organism. The activity should refer to an evolved function of the active gene product, i.e. one that was selected for. Exam |
| GO:0140299 | 0.000227955 | 364 / 3613 | "Binding to a small molecule and eliciting a change in the protein's activity in response to the intracellular level of that small molecule." [PMID:26328879] |
| GO:0140313 | 0.000227955 | 364 / 3613 | "Binding to a specific molecule to prevent it from interacting with other partners or to inhibit its localization to the area of the cell or complex where it is active." [PMID:13130076] |
| GO:0140489 | 0.000227955 | 364 / 3613 | "The action of a molecule that provides a shape or a sequence mimicking or complementary to the final product, providing template for copying the original molecule's shape or sequence." [GOC:pg] |
| GO:0140522 | 0.000227955 | 364 / 3613 | "The activity of joining two lipid bilayers to form a single membrane." [PMID:10332732, PMID:11493675, PMID:12600315, PMID:32100701, PMID:32641474] |
| GO:0140691 | 0.000227955 | 364 / 3613 | "Binding to an RNA or an RNA-containing complex to assist the folding process." [PMID:31165735] |
| GO:0140776 | 0.000227955 | 364 / 3613 | "A molecular function that involves direct binding to one of the subunits of a protein-containing complex and promoting the dissociation of one or many subunits. This often happens by changing the conformation of the protein being bound, which decreases its affinity for the rest of the complex." [PM |
| GO:0140777 | 0.000227955 | 364 / 3613 | "A molecular function that involves direct binding to one of the subunits of a protein-containing complex, thus preventing an interaction with a factor that would promote dissociation of the complex." [PMID:34040253] |
| GO:0140911 | 0.000227955 | 364 / 3613 | "An activity in which a protein is inserted into the membrane of another cell where it forms transmembrane pores. Pores disrupts the integrity of the cell membrane, resulting in deregulated ion homeostasis, cellular dysfunction, and can result in cell death." [PMID:1406491, PMID:25157079, PMID:34 |

| GO_Term | P_val | Counts | BioProc_Description |
|---|---|---|---|
| GO:0055085 | 3.3557E-08 | 79 / 423 | "The process in which a solute is transported across a lipid bilayer, from one side of a membrane to the other." [GOC:dph, GOC:jid] |
| GO:0045117 | 7.33348E-08 | 75 / 401 | "The directed movement of azoles, heterocyclic compounds found in many biologically important substances, across a lipid bilayer, across a membrane." [GOC:go_curators, ISBN:3527307206, Wikipedia:Azole] |
| GO:0097080 | 7.33348E-08 | 75 / 401 | "The directed movement of inorganic selenite (HSeO3-1 at physiological pH) across a plasma membrane." [GOC:mcc, PMID:20861301] |
| GO:0006855 | 8.14922E-08 | 75 / 402 | "The process in which a xenobiotic, a compound foreign to the organim exposed to it, is transported across a membrane. It may be synthesized by another organism (like ampicilin) or it can be a synthetic chemical." [GOC:ai, GOC:bf, GOC:krc] |
| GO:0015780 | 8.14922E-08 | 75 / 402 | "The directed movement of nucleotide-sugars into, out of or within a cell, or between cells, by means of some agent such as a transporter or pore. Nucleotide-sugars are any nucleotide in which the distal phosphoric residue of a nucleoside 5'-diphosphate is in glycosidic linkage with a monosaccharide c |
| GO:0006810 | 8.51766E-08 | 79 / 432 | "The directed movement of substances (such as macromolecules, small molecules, ions) or cellular components (such as complexes and organelles) into, out of or within a cell, or between cells, or within a multicellular organism by means of some agent such as a transporter or a transporter complex, a |
| GO:0034220 | 9.05028E-08 | 75 / 403 | "A process in which a monoatomic ion is transported across a membrane. Monatomic ions (also called simple ions) are ions consisting of exactly one atom." [GOC:mah] |
| GO:0046618 | 9.05028E-08 | 75 / 403 | "The directed movement of a xenobiotic from a cell, into the extracellular region. A xenobiotic is a compound foreign to the organim exposed to it. It may be synthesized by another organism (like ampicilin) or it can be a synthetic chemical." [GOC:go_curators, GOC:krc] |
| GO:0046967 | 9.05028E-08 | 75 / 403 | "The directed movement of substances from the cytosol to the endoplasmic reticulum of a cell." [GOC:ai] |
| GO:1902047 | 9.05028E-08 | 75 / 403 | "The process in which a polyamine macromolecule is transported across a membrane." [GOC:TermGenie, GOC:vw] |
| GO:0071806 | 1.07095E-07 | 76 / 412 | "The process in which a protein is transported across a membrane." [GOC:mah, GOC:vw] |
| GO:0140115 | 1.23525E-07 | 75 / 406 | "The directed movement of some substance from inside of a cell, across the plasma membrane and into the extracellular region." [GOC:pg] |
| GO:0001407 | 1.42381E-07 | 74 / 400 | "The process in which a glycerophosphodiester is transported across a membrane. Glycerophosphodiesters are small molecules composed of glycerol-3-phosphate and an alcohol, for example, glycerophosphoinositol." [GOC:mcc, PMID:12912892] |
| GO:0015794 | 1.42381E-07 | 74 / 400 | "The process in which glycerol-3-phosphate is transported across a membrane. Glycerol-3-phosphate is a phosphoric monoester of glycerol." [GOC:ai] |
| GO:0015897 | 1.42381E-07 | 74 / 400 | "The process in which an organomercurial compound is transported across a membrane. Organomercurial substances are any organic compound containing a mercury atom." [GOC:ai, PMID:18793329] |
| GO:0015904 | 1.42381E-07 | 74 / 400 | "The directed movement of tetracycline from one side of a membrane to the other. Tetracycline is a broad spectrum antibiotic that blocks binding of aminoacyl tRNA to the ribosomes of both Gram-positive and Gram-negative organisms (and those of organelles)." [GOC:curators] |
| GO:0015905 | 1.42381E-07 | 74 / 400 | "The directed movement of bicyclomycin across a lipid bilayer, from one side of a membrane to the other. Bicyclomycin (or bicozamycin) is an antibacterial drug often used as a livestock feed additive." [PMID:20067529] |
| GO:0035378 | 1.42381E-07 | 74 / 400 | "The process in which carbon dioxide (CO2) is transported across a membrane." [GOC:yaf] |
| GO:0035436 | 1.42381E-07 | 74 / 400 | "The process in which triose phosphate (glyceraldehyde 3-phosphate) is transported across a membrane. Glyceraldehyde 3-phosphate is any organic three carbon compound phosphate ester." [GOC:bf, ISBN:0198506732] |
| GO:0070811 | 1.42381E-07 | 74 / 400 | "The process in which glycerol-2-phosphate is transported across a membrane. Glycerol-2-phosphate is a phosphoric monoester of glycerol." [GOC:mah] |
| GO:0071918 | 1.42381E-07 | 74 / 400 | "The process in which urea, the water-soluble compound H2N-CO-NH2, is transported from one side of a membrane to the other by means of some agent such as a transporter or pore." [GOC:mah] |
| GO:0072489 | 1.42381E-07 | 74 / 400 | "The process in which methylammonium is transported across a membrane." [GOC:mah] |
| GO:0072531 | 1.42381E-07 | 74 / 400 | "The process in which a pyrimidine-containing compound is transported across a membrane. A pyrimidine-containing compound is any compound that contains pyrimidine or a formal derivative thereof." [GOC:mah] |
| GO:0080162 | 1.42381E-07 | 74 / 400 | "The directed movement of auxins from endoplasmic reticulum to cytosol." [PMID:19506555] |
| GO:0080170 | 1.42381E-07 | 74 / 400 | "The process in which hydrogen peroxide is transported across a membrane." [GOC:tb] |
| GO:0098660 | 1.42381E-07 | 74 / 400 | "The process in which an inorganic ion is transported across a membrane." [GOC:mah] |
| GO:0140317 | 1.42381E-07 | 74 / 400 | "The directed movement of a substance across the outer membrane in cells with two membranes." [PMID:15968039] |
| GO:0170006 | 1.42381E-07 | 74 / 400 | "The process in which lipoate is transported across a membrane." [PMID:20980265, PMID:25971966] |
| GO:1901642 | 1.42381E-07 | 74 / 400 | "The directed movement of nucleoside across a membrane." [GOC:pr, GOC:TermGenie] |
| GO:1901684 | 1.42381E-07 | 74 / 400 | "The process in which arsenate is transported across a membrane." [GOC:TermGenie] |
| GO:1901962 | 1.42381E-07 | 74 / 400 | "The directed movement of S-adenosyl-L-methionine across a membrane." [GOC:TermGenie, PMID:10497160] |
| GO:1902418 | 1.42381E-07 | 74 / 400 | "The process in which (+)-abscisic acid D-glucopyranosyl este is transported across a membrane." [GOC:TermGenie, PMID:24028845] |
| GO:1902603 | 1.42381E-07 | 74 / 400 | "The directed movement of carnitine across a membrane." [GO_REF:0000069, GOC:pr, GOC:TermGenie] |
| GO:1902616 | 1.42381E-07 | 74 / 400 | "The process in which acyl carnitine is transported across a membrane." [GO_REF:0000069, GOC:pr, GOC:TermGenie] |
| GO:1903825 | 1.42381E-07 | 74 / 400 | "The process in which an organic acid is transported across a membrane." [GO_REF:0000069, GOC:TermGenie] |
| GO:0071994 | 1.57782E-07 | 74 / 401 | "The process in which a phytochelatin is transported across a membrane. Phytochelatins are a group of peptides that bind metals (Cd, Zn, Cu, Pb, Hg) in thiolate coordination complexes." [GOC:mah, ISBN:0198506732] |
| GO:0140820 | 1.57782E-07 | 74 / 401 | "The directed movement of substances from the cytosol into the Golgi apparatus of a cell." [PMID:28373556] |
| GO:0015791 | 1.74746E-07 | 74 / 402 | "The directed movement of polyols, any polyhydric alcohol, across a membrane." [GOC:ai] |
| GO:0034219 | 1.74746E-07 | 74 / 402 | "The process in which a carbohydrate is transported across a membrane." [GOC:mah] |
| GO:0035382 | 1.74746E-07 | 74 / 402 | "The process in which a sterol is transported across a membrane. Sterols are steroids with one or more hydroxyl groups and a hydrocarbon side-chain in the molecule." [GOC:vw] |
| GO:0035461 | 1.74746E-07 | 74 / 402 | "The process in which a vitamin is transported across a membrane. A vitamin is one of a number of unrelated organic substances that occur in many foods in small amounts and that are necessary in trace amounts for the normal metabolic functioning of the body." [GOC:bf] |
| GO:0072530 | 1.74746E-07 | 74 / 402 | "The process in which a purine-containing compound is transported across a membrane. A purine-containing compound is any compound that contains purine or a formal derivative thereof." [GOC:mah] |
| GO:1901679 | 1.74746E-07 | 74 / 402 | "The directed movement of nucleotide across a membrane." [GOC:pr, GOC:TermGenie] |
| GO:1990542 | 1.85491E-07 | 75 / 410 | "The process in which a solute is transported from one side of a membrane to the other into, out of or within a mitochondrion." [PMID:20533899] |
| GO:0034486 | 1.93419E-07 | 74 / 403 | "The process in which a solute is transported from one side of the vacuolar membrane to the other." [GOC:mah] |
| GO:0035672 | 2.13962E-07 | 74 / 404 | "The process in which an oligopeptide is transported across a membrane. Oligopeptides are molecules that contain a small number (2 to 20) of amino-acid residues connected by peptide linkages." [GOC:vw, ISBN:0198506732] |
| GO:0098739 | 2.13962E-07 | 74 / 404 | "The directed movement of some substance from outside of a cell, across the plasma membrane and into the cytosol." [GOC:dos] |
| GO:0030950 | 2.6036E-07 | 10 / 15 | "Any cellular process that results in the specification, formation or maintenance of polarized actin-based cytoskeletal structures." [GOC:mah] |
| GO:0030866 | 3.45392E-07 | 12 / 22 | "A process that is carried out at the cellular level which results in the assembly, arrangement of constituent parts, or disassembly of actin-based cytoskeletal structures in the cell cortex, i.e. just beneath the plasma membrane." [GOC:dph, GOC:jl, GOC:mah, GOC:pf] |
| GO:0040007 | 4.55441E-07 | 346 / 3597 | "The increase in size or mass of an entire organism, a part of an organism or a cell." [GOC:bf, GOC:ma] |

| GO_Term | P_val | Counts | CellComp_Description |
|---|---|---|---|
| GO:0030120 | 3.40838E-05 | 6 / 8 | "A membrane coat found on a coated vesicle." [GOC:mah] |
| GO:0005761 | 0.000181495 | 0 / 75 | "A ribosome found in the mitochondrion of a eukaryotic cell; contains a characteristic set of proteins distinct from those of cytosolic ribosomes." [GOC:mah, ISBN:0198506732] |
| GO:0032991 | 0.000293017 | 477 / 4637 | "A stable assembly of two or more macromolecules, i.e. proteins, nucleic acids, carbohydrates or lipids, in which at least one component is a protein and the constituent parts function together." [GOC:dos, GOC:mah] |
| GO:0009987 | 0.000860131 | 36 / 508 | "Any process that is carried out at the cellular level, but not necessarily restricted to a single cell. For example, cell communication occurs among more than one cell, but occurs at the cellular level." [GOC:go_curators, GOC:isa_complete] |
| GO:0031981 | 0.001174032 | 36 / 501 | "The volume enclosed by the nuclear inner membrane." [GOC:mah, GOC:pz] |
| GO:0022414 | 0.001203556 | 33 / 468 | "A biological process that directly contributes to the process of producing new individuals by one or two organisms. The new individuals inherit some proportion of their genetic material from the parent or parents." [GOC:dph, GOC:isa_complete] |
| GO:0098754 | 0.001203556 | 33 / 468 | "Any process that reduces or removes the toxicity of a toxic substance. These may include transport of the toxic substance away from sensitive areas and to compartments or complexes whose purpose is sequestration of the toxic substance." [GOC:dos] |
| GO:0006869 | 0.001244094 | 3 / 3 | "The directed movement of lipids into, out of or within a cell, or between cells, by means of some agent such as a transporter or pore. Lipids are compounds soluble in an organic solvent but not, or sparingly, in an aqueous solvent." [ISBN:0198506732] |
| GO:0042176 | 0.001244094 | 3 / 3 | "Any process that modulates the frequency, rate or extent of the chemical reactions and pathways resulting in the breakdown of a protein by the destruction of the native, active configuration, with or without the hydrolysis of peptide bonds." [GOC:go_curators, GOC:jl] |
| GO:0050896 | 0.001316671 | 33 / 466 | "Any process that results in a change in state or activity of a cell or an organism (in terms of movement, secretion, enzyme production, gene expression, etc.) as a result of a stimulus. The process begins with detection of the stimulus and ends with a change in state or activity or the cell or organism." [G |
| GO:0000003 | 0.001376763 | 33 / 465 | "The production of new individuals that contain some portion of genetic material inherited from one or more parent organisms." [GOC:go_curators, GOC:isa_complete, GOC:jl, ISBN:0198506732] |
| GO:0002376 | 0.001376763 | 33 / 465 | "Any process involved in the development or functioning of the immune system, an organismal system for calibrated responses to potential internal or invasive threats." [GO_REF:0000022, GOC:add] |
| GO:0016032 | 0.001376763 | 33 / 465 | "A multi-organism process in which a virus is a participant. The other participant is the host. Includes infection of a host cell, replication of the viral genome, and assembly of progeny virus particles. In some cases the viral genetic material may integrate into the host genome and only subsequently, unde |
| GO:0032501 | 0.001376763 | 33 / 465 | "Any biological process, occurring at the level of a multicellular organism, pertinent to its function." [GOC:curators, GOC:dph, GOC:isa_complete, GOC:tb] |
| GO:0032502 | 0.001376763 | 33 / 465 | "A biological process whose specific outcome is the progression of an integrated living unit: an anatomical structure (which may be a subcellular structure, cell, tissue, or organ), or organism over time from an initial condition to a later condition." [GOC:isa_complete] |
| GO:0040011 | 0.001376763 | 33 / 465 | "Self-propelled movement of a cell or organism from one location to another." [GOC:dgh] |
| GO:0042592 | 0.001376763 | 33 / 465 | "Any biological process involved in the maintenance of an internal steady state." [GOC:jl, ISBN:0395825172] |
| GO:0043473 | 0.001376763 | 33 / 465 | "The accumulation of pigment in an organism, tissue or cell, either by increased deposition or by increased number of cells." [GOC:jl] |
| GO:0044419 | 0.001376763 | 33 / 465 | "Any process evolved to enable an interaction with an organism of a different species." [GOC:cc] |
| GO:0044848 | 0.001376763 | 33 / 465 | "A distinct period or stage in a biological process or cycle." [GOC:jl] |
| GO:0048511 | 0.001376763 | 33 / 465 | "Any process pertinent to the generation and maintenance of rhythms in the physiology of an organism." [GOC:jid] |
| GO:0051179 | 0.001376763 | 33 / 465 | "Any process in which a cell, a substance, or a cellular entity, such as a protein complex or organelle, is transported, tethered to or otherwise maintained in a specific location. In the case of substances, localization may also be achieved via selective degradation." [GOC:ai, GOC:dos] |
| GO:0051703 | 0.001376763 | 33 / 465 | "Any process in which an organism has an effect on an organism of the same species." [GOC:ai] |
| GO:0065007 | 0.001376763 | 33 / 465 | "Any process that modulates a measurable attribute of any biological process, quality or function." [GOC:dph, GOC:isa_complete, GOC:mah, GOC:pr, GOC:vw] |
| GO:0005575 | 0.001403565 | 475 / 4589 | "A location, relative to cellular compartments and structures, occupied by a macromolecular machine. There are three types of cellular components described in the gene ontology: (1) the cellular anatomical entity where a gene product carries out a molecular function (e.g., plasma membrane, cytoskele |
| GO:0044423 | 0.001403565 | 475 / 4589 | "Any constituent part of a virion, a complete fully infectious extracellular virus particle." [GOC:jl] |
| GO:0008150 | 0.00145826 | 36 / 496 | "A biological process is the execution of a genetically-encoded biological module or program. It consists of all the steps required to achieve the specific biological objective of the module. A biological process is accomplished by a particular set of molecular functions carried out by specific gene products ( |
| GO:0043232 | 0.001507872 | 1 / 76 | "Organized structure of distinctive morphology and function, not bounded by a lipid bilayer membrane and occurring within the cell. Includes ribosomes, the cytoskeleton and chromosomes." [GOC:go_curators] |
| GO:0001411 | 0.001856529 | 6 / 14 | "The end, or tip, of a fungal hypha, where polarized growth occurs during hyphal elongation." [GOC:mcc] |
| GO:0040007 | 0.001988031 | 35 / 478 | "The increase in size or mass of an entire organism, a part of an organism or a cell." [GOC:bf, GOC:ma] |
| GO:0140513 | 0.002062124 | 38 / 509 | "A stable assembly of two or more macromolecules, i.e. proteins, nucleic acids, carbohydrates or lipids, in which at least one component is a protein and the constituent parts function together in the nucleus." [GOC:pg] |
| GO:0008152 | 0.002421436 | 38 / 505 | "The chemical reactions and pathways, including anabolism and catabolism, by which living organisms transform chemical substances. Metabolic processes typically transform small molecules, but also include macromolecular processes such as DNA repair and replication, and protein synthesis and deg |
| GO:0000313 | 0.00325372 | 0 / 50 | "A ribosome contained within a subcellular membrane-bounded organelle." [GOC:mah, GOC:mcc] |
| GO:0016310 | 0.003323648 | 4 / 7 | "The process of introducing a phosphate group into a molecule, usually with the formation of a phosphoric ester, a phosphoric anhydride or a phosphoric amide." [ISBN:0198506732] |
| GO:0051286 | 0.003949191 | 6 / 16 | "The region at the end of the longest axis of a cylindrical or elongated cell." [GOC:ai, GOC:mah] |
| GO:0030663 | 0.004442883 | 3 / 4 | "The lipid bilayer surrounding a COPI-coated vesicle." [GOC:mah] |
| GO:0005576 | 0.005349881 | 23 / 130 | "The space external to the outermost structure of a cell. For cells without external protective or external encapsulating structures this refers to space outside of the plasma membrane. This term covers the host cell environment outside an intracellular parasite." [GOC:go_curators] |
| GO:0005602 | 0.005349881 | 23 / 130 | "A protein complex composed of six subunits of C1q, each formed of the three homologous polypeptide chains C1QA, C1QB, and C1QB, and tetramer of two C1QR and two C1QS polypeptide chains." [GOC:add, ISBN:0781735149] |
| GO:0005615 | 0.005349881 | 23 / 130 | "That part of a multicellular organism outside the cells proper, usually taken to be outside the plasma membranes, and occupied by fluid." [ISBN:0198547684] |
| GO:0010367 | 0.005349881 | 23 / 130 | "A protein complex whose composition varies amongst species; in rice it probably exists in a homo-tetramer to homo-hexamer form and in Gram negative bacteria as a dimer. Functions in the hydrolysis of alpha-(1,6)-D-glucosidic branch linkages. Isoamylases in animals are localized in the extracellular |
| GO:0031395 | 0.005349881 | 23 / 130 | "A neuropeptide hormone secreted by the central nervous system of insects that stimulates the tanning and sclerotization of the adult cuticle following eclosion. The active hormone consists of an obligate heterodimer of the alpha and beta subunits." [GOC:rc] |
| GO:0032311 | 0.005349881 | 23 / 130 | "A stable heterodimer of angiogenin and placental ribonuclease inhibitor; interaction between angiogenin and PRI prevents angiogenin binding to its receptor to stimulate angiogenesis." [PMID:2706246, PMID:3470787] |
| GO:0035182 | 0.005349881 | 23 / 130 | "An electron opaque backbone of the insect ovarian ring canal that is a part of or adjacent to the plasma membrane. The outer rim is established as the cleavage furrow is arrested, and contains F-actin, anillin, glycoproteins and at least one a protein with a high content of phosphorylated tyrosine residu |
| GO:0035183 | 0.005349881 | 23 / 130 | "A proteinaceous actin-rich layer of the insect ovarian ring canal that forms subcortically to the outer rim. The electron dense inner rim accumulates after the final mitotic division of each germline syncytia, and contains actin, a phosphotyrosine protein, and a number of cytoskeletal proteins." [PMID:10 |
| GO:0036117 | 0.005349881 | 23 / 130 | "A cable structure, surrounding some cell types (e.g. proximal or bronchial tubular epithelial cells), and composed of hyaluranon (HA), a ubiquitous connective tissue glycosaminoglycan." [GOC:yaf, PMID:16900089] |
| GO:0043083 | 0.005349881 | 23 / 130 | "The narrow gap that separates the presynaptic and postsynaptic membranes, into which neurotransmitter is released." [GOC:jl, http://synapses.mcg.edu/anatomy/chemical/synapse.stm] |
| GO:0043230 | 0.005349881 | 23 / 130 | "Organized structure of distinctive morphology and function, occurring outside the cell. Includes, for example, extracellular membrane vesicles (EMVs) and the cellulosomes of anaerobic bacteria and fungi." [GOC:jl, PMID:9914479] |
| GO:0043245 | 0.005349881 | 23 / 130 | "The environmental space outside of an organism; this may be a host organism in the case of parasitic and symbiotic organisms." [GOC:jl] |
| GO:0048046 | 0.005349881 | 23 / 130 | "The cell membranes and intracellular regions in a plant are connected through plasmodesmata, and plants may be described as having two major compartments: the living symplast and the non-living apoplast. The apoplast is external to the plasma membrane and includes cell walls, intercellular spaces |
| GO:0061696 | 0.005349881 | 23 / 130 | "A protein complex that is a protein hormone secreted by gonadotrope cells of the anterior pituitary of vertebrates. capable of regulating normal growth, sexual development, and reproductive function." [GOC:dph, PMID:11420129] |

#### Supplementary Data S3.4.10

| GO_Term | P_val | Counts | MolFunc_Description |
|---|---|---|---|
| GO:0003676 | 9.25791E-09 | 37 / 531 | "Binding to a nucleic acid." [GOC:jl] |
| GO:0003677 | 9.4624E-08 | 39 / 524 | "Any molecular function by which a gene product interacts selectively and non-covalently with DNA (deoxyribonucleic acid)." [GOC:dph, GOC:jl, GOC:tb, GOC:vw] |
| GO:0031490 | 3.07659E-07 | 19 / 324 | "Binding to DNA that is assembled into chromatin." [GOC:mah] |
| GO:0008270 | 4.25796E-07 | 37 / 489 | "Binding to a zinc ion (Zn)." [GOC:ai] |
| GO:0005198 | 1.23099E-06 | 500 / 3741 | "The action of a molecule that contributes to the structural integrity of a complex." [GOC:mah, GOC:vw] |
| GO:0016709 | 2.50105E-06 | 11 / 17 | "Catalysis of an oxidation-reduction (redox) reaction in which hydrogen or electrons are transferred from NADH or NADPH and one other donor, and one atom of oxygen is incorporated into one donor." [GOC:mah] |
| GO:0000981 | 3.65081E-06 | 16 / 270 | "A DNA-binding transcription factor activity that modulates the transcription of specific gene sets transcribed by RNA polymerase II." [GOC:txnOH-2018] |
| GO:0003697 | 5.49244E-06 | 21 / 314 | "Binding to single-stranded DNA." [GOC:elh, GOC:vw, PMID:22976174] |
| GO:0003735 | 5.93048E-06 | 3 / 123 | "The action of a molecule that contributes to the structural integrity of the ribosome." [GOC:mah] |
| GO:0003700 | 6.02499E-06 | 16 / 265 | "A transcription regulator activity that modulates transcription of gene sets via selective and non-covalent binding to a specific double-stranded genomic DNA sequence (sometimes referred to as a motif) within a cis-regulatory region. Regulatory regions include promoters (proximal and distal) and enha |
| GO:0043565 | 6.61447E-06 | 21 / 312 | "Binding to DNA of a specific nucleotide composition, e.g. GC-rich DNA binding, or with a specific sequence motif or type of DNA e.g. promotor binding or rDNA binding." [GOC:jl] |
| GO:0000217 | 6.65153E-06 | 19 / 293 | "Binding to a DNA secondary structure element such as a four-way junction, a bubble, a loop, Y-form DNA, or a double-strand/single-strand junction." [GOC:krc] |
| GO:0008301 | 8.03985E-06 | 19 / 291 | "The activity of binding selectively and non-covalently to and distorting the original structure of DNA, typically a straight helix, into a bend, or increasing the bend if the original structure was intrinsically bent due to its sequence." [GOC:krc, GOC:vw, PMID:10710711, PMID:19037758] |
| GO:0045027 | 8.03985E-06 | 19 / 291 | "Binding to DNA ends exposed by the creation of double-strand breaks (DSBs)." [GOC:jl] |
| GO:0070336 | 8.03985E-06 | 19 / 291 | "Binding to a flap structure in DNA. A DNA flap structure is one in which a single-stranded length of DNA or RNA protrudes from a double-stranded DNA molecule." [GOC:mah, PMID:15189154] |
| GO:0000497 | 8.83542E-06 | 19 / 290 | "Binding to nucleic acid via hydrogen bonds between the bases of a gene product molecule and the bases of a target DNA molecule." [GOC:krc] |
| GO:0003681 | 8.83542E-06 | 19 / 290 | "Binding to DNA in a bent conformation." [GOC:jl, PMID:12627977] |
| GO:0010844 | 8.83542E-06 | 19 / 290 | "Binding to a genomic region which promotes recombination." [GOC:dph, GOC:tb] |
| GO:0045142 | 8.83542E-06 | 19 / 290 | "Binding to a DNA triple helix. The formation of triple helical DNA has been evoked in several cellular processes including transcription, replication, and recombination." [PMID:10681538] |
| GO:0051880 | 8.83542E-06 | 19 / 290 | "Binding to G-quadruplex DNA structures, in which groups of four guanines adopt a flat, cyclic Hoogsteen hydrogen-bonding arrangement known as a guanine tetrad. The stacking of guanine tetrads results in G-quadruplex DNA structures. G-quadruplex DNA can form under physiological conditions fro |
| GO:0099077 | 8.83542E-06 | 19 / 290 | "DNA-binding activity that is dependent on binding to a histone." [PMID:11691835] |
| GO:0003690 | 1.28898E-05 | 22 / 314 | "Binding to double-stranded DNA." [GOC:elh, GOC:vw] |
| GO:0001228 | 2.95959E-05 | 15 / 239 | "A DNA-binding transcription factor activity that activates or increases transcription of specific gene sets transcribed by RNA polymerase II." [GOC:aruk, GOC:txnOH-2018, PMID:20737563, PMID:27145859] |
| GO:0003684 | 3.10976E-05 | 22 / 304 | "Binding to damaged DNA." [GOC:jl] |
| GO:0016491 | 3.5859E-05 | 127 / 658 | "Catalysis of an oxidation-reduction (redox) reaction, a reversible chemical reaction in which the oxidation state of an atom or atoms within a molecule is altered. One substrate acts as a hydrogen or electron donor and becomes oxidized, while the other acts as hydrogen or electron acceptor and becom |
| GO:0001227 | 3.9433E-05 | 15 / 236 | "A DNA-binding transcription factor activity that represses or decreases the transcription of specific gene sets transcribed by RNA polymerase II." [GOC:txnOH-2018] |
| GO:0004879 | 4.76688E-05 | 15 / 234 | "A DNA-binding transcription factor activity regulated by binding to a ligand that modulates the transcription of specific gene sets transcribed by RNA polymerase II. Nuclear receptor ligands are usually lipid-based (such as a steroid hormone) and the binding of the ligand to its receptor often occurs in th |
| GO:0140657 | 5.94441E-05 | 510 / 3753 | "A molecular function characterized by the coupling of ATP hydrolysis to other steps of a reaction mechanism to make the reaction energetically favorable, for example to catalyze a reaction or drive transport against a concentration gradient." [PMID:24878343, PMID:25750732, PMID:32933017, PMII |
| GO:0003943 | 6.07435E-05 | 5 / 5 | "Catalysis of the hydrolysis of the 4-sulfate groups of the N-acetyl-D-galactosamine 4-sulfate units of chondroitin sulfate and dermatan sulfate." [EC:3.1.6.12] |
| GO:0004065 | 6.07435E-05 | 5 / 5 | "Catalysis of the reaction: a phenol sulfate + H2O = a phenol + sulfate." [EC:3.1.6.1] |
| GO:0004098 | 6.07435E-05 | 5 / 5 | "Catalysis of the reaction: a cerebroside 3-sulfate + H2O = a cerebroside + sulfate." [EC:3.1.6.8] |
| GO:0004423 | 6.07435E-05 | 5 / 5 | "Catalysis of the hydrolysis of the 2-sulfate groups of the L-iduronate 2-sulfate units of dermatan sulfate, heparan sulfate and heparin." [EC:3.1.6.13] |
| GO:0004773 | 6.07435E-05 | 5 / 5 | "Catalysis of the reaction: 3-beta-hydroxyandrost-5-en-17-one 3-sulfate + H2O = 3-beta-hydroxyandrost-5-en-17-one + sulfate." [EC:3.1.6.2] |
| GO:0008449 | 6.07435E-05 | 5 / 5 | "Catalysis of the hydrolysis of the 6-sulfate group of the N-acetyl-D-glucosamine 6-sulfate units of heparan sulfate and keratan sulfate." [EC:3.1.6.14] |
| GO:0015024 | 6.07435E-05 | 5 / 5 | "Catalysis of the hydrolysis of the 2-sulfate groups of the 2-O-sulfo-D-glucuronate residues of chondroitin sulfate, heparin and heparitin sulfate." [EC:3.1.6.18] |
| GO:0018741 | 6.07435E-05 | 5 / 5 | "Catalysis of the reaction: a primary linear alkyl sulfate ester + H2O =>a primary alcohol + H(+) + sulfate." [RHEA:67908, UM-BBD_reactionID:r0602] |
| GO:0033886 | 6.07435E-05 | 5 / 5 | "Catalysis of the hydrolysis of the 2- and 3-sulfate groups of the polysulfates of cellulose and charonin." [EC:3.1.6.7] |
| GO:0033887 | 6.07435E-05 | 5 / 5 | "Catalysis of the reaction: 4-deoxy-beta-D-gluc-4-enuronosyl-(1->3)-N-acetyl-D-galactosamine 4-sulfate + H2O = 4-deoxy-beta-D-gluc-4-enuronosyl-(1->3)-N-acetyl-D-galactosamine + H+ + sulfate." [EC:3.1.6.9, RHEA:11444] |
| GO:0033888 | 6.07435E-05 | 5 / 5 | "Catalysis of the reaction: 4-deoxy-beta-D-gluc-4-enuronosyl-(1->3)-N-acetyl-D-galactosamine 6-sulfate + H2O = 4-deoxy-beta-D-gluc-4-enuronosyl-(1->3)-N-acetyl-D-galactosamine + H+ + sulfate." [EC:3.1.6.10, RHEA:10536] |
| GO:0033889 | 6.07435E-05 | 5 / 5 | "Catalysis of the hydrolysis of the 3-sulfate groups of the N-sulfo-D-glucosamine 3-O-sulfate units of heparin." [EC:3.1.6.15] |
| GO:0034889 | 6.07435E-05 | 5 / 5 | "Catalysis of the reaction: endosulfan hemisulfate + H2O = endosulfan monoalcohol + 2 H+ + sulfate." [UM-BBD_reactionID:r1384] |
| GO:0034902 | 6.07435E-05 | 5 / 5 | "Catalysis of the reaction: endosulfan sulfate + H2O = endosulfan diol + sulfite." [UM-BBD_reactionID:r1387] |
| GO:0043890 | 6.07435E-05 | 5 / 5 | "Catalysis of the hydrolysis of the 6-sulfate groups of the N-acetyl-D-galactosamine 6-sulfate units of chondroitin sulfate and of the D-galactose 6-sulfate units of keratan sulfate." [EC:3.1.6.4] |
| GO:0047753 | 6.07435E-05 | 5 / 5 | "Catalysis of the reaction: choline sulfate + H2O = choline + H+ + sulfate." [EC:3.1.6.6, RHEA:20820] |
| GO:0047825 | 6.07435E-05 | 5 / 5 | "Catalysis of the reaction: (R)-2-O-sulfolactate + H2O = (R)-lactate + H+ + sulfate." [EC:3.1.6.17, RHEA:20337] |
| GO:0047871 | 6.07435E-05 | 5 / 5 | "Catalysis of the reaction: N(2),6-disulfo-D-glucosamine + H2O = N-sulfo-D-glucosamine + H+ + sulfate." [EC:3.1.6.11, RHEA:15517] |
| GO:0047966 | 6.07435E-05 | 5 / 5 | "Catalysis of the reaction: D-glucose 6-sulfate + H2O = D-glucose + H+ + sulfate." [EC:3.1.6.3, RHEA:19145] |
| GO:0050106 | 6.07435E-05 | 5 / 5 | "Catalysis of the reaction: H2O + monomethyl sulfate = H+ + methanol + sulfate." [EC:3.1.6.16, RHEA:14221] |
| GO:0140110 | 8.66036E-05 | 498 / 3673 | "A molecular function that controls the rate, timing and/or magnitude of gene transcription. The function of transcriptional regulators is to modulate gene expression at the transcription step so that they are expressed in the right cell at the right time and in the right amount throughout the life of the cell |
| GO:0000099 | 0.000105337 | 8 / 13 | "Enables the transfer of sulfur amino acids from one side of a membrane to the other. Sulphur amino acids contain sulfur in the form of cystine, methionine or their derivatives." [GOC:ai, GOC:mtg_transport, ISBN:0815340729] |

| GO_Term | P_val | Counts | BioProc_Description |
|---|---|---|---|
| GO:0040007 | 1.75337E-10 | 471 / 3597 | "The increase in size or mass of an entire organism, a part of an organism or a cell." [GOC:bf, GOC:ma] |
| GO:0097080 | 2.89433E-10 | 102 / 401 | "The directed movement of inorganic selenite (HSeO3-1 at physiological pH) across a plasma membrane." [GOC:mcc, PMID:20861301] |
| GO:0055085 | 3.12194E-10 | 106 / 423 | "The process in which a solute is transported across a lipid bilayer, from one side of a membrane to the other." [GOC:dph, GOC:jid] |
| GO:0006855 | 3.36666E-10 | 102 / 402 | "The process in which a xenobiotic, a compound foreign to the organim exposed to it, is transported across a membrane. It may be synthesized by another organism (like ampicilin) or it can be a synthetic chemical." [GOC:ai, GOC:bf, GOC:krc] |
| GO:0015780 | 3.36666E-10 | 102 / 402 | "The directed movement of nucleotide-sugars into, out of or within a cell, or between cells, by means of some agent such as a transporter or pore. Nucleotide-sugars are any nucleotide in which the distal phosphoric residue of a nucleoside 5'-diphosphate is in glycosidic linkage with a monosaccharide c |
| GO:0034220 | 3.91261E-10 | 102 / 403 | "A process in which a monoatomic ion is transported across a membrane. Monatomic ions (also called simple ions) are ions consisting of exactly one atom." [GOC:mah] |
| GO:0046967 | 3.91261E-10 | 102 / 403 | "The directed movement of substances from the cytosol to the endoplasmic reticulum of a cell." [GOC:ai] |
| GO:1902047 | 3.91261E-10 | 102 / 403 | "The process in which a polyamine macromolecule is transported across a membrane." [GOC:TermGenie, GOC:vw] |
| GO:0006810 | 5.24851E-10 | 107 / 432 | "The directed movement of substances (such as macromolecules, small molecules, ions) or cellular components (such as complexes and organelles) into, out of or within a cell, or between cells, or within a multicellular organism by means of some agent such as a transporter or a transporter complex, a |
| GO:0001407 | 5.56962E-10 | 101 / 400 | "The process in which a glycerophosphodiester is transported across a membrane. Glycerophosphodiesters are small molecules composed of glycerol-3-phosphate and an alcohol, for example, glycerophosphoinositol." [GOC:mcc, PMID:12912892] |
| GO:0015794 | 5.56962E-10 | 101 / 400 | "The process in which glycerol-3-phosphate is transported across a membrane. Glycerol-3-phosphate is a phosphoric monoester of glycerol." [GOC:ai] |
| GO:0015897 | 5.56962E-10 | 101 / 400 | "The process in which an organomercurial compound is transported across a membrane. Organomercurial substances are any organic compound containing a mercury atom." [GOC:ai, PMID:18793329] |
| GO:0015904 | 5.56962E-10 | 101 / 400 | "The directed movement of tetracycline from one side of a membrane to the other. Tetracycline is a broad spectrum antibiotic that blocks binding of aminoacyl tRNA to the ribosomes of both Gram-positive and Gram-negative organisms (and those of organelles)." [GOC:curators] |
| GO:0015905 | 5.56962E-10 | 101 / 400 | "The directed movement of bicyclomycin across a lipid bilayer, from one side of a membrane to the other. Bicyclomycin (or bicozamycin) is an antibacterial drug often used as a livestock feed additive." [PMID:20067529] |
| GO:0035378 | 5.56962E-10 | 101 / 400 | "The process in which carbon dioxide (CO2) is transported across a membrane." [GOC:yaf] |
| GO:0035436 | 5.56962E-10 | 101 / 400 | "The process in which triose phosphate (glyceraldehyde 3-phosphate) is transported across a membrane. Glyceraldehyde 3-phosphate is any organic three carbon compound phosphate ester." [GOC:bf, ISBN:0198506732] |
| GO:0070811 | 5.56962E-10 | 101 / 400 | "The process in which glycerol-2-phosphate is transported across a membrane. Glycerol-2-phosphate is a phosphoric monoester of glycerol." [GOC:mah] |
| GO:0071918 | 5.56962E-10 | 101 / 400 | "The process in which urea, the water-soluble compound H2N-CO-NH2, is transported from one side of a membrane to the other by means of some agent such as a transporter or pore." [GOC:mah] |
| GO:0072489 | 5.56962E-10 | 101 / 400 | "The process in which methylammonium is transported across a membrane." [GOC:mah] |
| GO:0072531 | 5.56962E-10 | 101 / 400 | "The process in which a pyrimidine-containing compound is transported across a membrane. A pyrimidine-containing compound is any compound that contains pyrimidine or a formal derivative thereof." [GOC:mah] |
| GO:0080162 | 5.56962E-10 | 101 / 400 | "The directed movement of auxins from endoplasmic reticulum to cytosol." [PMID:19506555] |
| GO:0080170 | 5.56962E-10 | 101 / 400 | "The process in which hydrogen peroxide is transported across a membrane." [GOC:tb] |
| GO:0098660 | 5.56962E-10 | 101 / 400 | "The process in which an inorganic ion is transported across a membrane." [GOC:mah] |
| GO:0140317 | 5.56962E-10 | 101 / 400 | "The directed movement of a substance across the outer membrane in cells with two membranes." [PMID:15968039] |
| GO:0170006 | 5.56962E-10 | 101 / 400 | "The process in which lipoate is transported across a membrane." [PMID:20980265, PMID:25971966] |
| GO:1901642 | 5.56962E-10 | 101 / 400 | "The directed movement of nucleoside across a membrane." [GOC:pr, GOC:TermGenie] |
| GO:1901684 | 5.56962E-10 | 101 / 400 | "The process in which arsenate is transported across a membrane." [GOC:TermGenie] |
| GO:1901962 | 5.56962E-10 | 101 / 400 | "The directed movement of S-adenosyl-L-methionine across a membrane." [GOC:TermGenie, PMID:10497160] |
| GO:1902418 | 5.56962E-10 | 101 / 400 | "The process in which (+)-abscisic acid D-glucopyranosyl este is transported across a membrane." [GOC:TermGenie, PMID:24028845] |
| GO:1902603 | 5.56962E-10 | 101 / 400 | "The directed movement of carnitine across a membrane." [GO_REF:0000069, GOC:pr, GOC:TermGenie] |
| GO:1902616 | 5.56962E-10 | 101 / 400 | "The process in which acyl carnitine is transported across a membrane." [GO_REF:0000069, GOC:pr, GOC:TermGenie] |
| GO:1903825 | 5.56962E-10 | 101 / 400 | "The process in which an organic acid is transported across a membrane." [GO_REF:0000069, GOC:TermGenie] |
| GO:0045117 | 6.46065E-10 | 101 / 401 | "The directed movement of azoles, heterocyclic compounds found in many biologically important substances, across a lipid bilayer, across a membrane." [GOC:go_curators, ISBN:3527307206, Wikipedia:Azole] |
| GO:0071994 | 6.46065E-10 | 101 / 401 | "The process in which a phytochelatin is transported across a membrane. Phytochelatins are a group of peptides that bind metals (Cd, Zn, Cu, Pb, Hg) in thiolate coordination complexes." [GOC:mah, ISBN:0198506732] |
| GO:0140820 | 6.46065E-10 | 101 / 401 | "The directed movement of substances from the cytosol into the Golgi apparatus of a cell." [PMID:28373556] |
| GO:0015791 | 7.48765E-10 | 101 / 402 | "The directed movement of polyols, any polyhydric alcohol, across a membrane." [GOC:ai] |
| GO:0034219 | 7.48765E-10 | 101 / 402 | "The process in which a carbohydrate is transported across a membrane." [GOC:mah] |
| GO:0035382 | 7.48765E-10 | 101 / 402 | "The process in which a sterol is transported across a membrane. Sterols are steroids with one or more hydroxyl groups and a hydrocarbon side-chain in the molecule." [GOC:vw] |
| GO:0035461 | 7.48765E-10 | 101 / 402 | "The process in which a vitamin is transported across a membrane. A vitamin is one of a number of unrelated organic substances that occur in many foods in small amounts and that are necessary in trace amounts for the normal metabolic functioning of the body." [GOC:bf] |
| GO:0072530 | 7.48765E-10 | 101 / 402 | "The process in which a purine-containing compound is transported across a membrane. A purine-containing compound is any compound that contains purine or a formal derivative thereof." [GOC:mah] |
| GO:1901679 | 7.48765E-10 | 101 / 402 | "The directed movement of nucleotide across a membrane." [GOC:pr, GOC:TermGenie] |
| GO:0034486 | 8.67032E-10 | 101 / 403 | "The process in which a solute is transported from one side of the vacuolar membrane to the other." [GOC:mah] |
| GO:0046618 | 8.67032E-10 | 101 / 403 | "The directed movement of a xenobiotic from a cell, into the extracellular region. A xenobiotic is a compound foreign to the organim exposed to it. It may be synthesized by another organism (like ampicilin) or it can be a synthetic chemical." [GOC:go_curators, GOC:krc] |
| GO:0035672 | 1.00311E-09 | 101 / 404 | "The process in which an oligopeptide is transported across a membrane. Oligopeptides are molecules that contain a small number (2 to 20) of amino-acid residues connected by peptide linkages." [GOC:vw, ISBN:0198506732] |
| GO:0098739 | 1.00311E-09 | 101 / 404 | "The directed movement of some substance from outside of a cell, across the plasma membrane and into the cytosol." [GOC:dos] |
| GO:1990542 | 1.09333E-09 | 102 / 410 | "The process in which a solute is transported from one side of a membrane to the other into, out of or within a mitochondrion." [PMID:20533899] |
| GO:0008150 | 1.11014E-09 | 516 / 3820 | "A biological process is the execution of a genetically-encoded biological module or program. It consists of all the steps required to achieve the specific biological objective of the module. A biological process is accomplished by a particular set of molecular functions carried out by specific gene products ( |
| GO:0140115 | 1.3392E-09 | 101 / 406 | "The directed movement of some substance from inside of a cell, across the plasma membrane and into the extracellular region." [GOC:pg] |
| GO:0071806 | 1.45518E-09 | 102 / 412 | "The process in which a protein is transported across a membrane." [GOC:mah, GOC:vw] |
| GO:0002181 | 4.65012E-08 | 0 / 107 | "The chemical reactions and pathways resulting in the formation of a protein in the cytoplasm. This is a ribosome-mediated process in which the information in messenger RNA (mRNA) is used to specify the sequence of amino acids in the protein." [GOC:hjd] |

| GO_Term | P_val | Counts | CellComp_Description |
|---|---|---|---|
| GO:0031981 | 5.1247E-08 | 37 / 501 | "The volume enclosed by the nuclear inner membrane." [GOC:mah, GOC:pz] |
| GO:0043231 | 5.59399E-08 | 46 / 579 | "Organized structure of distinctive morphology and function, bounded by a single or double lipid bilayer membrane and occurring within the cell. Includes the nucleus, mitochondria, plastids, vacuoles, and vesicles. Excludes the plasma membrane." [GOC:go_curators] |
| GO:0140513 | 2.69169E-07 | 40 / 509 | "A stable assembly of two or more macromolecules, i.e. proteins, nucleic acids, carbohydrates or lipids, in which at least one component is a protein and the constituent parts function together in the nucleus." [GOC:pg] |
| GO:0000943 | 6.64442E-06 | 37 / 445 | "A complex of the retrotransposon RNA genome, reverse transcriptase, integrase, and associated molecules required for reproduction and integration of the retrotransposon into the host genome; the main structural molecule of the nucleocapsid is often a gag protein homolog." [GOC:clt, PMID:10861 |
| GO:0031039 | 6.64442E-06 | 37 / 445 | "A membrane-bounded organelle of ciliated protozoan cells that contains polyploid copies of a portion of the cell's complete genome. Transcription of genes occurs in macronuclei. Some ciliate species may contain multiple macronuclei per cell." [GOC:ns] |
| GO:0031040 | 6.64442E-06 | 37 / 445 | "A membrane-bounded organelle of ciliated protozoan cells that contains a diploid copy of the cell's complete genome. Sections of contiguous sequence in the macronucleus are often interrupted by internal eliminated sequences (IES), and may be permuted, in micronuclei. Genic transcription is not fou |
| GO:0042405 | 6.64442E-06 | 37 / 445 | "An intranuclear focus at which aggregated proteins have been sequestered." [GOC:jl] |
| GO:0043073 | 6.64442E-06 | 37 / 445 | "The nucleus of a germ cell, a reproductive cell in multicellular organisms." [CL:0000586, GOC:go_curators] |
| GO:0043076 | 6.64442E-06 | 37 / 445 | "The nucleus of a megasporocyte, a diploid cell that undergoes meiosis to produce four megaspores, and its descendents." [GOC:jl, ISBN:0618254153] |
| GO:0045120 | 6.64442E-06 | 37 / 445 | "The nucleus of either the ovum or the spermatozoon following fertilization. Thus, in the fertilized ovum, there are two pronuclei, one originating from the ovum, the other from the spermatozoon that brought about fertilization; they approach each other, but do not fuse until just before the first cleava |
| GO:0046818 | 6.64442E-06 | 37 / 445 | "A location in the host cell nucleus where viral proteins colocalize late in infection prior to the onset of viral DNA synthesis. More than one site can be present simultaneously." [PMID:10233976] |
| GO:0048353 | 6.64442E-06 | 37 / 445 | "Nucleus resulting from the fusion of the male gamete and two polar nuclei in the central cell of the embryo sac." [ISBN:0471245208] |
| GO:0048555 | 6.64442E-06 | 37 / 445 | "The nucleus of the generative cell, a cell contained within the pollen grain that will divide to produce two haploid sperm cells." [GOC:tair_curators] |
| GO:0048556 | 6.64442E-06 | 37 / 445 | "The nucleus of the microsporocyte. The microsporocyte is a diploid cell in which meiosis will occur, resulting in four microspores. A microspore is a spore that, in vascular plants, gives rise to a male gametophyte." [GOC:tair_curators, ISBN:047186840X] |
| GO:0097165 | 6.64442E-06 | 37 / 445 | "A dense aggregation in the nucleus composed of proteins and RNAs that appear when the cell is under stress." [GOC:ans, PMID:10359787, PMID:12865437] |
| GO:0097571 | 6.64442E-06 | 37 / 445 | "One of the two nuclei found in Giardia species (trophozoite stage). It is located on the left side of the cell when viewed from the dorsal side." [GOC:giardia, ISBN:0-444-81258-X] |
| GO:0097572 | 6.64442E-06 | 37 / 445 | "One of the two nuclei found in Giardia species (trophozoite stage). It is located on the right side of the cell when viewed from the dorsal side." [GOC:giardia, ISBN:0-444-81258-X] |
| GO:0110092 | 6.64442E-06 | 37 / 445 | "The area of a motile nucleus closest to the direction of movement." [GOC:al, GOC:mah, GOC:vw, PMID:15030757, PMID:24335254] |
| GO:0110093 | 6.64442E-06 | 37 / 445 | "The area of a motile nucleus furthest from the direction of movement." [GOC:kmv, PMID:24335254] |
| GO:0140510 | 6.64442E-06 | 37 / 445 | "A narrow constricted region of the nucleus that forms around the anaphase spindle during closed mitosis, and connects the main portions of the newly forming daughter nuclei." [PMID:32848252] |
| GO:1905754 | 6.64442E-06 | 37 / 445 | "Any nucleus that is part of a ascospore-type prospore." [GO_REF:0000064, GOC:TermGenie, PMID:26942678] |
| GO:0005759 | 9.35588E-06 | 6 / 152 | "The gel-like material, with considerable fine structure, that lies in the matrix space, or lumen, of a mitochondrion. It contains the enzymes of the tricarboxylic acid cycle and, in some organisms, the enzymes concerned with fatty acid oxidation." [GOC:as, ISBN:0198506732] |
| GO:0005634 | 9.49254E-06 | 38 / 449 | "A membrane-bounded organelle of eukaryotic cells in which chromosomes are housed and replicated. In most cells, the nucleus contains all of the cell's chromosomes except the organellar chromosomes, and is the site of RNA synthesis and processing. In some species, or in specialized cell types, RNA |
| GO:0005880 | 1.20116E-05 | 38 / 446 | "Any microtubule in the nucleus of a cell." [GOC:mah] |
| GO:0005635 | 1.68614E-05 | 39 / 450 | "The double lipid bilayer enclosing the nucleus and separating its contents from the rest of the cytoplasm; includes the intermembrane space, a gap of width 20-40 nm (also called the perinuclear space)." [ISBN:0198547684] |
| GO:0043232 | 6.35006E-05 | 1 / 76 | "Organized structure of distinctive morphology and function, not bounded by a lipid bilayer membrane and occurring within the cell. Includes ribosomes, the cytoskeleton and chromosomes." [GOC:go_curators] |
| GO:0008152 | 6.43587E-05 | 48 / 505 | "The chemical reactions and pathways, including anabolism and catabolism, by which living organisms transform chemical substances. Metabolic processes typically transform small molecules, but also include macromolecular processes such as DNA repair and replication, and protein synthesis and deg |
| GO:0008150 | 6.8647E-05 | 47 / 496 | "A biological process is the execution of a genetically-encoded biological module or program. It consists of all the steps required to achieve the specific biological objective of the module. A biological process is accomplished by a particular set of molecular functions carried out by specific gene products ( |
| GO:0005761 | 7.37142E-05 | 1 / 75 | "A ribosome found in the mitochondrion of a eukaryotic cell; contains a characteristic set of proteins distinct from those of cytosolic ribosomes." [GOC:mah, ISBN:0198506732] |
| GO:0009987 | 8.96506E-05 | 49 / 508 | "Any process that is carried out at the cellular level, but not necessarily restricted to a single cell. For example, cell communication occurs among more than one cell, but occurs at the cellular level." [GOC:go_curators, GOC:isa_complete] |
| GO:0040007 | 0.000132961 | 46 / 478 | "The increase in size or mass of an entire organism, a part of an organism or a cell." [GOC:bf, GOC:ma] |
| GO:0022414 | 0.000151546 | 45 / 468 | "A biological process that directly contributes to the process of producing new individuals by one or two organisms. The new individuals inherit some proportion of their genetic material from the parent or parents." [GOC:dph, GOC:isa_complete] |
| GO:0098754 | 0.000151546 | 45 / 468 | "Any process that reduces or removes the toxicity of a toxic substance. These may include transport of the toxic substance away from sensitive areas and to compartments or complexes whose purpose is sequestration of the toxic substance." [GOC:dos] |
| GO:0032991 | 0.000155681 | 659 / 4637 | "A stable assembly of two or more macromolecules, i.e. proteins, nucleic acids, carbohydrates or lipids, in which at least one component is a protein and the constituent parts function together." [GOC:dos, GOC:mah] |
| GO:0050896 | 0.000172541 | 45 / 466 | "Any process that results in a change in state or activity of a cell or an organism (in terms of movement, secretion, enzyme production, gene expression, etc.) as a result of a stimulus. The process begins with detection of the stimulus and ends with a change in state or activity or the cell or organism." [G |
| GO:0000003 | 0.000184032 | 45 / 465 | "The production of new individuals that contain some portion of genetic material inherited from one or more parent organisms." [GOC:go_curators, GOC:isa_complete, GOC:jl, ISBN:0198506732] |
| GO:0002376 | 0.000184032 | 45 / 465 | "Any process involved in the development or functioning of the immune system, an organismal system for calibrated responses to potential internal or invasive threats." [GO_REF:0000022, GOC:add] |
| GO:0016032 | 0.000184032 | 45 / 465 | "A multi-organism process in which a virus is a participant. The other participant is the host. Includes infection of a host cell, replication of the viral genome, and assembly of progeny virus particles. In some cases the viral genetic material may integrate into the host genome and only subsequently, unde |
| GO:0032501 | 0.000184032 | 45 / 465 | "Any biological process, occurring at the level of a multicellular organism, pertinent to its function." [GOC:curators, GOC:dph, GOC:isa_complete, GOC:tb] |
| GO:0032502 | 0.000184032 | 45 / 465 | "A biological process whose specific outcome is the progression of an integrated living unit: an anatomical structure (which may be a subcellular structure, cell, tissue, or organ), or organism over time from an initial condition to a later condition." [GOC:isa_complete] |
| GO:0040011 | 0.000184032 | 45 / 465 | "Self-propelled movement of a cell or organism from one location to another." [GOC:dgh] |
| GO:0042592 | 0.000184032 | 45 / 465 | "Any biological process involved in the maintenance of an internal steady state." [GOC:jl, ISBN:0395825172] |
| GO:0043473 | 0.000184032 | 45 / 465 | "The accumulation of pigment in an organism, tissue or cell, either by increased deposition or by increased number of cells." [GOC:jl] |
| GO:0044419 | 0.000184032 | 45 / 465 | "Any process evolved to enable an interaction with an organism of a different species." [GOC:cc] |
| GO:0044848 | 0.000184032 | 45 / 465 | "A distinct period or stage in a biological process or cycle." [GOC:jl] |
| GO:0048511 | 0.000184032 | 45 / 465 | "Any process pertinent to the generation and maintenance of rhythms in the physiology of an organism." [GOC:jid] |
| GO:0051179 | 0.000184032 | 45 / 465 | "Any process in which a cell, a substance, or a cellular entity, such as a protein complex or organelle, is transported, tethered to or otherwise maintained in a specific location. In the case of substances, localization may also be achieved via selective degradation." [GOC:ai, GOC:dos] |
| GO:0051703 | 0.000184032 | 45 / 465 | "Any process in which an organism has an effect on an organism of the same species." [GOC:ai] |
| GO:0065007 | 0.000184032 | 45 / 465 | "Any process that modulates a measurable attribute of any biological process, quality or function." [GOC:dph, GOC:isa_complete, GOC:mah, GOC:pr, GOC:vw] |
| GO:0030120 | 0.00020975 | 6 / 8 | "A membrane coat found on a coated vesicle." [GOC:mah] |

#### Supplementary Data S3.4.11

| GO_Term | P_val | Counts | MolFunc_Description |
|---|---|---|---|
| GO:0003676 | 1.03615E-09 | 45 / 531 | "Binding to a nucleic acid." [GOC:jl] |
| GO:0003677 | 5.45572E-08 | 49 / 524 | "Any molecular function by which a gene product interacts selectively and non-covalently with DNA (deoxyribonucleic acid)." [GOC:dph, GOC:jl, GOC:tb, GOC:vw] |
| GO:0005198 | 5.82686E-07 | 593 / 374 | "The action of a molecule that contributes to the structural integrity of a complex." [GOC:mah, GOC:vw] |
| GO:0003735 | 1.46702E-06 | 4 / 123 | "The action of a molecule that contributes to the structural integrity of the ribosome." [GOC:mah] |
| GO:0031490 | 1.56297E-06 | 27 / 324 | "Binding to DNA that is assembled into chromatin." [GOC:mah] |
| GO:0003697 | 9.86296E-06 | 28 / 314 | "Binding to single-stranded DNA." [GOC:elh, GOC:vw, PMID:22976174] |
| GO:0008270 | 1.81235E-05 | 53 / 489 | "Binding to a zinc ion (Zn)." [GOC:ai] |
| GO:0140657 | 3.47334E-05 | 604 / 375 | "A molecular function characterized by the coupling of ATP hydrolysis to other steps of a reaction mechanism to make the reaction energetically favorable, for example to catalyze a reaction or drive transport against a concentration gradient." [PMID:24878343, PMID:25750732, PMID:32933017, PMID |
| GO:0019843 | 3.68928E-05 | 5 / 112 | "Binding to a ribosomal RNA." [GOC:jl] |
| GO:0000217 | 3.68946E-05 | 27 / 293 | "Binding to a DNA secondary structure element such as a four-way junction, a bubble, a loop, Y-form DNA, or a double-strand/single-strand junction." [GOC:krc] |
| GO:0000981 | 3.79418E-05 | 24 / 270 | "A DNA-binding transcription factor activity that modulates the transcription of specific gene sets transcribed by RNA polymerase II." [GOC:txnOH-2018] |
| GO:0000099 | 3.83164E-05 | 9 / 13 | "Enables the transfer of sulfur amino acids from one side of a membrane to the other. Sulphur amino acids contain sulfur in the form of cystine, methionine or their derivatives." [GOC:ai, GOC:mtg_transport, ISBN:0815340729] |
| GO:0003690 | 4.20271E-05 | 30 / 314 | "Binding to double-stranded DNA." [GOC:elh, GOC:vw] |
| GO:0008301 | 4.46845E-05 | 27 / 291 | "The activity of binding selectively and non-covalently to and distorting the original structure of DNA, typically a straight helix, into a bend, or increasing the bend if the original structure was intrinsically bent due to its sequence." [GOC:krc, GOC:vw, PMID:10710711, PMID:19037758] |
| GO:0045027 | 4.46845E-05 | 27 / 291 | "Binding to DNA ends exposed by the creation of double-strand breaks (DSBs)." [GOC:jl] |
| GO:0070336 | 4.46845E-05 | 27 / 291 | "Binding to a flap structure in DNA. A DNA flap structure is one in which a single-stranded length of DNA or RNA protrudes from a double-stranded DNA molecule." [GOC:mah, PMID:15189154] |
| GO:0000497 | 4.91464E-05 | 27 / 290 | "Binding to nucleic acid via hydrogen bonds between the bases of a gene product molecule and the bases of a target DNA molecule." [GOC:krc] |
| GO:0003681 | 4.91464E-05 | 27 / 290 | "Binding to DNA in a bent conformation." [GOC:jl, PMID:12627977] |
| GO:0010844 | 4.91464E-05 | 27 / 290 | "Binding to a genomic region which promotes recombination." [GOC:dph, GOC:tb] |
| GO:0045142 | 4.91464E-05 | 27 / 290 | "Binding to a DNA triple helix. The formation of triple helical DNA has been evoked in several cellular processes including transcription, replication, and recombination." [PMID:10681538] |
| GO:0051880 | 4.91464E-05 | 27 / 290 | "Binding to G-quadruplex DNA structures, in which groups of four guanines adopt a flat, cyclic Hoogsteen hydrogen-bonding arrangement known as a guanine tetrad. The stacking of guanine tetrads results in G-quadruplex DNA structures. G-quadruplex DNA can form under physiological conditions fron |
| GO:0099077 | 4.91464E-05 | 27 / 290 | "DNA-binding activity that is dependent on binding to a histone." [PMID:11691835] |
| GO:0043565 | 5.05423E-05 | 30 / 312 | "Binding to DNA of a specific nucleotide composition, e.g. GC-rich DNA binding, or with a specific sequence motif or type of DNA e.g. promotor binding or rDNA binding." [GOC:jl] |
| GO:0003700 | 6.20643E-05 | 24 / 265 | "A transcription regulator activity that modulates transcription of gene sets via selective and non-covalent binding to a specific double-stranded genomic DNA sequence (sometimes referred to as a motif) within a cis-regulatory region. Regulatory regions include promoters (proximal and distal) and enhar |
| GO:0003723 | 7.51866E-05 | 35 / 344 | "Binding to an RNA molecule or a portion thereof." [GOC:jl, GOC:mah] |
| GO:0030515 | 0.000100211 | 5 / 105 | "Binding to a small nucleolar RNA." [GOC:mah] |
| GO:0015179 | 0.000100918 | 10 / 17 | "Enables the transfer of an L-amino acid from one side of a membrane to the other. L-amino acids are the L-enantiomers of amino acids." [GOC:ai, GOC:jsg, GOC:mah, GOC:mtg_transport, ISBN:0815340729] |
| GO:0016709 | 0.000100918 | 10 / 17 | "Catalysis of an oxidation-reduction (redox) reaction in which hydrogen or electrons are transferred from NADH or NADPH and one other donor, and one atom of oxygen is incorporated into one donor." [GOC:mah] |
| GO:0003684 | 0.000104053 | 30 / 304 | "Binding to damaged DNA." [GOC:jl] |
| GO:0140110 | 0.000117037 | 592 / 367 | "A molecular function that controls the rate, timing and/or magnitude of gene transcription. The function of transcriptional regulators is to modulate gene expression at the transcription step so that they are expressed in the right cell at the right time and in the right amount throughout the life of the cell a |
| GO:0003943 | 0.000139005 | 5 / 5 | "Catalysis of the hydrolysis of the 4-sulfate groups of the N-acetyl-D-galactosamine 4-sulfate units of chondroitin sulfate and dermatan sulfate." [EC:3.1.6.12] |
| GO:0004065 | 0.000139005 | 5 / 5 | "Catalysis of the reaction: a phenol sulfate + H2O = a phenol + sulfate." [EC:3.1.6.1] |
| GO:0004098 | 0.000139005 | 5 / 5 | "Catalysis of the reaction: a cerebroside 3-sulfate + H2O = a cerebroside + sulfate." [EC:3.1.6.8] |
| GO:0004423 | 0.000139005 | 5 / 5 | "Catalysis of the hydrolysis of the 2-sulfate groups of the L-iduronate 2-sulfate units of dermatan sulfate, heparan sulfate and heparin." [EC:3.1.6.13] |
| GO:0004773 | 0.000139005 | 5 / 5 | "Catalysis of the reaction: 3-beta-hydroxyandrost-5-en-17-one 3-sulfate + H2O = 3-beta-hydroxyandrost-5-en-17-one + sulfate." [EC:3.1.6.2] |
| GO:0008449 | 0.000139005 | 5 / 5 | "Catalysis of the hydrolysis of the 6-sulfate group of the N-acetyl-D-glucosamine 6-sulfate units of heparan sulfate and keratan sulfate." [EC:3.1.6.14] |
| GO:0015024 | 0.000139005 | 5 / 5 | "Catalysis of the hydrolysis of the 2-sulfate groups of the 2-O-sulfo-D-glucuronate residues of chondroitin sulfate, heparin and heparitin sulfate." [EC:3.1.6.18] |
| GO:0018741 | 0.000139005 | 5 / 5 | "Catalysis of the reaction: a primary linear alkyl sulfate ester + H2O =>a primary alcohol + H(+) + sulfate." [RHEA:67908, UM-BBD_reactionID:r0602] |
| GO:0033886 | 0.000139005 | 5 / 5 | "Catalysis of the hydrolysis of the 2- and 3-sulfate groups of the polysulfates of cellulose and charonin." [EC:3.1.6.7] |
| GO:0033887 | 0.000139005 | 5 / 5 | "Catalysis of the reaction: 4-deoxy-beta-D-gluc-4-enuronosyl-(1->3)-N-acetyl-D-galactosamine 4-sulfate + H2O = 4-deoxy-beta-D-gluc-4-enuronosyl-(1->3)-N-acetyl-D-galactosamine + H+ + sulfate." [EC:3.1.6.9, RHEA:11444] |
| GO:0033888 | 0.000139005 | 5 / 5 | "Catalysis of the reaction: 4-deoxy-beta-D-gluc-4-enuronosyl-(1->3)-N-acetyl-D-galactosamine 6-sulfate + H2O = 4-deoxy-beta-D-gluc-4-enuronosyl-(1->3)-N-acetyl-D-galactosamine + H+ + sulfate." [EC:3.1.6.10, RHEA:10536] |
| GO:0033889 | 0.000139005 | 5 / 5 | "Catalysis of the hydrolysis of the 3-sulfate groups of the N-sulfo-D-glucosamine 3-O-sulfate units of heparin." [EC:3.1.6.15] |
| GO:0034889 | 0.000139005 | 5 / 5 | "Catalysis of the reaction: endosulfan hemisulfate + H2O = endosulfan monoalcohol + 2 H+ + sulfate." [UM-BBD_reactionID:r1384] |
| GO:0034902 | 0.000139005 | 5 / 5 | "Catalysis of the reaction: endosulfan sulfate + H2O = endosulfan diol + sulfite." [UM-BBD_reactionID:r1387] |
| GO:0043890 | 0.000139005 | 5 / 5 | "Catalysis of the hydrolysis of the 6-sulfate groups of the N-acetyl-D-galactosamine 6-sulfate units of chondroitin sulfate and of the D-galactose 6-sulfate units of keratan sulfate." [EC:3.1.6.4] |
| GO:0047753 | 0.000139005 | 5 / 5 | "Catalysis of the reaction: choline sulfate + H2O = choline + H+ + sulfate." [EC:3.1.6.6, RHEA:20820] |
| GO:0047825 | 0.000139005 | 5 / 5 | "Catalysis of the reaction: (R)-2-O-sulfolactate + H2O = (R)-lactate + H+ + sulfate." [EC:3.1.6.17, RHEA:20337] |
| GO:0047871 | 0.000139005 | 5 / 5 | "Catalysis of the reaction: N(2),6-disulfo-D-glucosamine + H2O = N-sulfo-D-glucosamine + H+ + sulfate." [EC:3.1.6.11, RHEA:15517] |
| GO:0047966 | 0.000139005 | 5 / 5 | "Catalysis of the reaction: D-glucose 6-sulfate + H2O = D-glucose + H+ + sulfate." [EC:3.1.6.3, RHEA:19145] |
| GO:0050106 | 0.000139005 | 5 / 5 | "Catalysis of the reaction: H2O + monomethyl sulfate = H+ + methanol + sulfate." [EC:3.1.6.16, RHEA:14221] |

| GO_Term | P_val | Counts | BioProc_Description |
|---|---|---|---|
| GO:0040007 | 1.75592E-08 | 555 / 359 | "The increase in size or mass of an entire organism, a part of an organism or a cell." [GOC:bf, GOC:ma] |
| GO:0006414 | 3.65696E-08 | 0 / 93 | "The successive addition of amino acid residues to a nascent polypeptide chain during protein biosynthesis." [GOC:ems] |
| GO:0032543 | 4.40659E-08 | 0 / 92 | "The chemical reactions and pathways resulting in the formation of a protein in a mitochondrion. This is a ribosome-mediated process in which the information in messenger RNA (mRNA) is used to specify the sequence of amino acids in the protein; the mitochondrion has its own ribosomes and transfer |
| GO:0002181 | 5.88792E-08 | 1 / 107 | "The chemical reactions and pathways resulting in the formation of a protein in the cytoplasm. This is a ribosome-mediated process in which the information in messenger RNA (mRNA) is used to specify the sequence of amino acids in the protein." [GOC:hjd] |
| GO:0030950 | 1.09277E-07 | 12 / 15 | "Any cellular process that results in the specification, formation or maintenance of polarized actin-based cytoskeletal structures." [GOC:mah] |
| GO:0032544 | 3.41532E-07 | 0 / 81 | "The chemical reactions and pathways resulting in the formation of a protein in a plastid. This is a ribosome-mediated process in which the information in messenger RNA (mRNA) is used to specify the sequence of amino acids in the protein; the plastid has its own ribosomes and transfer RNAs, and uses a |
| GO:0071891 | 3.41532E-07 | 0 / 81 | "An N-terminal peptidyl-proline dimethylation process that contributes to translation." [GOC:mah] |
| GO:0140241 | 3.41532E-07 | 0 / 81 | "Translation that occurs at the synapse." [PMID:23083742] |
| GO:0097080 | 3.52035E-07 | 103 / 401 | "The directed movement of inorganic selenite (HSeO3-1 at physiological pH) across a plasma membrane." [GOC:mcc, PMID:20861301] |
| GO:0006855 | 3.99322E-07 | 103 / 402 | "The process in which a xenobiotic, a compound foreign to the organim exposed to it, is transported across a membrane. It may be synthesized by another organism (like ampicilin) or it can be a synthetic chemical." [GOC:ai, GOC:bf, GOC:krc] |
| GO:0015780 | 3.99322E-07 | 103 / 402 | "The directed movement of nucleotide-sugars into, out of or within a cell, or between cells, by means of some agent such as a transporter or pore. Nucleotide-sugars are any nucleotide in which the distal phosphoric residue of a nucleoside 5'-diphosphate is in glycosidic linkage with a monosaccharide or |
| GO:0006810 | 4.05463E-07 | 109 / 432 | "The directed movement of substances (such as macromolecules, small molecules, ions) or cellular components (such as complexes and organelles) into, out of or within a cell, or between cells, or within a multicellular organism by means of some agent such as a transporter or a transporter complex, a p |
| GO:0006259 | 4.11559E-07 | 1 / 96 | "Any cellular metabolic process involving deoxyribonucleic acid. This is one of the two main types of nucleic acid, consisting of a long, unbranched macromolecule formed from one, or more commonly, two, strands of linked deoxyribonucleotides." [ISBN:0198506732] |
| GO:0009059 | 4.11559E-07 | 1 / 96 | "The chemical reactions and pathways resulting in the formation of a macromolecule, any molecule of high relative molecular mass, the structure of which essentially comprises the multiple repetition of units derived, actually or conceptually, from molecules of low relative molecular mass." [GOC:mah] |
| GO:0034220 | 4.52552E-07 | 103 / 403 | "A process in which a monoatomic ion is transported across a membrane. Monatomic ions (also called simple ions) are ions consisting of exactly one atom." [GOC:mah] |
| GO:0034486 | 4.52552E-07 | 103 / 403 | "The process in which a solute is transported from one side of the vacuolar membrane to the other." [GOC:mah] |
| GO:0046967 | 4.52552E-07 | 103 / 403 | "The directed movement of substances from the cytosol to the endoplasmic reticulum of a cell." [GOC:ai] |
| GO:1902047 | 4.52552E-07 | 103 / 403 | "The process in which a polyamine macromolecule is transported across a membrane." [GOC:TermGenie, GOC:vw] |
| GO:0055085 | 4.5746E-07 | 107 / 423 | "The process in which a solute is transported across a lipid bilayer, from one side of a membrane to the other." [GOC:dph, GOC:jid] |
| GO:0001407 | 5.77632E-07 | 102 / 40C | "The process in which a glycerophosphodiester is transported across a membrane. Glycerophosphodiesters are small molecules composed of glycerol-3-phosphate and an alcohol, for example, glycerophosphoinositol." [GOC:mcc, PMID:12912892] |
| GO:0015794 | 5.77632E-07 | 102 / 40C | "The process in which glycerol-3-phosphate is transported across a membrane. Glycerol-3-phosphate is a phosphoric monoester of glycerol." [GOC:ai] |
| GO:0015897 | 5.77632E-07 | 102 / 40C | "The process in which an organomercurial compound is transported across a membrane. Organomercurial substances are any organic compound containing a mercury atom." [GOC:ai, PMID:18793329] |
| GO:0015904 | 5.77632E-07 | 102 / 40C | "The directed movement of tetracycline from one side of a membrane to the other. Tetracycline is a broad spectrum antibiotic that blocks binding of aminoacyl tRNA to the ribosomes of both Gram-positive and Gram-negative organisms (and those of organelles)." [GOC:curators] |
| GO:0015905 | 5.77632E-07 | 102 / 40C | "The directed movement of bicyclomycin across a lipid bilayer, from one side of a membrane to the other. Bicyclomycin (or bicozamycin) is an antibacterial drug often used as a livestock feed additive." [PMID:20067529] |
| GO:0035378 | 5.77632E-07 | 102 / 40C | "The process in which carbon dioxide (CO2) is transported across a membrane." [GOC:yaf] |
| GO:0035436 | 5.77632E-07 | 102 / 40C | "The process in which triose phosphate (glyceraldehyde 3-phosphate) is transported across a membrane. Glyceraldehyde 3-phosphate is any organic three carbon compound phosphate ester." [GOC:bf, ISBN:0198506732] |
| GO:0070811 | 5.77632E-07 | 102 / 40C | "The process in which glycerol-2-phosphate is transported across a membrane. Glycerol-2-phosphate is a phosphoric monoester of glycerol." [GOC:mah] |
| GO:0071918 | 5.77632E-07 | 102 / 40C | "The process in which urea, the water-soluble compound H2N-CO-NH2, is transported from one side of a membrane to the other by means of some agent such as a transporter or pore." [GOC:mah] |
| GO:0072489 | 5.77632E-07 | 102 / 40C | "The process in which methylammonium is transported across a membrane." [GOC:mah] |
| GO:0072531 | 5.77632E-07 | 102 / 40C | "The process in which a pyrimidine-containing compound is transported across a membrane. A pyrimidine-containing compound is any compound that contains pyrimidine or a formal derivative thereof." [GOC:mah] |
| GO:0080162 | 5.77632E-07 | 102 / 40C | "The directed movement of auxins from endoplasmic reticulum to cytosol." [PMID:19506555] |
| GO:0080170 | 5.77632E-07 | 102 / 40C | "The process in which hydrogen peroxide is transported across a membrane." [GOC:tb] |
| GO:0098660 | 5.77632E-07 | 102 / 40C | "The process in which an inorganic ion is transported across a membrane." [GOC:mah] |
| GO:0140317 | 5.77632E-07 | 102 / 40C | "The directed movement of a substance across the outer membrane in cells with two membranes." [PMID:15968039] |
| GO:0170006 | 5.77632E-07 | 102 / 40C | "The process in which lipoate is transported across a membrane." [PMID:20980265, PMID:25971966] |
| GO:1901642 | 5.77632E-07 | 102 / 40C | "The directed movement of nucleoside across a membrane." [GOC:pr, GOC:TermGenie] |
| GO:1901684 | 5.77632E-07 | 102 / 40C | "The process in which arsenate is transported across a membrane." [GOC:TermGenie] |
| GO:1901962 | 5.77632E-07 | 102 / 40C | "The directed movement of S-adenosyl-L-methionine across a membrane." [GOC:TermGenie, PMID:10497160] |
| GO:1902418 | 5.77632E-07 | 102 / 40C | "The process in which (+)-abscisic acid D-glucopyranosyl este is transported across a membrane." [GOC:TermGenie, PMID:24028845] |
| GO:1902603 | 5.77632E-07 | 102 / 40C | "The directed movement of carnitine across a membrane." [GO_REF:0000069, GOC:pr, GOC:TermGenie] |
| GO:1902616 | 5.77632E-07 | 102 / 40C | "The process in which acyl carnitine is transported across a membrane." [GO_REF:0000069, GOC:pr, GOC:TermGenie] |
| GO:1903825 | 5.77632E-07 | 102 / 40C | "The process in which an organic acid is transported across a membrane." [GO_REF:0000069, GOC:TermGenie] |
| GO:0098754 | 6.14011E-07 | 547 / 352 | "Any process that reduces or removes the toxicity of a toxic substance. These may include transport of the toxic substance away from sensitive areas and to compartments or complexes whose purpose is sequestration of the toxic substance." [GOC:dos] |
| GO:0022414 | 6.23259E-07 | 548 / 353 | "A biological process that directly contributes to the process of producing new individuals by one or two organisms. The new individuals inherit some proportion of their genetic material from the parent or parents." [GOC:dph, GOC:isa_complete] |
| GO:0045117 | 6.53414E-07 | 102 / 401 | "The directed movement of azoles, heterocyclic compounds found in many biologically important substances, across a lipid bilayer, across a membrane." [GOC:go_curators, ISBN:3527307206, Wikipedia:Azole] |
| GO:0071994 | 6.53414E-07 | 102 / 401 | "The process in which a phytochelatin is transported across a membrane. Phytochelatins are a group of peptides that bind metals (Cd, Zn, Cu, Pb, Hg) in thiolate coordination complexes." [GOC:mah, ISBN:0198506732] |
| GO:0140820 | 6.53414E-07 | 102 / 401 | "The directed movement of substances from the cytosol into the Golgi apparatus of a cell." [PMID:28373556] |
| GO:0015791 | 7.38474E-07 | 102 / 402 | "The directed movement of polyols, any polyhydric alcohol, across a membrane." [GOC:ai] |
| GO:0034219 | 7.38474E-07 | 102 / 402 | "The process in which a carbohydrate is transported across a membrane." [GOC:mah] |
| GO:0035382 | 7.38474E-07 | 102 / 402 | "The process in which a sterol is transported across a membrane. Sterols are steroids with one or more hydroxyl groups and a hydrocarbon side-chain in the molecule." [GOC:vw] |

| GO_Term | P_val | Counts | CellComp_Description |
|---|---|---|---|
| GO:0043231 | 8.11638E-08 | 57 / 579 | "Organized structure of distinctive morphology and function, bounded by a single or double lipid bilayer membrane and occurring within the cell. Includes the nucleus, mitochondria, plastids, vacuoles, and vesicles. Excludes the plasma membrane." [GOC:go_curators] |
| GO:0031981 | 5.0711E-07 | 49 / 501 | "The volume enclosed by the nuclear inner membrane." [GOC:mah, GOC:pz] |
| GO:0140513 | 3.30889E-06 | 53 / 509 | "A stable assembly of two or more macromolecules, i.e. proteins, nucleic acids, carbohydrates or lipids, in which at least one component is a protein and the constituent parts function together in the nucleus." [GOC:pg] |
| GO:0005759 | 2.33485E-05 | 9 / 152 | "The gel-like material, with considerable fine structure, that lies in the matrix space, or lumen, of a mitochondrion. It contains the enzymes of the tricarboxylic acid cycle and, in some organisms, the enzymes concerned with fatty acid oxidation." [GOC:as, ISBN:0198506732] |
| GO:0000943 | 6.12555E-05 | 49 / 445 | "A complex of the retrotransposon RNA genome, reverse transcriptase, integrase, and associated molecules required for reproduction and integration of the retrotransposon into the host genome; the main structural molecule of the nucleocapsid is often a gag protein homolog." [GOC:clt, PMID:108619( |
| GO:0031039 | 6.12555E-05 | 49 / 445 | "A membrane-bounded organelle of ciliated protozoan cells that contains polyploid copies of a portion of the cell's complete genome. Transcription of genes occurs in macronuclei. Some ciliate species may contain multiple macronuclei per cell." [GOC:ns] |
| GO:0031040 | 6.12555E-05 | 49 / 445 | "A membrane-bounded organelle of ciliated protozoan cells that contains a diploid copy of the cell's complete genome. Sections of contiguous sequence in the macronucleus are often interrupted by internal eliminated sequences (IES), and may be permuted, in micronuclei. Genic transcription is not foun |
| GO:0042405 | 6.12555E-05 | 49 / 445 | "An intranuclear focus at which aggregated proteins have been sequestered." [GOC:jl] |
| GO:0043073 | 6.12555E-05 | 49 / 445 | "The nucleus of a germ cell, a reproductive cell in multicellular organisms." [CL:0000586, GOC:go_curators] |
| GO:0043076 | 6.12555E-05 | 49 / 445 | "The nucleus of a megasporocyte, a diploid cell that undergoes meiosis to produce four megaspores, and its descendents." [GOC:jl, ISBN:0618254153] |
| GO:0045120 | 6.12555E-05 | 49 / 445 | "The nucleus of either the ovum or the spermatozoon following fertilization. Thus, in the fertilized ovum, there are two pronuclei, one originating from the ovum, the other from the spermatozoon that brought about fertilization; they approach each other, but do not fuse until just before the first cleavag |
| GO:0046818 | 6.12555E-05 | 49 / 445 | "A location in the host cell nucleus where viral proteins colocalize late in infection prior to the onset of viral DNA synthesis. More than one site can be present simultaneously." [PMID:10233976] |
| GO:0048353 | 6.12555E-05 | 49 / 445 | "Nucleus resulting from the fusion of the male gamete and two polar nuclei in the central cell of the embryo sac." [ISBN:0471245208] |
| GO:0048555 | 6.12555E-05 | 49 / 445 | "The nucleus of the generative cell, a cell contained within the pollen grain that will divide to produce two haploid sperm cells." [GOC:tair_curators] |
| GO:0048556 | 6.12555E-05 | 49 / 445 | "The nucleus of the microsporocyte. The microsporocyte is a diploid cell in which meiosis will occur, resulting in four microspores. A microspore is a spore that, in vascular plants, gives rise to a male gametophyte." [GOC:tair_curators, ISBN:047186840X] |
| GO:0097165 | 6.12555E-05 | 49 / 445 | "A dense aggregation in the nucleus composed of proteins and RNAs that appear when the cell is under stress." [GOC:ans, PMID:10359787, PMID:12865437] |
| GO:0097571 | 6.12555E-05 | 49 / 445 | "One of the two nuclei found in Giardia species (trophozoite stage). It is located on the left side of the cell when viewed from the dorsal side." [GOC:giardia, ISBN:0-444-81258-X] |
| GO:0097572 | 6.12555E-05 | 49 / 445 | "One of the two nuclei found in Giardia species (trophozoite stage). It is located on the right side of the cell when viewed from the dorsal side." [GOC:giardia, ISBN:0-444-81258-X] |
| GO:0110092 | 6.12555E-05 | 49 / 445 | "The area of a motile nucleus closest to the direction of movement." [GOC:al, GOC:mah, GOC:vw, PMID:15030757, PMID:24335254] |
| GO:0110093 | 6.12555E-05 | 49 / 445 | "The area of a motile nucleus furthest from the direction of movement." [GOC:kmv, PMID:24335254] |
| GO:0140510 | 6.12555E-05 | 49 / 445 | "A narrow constricted region of the nucleus that forms around the anaphase spindle during closed mitosis, and connects the main portions of the newly forming daughter nuclei." [PMID:32848252] |
| GO:1905754 | 6.12555E-05 | 49 / 445 | "Any nucleus that is part of a ascospore-type prospore." [GO_REF:0000064, GOC:TermGenie, PMID:26942678] |
| GO:0043232 | 7.55633E-05 | 2 / 76 | "Organized structure of distinctive morphology and function, not bounded by a lipid bilayer membrane and occurring within the cell. Includes ribosomes, the cytoskeleton and chromosomes." [GOC:go_curators] |
| GO:0005634 | 7.67917E-05 | 50 / 449 | "A membrane-bounded organelle of eukaryotic cells in which chromosomes are housed and replicated. In most cells, the nucleus contains all of the cell's chromosomes except the organellar chromosomes, and is the site of RNA synthesis and processing. In some species, or in specialized cell types, RNA r |
| GO:0000313 | 8.34774E-05 | 0 / 50 | "A ribosome contained within a subcellular membrane-bounded organelle." [GOC:mah, GOC:mcc] |
| GO:0005880 | 9.62322E-05 | 50 / 446 | "Any microtubule in the nucleus of a cell." [GOC:mah] |
| GO:0005635 | 0.000119292 | 51 / 450 | "The double lipid bilayer enclosing the nucleus and separating its contents from the rest of the cytoplasm; includes the intermembrane space, a gap of width 20-40 nm (also called the perinuclear space)." [ISBN:0198547684] |
| GO:0098798 | 0.000126501 | 5 / 103 | "A protein complex that is part of a mitochondrion." [GOC:dos] |
| GO:0008152 | 0.0001858 | 60 / 505 | "The chemical reactions and pathways, including anabolism and catabolism, by which living organisms transform chemical substances. Metabolic processes typically transform small molecules, but also include macromolecular processes such as DNA repair and replication, and protein synthesis and degr |
| GO:0030874 | 0.00021448 | 0 / 45 | "The portion of nuclear chromatin associated with the nucleolus; includes the DNA encoding the ribosomal RNA." [GOC:mah] |
| GO:0034388 | 0.00021448 | 0 / 45 | "A protein complex that forms a subcomplex of the 90S preribosome and can interact directly with the 5' External Transcribed Spacer (ETS) of the full length pre-rRNA transcript. In S. cerevisiae, it sediments at 25-30 S and is composed of Pwp2p, Dip2p, Utp21p, Utp13p, Utp18p, and Utp6p." [GOC:krc, PI |
| GO:0070545 | 0.00021448 | 0 / 45 | "A protein complex that is involved in coordinating ribosome biogenesis with cell cycle progression. In human, it is composed of Pes1, Bop1, and WDR12; in Saccharomyces the proteins are known as Nop7p, Erb1 and Ytm1 respectively." [GOC:ab, GOC:mah, PMID:16043514, PMID:17353269] |
| GO:0008150 | 0.000216181 | 59 / 496 | "A biological process is the execution of a genetically-encoded biological module or program. It consists of all the steps required to achieve the specific biological objective of the module. A biological process is accomplished by a particular set of molecular functions carried out by specific gene products (o |
| GO:0022414 | 0.000230776 | 55 / 468 | "A biological process that directly contributes to the process of producing new individuals by one or two organisms. The new individuals inherit some proportion of their genetic material from the parent or parents." [GOC:dph, GOC:isa_complete] |
| GO:0098754 | 0.000230776 | 55 / 468 | "Any process that reduces or removes the toxicity of a toxic substance. These may include transport of the toxic substance away from sensitive areas and to compartments or complexes whose purpose is sequestration of the toxic substance." [GOC:dos] |
| GO:0009987 | 0.00023805 | 61 / 508 | "Any process that is carried out at the cellular level, but not necessarily restricted to a single cell. For example, cell communication occurs among more than one cell, but occurs at the cellular level." [GOC:go_curators, GOC:isa_complete] |
| GO:0005740 | 0.000255609 | 6 / 107 | "The double lipid bilayer enclosing the mitochondrion and separating its contents from the cell cytoplasm; includes the intermembrane space." [GOC:ai, GOC:pz] |
| GO:0034456 | 0.000258998 | 0 / 44 | "A protein complex that forms a subcomplex of the 90S preribosome. In S. cerevisiae, it is composed of Rrp7p, Utp22p, Ckb1p, Cka1p, Ckb2p and Cka2p." [GOC:mah, PMID:17515605] |
| GO:0034457 | 0.000258998 | 0 / 44 | "A protein complex that forms a subcomplex of the 90S preribosome. In S. cerevisiae, it is composed of Mpp10p, Imp3p and Imp4p." [GOC:mah, PMID:17515605] |
| GO:0050896 | 0.000264393 | 55 / 466 | "Any process that results in a change in state or activity of a cell or an organism (in terms of movement, secretion, enzyme production, gene expression, etc.) as a result of a stimulus. The process begins with detection of the stimulus and ends with a change in state or activity or the cell or organism." [GO |
| GO:0000003 | 0.000282855 | 55 / 465 | "The production of new individuals that contain some portion of genetic material inherited from one or more parent organisms." [GOC:go_curators, GOC:isa_complete, GOC:jl, ISBN:0198506732] |
| GO:0002376 | 0.000282855 | 55 / 465 | "Any process involved in the development or functioning of the immune system, an organismal system for calibrated responses to potential internal or invasive threats." [GO_REF:0000022, GOC:add] |
| GO:0016032 | 0.000282855 | 55 / 465 | "A multi-organism process in which a virus is a participant. The other participant is the host. Includes infection of a host cell, replication of the viral genome, and assembly of progeny virus particles. In some cases the viral genetic material may integrate into the host genome and only subsequently, under |
| GO:0032501 | 0.000282855 | 55 / 465 | "Any biological process, occurring at the level of a multicellular organism, pertinent to its function." [GOC:curators, GOC:dph, GOC:isa_complete, GOC:tb] |
| GO:0032502 | 0.000282855 | 55 / 465 | "A biological process whose specific outcome is the progression of an integrated living unit: an anatomical structure (which may be a subcellular structure, cell, tissue, or organ), or organism over time from an initial condition to a later condition." [GOC:isa_complete] |
| GO:0040011 | 0.000282855 | 55 / 465 | "Self-propelled movement of a cell or organism from one location to another." [GOC:dgh] |
| GO:0042592 | 0.000282855 | 55 / 465 | "Any biological process involved in the maintenance of an internal steady state." [GOC:jl, ISBN:0395825172] |
| GO:0043473 | 0.000282855 | 55 / 465 | "The accumulation of pigment in an organism, tissue or cell, either by increased deposition or by increased number of cells." [GOC:jl] |
| GO:0044419 | 0.000282855 | 55 / 465 | "Any process evolved to enable an interaction with an organism of a different species." [GOC:cc] |
| GO:0044848 | 0.000282855 | 55 / 465 | "A distinct period or stage in a biological process or cycle." [GOC:jl] |

#### Supplementary Data S3.4.12

| GO_Term | P_val | Counts | MolFunc_Description |
|---|---|---|---|
| GO:0003676 | 3.00908E-11 | 75 / 531 | "Binding to a nucleic acid." [GOC:jl] |
| GO:0003677 | 1.89051E-10 | 76 / 524 | "Any molecular function by which a gene product interacts selectively and non-covalently with DNA (deoxyribonucleic acid)." [GOC:dph, GOC:jl, GOC:tb, GOC:vw] |
| GO:0031490 | 7.01903E-08 | 44 / 324 | "Binding to DNA that is assembled into chromatin." [GOC:mah] |
| GO:0003690 | 6.68764E-07 | 45 / 314 | "Binding to double-stranded DNA." [GOC:elh, GOC:vw] |
| GO:0003697 | 6.68764E-07 | 45 / 314 | "Binding to single-stranded DNA." [GOC:elh, GOC:vw, PMID:22976174] |
| GO:0043565 | 8.91849E-07 | 45 / 312 | "Binding to DNA of a specific nucleotide composition, e.g. GC-rich DNA binding, or with a specific sequence motif or type of DNA e.g. promotor binding or rDNA binding." [GOC:jl] |
| GO:0003684 | 1.36906E-06 | 44 / 304 | "Binding to damaged DNA." [GOC:jl] |
| GO:0000217 | 1.57669E-06 | 42 / 293 | "Binding to a DNA secondary structure element such as a four-way junction, a bubble, a loop, Y-form DNA, or a double-strand/single-strand junction." [GOC:krc] |
| GO:0008301 | 2.099E-06 | 42 / 291 | "The activity of binding selectively and non-covalently to and distorting the original structure of DNA, typically a straight helix, into a bend, or increasing the bend if the original structure was intrinsically bent due to its sequence." [GOC:krc, GOC:vw, PMID:10710711, PMID:1903 |
| GO:0045027 | 2.099E-06 | 42 / 291 | "Binding to DNA ends exposed by the creation of double-strand breaks (DSBs)." [GOC:jl] |
| GO:0070336 | 2.099E-06 | 42 / 291 | "Binding to a flap structure in DNA. A DNA flap structure is one in which a single-stranded length of DNA or RNA protrudes from a double-stranded DNA molecule." [GOC:mah, PMID:15189154] |
| GO:0000497 | 2.41944E-06 | 42 / 290 | "Binding to nucleic acid via hydrogen bonds between the bases of a gene product molecule and the bases of a target DNA molecule." [GOC:krc] |
| GO:0003681 | 2.41944E-06 | 42 / 290 | "Binding to DNA in a bent conformation." [GOC:jl, PMID:12627977] |
| GO:0010844 | 2.41944E-06 | 42 / 290 | "Binding to a genomic region which promotes recombination." [GOC:dph, GOC:tb] |
| GO:0045142 | 2.41944E-06 | 42 / 290 | "Binding to a DNA triple helix. The formation of triple helical DNA has been evoked in several cellular processes including transcription, replication, and recombination." [PMID:10681538] |
| GO:0051880 | 2.41944E-06 | 42 / 290 | "Binding to G-quadruplex DNA structures, in which groups of four guanines adopt a flat, cyclic Hoogsteen hydrogen-bonding arrangement known as a guanine tetrad. The stacking of guanine tetrads results in G-quadruplex DNA structures. G-quadruplex DNA can form under pl |
| GO:0099077 | 2.41944E-06 | 42 / 290 | "DNA-binding activity that is dependent on binding to a histone." [PMID:11691835] |
| GO:0003723 | 4.66055E-06 | 54 / 344 | "Binding to an RNA molecule or a portion thereof." [GOC:jl, GOC:mah] |
| GO:0005198 | 9.46738E-06 | 895 / 374 | "The action of a molecule that contributes to the structural integrity of a complex." [GOC:mah, GOC:vw] |
| GO:0030515 | 9.60221E-06 | 9 / 105 | "Binding to a small nucleolar RNA." [GOC:mah] |
| GO:0019843 | 2.50226E-05 | 11 / 112 | "Binding to a ribosomal RNA." [GOC:jl] |
| GO:0003725 | 3.90904E-05 | 9 / 98 | "Binding to double-stranded RNA." [GOC:jl] |
| GO:0036002 | 3.90904E-05 | 9 / 98 | "Binding to a pre-messenger RNA (pre-mRNA), an intermediate molecule between DNA and protein that may contain introns and, at least in part, encodes one or more proteins. Introns are removed from pre-mRNA to form a mRNA molecule." [GOC:bf, GOC:kmv, PMID:219011 |
| GO:0017069 | 4.75695E-05 | 9 / 97 | "Binding to a small nuclear RNA (snRNA)." [GOC:mah] |
| GO:0033204 | 4.75695E-05 | 9 / 97 | "Binding to RNA subunit of ribonuclease P." [GOC:pg, PMID:11455963] |
| GO:0034336 | 4.75695E-05 | 9 / 97 | "Binding to an RNA molecule that has assumed an incorrect conformation." [GOC:mah, PMID:10393192] |
| GO:0140657 | 5.61994E-05 | 903 / 375 | "A molecular function characterized by the coupling of ATP hydrolysis to other steps of a reaction mechanism to make the reaction energetically favorable, for example to catalyze a reaction or drive transport against a concentration gradient." [PMID:24878343, PMID:25750732 |
| GO:0001069 | 5.78228E-05 | 9 / 96 | "Binding to a RNA region that regulates a nucleic acid-based process. Such processes include transcription, DNA replication, and DNA repair." [GOC:txnOH] |
| GO:0002151 | 5.78228E-05 | 9 / 96 | "Binding to a G-quadruplex RNA structure, in which groups of four guanines adopt a flat, cyclic hydrogen-bonding arrangement known as a guanine tetrad." [PMID:18294969, PMID:18568163, PMID:19330720] |
| GO:0030555 | 5.78228E-05 | 9 / 96 | "Specifies the site of a posttranscriptional modification in an RNA molecule by base pairing with a short sequence around the target residue." [GOC:mah, PMID:12457565] |
| GO:0034583 | 5.78228E-05 | 9 / 96 | "Binding to a 21U-RNA, a 21-nucleotide RNA characterized by a uridine 5'-monophosphate and a modified 3' end resistant to periodate degradation. 21U-RNAs are derived from distinct, autonomously expressed loci within the genome." [GOC:kmv] |
| GO:0035613 | 5.78228E-05 | 9 / 96 | "Binding to a stem-loop in an RNA molecule. An RNA stem-loop is a secondary RNA structure consisting of a double-stranded RNA (dsRNA) stem and a terminal loop." [GOC:sart, PMID:16568238, PMID:20455544] |
| GO:0042835 | 5.78228E-05 | 9 / 96 | "Binding to a BRE RNA element (Bruno response element)." [PMID:10893231] |
| GO:0061752 | 5.78228E-05 | 9 / 96 | "Binding to long non-coding RNA molecules transcribed from subtelomeric regions in most eukaryotes. Telomeric repeat-containing RNA (TERRA) molecules consist of subtelomeric-derived sequences and G-rich telomeric repeats." [GOC:BHF, GOC:BHF_telomere, GOC:dph, GOC |
| GO:0061980 | 5.78228E-05 | 9 / 96 | "Binding to a small regulatory RNA, a short RNA (usually 50-200 nt long) that is either independently transcribed or processed from a longer RNA by an RNAse enzyme." [PMID:14622403, PMID:23475961] |
| GO:0062104 | 5.78228E-05 | 9 / 96 | "Binding to a region of RNA containing a Pumilio-response element element. The consensus sequence for the element is UGUAAAUA." [PMID:30601114] |
| GO:0070034 | 5.78228E-05 | 9 / 96 | "Binding to the telomerase RNA template." [GOC:krc, PMID:16884717] |
| GO:0070878 | 5.78228E-05 | 9 / 96 | "Binding to a primary microRNA (pri-miRNA) transcript, an RNA molecule that is processed into a short hairpin-shaped structure called a pre-miRNA and finally into a functional miRNA. Both double-stranded and single-stranded regions of a pri-miRNA are required for binding. |
| GO:0070883 | 5.78228E-05 | 9 / 96 | "Binding to a precursor microRNA (pre-miRNA) transcript, a stem-loop-containing precursor of microRNA." [PMID:18951094] |
| GO:0071208 | 5.78228E-05 | 9 / 96 | "Binding to the downstream cleavage product (DCP) generated by histone pre-mRNA 3'-end processing." [PMID:19470752] |
| GO:0106222 | 5.78228E-05 | 9 / 96 | "Binding to a long noncoding RNA (lncRNA)." [PMID:25578728] |
| GO:1904678 | 5.78228E-05 | 9 / 96 | "Binding to an alpha-aminoacyl-tRNA." [GO_REF:0000067, GOC:TermGenie, ISBN:155581073X] |
| GO:1990605 | 5.78228E-05 | 9 / 96 | "Binding to an RNA molecule containing GU repeats." [PMID:20081200] |
| GO:0000049 | 6.39286E-05 | 11 / 107 | "Binding to a transfer RNA." [GOC:ai] |
| GO:0003727 | 6.74684E-05 | 10 / 101 | "Binding to single-stranded RNA." [GOC:jl] |
| GO:0000339 | 9.82091E-05 | 10 / 99 | "Binding to a 7-methylguanosine (m7G) group or derivative located at the 5' end of an RNA molecule." [GOC:krc] |
| GO:0140110 | 0.000100983 | 883 / 367 | "A molecular function that controls the rate, timing and/or magnitude of gene transcription. The function of transcriptional regulators is to modulate gene expression at the transcription step so that they are expressed in the right cell at the right time and in the right amount |
| GO:0000099 | 0.000112746 | 10 / 13 | "Enables the transfer of sulfur amino acids from one side of a membrane to the other. Sulphur amino acids contain sulfur in the form of cystine, methionine or their derivatives." [GOC:ai, GOC:mtg_transport, ISBN:0815340729] |
| GO:0009055 | 0.000124508 | 878 / 365 | "A molecular function representing the directed movement of electrons from one molecular entity to another, typically mediated by electron carriers or acceptors, resulting in the transfer of energy and/or the reduction-oxidation (redox) transformation of chemical species. Th |
| GO:0016491 | 0.000144278 | 199 / 657 | "Catalysis of an oxidation-reduction (redox) reaction, a reversible chemical reaction in which the oxidation state of an atom or atoms within a molecule is altered. One substrate acts as a hydrogen or electron donor and becomes oxidized, while the other acts as hydrogen or e |

| GO_Term | P_val | Counts | BioProc_Description |
|---|---|---|---|
| GO:0006259 | 4.68018E-09 | 3 / 96 | "Any cellular metabolic process involving deoxyribonucleic acid. This is one of the two main types of nucleic acid, consisting of a long, unbranched macromolecule formed from one, or more commonly, two, strands of linked deoxyribonucleotides." [ISBN:0198506732] |
| GO:0010467 | 2.45061E-07 | 32 / 255 | "The process in which a gene's sequence is converted into a mature gene product (protein or RNA). This includes the production of an RNA transcript and its processing, as well as translation and maturation for protein-coding genes." [GOC:txnOH-2018, PMID:25934543, PMID: |
| GO:0070914 | 7.19449E-06 | 1 / 51 | "A DNA repair process that is initiated by an endonuclease that introduces a single-strand incision immediately 5' of a UV-induced damage site. UV-damage excision repair acts on both cyclobutane pyrimidine dimers (CPDs) and pyrimidine-pyrimidone 6-4 photoproducts (6-4P |
| GO:0055085 | 8.43651E-06 | 141 / 423 | "The process in which a solute is transported across a lipid bilayer, from one side of a membrane to the other." [GOC:dph, GOC:jid] |
| GO:0035672 | 1.13236E-05 | 135 / 404 | "The process in which an oligopeptide is transported across a membrane. Oligopeptides are molecules that contain a small number (2 to 20) of amino-acid residues connected by peptide linkages." [GOC:vw, ISBN:0198506732] |
| GO:0097080 | 1.21549E-05 | 134 / 401 | "The directed movement of inorganic selenite (HSeO3-1 at physiological pH) across a plasma membrane." [GOC:mcc, PMID:20861301] |
| GO:0000731 | 1.23368E-05 | 1 / 49 | "Synthesis of DNA that proceeds from the broken 3' single-strand DNA end and uses the homologous intact duplex as the template." [PMID:10357855] |
| GO:0006351 | 1.27633E-05 | 24 / 188 | "The synthesis of an RNA transcript from a DNA template." [GOC:jl, GOC:txnOH] |
| GO:0015780 | 1.38702E-05 | 134 / 402 | "The directed movement of nucleotide-sugars into, out of or within a cell, or between cells, by means of some agent such as a transporter or pore. Nucleotide-sugars are any nucleotide in which the distal phosphoric residue of a nucleoside 5'-diphosphate is in glycosidic linka |
| GO:0035461 | 1.38702E-05 | 134 / 402 | "The process in which a vitamin is transported across a membrane. A vitamin is one of a number of unrelated organic substances that occur in many foods in small amounts and that are necessary in trace amounts for the normal metabolic functioning of the body." [GOC:bf] |
| GO:0034220 | 1.58066E-05 | 134 / 403 | "A process in which a monoatomic ion is transported across a membrane. Monatomic ions (also called simple ions) are ions consisting of exactly one atom." [GOC:mah] |
| GO:0046618 | 1.58066E-05 | 134 / 403 | "The directed movement of a xenobiotic from a cell, into the extracellular region. A xenobiotic is a compound foreign to the organim exposed to it. It may be synthesized by another organism (like ampicilin) or it can be a synthetic chemical." [GOC:go_curators, GOC:krc] |
| GO:0006281 | 1.59119E-05 | 8 / 97 | "The process of restoring DNA after damage. Genomes are subject to damage by chemical and physical agents in the environment (e.g. UV and ionizing radiations, chemical mutagens, fungal and bacterial toxins, etc.) and by free radicals or alkylating agents endogenously gene |
| GO:0001407 | 1.69619E-05 | 133 / 400 | "The process in which a glycerophosphodiester is transported across a membrane. Glycerophosphodiesters are small molecules composed of glycerol-3-phosphate and an alcohol, for example, glycerophosphoinositol." [GOC:mcc, PMID:12912892] |
| GO:0015794 | 1.69619E-05 | 133 / 400 | "The process in which glycerol-3-phosphate is transported across a membrane. Glycerol-3-phosphate is a phosphoric monoester of glycerol." [GOC:ai] |
| GO:0015897 | 1.69619E-05 | 133 / 400 | "The process in which an organomercurial compound is transported across a membrane. Organomercurial substances are any organic compound containing a mercury atom." [GOC:ai, PMID:18793329] |
| GO:0015904 | 1.69619E-05 | 133 / 400 | "The directed movement of tetracycline from one side of a membrane to the other. Tetracycline is a broad spectrum antibiotic that blocks binding of aminoacyl tRNA to the ribosomes of both Gram-positive and Gram-negative organisms (and those of organelles)." [GOC:curat |
| GO:0015905 | 1.69619E-05 | 133 / 400 | "The directed movement of bicyclomycin across a lipid bilayer, from one side of a membrane to the other. Bicyclomycin (or bicozamycin) is an antibacterial drug often used as a livestock feed additive." [PMID:20067529] |
| GO:0035378 | 1.69619E-05 | 133 / 400 | "The process in which carbon dioxide (CO2) is transported across a membrane." [GOC:yaf] |
| GO:0035436 | 1.69619E-05 | 133 / 400 | "The process in which triose phosphate (glyceraldehyde 3-phosphate) is transported across a membrane. Glyceraldehyde 3-phosphate is any organic three carbon compound phosphate ester." [GOC:bf, ISBN:0198506732] |
| GO:0070811 | 1.69619E-05 | 133 / 400 | "The process in which glycerol-2-phosphate is transported across a membrane. Glycerol-2-phosphate is a phosphoric monoester of glycerol." [GOC:mah] |
| GO:0071918 | 1.69619E-05 | 133 / 400 | "The process in which urea, the water-soluble compound H2N-CO-NH2, is transported from one side of a membrane to the other by means of some agent such as a transporter or pore." [GOC:mah] |
| GO:0072489 | 1.69619E-05 | 133 / 400 | "The process in which methylammonium is transported across a membrane." [GOC:mah] |
| GO:0072531 | 1.69619E-05 | 133 / 400 | "The process in which a pyrimidine-containing compound is transported across a membrane. A pyrimidine-containing compound is any compound that contains pyrimidine or a formal derivative thereof." [GOC:mah] |
| GO:0080162 | 1.69619E-05 | 133 / 400 | "The directed movement of auxins from endoplasmic reticulum to cytosol." [PMID:19506555] |
| GO:0080170 | 1.69619E-05 | 133 / 400 | "The process in which hydrogen peroxide is transported across a membrane." [GOC:tb] |
| GO:0098660 | 1.69619E-05 | 133 / 400 | "The process in which an inorganic ion is transported across a membrane." [GOC:mah] |
| GO:0140317 | 1.69619E-05 | 133 / 400 | "The directed movement of a substance across the outer membrane in cells with two membranes." [PMID:15968039] |
| GO:0170006 | 1.69619E-05 | 133 / 400 | "The process in which lipoate is transported across a membrane." [PMID:20980265, PMID:25971966] |
| GO:1901642 | 1.69619E-05 | 133 / 400 | "The directed movement of nucleoside across a membrane." [GOC:pr, GOC:TermGenie] |
| GO:1901684 | 1.69619E-05 | 133 / 400 | "The process in which arsenate is transported across a membrane." [GOC:TermGenie] |
| GO:1901962 | 1.69619E-05 | 133 / 400 | "The directed movement of S-adenosyl-L-methionine across a membrane." [GOC:TermGenie, PMID:10497160] |
| GO:1902418 | 1.69619E-05 | 133 / 400 | "The process in which (+)-abscisic acid D-glucopyranosyl este is transported across a membrane." [GOC:TermGenie, PMID:24028845] |
| GO:1902603 | 1.69619E-05 | 133 / 400 | "The directed movement of carnitine across a membrane." [GO_REF:0000069, GOC:pr, GOC:TermGenie] |
| GO:1902616 | 1.69619E-05 | 133 / 400 | "The process in which acyl carnitine is transported across a membrane." [GO_REF:0000069, GOC:pr, GOC:TermGenie] |
| GO:1903825 | 1.69619E-05 | 133 / 400 | "The process in which an organic acid is transported across a membrane." [GO_REF:0000069, GOC:TermGenie] |
| GO:0006366 | 1.91514E-05 | 27 / 201 | "The synthesis of RNA from a DNA template by RNA polymerase II (RNAP II), originating at an RNA polymerase II promoter. Includes transcription of messenger RNA (mRNA) and certain small nuclear RNAs (snRNAs)." [GOC:jl, GOC:txnOH, ISBN:0321000382] |
| GO:0045117 | 1.93021E-05 | 133 / 401 | "The directed movement of azoles, heterocyclic compounds found in many biologically important substances, across a lipid bilayer, across a membrane." [GOC:go_curators, ISBN:3527307206, Wikipedia:Azole] |
| GO:0071994 | 1.93021E-05 | 133 / 401 | "The process in which a phytochelatin is transported across a membrane. Phytochelatins are a group of peptides that bind metals (Cd, Zn, Cu, Pb, Hg) in thiolate coordination complexes." [GOC:mah, ISBN:0198506732] |
| GO:0140820 | 1.93021E-05 | 133 / 401 | "The directed movement of substances from the cytosol into the Golgi apparatus of a cell." [PMID:28373556] |
| GO:0006855 | 2.19361E-05 | 133 / 402 | "The process in which a xenobiotic, a compound foreign to the organim exposed to it, is transported across a membrane. It may be synthesized by another organism (like ampicilin) or it can be a synthetic chemical." [GOC:ai, GOC:bf, GOC:krc] |
| GO:0015791 | 2.19361E-05 | 133 / 402 | "The directed movement of polyols, any polyhydric alcohol, across a membrane." [GOC:ai] |
| GO:0034219 | 2.19361E-05 | 133 / 402 | "The process in which a carbohydrate is transported across a membrane." [GOC:mah] |
| GO:0035382 | 2.19361E-05 | 133 / 402 | "The process in which a sterol is transported across a membrane. Sterols are steroids with one or more hydroxyl groups and a hydrocarbon side-chain in the molecule." [GOC:vw] |
| GO:0072530 | 2.19361E-05 | 133 / 402 | "The process in which a purine-containing compound is transported across a membrane. A purine-containing compound is any compound that contains purine or a formal derivative thereof." [GOC:mah] |
| GO:1901679 | 2.19361E-05 | 133 / 402 | "The directed movement of nucleotide across a membrane." [GOC:pr, GOC:TermGenie] |
| GO:0006974 | 2.21047E-05 | 2 / 55 | "Any process that results in a change in state or activity of a cell (in terms of movement, secretion, enzyme production, gene expression, etc.) as a result of a stimulus indicating damage to its DNA from environmental insults or errors during metabolism." [GOC:go_curators] |
| GO:0034470 | 2.21047E-05 | 2 / 55 | "Any process that results in the conversion of one or more primary non-coding RNA (ncRNA) transcripts into one or more mature ncRNA molecules." [GOC:mah, PMID:27573892] |
| GO:0140115 | 2.32101E-05 | 134 / 406 | "The directed movement of some substance from inside of a cell, across the plasma membrane and into the extracellular region." [GOC:pg] |
| GO:1990542 | 2.4531E-05 | 135 / 410 | "The process in which a solute is transported from one side of a membrane to the other into, out of or within a mitochondrion." [PMID:20533899] |

| GO_Term | P_val | Counts | CellComp_Description |
|---|---|---|---|
| GO:0043231 | 1.05697E-07 | 97 / 579 | "Organized structure of distinctive morphology and function, bounded by a single or double lipid bilayer membrane and occurring within the cell. Includes the nucleus, mitochondria, plastids, vacuoles, and vesicles. Excludes the plasma membrane." [GOC:go_curators] |
| GO:0031981 | 4.58208E-07 | 83 / 501 | "The volume enclosed by the nuclear inner membrane." [GOC:mah, GOC:pz] |
| GO:0005761 | 8.79075E-07 | 3 / 75 | "A ribosome found in the mitochondrion of a eukaryotic cell; contains a characteristic set of proteins distinct from those of cytosolic ribosomes." [GOC:mah, ISBN:0198506732] |
| GO:0030874 | 2.13498E-06 | 0 / 45 | "The portion of nuclear chromatin associated with the nucleolus; includes the DNA encoding the ribosomal RNA." [GOC:mah] |
| GO:0034388 | 2.13498E-06 | 0 / 45 | "A protein complex that forms a subcomplex of the 90S preribosome and can interact directly with the 5' External Transcribed Spacer (ETS) of the full length pre-rRNA transcript. In S. cerevisiae, it sediments at 25-30 S and is composed of Pwp2p, Dip2p, Utp21p, Utp13p, Utp18 |
| GO:0070545 | 2.13498E-06 | 0 / 45 | "A protein complex that is involved in coordinating ribosome biogenesis with cell cycle progression. In human, it is composed of Pes1, Bop1, and WDR12; in Saccharomyces the proteins are known as Nop7p, Erb1 and Ytm1 respectively." [GOC:ab, GOC:mah, PMID:16043514, PM |
| GO:0034456 | 2.85782E-06 | 0 / 44 | "A protein complex that forms a subcomplex of the 90S preribosome. In S. cerevisiae, it is composed of Rrp7p, Utp22p, Ckb1p, Cka1p, Ckb2p and Cka2p." [GOC:mah, PMID:17515605] |
| GO:0034457 | 2.85782E-06 | 0 / 44 | "A protein complex that forms a subcomplex of the 90S preribosome. In S. cerevisiae, it is composed of Mpp10p, Imp3p and Imp4p." [GOC:mah, PMID:17515605] |
| GO:0005759 | 2.88949E-06 | 16 / 152 | "The gel-like material, with considerable fine structure, that lies in the matrix space, or lumen, of a mitochondrion. It contains the enzymes of the tricarboxylic acid cycle and, in some organisms, the enzymes concerned with fatty acid oxidation." [GOC:as, ISBN:0198506732] |
| GO:0005730 | 5.11958E-06 | 0 / 42 | "A small, dense body one or more of which are present in the nucleus of eukaryotic cells. It is rich in RNA and protein, is not bounded by a limiting membrane, and is not seen during mitosis. Its prime function is the transcription of the nucleolar DNA into 45S ribosomal-prec |
| GO:0005763 | 6.85158E-06 | 0 / 41 | "The smaller of the two subunits of a mitochondrial ribosome." [GOC:mcc] |
| GO:0101019 | 6.85158E-06 | 0 / 41 | "A ribonuclease complex that has 3-prime to 5-prime distributive hydrolytic exoribonuclease activity and in some taxa (e.g. yeast) endoribonuclease activity, producing 5-prime-phosphomonoesters. Participates in a multitude of cellular RNA processing and degradation event |
| GO:0005762 | 9.16891E-06 | 0 / 40 | "The larger of the two subunits of a mitochondrial ribosome. Two sites on the ribosomal large subunit are involved in translation: the aminoacyl site (A site) and peptidyl site (P site)." [GOC:mcc] |
| GO:0000120 | 1.22692E-05 | 0 / 39 | "A transcription factor complex that acts at a regulatory region of a gene transcribed by RNA polymerase I." [GOC:mah] |
| GO:0001650 | 1.22692E-05 | 0 / 39 | "A structure found most metazoan nucleoli, but not usually found in lower eukaryotes; surrounded by the dense fibrillar component; the zone of transcription from multiple copies of the pre-rRNA genes is in the border region between these two structures." [PMID:10754561 |
| GO:0001651 | 1.22692E-05 | 0 / 39 | "A structure found in the nucleolus, which contains newly synthesized preribosomal RNA (pre-rRNA) and a collection of proteins." [PMID:10754561] |
| GO:0001652 | 1.22692E-05 | 0 / 39 | "A structure found in the nucleolus, which contains nearly completed preribosomal particles destined for the cytoplasm." [PMID:10754561] |
| GO:0005655 | 1.22692E-05 | 0 / 39 | "A ribonuclease P complex located in the nucleolus of a eukaryotic cell, where it catalyzes the 5' endonucleolytic cleavage of precursor tRNAs to yield mature tRNAs. Eukaryotic nucleolar ribonuclease P complexes generally contain a single RNA molecule that is necessary but r |
| GO:0005736 | 1.22692E-05 | 0 / 39 | "RNA polymerase I, one of three nuclear DNA-directed RNA polymerases found in all eukaryotes, is a multisubunit complex; typically it produces rRNAs. Two large subunits comprise the most conserved portion including the catalytic site and share similarity with other eukary |
| GO:0030685 | 1.22692E-05 | 0 / 39 | "Any complex of pre-rRNAs, ribosomal proteins, and associated proteins formed in the nucleolus during ribosome biogenesis." [PMID:10567516] |
| GO:0030869 | 1.22692E-05 | 0 / 39 | "A protein complex that mediates transcriptional silencing at the rDNA locus (the name derives from regulator of nucleolar silencing and telophase). In Saccharomyces the complex contains Net1p, Sir2p, Cdc14p, and at least one more subunit." [PMID:12196389] |
| GO:0031429 | 1.22692E-05 | 0 / 39 | "A box H/ACA RNP complex that is located in the nucleolus." [GOC:vw, ISBN:0879695897, PMID:17284456, PMID:20227365] |
| GO:0034455 | 1.22692E-05 | 0 / 39 | "A protein complex that forms a subcomplex of the 90S preribosome and is required for the subsequent assembly of the rest of the preribosome. In S. cerevisiae, it is composed of Utp5p, Utp4p, Nan1p, Utp8p, Utp9p, Utp10 and Utp15p." [GOC:krc, GOC:mah, GOC:vw, PMID:1 |
| GO:0061773 | 1.22692E-05 | 0 / 39 | "A chromatin silencing complex that recruits histone-modifying enzymes and upregulates silencing of rDNA in response to glucose starvation." [GOC:BHM, Intact:EBI-11789632, Intact:EBI-11790279, PMID:18485871] |
| GO:0110016 | 1.22692E-05 | 0 / 39 | "A chromatin remodeling complex that positively regulates histone H3 acetylation, in particular H3K9, by recruiting histone acetyltransferases to rDNA gene regions. Located in the nucleolus where it assembles on RNA Polymerase I (Pol I) and possibly on RNA Polymerase III (F |
| GO:0140513 | 1.85138E-05 | 92 / 509 | "A stable assembly of two or more macromolecules, i.e. proteins, nucleic acids, carbohydrates or lipids, in which at least one component is a protein and the constituent parts function together in the nucleus." [GOC:pg] |
| GO:0000943 | 4.74575E-05 | 80 / 445 | "A complex of the retrotransposon RNA genome, reverse transcriptase, integrase, and associated molecules required for reproduction and integration of the retrotransposon into the host genome; the main structural molecule of the nucleocapsid is often a gag protein homo |
| GO:0031039 | 4.74575E-05 | 80 / 445 | "A membrane-bounded organelle of ciliated protozoan cells that contains polyploid copies of a portion of the cell's complete genome. Transcription of genes occurs in macronuclei. Some ciliate species may contain multiple macronuclei per cell." [GOC:ns] |
| GO:0031040 | 4.74575E-05 | 80 / 445 | "A membrane-bounded organelle of ciliated protozoan cells that contains a diploid copy of the cell's complete genome. Sections of contiguous sequence in the macronucleus are often interrupted by internal eliminated sequences (IES), and may be permuted, in micronuclei. |
| GO:0042405 | 4.74575E-05 | 80 / 445 | "An intranuclear focus at which aggregated proteins have been sequestered." [GOC:jl] |
| GO:0043073 | 4.74575E-05 | 80 / 445 | "The nucleus of a germ cell, a reproductive cell in multicellular organisms." [CL:0000586, GOC:go_curators] |
| GO:0043076 | 4.74575E-05 | 80 / 445 | "The nucleus of a megasporocyte, a diploid cell that undergoes meiosis to produce four megaspores, and its descendents." [GOC:jl, ISBN:0618254153] |
| GO:0045120 | 4.74575E-05 | 80 / 445 | "The nucleus of either the ovum or the spermatozoon following fertilization. Thus, in the fertilized ovum, there are two pronuclei, one originating from the ovum, the other from the spermatozoon that brought about fertilization; they approach each other, but do not fuse u |
| GO:0046818 | 4.74575E-05 | 80 / 445 | "A location in the host cell nucleus where viral proteins colocalize late in infection prior to the onset of viral DNA synthesis. More than one site can be present simultaneously." [PMID:10233976] |
| GO:0048353 | 4.74575E-05 | 80 / 445 | "Nucleus resulting from the fusion of the male gamete and two polar nuclei in the central cell of the embryo sac." [ISBN:0471245208] |
| GO:0048555 | 4.74575E-05 | 80 / 445 | "The nucleus of the generative cell, a cell contained within the pollen grain that will divide to produce two haploid sperm cells." [GOC:tair_curators] |
| GO:0048556 | 4.74575E-05 | 80 / 445 | "The nucleus of the microsporocyte. The microsporocyte is a diploid cell in which meiosis will occur, resulting in four microspores. A microspore is a spore that, in vascular plants, gives rise to a male gametophyte." [GOC:tair_curators, ISBN:047186840X] |
| GO:0097165 | 4.74575E-05 | 80 / 445 | "A dense aggregation in the nucleus composed of proteins and RNAs that appear when the cell is under stress." [GOC:ans, PMID:10359787, PMID:12865437] |
| GO:0097571 | 4.74575E-05 | 80 / 445 | "One of the two nuclei found in Giardia species (trophozoite stage). It is located on the left side of the cell when viewed from the dorsal side." [GOC:giardia, ISBN:0-444-81258-X] |
| GO:0097572 | 4.74575E-05 | 80 / 445 | "One of the two nuclei found in Giardia species (trophozoite stage). It is located on the right side of the cell when viewed from the dorsal side." [GOC:giardia, ISBN:0-444-81258-X] |
| GO:0110092 | 4.74575E-05 | 80 / 445 | "The area of a motile nucleus closest to the direction of movement." [GOC:al, GOC:mah, GOC:vw, PMID:15030757, PMID:24335254] |
| GO:0110093 | 4.74575E-05 | 80 / 445 | "The area of a motile nucleus furthest from the direction of movement." [GOC:kmv, PMID:24335254] |
| GO:0140510 | 4.74575E-05 | 80 / 445 | "A narrow constricted region of the nucleus that forms around the anaphase spindle during closed mitosis, and connects the main portions of the newly forming daughter nuclei." [PMID:32848252] |
| GO:1905754 | 4.74575E-05 | 80 / 445 | "Any nucleus that is part of a ascospore-type prospore." [GO_REF:0000064, GOC:TermGenie, PMID:26942678] |
| GO:0005880 | 6.72842E-05 | 81 / 446 | "Any microtubule in the nucleus of a cell." [GOC:mah] |
| GO:0005635 | 7.08284E-05 | 82 / 450 | "The double lipid bilayer enclosing the nucleus and separating its contents from the rest of the cytoplasm; includes the intermembrane space, a gap of width 20-40 nm (also called the perinuclear space)." [ISBN:0198547684] |
| GO:0005634 | 7.79865E-05 | 82 / 449 | "A membrane-bounded organelle of eukaryotic cells in which chromosomes are housed and replicated. In most cells, the nucleus contains all of the cell's chromosomes except the organellar chromosomes, and is the site of RNA synthesis and processing. In some species, or ir |
| GO:0098798 | 0.000128453 | 11 / 103 | "A protein complex that is part of a mitochondrion." [GOC:dos] |
| GO:0009987 | 0.000236916 | 98 / 508 | "Any process that is carried out at the cellular level, but not necessarily restricted to a single cell. For example, cell communication occurs among more than one cell, but occurs at the cellular level." [GOC:go_curators, GOC:isa_complete] |
| GO:0000313 | 0.000377854 | 3 / 50 | "A ribosome contained within a subcellular membrane-bounded organelle." [GOC:mah, GOC:mcc] |

#### S4 - Comparison of DPoP Results With Published Findings From *S. cerevisiae*.

Following lists of phosphosites were identified as differentially phosphorylated during high osmolarity conditions in S. cerevisiae, via analysis by either DPoP, or by published literature (Kanshin et al, 2015).

| Method of Analysis → | DPoP<br>803 Differential | Kanshin et al., 2015<br>596 Differential |
|---|---|---|
| List of Significant Signals ↓<br>(Uniprot ID, P-site) | P31376 S331/T335<br>P31376 T366<br>P31383 T23<br>P31380 S129<br>P31380 T131<br>P31381 S27<br>P31381 S58/S60<br>P07866 S200/S203<br>P07866 S559<br>P07866 T1125<br>P39732 S594/S604/S609<br>P00549 S53<br>Q01329 S579<br>P39723 S48<br>P39717 S633<br>P39705 S246<br>P27637 S507<br>P27637 S655<br>P35845 S691/T694<br>P35845 T694<br>P32790 S454<br>P38199 S358/S362<br>P38193 S413<br>P34216 S1012<br>P34216 S628<br>P34216 T251<br>P34216 T948<br>P34217 S191/S194<br>P34217 S389/T390/S393<br>P34217 S653/S655<br>P34219 S318<br>P34219 S318/S329<br>P34219 S333<br>P34225 S306/S308<br>P34225 T290/S293<br>P38187 T730<br>P38041 S348<br>P38041 S359<br>P38041 S626<br>P38041 S655/S659<br>P38041 S86<br>P38041 T158<br>P38041 T670<br>P38177 S433/S439<br>P38174 T2/S9<br>P38167 T1028/S1030<br>P38165 T379<br>P05316 S6<br>P29465 S32<br>P29465 S600<br>P10664 T287<br>P10664 T60<br>P38079 S293<br>P38080 S531/S541<br>P38084 S24/S26 | P47082 S50<br>P33748 S304<br>P47082 S115<br>Q03656 S449<br>P34219 S280<br>Q12427 S254<br>P40986 S9<br>Q06108 S966<br>P38970 S233<br>P33748 S633<br>P39732 S536<br>P32793 S374<br>P53191 S6<br>P50944 S34<br>P47082 T181<br>P12688 S653<br>P38691 S845<br>P38970 S19<br>P38691 S591<br>P53191 S113<br>P25339 T205<br>P33749 S279<br>Q08206 T353<br>P40059 S313<br>P32568 T53<br>P47082 T117<br>P11792 T723/S726<br>P47013 T16/S18<br>P27692 S1009/S1015<br>P53273 S842/T847<br>P38691 S624/S631<br>P40059 S313/S318<br>P40528 S859/S860<br>P43638 S81/S85<br>P04821 S629/S632<br>P38970 S229/S233<br>Q12256 T606/T608<br>P40059 S313/S318<br>P34219 S317/S318/S329<br>P26570 T171<br>P43633 S258<br>P38329 S870<br>P32790 S449<br>P47035 S1082<br>P39016 S834<br>P32798 S407<br>P33401 S120<br>P47082 S187<br>P40021 S670<br>P53950 S689<br>P32862 T279<br>P20107 S387<br>P34219 S298<br>Q12043 S645<br>Q02208 S527 |

| | | |
|---|---|---|
| | P38085 S13<br>P38249 T938<br>P18412 T273/T276<br>P38257 S222<br>P38261 S523<br>P38266 S843/S847<br>P14922 S790<br>P14922 S815/S817<br>P14922 T938<br>P02994 S289<br>P02994 S6<br>P38272 S343/S348<br>P38283 S73<br>P38284 S120<br>P38284 S217<br>P38302 S139<br>P05755 S162<br>P29366 S137<br>P38310 S449/S452<br>P38313 S41/S45<br>P32327 S513<br>P38321 S156<br>P38329 S1028<br>P38329 S26/S29<br>P38140 S126<br>P38346 S137<br>P29539 S133/S134/S138<br>P29539 S1601<br>P27344 S186/S188<br>P38356 S47<br>P25587 T180/T184<br>P25555 S13<br>P25558 S904<br>P25558 T1157/S1160<br>P25389 S666<br>P25389 S729/S732<br>P25579 T110/S112<br>P25588 S215<br>P25588 S605/S607<br>P25588 S605/S607/T609<br>P25591 S269<br>P25342 S316<br>P25621 S17/S19<br>P25621 S7<br>P25623 S405<br>P25635 S912/S913<br>P25364 T428<br>P25644 S456/S457<br>P15891 T181<br>P24279 T791<br>P24279 T798/S801<br>P32618 S520<br>P32618 S885<br>P32618 T926/S930<br>P37303 S369<br>P33332 S32<br>P33332 S6<br>P40013 S131<br>P40013 S165/S166/S172<br>P40013 T249<br>P08456 S4/S34<br>P40021 S152<br>P40021 S343/S352<br>P40021 S670<br>P40021 T151<br>P18494 S552/S562 | P39962 S403<br>P50942 S1082<br>P40061 S19<br>P18494 S267<br>P46970 S1024<br>P43574 S399<br>P40463 S325<br>P40210 S436<br>P32578 S397<br>Q12494 S664<br>P50089 S642<br>P38814 S242<br>P37012 S119<br>P36003 T161<br>Q99252 T381<br>P42845 T419<br>P24279 T791<br>Q12697 S161<br>Q12296 S688<br>P22211 S317/S320<br>P47013 T16/S18<br>P38903 S263/S264<br>Q12427 S345/S350<br>P40210 T433/S434<br>Q12221 S902/S905<br>Q99271 T874/T876<br>P36169 S276/S280<br>Q08206 T101/S104<br>P18888 S122/T125<br>P34219 S308/S314<br>P32864 S3/S9<br>Q12407 S64/S65<br>P20107 S387/S393/S395<br>P38691 S591/S592/S599<br>Q12256 T606/T608/S614<br>P53273 S149/S152/S153<br>P38970 S217<br>P32793 S274<br>P41911 S72<br>P39998 S257<br>P37262 S42<br>P27692 S1032<br>Q00055 S24<br>P38691 S827<br>P39705 S10<br>Q05785 S167<br>P38968 S988<br>P40956 S240<br>Q06315 S580<br>P22211 S249<br>P06634 S539<br>P33399 S19<br>P27692 S1043<br>P40013 S165<br>Q05518 S202<br>P53094 S781<br>Q12433 S505<br>Q12321 S404<br>P20484 S434<br>P15436 S30<br>Q08816 S131<br>P27705 T371/S377<br>P41913 S359/T363<br>P32578 S181/S182<br>P06102 S442/S446<br>P53939 S123/S125 |

| | | |
|---|---|---|
| | P18494 S619 | P40528 S868 |
| | P10869 T333 | Q07655 S274 |
| | P40048 S248 | Q03208 S72 |
| | P40052 S189/S192 | P32864 S9 |
| | P40059 S318 | P32618 T154 |
| | P40059 S491 | P32356 S60 |
| | P39966 T380 | P46958 S148 |
| | P39969 S519/S523/S528 | Q04991 S249 |
| | P38682 S170 | Q08206 S371 |
| | P38682 S398 | P34761 S225 |
| | P38990 S36 | P53735 S498 |
| | P32634 S1355/S1356 | P34216 S241 |
| | P32634 S1521 | P38070 S90 |
| | P32634 S877/S881 | P22211 S353 |
| | P40091 S226/S230 | P22211 S51 |
| | Q01477 S339/S341 | P41696 S227 |
| | Q01477 S404 | P38167 S775 |
| | P39960 S1012 | P12688 T57 |
| | P06843 S126 | P38970 S275 |
| | P32657 S989 | P53191 S174 |
| | P32660 S1552 | P32873 S488 |
| | P39956 S590 | P11792 T737 |
| | P29311 S212 | P46678 S178 |
| | Q03497 S169/T172 | P47035 S439 |
| | P38741 S320 | Q12523 S12 |
| | P18888 S122/T125 | P46678 S164 |
| | P38739 S498/S502 | P32578 S33 |
| | P38767 S15 | P40917 S99 |
| | P38779 T11 | P33749 S263 |
| | P38788 S477/S480 | P39962 S431 |
| | P38713 S206 | P32432 S170 |
| | P38713 S605/S608 | P24870 S33 |
| | P38691 S815 | P39520 S1041 |
| | P38691 T501/T504 | P53916 T149 |
| | P38692 S735 | Q02725 T183 |
| | P38692 T1073 | Q06628 S280 |
| | P38692 T625 | P53599 S118 |
| | P38692 T645 | P32634 S1307 |
| | P38835 T382 | P35724 S104 |
| | P23291 S523 | Q06673 S1053 |
| | P38845 S332/S335 | P43560 S140 |
| | P38845 T182/S191 | P38165 S246 |
| | P32900 S126 | Q12221 S902 |
| | P32900 S167/T169 | Q06251 T235 |
| | P32900 S219/T221/T232 | P40501 T375 |
| | P38853 S494/S503 | P38167 S527 |
| | P38854 S282 | Q04749 S273 |
| | P38854 S371 | P34216 T251 |
| | P38870 S771 | P38854 S92 |
| | P11792 T737 | P25351 S313 |
| | P40318 S339/S342 | P53191 S82 |
| | P07278 S84 | P40460 S37 |
| | P40535 S557 | P38314 S524 |
| | P40535 T401 | Q12416 S88 |
| | P06102 S111 | P25371 S702 |
| | P06102 S322 | P18494 S619 |
| | P06102 S322/S324 | P34219 S333 |
| | P40529 S207 | Q04749 S305 |
| | P40187 S196 | P53258 S862 |
| | P40528 S868 | P46974 S231 |
| | P40528 T834 | P50089 S667 |
| | P40527 T90 | P43565 S1337 |
| | P04801 S195 | Q08930 S343 |
| | P40498 S155 | Q06479 S627 |
| | P40484 S80/T85 | P50111 S229 |
| | P40433 T157 | Q06833 S714 |

| | | |
|---|---|---|
| | P40433 T158 | P13382 S298 |
| | P40480 T688/S690 | P16649 T275 |
| | P40477 S1073 | P25357 S318 |
| | P40477 S940/S945 | P53120 S1167 |
| | P40477 T870/S874 | P43638 S81 |
| | P40476 S276 | P48415 S759 |
| | P23250 T274 | P22211 S125 |
| | P40474 S6/S7 | P38931 S608 |
| | P26784 S163 | Q08269 S185 |
| | P40463 S159 | P47082 S50/S58 |
| | P40463 T160 | P53273 S37/S42 |
| | P40460 S95/S100 | P53191 S8/T10 |
| | P39928 T274/S278 | Q12296 S614/T617 |
| | P32351 S60 | P40210 T433/S436 |
| | P40563 S167 | Q12256 T606/T608 |
| | P36101 S420 | P34219 S300/S303 |
| | P32861 Y13 | P47082 T181/S187 |
| | P32862 S1130 | P53094 S1202/T1203 |
| | P32862 S492 | Q06820 S18/S21 |
| | P32862 T214/S229 | P23900 S181/S185 |
| | P32857 T498/S518 | P12688 S644/S653 |
| | P32801 S516/S519 | Q12494 S664/S670 |
| | P32801 S581 | P23900 T168/S172/S175 |
| | P35735 Y108 | Q03497 S169 |
| | P14540 Y310 | Q12296 S637 |
| | P33749 S263 | P53943 T28 |
| | P33749 S319 | Q04991 S269 |
| | P33749 S531/S532 | P25351 S321 |
| | P32364 S566/S570 | P47082 S37 |
| | P36076 S233 | Q9P305 S119 |
| | P35201 S325 | Q06324 S37 |
| | P34244 S1045 | Q06836 S356 |
| | P34244 S396/S407 | Q03210 S62 |
| | P34244 S866 | P38970 T157 |
| | P34250 T503/S507 | P40091 T150 |
| | P14164 S215 | P35724 S472 |
| | P32330 S93/S96 | P38835 S511 |
| | P34163 S466 | P53968 S385 |
| | P36003 T161 | P50089 S234 |
| | P36049 S162/S177/S183 | Q03361 T662 |
| | P36049 S177 | P50896 S238 |
| | P36048 S85/T88 | P19659 S783 |
| | P05740 S142 | P32047 S228 |
| | P34233 S253 | P04821 S629 |
| | P34233 T450/S452 | Q07653 S793 |
| | P36015 S178 | P53273 S84 |
| | P36002 S75 | P12688 S644 |
| | Q02208 S653 | P38227 T425 |
| | P36114 S380 | P38767 S15 |
| | P36115 S157 | P51862 S396 |
| | P36117 S569/S573 | P33122 T237 |
| | P36126 S265 | P38287 S354 |
| | P36022 T2202/T2206/T2208 | P39962 S408 |
| | P36150 S281 | Q03465 S326 |
| | P36024 S54/S56 | P32047 S322 |
| | P36165 S55 | P40433 S644 |
| | P36165 S737 | P40361 S9 |
| | P36166 S106/T108 | Q04749 S297 |
| | P36166 S128 | P53273 S842 |
| | P36166 S43/S46 | P36000 T73 |
| | P32901 S594 | P38346 S137 |
| | P36169 S250 | P50089 S232 |
| | P36169 S276 | P24583 S623 |
| | P43563 S56/S59 | P34161 S302 |
| | P43564 S24/T28 | P32583 S394 |
| | P43564 T429 | P07278 S84 |

| | | |
|---|---|---|
| | P43565 S1542<br>P43574 S270<br>P22943 S21/S24<br>P43583 S62/T64<br>P07560 S201/S204<br>P43587 S131/S133<br>P43598 S64/T65<br>Q12451 S7<br>Q12451 T723<br>Q07395 S23<br>Q07442 S50/S54<br>P05318 S96<br>Q12066 S439<br>Q12066 S441<br>Q07533 S117/S122<br>Q07533 S354<br>P25694 S599<br>P04050 S1293<br>P04050 T1471<br>Q12342 S463/S466<br>Q12136 S336/S339<br>Q12136 S53/S72<br>Q12518 T346/S350<br>P04819 S119<br>Q12515 S141<br>Q12515 S36<br>Q12515 S39/S47<br>P23595 S64<br>P38968 S974<br>P38968 S980/S988<br>P38968 T996<br>Q12424 S476/S479<br>Q07655 S236<br>Q07657 S408/S416<br>P35197 S168/T170<br>P35197 T217<br>Q07684 S417<br>P40317 S191/S193<br>P32568 S80/S86<br>Q00816 S572<br>Q00816 S922<br>Q12117 T295<br>P38966 T663/S668<br>Q12675 S1542/T1545<br>Q12675 S396/S403<br>Q03833 S690<br>Q03834 T42<br>O13329 Y460/S467<br>Q04602 S142<br>P13185 S951<br>Q03897 S922<br>P16664 S351<br>Q00402 S1464<br>Q12427 S138/S141<br>Q12427 S345/S350<br>P11075 S434/S437<br>P11075 S779/S782<br>P05453 S148<br>P32494 S134<br>P40986 S9<br>Q04004 S222/S224/S226<br>Q04007 S334<br>Q04007 S399<br>P49686 S137<br>P42073 S245<br>P38994 S267 | P40013 T249<br>P50896 T334<br>Q04991 S259<br>P38835 T382<br>P53274 T93<br>P38284 S217<br>P38703 S23/S24<br>P40463 S159/T160<br>Q03656 S449/S453<br>P38691 S1006/S1014<br>Q04991 S238/S249<br>P07278 S83/S84<br>P32330 S93/S96<br>P18494 S552/S562<br>Q03210 T74<br>P08018 S68<br>P40059 S423<br>Q05518 S416<br>Q03088 S662<br>P46675 S813<br>Q07807 S86<br>Q02831 S95<br>P47068 S565<br>P36076 S42<br>P37304 S659/S664<br>P06102 S450/T454<br>P43564 S1037/S1045<br>P40494 T553<br>P00549 S22<br>P36062 S119<br>P39016 S838<br>P40463 S159<br>Q12494 S55<br>P47029 S838<br>P47068 S631<br>P48415 S1611<br>P25623 S347<br>Q07825 S56<br>P38590 T170<br>P33202 T135<br>P33749 S531/S532<br>P34072 T553/S554<br>Q99271 T874/T876<br>P40095 S28<br>Q12476 S256<br>Q12424 S364<br>P38928 S676<br>P32944 S379<br>P32790 T993<br>Q99271 S568<br>P33314 S1071<br>P32900 S369<br>P53274 T152<br>P30605 S55<br>P34244 S1045<br>P34237 S453<br>P38261 S313<br>P39109 S873<br>P53064 S435<br>P38692 T645<br>P38854 S55<br>P36115 S195<br>P08678 S234<br>Q08157 S255<br>P40433 T157<br>P32599 T111 |

| | | |
|---|---|---|
| | P38994 T222 | P29465 S40 |
| | P32501 S538 | P38835 S616/T620 |
| | P07248 S230 | P38284 S141/S145 |
| | P07248 S299/S302 | Q12071 S66/S69 |
| | P14737 S83 | Q12427 S285/S286 |
| | P11978 S1134 | P47035 S615/S619 |
| | P11978 S516 | P26570 Y221/T235 |
| | P11978 T62/S63 | Q92317 S135 |
| | Q04934 S419 | P38165 T56 |
| | Q04934 S59 | P48415 S1515 |
| | Q04947 S206 | P43585 S187 |
| | Q03780 S342/S345/S348 | Q04602 S106 |
| | P37304 T755 | P38329 S34 |
| | Q05543 S294/S297 | P33300 S349 |
| | P24276 S154/S155 | P32833 T187/S188 |
| | P24276 S319/S322/S325 | P40468 S2352/S2355 |
| | P24276 S578 | P32047 S257 |
| | P24276 T261 | P34217 T512 |
| | Q06631 S10 | Q06146 S197 |
| | Q06631 S41/S44 | P53244 S158 |
| | Q05518 T57 | Q04947 S206 |
| | P29509 T251 | P08678 S241 |
| | Q06340 S125/S126/S128 | P32909 S96 |
| | Q06340 S71 | P06634 S37 |
| | P33301 S533/S534 | P40048 T75 |
| | Q06407 S375 | P05453 S148 |
| | Q06407 S523 | P20485 S14 |
| | Q06407 S524/S527 | P38713 T210 |
| | Q06407 S702/S707 | P25587 S241 |
| | P17121 S99 | P40468 S300 |
| | Q92317 S137 | Q03833 S425 |
| | Q04177 S365 | Q04279 S217 |
| | Q04183 S364 | P40474 S40 |
| | Q04031 S113/S116/S122 | P20485 S12 |
| | P32578 S181/S182 | Q02866 S475 |
| | P32578 S299/S302 | Q02100 T215 |
| | P32578 S81 | P53950 S260 |
| | P33329 S203 | P39104 S396 |
| | P33329 T93 | Q03769 S394 |
| | Q04087 S307 | P48361 S948 |
| | P14127 S96 | P38080 S985 |
| | Q03281 S141 | Q06636 S16 |
| | Q03306 T760 | Q07825 S69 |
| | Q03361 S145 | P53950 T258 |
| | Q03361 S552/S568 | P47061 S790 |
| | Q03361 S851 | P53739 S414 |
| | P11491 S123 | Q12753 S439 |
| | P50896 T523 | Q03361 S510 |
| | Q12263 S382/S385/S389 | P36041 S387 |
| | Q12263 S750 | P36004 S131 |
| | P48813 S111/S112 | P38590 S98 |
| | Q04412 S450 | Q12494 S27 |
| | P05030 S911/T912 | Q12502 S798 |
| | P07264 S488 | P25623 S405 |
| | P43633 S258 | P53739 S414/S426 |
| | P43633 S440/S443 | Q06146 S197/S200 |
| | P43633 S76 | Q07824 S72/S76 |
| | P53191 S6 | P39962 S404/S408 |
| | P53191 S8/T10 | Q02100 S108/T113 |
| | P53191 S82 | P53599 S54/S57 |
| | P05737 S228 | Q06108 S1048/S1059 |
| | P53148 T135 | P08678 T389 |
| | P53131 S38 | Q08269 S383 |
| | P53129 S130 | P35209 S627 |
| | P0CX44 S6 | Q02100 S54 |
| | P0CX44 S7 | P38968 S974 |

| | | |
|---|---|---|
| | P22147 S1510 | P25341 S269 |
| | P53104 S647 | Q02875 S651 |
| | P53094 S618/S621 | P38623 S61 |
| | P53094 S702 | P48361 S973 |
| | P53094 S781/T785 | Q02875 S644 |
| | P53094 S900/S904 | Q06108 T1046 |
| | P53094 S919 | Q08831 S311 |
| | P53090 S272 | P39730 S257 |
| | P53090 T311 | Q9P305 S107 |
| | P06774 S19 | Q03533 S449 |
| | P06774 S31 | P47072 T25 |
| | P04807 S15 | P32909 S84 |
| | P53201 S354 | P38692 T625 |
| | P16965 S69 | P43565 S1502 |
| | P40357 S186 | P53115 S65 |
| | P40357 S220 | Q12515 S47 |
| | P32334 S1300 | P33332 S43 |
| | P53238 S83 | Q12236 T74 |
| | P53241 T53/T56 | P34761 T479 |
| | P53244 S158 | P40563 S476 |
| | P53250 S308 | P41697 T12 |
| | P53252 T233 | Q99247 S668 |
| | P48361 S948 | P48562 S367 |
| | P48361 S969 | P53076 S248 |
| | P53267 S20/S23 | Q02785 S56 |
| | P53267 S31 | P43565 T1360 |
| | P23615 S94 | P19659 T793 |
| | P48837 S479 | P06786 S629 |
| | P49090 T346/S352 | Q02875 S657 |
| | P53273 S842 | P46676 S738 |
| | P53274 S85 | P40529 S183 |
| | P53274 T93 | P10961 T97 |
| | P53274 T93/S106 | P38845 S332 |
| | P53276 T142 | P38623 S520 |
| | P53278 T342/S343/S347 | P32325 S623 |
| | P53280 S1121 | P38623 S46/S50 |
| | P32504 S506/S509 | Q12256 T608/S614 |
| | P33336 S200 | Q03088 S542/S545 |
| | P27476 S405 | P43598 S64/T65 |
| | P32893 S183/S187 | P50942 S1081/S1082 |
| | P41895 S198 | P34217 S653/S655 |
| | P00359 S201 | Q05518 T23/S26 |
| | P32566 S381/S394 | P48415 S762/S765/S768 |
| | P32566 S394/S400 | P38427 S147 |
| | P32566 S488 | P38968 T1050 |
| | P50089 S232 | Q06146 S137 |
| | P16861 S189/S192 | P38427 S161 |
| | P53049 S18/S24 | P38853 S503 |
| | O13535 S1093 | P06101 S17 |
| | P08678 S241 | P32909 S125 |
| | P08678 S75/S78 | P32770 S455 |
| | P47072 T25 | P38080 S458 |
| | P47068 S631/S634 | P53191 S300 |
| | P47068 T514 | P36165 S836 |
| | P47068 T88 | P53898 S81 |
| | P47068 T895 | P38165 S227 |
| | P47061 S790 | Q06385 S14 |
| | P20448 S710/S714 | P38426 S181 |
| | P14907 S317 | P53925 S244 |
| | P14907 S631 | Q08831 S14 |
| | P43638 S174 | Q08732 S139 |
| | P43638 S85 | Q12071 S69/S72 |
| | P47047 T38 | P53890 S270/T274 |
| | P47041 S71 | P48524 S54/S56/S58 |
| | P40361 S34/S41 | P53894 T559 |
| | P47035 S1082 | P32485 T174 |

| | | |
|---|---|---|
| | P47035 S452<br>P47035 T304<br>P47035 T357<br>P06105 T50<br>P80428 S349<br>P80428 T338<br>P32447 T265<br>P08018 S9<br>P10081 S129<br>P10081 T146<br>P21268 S361<br>P38970 S336<br>P38970 S66/S68<br>P38970 S68<br>P38970 T157<br>P32525 S436<br>P47082 S37<br>P47082 T117<br>P47098 S38<br>P21373 S67<br>P47113 T216/T219<br>P47116 S752/S755<br>P47116 S781/S784<br>P47116 S784<br>P17106 S156<br>P17106 S45/S48<br>P47134 S383<br>P47134 T604<br>P47135 S951/S954<br>P47136 S367<br>P47136 S538/S539<br>P47136 S545<br>P47139 S60/S69<br>P47170 S737<br>P14772 S885<br>P14772 S931/T934<br>P04802 S214<br>P23201 S646<br>Q12373 S174<br>Q07824 T89/S91<br>Q07872 S213<br>P23900 S212<br>Q12244 S54<br>Q04673 S7/S9<br>Q07950 S73/S76<br>P46675 S280<br>P46675 S812/S813/S815<br>P46675 S812/S815<br>Q12019 S4353<br>P32485 T174/Y176<br>Q12500 S513<br>Q12436 S236/S245<br>P21192 S709<br>P21192 S709/S714<br>P20485 S12<br>Q99271 T765/S768<br>Q99271 T874/T876<br>Q99296 S651<br>P33322 S416<br>P21304 S131<br>Q05785 T450<br>Q05785 T472/T479<br>Q05812 S108/S123<br>Q05812 S127/S130<br>Q05812 S587<br>P47818 S83 | P53894 T743<br>P53836 S881/S885<br>P39960 S303<br>P32485 Y176<br>P38080 T1012<br>Q12446 T380<br>P53309 S451<br>Q9P305 S115<br>P15891 S357<br>Q04279 S685<br>Q05812 S57<br>P34250 S45<br>P25623 S297<br>Q04279 S645<br>P34250 S236<br>Q06407 S277<br>P47116 T160<br>P47113 S180<br>P47041 S139<br>P47068 T88<br>P43638 S85<br>P39524 S102<br>Q06407 S375<br>P43597 S201<br>P32568 S80<br>P38080 T857<br>P38041 S525<br>P38814 S305<br>P39101 S266<br>P47136 S367<br>P47068 T514<br>Q02866 S173/S176<br>Q08229 S117/S120<br>P48415 S840/S841<br>P24276 S154/S155<br>P36027 S352/S355<br>P32895 T40/S47<br>Q04279 S235/S238<br>Q04279 S235/S238<br>P40494 S484/T488<br>P53191 S381<br>Q04279 S48<br>Q12009 S59<br>P47136 S167<br>P38970 S391/S395<br>P40072 T66/S67<br>P38041 S525/S528<br>P48361 S1045<br>P08018 S514<br>P38691 S599<br>P34232 T125<br>P38623 T379<br>P37370 S695<br>P32909 T70<br>Q04279 S118<br>P48361 T808<br>P53274 S155<br>Q03758 T22<br>P23900 T168<br>P38080 S10<br>P40986 S12<br>P33749 S319<br>P53897 S64<br>P38080 S12<br>Q05785 T479<br>Q06108 T21 |

| | | |
|---|---|---|
| | P39520 S41<br>P39520 S972<br>Q05958 S451/S464<br>P38623 S46/S50<br>P38623 S520<br>P38623 S61<br>P38623 T379<br>Q06146 S129<br>Q06146 S197<br>P19882 S22/S23<br>P04821 S629/S632<br>P32457 S41/T44/T47<br>P32457 S503/S509<br>P41697 T352<br>P32499 S399<br>P32499 S540<br>P37370 S709<br>Q06144 S9<br>P35817 S615/T617<br>Q06685 S77<br>Q06440 S443<br>Q06440 S463<br>P06701 T425<br>Q06205 S82<br>P52919 S260/S265<br>Q04233 S718<br>P25087 S375<br>P26570 Y221/T235<br>P50109 S150<br>P50109 S238<br>P50109 S376<br>P50109 T298<br>Q03718 S159<br>Q03714 S376/S379<br>Q03707 S231<br>Q03707 T241<br>Q04711 S1095<br>Q04711 S47<br>P54003 S293/S301<br>P23248 S80<br>Q03640 S1371/S1373/T1379<br>Q03640 S1400/S1403/S1404<br>P38911 S80/S81<br>Q04511 T514<br>Q04500 S142<br>Q04500 S34/S35<br>Q04500 T628<br>P54786 S575/S577<br>Q03758 T22<br>Q03735 S467<br>Q03208 S180<br>Q03208 S2<br>P50105 T19<br>Q03690 T1253/S1259<br>Q05050 S791/S795<br>P23748 S415/S417<br>P33748 S304/S308<br>Q04749 S297<br>Q04779 S17<br>Q04279 S450<br>Q04279 S656<br>Q04279 S669/T675<br>Q04461 S237/S249<br>P39523 S649<br>Q04216 S335/S338<br>P40210 T433/S434 | P40059 S318<br>P32790 S996<br>P38835 S365<br>Q05518 T153<br>Q04279 S217/S220<br>P19882 S22/S23<br>P19659 S746/T750<br>P19659 S789/T793<br>P32485 T174/Y176<br>P34217 S653/S655<br>P40494 T553/S556<br>P32790 S785<br>P11792 S726<br>Q06407 S707<br>P17121 S70<br>Q03780 S363<br>Q04279 S85<br>P19659 S746<br>P34219 S318<br>Q06108 S975<br>Q03656 S393<br>P48524 T111<br>Q06146 S48<br>P47113 S150<br>P22137 T1651<br>Q08732 S37<br>P23201 T1065<br>P53894 T751<br>P40059 S322<br>P22211 S316<br>Q9P305 S63<br>P22211 S357<br>P23255 T158<br>P27705 S179<br>P33748 S629<br>Q12494 S646<br>P38249 T938<br>P33748 S625<br>P39732 S609<br>Q04007 S392<br>P43565 S1557<br>Q06451 S55<br>P43560 S149<br>Q05785 T450<br>Q05775 T75<br>P48524 S58<br>P50944 S71<br>P08018 S9<br>P40048 S445<br>P46949 S92<br>P27637 S655/S658<br>P48415 S678/S681<br>Q03497 T203/S206<br>P48361 S804/T808<br>Q02805 S722/S725<br>P53009 S704/T708<br>P39732 S605/S609<br>Q03780 S342/S345<br>P48524 T878/T881<br>P34217 S389/T390/S393<br>P48361 S1070<br>Q01389 S266<br>Q9P305 S128<br>P24276 S126<br>Q01389 S411<br>Q12515 S246 |

| | | |
|---|---|---|
| | Q12674 S661 | P24276 S228 |
| | Q12674 S677 | P48415 T1691 |
| | P32567 S773/S774 | P34217 S655 |
| | P35209 S627 | P38968 S836 |
| | P35209 S743/S755 | P23201 S599 |
| | P49095 S29 | P17121 S37 |
| | Q04322 S73 | Q12518 T366 |
| | Q03656 S449/S453 | P43603 S375 |
| | Q03656 S453 | P53901 S376 |
| | Q03661 S1527/S1528 | P23293 S417 |
| | Q03661 S1539 | P48415 S1511/S1515 |
| | Q03661 S579/S583 | P36166 S43/S46 |
| | Q03661 S865/S866 | P53894 S63/S66 |
| | Q03661 T593 | |
| | P32829 S578 | |
| | P20107 S387 | |
| | P38426 S373/S374 | |
| | P38426 T265/S267/S273 | |
| | P49960 S19 | |
| | P50111 S229 | |
| | P48524 S54/S56 | |
| | P48524 S54/S56/S58 | |
| | Q03262 S158 | |
| | Q03533 S449 | |
| | Q03559 S14/S23 | |
| | P41818 S158 | |
| | P38590 S98 | |
| | P53950 S260 | |
| | P53950 T258 | |
| | P53947 S276 | |
| | P32047 S257 | |
| | P32047 S265 | |
| | P32047 S322 | |
| | P32047 T173 | |
| | P06786 S1370 | |
| | P53935 S960 | |
| | P53935 T1063 | |
| | P01120 S273/T276 | |
| | P50944 S71 | |
| | P13382 S240 | |
| | P13382 S298 | |
| | P32389 T206 | |
| | P53550 S725/S728 | |
| | P53550 S747/S751 | |
| | P53917 S524/S527 | |
| | P53911 S143 | |
| | P53911 S200 | |
| | P53903 S119 | |
| | P53901 S364/S376 | |
| | P53898 S49/S52 | |
| | P53890 S270/T274 | |
| | P53890 S346 | |
| | Q02100 S558 | |
| | P53885 S288/S291 | |
| | P22211 S317/S320 | |
| | P22211 S353/S356 | |
| | P22211 S353/S356/S357 | |
| | P22211 S357 | |
| | P22211 T247 | |
| | P34761 T479 | |
| | P41812 S503 | |
| | P53860 S346/S349 | |
| | P53858 S278/S281 | |
| | P53858 S746 | |
| | P33338 T294/T296/S308 | |
| | P33338 T296 | |

| | |
|---|---|
| | P53617 S263/S265<br>P41832 S261<br>P53836 S24<br>P53836 S518<br>P53836 S881/S885<br>P53836 T576<br>P48562 S367<br>P21965 S196<br>P21965 Y199<br>P42845 T209<br>P42839 S121<br>P42839 T14<br>P42836 T318/S322<br>P53628 S13<br>P53599 S118<br>P53599 S39<br>P53599 S54/S57<br>P04786 S14/S15<br>Q08109 S528<br>Q08157 S255<br>Q08157 S334/S337<br>P25040 T270<br>P05319 S96<br>P19659 T793<br>Q12296 S688<br>Q12265 T120/T127<br>Q08229 S154<br>Q08229 S204/S205<br>Q08229 S66<br>Q08236 S1024/S1027<br>P35193 S232/S243/S246<br>Q99247 S668<br>Q12236 S1001<br>Q12236 S686<br>Q12469 S169<br>Q99383 S462<br>Q08269 S176<br>Q08269 S383<br>Q08280 S33/S41<br>Q08281 S1087/S1089<br>P89105 S1015/S1017<br>P50875 T516<br>Q08299 S16/T17<br>Q08299 T17<br>P54867 T356<br>Q02805 S436<br>Q02805 S722/S725<br>Q08412 S91<br>Q08412 T364<br>Q08438 T143<br>Q08438 T90<br>Q92331 S32/S37<br>Q12416 S154/S156<br>Q12416 S88<br>Q12275 S58<br>Q12271 S1031/S1035<br>P39083 S388<br>P33302 S837/S840/S841<br>P33302 S849/S850/S854<br>Q12216 S489<br>Q12246 S154/S158<br>Q12123 S341<br>P21339 S33<br>P06634 S535/S539<br>P06634 S585<br>Q01919 S365 |

| | |
|---|---|
| | Q08641 S93<br>Q08650 S17<br>Q08732 S37<br>Q12256 S52<br>Q12256 S549<br>Q12256 S633/S636<br>Q12256 T608<br>Q12256 T608/S614<br>P12962 S91<br>Q12697 S161<br>Q12697 T95<br>O14468 T22/S24<br>Q08773 T1079<br>Q12382 S44/S45<br>Q12507 T45<br>P32798 S225<br>Q12502 S798<br>Q08831 S14<br>P32336 T78<br>Q08904 S3<br>Q12200 S882/S885<br>Q02725 S198<br>Q02725 T183<br>Q03088 S542/S545<br>Q03088 S702<br>Q03063 S126/S129<br>Q03063 S142<br>Q02770 S267<br>P48415 S313<br>P48415 S472/S483<br>P48415 S660/S663<br>P48415 S681<br>P48415 S969<br>Q02884 S222<br>Q02875 S644<br>Q02979 S965<br>Q02969 S49<br>P32873 S229<br>P32873 S324/S327<br>Q02959 S533<br>Q02950 S135<br>P33419 T240<br>P53551 S174<br>Q03016 S66<br>Q03002 S766<br>P05744 S97<br>Q12152 S8<br>P26637 T152<br>P32945 S208<br>Q08921 S141/S144<br>Q08921 S227/S234<br>Q08923 S267<br>Q08926 S98<br>Q08951 S700/S703<br>Q08952 S178<br>Q08966 S105<br>Q08967 S635<br>Q12344 S66<br>Q12365 S61<br>Q12365 T51/S54<br>P53388 S18<br>Q12522 S233<br>Q12495 S515<br>P30665 S145<br>P30665 S52/S56<br>P38431 T187/S189 |

| | |
|---|---|
| | Q12221 S902/S905<br>Q06815 S281<br>Q06833 S714<br>Q06833 S717/S720/S723<br>Q06833 S720/S723<br>P32328 S71/S75<br>Q06108 S975<br>Q06523 T343<br>Q12207 T162<br>P32486 S134/S136<br>P06738 S19<br>P34167 T370<br>Q06604 S39/S43/S46<br>Q06616 S170/S175<br>Q06625 S155/S162<br>Q06628 S346/S348<br>Q06628 S428<br>P32349 S455<br>Q9P305 S106/S107<br>Q3E829 S60 |

.
